# Functional differentiation of Sec13 paralogs in the euglenozoan protists

**DOI:** 10.1101/2022.11.08.515606

**Authors:** Drahomíra Faktorová, Kristína Záhonová, Corinna Benz, Joel B. Dacks, Mark C. Field, Julius Lukeš

## Abstract

The β-propeller protein Sec13 plays roles in at least three distinct processes by virtue of being a component of the COPII endoplasmic reticulum export vesicle coat, the nuclear pore complex (NPC) and the Seh1-associated (SEA)/GATOR nutrient-sensing complex, suggesting that regulatory mechanisms coordinating these cellular activities may operate *via* Sec13. The NPC, COPII and SEA/GATOR are all ancient features of eukaryotic cells. In the vast majority of eukaryotes, a single Sec13 gene is present, but here we report that the Euglenozoa, a lineage encompassing the diplonemid, kinetoplastid and euglenid protists contain two Sec13 paralogs. Furthermore, based on protein interactions and localization studies we show that Sec13 functions in diplonemids are divided between the Sec13a and Sec13b paralogs. Specifically, Sec13a interacts with COPII and the NPC, but Sec13b interacts with Sec16 and components of the SEA/GATOR complex. We infer that euglenozoan Sec13a is responsible for NPC functions and canonical anterograde transport activities while Sec13b acts within nutrient and autophagy-related pathways, indicating a fundamentally distinct organization of coatomer complexes in the euglenozoan flagellates.

## Introduction

Adaptation requires the development or modification of specific functions and arises by multiple mechanisms. Paralog gene expansion is a common mechanism, which we and others have argued is a major driver behind eukaryogenesis and the origins of intracellular compartments (Dacks and Field, 2007; Dacks and Field, 2018; Wideman and Muñoz-Gómez, 2016). While many events occurred during eukaryogenesis, this is an ongoing process, with evolution of lineage-specific genes frequently associated with paralog expansions (Carlton et al., 2007; Rutherford and Moore, 2002). Membrane-trafficking and related processes are examples of a highly evolvable system featuring many expansions and losses between the configurations in different organisms, likely reflecting a central importance of these organelles and a high degree of evolvability. The membrane coating/deforming protocoatomer complexes that make up the vesicle coats and nuclear pore complex (NPC) are ancient, with reconstructions placing the majority as present in the last eukaryotic common ancestor (LECA) (Neumann et al., 2010). Elaboration within individual lineages, by creation of paralogs, losses of components or entire complexes has been reported extensively, demonstrating that despite an ancient origin and central role within eukaryogenesis, considerable plasticity allows for ongoing modifications to vesicle transport and organellar complexity. Furthermore, it has been argued that the protocoatomer architecture, based around an evolvable β-propeller/a-solenoid protein is exceptionally well suited for the evolution of new functionality (Rout and Field, 2022).

The COPII complex is central to anterograde membrane-trafficking from the endoplasmic reticulum (ER) and was present in LECA. Canonically, it is composed of seven core subunits, of which five, Sar1, Sec13, Sec23, Sec24 and Sec31 are near ubiquitous. The two other subunits, Sec12 and Sec16, are less well conserved (Schlacht and Dacks, 2015). Sec12 is guanine nucleotide exchange factor that activates Sar1 by enhancing exchange of GDP for GTP and is located at the ER membrane. Sar1:GTP recruits the Sec23/Sec24 complex and subsequent recruitment of Sec13/Sec31 heterodimers generates the COPII coat. A previously unrecognised pan-eukaryotic paralog of Sar1, SarB, was recently reported, which may suggest differentiation within COPII recruitment mechanisms (Vargová et al., 2021). Sec16 is part of a scaffold, which locates to ER exit sites (ERES) and is essential for COPII recruitment but is also implicated in non-conventional exocytosis and autophagy (Tang, 2017; Yorimitsu and Sato, 2020). However, the absence of Sec16 from many lineages questions the essentiality of Sec16 towards COPII transport (Schlacht and Dacks, 2015).

Sec13, a β-propeller protein and a member of the extensive protocoatomer family, is a component of at least three distinct complexes, and these promiscuous interactions may reflect a deeper role in the coordination of multiple processes, *albeit* with details presently unclear. Besides a role in ER-derived transport, Sec13 is also a component of the NPC and the SEA/GATOR protein complex (Dokudovskaya et al., 2011; Fontoura et al., 1999). Sec13 provides positive regulation to TORC1 signaling and is instrumental in assembly of the COPII membrane-deforming coat (Barlowe et al., 1994). Sec13, COPII, the NPC and SEA/GATOR are well conserved throughout eukaryotes, with only one Sec13 paralog is found in most lineages.

Diplonemids are highly diverse heterotrophic flagellates, and very abundant in the oceans (Flegontova et al., 2020). They form a sister group to the mostly parasitic kinetoplastids and represent the third arm of the Euglenozoa, with representative genome and/or transcriptomes available from all lineages. Several diplonemids are bacterivorous (Prokopchuk et al., 2022), although the lifestyle of most species remains unknown, and likely ranges from phagotrophy through predation to parasitism (Tashyreva et al., 2022). Importantly, these ecologically and evolutionary highly relevant protists recently joined a rather narrow group of genetically tractable organisms (Faktorová et al., 2020a), and functions can now be studied in the model species *Paradiplonema papillatum* (renamed from *Diplonema papillatum*) (Faktorová et al., 2020b; Tashyreva et al., 2022).

As the COPII complex is well-conserved, we considered this as a good target for study in *P. papillatum* and selected PpSec13 for analysis, to address the questions of how conserved the role of Sec13 in this divergent organism is. Importantly, there are two Sec13 paralogs in kinetoplastid flagellates, and with the availability of diplonemid genomes and/or transcriptomes we asked whether the same feature occurred also in this sister lineage. We found that Sec13 is indeed present as two paralogs, PpSec13a and PpSec13b, but unexpectedly the functionality of Sec13 in *P. papillatum* is divided, with PpSec13a involved in conventional COPII and NPC functions, while PpSec13b participates in autophagy and nutrient-sensitive functions. Sec13 division of labor is an early feature of the whole Euglenozoa lineage, suggesting a distinct strategy for regulating the multiple functions of Sec13.

## Results

### COPII components in diplonemids

Using sequences of COPII subunits previously identified in other members of the euglenozoans and *Naegleria gruberi*, a representative of their sister lineage Heterolobosea (Schlacht and Dacks, 2015), we used homology-searching tools to identify genes encoding these proteins in the genome of *Paradiplonema papillatum* and recently available transcriptomes from six additional diplonemid species, namely *Diplonema japonicum, Rhynchopus humris, Lacrimia lanifica, Sulcionema specki, Artemidia motanka* and *Namystynia karyoxenos* (Kaur et al., 2020). We identified orthologs for Sar1, Sec13, Sec23, Sec24, and Sec31 (**Fig. 1**; **Suppl. Tables 1 and 2**), whereas the less conserved Sec12 and Sec16 were not found, even with more sensitive hidden Markov model searches. Several of the conserved subunits were duplicated in all or most diplonemid species. To investigate these duplications, we performed phylogenetic analyses including sequences of relatives from the kinetoplastid and euglenid lineages.

**Fig. 1.**
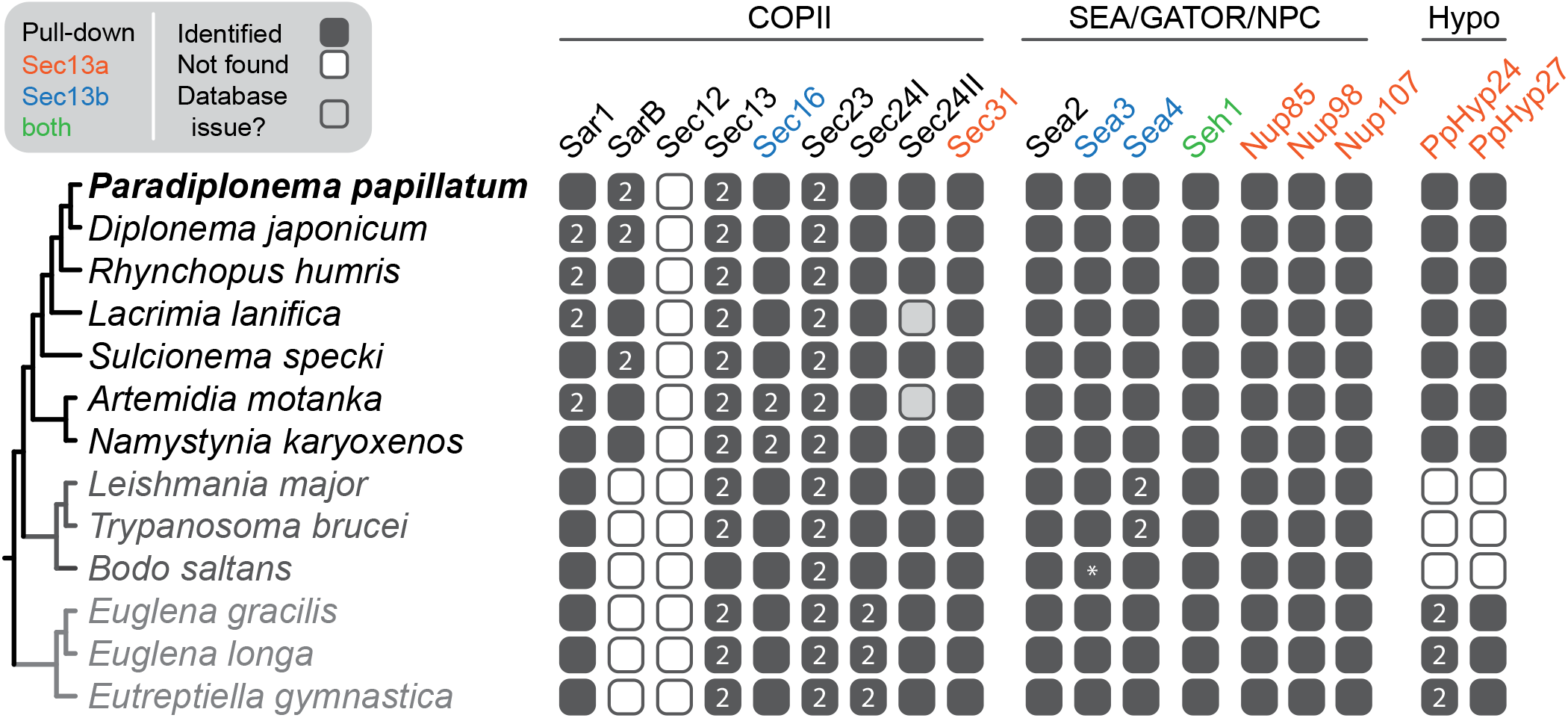
Overview of presence of COPII, SEA/GATOR/NPC and hypothetical (Hypo) subunits/proteins in Euglenozoa. Proteins pulled down with PpSec13a or PpSec13b proteins or both are shown in orange, blue or green, respectively. The annotation of sequences was confirmed by reverse best hit and/or phylogenetic analyses. The only notable exception was *B. saltans* Sea3 (marked by asterisk), which retrieved Sea4 in BLAST against *P. papillatum* proteins, but Sea3 against heterolobosean *N. gruberi* (**Suppl. Table 7**). In the phylogenetic tree, it clustered together with Sea3 sequences from trypanosomatids, and thus we concluded that it is in fact Sea3 as well. For individual phylogenetic trees, see **Suppl. Figs. 1 and 2**.

COPII components are present in the parasitic trypanosomatids *Trypanosoma brucei*, *Leptomonas pyrrhocoris* and *Leishmania major* (Berriman et al., 2005; Flegontov et al., 2016; Ivens et al., 2005) and in the free-living euglenid *Euglena gracilis* (Ebenezer et al., 2019). Moreover, these components are functionally equivalent to orthologs in animal and fungal model organisms based on localization and genetic manipulations (Demmel et al., 2011; Kruzel et al., 2017; Sevova and Bangs, 2009). We expanded sampling by including the free-living kinetoplastid *Bodo saltans* and the euglenids *E. longa* and *Eutreptiella gymnastica*, to represent the diversity of euglenozoans. Sar1 was duplicated within the diplonemid lineage but only one ortholog was identified in *P. papillatum, S. specki*, and *N. karyoxenos* (**Fig. 1; Suppl. Fig. 1A**). Additionally, SarB was identified as a highly divergent ortholog in all diplonemids (**Suppl. Fig. 1A**). Duplication of Sec23 clearly took place in the euglenozoan ancestor (**Suppl. Fig. 1B**). Euglenozoan also possess both LECA paralogs (Sec24I and Sec24II) of Sec24 with euglenids having duplicated Sec24I (**Fig. 1; Suppl. Fig. 1C**). Sec31 is present as a single paralog in all euglenozoans (**Suppl. Fig. 1D**). Sec12 is missing in the entire euglenozoan clade (**Fig. 1**). Although Sec16 was identified in *T. brucei* and *L. major* and its homolog was present in *B. saltans*, we did not find orthologs in euglenids or diplonemids by these bioinformatics methods (but see below).

Furthermore, all euglenozoans contain two paralogs of Sec13 (hereafter Sec13a and Sec13b) except for *B. saltans*, where only Sec13b was identified (**Fig. 1**). However, because of low statistical support at the backbone of the phylogenetic tree (**Fig. 2A**), we performed the approximately unbiased (AU) test constraining monophyly of all euglenozoan Sec13 paralogs consistent with a hypothesis of duplication of Sec13 at the base of the euglenozoan clade. Indeed, this alternative topology was not rejected (**Fig. 2B**). Significantly, in *P. papillatum* the Sec13 paralogs share only 28% sequence similarity (**Fig. 3A**), even though they retain a predicted β-propeller architecture, *albeit* with clear regions of predicted structural divergence (**Fig. 3B**). Next, we took advantage of the recently established genetic modification system (Faktorová et al., 2020a; Faktorová et al., 2020b) to investigate Sec13 function in *P. papillatum*.

**Fig. 2.**
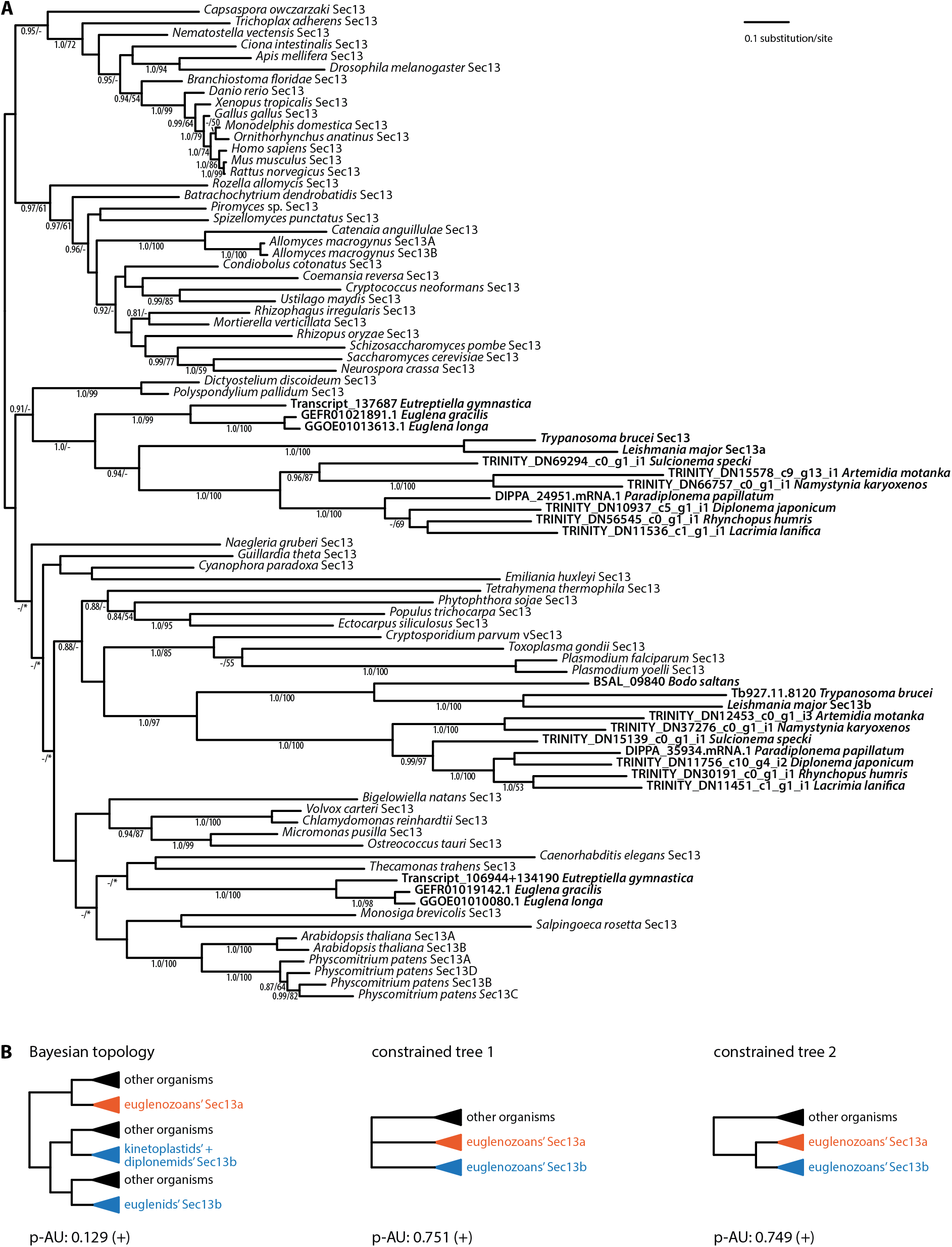
Phylogenetic analyses of Sec13. **(A)** MrBayes topology of Sec13 phylogenetic tree is shown, onto which posterior probabilities (PP) and bootstrap support (BS) values from RAxML are overlayed. Support values for PP < 0.8 and BS < 50% are denoted by a dash (-), whereas an asterisk (*) marks a topology that was not retrieved in a particular analysis. **(B)** Alternative topologies constraining monophyly of euglenozoan Sec13 are shown together with p-value. Sign + indicates that the topology was not rejected by the AU test.

**Fig. 3.**
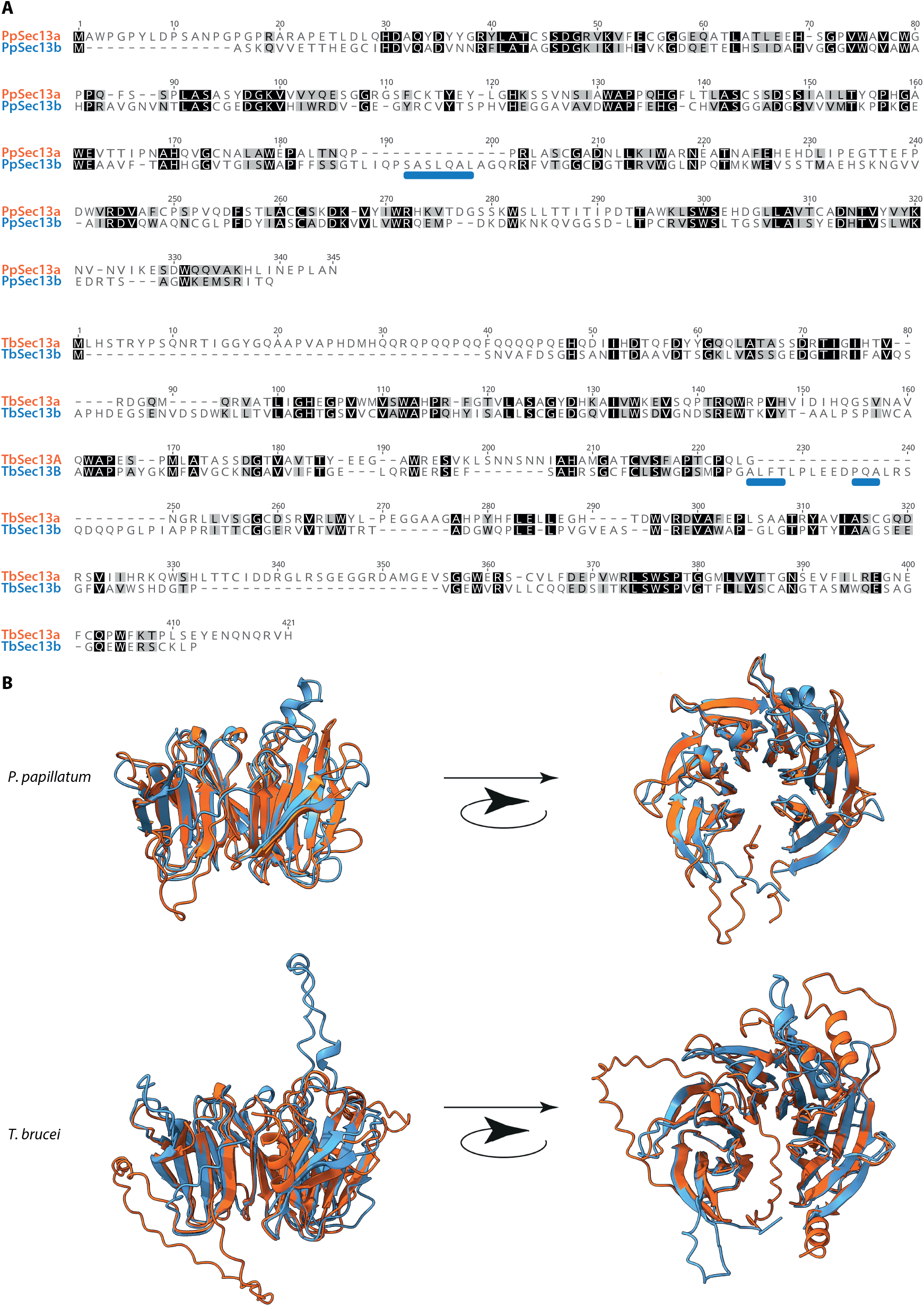
Structural divergence between Sec13a and Sec13b. **(A)** Alignments of Sec13 paralogs for *P. papillatum* and *T. brucei*. **(B)** AlphaFold predictions for Sec13a and Sec13b (colors correspond to **Fig. 1**) for *P. papillatum* and *T. brucei* to illustrate the positions of indel peptide sequences (marked by blue rectangles in A). Note that the positions of indels are distinct, but likely provide significantly unique binding sites to discriminate between protein interactions ad hence complex association.

### Localization of PpSec13a in *Paradiplonema papillatum*

The PpSec13a gene of *P. papillatum* was tagged to determine the intracellular localization and the identify of interaction partners for PpSec13a. Plasmid pDP002 (Faktorová et al., 2020a), containing a protein A (PrA) tag and a neomycin as a selectable marker was used for endogenous tagging at the C-terminus (see Methods and **Suppl. Table 3**). Expression of PrA-tagged PpSec13a was verified by immunoblotting (**Fig. 4A**). We used indirect immunofluorescence microscopy to determine the localization of PpSec13a using an anti-PrA antibody. This revealed a punctate pattern in the cytoplasm and around the nuclear envelope (**Fig. 5A**), suggesting the presence of PpSec13a in both the ER and the NPC, respectively.

**Fig. 4.**
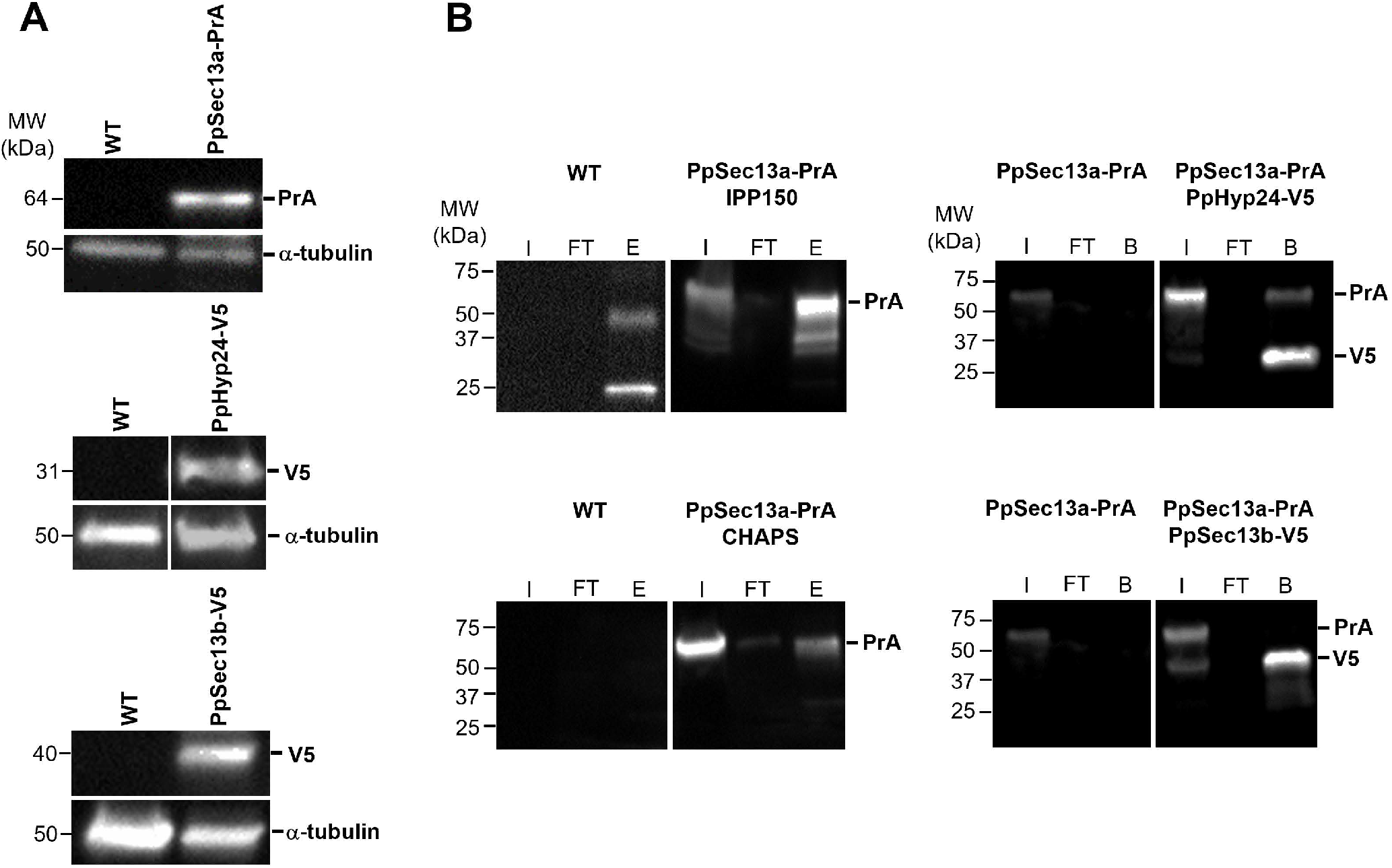
Immunoblot analyses and immunoprecipitations of Pr-A or V5-tagged cell lines. **(A)** From top to bottom: PpSec13a-PrA, PpHyp24-V5 and PpSec13b-V5. Rabbit anti-prA (1:10,000) or mouse anti-V5 (1:2,000) antibodies were used for verification of expression of tagged proteins. Mouse anti-α-tubulin (1:5,000) was used as a loading control. **(B)** Immunoprecipitation of PrA- or V5-tagged cell lines using following conditions: PpSec13a-PrA (IPP150 buffer) + IgG Sepharose beads; PpSec13a-PrA + PpHyp24-V5 (IPP150 buffer) + V5-trapped magnetic beads; PpSec13a-PrA (CHAPS buffer) + IgG Sepharose beads and PpSec13a-PrA + PpSec13b-V5 (IPP150 buffer) + V5-trapped magnetic beads. Wild type (WT) strain (for left panels) or PpSec13a-PrA (for right panels) were used as a controls. I, input; FT, flow-through; E, eluate; B, beads.

**Fig. 5.**
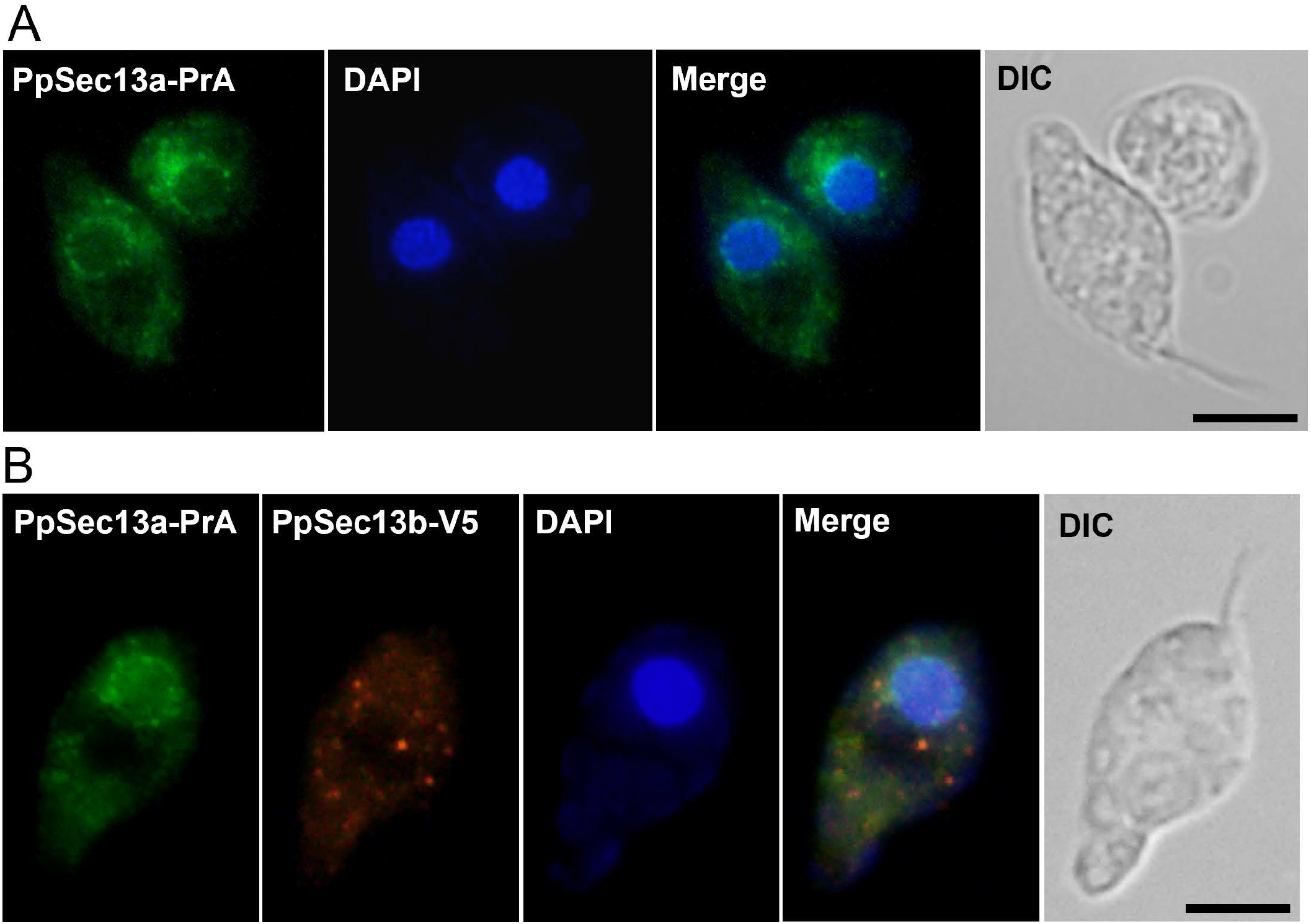
Localization of PpSec13a-PrA (A) and of PpSec13a-PrA + PpSec13b-V5 (B) proteins in *P. papillatum*. Immunofluorescence assay of PrA-tagged PpSec13a (A) and PrA-tagged PpSec13a together with V5-tagged PpSec13b cells (B), stained with polyclonal anti-PrA antibodies (green) or monoclonal anti-V5 antibodies (red). DNA was stained with DAPI (blue). DIC, Differential interference contrast. Scale bar is 5 μm.

### Interacting partners of PpSec13a::PrA

To purify PpSec13a together with its interacting partners, we performed immunoisolations with PpSec13a::PrA-tagged cell lysates with parental/wild type (WT) cell lysates serving as a control. Immunoblot analysis confirmed the presence of PpSec13a-PrA in the eluate fraction following immunoprecipitation (IP) (**Fig. 4B**). Subsequent analysis by mass spectrometry (MS) identified PpSec31 (DIPPA_09941) as well as two hypothetical proteins with so far unknown function that we name here based on their predicted molecular weights PpHyp24 (DIPPA_09665) and PpHyp27 (DIPPA_31723) (**Fig. 6; Suppl. Fig. 2A,B; Suppl. Table 4**). To confirm these interactions, we tagged the PpHyp24 gene to perform a reciprocal IP. Plasmid pDP011 was designed with a 3xV5 tag and the hygromycin B resistance gene (**Suppl. Fig. 3**) as a backbone for PCR amplification (**Suppl. Table 3**) and electroporated into *P. papillatum*. The expression of V5-tagged PpHyp24 in PpSec13::PrA cells was verified by immunoblotting (**Fig. 4A**).

**Fig. 6.**
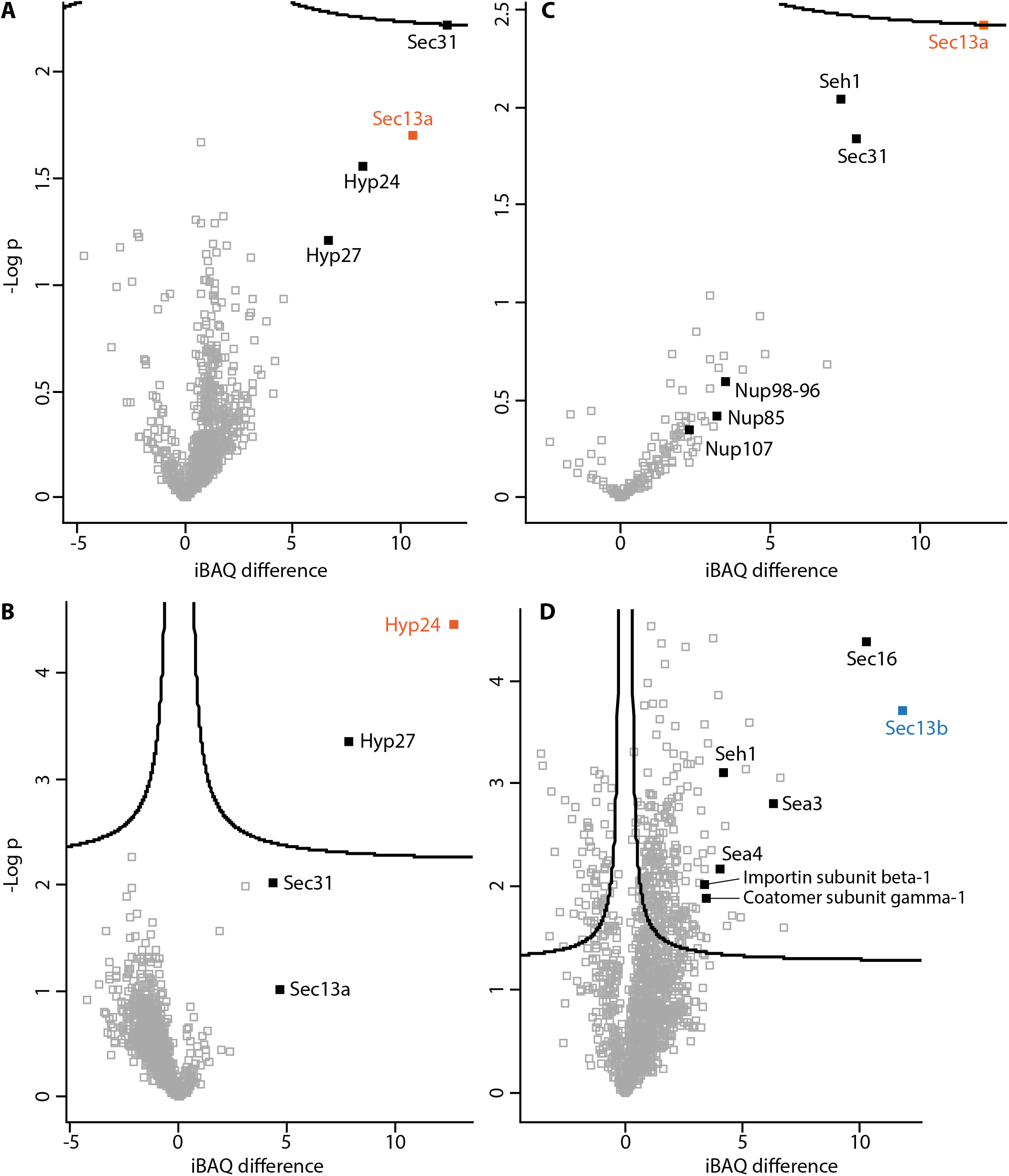
Volcano plots of analyzed proteins. **(A)** PpSec13a-PrA (IPP150 buffer) + IgG Sepharose beads; **(B)** PpSec13a-PrA + PpHyp24-V5 (IPP150 buffer) + V5-trapped magnetic beads; **(C)** PpSec13a-PrA (CHAPS buffer) + IgG Sepharose beads; **(D)** PpSec13a-PrA + PpSec13b-V5 (IPP150 buffer) + V5-trapped magnetic beads. The tagged proteins are show in bold, colors correspond to **Fig. 1**.

Immunoprecipitates of PpHyp24-tagged cell lysates were prepared with V5-derivatized magnetic beads and subjected to liquid chromatography-tandem mass spectrometry (LC-MS/MS). The presence of PpHyp24::V5 in the bound fraction was verified by immunoblot analysis, which also showed the presence of PpSec13a::PrA (**Fig. 4B**). LC-MS/MS of the IP revealed enrichment of all four proteins previously identified by PpSec13a::PrA immunoisolation, validating the composition of this complex (**Fig. 6; Suppl. Table 4**).

The observed immunofluorescence of PpSec13a was consistent with NPC localization (**Fig. 5A**). To confirm this, we used essentially the procedure described above but with a buffer successfully used for immunoisolation of the NPC from the related flagellate *T. brucei* (Obado et al., 2016). Following verification by immunoblotting (**Fig. 4B**), these immunoprecipitates were subjected to LC-MS/MS, which confirmed the presence of PpSec31, whereas PpHyp24 and PpHyp27 were not detected. Moreover, the modified conditions also identified three NPC components, namely PpNup107, PpNup98-96, PpNup85 and PpSeh1 (**Fig. 6**; **Suppl. Tables 4 and 5; Suppl. Fig. 2C**).

### Localization and interacting partners of PpSec13b

To identify interacting partners of the second Sec13 paralog PpSec13b, we again used the pDP011 plasmid for tagging (**Suppl. Table 3**). Similarly, as with the PpHyp24::V5 cell line, we established a cell line expressing both PpSec13b::V5 and PpSec13a::PrA. Expression of PpSec13b::V5 was confirmed by immunoblot analysis (**Fig. 4A**). We determined the localization of both PpSec13a and PpSec13b by performing double labeling immunofluorescence using anti-PrA and anti-V5 antibodies, respectively (**Fig. 5B**). Interestingly, the localizations of PpSec13a and PpSec13b were distinct, and PpSec13b was confined to the cytoplasm with a punctuate distribution (**Fig. 5B**).

The co-immunoprecipitation (co-IP) using V5-trapped magnetic beads also indicated lack of PpSec13a and PpSec13b interaction (**Fig. 4B**). Analysis of the PpSec13b IP revealed enrichment of DIPPA_10282, DIPPA_06252 and DIPPA_21857 gene products in addition to the affinity handle (**Fig. 6; Suppl. Table 4**). DIPPA_10282 is a Sec16 ortholog and, using DIPPA_10282 as query, we identified orthologs in other diplonemid (**Suppl. Table 2**) and euglenid transcriptomes, which by phylogenetic analysis form a robust clade with the kinetoplastids (**Suppl. Fig. 1E**). Given the identification of this divergent PpSec16, we revisited the distribution of Sec16 across eukaryotes, identifying orthologs in the rotoshaerid *Fonticula alba*, the red alga *Cyanidioschyzon merolae*, the apicomplexan parasites *Cryptosporidium parvum, Babesia bovis* and *Theileria equi* and multiple haptophytes (**Suppl. Table 6**), indicating that while still poorly retained Sec16 is present more broadly than thought previously. BLAST searches of DIPPA_21857 and orthologs from other euglenozoans revealed DIPPA_21857 as Sea4. However, Sea2, Sea3 and Sea4 are themselves paralogs and BLAST searches of DIPPA_06252 and euglenozoan orthologs identified these proteins as components of the Seh1-associated (SEA)/GATOR complex but without sufficient differences in E-values to discriminate between Sea2, Sea3 or Sea4 (**Suppl. Table 7**). Moreover, we identified an additional Sea2 gene in the genomes and transcriptomes of euglenozoans (DIPPA_06078 in *P. papillatum*) (**Fig. 1; Suppl. Table 7**), suggesting that DIPPA_06252 (and the euglenozoan orthologs) represents Sea3.

Phylogenetic analyses of euglenozoan and selected eukaryote sequences clearly distinguished Sea2, Sea3 and Sea4 clades (**Suppl. Fig. 2D**), but paralogs of kinetoplastid Sea2 and Sea3 were not reconstructed within any of these clades. We therefore used the kinetoplastid candidates as queries in reverse BLASTs against *P. papillatum*. In nearly every case this confirmed their initial annotations as Sea2 and Sea3 (**Suppl. Table 7**). In animals and fungi, Sea2, Sea3 and Sea4 are components of the SEA/GATOR complex and regulate the Target of Rapamycin Complex (TORC) signaling pathway involved in controlling cell growth, stress responses and other processes. These complexes are located within the endosomal system, and hence their identification is consistent with the localization of PpSec13b.

### Ultrastructure and organization of *P. papillatum* organelles

To understand the organization of the endomembrane system in *P. papillatum* and possible distribution of PpSec13 paralogs, we analyzed its ultrastructure by transmission electron microscopy (**Fig. 7**). Prominent structures identified include the flagellar pocket, cytopharynx, array of subpellicular microtubules, terminal endosomes/lysosomes and a large nucleus with prominent nuclear pores, nucleolus, and regions of heterochromatin (**Fig. 7A**).

**Fig. 7.**
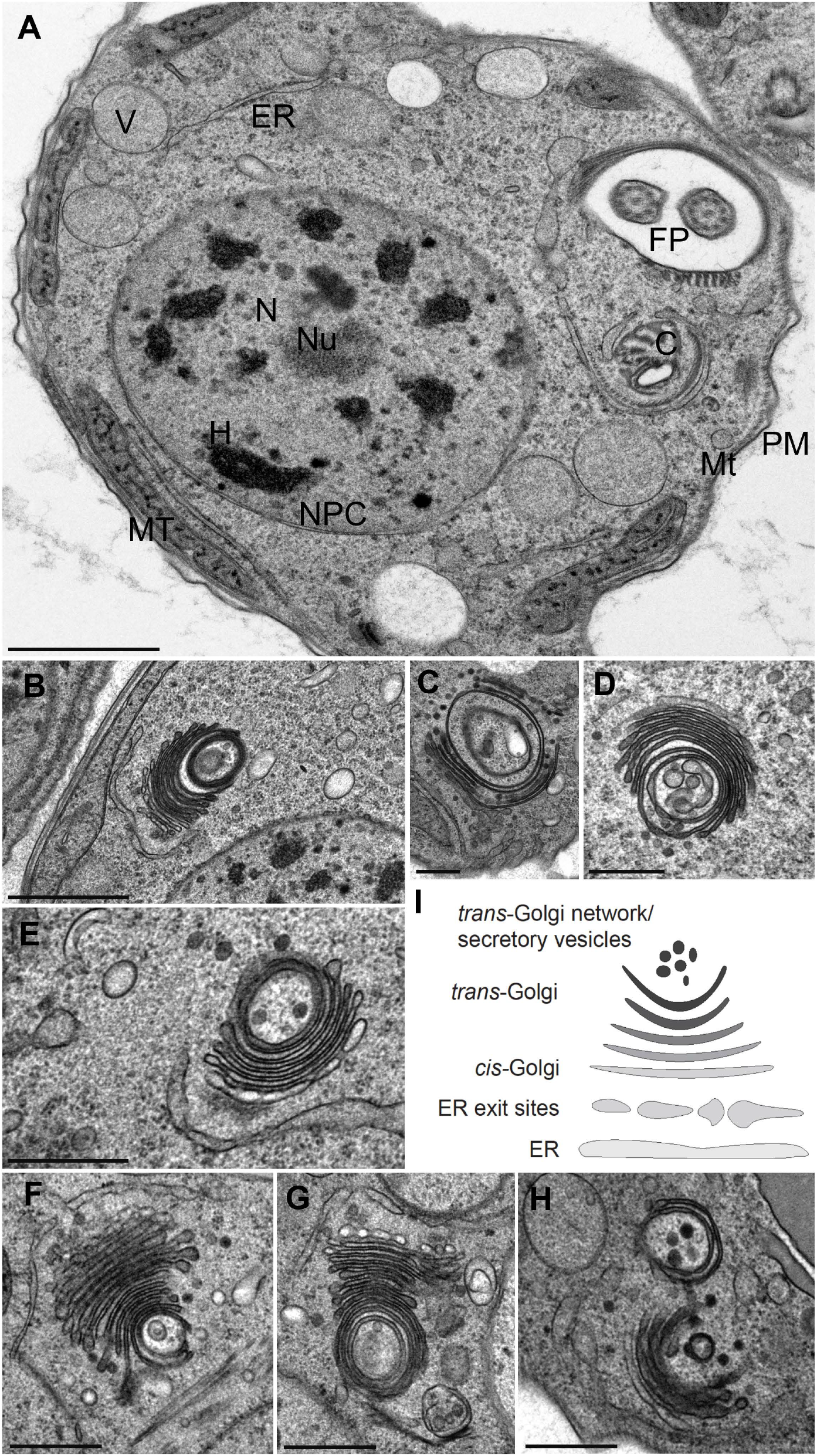
Transmission electron micrographs of *P. papillatum*. **(A)** Cross section of the whole cell showing position of subcellular structures and organelles. N, nucleus; Nu, nucleolus; H, heterochromatin; NPC, nuclear pore complex; PM, plasma membrane; Mt, microtubules below PM; MT, mitochondrion; ER, endoplasmic reticulum; V, vacuole; C, cytostome; FP, flagellar pocket. **(B - H)** Details of various Golgi complex in *P. papillatum*. **(I)** The scheme of ER-Golgi structure in *P. papillatum*. Scale bars: 1 μm (A–D), and 500 nm (E-H).

Moreover, we observed a complex series of additional membrane-bound structures, which include ER, visualized as tubular vesicles of varying length associated with the Golgi complex (**Fig. 6B-I**). Endosome-like structures, manifested frequently as vesicles within vesicles, may represent either true endosomes, multivesicular bodies or autophagosomes. We also noted that the Golgi complex is particularly prominent and is structurally highly coordinated with the ER and ER exit sites (ERES). This organelle is highly distinctive, with an extremely concave morphology that includes circular *trans-most* cisternal profiles and which are associated with vesicular structures also harboring internal membrane profiles. Significantly, the *cis*-Golgi cisternae are associated with a putative ER tubule, which may be the location of ERES. As the cisternae progress from the *cis* to *trans* faces, their content becomes increasingly electron dense, reflecting concentration of cargo for export to the cell surface. In some sections (**Fig. 6D,E,H**) we observe likely transport vesicles of ~10 nm diameter associated with the Golgi complex and of similar electron density to the *trans-* cisternal compartments, which we propose represent vesicles destined for the surface.

The morphology clearly demonstrates the presence of a highly organized ERES, a stacked Golgi complex and the NPCs. The organization of the anterograde pathway is quite striking with a clear high order organization between the ERES and the *cis*-face of the Golgi complex. Moreover, it indicates that the expected compartments and transport events predicted by affinity isolation are supported by the presence of the pre-requisite organelles. Specifically, a stacked Golgi complex (Sec16), ERES (Sec31), endosomal degradative compartments (SEA/GATOR) and the NPC are all clear and present, also consistent with the contributions of PpSec13 paralogs as observed at the light level; specifically punctate nuclear rim staining and cytoplasmic puncta that likely correspond to the ERES, possible Golgi association and endosomes. Furthermore, the highly organized architecture of the *P. papillatum* cell may also suggest a significant level of precision with respect to regulation and organellar morphology in these protists.

## Discussion

Evolution and diversification represent an ongoing process, and for eukaryogenesis began over a billion years ago (Eme et al., 2017; Koonin, 2015; Roger et al., 2021). The eukaryotic cellular plan achieved considerable complexity by the time of the LECA and continues to be modified in extant lineages. Part of the machinery facilitating evolution of multiple compartments arose through expansion of paralog families (Dacks and Field, 2007; Devos et al., 2004). Here we demonstrate that Sec13, a component of at least three protocoatomer complexes, achieved unique diversity within the Euglenozoa, an early diverging lineage, with two Sec13 paralogs, each of which has distinct subfunctions. This is supported by interactome proteomics and localization using epitope-tagged gene products. Given that reconstructions of the LECA cannot always determine if there were more than one paralog of a gene present, we cannot determine if the Euglenozoa represent a lineage specific expansion, or secondary loss of one Sec13 paralog in the main eukaryotic lineage, shortly following divergence early in post-LECA evolution.

Significantly, Sec13a has been analyzed quite extensively in *Trypanosoma brucei*, and is a component of the NPC, localized into the ERES (DeGrasse et al., 2009). Similarly, TbSec13a possesses a nuclear localization signal, while TbSec13b prominently lacks it. In *T. brucei* immunoisolation of TbSec13a does not capture TbSec16 or the SEA/GATOR complex components (Obado et al., 2016). Consistently, in *P. papillatum* we identified PpSec16 and Sea proteins only from immunoisolations of PpSec13b but not PpSec13a, suggesting a conserved distinction between Sec13a and Sec13b across the Euglenozoa.

The arrangement and coordination of the ERES and NPC transport is clearly divergent in trypanosomes and related flagellates. The lineage is characterized by a cell surface dominated by lipid-anchored proteins and glycoconjugates. The two paralogs of Sec23 and Sec24 form distinct heterodimers of TbSec23b/TbSec24a and TbSec23a/TbSec24b (TbSec24a corresponds to LECA Sec24I and TbSec24b to LECA Sec24II) and, importantly, TbSec23b/TbSec24a specifically mediates anterograde transport of GPI-anchored proteins (Sharif and Bangs, 2022), indicating a complex mechanism regulating ER export. Further, Sec12, the guanine nucleotide exchange factor for Sar1 in yeasts and animals is absent (Wang et al., 2010), while there is considerable heterogeneity in the Sar1/SarB paralog cohort across the Euglenozoa in general. TbSec31 is regulated by the cell cycle-dependent kinase CDK1 (Hu et al., 2016) and, significantly, both replication of the Golgi complex and the ERES is highly coordinated with entry to G1 phase of the cell cycle. Finally, direct interaction between Sec13 and Sec16 indicates a role for the latter in both anterograde transport, as well as in the maintenance of Golgi morphology (Sealey-Cardona et al., 2014), and given the highly organized *P. papillatum* Golgi complex, we speculate that Sec16 may have a similar role in this protist. Indeed, ultrastructurally very prominent Golgi complexes have been described in other diplonemids, namely *Rhynchopus humris*, *Lacrimia lanifica*, *Sulcionema specki, Flectonema neradi* (Tashyreva et al., 2018) and *Namystynia karyoxenos* (Prokopchuk et al., 2019). It is worth to mention here that based on recent environmental DNA sequencing, diplonemids represent the most diverse and 5^th^ most abundant marine protists (Tashyreva et al., 2022), which shows their enormous success in the extant oceans.

Overall, while the parasitic trypanosomes are distinct from the free-living diplonemids, conservation of a complex COPII/Sec13 protein cohort suggests likely cell cycle and cargo-dependent coordination of anterograde function. We speculate that the need to maintain distinct aspects of this critical part of the cell may explain the retention and division of labor of two Sec13 paralogs.

## Material and Methods

### Sequence searches, structure predictions, and phylogenetic analyses

COPII subunits were retrieved by BLAST+ v2.8.1 (Altschul et al., 1990) searches using discobid sequences (Schlacht and Dacks, 2015) as queries in the genome of *Paradiplonema papillatum* (our unpublished data), transcriptomes of diplonemids *Diplonema japonicum*, *Rhynchopus humris*, *Lacrimia lanifica*, *Sulcionema specki*, *Artemidia motanka*, and *Namystynia karyoxenos* (Kaur et al., 2020), genome of the kinetoplastid *Bodo saltans* (Jackson et al., 2016), and transcriptomes of the euglenids *Euglena longa* (Záhonová et al., 2018) and *Eutreptiella gymnastica* (Keeling et al., 2014; reassembly available at https://doi.org/10.5281/zenodo.257410). To identify more divergent sequences, more sensitive searches by HMMER v3.3 based on profile hidden Markov models (Eddy, 2009) were performed in parallel. Protein domains were predicted by InterProScan (Jones et al., 2014) implemented in Geneious v2020.2.5 (Kearse et al., 2012). Secondary structures of proteins were de novo predicted by AlphaFold2 (Jumper et al., 2021), visualized and overlayed by ChimeraX v1.4 (Pettersen et al., 2021).

Datasets of COPII components were obtained from previous studies (DeGrasse et al., 2009; Schlacht and Dacks, 2015; Vargová et al., 2021). As PpHyp27 did not retrieve any orthologs in reverse BLAST searches against NCBI, both PpHyp24 and PpHyp27 were subjected to more sensitive profile searches using HHBlits (Remmert et al., 2012) and best hits served as datasets for phylogenetic analyses. Found sequences were added to the datasets, aligned by MAFFT v7.458 under L-INS-i strategy (Katoh and Standley, 2013), and poorly aligned positions were removed by trimAl using -gt 0.8 option (Capella-Gutiérrez et al., 2009). Maximum likelihood (ML) phylogenetic analysis was performed by RAxML v8.2.8 (Stamatakis, 2014) under the LG4X mixture model using 100 rapid bootstrap replicates (-f a). Bayesian inference was performed by MrBayes v3.2.7a (Ronquist et al., 2012) under the LG4X mixture model, with at least 10 million Markov Chain Monte Carlo generations and 4 gamma rate categories. Sampling frequency was set to every 1,000 generations with the first 25% of the runs discarded as burn-in. Bootstrap support values were overlaid onto the MrBayes tree topology with posterior probabilities.

To test the monophyly of euglenozoan Sec13 sequences, the AU test was performed. We constructed ML trees using IQ-TREE v2.2.0 (Minh et al., 2020) under the LG+C20+F+G model constraining the monophyly of euglenozoan Sec13 using -g option. The AU test was ran in IQ-TREE on the constrained trees, the topology retrieved from the Bayesian analysis, and 1,000 distinct local topologies saved during ML analysis. Topologies that returned an AU-test p-value<0.05 were rejected.

### Strain and cultivation of *P. papillatum*

*P. papillatum* (ATCC 50162) was cultivated axenically at 27 °C in an artificial sea salt medium as described previously (Faktorová et al., 2020b) and cell density was measured manually using the Neubauer cell chamber.

### Endogenous C-terminal gene tagging and used cell lines

All cassettes were designed and prepared by fusion PCR approach using Phusion polymerase (NEB Biolabs, M0530S) as described elsewhere (Faktorová et al., 2020b). For PrA tagging, PrA+Neo^R^ cassette was amplified from pDP002 plasmid, while for V5 tagging 3xV5+Hyg^R^ cassette was amplified from newly designed pDP011 plasmid (**Suppl. Fig. 3**). Used primers and PCR product sizes are listed in **Suppl. Table 3.** The gel-purified cassettes were ethanol-precipitated and sequentially electroporated into the *P. papillatum* cells (Faktorová et al., 2020b; Kaur et al., 2018). For transformation, a total of 5 x 10^7^ cells were harvested and electroporated with appropriate DNA constructs (cassettes; see below) as described previously (Faktorová et al., 2020b).

Eighteen to 24 hours after electroporation, transfectants were subjected to selection in 24 well plates at 27 °C with increasing concentrations of either G418 (25–80 μg.ml^-1^) for establishing the PpSec13a-PrA cell line, or hygromycin (100–225 μg.ml^-1^) for creating the PpHyp24-V5 cell line, or both (G418 and hygromycin) at the same time for establishing the PpSec13a-PrA+PpSec13b-V5 double-tagged cell line. After two to three weeks, successful transfectants were obtained and each clone was expanded to a volume of 20 ml. The expression and verification of expected size of the tagged proteins in obtained resistant cell lines was verified by immunoblot analysis.

In this study we selected and further used the following cell lines: 1/ PpSec13a-PrA+Neo^R^ cell line growing in media supplemented with 67.5 μg.ml^-1^ G418; 2/ PpHyp24-V5+Hyg^R^ growing in media supplemented with 175 μg.ml^-1^ of hygromycin and 3/ PpSec13a-PrA+Neo^R^/PpSec13b-V5+Hyg^R^ growing in media supplemented with 67.5 ug.ml^-1^ G418 and 100 μg.ml^-1^ of hygromycin.

### Immunoblot analysis

Immunoblot (western blot) analysis of *P. papillatum* samples was performed as described previously (Faktorová et al., 2020b). Membranes were first incubated with rabbit anti-PrA (1:10,000; Sigma-Aldrich, P3775) or mouse anti-V5 (1:2,000; ThermoFisher Scientific, 37-7500) primary antibodies at 4 °C overnight or at room temperature for 2 hours. After five washes in phosphate buffered saline supplemented with Tween (PBS-T; 0.05% (v/v) Tween in 1× PBS), membranes were incubated with HRP-coupled anti-rabbit (1:1,000; Sigma-Aldrich, A21428) or anti-mouse (1:1,000; Sigma-Aldrich, A9044) secondary antibodies at room temperature for 1 hour. Next, they were washed five times in PBS-T, and the signal was developed using Clarity Western ECL Substrate (Bio-Rad). The anti-α-tubulin antibody (produced in mouse; 1:10,000; Sigma-Aldrich, T9026) was used as a loading control.

### Immunofluorescence assay

20-30 ml of a log phase culture was centrifuged at 1,000 g for 5 min in order to visualize localization of PpSec13a and/or PpSec13b in *P. papillatum*. Cells were resuspended in 500 μl of 4% paraformaldehyde (dissolved in sea water) and fixed for 20 min on Superfrost plus slides (Thermo Scientific, J1800AMNZ) at room temperature. The fixative was washed out from cells with 1× PBS. For antibody staining, cells were permeabilized in ice-cold methanol for 20 min. The slides were kept in a humid chamber throughout the procedure. Afterwards, the slides were washed with 1× PBS, and blocked for 45 min in 5.5% (w/v) fetal bovine serum in PBS-T. The blocking solution was removed, and cells were washed with 1× PBS. Rabbit anti-PrA (1:2,000; Sigma, P3775) and/or mouse anti-V5 (1:150; ThermoFisher Scientific, 37-7500) primary antibodies diluted in 3% (w/v) BSA (Bovine serum albumin, Sigma, A4503) in PBS-T was added on slides and incubated either for 2 hours at room temperature or at 4 °C overnight, covered with parafilm. Next, the primary antibody was removed, and slides were washed three times with PBS-T and twice with 1× PBS. AlexaFluor488-labeled goat anti-rabbit (1:1,000; Invitrogen, A11034) and/or AlexaFluor555-labeled goat anti-mouse (1:1,000; Invitrogen, A21422) secondary antibody was added and incubated at room temperature for 1 hour in the dark, covered with parafilm. All slides were then rinsed three times with PBS-T and twice with 1× PBS and coated with 4’,6-diamidino-2-phenylindole (DAPI) containing the antifade reagent ProlongGold (Life Technologies). Images were acquired using an Olympus BX63 automated fluorescence microscope equipped with an Olympus DP74 digital camera and evaluated with cellSens Dimension software (Olympus).

### Immunoprecipitation

For prA-tagged cell line, 5 x 10^8^ cells (PpSec13a-PrA+Neo^R^) were grown at 27 °C in media with selection antibiotics (see above). Cells were harvested at 1,000 g for 10 min and subsequently resuspended either in 1 ml ice cold IPP150 buffer (10 mM Tris-HCl pH 6.8, 150 mM NaCl, 0.1% IGEPAL CA-630; Sigma I8896) or in CHAPS buffer (20 mM HEPES pH 7.4, 100 mM NaCl, 0.5 % CHAPS; Roche 10810118001), both supplemented with 1× cOmplete EDTA-free protease inhibitors (Sigma, 11873580001) and five times passed through a 30-gauge needle. The lysate was subsequently cleared twice by centrifugation (12,000 g, 10 min, 4 °C) and the supernatant was incubated with 75 μl of IgG Sepharose 6 Fastbeads (GE Healthcare, 52-2083-00 AH) by rotating at 4 °C for 2 to 3 hours to enable binding of the tagged protein. The beads were washed five times using the same buffer as for cell lysis and the complex of PpSec13a-PrA together with its potential interaction partners were eluted using 100 μl of 0.1 M glycine (pH 3.0) by rotating for 5 min at room temperature and immediately neutralized with 10 μl of 1 M Tris-HCl (pH 9.0). Aliquots of input, flow through and elution fractions were processed for immunoblotting. The elution fraction was subsequently sent for MS analysis.

For V5-tagged cell lines, 5 x 10^8^ cells (PpHyp24-V5+Hyg^R^ or PpSec13a-PrA+Neo^R^/PpSec13b-V5+Hyg^R^) were grown at 27 °C in media with appropriate selection antibiotics. Cells were harvested at 1,000 g for 10 min, resuspended in 1 ml ice cold IPP150 buffer and subsequently processed using similar protocol as above with the following exceptions: 1/ supernatant was incubated with 50 μl of V5-Trap magnetic Particles M-270 (Chromotek, v5td-20; the advantage of these beads is that there are no heavy and light antibody chains present in the bound fraction and therefore even proteins of 25/50kDa can be seen on immunoblot), 2/ last two washes of beads were done using a buffer without detergent and the beads were sent for MS analysis. For all IP experiments three replicates of each sample were processed by MS. Wild type cells were used as a control.

### Mass spectrometry and data analysis of immunoprecipitated samples

Trypsin-digestion of the eluted prA-tagged bait and wild type controls or the V5-paramagnetic beads incubated with V5-tagged bait and wild type controls was performed prior to LC-MS/MS as follows. IP samples were resuspended in 100 mM TEAB containing 2% SDC, and cysteines were reduced with 10 mM final concentration of TCEP and blocked with 40 mM final concentration of chloroacetamide (60 °C for 30 min). Samples were cleaved on beads with 1 μg of trypsin at 37 °C overnight. After digestion, samples were centrifuged, and supernatants were collected and acidified with TFA to 1% final concentration. SDC was removed by extraction with ethylacetate, and peptides were desalted using in-house made stage tips packed with C18 disks (Empore). Nano Reversed phase column (EASY-Spray column, 50 cm × 75 μm ID, PepMap C18, 2 μm particles, 100 Å pore size) was used for LC/MS analysis. Mobile phase buffer A was composed of water and 0.1% formic acid, and mobile phase B was composed of acetonitrile and 0.1% formic acid. Samples were loaded onto the trap column (Acclaim PepMap300, C18, 5 μm, 300 Å Wide Pore, 300 μm x 5 mm, 5 Cartridges) for 4 min at 15 μl.min^-1^. Loading buffer was composed of water, 2% acetonitrile and 0.1% trifluoroacetic acid. Peptides were eluted with Mobile phase B gradient from 4% to 35% B in 60 min. Eluting peptide cations were converted to gas-phase ions by electrospray ionization and analyzed on a Thermo Orbitrap Fusion (Q-OT-qIT, Thermo). Survey scans of peptide precursors from 350 to 1,400 m/z were performed at 120K resolution (at 200 m/z) with a 5× 10^5^ ion count target. Tandem MS was performed by isolation at 1,5 Th with the quadrupole, HCD fragmentation with normalized collision energy of 30, and rapid scan MS analysis in the ion trap. The MS/MS ion count target was set to 104 and the max injection time was 35 ms. Only those precursors with charge state 2–6 were sampled for MS/MS. The dynamic exclusion duration was set to 45 s with a 10 ppm tolerance around the selected precursor and its isotopes. Monoisotopic precursor selection was turned on and the instrument was run in top speed mode with 2 s cycles.

Data were processed using MaxQuant v1.6.14, which incorporates the Andromeda search engine (Cox et al., 2011). A custom protein sequence database of *P. papillatum* proteins (43,871 sequences) supplemented with frequently observed contaminants was used to identify proteins. Search parameters were the default ones employed by MaxQuant for Orbitrap analyzers with full trypsin specificity and allowing for up to two missed cleavages. Carbamidomethylation of cysteine was set as a fixed modification and oxidation of methionine and N-terminal protein acetylation were allowed as variable modifications. Match between runs for biological replicates was part of the experimental design. Peptides were required to be at least seven amino acids long, with false discovery rates of 0.01 calculated at the levels of peptides, proteins, and modification sites based on the number of hits against the reversed sequence database. iBAQ indices (raw intensities divided by the number of theoretical peptides) were used for protein quantification, which allows comparing of protein abundances both within samples and between them. After filtering to remove any protein IDs with less than two unique peptides, an Andromeda score of less than 20 and less than two valid values in the respective bait replicates, the obtained data were processed in Perseus v1.6.14 as described previously (Zoltner et al., 2020).

### Transmission electron microscopy

Transmission electron microscopy samples were prepared by high pressure freezing technique (HPF) as described previously (Yurchenko et al., 2014). Ultrathin sections were observed using a JEOL 1010 microscope at accelerating voltage of 80 kV. Images were captured with an Olympus Mega View III camera.

## Supporting information

Supplementary data - all

## Acknowledgements

We thank Binnypreet Kaur (Texas A&M University) for help in the initial stages of this project, Matus Valach and Gertraud Burger (University of Montreal) for help with codon optimalisation of the pDP011 construct. We also acknowledge Karel Harant and Pavel Talacko (Biocev, Prague) for mass spectrometry analysis, Daria Tashyreva, Martina Tesařová and Galina Prokopchuk (Institute of Parasitology) for discussions and help with transmission electron microscopy, and Courtney W. Stairs (Lund University) for help with the AU test.

## Data availability

All data are contained within manuscript and its supplementary material. The DNA sequence of pDP011 was deposited in GenBank under the XXX accession number. The mass spectrometry proteomics data have been deposited to the ProteomeXchange Consortium via the PRIDE (Perez-Riverol et al., 2022) partner repository with the dataset identifier PXD037122.

## Author Contributions

D.F.: construct design; cell cultivation; cassettes amplification, electroporation and cell lines selection; immunoblotting; fluorescence microscopy; writing of the initial manuscript draft; C.B. Mass spec analysis; D.F., K.Z., C.B., M.C.F.: visualization; K.Z, J.B.D., M.C.F.: bioinformatic and phylogenetic analysis; K.Z, C.B., M.C.F., J.B.D., J.L.: writing – review and editing; J.L.: project administration; supervision; funding acquisition; resources.

## Supplementary Material Legends

**Supplementary Figure 1. Phylogenetic analyses of COPII components Sar1 and SarB (A), Sec23 (B), Sec24 (C), Sec31 (D), and Sec16 (E).** MrBayes topology of phylogenetic tree is shown, onto which posterior probabilities (PP) and bootstrap support (BS) values from RAxML are overlayed. Support values for PP < 0.8 and BS < 50% are denoted by a dash (-), whereas an asterisk (*) marks a topology that was not retrieved in a particular analysis.

**Supplementary Figure 2. Phylogenetic analyses of hypothetical proteins PpHyp24 (A) and PpHyp27 (B), NUP component Seh1 (C), and SEA/GATOR components Sea2, Sea3, and Sea4 (D).** MrBayes topology of phylogenetic tree is shown, onto which posterior probabilities (PP) and bootstrap support (BS) values from RAxML are overlayed. Support values for PP < 0.8 and BS < 50% are denoted by a dash (-).

**Supplementary Figure 3.** Scheme of pDP011 plasmid.

**Supplementary Table 1.** List of COPII genes identified in *P. papillatum* genome.

**Supplementary Table 2.** List of COPII components found in studied diplonemids and their reverse best hits.

**Supplementary Table 3.** List of primers used in this study. Sequences in bold correspond to parts aligning to pDP002 (A) or pDP011 (B and C) plasmid.

**Supplementary Table 4.** Mass spectrometry (MS) results from triplicate experiments for PpSec13a-PrA (IPP150 buffer) (A), PpHyp24-V5 (IPP150 buffer) (B), PpSec13a-PrA (CHAPS buffer) (C), PpSec13b-V5 (IPP150 buffer) (D). For whole MS data see **Suppl. Data 1**.

**Supplementary Table 5.** List of Seh1, Nup85, Nup98, and Nup107 proteins in studied diplonemids and their reverse best hits.

**Supplementary Table 6.** Identified Sec16 proteins in organisms in EukProt database.

**Supplementary Table 7.** List of Sea2, Sea3, and Sea4 proteins in studied euglenozoans and their reverse best hits.

**Supplementary Data 1.** Whole MS data.

## References

Altschul, S. F., Gish, W., Miller, W., Myers, E. W. and Lipman, D. J. (1990). Basic local alignment search tool. J. Mol. Biol. 215, 403–410.

Barlowe, C., Orci, L., Yeung, T., Hosobuchi, M., Hamamoto, S., Salama, N., Rexach, M. F., Ravazzola, M., Amherdt, M. and Schekman, R. (1994). COPII: A membrane coat formed by Sec proteins that drive vesicle budding from the endoplasmic reticulum. Cell 77, 895–907.

Berriman, M., Ghedin, E., Hertz-Fowler, C., Blandin, G., Renauld, H., Bartholomeu, D. C., Lennard, N. J., Caler, E., Hamlin, N. E., Haas, B., et al. (2005). The genome of the African trypanosome Trypanosoma brucei. Science 309, 416–422.

Capella-Gutiérrez, S., Silla-Martínez, J. M. and Gabaldón, T. (2009). trimAl: a tool for automated alignment trimming in large-scale phylogenetic analyses. Bioinformatics 25, 1972–1973.

Carlton, J. M., Hirt, R. P., Silva, J. C., Delcher, A. L., Schatz, M., Zhao, Q., Wortman, J. R., Bidwell, S. L., Alsmark, U. C. M., Besteiro, S., et al. (2007). Draft genome sequence of the sexually transmitted pathogen Trichomonas vaginalis. Science 315, 207–212.

Cox, J., Neuhauser, N., Michalski, A., Scheltema, R. A., Olsen, J. V. and Mann, M. (2011). Andromeda: A peptide search engine integrated into the MaxQuant environment. J. Proteome Res. 10, 1794–1805.

Dacks, J. B. and Field, M. C. (2007). Evolution of the eukaryotic membrane-trafficking system: Origins, tempo and mode. J. Cell Sci. 120, 2977–2985.

Dacks, J. B. and Field, M. C. (2018). Evolutionary origins and specialisation of membrane transport. Curr. Opin. Cell Biol. 53, 70–76.

DeGrasse, J. A., Dubois, K. N., Devos, D., Siegel, T. N., Sali, A., Field, M. C., Rout, M. P. and Chait, B. T. (2009). Evidence for a shared nuclear pore complex architecture that is conserved from the last common eukaryotic ancestor. Mol. Cell. Proteomics 8, 2119–2130.

Demmel, L., Melak, M., Kotisch, H., Fendos, J., Reipert, S. and Warren, G. (2011). Differential selection of golgi proteins by COPII Sec24 isoforms in procyclic Trypanosoma brucei. Traffic 12, 1575–1591.

Devos, D., Dokudovskaya, S., Alber, F., Williams, R., Chait, B. T., Sali, A. and Rout, M. P. (2004). Components of coated vesicles and nuclear pore complexes share a common molecular architecture. PLoS Biol. 2, e380.

Dokudovskaya, S., Waharte, F., Schlessinger, A., Pieper, U., Devos, D. P., Cristea, I. M., Williams, R., Salamero, J., Chait, B. T., Sali, A., et al. (2011). A conserved coatomer-related complex containing Sec13 and Seh1 dynamically associates with the vacuole in Saccharomyces cerevisiae. Mol. Cell. Proteomics 10, M110.006478.

Ebenezer, T. E., Zoltner, M., Burrell, A., Nenarokova, A., Novák Vanclová, A. M. G., Prasad, B., Soukal, P., Santana-Molina, C., O’Neill, E., Nankissoor, N. N., et al. (2019). Transcriptome, proteome and draft genome of Euglena gracilis. BMC Biol. 17, 11.

Eddy, S. R. (2009). A new generation of homology search tools based on probabilistic inference. Genome Inf. 23, 205–211.

Eme, L., Spang, A., Lombard, J., Stairs, C. W. and Ettema, T. J. G. (2017). Archaea and the origin of eukaryotes. Nat. Rev. Microbiol. 15, 711–723.

Faktorová, D., Nisbet, R. E. R., Fernández Robledo, J. A., Casacuberta, E., Sudek, L., Allen, A. E., Ares, M., Aresté, C., Balestreri, C., Barbrook, A. C., et al. (2020a). Genetic tool development in marine protists: emerging model organisms for experimental cell biology. Nat. Methods 17, 481–494.

Faktorová, D., Kaur, B., Valach, M., Graf, L., Benz, C., Burger, G. and Lukeš, J. (2020b). Targeted integration by homologous recombination enables in situ tagging and replacement of genes in the marine microeukaryote Diplonema papillatum. Environ. Microbiol. 22, 3660–3670.

Flegontov, P., Butenko, A., Firsov, S., Kraeva, N., Eliáš, M., Field, M. C., Filatov, D., Flegontova, O., Gerasimov, E. S., Hlaváčova, J., et al. (2016). Genome of Leptomonas pyrrhocoris: a high-quality reference for monoxenous trypanosomatids and new insights into evolution of Leishmania. Sci. Rep. 6, 23704.

Flegontova, O., Flegontov, P., Londoño, P. A. C., Walczowski, W., Šantić, D., Edgcomb, V. P., Lukeš, J. and Horák, A. (2020). Environmental determinants of the distribution of planktonic diplonemids and kinetoplastids in the oceans. Environ. Microbiol. 22, 4014–4031.

Fontoura, B. M. A., Blobel, G. and Matunis, M. J. (1999). A conserved biogenesis pathway for nucleoporins: Proteolytic processing of a 186-kilodalton precursor generates Nup98 and the novel nucleoporin, Nup96. J. Cell Biol. 144, 1097–1112.

Hu, H., Gourguechon, S., Wang, C. C. and Li, Z. (2016). The G1 cyclin-dependent kinase CRK1 in Trypanosoma brucei regulates anterograde protein transport by phosphorylating the COPII subunit Sec31. J. Biol. Chem. 291, 15527–15539.

Ivens, A. C., Peacock, C. S., Worthey, E. A., Murphy, L., Aggarwal, G., Berriman, M., Sisk, E., Rajandream, M. A., Adlem, E., Aert, R., et al. (2005). The genome of the kinetoplastid parasite, Leishmania major. Science 309, 436–442.

Jackson, A. P., Otto, T. D., Aslett, M., Armstrong, S. D., Bringaud, F., Schlacht, A., Hartley, C., Sanders, M., Wastling, J. M., Dacks, J. B., et al. (2016). Kinetoplastid phylogenomics reveals the evolutionary innovations associated with the origins of parasitism. Curr. Biol. 26, 161–172.

Jones, P., Binns, D., Chang, H. Y., Fraser, M., Li, W., McAnulla, C., McWilliam, H., Maslen, J., Mitchell, A., Nuka, G., et al. (2014). InterProScan 5: genome-scale protein function classification. Bioinformatics 30, 1236–1240.

Jumper, J., Evans, R., Pritzel, A., Green, T., Figurnov, M., Ronneberger, O., Tunyasuvunakool, K., Bates, R., Žídek, A., Potapenko, A., et al. (2021). Highly accurate protein structure prediction with AlphaFold. Nature 596, 583–589.

Katoh, K. and Standley, D. M. (2013). MAFFT multiple sequence alignment software version 7: improvements in performance and usability. Mol. Biol. Evol. 30, 772–780.

Kaur, B., Valach, M., Peña-Diaz, P., Moreira, S., Keeling, P. J., Burger, G., Lukeš, J. and Faktorová, D. (2018). Transformation of Diplonema papillatum, the type species of the highly diverse and abundant marine microeukaryotes Diplonemida (Euglenozoa). Environ. Microbiol. 20, 1030–1040.

Kaur, B., Záhonová, K., Valach, M., Faktorová, D., Prokopchuk, G., Burger, G. and Lukeš, J. (2020). Gene fragmentation and RNA editing without borders: Eccentric mitochondrial genomes of diplonemids. Nucleic Acids Res. 48, 2694–2708.

Kearse, M., Moir, R., Wilson, A., Stones-Havas, S., Cheung, M., Sturrock, S., Buxton, S., Cooper, A., Markowitz, S., Duran, C., et al. (2012). Geneious Basic: an integrated and extendable desktop software platform for the organization and analysis of sequence data. Bioinformatics 28, 1647–1649.

Keeling, P. J., Burki, F., Wilcox, H. M., Allam, B., Allen, E. E., Amaral-Zettler, L. A., Armbrust, E. V, Archibald, J. M., Bharti, A. K., Bell, C. J., et al. (2014). The Marine Microbial Eukaryote Transcriptome Sequencing Project (MMETSP): illuminating the functional diversity of eukaryotic life in the oceans through transcriptome sequencing. PLoS Biol. 12, e1001889.

Koonin, E. V. (2015). Origin of eukaryotes from within archaea, archaeal eukaryome and bursts of gene gain: Eukaryogenesis just made easier? Philos. Trans. R. Soc. B Biol. Sci. 370, 20140333.

Kruzel, E. K., Zimmett, G. P. and Bangs, J. D. (2017). Life stage-specific cargo receptors facilitate glycosylphosphatidylinositol-anchored surface coat protein transport in Trypanosoma brucei. mSphere 2, e00282–17.

Minh, B. Q., Schmidt, H. A., Chernomor, O., Schrempf, D., Woodhams, M. D., Von Haeseler, A., Lanfear, R. and Teeling, E. (2020). IQ-TREE 2: New models and efficient methods for phylogenetic inference in the genomic era. Mol. Biol. Evol. 37, 1530–1534.

Neumann, N., Lundin, D. and Poole, A. M. (2010). Comparative genomic evidence for a complete nuclear pore complex in the last eukaryotic common ancestor. PLoS One 5, e13241.

Obado, S. O., Brillantes, M., Uryu, K., Zhang, W., Ketaren, N. E., Chait, B. T., Field, M. C. and Rout, M. P. (2016). Interactome mapping reveals the evolutionary history of the nuclear pore complex. PLoS Biol. 14, e1002365.

Perez-Riverol, Y., Bai, J., Bandla, C., García-Seisdedos, D., Hewapathirana, S., Kamatchinathan, S., Kundu, D. J., Prakash, A., Frericks-Zipper, A., Eisenacher, M., et al. (2022). The PRIDE database resources in 2022: A hub for mass spectrometry-based proteomics evidences. Nucleic Acids Res. 50, D543–D552.

Pettersen, E. F., Goddard, T. D., Huang, C. C., Meng, E. C., Couch, G. S., Croll, T. I., Morris, J. H. and Ferrin, T. E. (2021). UCSF ChimeraX: Structure visualization for researchers, educators, and developers. Protein Sci. 30, 70–82.

Prokopchuk, G., Tashyreva, D., Yabuki, A., Horák, A., Masařová, P. and Lukeš, J. (2019). Morphological, ultrastructural, motility and evolutionary characterization of two new Hemistasiidae species. Protist 170, 259–282.

Prokopchuk, G., Korytář, T., Juricová, V., Majstorović, J., Horák, A., Šimek, K. and Lukeš, J. (2022). Trophic flexibility of marine diplonemids - switching from osmotrophy to bacterivory. ISME J. 16, 1409–1419.

Remmert, M., Biegert, A., Hauser, A. and Söding, J. (2012). HHblits: Lightning-fast iterative protein sequence searching by HMM-HMM alignment. Nat. Methods 9, 173–175.

Roger, A. J., Susko, E. and Leger, M. M. (2021). Evolution: Reconstructing the timeline of eukaryogenesis. Curr. Biol. 31, R193–R196.

Ronquist, F., Teslenko, M., van der Mark, P., Ayres, D. L., Darling, A., Hohna, S., Larget, B., Liu, L., Suchard, M. A. and Huelsenbeck, J. P. (2012). MrBayes 3.2: efficient Bayesian phylogenetic inference and model choice across a large model space. Syst Biol 61, 539–542.

Rout, M. P. and Field, M. C. (2022). Coatomer in the universe of cellular complexity. Mol Biol Cell accepted.

Rutherford, S. and Moore, I. (2002). The Arabidopsis Rab GTPase family: Another enigma variation. Curr. Opin. Plant Biol. 5, 518–528.

Schlacht, A. and Dacks, J. B. (2015). Unexpected ancient paralogs and an evolutionary model for the COPII coat complex. Genome Biol. Evol. 7, 1098–1109.

Sealey-Cardona, M., Schmidt, K., Demmel, L., Hirschmugl, T., Gesell, T., Dong, G. and Warren, G. (2014). Sec16 determines the size and functioning of the Golgi in the protist parasite, Trypanosoma brucei. Traffic 15, 613–629.

Sevova, E. S. and Bangs, J. D. (2009). Streamlined architecture and glycosylphosphatidylinositol-dependent trafficking in the early secretory pathway of African trypanosomes. Mol. Biol. Cell 20, 4739–4750.

Sharif, M. and Bangs, J. D. (2022). Stage-specific COPII-mediated cargo selectivity in african trypanosomes. mSphere 7, e0018822.

Stamatakis, A. (2014). RAxML version 8: A tool for phylogenetic analysis and post-analysis of large phylogenies. Bioinformatics 30, 1312–1313.

Tang, B. L. (2017). Sec16 in conventional and unconventional exocytosis: Working at the interface of membrane traffic and secretory autophagy? J. Cell. Physiol. 232, 3234–3243.

Tashyreva, D., Prokopchuk, G., Yabuki, A., Kaur, B., Faktorová, D., Votýpka, J., Kusaka, C., Fujikura, K., Shiratori, T., Ishida, K.-I., et al. (2018). Phylogeny and morphology of new diplonemids from Japan. Protist 169, 158–179.

Tashyreva, D., Simpson, A. G. B., Prokopchuk, G., Škodová-Sveráková, I., Butenko, A., Hammond, M., George, E. E., Flegontova, O., Záhonová, K., Faktorová, D., et al. (2022). Diplonemids – A review on “new” flagellates on the oceanic block. Protist 173, 125868.

Vargová, R., Wideman, J. G., Derelle, R., Klimeš, V., Kahn, R. A., Dacks, J. B. and Eliáš, M. (2021). A eukaryotewide perspective on the diversity and evolution of the ARF GTPase protein family. Genome Biol. Evol. 13, evab157.

Wang, Y. N., Wang, M. and Field, M. C. (2010). Trypanosoma brucei: Trypanosome-specific endoplasmic reticulum proteins involved in variant surface glycoprotein expression. Exp. Parasitol. 125, 208–221.

Wideman, J. G. and Muñoz-Gómez, S. A. (2016). The evolution of ERMIONE in mitochondrial biogenesis and lipid homeostasis: An evolutionary view from comparative cell biology. Biochim. Biophys. Acta - Mol. Cell Biol. Lipids 1861, 900–912.

Yorimitsu, T. and Sato, K. (2020). Sec16 function in ER export and autophagy is independent of its phosphorylation in Saccharomyces cerevisiae. Mol. Biol. Cell 31, 149–156.

Yurchenko, V., Votýpka, J., Tesařová, M., Klepetková, H., Kraeva, N., Jirku, M. and Lukeš, J. (2014). Ultrastructure and molecular phylogeny of four new species of monoxenous trypanosomatids from flies (Diptera: Brachycera) with redefinition of the genus Wallaceina. Folia Parasitol. 61, 97–112.

Záhonová, K., Füssy, Z., Birčák, E., Novák Vanclová, A. M. G., Klimeš, V., Vesteg, M., Krajčovič, J., Oborník, M. and Eliáš, M. (2018). Peculiar features of the plastids of the colourless alga Euglena longa and photosynthetic euglenophytes unveiled by transcriptome analyses. Sci. Rep. 8, 17012.

Zoltner, M., Campagnaro, G. D., Taleva, G., Burrell, A., Cerone, M., Leung, K. F., Achcar, F., Horn, D., Vaughan, S., Gadelha, C., et al. (2020). Suramin exposure alters cellular metabolism and mitochondrial energy production in African trypanosomes. J. Biol. Chem. 295, 8331–8347.

