## Supplementary data - all for "Functional differentiation of Sec13 paralogs in the euglenozoan protists"

**Supplementary Figures 1-3**

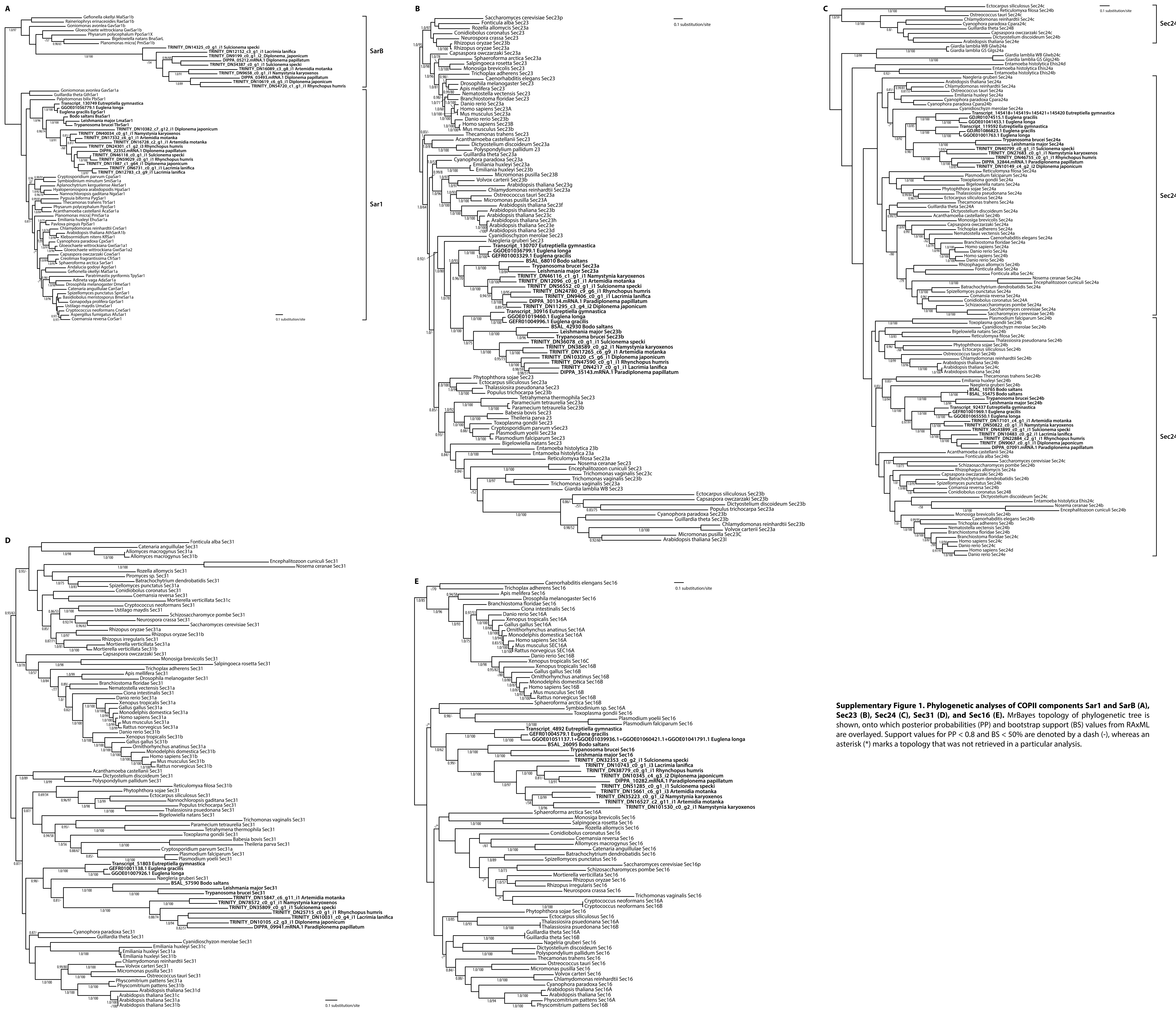


**Supplementary Figure 1. Phylogenetic analyses of COPII components Sar1 and SarB (A), Sec23 (B), Sec24 (C), Sec31 (D), and Sec16 (E).** MrBayes topology of phylogenetic tree is shown, onto which posterior probabilities (PP) and bootstrap support (BS) values from RAxML are overlayed. Support values for PP < 0.8 and BS < 50% are denoted by a dash (-), whereas an asterisk (*) marks a topology that was not retrieved in a particular analysis.

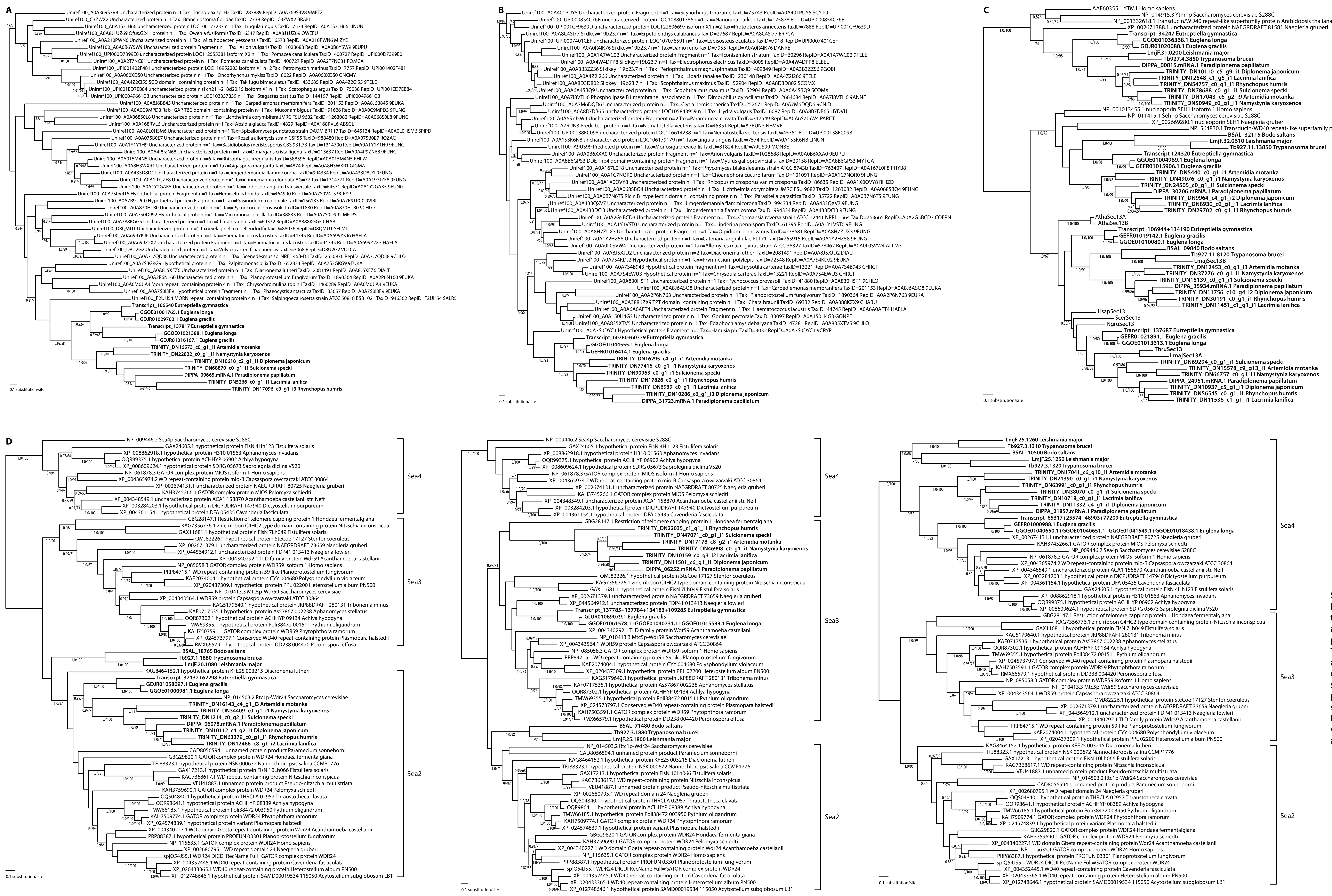


**Supplementary Figure 2. Phylogenetic analyses of hypothetical proteins PpHyp24 (A) and PpHyp27 (B), NUP component Seh1 (C), and SEA/GATOR components Sea2, Sea3, and Sea4 (D).** MrBayes topology of phylogenetic tree is shown, onto which posterior probabilities (PP) and bootstrap support (BS) values from RAxML are overlayed. Support values for PP < 0.8 and BS < 50% are denoted by a dash (-).

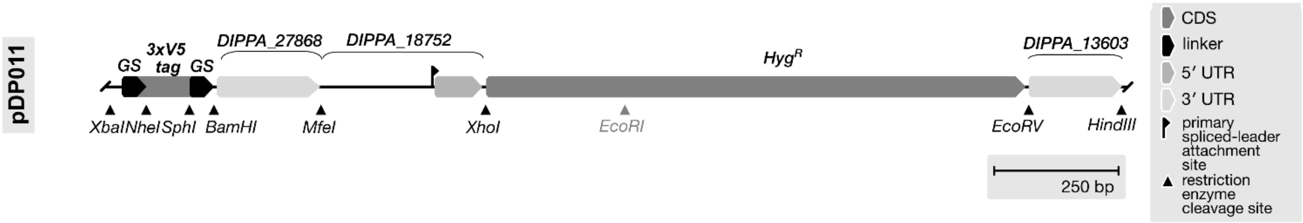


**Suppl. Figure 3: Scheme of pDP011 plasmid.**

**Functional differentiation of Sec13 paralogs in the euglenozoan protists**

Drahomíra Faktorová[1,2,#,*], Kristína Záhonová[1,3,4,*], Corinna Benz[1], Joel B. Dacks[1,5,6], Mark C. Field[1,7] and Julius Lukeš[1,2,#]

[1] *Institute of Parasitology, Biology Centre, Czech Academy of Sciences, České Budějovice, Czech Republic*
[2] *Faculty of Sciences, University of South Bohemia, České Budějovice, Czech Republic*
[3] *Faculty of Science, Charles University, BIOCEV, Vestec, Czech Republic*
[4] *Life Science Research Centre, Department of Biology and Ecology, Faculty of Science, University of Ostrava, Ostrava, Czech Republic*
[5] *Division of Infectious Disease, Department of Medicine, Faculty of Medicine and Dentistry, University of Alberta, Edmonton, Canada*
[6] *Centre for Life's Origins and Evolution, Department of Genetics, Evolution, and Environment, University College London, London, UK*
[7] *School of Life Sciences, University of Dundee, Dundee, UK*

*These authors contributed equally to this work.

**Supplementary Tables 1-7**

**Supplementary Table 1.** List of COPII genes identified in *P. papillatum* genome.

| COPII gene | *P. papiilatum* accession |
|---|---|
| **Sar1** | DIPPA_22352 |
| **SarB** | DIPPA_05212 |
| | DIPPA_03493 |
| **Sec12** | - |
| **Sec13a** | DIPPA_24951 |
| **Sec13b** | DIPPA_35934 |
| **Sec16** | DIPPA_10282 |
| **Sec23a** | DIPPA_30134 |
| **Sec23b** | DIPPA_35143 |
| **Sec24I** | DIPPA_32844 |
| **Sec24II** | DIPPA_07091 |
| **Sec31** | DIPPA_09941 |

**Supplementary Table 2.** List of COPII components identified in studied diplonemids and their reverse best hits.

| Query | | | FWD BLAST hit | | | | REV BLAST hit - *H. sapiens* | | | REV BLAST hit - *S. cerevisiae* | | | |
|---|---|---|---|---|---|---|---|---|---|---|---|---|---|
| **Protein** | **ID** | **Organism** | **Organism** | **ID** | **Sequence** | **E-value** | **ID** | **Description** | **E-value** | **ID** | **Description** | **E-value** | **Note** |
| **Sar1** | XP 845120.1 | *Trypanosoma brucei* | *Paradiplonema papillatum* | DIPPA 22352.mRNA.1 | MGWFDWLSQT | 1,86E-73 | NP 057187.1 | GTP-binding protein SAR1b [Homo sapiens] | 1,89E-60 | NP 015106.1 | Arf family GTPase SAR1 [Saccharomyces cerevisiae S288C] | 1,14E-66 | confirmed by tree |
| **SarB** | QXF29078.1 | *Planomonas micra* | *Paradiplonema papillatum* | DIPPA 05212.mRNA.1 | MGCSDSKHDGF | 4.28E-05 | no SarB in *H. sapiens* | | | no SarB in *S. cerevisiae* | | | confirmed by tree |
| **SarB** | QXF29078.1 | *Planomonas micra* | *Paradiplonema papillatum* | DIPPA 03493.mRNA.1 | MGCTGSKGPPC | 3,24E-05 | no SarB in *H. sapiens* | | | no SarB in *S. cerevisiae* | | | confirmed by tree |
| **Sec12** | XP 002672421.1 | *Naegleria gruberi* | *Paradiplonema papillatum* | DIPPA 16358.mRNA.1 | MLAHPPAPQW( | 1.71E-01 | no hit found | | | NP 011568.3 | uncharacterized protein YGR054W [Saccharomyces cerevisiae S288C] | 4.41E-01 | |
| **Sec13a** | XP 827898.1 | *Trypanosoma brucei* | *Paradiplonema papillatum* | DIPPA 24951.mRNA.1 | MAWPGPYLDPS | 3,36E-39 | XP 005265436.1 | protein SEC13 homolog isoform X3 [Homo sapiens] | 4,36E-65 | NP 013309.1 | GTPase-activating protein SEC13 [Saccharomyces cerevisiae S288C] | 1,60E-54 | confirmed by tree |
| **Sec13b** | XP 828886.1 | *Trypanosoma brucei* | *Paradiplonema papillatum* | DIPPA 35934.mRNA.1 | MASKQVVETTH | 1.99E-28 | NP 899195.1 | protein SEC13 homolog isoform 1 [Homo sapiens] | 1.16E-47 | NP 013309.1 | GTPase-activating protein SEC13 [Saccharomyces cerevisiae S288C] | 8.37E-46 | confirmed by tree |
| **Sec16** | XP 843977.1 | *Trypanosoma brucei* | *Paradiplonema papillatum* | DIPPA 10282.mRNA.1 | MPSSQLQTGAK | found by pull-down of Sec13b (DIPPA 35934.mRNA.1) | XP 011536442.1 | pre-mRNA-processing factor 40 homolog B isoform X8 [Homo sapiens] | 6,97E-01 | NP 015478.1 | Urn1p [Saccharomyces cerevisiae S288C] | 2,46E-02 | confirmed by tree |
| **Sec23** | XP 823008.1 | *Trypanosoma brucei* | *Paradiplonema papillatum* | DIPPA 30134.mRNA.1 | MGDRSADEIKW | 5.01E-171 | XP 016883082.1 | protein transport protein Sec23B isoform X1 [Homo sapiens] | 0 | NP 015507.1 | GTPase-activating protein SEC23 [Saccharomyces cerevisiae S288C] | 0 | confirmed by tree |
| **Sec23** | XP 847197.1 | *Trypanosoma brucei* | *Paradiplonema papillatum* | DIPPA 35143.mRNA.1 | MAEYFQDMEDI | 1.74E-124 | NP 006355.2 | protein transport protein Sec23A [Homo sapiens] | ######## | NP 015507.1 | GTPase-activating protein SEC23 [Saccharomyces cerevisiae S288C] | ######## | confirmed by tree |
| **Sec24I** | XP 844132.1 | *Trypanosoma brucei* | *Paradiplonema papillatum* | DIPPA 32844.mRNA.1 | MAGNGYPAMP | 2,87E-105 | XP 006714586.1 | protein transport protein Sec24A isoform X2 [Homo sapiens] | ######## | NP 012157.3 | COPII subunit SEC24 [Saccharomyces cerevisiae S288C] | 4,30E-83 | confirmed by tree |
| **Sec24II** | XP 843715.1 | *Trypanosoma brucei* | *Paradiplonema papillatum* | DIPPA 07091.mRNA.1 | MSGAPPGAMG | 2.42E-122 | XP 011538683.1 | protein transport protein Sec24C isoform X1 [Homo sapiens] | ######## | NP 011966.1 | Sfb3p [Saccharomyces cerevisiae S288C] | 2.63E-71 | confirmed by tree |
| **Sec31** | XP 828691.1 | *Trypanosoma brucei* | *Paradiplonema papillatum* | DIPPA 09941.mRNA.1 | MEGPGSSVFAW | 4,52E-28 | NP 001287674.1 | protein transport protein Sec31A isoform 7 [Homo sapiens] | 4,57E-40 | NP 010086.2 | Sec31p [Saccharomyces cerevisiae S288C] | 1,14E-18 | confirmed by tree |
| **Sar1** | DIPPA 22352.mRNA.1 | *Paradiplonema papillatum* | *Diplonema japonicum* | TRINITY DN11987 c1 g64 i1 | MGWFDWMW( | 3.24E-92 | NP 057187.1 | GTP-binding protein SAR1b [Homo sapiens] | 6.85E-67 | NP 015106.1 | Arf family GTPase SAR1 [Saccharomyces cerevisiae S288C] | 2.07E-65 | confirmed by tree |
| **Sar1** | DIPPA 22352.mRNA.1 | *Paradiplonema papillatum* | *Diplonema japonicum* | TRINITY DN10382 c7 g12 i1 | MDSLFSMFQSV | 1,67E-64 | NP 057187.1 | GTP-binding protein SAR1b [Homo sapiens] | 8,85E-57 | NP 015106.1 | Arf family GTPase SAR1 [Saccharomyces cerevisiae S288C] | 1,02E-56 | confirmed by tree |
| **SarB** | DIPPA 05212.mRNA.1 | *Paradiplonema papillatum* | *Diplonema japonicum* | TRINITY DN9199 c0 g1 i2 | MGCTRSKESSR( | 6.17E-60 | no SarB in *H. sapiens* | | | no SarB in *S. cerevisiae* | | | confirmed by tree |
| **SarB** | DIPPA 03493.mRNA.1 | *Paradiplonema papillatum* | *Diplonema japonicum* | TRINITY DN10619 c6 g5 i1 | MGCSESKPPPRI | 1,31E-53 | no SarB in *H. sapiens* | | | no SarB in *S. cerevisiae* | | | confirmed by tree |
| **Sec12** | XP 002672421.1 | *Naegleria gruberi* | *Diplonema japonicum* | - | | | | | | | | | |
| **Sec13a** | DIPPA 24951.mRNA.1 | *Paradiplonema papillatum* | *Diplonema japonicum* | TRINITY DN10937 c5 g1 i1 | MANGVVTEMLI | 4.59E-126 | XP 005265436.1 | protein SEC13 homolog isoform X3 [Homo sapiens] | 1.15E-65 | NP 013309.1 | GTPase-activating protein SEC13 [Saccharomyces cerevisiae S288C] | 7.50E-57 | confirmed by tree |
| **Sec13b** | DIPPA 35934.mRNA.1 | *Paradiplonema papillatum* | *Diplonema japonicum* | TRINITY DN11756 c10 g4 i2 | MSEAAQRVDIAT | 1,67E-149 | NP 899195.1 | protein SEC13 homolog isoform 1 [Homo sapiens] | 2,41E-52 | NP 013309.1 | GTPase-activating protein SEC13 [Saccharomyces cerevisiae S288C] | 9,35E-47 | confirmed by tree |
| **Sec16** | DIPPA 10282.mRNA.1 | *Paradiplonema papillatum* | *Diplonema japonicum* | TRINITY DN10345 c4 g3 i2 | MPVAVGSKVRL | 1.25E-107 | XP 011517565.1 | protein transport protein Sec16A isoform X16 [Homo sapiens] | 1.87E-08 | NP 015240.1 | Sec16p [Saccharomyces cerevisiae S288C] | 6.75E-03 | confirmed by tree |
| **Sec23** | DIPPA 30134.mRNA.1 | *Paradiplonema papillatum* | *Diplonema japonicum* | TRINITY DN11295 c3 g4 i2 | MGEVGQEAKW | 0 | XP 016883082.1 | protein transport protein Sec23B isoform X1 [Homo sapiens] | 0 | NP 015507.1 | GTPase-activating protein SEC23 [Saccharomyces cerevisiae S288C] | 0 | confirmed by tree |
| **Sec23** | DIPPA 35143.mRNA.1 | *Paradiplonema papillatum* | *Diplonema japonicum* | TRINITY DN10320 c5 g6 i1 | MADYFQTMEDI | 0 | NP 006355.2 | protein transport protein Sec23A [Homo sapiens] | 0 | NP 015507.1 | GTPase-activating protein SEC23 [Saccharomyces cerevisiae S288C] | ######## | confirmed by tree |
| **Sec24I** | DIPPA 32844.mRNA.1 | *Paradiplonema papillatum* | *Diplonema japonicum* | TRINITY DN10149 c4 g2 i2 | MSYPYANQHQH | 0 | XP 011529842.1 | protein transport protein Sec24B isoform X8 [Homo sapiens] | 0 | NP 012157.3 | COPII subunit SEC24 [Saccharomyces cerevisiae S288C] | 1,76E-89 | confirmed by tree |
| **Sec24II** | DIPPA 07091.mRNA.1 | *Paradiplonema papillatum* | *Diplonema japonicum* | TRINITY DN9067 c0 g1 i1 | MSAPPPGALPP( | 0 | XP 011538683.1 | protein transport protein Sec24C isoform X1 [Homo sapiens] | 0 | NP 011966.1 | Sfb3p [Saccharomyces cerevisiae S288C] | 1.35E-78 | confirmed by tree |
| **Sec31** | DIPPA 09941.mRNA.1 | *Paradiplonema papillatum* | *Diplonema japonicum* | TRINITY DN10105 c2 g3 i1 | MEGPGSSVFAW | 0 | NP 057295.2 | protein transport protein Sec31A isoform 2 [Homo sapiens] | 0 | NP 010086.2 | Sec31p [Saccharomyces cerevisiae S288C] | 2.04E-30 | confirmed by tree |
| **Sar1** | DIPPA 22352.mRNA.1 | *Paradiplonema papillatum* | *Rhynchopus humris* | TRINITY DN24301 c1 g2 i3 | MGWFDWLTST | 4,18E-82 | NP 057187.1 | GTP-binding protein SAR1b [Homo sapiens] | 1,72E-69 | NP 015106.1 | Arf family GTPase SAR1 [Saccharomyces cerevisiae S288C] | 1,37E-65 | confirmed by tree |
| **SarB** | DIPPA 05212.mRNA.1 | *Paradiplonema papillatum* | *Rhynchopus humris* | - | | | | | | | | | confirmed by tree |
| **SarB** | DIPPA 03493.mRNA.1 | *Paradiplonema papillatum* | *Rhynchopus humris* | TRINITY DN54720 c1 g1 i1 | MLRRLSSNDSVL | 6,80E-38 | no SarB in *H. sapiens* | | | no SarB in *S. cerevisiae* | | | confirmed by tree |
| **Sec12** | XP 002672421.1 | *Naegleria gruberi* | *Rhynchopus humris* | - | | | | | | | | | |
| **Sec13a** | DIPPA 24951.mRNA.1 | *Paradiplonema papillatum* | *Rhynchopus humris* | TRINITY DN56545 c0 g1 i1 | MAAEVLHTGHT | 7,58E-121 | NP 899195.1 | protein SEC13 homolog isoform 1 [Homo sapiens] | 5,78E-73 | NP 013309.1 | GTPase-activating protein SEC13 [Saccharomyces cerevisiae S288C] | 3,92E-56 | confirmed by tree |
| **Sec13b** | DIPPA 35934.mRNA.1 | *Paradiplonema papillatum* | *Rhynchopus humris* | TRINITY DN30191 c0 g1 i1 | MSDRPQKQVID | 7.38E-141 | XP 005265436.1 | protein SEC13 homolog isoform X3 [Homo sapiens] | 2.20E-53 | NP 013309.1 | GTPase-activating protein SEC13 [Saccharomyces cerevisiae S288C] | 3.60E-51 | confirmed by tree |
| **Sec16** | DIPPA 10282.mRNA.1 | *Paradiplonema papillatum* | *Rhynchopus humris* | TRINITY DN38779 c0 g1 i1 | AAMLPDAVLPP/ | 2,36E-54 | NP 596869.4 | titin isoform N2-A [Homo sapiens] | 3,54E-02 | NP 015478.1 | Urn1p [Saccharomyces cerevisiae S288C] | 8,16E-02 | confirmed by tree |
| **Sec23** | DIPPA 30134.mRNA.1 | *Paradiplonema papillatum* | *Rhynchopus humris* | TRINITY DN24780 c9 g6 i1 | MDEGQQAAAAI | 0 | XP 016883082.1 | protein transport protein Sec23B isoform X1 [Homo sapiens] | 0 | NP 015507.1 | GTPase-activating protein SEC23 [Saccharomyces cerevisiae S288C] | ######## | confirmed by tree |
| **Sec23** | DIPPA 35143.mRNA.1 | *Paradiplonema papillatum* | *Rhynchopus humris* | TRINITY DN47590 c0 g1 i1 | MAEYFQQMEDI | 0 | NP 006355.2 | protein transport protein Sec23A [Homo sapiens] | 0 | NP 015507.1 | GTPase-activating protein SEC23 [Saccharomyces cerevisiae S288C] | ######## | confirmed by tree |
| **Sec24I** | DIPPA 32844.mRNA.1 | *Paradiplonema papillatum* | *Rhynchopus humris* | TRINITY DN46755 c0 g1 i1 | GYAAGNGQQQ' | 0 | XP 011529842.1 | protein transport protein Sec24B isoform X8 [Homo sapiens] | 0 | NP 012157.3 | COPII subunit SEC24 [Saccharomyces cerevisiae S288C] | 1,96E-78 | confirmed by tree |
| **Sec24II** | DIPPA 07091.mRNA.1 | *Paradiplonema papillatum* | *Rhynchopus humris* | TRINITY DN22884 c2 g1 i1 | GPPAMPPGGGI | 0 | XP 016872457.1 | protein transport protein Sec24C isoform X3 [Homo sapiens] | 0 | NP 011966.1 | Sfb3p [Saccharomyces cerevisiae S288C] | 1.07E-65 | confirmed by tree |
| **Sec31** | DIPPA 09941.mRNA.1 | *Paradiplonema papillatum* | *Rhynchopus humris* | TRINITY DN25715 c0 g1 i1 | MEGPSSCVCAW | 8,85E-147 | NP 001287674.1 | protein transport protein Sec31A isoform 7 [Homo sapiens] | 1,16E-33 | NP 010086.2 | Sec31p [Saccharomyces cerevisiae S288C] | 4,41E-15 | confirmed by tree |
| **Sar1** | DIPPA 22352.mRNA.1 | *Paradiplonema papillatum* | *Lacrimia lanifica* | TRINITY DN12783 c3 g9 i1 | MGWFDWVGS/ | 2.21E-89 | NP 057187.1 | GTP-binding protein SAR1b [Homo sapiens] | 3.62E-66 | NP 015106.1 | Arf family GTPase SAR1 [Saccharomyces cerevisiae S288C] | 8.78E-63 | confirmed by tree |
| **Sar1** | DIPPA 22352.mRNA.1 | *Paradiplonema papillatum* | *Lacrimia lanifica* | TRINITY DN6731 c0 g1 i1 | MGLLDWVASA\ | 8,99E-83 | NP 057187.1 | GTP-binding protein SAR1b [Homo sapiens] | 1,49E-66 | NP 015106.1 | Arf family GTPase SAR1 [Saccharomyces cerevisiae S288C] | 9,96E-62 | confirmed by tree |
| **SarB** | DIPPA 05212.mRNA.1 | *Paradiplonema papillatum* | *Lacrimia lanifica* | TRINITY DN12152 c3 g1 i1 | MRPPRPVVSSV\ | 1.60E-35 | no SarB in *H. sapiens* | | | no SarB in *S. cerevisiae* | | | confirmed by tree |
| **SarB** | DIPPA 03493.mRNA.1 | *Paradiplonema papillatum* | *Lacrimia lanifica* | - | | | | | | | | | confirmed by tree |
| **Sec12** | XP 002672421.1 | *Naegleria gruberi* | *Lacrimia lanifica* | - | | | | | | | | | |
| **Sec13a** | DIPPA 24951.mRNA.1 | *Paradiplonema papillatum* | *Lacrimia lanifica* | TRINITY DN11536 c1 g1 i1 | MATTVDHFNTG | 5.18E-112 | NP 899195.1 | protein SEC13 homolog isoform 1 [Homo sapiens] | 1.36E-69 | NP 013309.1 | GTPase-activating protein SEC13 [Saccharomyces cerevisiae S288C] | 8.95E-55 | confirmed by tree |
| **Sec13b** | DIPPA 35934.mRNA.1 | *Paradiplonema papillatum* | *Lacrimia lanifica* | TRINITY DN11451 c1 g1 i1 | MAEHPQKQVIQ | 2,63E-135 | NP 899195.1 | protein SEC13 homolog isoform 1 [Homo sapiens] | 2,13E-53 | NP 013309.1 | GTPase-activating protein SEC13 [Saccharomyces cerevisiae S288C] | 1,04E-44 | confirmed by tree |
| **Sec16** | DIPPA 10282.mRNA.1 | *Paradiplonema papillatum* | *Lacrimia lanifica* | TRINITY DN10743 c0 g1 i3 | MAAGFQVGAH' | 9.13E-45 | no hit found | | | NP 013245.1 | endo-1.3(4)-beta-glucanase [Saccharomyces cerevisiae S288C] | 6.64E-01 | confirmed by tree |
| **Sec23** | DIPPA 30134.mRNA.1 | *Paradiplonema papillatum* | *Lacrimia lanifica* | TRINITY DN9406 c0 g1 i1 | MGAIGDDYVPK | 0 | XP 016883082.1 | protein transport protein Sec23B isoform X1 [Homo sapiens] | 0 | NP 015507.1 | GTPase-activating protein SEC23 [Saccharomyces cerevisiae S288C] | ######## | confirmed by tree |
| **Sec23** | DIPPA 35143.mRNA.1 | *Paradiplonema papillatum* | *Lacrimia lanifica* | TRINITY DN4217 c0 g1 i1 | MEEFKKIEEEHG | 0 | NP 006355.2 | protein transport protein Sec23A [Homo sapiens] | 0 | NP 015507.1 | GTPase-activating protein SEC23 [Saccharomyces cerevisiae S288C] | ######## | confirmed by tree |
| **Sec24I** | DIPPA 32844.mRNA.1 | *Paradiplonema papillatum* | *Lacrimia lanifica* | - | | | | | | | | 1,25E-75 | confirmed by tree |
| **Sec24II** | DIPPA 07091.mRNA.1 | *Paradiplonema papillatum* | *Lacrimia lanifica* | TRINITY DN10483 c0 g2 i1 | TGAPGMPPGM | 1,22E-67 | XP 016872456.1 | protein transport protein Sec24C isoform X2 [Homo sapiens] | 0 | NP 011966.1 | Sfb3p [Saccharomyces cerevisiae S288C] | 1,25E-75 | confirmed by tree |
| **Sec31** | DIPPA 09941.mRNA.1 | *Paradiplonema papillatum* | *Lacrimia lanifica* | TRINITY DN10031 c0 g4 i1 | VRWGGGGDGV | 4.34E-83 | NP 001287673.1 | protein transport protein Sec31A isoform 6 [Homo sapiens] | 4.12E-25 | NP 010086.2 | Sec31p [Saccharomyces cerevisiae S288C] | 1.86E-22 | confirmed by tree |
| **Sar1** | DIPPA 22352.mRNA.1 | *Paradiplonema papillatum* | *Sulcionema specki* | TRINITY DN46118 c0 g1 i1 | MGWFDWMWI | 5,10E-91 | NP 057187.1 | GTP-binding protein SAR1b [Homo sapiens] | 1,43E-67 | NP 015106.1 | Arf family GTPase SAR1 [Saccharomyces cerevisiae S288C] | 1,76E-66 | confirmed by tree |
| **SarB** | DIPPA 05212.mRNA.1 | *Paradiplonema papillatum* | *Sulcionema specki* | TRINITY DN14325 c0 g1 i1 | MGCSKSKQHEP | 3.51E-38 | no SarB in *H. sapiens* | | | no SarB in *S. cerevisiae* | | | confirmed by tree |
| **SarB** | DIPPA 03493.mRNA.1 | *Paradiplonema papillatum* | *Sulcionema specki* | TRINITY DN34387 c0 g1 i1 | MGCKGSHEASV | 4,51E-62 | no SarB in *H. sapiens* | | | no SarB in *S. cerevisiae* | | | confirmed by tree |
| **Sec12** | XP 002672421.1 | *Naegleria gruberi* | *Sulcionema specki* | - | | | | | | | | | |
| **Sec13a** | DIPPA 24951.mRNA.1 | *Paradiplonema papillatum* | *Sulcionema specki* | TRINITY DN69294 c0 g1 i1 | MAGQEINTAHE | 8,99E-90 | XP 005265436.1 | protein SEC13 homolog isoform X3 [Homo sapiens] | 9,38E-74 | NP 013309.1 | GTPase-activating protein SEC13 [Saccharomyces cerevisiae S288C] | 1,14E-59 | confirmed by tree |
| **Sec13b** | DIPPA 35934.mRNA.1 | *Paradiplonema papillatum* | *Sulcionema specki* | TRINITY DN15139 c0 g1 i1 | MSESTAARQSID | 9.33E-126 | XP 005265436.1 | protein SEC13 homolog isoform X3 [Homo sapiens] | 1.82E-58 | NP 013309.1 | GTPase-activating protein SEC13 [Saccharomyces cerevisiae S288C] | 5.75E-50 | confirmed by tree |
| **Sec16** | DIPPA 10282.mRNA.1 | *Paradiplonema papillatum* | *Sulcionema specki* | TRINITY DN32353 c0 g2 i1 | QHQPSQQPFQA | 2.98E-24 | XP 011517559.1 | protein transport protein Sec16A isoform X11 [Homo sapiens] | 5.16E-13 | no hit found | | | confirmed by tree |
| **Sec16** | DIPPA 10282.mRNA.1 | *Paradiplonema papillatum* | *Sulcionema specki* | TRINITY DN51285 c0 g1 i1 | EHTPVPTTTTTTI | 4,30E-33 | XP 011517563.1 | protein transport protein Sec16A isoform X14 [Homo sapiens] | 2,07E-03 | no hit found | | | confirmed by tree |
| **Sec23** | DIPPA 30134.mRNA.1 | *Paradiplonema papillatum* | *Sulcionema specki* | TRINITY DN56552 c0 g1 i1 | PLKMSAQQHDN | 0 | XP 016883082.1 | protein transport protein Sec23B isoform X1 [Homo sapiens] | 0 | NP 015507.1 | GTPase-activating protein SEC23 [Saccharomyces cerevisiae S288C] | ######## | confirmed by tree |
| **Sec23** | DIPPA 35143.mRNA.1 | *Paradiplonema papillatum* | *Sulcionema specki* | TRINITY DN36078 c0 g1 i1 | MDYFKDEEARD | 0 | NP 006355.2 | protein transport protein Sec23A [Homo sapiens] | 0 | NP 015507.1 | GTPase-activating protein SEC23 [Saccharomyces cerevisiae S288C] | ######## | confirmed by tree |
| **Sec24I** | DIPPA 32844.mRNA.1 | *Paradiplonema papillatum* | *Sulcionema specki* | TRINITY DN40799 c0 g1 i1 | APHHSAPPPGPI | 0 | XP 006714586.1 | protein transport protein Sec24A isoform X2 [Homo sapiens] | 0 | NP 014349.1 | COPII subunit SFB2 [Saccharomyces cerevisiae S288C] | 6.79E-86 | confirmed by tree |
| **Sec24II** | DIPPA 07091.mRNA.1 | *Paradiplonema papillatum* | *Sulcionema specki* | TRINITY DN43899 c0 g1 i1 | PPGMPPGGPP( | 0 | NP 940999.1 | protein transport protein Sec24C [Homo sapiens] | 0 | NP 011966.1 | Sfb3p [Saccharomyces cerevisiae S288C] | 2,73E-71 | confirmed by tree |
| **Sec31** | DIPPA 09941.mRNA.1 | *Paradiplonema papillatum* | *Sulcionema specki* | TRINITY DN35809 c0 g1 i1 | MELASTNVSCVI | 7.00E-129 | NP 001287674.1 | protein transport protein Sec31A isoform 7 [Homo sapiens] | 1.49E-41 | NP 010086.2 | Sec31p [Saccharomyces cerevisiae S288C] | 6.14E-29 | confirmed by tree |
| **Sar1** | DIPPA 22352.mRNA.1 | *Paradiplonema papillatum* | *Artemidia motanka* | TRINITY DN17332 c6 g1 i1 | MGWFDWLEGA | 1,01E-90 | NP 057187.1 | GTP-binding protein SAR1b [Homo sapiens] | 1,12E-69 | NP 015106.1 | Arf family GTPase SAR1 [Saccharomyces cerevisiae S288C] | 9,74E-69 | confirmed by tree |
| **Sar1** | DIPPA 22352.mRNA.1 | *Paradiplonema papillatum* | *Artemidia motanka* | TRINITY DN16728 c2 g1 i1 | MSFLAAAKAWL | 1.89E-77 | NP 057187.1 | GTP-binding protein SAR1b [Homo sapiens] | 2.36E-58 | NP 015106.1 | Arf family GTPase SAR1 [Saccharomyces cerevisiae S288C] | 3.74E-59 | confirmed by tree |
| **SarB** | DIPPA 05212.mRNA.1 | *Paradiplonema papillatum* | *Artemidia motanka* | - | | | | | | | | | confirmed by tree |
| **SarB** | DIPPA 03493.mRNA.1 | *Paradiplonema papillatum* | *Artemidia motanka* | TRINITY DN16089 c3 g8 i1 | MGCCGSAQPKF | 2,82E-30 | no SarB in *H. sapiens* | | | no SarB in *S. cerevisiae* | | | confirmed by tree |
| **Sec12** | XP 002672421.1 | *Naegleria gruberi* | *Artemidia motanka* | - | | | | | | | | | |
| **Sec13a** | DIPPA 24951.mRNA.1 | *Paradiplonema papillatum* | *Artemidia motanka* | TRINITY DN15578 c9 g13 i1 | MAERMLNTGHI | 4,99E-66 | NP 899195.1 | protein SEC13 homolog isoform 1 [Homo sapiens] | 2,27E-49 | NP 013309.1 | GTPase-activating protein SEC13 [Saccharomyces cerevisiae S288C] | 7,67E-38 | confirmed by tree |
| **Sec13b** | DIPPA 35934.mRNA.1 | *Paradiplonema papillatum* | *Artemidia motanka* | TRINITY DN12453 c0 g1 i3 | MSEGGRQTFET | 2.23E-90 | NP 899195.1 | protein SEC13 homolog isoform 1 [Homo sapiens] | 2.85E-54 | NP 013309.1 | GTPase-activating protein SEC13 [Saccharomyces cerevisiae S288C] | 6.11E-47 | confirmed by tree |
| **Sec16** | DIPPA 10282.mRNA.1 | *Paradiplonema papillatum* | *Artemidia motanka* | TRINITY DN15661 c6 g1 i3 | MADPFAGDPQ/ | 1,08E-26 | NP 149118.2 | protein transport protein Sec16B isoform 1 [Homo sapiens] | 1,97E-07 | NP 012913.3 | Prp40p [Saccharomyces cerevisiae S288C] | 3,43E-05 | confirmed by tree |
| **Sec16** | DIPPA 10282.mRNA.1 | *Paradiplonema papillatum* | *Artemidia motanka* | TRINITY DN16527 c2 g11 i1 | DPFSPPPAGVD/ | 5.42E-33 | XP 005260627.1 | E3 ubiquitin-protein ligase Itchy homolog isoform X4 [Homo sapiens] | 4.93E-06 | NP 012913.3 | Prp40p [Saccharomyces cerevisiae S288C] | 5.44E-07 | confirmed by tree |
| **Sec23** | DIPPA 30134.mRNA.1 | *Paradiplonema papillatum* | *Artemidia motanka* | TRINITY DN12096 c0 g1 i1 | MAYQYEQQAYA | 0 | XP 016883082.1 | protein transport protein Sec23B isoform X1 [Homo sapiens] | 0 | NP 015507.1 | GTPase-activating protein SEC23 [Saccharomyces cerevisiae S288C] | 0 | confirmed by tree |
| **Sec23** | DIPPA 35143.mRNA.1 | *Paradiplonema papillatum* | *Artemidia motanka* | TRINITY DN17265 c6 g9 i1 | MAEYFEKMEEA | 0 | NP 006355.2 | protein transport protein Sec23A [Homo sapiens] | 0 | NP 015507.1 | GTPase-activating protein SEC23 [Saccharomyces cerevisiae S288C] | ######## | confirmed by tree |
| **Sec24I** | DIPPA 32844.mRNA.1 | *Paradiplonema papillatum* | *Artemidia motanka* | - | | | | | | | | | |
| **Sec24II** | DIPPA 07091.mRNA.1 | *Paradiplonema papillatum* | *Artemidia motanka* | TRINITY DN17101 c4 g1 i1 | MLPPPPAGVPP' | 4,49E-72 | NP 940999.1 | protein transport protein Sec24C [Homo sapiens] | 0 | NP 011966.1 | Sfb3p [Saccharomyces cerevisiae S288C] | 1,30E-68 | confirmed by tree |
| **Sec31** | DIPPA 09941.mRNA.1 | *Paradiplonema papillatum* | *Artemidia motanka* | TRINITY DN15847 c6 g11 i1 | MSELAGSALSC\ | 4.33E-83 | NP 057295.2 | protein transport protein Sec31A isoform 2 [Homo sapiens] | 7.49E-45 | NP 010086.2 | Sec31p [Saccharomyces cerevisiae S288C] | 1.90E-38 | confirmed by tree |
| **Sar1** | DIPPA 22352.mRNA.1 | *Paradiplonema papillatum* | *Namystynia karyoxenos* | TRINITY DN40034 c0 g1 i1 | MGWFDWMW( | 1,15E-91 | NP 057187.1 | GTP-binding protein SAR1b [Homo sapiens] | 3,28E-71 | NP 015106.1 | Arf family GTPase SAR1 [Saccharomyces cerevisiae S288C] | 4,86E-67 | confirmed by tree |
| **SarB** | DIPPA 05212.mRNA.1 | *Paradiplonema papillatum* | *Namystynia karyoxenos* | TRINITY DN9658 c0 g1 i1 | MGCCESADKPQ | 3.02E-31 | no SarB in *H. sapiens* | | | no SarB in *S. cerevisiae* | | | confirmed by tree |
| **SarB** | DIPPA 03493.mRNA.1 | *Paradiplonema papillatum* | *Namystynia karyoxenos* | - | | | | | | | | | confirmed by tree |
| **Sec12** | XP 002672421.1 | *Naegleria gruberi* | *Namystynia karyoxenos* | - | | | | | | | | | |
| **Sec13a** | DIPPA 24951.mRNA.1 | *Paradiplonema papillatum* | *Namystynia karyoxenos* | TRINITY DN66757 c0 g1 i1 | MVEVMNTGHE/ | 2.14E-64 | NP 899195.1 | protein SEC13 homolog isoform 1 [Homo sapiens] | 2.47E-61 | NP 013309.1 | GTPase-activating protein SEC13 [Saccharomyces cerevisiae S288C] | 4.00E-52 | confirmed by tree |
| **Sec13b** | DIPPA 35934.mRNA.1 | *Paradiplonema papillatum* | *Namystynia karyoxenos* | TRINITY DN37276 c0 g1 i1 | MADGSARLSFD | 4,27E-93 | NP 899195.1 | protein SEC13 homolog isoform 1 [Homo sapiens] | 8,48E-54 | NP 013309.1 | GTPase-activating protein SEC13 [Saccharomyces cerevisiae S288C] | 1,54E-46 | confirmed by tree |
| **Sec16** | DIPPA 10282.mRNA.1 | *Paradiplonema papillatum* | *Namystynia karyoxenos* | TRINITY DN35223 c0 g1 i2 | MADPAAAASAP | 9.20E-38 | XP 011517551.1 | protein transport protein Sec16A isoform X2 [Homo sapiens] | 1.63E-07 | NP 012913.3 | Prp40p [Saccharomyces cerevisiae S288C] | 1.17E-05 | confirmed by tree |
| **Sec16** | DIPPA 10282.mRNA.1 | *Paradiplonema papillatum* | *Namystynia karyoxenos* | TRINITY DN101530 c0 g2 i1 | FGSPLSAQEPPA | 4,70E-25 | no hit found | | | no hit found | | | confirmed by tree |
| **Sec23** | DIPPA 30134.mRNA.1 | *Paradiplonema papillatum* | *Namystynia karyoxenos* | TRINITY DN46116 c1 g1 i1 | MVVPLGCLYPPL | 0 | XP 016883082.1 | protein transport protein Sec23B isoform X1 [Homo sapiens] | 0 | NP 015507.1 | GTPase-activating protein SEC23 [Saccharomyces cerevisiae S288C] | 0 | confirmed by tree |
| **Sec23** | DIPPA 35143.mRNA.1 | *Paradiplonema papillatum* | *Namystynia karyoxenos* | TRINITY DN38589 c0 g2 i1 | PSMGAPPGPSN | 0 | XP 016883082.1 | protein transport protein Sec23B isoform X1 [Homo sapiens] | 0 | NP 015507.1 | GTPase-activating protein SEC23 [Saccharomyces cerevisiae S288C] | ######## | confirmed by tree |
| **Sec24I** | DIPPA 32844.mRNA.1 | *Paradiplonema papillatum* | *Namystynia karyoxenos* | TRINITY DN27683 c0 g1 i1 | MARLPPPPAPG | 0 | XP 006714586.1 | protein transport protein Sec24A isoform X2 [Homo sapiens] | 0 | NP 014349.1 | COPII subunit SFB2 [Saccharomyces cerevisiae S288C] | 9.39E-82 | confirmed by tree |
| **Sec24II** | DIPPA 07091.mRNA.1 | *Paradiplonema papillatum* | *Namystynia karyoxenos* | TRINITY DN50822 c0 g1 i1 | QQGGWARPPPI | 0 | XP 011538683.1 | protein transport protein Sec24C isoform X1 [Homo sapiens] | 0 | NP 011966.1 | Sfb3p [Saccharomyces cerevisiae S288C] | 1,48E-75 | confirmed by tree |
| **Sec31** | DIPPA 09941.mRNA.1 | *Paradiplonema papillatum* | *Namystynia karyoxenos* | TRINITY DN78572 c0 g1 i1 | MSEVAGAPCSC | 2,16E-107 | NP 001287674.1 | protein transport protein Sec31A isoform 7 [Homo sapiens] | 1,02E-44 | NP 010086.2 | Sec31p [Saccharomyces cerevisiae S288C] | 1,74E-31 | confirmed by tree |

**Supplementary Table 3.** List of primers used in this study. Sequences in bold correspond to parts aligning to pDP002 (A) or pDP011 (B and C) plasmid.

**A. C-terminal tagging of PpSec13a with ProtA-tag + Neo$^R$ (template: pDP002 plasmid)**

| PCR reaction | Primer name | Primer sequence (5′ to 3′) | PCR product size (bp) |
|---|---|---|---|
| PCR A | A-Sec13a-Fw | GGAAAGAGGCCGCGACGGCACGAGCAAACCAGGAAGGAGGGAGCGCAAAC | 1791 |
| | A-Sec13a-Rv | **CCCGACCCGGAGCCGCTGCC**GTTCGCCAGCGGTTCGTTGA | |
| PCR B | B-Sec13a-Fw | **CTTGCCCGAGCGGGGTGTTACGTGCTGCAAGTTTAAGCTT**AACTGAGACCGGAGT CTCCTGTCCTGT | 2292 |
| | B-Sec13a-Rv | TGGATCGAGATGCATTCCCTCATATGCACACGGGACCTGAGTCCT | |
| PCR C | C-tag-Fw | GGCAGCGGCTCCGGGTCGGGTAG | 2276 |
| | C-tag-Rv | AAGCTTAAACTTGCAGCACGTAACACCCCGCTCGG | |
| PCR D (Nested PCR) | D-Sec13a-Fw | CCAAAGTACTCGACGTGGC | 6145 |

**B. C-terminal tagging of PpHyp24 with V5-tag + Hyg$^R$ (template: pDP011 plasmid)**

| PCR reaction | Primer name | Primer sequence (5′ to 3′) | PCR product size (bp) |
|---|---|---|---|
| PCR A | A-Hyp24-Fw | GGCCCAACGGGCGTATCAGTGAGT | 2040 |
| | A-Hyp24-Rv | **CGATCCGCTACCCGACCCGGAGCCGCTGCC**TGCATTAAAGAAGGACTGCACGTGC | |
| PCR B | B-Hyp24-Fw | **CTTGCCCGAGCGGGGTGTTACGTGCTGCAAGTTTAAGCTT**AACGCCGTTCGGCGA CCCAAACG | 1983 |
| | B-Hyp24-Rv | GCAGCCCCTCGAGTATTTCCGGC | |
| PCR C | C-tag-Fw | GGCAGCGGCTCCGGGTCGGGTAG | 1949 |
| | C-tag-Rv | AAGCTTAAACTTGCAGCACGTAACACCCCGCTCGG | |
| PCR D (Nested PCR) | D-Hyp24-Fw | GTTTGCAGAACATCGCGACCCTGGC | 5486 |
| | D-Hyp24-Rv | GTTGTACGGTGGGAGGGGTGCTG | |

**C. C-terminal tagging of PpSec13b with V5-tag + Hyg$^R$ (template: pDP011 plasmid)**

| PCR reaction | Primer name | Primer sequence (5′ to 3′) | PCR product size (bp) |
|---|---|---|---|
| PCR A | A-Sec13b-Fw | GCGCGCCAAGATGTGTCCCCTC | 1911 |
| | A-Sec13b-Rv | **CGATCCGCTACCCGACCCGGAGCCGCTGCC**CTGCGTGATGCGGGACATTTCCTTC | |
| PCR B | B-Sec13b-Fw | **CTTGCCCGAGCGGGGTGTTACGTGCTGCAAGTTTAAGCTT**AACCGTGTGTCCACC CGTTAAGTGTCGA | 2035 |
| | B-Sec13b-Rv | TGGGTGGACGCCGGATTTCTTATCAGAG | |
| PCR C | C-tag-Fw | GGCAGCGGCTCCGGGTCGGGTAG | 1949 |
| | C-tag-Rv | AAGCTTAAACTTGCAGCACGTAACACCCCGCTCGG | |
| PCR D (Nested PCR) | D-Sec13b-Fw | GCATTCACGACGTCCAGGTGCGTTCA | 5507 |
| | D-Sec13b-Rv | GAGTGTGGTCGGCTTGTGAACA | |

**Supplementary Table 4.** Mass spectrometry results from triplicate experiments for PpSec13a-PrA (IPP150 buffer) (A), PpHyp24-V5 (IPP150 buffer) (B), PpSec13a-PrA (CHAPS buffer) (C), PpSec13b-V5 (IPP150 buffer) (D). For whole MS data see Suppl. Data 1.

**A. Sec13a-PrA (IPP150 buffer) + IgG Sepharose beads**

| Locus | Name | Mol. weight [kDa] | Unique peptides | Sequence coverage [%]* |
|---|---|---|---|---|
| DIPPA_24951 | **PpSec13a** | 34,296 | 12 | 49,5 |
| DIPPA_09941 | PpSec31 | 130,25 | 63 | 58,2 |
| DIPPA_09665 | PpHyp24 | 24,214 | 12 | 65,5 |
| DIPPA_31723 | PpHyp27 | 26,81 | 10 | 39,2 |

**B.PpHyp24-V5 (IPP150 buffer) + IgG Sepharose beads**

| Locus | Name | Mol. weight [kDa] | Unique peptides | Sequence coverage [%]* |
|---|---|---|---|---|
| DIPPA_09665 | **PpHyp24** | 24,214 | 18 | 80,1 |
| DIPPA_24951 | PpSec13a | 34,296 | 3 | 22,8 |
| DIPPA_09941 | PpSec31 | 130,25 | 9 | 20,2 |
| DIPPA_31723 | PpHyp27 | 26,81 | 12 | 52,1 |

**C. Sec13a-PrA (CHAPS buffer) + IgG Sepharose beads**

| Locus | Name | Mol. weight [kDa] | Unique peptides | Sequence coverage [%]* |
|---|---|---|---|---|
| DIPPA_24951 | **PpSec13a** | 35,549 | 7 | 35,5 |
| DIPPA_09941 | PpSec31 | 130,25 | 27 | 35,2 |
| DIPPA_22976 | PpNup107 | 108,64 | 17 | 27,8 |
| DIPPA_27525 | PpNup98-Nup96 | 73,667 | 17 | 36,8 |
| DIPPA_16310 | PpNup85 | 82,647 | 11 | 24,4 |
| DIPPA_30206 | PpSeh1 | 40,36 | 12 | 50,8 |

**D. Sec13b-V5 (IPP150 buffer) + V5-trapped magnetic beads**

| Locus | Name | Mol. weight [kDa] | Unique peptides | Sequence coverage [%]* |
|---|---|---|---|---|
| DIPPA_35934 | **PpSec13b** | 34,042 | 17 | 74,2 |
| DIPPA_10282 | PpSec16 | 193,19 | 26 | 20,7 |
| DIPPA_06252 | PpSea3 | 95,466 | 12 | 22,8 |
| DIPPA_21857 | PpSea4 | 119,14 | 8 | 17,7 |

* maximum observed peptide coverage

**Supplementary Table 5.** List of Seh1, Nup85, Nup98, and Nup107 proteins in studied diplonemids and their reverse best hits.

| Query | | | FWD BLAST hit | | | | REV BLAST hit - *T. brucei* | | | REV BLAST hit - *H. sapiens* | | | REV BLAST hit - *S. cerevisiae* | | | Predicted protein |
|---|---|---|---|---|---|---|---|---|---|---|---|---|---|---|---|---|
| **Protein** | **ID** | **Organism** | **Organism** | **ID** | **Sequence** | **E-value** | **ID** | **Description** | **E-value** | **ID** | **Description** | **E-value** | **ID** | **Description** | **E-value** | |
| **Seh1** | - | - | *Paradiplonema papillatum* | DIPPA_30206.mRNA.1 | MEPTPVGDGFS | found by pull-down | Tb927.10.14180 | gene_product=protein transport protein SEC13 | 8.80E-14 | NP_112493.2 | nucleoporin SEH1 isoform 2 [Homo sapiens] | 6.16E-46 | NP_011415.1 | Seh1p [Saccharomyces cerevisiae S288C] | 1.33E-37 | Seh1 |
| **Nup85** | - | - | *Paradiplonema papillatum* | DIPPA_16310.mRNA.1 | MEPKQFSQLLSI | found by pull-down | no hit found | | | no hit found | | | no hit found | | | Nup85 |
| **Nup98-96** | - | - | *Paradiplonema papillatum* | DIPPA_27525.mRNA.1 | MVRIGLVRKRA | found by pull-down | Tb927.3.4190 | gene_product=endosomal integral membrane protein, putative | 3.87E-02 | NP_001352054.1 | nuclear pore complex protein Nup98-Nup96 isoform 5 [Homo sapiens] | 4.60E-11 | no hit found | | | Nup96 |
| **Nup107** | - | - | *Paradiplonema papillatum* | DIPPA_22976.mRNA.1 | MYTPGKFDSIVI | found by pull-down | no hit found | | | NP_065134.1 | nuclear pore complex protein Nup107 isoform 1 [Homo sapiens] | 1.22E-08 | NP_015247.1 | F1F0 ATP synthase subunit 4 [Saccharomyces cerevisiae S288C] | 9.31E-01 | Nup107 |
| **Seh1** | DIPPA_30206.mRNA.1 | *Paradiplonema papillatum* | *Diplonema japonicum* | TRINITY_DN9964_c4_g1_i2 | MDIDEGHPSGH | 1.76E-137 | Tb927.10.14180 | gene_product=protein transport protein SEC13 | 9.88E-16 | NP_112493.2 | nucleoporin SEH1 isoform 2 [Homo sapiens] | 5.85E-43 | NP_011415.1 | Seh1p [Saccharomyces cerevisiae S288C] | 1.27E-37 | Seh1 |
| **Nup85** | DIPPA_16310.mRNA.1 | *Paradiplonema papillatum* | *Diplonema japonicum* | TRINITY_DN10163_c6_g9_i1 | MANKSLAAVLC | 0 | no hit found | | | no hit found | | | NP_014699.1 | rRNA-binding endoribonuclease [Saccharomyces cerevisiae S288C] | 4.05E-01 | - |
| **Nup98-96** | DIPPA_27525.mRNA.1 | *Paradiplonema papillatum* | *Diplonema japonicum* | TRINITY_DN10246_c2_g5_i1 | MVRIGLLKTARE | 2.77E-119 | Tb11.0590 | gene_product=variant surface glycoprotein (VSG), putative | 1.58E-01 | NP_057404.2 | nuclear pore complex protein Nup98-Nup96 isoform 1 precursor [Homo sapiens] | 4.17E-08 | no hit found | | | Nup96 |
| **Nup107** | DIPPA_22976.mRNA.1 | *Paradiplonema papillatum* | *Diplonema japonicum* | TRINITY_DN9997_c4_g2_i1 | MEFTPSHGLASC | 0 | no hit found | | | XP_005269094.1 | nuclear pore complex protein Nup107 isoform X1 [Homo sapiens] | 4.28E-03 | no hit found | | | Nup107 |
| **Seh1** | DIPPA_30206.mRNA.1 | *Paradiplonema papillatum* | *Rhynchopus humris* | TRINITY_DN29702_c0_g1_i1 | MGYPPPFSSNV | 2.20E-132 | Tb927.11.13850 | gene_product=hypothetical protein, conserved | 1.05E-11 | XP_011524046.1 | nucleoporin SEH1 isoform X3 [Homo sapiens] | 3.36E-40 | NP_013309.1 | GTPase-activating protein SEC13 [Saccharomyces cerevisiae S288C] | 8.40E-30 | Seh1 |
| **Nup85** | DIPPA_16310.mRNA.1 | *Paradiplonema papillatum* | *Rhynchopus humris* | TRINITY_DN10810_c0_g1_i1 | MCAGTGSVPAV | 4.00E-141 | Tb927.5.120 | gene_product=expression site-associated gene 9 (ESAG9) protein, putative | 5.43E-01 | no hit found | | | NP_012511.1 | Pet130p [Saccharomyces cerevisiae S288C] | 4.56E-02 | Nup85 |
| **Nup98-96** | DIPPA_27525.mRNA.1 | *Paradiplonema papillatum* | *Rhynchopus humris* | TRINITY_DN63797_c0_g1_i1 | VQVRAANGAFA | 5.37E-44 | Tb927.11.14700 | gene_product=hypothetical protein, conserved | 5.94E-01 | NP_001352058.1 | nuclear pore complex protein Nup98-Nup96 isoform 9 [Homo sapiens] | 1.94E-08 | no hit found | | | Nup96 |
| **Nup107** | DIPPA_22976.mRNA.1 | *Paradiplonema papillatum* | *Rhynchopus humris* | TRINITY_DN57352_c0_g1_i1 | MASEAITMCED | 2.48E-173 | Tb927.11.2950 | gene_product=Nucleoporin NUP89 | 1.63E-03 | NP_065134.1 | nuclear pore complex protein Nup107 isoform 1 [Homo sapiens] | 1.74E-09 | no hit found | | | Nup107 |
| **Seh1** | DIPPA_30206.mRNA.1 | *Paradiplonema papillatum* | *Lacrimia lanifica* | TRINITY_DN8930_c0_g1_i1 | MNATAAAVAA | 3.89E-124 | Tb927.11.8120 | gene_product=protein transport protein SEC13, putative | 4.35E-19 | NP_112493.2 | nucleoporin SEH1 isoform 2 [Homo sapiens] | 2.99E-45 | NP_011415.1 | Seh1p [Saccharomyces cerevisiae S288C] | 2.57E-32 | Seh1 |
| **Nup85** | DIPPA_16310.mRNA.1 | *Paradiplonema papillatum* | *Lacrimia lanifica* | TRINITY_DN9810_c0_g1_i1 | MSQDMDAETM | 4.32E-120 | no hit found | | | no hit found | | | no hit found | | | - |
| **Nup98-96** | DIPPA_27525.mRNA.1 | *Paradiplonema papillatum* | *Lacrimia lanifica* | TRINITY_DN11343_c0_g11_i1 | MVRIGLRKKGPI | 1.18E-62 | Tb10.v4.0088 | gene_product=variant surface glycoprotein (VSG), putative | 2.03E-01 | NP_001352057.1 | nuclear pore complex protein Nup98-Nup96 isoform 8 precursor [Homo sapiens] | 2.59E-08 | no hit found | | | Nup96 |
| **Nup107** | DIPPA_22976.mRNA.1 | *Paradiplonema papillatum* | *Lacrimia lanifica* | TRINITY_DN10186_c0_g1_i1 | MMSAMSATPLA | 4.97E-179 | Tb927.6.1200 | gene_product=Polyadenylation mediator complex protein PAMC2, putative | 5.70E-01 | XP_005269094.1 | nuclear pore complex protein Nup107 isoform X1 [Homo sapiens] | 1.13E-07 | NP_011497.1 | J-type chaperone JAC1 [Saccharomyces cerevisiae S288C] | 5.30E-01 | Nup107 |
| **Seh1** | DIPPA_30206.mRNA.1 | *Paradiplonema papillatum* | *Sulcionema specki* | TRINITY_DN24505_c0_g1_i1 | MPAADPSTGTP | 2.37E-109 | Tb927.11.8120 | gene_product=protein transport protein SEC13, putative | 3.00E-17 | XP_011524046.1 | nucleoporin SEH1 isoform X3 [Homo sapiens] | 2.25E-47 | NP_011415.1 | Seh1p [Saccharomyces cerevisiae S288C] | 5.72E-36 | Seh1 |
| **Nup85** | DIPPA_16310.mRNA.1 | *Paradiplonema papillatum* | *Sulcionema specki* | TRINITY_DN101444_c0_g1_i1 | MQGITGSSRLCI | 1.10E-171 | Tb927.2.5750 | gene_product=tyrosyl-DNA Phosphodiesterase (Tdp1), putative | 2.58E-02 | XP_016880599.1 | nuclear pore complex protein Nup85 isoform X3 [Homo sapiens] | 9.88E-03 | NP_116687.1 | condensin subunit SMC2 [Saccharomyces cerevisiae S288C] | 2.09E-01 | - |
| **Nup98-96** | DIPPA_27525.mRNA.1 | *Paradiplonema papillatum* | *Sulcionema specki* | TRINITY_DN79343_c0_g1_i1 | MVRIGFVKKRT( | 2.85E-47 | no hit found | | | NP_001352054.1 | nuclear pore complex protein Nup98-Nup96 isoform 5 [Homo sapiens] | 4.75E-13 | NP_013652.1 | lysophospholipase [Saccharomyces cerevisiae S288C] | 5.00E-01 | Nup96 |
| **Nup107** | DIPPA_22976.mRNA.1 | *Paradiplonema papillatum* | *Sulcionema specki* | TRINITY_DN66982_c0_g1_i1 | MQTPRVDSALV | 3.18E-156 | no hit found | | | NP_065134.1 | nuclear pore complex protein Nup107 isoform 1 [Homo sapiens] | 1.82E-05 | no hit found | | | Nup107 |
| **Seh1** | DIPPA_30206.mRNA.1 | *Paradiplonema papillatum* | *Artemidia motanka* | TRINITY_DN5440_c0_g1_i1 | MFAEAVPAGAG | 3.40E-70 | Tb927.10.14180 | gene_product=protein transport protein SEC13 | 3.70E-20 | NP_112493.2 | nucleoporin SEH1 isoform 2 [Homo sapiens] | 1.01E-45 | NP_011415.1 | Seh1p [Saccharomyces cerevisiae S288C] | 4.01E-35 | Seh1 |
| **Nup85** | DIPPA_16310.mRNA.1 | *Paradiplonema papillatum* | *Artemidia motanka* | TRINITY_DN22359_c0_g1_i1 | MELDRVRAMP( | 3.45E-29 | no hit found | | | no hit found | | | no hit found | | | - |
| **Nup98-96** | DIPPA_27525.mRNA.1 | *Paradiplonema papillatum* | *Artemidia motanka* | TRINITY_DN8787_c0_g1_i2 | MGRKVGLGSAP | 1.19E-12 | Tb927.10.3800 | gene_product=predicted RING finger protein | 8.09E-01 | NP_057404.2 | nuclear pore complex protein Nup98-Nup96 isoform 1 precursor [Homo sapiens] | 5.89E-04 | no hit found | | | Nup96 |
| **Nup107** | DIPPA_22976.mRNA.1 | *Paradiplonema papillatum* | *Artemidia motanka* | TRINITY_DN13201_c0_g1_i1 | MLAPAARLCPA | 5.30E-41 | Tb927.11.3390 | gene_product=Concanavalin A-like lectin/glucanases superfamily/SPRY domain/HECT-( | 2.26E-02 | XP_016862507.1 | E3 ubiquitin-protein ligase RNF123 isoform X1 [Homo sapiens] | 9.05E-03 | no hit found | | | Nup107: SPRY |
| **Seh1** | DIPPA_30206.mRNA.1 | *Paradiplonema papillatum* | *Namystynia karyoxenos* | TRINITY_DN49076_c0_g1_i1 | MNAPQNAGGV | 4.09E-68 | Tb927.10.14180 | gene_product=protein transport protein SEC13 | 3.43E-13 | NP_112493.2 | nucleoporin SEH1 isoform 2 [Homo sapiens] | 7.26E-42 | NP_011415.1 | Seh1p [Saccharomyces cerevisiae S288C] | 9.90E-38 | Seh1 |
| **Nup85** | DIPPA_16310.mRNA.1 | *Paradiplonema papillatum* | *Namystynia karyoxenos* | TRINITY_DN9585_c0_g1_i1 | MALHLGTALGS | 9.73E-08 | no hit found | | | no hit found | | | no hit found | | | - |
| **Nup98-96** | DIPPA_27525.mRNA.1 | *Paradiplonema papillatum* | *Namystynia karyoxenos* | TRINITY_DN22878_c0_g1_i1 | MEPDSLPTPAPI | 1.87E-12 | Tb927.8.3290 | gene_product=DNA polymerase zeta catalytic subunit, putative | 5.31E-01 | NP_001352055.1 | nuclear pore complex protein Nup98-Nup96 isoform 6 precursor [Homo sapiens] | 3.21E-07 | no hit found | | | Nup96 |
| **Nup107** | DIPPA_22976.mRNA.1 | *Paradiplonema papillatum* | *Namystynia karyoxenos* | TRINITY_DN78419_c0_g1_i1 | MFADPIETPPPV | 3.15E-42 | Tb927.11.2950 | gene_product=Nucleoporin NUP89 | 1.06E-03 | no hit found | | | NP_010539.1 | Met32p [Saccharomyces cerevisiae S288C] | 8.56E-01 | Nup107: SPRY |

**Supplementary Table 6.** Identified Sec16 proteins in organisms in EukProt database.

| | HMM hit | | | | REV BLAST hit | | | | | |
|---|---|---|---|---|---|---|---|---|---|---|
| **EukProt number** | **Lineage** | **ID** | **evalue** | **length** | **ID** | **description** | **length** | **evalue** | **bitscore** | **% identity** |
| EP00075 | Obazoa;Opisthokonta;Holozoa;Choanozoa;Metazoa | EP00075 Anas platyrhynchos 37287 | 1,60E-250 | 2205 | XP 027326540.1 | protein transport protein Sec16A isoform X6 [Anas platyrhynchos] | 2205 | 0 | 4521,07 | 100 |
| EP00076 | Obazoa;Opisthokonta;Holozoa;Choanozoa;Metazoa | EP00076 Gallus gallus 10894 | 2,90E-250 | 2201 | XP 004946034.1 | protein transport protein Sec16A isoform X8 [Gallus gallus] | 2201 | 0 | 4518,76 | 100 |
| EP00072 | Obazoa;Opisthokonta;Holozoa;Choanozoa;Metazoa | EP00072 Mus musculus 31702 | 1,10E-248 | 2161 | XP 006497977.1 | protein transport protein Sec16A isoform X5 [Mus musculus] | 2161 | 0 | 4408,59 | 100 |
| EP00072 | Obazoa;Opisthokonta;Holozoa;Choanozoa;Metazoa | EP00072 Mus musculus 58345 | 2,00E-248 | 2125 | XP 011237375.1 | protein transport protein Sec16A isoform X6 [Mus musculus] | 2125 | 0 | 4333,1 | 100 |
| EP00076 | Obazoa;Opisthokonta;Holozoa;Choanozoa;Metazoa | EP00076 Gallus gallus 31626 | 5,50E-248 | 2420 | XP 024997164.1 | protein transport protein Sec16A isoform X6 [Gallus gallus] | 2420 | 0 | 4957,5 | 100 |
| EP00076 | Obazoa;Opisthokonta;Holozoa;Choanozoa;Metazoa | EP00076 Gallus gallus 10891 | 7,00E-247 | 2437 | XP 004946030.1 | protein transport protein Sec16A isoform X4 [Gallus gallus] | 2437 | 0 | 4990,63 | 100 |
| EP00073 | Obazoa;Opisthokonta;Holozoa;Choanozoa;Metazoa | EP00073 Rattus norvegicus 21819 | 1,50E-246 | 2136 | XP 006233728.1 | PREDICTED: protein transport protein Sec16A isoform X2 [Rattus norvegicus] | 2136 | 0 | 4355,82 | 100 |
| EP00076 | Obazoa;Opisthokonta;Holozoa;Choanozoa;Metazoa | EP00076 Gallus gallus 10890 | 3,40E-246 | 2441 | XP 004946029.1 | protein transport protein Sec16A isoform X3 [Gallus gallus] | 2441 | 0 | 5000,65 | 100 |
| EP00075 | Obazoa;Opisthokonta;Holozoa;Choanozoa;Metazoa | EP00075 Anas platyrhynchos 37284 | 5,20E-246 | 2436 | XP 027326537.1 | protein transport protein Sec16A isoform X3 [Anas platyrhynchos] | 2436 | 0 | 4982,54 | 100 |
| EP00076 | Obazoa;Opisthokonta;Holozoa;Choanozoa;Metazoa | EP00076 Gallus gallus 46737 | 9,70E-246 | 2457 | XP 415419.4 | protein transport protein Sec16A isoform X1 [Gallus gallus] | 2457 | 0 | 5030,69 | 100 |
| EP00076 | Obazoa;Opisthokonta;Holozoa;Choanozoa;Metazoa | EP00076 Gallus gallus 16070 | 1,10E-245 | 2456 | XP 015135281.1 | protein transport protein Sec16A isoform X2 [Gallus gallus] | 2456 | 0 | 5029,15 | 100 |
| EP00076 | Obazoa;Opisthokonta;Holozoa;Choanozoa;Metazoa | EP00076 Gallus gallus 10893 | 2,80E-245 | 2416 | XP 004946033.1 | protein transport protein Sec16A isoform X7 [Gallus gallus] | 2416 | 0 | 4945,95 | 100 |
| EP00072 | Obazoa;Opisthokonta;Holozoa;Choanozoa;Metazoa | EP00072 Mus musculus 31697 | 4,00E-245 | 2393 | XP 006497972.1 | protein transport protein Sec16A isoform X1 [Mus musculus] >gi\|568913417\|ref\|XP 006 | 2393 | 0 | 4871,6 | 100 |
| EP00072 | Obazoa;Opisthokonta;Holozoa;Choanozoa;Metazoa | EP00072 Mus musculus 31698 | 4,00E-245 | 2393 | XP 006497972.1 | protein transport protein Sec16A isoform X1 [Mus musculus] >gi\|568913417\|ref\|XP 006 | 2393 | 0 | 4871,6 | 100 |
| EP00075 | Obazoa;Opisthokonta;Holozoa;Choanozoa;Metazoa | EP00075 Anas platyrhynchos 37282 | 5,70E-245 | 2440 | XP 027326535.1 | protein transport protein Sec16A isoform X2 [Anas platyrhynchos] | 2440 | 0 | 4990,63 | 100 |
| EP00076 | Obazoa;Opisthokonta;Holozoa;Choanozoa;Metazoa | EP00076 Gallus gallus 10892 | 6,00E-245 | 2437 | XP 004946031.1 | protein transport protein Sec16A isoform X5 [Gallus gallus] | 2437 | 0 | 4986,01 | 100 |
| EP00075 | Obazoa;Opisthokonta;Holozoa;Choanozoa;Metazoa | EP00075 Anas platyrhynchos 37285 | 7,10E-244 | 2415 | XP 027326538.1 | protein transport protein Sec16A isoform X4 [Anas platyrhynchos] | 2415 | 0 | 4937,47 | 100 |
| EP00075 | Obazoa;Opisthokonta;Holozoa;Choanozoa;Metazoa | EP00075 Anas platyrhynchos 37286 | 3,50E-243 | 2410 | XP 027326539.1 | protein transport protein Sec16A isoform X5 [Anas platyrhynchos] | 2410 | 0 | 4932,85 | 100 |
| EP00075 | Obazoa;Opisthokonta;Holozoa;Choanozoa;Metazoa | EP00075 Anas platyrhynchos 2410 | 1,80E-242 | 2456 | XP 027326534.1 | protein transport protein Sec16A isoform X1 [Anas platyrhynchos] >gi\|1540399886\|ref\|X | 2456 | 0 | 5018,75 | 100 |
| EP00075 | Obazoa;Opisthokonta;Holozoa;Choanozoa;Metazoa | EP00075 Anas platyrhynchos 2411 | 1,80E-242 | 2456 | XP 027326534.1 | protein transport protein Sec16A isoform X1 [Anas platyrhynchos] >gi\|1540399886\|ref\|X | 2456 | 0 | 5018,75 | 100 |
| EP00075 | Obazoa;Opisthokonta;Holozoa;Choanozoa;Metazoa | EP00075 Anas platyrhynchos 37281 | 1,80E-242 | 2456 | XP 027326534.1 | protein transport protein Sec16A isoform X1 [Anas platyrhynchos] >gi\|1540399886\|ref\|X | 2456 | 0 | 5018,75 | 100 |
| EP00073 | Obazoa;Opisthokonta;Holozoa;Choanozoa;Metazoa | EP00073 Rattus norvegicus 21816 | 1,80E-241 | 2366 | XP 006233724.1 | PREDICTED: protein transport protein Sec16A isoform X1 [Rattus norvegicus] >gi\|564339 | 2366 | 0 | 4821,53 | 100 |
| EP00073 | Obazoa;Opisthokonta;Holozoa;Choanozoa;Metazoa | EP00073 Rattus norvegicus 21817 | 1,80E-241 | 2366 | XP 006233724.1 | PREDICTED: protein transport protein Sec16A isoform X1 [Rattus norvegicus] >gi\|564339 | 2366 | 0 | 4821,53 | 100 |
| EP00073 | Obazoa;Opisthokonta;Holozoa;Choanozoa;Metazoa | EP00073 Rattus norvegicus 21818 | 1,80E-241 | 2366 | XP 006233724.1 | PREDICTED: protein transport protein Sec16A isoform X1 [Rattus norvegicus] >gi\|564339 | 2366 | 0 | 4821,53 | 100 |
| EP00072 | Obazoa;Opisthokonta;Holozoa;Choanozoa;Metazoa | EP00072 Mus musculus 31701 | 2,60E-241 | 2331 | XP 006497976.1 | protein transport protein Sec16A isoform X4 [Mus musculus] | 2331 | 0 | 4743,72 | 100 |
| EP00072 | Obazoa;Opisthokonta;Holozoa;Choanozoa;Metazoa | EP00072 Mus musculus 26093 | 3,90E-240 | 2357 | NP 694765.2 | protein transport protein Sec16A [Mus musculus] | 2357 | 0 | 4794,95 | 100 |
| EP00072 | Obazoa;Opisthokonta;Holozoa;Choanozoa;Metazoa | EP00072 Mus musculus 31700 | 9,60E-240 | 2373 | XP 006497975.1 | protein transport protein Sec16A isoform X3 [Mus musculus] | 2373 | 0 | 4835,01 | 100 |
| EP00072 | Obazoa;Opisthokonta;Holozoa;Choanozoa;Metazoa | EP00072 Mus musculus 31699 | 1,60E-239 | 2377 | XP 006497974.1 | protein transport protein Sec16A isoform X2 [Mus musculus] | 2377 | 0 | 4836,17 | 100 |
| EP00071 | Obazoa;Opisthokonta;Holozoa;Choanozoa;Metazoa | EP00071 Canis lupus 45730 | 1,20E-238 | 2098 | XP 022279287.1 | protein transport protein Sec16A isoform X9 [Canis lupus familiaris] | 2098 | 0 | 4227,94 | 100 |
| EP00071 | Obazoa;Opisthokonta;Holozoa;Choanozoa;Metazoa | EP00071 Canis lupus 8232 | 4,50E-238 | 2120 | XP 005625163.1 | protein transport protein Sec16A isoform X8 [Canis lupus familiaris] | 2120 | 0 | 4273,39 | 100 |
| EP00074 | Obazoa;Opisthokonta;Holozoa;Choanozoa;Metazoa | EP00074 Homo sapiens 67256 | 2,60E-236 | 2142 | XP 011517564.1 | protein transport protein Sec16A isoform X15 [Homo sapiens] | 2142 | 0 | 4326,93 | 100 |
| EP00078 | Obazoa;Opisthokonta;Holozoa;Choanozoa;Metazoa | EP00078 Anolis carolinensis 7745 | 1,20E-234 | 2223 | XP 003229913.2 | PREDICTED: protein transport protein Sec16A [Anolis carolinensis] | 2223 | 0 | 4539,95 | 100 |
| EP00071 | Obazoa;Opisthokonta;Holozoa;Choanozoa;Metazoa | EP00071 Canis lupus 45729 | 2,30E-233 | 2297 | XP 022279286.1 | protein transport protein Sec16A isoform X7 [Canis lupus familiaris] | 2297 | 0 | 4627,39 | 100 |
| EP00077 | Obazoa;Opisthokonta;Holozoa;Choanozoa;Metazoa | EP00077 Pelodiscus sinensis 19322 | 6,10E-231 | 2480 | XP 014427834.1 | protein transport protein Sec16A [Pelodiscus sinensis] | 2480 | 0 | 5074,22 | 100 |
| EP00073 | Obazoa;Opisthokonta;Holozoa;Choanozoa;Metazoa | EP00073 Rattus norvegicus 11957 | 4,80E-230 | 2346 | NP 001263346.1 | protein transport protein Sec16A [Rattus norvegicus] | 2346 | 0 | 4778 | 100 |
| EP00071 | Obazoa;Opisthokonta;Holozoa;Choanozoa;Metazoa | EP00071 Canis lupus 22287 | 6,40E-230 | 2336 | XP 013972403.1 | protein transport protein Sec16A isoform X6 [Canis lupus familiaris] | 2336 | 0 | 4700,58 | 100 |
| EP00074 | Obazoa;Opisthokonta;Holozoa;Choanozoa;Metazoa | EP00074 Homo sapiens 67247 | 6,70E-230 | 2356 | XP 011517555.1 | protein transport protein Sec16A isoform X6 [Homo sapiens] | 2356 | 0 | 4754,5 | 100 |
| EP00071 | Obazoa;Opisthokonta;Holozoa;Choanozoa;Metazoa | EP00071 Canis lupus 8231 | 8,50E-230 | 2337 | XP 005625162.1 | protein transport protein Sec16A isoform X5 [Canis lupus familiaris] | 2337 | 0 | 4702,12 | 100 |
| EP00071 | Obazoa;Opisthokonta;Holozoa;Choanozoa;Metazoa | EP00071 Canis lupus 8228 | 1,00E-229 | 2356 | XP 005625159.1 | protein transport protein Sec16A isoform X2 [Canis lupus familiaris] | 2356 | 0 | 4746,42 | 100 |
| EP00071 | Obazoa;Opisthokonta;Holozoa;Choanozoa;Metazoa | EP00071 Canis lupus 8229 | 8,10E-229 | 2356 | XP 005625160.1 | protein transport protein Sec16A isoform X3 [Canis lupus familiaris] | 2356 | 0 | 4746,8 | 100 |
| EP00071 | Obazoa;Opisthokonta;Holozoa;Choanozoa;Metazoa | EP00071 Canis lupus 8230 | 8,20E-229 | 2355 | XP 005625161.1 | protein transport protein Sec16A isoform X4 [Canis lupus familiaris] | 2355 | 0 | 4744,87 | 100 |
| EP00071 | Obazoa;Opisthokonta;Holozoa;Choanozoa;Metazoa | EP00071 Canis lupus 22286 | 1,10E-228 | 2357 | XP 005625154.1 | protein transport protein Sec16A isoform X1 [Canis lupus familiaris] >gi\|545512618\|ref\|X | 2357 | 0 | 4748,34 | 100 |
| EP00071 | Obazoa;Opisthokonta;Holozoa;Choanozoa;Metazoa | EP00071 Canis lupus 8225 | 1,10E-228 | 2357 | XP 005625154.1 | protein transport protein Sec16A isoform X1 [Canis lupus familiaris] >gi\|545512618\|ref\|X | 2357 | 0 | 4748,34 | 100 |
| EP00071 | Obazoa;Opisthokonta;Holozoa;Choanozoa;Metazoa | EP00071 Canis lupus 8226 | 1,10E-228 | 2357 | XP 005625154.1 | protein transport protein Sec16A isoform X1 [Canis lupus familiaris] >gi\|545512618\|ref\|X | 2357 | 0 | 4748,34 | 100 |
| EP00071 | Obazoa;Opisthokonta;Holozoa;Choanozoa;Metazoa | EP00071 Canis lupus 8227 | 1,10E-228 | 2357 | XP 005625154.1 | protein transport protein Sec16A isoform X1 [Canis lupus familiaris] >gi\|545512618\|ref\|X | 2357 | 0 | 4748,34 | 100 |
| EP00079 | Obazoa;Opisthokonta;Holozoa;Choanozoa;Metazoa | EP00079 Xenopus tropicalis 17056 | 5,80E-228 | 2362 | XP 004916695.1 | PREDICTED: protein transport protein Sec16A isoform X2 [Xenopus tropicalis] | 2362 | 0 | 4804,19 | 100 |
| EP00079 | Obazoa;Opisthokonta;Holozoa;Choanozoa;Metazoa | EP00079 Xenopus tropicalis 33294 | 6,00E-228 | 2365 | XP 017947065.1 | PREDICTED: protein transport protein Sec16A isoform X1 [Xenopus tropicalis] >gi\|106283 | 2365 | 0 | 4805,35 | 100 |
| EP00079 | Obazoa;Opisthokonta;Holozoa;Choanozoa;Metazoa | EP00079 Xenopus tropicalis 33295 | 6,00E-228 | 2365 | XP 017947065.1 | PREDICTED: protein transport protein Sec16A isoform X1 [Xenopus tropicalis] >gi\|106283 | 2365 | 0 | 4805,35 | 100 |
| EP00074 | Obazoa;Opisthokonta;Holozoa;Choanozoa;Metazoa | EP00074 Homo sapiens 67257 | 1,70E-225 | 2100 | XP 011517565.1 | protein transport protein Sec16A isoform X16 [Homo sapiens] | 2100 | 0 | 4240,65 | 100 |
| EP00068 | Obazoa;Opisthokonta;Holozoa;Choanozoa;Metazoa | EP00068 Latimeria chalumnae 25885 | 1,40E-221 | 2504 | XP 014346202.1 | PREDICTED: protein transport protein Sec16A [Latimeria chalumnae] | 2504 | 0 | 5134,7 | 100 |
| EP00079 | Obazoa;Opisthokonta;Holozoa;Choanozoa;Metazoa | EP00079 Xenopus tropicalis 10427 | 1,80E-221 | 2342 | XP 002935278.2 | PREDICTED: protein transport protein Sec16A isoform X4 [Xenopus tropicalis] | 2342 | 0 | 4754,12 | 100 |
| EP00079 | Obazoa;Opisthokonta;Holozoa;Choanozoa;Metazoa | EP00079 Xenopus tropicalis 33296 | 3,50E-221 | 2345 | XP 017947067.1 | PREDICTED: protein transport protein Sec16A isoform X3 [Xenopus tropicalis] | 2345 | 0 | 4762,59 | 100 |
| EP00074 | Obazoa;Opisthokonta;Holozoa;Choanozoa;Metazoa | EP00074 Homo sapiens 67251 | 6,20E-221 | 2332 | XP 011517559.1 | protein transport protein Sec16A isoform X11 [Homo sapiens] | 2332 | 0 | 4710,98 | 100 |
| EP00074 | Obazoa;Opisthokonta;Holozoa;Choanozoa;Metazoa | EP00074 Homo sapiens 67253 | 8,90E-221 | 2312 | XP 011517561.1 | protein transport protein Sec16A isoform X13 [Homo sapiens] | 2312 | 0 | 4666,29 | 100 |
| EP00074 | Obazoa;Opisthokonta;Holozoa;Choanozoa;Metazoa | EP00074 Homo sapiens 14770 | 2,20E-220 | 2334 | NP 001263347.1 | protein transport protein Sec16A isoform 2 [Homo sapiens] >gi\|767958873\|ref\|XP 0115 | 2334 | 0 | 4710,21 | 100 |
| EP00074 | Obazoa;Opisthokonta;Holozoa;Choanozoa;Metazoa | EP00074 Homo sapiens 67250 | 2,20E-220 | 2334 | NP 001263347.1 | protein transport protein Sec16A isoform 2 [Homo sapiens] >gi\|767958873\|ref\|XP 0115 | 2334 | 0 | 4710,21 | 100 |
| EP00073 | Obazoa;Opisthokonta;Holozoa;Choanozoa;Metazoa | EP00073 Rattus norvegicus 39807 | 3,10E-220 | 1057 | XP 008767904.1 | PREDICTED: protein transport protein Sec16B isoform X1 [Rattus norvegicus] >gi\|6720745 | 1057 | 0 | 2185,22 | 100 |
| EP00073 | Obazoa;Opisthokonta;Holozoa;Choanozoa;Metazoa | EP00073 Rattus norvegicus 39808 | 3,10E-220 | 1057 | XP 008767904.1 | PREDICTED: protein transport protein Sec16B isoform X1 [Rattus norvegicus] >gi\|6720745 | 1057 | 0 | 2185,22 | 100 |
| EP00073 | Obazoa;Opisthokonta;Holozoa;Choanozoa;Metazoa | EP00073 Rattus norvegicus 52053 | 3,10E-220 | 1057 | XP 008767904.1 | PREDICTED: protein transport protein Sec16B isoform X1 [Rattus norvegicus] >gi\|6720745 | 1057 | 0 | 2185,22 | 100 |
| EP00073 | Obazoa;Opisthokonta;Holozoa;Choanozoa;Metazoa | EP00073 Rattus norvegicus 52054 | 3,10E-220 | 1057 | XP 008767904.1 | PREDICTED: protein transport protein Sec16B isoform X1 [Rattus norvegicus] >gi\|6720745 | 1057 | 0 | 2185,22 | 100 |
| EP00073 | Obazoa;Opisthokonta;Holozoa;Choanozoa;Metazoa | EP00073 Rattus norvegicus 52055 | 3,10E-220 | 1057 | XP 008767904.1 | PREDICTED: protein transport protein Sec16B isoform X1 [Rattus norvegicus] >gi\|6720745 | 1057 | 0 | 2185,22 | 100 |
| EP00073 | Obazoa;Opisthokonta;Holozoa;Choanozoa;Metazoa | EP00073 Rattus norvegicus 52056 | 3,10E-220 | 1057 | XP 008767904.1 | PREDICTED: protein transport protein Sec16B isoform X1 [Rattus norvegicus] >gi\|6720745 | 1057 | 0 | 2185,22 | 100 |
| EP00062 | Obazoa;Opisthokonta;Holozoa;Choanozoa;Metazoa | EP00062 Callorhinchus milii 17239 | 5,00E-220 | 2357 | XP 007901056.1 | PREDICTED: protein transport protein Sec16A isoform X4 [Callorhinchus milii] | 2357 | 0 | 4792,25 | 100 |
| EP00074 | Obazoa;Opisthokonta;Holozoa;Choanozoa;Metazoa | EP00074 Homo sapiens 93405 | 6,90E-220 | 2351 | XP 016870852.1 | protein transport protein Sec16A isoform X8 [Homo sapiens] | 2351 | 0 | 4746,42 | 100 |
| EP00074 | Obazoa;Opisthokonta;Holozoa;Choanozoa;Metazoa | EP00074 Homo sapiens 43974 | 2,20E-219 | 1060 | NP 149118.2 | protein transport protein Sec16B isoform 1 [Homo sapiens] | 1060 | 0 | 2180,99 | 100 |
| EP00074 | Obazoa;Opisthokonta;Holozoa;Choanozoa;Metazoa | EP00074 Homo sapiens 67249 | 2,20E-219 | 2337 | XP 011517557.1 | protein transport protein Sec16A isoform X9 [Homo sapiens] | 2337 | 0 | 4714,83 | 100 |
| EP00074 | Obazoa;Opisthokonta;Holozoa;Choanozoa;Metazoa | EP00074 Homo sapiens 31140 | 3,20E-219 | 1061 | NP 001343435.1 | protein transport protein Sec16B [Homo sapiens] >gi\|1251775589\|ref\|NP 001343434.1\| | 1061 | 0 | 2184,07 | 100 |
| EP00074 | Obazoa;Opisthokonta;Holozoa;Choanozoa;Metazoa | EP00074 Homo sapiens 31141 | 3,20E-219 | 1061 | NP 001343435.1 | protein transport protein Sec16B [Homo sapiens] >gi\|1251775589\|ref\|NP 001343434.1\| | 1061 | 0 | 2184,07 | 100 |
| EP00074 | Obazoa;Opisthokonta;Holozoa;Choanozoa;Metazoa | EP00074 Homo sapiens 37981 | 3,50E-219 | 2357 | NP 055681.1 | protein transport protein Sec16A isoform 1 [Homo sapiens] >gi\|767958865\|ref\|XP 0115 | 2357 | 0 | 4753,73 | 100 |
| EP00074 | Obazoa;Opisthokonta;Holozoa;Choanozoa;Metazoa | EP00074 Homo sapiens 67246 | 3,50E-219 | 2357 | NP 055681.1 | protein transport protein Sec16A isoform 1 [Homo sapiens] >gi\|767958865\|ref\|XP 0115 | 2357 | 0 | 4753,73 | 100 |
| EP00074 | Obazoa;Opisthokonta;Holozoa;Choanozoa;Metazoa | EP00074 Homo sapiens 67248 | 5,40E-219 | 2354 | XP 011517556.1 | protein transport protein Sec16A isoform X7 [Homo sapiens] | 2354 | 0 | 4750,65 | 100 |
| EP00074 | Obazoa;Opisthokonta;Holozoa;Choanozoa;Metazoa | EP00074 Homo sapiens 67244 | 2,80E-218 | 2374 | XP 011517552.1 | protein transport protein Sec16A isoform X3 [Homo sapiens] | 2374 | 0 | 4788,4 | 100 |
| EP00074 | Obazoa;Opisthokonta;Holozoa;Choanozoa;Metazoa | EP00074 Homo sapiens 55243 | 5,80E-218 | 2379 | XP 005266191.1 | protein transport protein Sec16A isoform X1 [Homo sapiens] >gi\|767958853\|ref\|XP 011 | 2379 | 0 | 4795,72 | 100 |
| EP00074 | Obazoa;Opisthokonta;Holozoa;Choanozoa;Metazoa | EP00074 Homo sapiens 67240 | 5,80E-218 | 2379 | XP 005266191.1 | protein transport protein Sec16A isoform X1 [Homo sapiens] >gi\|767958853\|ref\|XP 011 | 2379 | 0 | 4795,72 | 100 |
| EP00074 | Obazoa;Opisthokonta;Holozoa;Choanozoa;Metazoa | EP00074 Homo sapiens 67241 | 5,80E-218 | 2379 | XP 005266191.1 | protein transport protein Sec16A isoform X1 [Homo sapiens] >gi\|767958853\|ref\|XP 011 | 2379 | 0 | 4795,72 | 100 |
| EP00074 | Obazoa;Opisthokonta;Holozoa;Choanozoa;Metazoa | EP00074 Homo sapiens 67242 | 5,80E-218 | 2379 | XP 005266191.1 | protein transport protein Sec16A isoform X1 [Homo sapiens] >gi\|767958853\|ref\|XP 011 | 2379 | 0 | 4795,72 | 100 |
| EP00074 | Obazoa;Opisthokonta;Holozoa;Choanozoa;Metazoa | EP00074 Homo sapiens 67254 | 5,80E-218 | 2379 | XP 005266191.1 | protein transport protein Sec16A isoform X1 [Homo sapiens] >gi\|767958853\|ref\|XP 011 | 2379 | 0 | 4795,72 | 100 |
| EP00079 | Obazoa;Opisthokonta;Holozoa;Choanozoa;Metazoa | EP00079 Xenopus tropicalis 33298 | 7,40E-218 | 2315 | XP 017947069.1 | PREDICTED: protein transport protein Sec16A isoform X6 [Xenopus tropicalis] | 2315 | 0 | 4697,88 | 100 |
| EP00074 | Obazoa;Opisthokonta;Holozoa;Choanozoa;Metazoa | EP00074 Homo sapiens 67243 | 8,50E-218 | 2376 | XP 011517551.1 | protein transport protein Sec16A isoform X2 [Homo sapiens] | 2376 | 0 | 4789,94 | 100 |
| EP00074 | Obazoa;Opisthokonta;Holozoa;Choanozoa;Metazoa | EP00074 Homo sapiens 67245 | 1,50E-217 | 2359 | XP 011517553.1 | protein transport protein Sec16A isoform X4 [Homo sapiens] | 2359 | 0 | 4760,67 | 100 |
| EP00070 | Obazoa;Opisthokonta;Holozoa;Choanozoa;Metazoa | EP00070 Physeter macrocephalus 46553 | 1,40E-216 | 2323 | XP 028353447.1 | protein transport protein Sec16A isoform X5 [Physeter catodon] | 2323 | 0 | 4672,46 | 100 |
| EP00062 | Obazoa;Opisthokonta;Holozoa;Choanozoa;Metazoa | EP00062 Callorhinchus milii 17240 | 1,90E-216 | 2436 | XP 007901053.1 | PREDICTED: protein transport protein Sec16A isoform X2 [Callorhinchus milii] | 2436 | 0 | 4950,19 | 100 |
| EP00067 | Obazoa;Opisthokonta;Holozoa;Choanozoa;Metazoa | EP00067 Danio rerio 26211 | 2,00E-216 | 2163 | XP 009290988.1 | protein transport protein Sec16A isoform X3 [Danio rerio] | 2163 | 0 | 4342,34 | 100 |
| EP00079 | Obazoa;Opisthokonta;Holozoa;Choanozoa;Metazoa | EP00079 Xenopus tropicalis 33297 | 2,20E-216 | 2335 | XP 017947068.1 | PREDICTED: protein transport protein Sec16A isoform X5 [Xenopus tropicalis] | 2335 | 0 | 4744,1 | 100 |
| EP00067 | Obazoa;Opisthokonta;Holozoa;Choanozoa;Metazoa | EP00067 Danio rerio 26209 | 1,10E-215 | 2183 | XP 009290986.1 | protein transport protein Sec16A isoform X1 [Danio rerio] | 2183 | 0 | 4382,79 | 100 |
| EP00062 | Obazoa;Opisthokonta;Holozoa;Choanozoa;Metazoa | EP00062 Callorhinchus milii 17237 | 1,70E-215 | 2426 | XP 007901054.1 | PREDICTED: protein transport protein Sec16A isoform X3 [Callorhinchus milii] >gi\|632969 | 2426 | 0 | 4906,66 | 100 |
| EP00062 | Obazoa;Opisthokonta;Holozoa;Choanozoa;Metazoa | EP00062 Callorhinchus milii 17238 | 1,70E-215 | 2426 | XP 007901054.1 | PREDICTED: protein transport protein Sec16A isoform X3 [Callorhinchus milii] >gi\|632969 | 2426 | 0 | 4906,66 | 100 |
| EP00072 | Obazoa;Opisthokonta;Holozoa;Choanozoa;Metazoa | EP00072 Mus musculus 22993 | 1,80E-215 | 1051 | NP 203505.3 | protein transport protein Sec16B [Mus musculus] >gi\|229606101\|ref\|NP 001153458.1\| | 1051 | 0 | 2167,5 | 100 |
| EP00072 | Obazoa;Opisthokonta;Holozoa;Choanozoa;Metazoa | EP00072 Mus musculus 4805 | 1,80E-215 | 1051 | NP 203505.3 | protein transport protein Sec16B [Mus musculus] >gi\|229606101\|ref\|NP 001153458.1\| | 1051 | 0 | 2167,5 | 100 |
| EP00072 | Obazoa;Opisthokonta;Holozoa;Choanozoa;Metazoa | EP00072 Mus musculus 66709 | 1,80E-215 | 1051 | NP 203505.3 | protein transport protein Sec16B [Mus musculus] >gi\|229606101\|ref\|NP 001153458.1\| | 1051 | 0 | 2167,5 | 100 |
| EP00072 | Obazoa;Opisthokonta;Holozoa;Choanozoa;Metazoa | EP00072 Mus musculus 66710 | 1,80E-215 | 1051 | NP 203505.3 | protein transport protein Sec16B [Mus musculus] >gi\|229606101\|ref\|NP 001153458.1\| | 1051 | 0 | 2167,5 | 100 |
| EP00070 | Obazoa;Opisthokonta;Holozoa;Choanozoa;Metazoa | EP00070 Physeter macrocephalus 46552 | 1,80E-215 | 2343 | XP 028353446.1 | protein transport protein Sec16A isoform X4 [Physeter catodon] | 2343 | 0 | 4722,92 | 100 |
| EP00067 | Obazoa;Opisthokonta;Holozoa;Choanozoa;Metazoa | EP00067 Danio rerio 26210 | 1,90E-215 | 2182 | XP 009290987.1 | protein transport protein Sec16A isoform X2 [Danio rerio] | 2182 | 0 | 4380,86 | 100 |
| EP00062 | Obazoa;Opisthokonta;Holozoa;Choanozoa;Metazoa | EP00062 Callorhinchus milii 17241 | 2,50E-215 | 2452 | XP 007901052.1 | PREDICTED: protein transport protein Sec16A isoform X1 [Callorhinchus milii] | 2452 | 0 | 4978,69 | 100 |
| EP00070 | Obazoa;Opisthokonta;Holozoa;Choanozoa;Metazoa | EP00070 Physeter macrocephalus 20863 | 3,70E-214 | 2368 | XP 023981966.1 | protein transport protein Sec16A isoform X1 [Physeter catodon] >gi\|1348484315\|ref\|XP | 2368 | 0 | 4762,98 | 100 |
| EP00070 | Obazoa;Opisthokonta;Holozoa;Choanozoa;Metazoa | EP00070 Physeter macrocephalus 20864 | 3,70E-214 | 2368 | XP 023981966.1 | protein transport protein Sec16A isoform X1 [Physeter catodon] >gi\|1348484315\|ref\|XP | 2368 | 0 | 4762,98 | 100 |
| EP00070 | Obazoa;Opisthokonta;Holozoa;Choanozoa;Metazoa | EP00070 Physeter macrocephalus 20865 | 9,70E-213 | 2367 | XP 023981968.1 | protein transport protein Sec16A isoform X2 [Physeter catodon] | 2367 | 0 | 4760,67 | 100 |
| EP00070 | Obazoa;Opisthokonta;Holozoa;Choanozoa;Metazoa | EP00070 Physeter macrocephalus 8219 | 1,40E-212 | 1065 | XP 007117743.2 | protein transport protein Sec16B [Physeter catodon] | 1065 | 0 | 2187,15 | 100 |
| EP00070 | Obazoa;Opisthokonta;Holozoa;Choanozoa;Metazoa | EP00070 Physeter macrocephalus 20866 | 2,50E-212 | 2348 | XP 023981970.1 | protein transport protein Sec16A isoform X3 [Physeter catodon] | 2348 | 0 | 4721,38 | 100 |
| EP00073 | Obazoa;Opisthokonta;Holozoa;Choanozoa;Metazoa | EP00073 Rattus norvegicus 39809 | 5,40E-212 | 949 | XP 008767908.1 | PREDICTED: protein transport protein Sec16B isoform X2 [Rattus norvegicus] | 949 | 0 | 1963,73 | 100 |
| EP00072 | Obazoa;Opisthokonta;Holozoa;Choanozoa;Metazoa | EP00072 Mus musculus 31156 | 3,40E-210 | 957 | XP 006497112.1 | protein transport protein Sec16B isoform X2 [Mus musculus] | 957 | 0 | 1977,99 | 100 |
| EP00071 | Obazoa;Opisthokonta;Holozoa;Choanozoa;Metazoa | EP00071 Canis lupus 21344 | 3,50E-210 | 1046 | XP 005622544.1 | protein transport protein Sec16B isoform X1 [Canis lupus familiaris] >gi\|928139773\|ref\|X | 1046 | 0 | 2151,71 | 100 |
| EP00071 | Obazoa;Opisthokonta;Holozoa;Choanozoa;Metazoa | EP00071 Canis lupus 21345 | 3,50E-210 | 1046 | XP 005622544.1 | protein transport protein Sec16B isoform X1 [Canis lupus familiaris] >gi\|928139773\|ref\|X | 1046 | 0 | 2151,71 | 100 |
| EP00071 | Obazoa;Opisthokonta;Holozoa;Choanozoa;Metazoa | EP00071 Canis lupus 21346 | 3,50E-210 | 1046 | XP 005622544.1 | protein transport protein Sec16B isoform X1 [Canis lupus familiaris] >gi\|928139773\|ref\|X | 1046 | 0 | 2151,71 | 100 |
| EP00071 | Obazoa;Opisthokonta;Holozoa;Choanozoa;Metazoa | EP00071 Canis lupus 21347 | 3,50E-210 | 1046 | XP 005622544.1 | protein transport protein Sec16B isoform X1 [Canis lupus familiaris] >gi\|928139773\|ref\|X | 1046 | 0 | 2151,71 | 100 |
| EP00071 | Obazoa;Opisthokonta;Holozoa;Choanozoa;Metazoa | EP00071 Canis lupus 43161 | 3,50E-210 | 1046 | XP 005622544.1 | protein transport protein Sec16B isoform X1 [Canis lupus familiaris] >gi\|928139773\|ref\|X | 1046 | 0 | 2151,71 | 100 |
| EP00071 | Obazoa;Opisthokonta;Holozoa;Choanozoa;Metazoa | EP00071 Canis lupus 6964 | 3,50E-210 | 1046 | XP 005622544.1 | protein transport protein Sec16B isoform X1 [Canis lupus familiaris] >gi\|928139773\|ref\|X | 1046 | 0 | 2151,71 | 100 |
| EP00101 | Obazoa;Opisthokonta;Holozoa;Choanozoa;Metazoa | EP00101 Apis mellifera 2595 | 2,20E-208 | 2173 | XP 006558052.1 | uncharacterized protein LOC409888 isoform X2 [Apis mellifera] | 2173 | 0 | 4481,4 | 100 |
| EP00101 | Obazoa;Opisthokonta;Holozoa;Choanozoa;Metazoa | EP00101 Apis mellifera 9755 | 6,50E-208 | 2164 | XP 016766844.1 | uncharacterized protein LOC409888 isoform X4 [Apis mellifera] | 2164 | 0 | 4464,45 | 100 |
| EP00101 | Obazoa;Opisthokonta;Holozoa;Choanozoa;Metazoa | EP00101 Apis mellifera 9756 | 1,10E-207 | 2146 | XP 016766845.1 | uncharacterized protein LOC409888 isoform X5 [Apis mellifera] | 2146 | 0 | 4425,93 | 100 |
| EP00070 | Obazoa;Opisthokonta;Holozoa;Choanozoa;Metazoa | EP00070 Physeter macrocephalus 20867 | 2,40E-207 | 2293 | XP 023981971.1 | protein transport protein Sec16A isoform X6 [Physeter catodon] | 2293 | 0 | 4599,27 | 100 |
| EP00101 | Obazoa;Opisthokonta;Holozoa;Choanozoa;Metazoa | EP00101 Apis mellifera 2594 | 4,60E-207 | 2175 | XP 006558051.1 | uncharacterized protein LOC409888 isoform X1 [Apis mellifera] | 2175 | 0 | 4485,63 | 100 |
| EP00101 | Obazoa;Opisthokonta;Holozoa;Choanozoa;Metazoa | EP00101 Apis mellifera 9759 | 2,30E-206 | 2129 | XP 016766848.1 | uncharacterized protein LOC409888 isoform X7 [Apis mellifera] >gi\|1032017843\|ref\|XP ( | 2129 | 0 | 4392,03 | 100 |
| EP00101 | Obazoa;Opisthokonta;Holozoa;Choanozoa;Metazoa | EP00101 Apis mellifera 9760 | 2,30E-206 | 2129 | XP 016766848.1 | uncharacterized protein LOC409888 isoform X7 [Apis mellifera] >gi\|1032017843\|ref\|XP ( | 2129 | 0 | 4392,03 | 100 |
| EP00064 | Obazoa;Opisthokonta;Holozoa;Choanozoa;Metazoa | EP00064 Thunnus orientalis 2733 | 2,40E-206 | 1923 | XP 019937655.1 | PREDICTED: protein transport protein Sec16A isoform X2 [Paralichthys olivaceus] | 2360 | 0 | 2974,88 | 87,3752495 |
| EP00101 | Obazoa;Opisthokonta;Holozoa;Choanozoa;Metazoa | EP00101 Apis mellifera 9757 | 3,10E-206 | 2130 | XP 016766846.1 | uncharacterized protein LOC409888 isoform X6 [Apis mellifera] >gi\|1032017839\|ref\|XP ( | 2130 | 0 | 4393,57 | 100 |
| EP00101 | Obazoa;Opisthokonta;Holozoa;Choanozoa;Metazoa | EP00101 Apis mellifera 9758 | 3,10E-206 | 2130 | XP 016766846.1 | uncharacterized protein LOC409888 isoform X6 [Apis mellifera] >gi\|1032017839\|ref\|XP ( | 2130 | 0 | 4393,57 | 100 |
| EP00071 | Obazoa;Opisthokonta;Holozoa;Choanozoa;Metazoa | EP00071 Canis lupus 21348 | 6,60E-206 | 931 | XP 013970641.1 | protein transport protein Sec16B isoform X2 [Canis lupus familiaris] | 931 | 0 | 1915,58 | 100 |
| EP00073 | Obazoa;Opisthokonta;Holozoa;Choanozoa;Metazoa | EP00073 Rattus norvegicus 15355 | 3,10E-205 | 1058 | NP 446023.1 | protein transport protein Sec16B [Rattus norvegicus] | 1058 | 0 | 2185,61 | 100 |
| EP00066 | Obazoa;Opisthokonta;Holozoa;Choanozoa;Metazoa | EP00066 Takifugu rubripes 13661 | 3,40E-204 | 2392 | XP 011602927.2 | protein transport protein Sec16A isoform X1 [Takifugu rubripes] >gi\|1698335776\|ref\|XP | 2386 | 0 | 4819,22 | 99,91638796 |
| EP00066 | Obazoa;Opisthokonta;Holozoa;Choanozoa;Metazoa | EP00066 Takifugu rubripes 13662 | 3,40E-204 | 2392 | XP 011602927.2 | protein transport protein Sec16A isoform X1 [Takifugu rubripes] >gi\|1698335776\|ref\|XP | 2386 | 0 | 4819,22 | 99,91638796 |
| EP00072 | Obazoa;Opisthokonta;Holozoa;Choanozoa;Metazoa | EP00072 Mus musculus 66712 | 1,60E-203 | 888 | XP 017168475.1 | protein transport protein Sec16B isoform X3 [Mus musculus] | 888 | 0 | 1849,33 | 100 |
| EP00066 | Obazoa;Opisthokonta;Holozoa;Choanozoa;Metazoa | EP00066 Takifugu rubripes 13664 | 4,20E-203 | 2197 | XP 011602930.2 | protein transport protein Sec16A isoform X4 [Takifugu rubripes] | 2191 | 0 | 4426,31 | 99,90896677 |
| EP00104 | Obazoa;Opisthokonta;Holozoa;Choanozoa;Metazoa | EP00104 Lingula anatina 31389 | 4,00E-201 | 1621 | XP 013415164.1 | protein transport protein Sec16A isoform X5 [Lingula anatina] | 1621 | 0 | 3325,8 | 100 |
| EP00104 | Obazoa;Opisthokonta;Holozoa;Choanozoa;Metazoa | EP00104 Lingula anatina 31390 | 5,30E-201 | 1620 | XP 013415165.1 | protein transport protein Sec16A isoform X6 [Lingula anatina] | 1620 | 0 | 3325,8 | 100 |
| EP00104 | Obazoa;Opisthokonta;Holozoa;Choanozoa;Metazoa | EP00104 Lingula anatina 31387 | 8,10E-199 | 1664 | XP 013415162.1 | protein transport protein Sec16A isoform X4 [Lingula anatina] | 1664 | 0 | 3416,71 | 100 |
| EP00104 | Obazoa;Opisthokonta;Holozoa;Choanozoa;Metazoa | EP00104 Lingula anatina 31388 | 5,60E-198 | 1664 | XP 013415163.1 | protein transport protein Sec16A isoform X3 [Lingula anatina] | 1664 | 0 | 3418,25 | 100 |
| EP00104 | Obazoa;Opisthokonta;Holozoa;Choanozoa;Metazoa | EP00104 Lingula anatina 31385 | 6,30E-198 | 1684 | XP 013415160.1 | protein transport protein Sec16A isoform X1 [Lingula anatina] | 1684 | 0 | 3458,69 | 100 |
| EP00104 | Obazoa;Opisthokonta;Holozoa;Choanozoa;Metazoa | EP00104 Lingula anatina 31386 | 1,30E-197 | 1683 | XP 013415161.1 | protein transport protein Sec16A isoform X2 [Lingula anatina] | 1683 | 0 | 3456,38 | 100 |
| EP00104 | Obazoa;Opisthokonta;Holozoa;Choanozoa;Metazoa | EP00104 Lingula anatina 5133 | 1,10E-196 | 2185 | XP 013384481.1 | uncharacterized protein LOC106154623 [Lingula anatina] | 2185 | 0 | 4504,89 | 100 |
| EP00069 | Obazoa;Opisthokonta;Holozoa;Choanozoa;Metazoa | EP00069 Balaena mysticetus 859 | 6,80E-196 | 2581 | XP 028017720.1 | protein transport protein Sec16A [Balaenoptera acutorostrata scammoni] | 2431 | 0 | 2324,67 | 95,33834586 |
| EP00066 | Obazoa;Opisthokonta;Holozoa;Choanozoa;Metazoa | EP00066 Takifugu rubripes 13663 | 3,90E-194 | 2372 | XP 011602929.2 | protein transport protein Sec16A isoform X3 [Takifugu rubripes] | 2366 | 0 | 4778,77 | 99,91568297 |
| EP00063 | Obazoa;Opisthokonta;Holozoa;Choanozoa;Metazoa | EP00063 Salmo salar 4395 | 2,10E-192 | 2362 | XP 013979230.1 | PREDICTED: protein transport protein Sec16A [Salmo salar] | 2362 | 0 | 4762,21 | 100 |
| EP00065 | Obazoa;Opisthokonta;Holozoa;Choanozoa;Metazoa | EP00065 Mola mola 5948 | 1,20E-191 | 2038 | XP 006796819.1 | PREDICTED: protein transport protein Sec16A-like isoform X3 [Neolamprologus brichardi] | 2312 | 0 | 2389,76 | 76,98838248 |
| EP00058 | Obazoa;Opisthokonta;Holozoa;Choanozoa;Metazoa | EP00058 Branchiostoma floridae 14780 | 2,80E-189 | 2645 | XP 002600268.1 | hypothetical protein BRAFLDRAFT 66773 [Branchiostoma floridae] | 2645 | 0 | 5398,17 | 100 |
| EP00069 | Obazoa;Opisthokonta;Holozoa;Choanozoa;Metazoa | EP00069 Balaena mysticetus 14077 | 7,30E-187 | 1231 | XP 007117743.2 | protein transport protein Sec16B [Physeter catodon] | 1065 | 0 | 1492,63 | 90 |
| EP00070 | Obazoa;Opisthokonta;Holozoa;Choanozoa;Metazoa | EP00070 Physeter macrocephalus 46554 | 6,40E-186 | 2111 | XP 028353448.1 | protein transport protein Sec16A isoform X7 [Physeter catodon] | 2111 | 0 | 4259,14 | 100 |
| EP00076 | Obazoa;Opisthokonta;Holozoa;Choanozoa;Metazoa | EP00076 Gallus gallus 23846 | 1,20E-185 | 979 | XP 015145778.1 | protein transport protein Sec16B isoform X3 [Gallus gallus] | 979 | 0 | 1998,02 | 100 |
| EP00076 | Obazoa;Opisthokonta;Holozoa;Choanozoa;Metazoa | EP00076 Gallus gallus 43046 | 1,70E-185 | 983 | XP 025008584.1 | protein transport protein Sec16B isoform X2 [Gallus gallus] | 983 | 0 | 2011,88 | 100 |
| EP00075 | Obazoa;Opisthokonta;Holozoa;Choanozoa;Metazoa | EP00075 Anas platyrhynchos 29380 | 2,10E-185 | 956 | XP 027318633.1 | protein transport protein Sec16B isoform X5 [Anas platyrhynchos] >gi\|1540448169\|ref\|X | 956 | 0 | 1944,47 | 100 |
| EP00075 | Obazoa;Opisthokonta;Holozoa;Choanozoa;Metazoa | EP00075 Anas platyrhynchos 29382 | 2,10E-185 | 956 | XP 027318633.1 | protein transport protein Sec16B isoform X5 [Anas platyrhynchos] >gi\|1540448169\|ref\|X | 956 | 0 | 1944,47 | 100 |
| EP00076 | Obazoa;Opisthokonta;Holozoa;Choanozoa;Metazoa | EP00076 Gallus gallus 43047 | 2,20E-185 | 966 | XP 025008585.1 | protein transport protein Sec16B isoform X4 [Gallus gallus] | 966 | 0 | 1977,6 | 100 |
| EP00074 | Obazoa;Opisthokonta;Holozoa;Choanozoa;Metazoa | EP00074 Homo sapiens 67258 | 2,30E-185 | 1953 | XP 011517566.1 | protein transport protein Sec16A isoform X17 [Homo sapiens] | 1953 | 0 | 3962,15 | 100 |
| EP00076 | Obazoa;Opisthokonta;Holozoa;Choanozoa;Metazoa | EP00076 Gallus gallus 43045 | 2,40E-185 | 984 | XP 025008583.1 | protein transport protein Sec16B isoform X1 [Gallus gallus] | 984 | 0 | 2013,81 | 100 |
| EP00076 | Obazoa;Opisthokonta;Holozoa;Choanozoa;Metazoa | EP00076 Gallus gallus 43048 | 2,80E-185 | 928 | XP 025008586.1 | protein transport protein Sec16B isoform X5 [Gallus gallus] >gi\|1390121326\|ref\|XP 025( | 928 | 0 | 1896,32 | 100 |
| EP00076 | Obazoa;Opisthokonta;Holozoa;Choanozoa;Metazoa | EP00076 Gallus gallus 43049 | 2,80E-185 | 928 | XP 025008586.1 | protein transport protein Sec16B isoform X5 [Gallus gallus] >gi\|1390121326\|ref\|XP 025( | 928 | 0 | 1896,32 | 100 |
| EP00076 | Obazoa;Opisthokonta;Holozoa;Choanozoa;Metazoa | EP00076 Gallus gallus 1834 | 3,70E-185 | 929 | NP 001026440.1 | protein transport protein Sec16B [Gallus gallus] | 929 | 0 | 1897,86 | 100 |
| EP00075 | Obazoa;Opisthokonta;Holozoa;Choanozoa;Metazoa | EP00075 Anas platyrhynchos 29379 | 5,40E-185 | 964 | XP 027318632.1 | protein transport protein Sec16B isoform X4 [Anas platyrhynchos] | 964 | 0 | 1959,88 | 100 |
| EP00077 | Obazoa;Opisthokonta;Holozoa;Choanozoa;Metazoa | EP00077 Pelodiscus sinensis 34128 | 7,20E-185 | 827 | XP 025042393.1 | protein transport protein Sec16B [Pelodiscus sinensis] | 827 | 0 | 1713,35 | 100 |
| EP00075 | Obazoa;Opisthokonta;Holozoa;Choanozoa;Metazoa | EP00075 Anas platyrhynchos 29376 | 7,80E-185 | 965 | XP 027318629.1 | protein transport protein Sec16B isoform X1 [Anas platyrhynchos] >gi\|1540448161\|ref\|X | 965 | 0 | 1961,42 | 100 |
| EP00075 | Obazoa;Opisthokonta;Holozoa;Choanozoa;Metazoa | EP00075 Anas platyrhynchos 29377 | 7,80E-185 | 965 | XP 027318629.1 | protein transport protein Sec16B isoform X1 [Anas platyrhynchos] >gi\|1540448161\|ref\|X | 965 | 0 | 1961,42 | 100 |
| EP00075 | Obazoa;Opisthokonta;Holozoa;Choanozoa;Metazoa | EP00075 Anas platyrhynchos 29378 | 7,80E-185 | 965 | XP 027318629.1 | protein transport protein Sec16B isoform X1 [Anas platyrhynchos] >gi\|1540448161\|ref\|X | 965 | 0 | 1961,42 | 100 |
| EP00079 | Obazoa;Opisthokonta;Holozoa;Choanozoa;Metazoa | EP00079 Xenopus tropicalis 15913 | 6,60E-184 | 978 | XP 004914083.1 | PREDICTED: protein transport protein Sec16B isoform X1 [Xenopus tropicalis] | 978 | 0 | 2048,86 | 100 |
| EP00079 | Obazoa;Opisthokonta;Holozoa;Choanozoa;Metazoa | EP00079 Xenopus tropicalis 24435 | 1,60E-183 | 963 | XP 012817075.1 | PREDICTED: protein transport protein Sec16B isoform X2 [Xenopus tropicalis] | 963 | 0 | 2019,97 | 100 |
| EP00079 | Obazoa;Opisthokonta;Holozoa;Choanozoa;Metazoa | EP00079 Xenopus tropicalis 24436 | 3,60E-183 | 950 | XP 012817076.1 | PREDICTED: protein transport protein Sec16B isoform X3 [Xenopus tropicalis] | 950 | 0 | 1990,31 | 100 |
| EP00076 | Obazoa;Opisthokonta;Holozoa;Choanozoa;Metazoa | EP00076 Gallus gallus 23848 | 3,00E-182 | 855 | XP 015145784.1 | protein transport protein Sec16B isoform X6 [Gallus gallus] | 855 | 0 | 1751,87 | 100 |
| EP00083 | Obazoa;Opisthokonta;Holozoa;Choanozoa;Metazoa | EP00083 Stegodyphus mimosarum 3631 | 5,20E-182 | 1238 | XP 015903914.2 | COPII coat assembly protein sec16 [Parasteatoda tepidariorum] | 2031 | 0 | 1429,46 | 71,59863946 |
| EP00106 | Obazoa;Opisthokonta;Holozoa;Choanozoa;Metazoa | EP00106 Octopus bimaculoides 1675 | 2,80E-180 | 2118 | XP 014769117.1 | PREDICTED: uncharacterized protein LOC106868398 isoform X7 [Octopus bimaculoides] | 2118 | 0 | 4403,59 | 100 |
| EP00074 | Obazoa;Opisthokonta;Holozoa;Choanozoa;Metazoa | EP00074 Homo sapiens 67252 | 3,00E-180 | 2332 | XP 011517560.1 | protein transport protein Sec16A isoform X12 [Homo sapiens] | 2332 | 0 | 4695,57 | 100 |
| EP00075 | Obazoa;Opisthokonta;Holozoa;Choanozoa;Metazoa | EP00075 Anas platyrhynchos 29383 | 8,70E-180 | 817 | XP 027318636.1 | protein transport protein Sec16B isoform X7 [Anas platyrhynchos] | 817 | 0 | 1662,12 | 100 |
| EP00106 | Obazoa;Opisthokonta;Holozoa;Choanozoa;Metazoa | EP00106 Octopus bimaculoides 1674 | 1,70E-179 | 2140 | XP 014769116.1 | PREDICTED: uncharacterized protein LOC106868398 isoform X6 [Octopus bimaculoides] | 2140 | 0 | 4445,57 | 100 |

| | | | | | | | | | | |
|---|---|---|---|---|---|---|---|---|---|---|
| EP00106 | Obazoa;Opisthokonta;Holozoa;Choanozoa;Metazoa | EP00106 Octopus bimaculoides 1671 | 2,40E-179 | 2159 | XP 014769113.1 | PREDICTED: uncharacterized protein LOC106868398 isoform X4 [Octopus bimaculoides] | 2159 | 0 | 4480,63 | 100 |
| EP00076 | Obazoa;Opisthokonta;Holozoa;Choanozoa;Metazoa | EP00076 Gallus gallus 43050 | 6,10E-179 | 816 | XP 025008588.1 | protein transport protein Sec16B isoform X7 [Gallus gallus] | 816 | 0 | 1677,92 | 100 |
| EP00106 | Obazoa;Opisthokonta;Holozoa;Choanozoa;Metazoa | EP00106 Octopus bimaculoides 1668 | 5,40E-178 | 2181 | XP 014769110.1 | PREDICTED: uncharacterized protein LOC106868398 isoform X1 [Octopus bimaculoides] | 2181 | 0 | 4528,01 | 100 |
| EP00106 | Obazoa;Opisthokonta;Holozoa;Choanozoa;Metazoa | EP00106 Octopus bimaculoides 1673 | 1,20E-177 | 2142 | XP 014769115.1 | PREDICTED: uncharacterized protein LOC106868398 isoform X5 [Octopus bimaculoides] | 2142 | 0 | 4445,96 | 100 |
| EP00106 | Obazoa;Opisthokonta;Holozoa;Choanozoa;Metazoa | EP00106 Octopus bimaculoides 1670 | 2,10E-177 | 2174 | XP 014769112.1 | PREDICTED: uncharacterized protein LOC106868398 isoform X3 [Octopus bimaculoides] | 2174 | 0 | 4512,21 | 100 |
| EP00114 | Obazoa;Opisthokonta;Holozoa;Choanozoa;Metazoa | EP00114 Trichoplax sp H2 9654 | 3,50E-177 | 1692 | XP 002109984.1 | hypothetical protein TRIADDRAFT 53450 [Trichoplax adhaerens] | 1642 | 0 | 3341,59 | 99,52718676 |
| EP00079 | Obazoa;Opisthokonta;Holozoa;Choanozoa;Metazoa | EP00079 Xenopus tropicalis 11504 | 1,10E-176 | 849 | XP 002937408.2 | PREDICTED: protein transport protein Sec16B isoform X4 [Xenopus tropicalis] | 849 | 0 | 1774,6 | 100 |
| EP00113 | Obazoa;Opisthokonta;Holozoa;Choanozoa;Metazoa | EP00113 Trichoplax adhaerens 2397 | 5,30E-176 | 1642 | XP 002109984.1 | hypothetical protein TRIADDRAFT 53450 [Trichoplax adhaerens] | 1642 | 0 | 3384,73 | 100 |
| EP00107 | Obazoa;Opisthokonta;Holozoa;Choanozoa;Metazoa | EP00107 Lottia gigantea 6265 | 5,70E-174 | 1958 | XP 009050357.1 | hypothetical protein LOTGIDRAFT 231137 [Lottia gigantea] | 1958 | 0 | 4061,92 | 100 |
| EP00098 | Obazoa;Opisthokonta;Holozoa;Choanozoa;Metazoa | EP00098 Tribolium castaneum 16143 | 8,70E-174 | 1357 | XP 015839570.1 | PREDICTED: protein transport protein Sec16A isoform X3 [Tribolium castaneum] >gi\|1008 | 1357 | 0 | 2827,35 | 100 |
| EP00098 | Obazoa;Opisthokonta;Holozoa;Choanozoa;Metazoa | EP00098 Tribolium castaneum 16144 | 8,70E-174 | 1357 | XP 015839570.1 | PREDICTED: protein transport protein Sec16A isoform X3 [Tribolium castaneum] >gi\|1008 | 1357 | 0 | 2827,35 | 100 |
| EP00098 | Obazoa;Opisthokonta;Holozoa;Choanozoa;Metazoa | EP00098 Tribolium castaneum 16145 | 1,10E-173 | 1354 | XP 015839568.1 | PREDICTED: protein transport protein Sec16A isoform X1 [Tribolium castaneum] >gi\|1008 | 1376 | 0 | 2821,96 | 99,85228951 |
| EP00078 | Obazoa;Opisthokonta;Holozoa;Choanozoa;Metazoa | EP00078 Anolis carolinensis 31665 | 1,10E-173 | 990 | XP 016851775.1 | PREDICTED: protein transport protein Sec16B [Anolis carolinensis] | 990 | 0 | 2055,41 | 100 |
| EP00098 | Obazoa;Opisthokonta;Holozoa;Choanozoa;Metazoa | EP00098 Tribolium castaneum 8131 | 1,20E-173 | 1366 | XP 008199877.1 | PREDICTED: protein transport protein Sec16B isoform X2 [Tribolium castaneum] | 1366 | 0 | 2850,08 | 100 |
| EP00098 | Obazoa;Opisthokonta;Holozoa;Choanozoa;Metazoa | EP00098 Tribolium castaneum 16141 | 1,50E-173 | 1376 | XP 015839568.1 | PREDICTED: protein transport protein Sec16A isoform X1 [Tribolium castaneum] >gi\|1008 | 1376 | 0 | 2869,72 | 100 |
| EP00098 | Obazoa;Opisthokonta;Holozoa;Choanozoa;Metazoa | EP00098 Tribolium castaneum 16142 | 1,50E-173 | 1376 | XP 015839568.1 | PREDICTED: protein transport protein Sec16A isoform X1 [Tribolium castaneum] >gi\|1008 | 1376 | 0 | 2869,72 | 100 |
| EP00098 | Obazoa;Opisthokonta;Holozoa;Choanozoa;Metazoa | EP00098 Tribolium castaneum 16146 | 5,40E-172 | 1339 | XP 015839573.1 | PREDICTED: protein transport protein Sec16A isoform X5 [Tribolium castaneum] | 1339 | 0 | 2791,53 | 100 |
| EP00101 | Obazoa;Opisthokonta;Holozoa;Choanozoa;Metazoa | EP00101 Apis mellifera 14015 | 6,20E-172 | 2167 | XP 026295583.1 | uncharacterized protein LOC409888 isoform X3 [Apis mellifera] | 2167 | 0 | 4474,08 | 100 |
| EP00068 | Obazoa;Opisthokonta;Holozoa;Choanozoa;Metazoa | EP00068 Latimeria chalumnae 22439 | 2,70E-171 | 878 | XP 014342756.1 | PREDICTED: protein transport protein Sec16B [Latimeria chalumnae] | 878 | 0 | 1820,82 | 100 |
| EP00247 | Archaeplastida;Chloroplastida;Streptophyta;"core s | EP00247 Physcomitrella patens 17088 | 3,20E-171 | 1264 | XP 024373339.1 | protein transport protein SEC16B homolog isoform X2 [Physcomitrella patens] >gi\|13738! | 1264 | 0 | 2595,85 | 100 |
| EP00247 | Archaeplastida;Chloroplastida;Streptophyta;"core s | EP00247 Physcomitrella patens 17095 | 3,20E-171 | 1264 | XP 024373339.1 | protein transport protein SEC16B homolog isoform X2 [Physcomitrella patens] >gi\|13738! | 1264 | 0 | 2595,85 | 100 |
| EP00247 | Archaeplastida;Chloroplastida;Streptophyta;"core s | EP00247 Physcomitrella patens 17087 | 8,70E-171 | 1444 | XP 024373338.1 | protein transport protein SEC16A homolog isoform X1 [Physcomitrella patens] >gi\|13738! | 1444 | 0 | 2956,01 | 100 |
| EP00247 | Archaeplastida;Chloroplastida;Streptophyta;"core s | EP00247 Physcomitrella patens 17093 | 8,70E-171 | 1444 | XP 024373338.1 | protein transport protein SEC16A homolog isoform X1 [Physcomitrella patens] >gi\|13738! | 1444 | 0 | 2956,01 | 100 |
| EP00060 | Obazoa;Opisthokonta;Holozoa;Choanozoa;Metazoa | EP00060 Petromyzon marinus 38035 | 9,60E-171 | 1326 | XP 015734342.1 | PREDICTED: protein transport protein Sec16A isoform X7 [Coturnix japonica] | 2200 | 0 | 777,704 | 52,59048971 |
| EP00067 | Obazoa;Opisthokonta;Holozoa;Choanozoa;Metazoa | EP00067 Danio rerio 29924 | 1,00E-170 | 935 | XP 685029.5 | protein transport protein Sec16B isoform X1 [Danio rerio] >gi\|688560164\|ref\|XP 00930C | 935 | 0 | 1949,48 | 100 |
| EP00067 | Obazoa;Opisthokonta;Holozoa;Choanozoa;Metazoa | EP00067 Danio rerio 51441 | 1,00E-170 | 935 | XP 685029.5 | protein transport protein Sec16B isoform X1 [Danio rerio] >gi\|688560164\|ref\|XP 00930C | 935 | 0 | 1949,48 | 100 |
| EP00067 | Obazoa;Opisthokonta;Holozoa;Choanozoa;Metazoa | EP00067 Danio rerio 41534 | 2,90E-170 | 935 | XP 021327437.1 | protein transport protein Sec16B isoform X1 [Danio rerio] | 935 | 0 | 1949,48 | 100 |
| EP00260 | Archaeplastida;Chloroplastida;Streptophyta;"core s | EP00260 Arabidopsis thaliana 37518 | 2,10E-169 | 1361 | NP 199560.1 | RGPR-like protein [Arabidopsis thaliana] >gi\|1063736704\|ref\|NP 001318753.1\| RGPR-lik | 1361 | 0 | 2823,88 | 100 |
| EP00260 | Archaeplastida;Chloroplastida;Streptophyta;"core s | EP00260 Arabidopsis thaliana 6911 | 2,10E-169 | 1361 | NP 199560.1 | RGPR-like protein [Arabidopsis thaliana] >gi\|1063736704\|ref\|NP 001318753.1\| RGPR-lik | 1361 | 0 | 2823,88 | 100 |
| EP00260 | Archaeplastida;Chloroplastida;Streptophyta;"core s | EP00260 Arabidopsis thaliana 37517 | 4,10E-169 | 1350 | NP 199559.1 | RGPR-like protein [Arabidopsis thaliana] | 1350 | 0 | 2787,29 | 100 |
| EP00106 | Obazoa;Opisthokonta;Holozoa;Choanozoa;Metazoa | EP00106 Octopus bimaculoides 1669 | 7,50E-169 | 2174 | XP 014769111.1 | PREDICTED: uncharacterized protein LOC106868398 isoform X2 [Octopus bimaculoides] | 2174 | 0 | 4511,06 | 100 |
| EP00247 | Archaeplastida;Chloroplastida;Streptophyta;"core s | EP00247 Physcomitrella patens 35259 | 1,60E-168 | 1438 | XP 024391510.1 | protein transport protein SEC16A homolog isoform X2 [Physcomitrella patens] >gi\|13739: | 1438 | 0 | 2942,14 | 100 |
| EP00247 | Archaeplastida;Chloroplastida;Streptophyta;"core s | EP00247 Physcomitrella patens 35260 | 1,60E-168 | 1438 | XP 024391510.1 | protein transport protein SEC16A homolog isoform X2 [Physcomitrella patens] >gi\|13739: | 1438 | 0 | 2942,14 | 100 |
| EP00247 | Archaeplastida;Chloroplastida;Streptophyta;"core s | EP00247 Physcomitrella patens 35261 | 1,60E-168 | 1438 | XP 024391510.1 | protein transport protein SEC16A homolog isoform X2 [Physcomitrella patens] >gi\|13739: | 1438 | 0 | 2942,14 | 100 |
| EP00062 | Obazoa;Opisthokonta;Holozoa;Choanozoa;Metazoa | EP00062 Callorhinchus milii 17296 | 1,60E-168 | 1012 | XP 007901208.1 | PREDICTED: protein transport protein Sec16B [Callorhinchus milii] | 1012 | 0 | 2095,47 | 100 |
| EP00247 | Archaeplastida;Chloroplastida;Streptophyta;"core s | EP00247 Physcomitrella patens 35252 | 2,00E-168 | 1457 | XP 024391503.1 | protein transport protein SEC16A homolog isoform X1 [Physcomitrella patens] >gi\|13739: | 1457 | 0 | 2980,66 | 100 |
| EP00247 | Archaeplastida;Chloroplastida;Streptophyta;"core s | EP00247 Physcomitrella patens 35254 | 2,00E-168 | 1457 | XP 024391503.1 | protein transport protein SEC16A homolog isoform X1 [Physcomitrella patens] >gi\|13739: | 1457 | 0 | 2980,66 | 100 |
| EP00247 | Archaeplastida;Chloroplastida;Streptophyta;"core s | EP00247 Physcomitrella patens 35255 | 2,00E-168 | 1457 | XP 024391503.1 | protein transport protein SEC16A homolog isoform X1 [Physcomitrella patens] >gi\|13739: | 1457 | 0 | 2980,66 | 100 |
| EP00247 | Archaeplastida;Chloroplastida;Streptophyta;"core s | EP00247 Physcomitrella patens 35256 | 2,00E-168 | 1457 | XP 024391503.1 | protein transport protein SEC16A homolog isoform X1 [Physcomitrella patens] >gi\|13739: | 1457 | 0 | 2980,66 | 100 |
| EP00247 | Archaeplastida;Chloroplastida;Streptophyta;"core s | EP00247 Physcomitrella patens 35257 | 2,00E-168 | 1457 | XP 024391503.1 | protein transport protein SEC16A homolog isoform X1 [Physcomitrella patens] >gi\|13739: | 1457 | 0 | 2980,66 | 100 |
| EP00247 | Archaeplastida;Chloroplastida;Streptophyta;"core s | EP00247 Physcomitrella patens 35258 | 2,00E-168 | 1457 | XP 024391503.1 | protein transport protein SEC16A homolog isoform X1 [Physcomitrella patens] >gi\|13739: | 1457 | 0 | 2980,66 | 100 |
| EP00063 | Obazoa;Opisthokonta;Holozoa;Choanozoa;Metazoa | EP00063 Salmo salar 3776 | 1,60E-167 | 993 | XP 013978611.1 | PREDICTED: protein transport protein Sec16B [Salmo salar] | 993 | 0 | 2055,03 | 100 |
| EP00259 | Archaeplastida;Chloroplastida;Streptophyta;"core s | EP00259 Arabidopsis lyrata 1101 | 9,40E-167 | 1358 | XP 002865115.1 | protein transport protein SEC16A homolog isoform X1 [Arabidopsis lyrata subsp. lyrata] | 1358 | 0 | 2810,02 | 100 |
| EP00074 | Obazoa;Opisthokonta;Holozoa;Choanozoa;Metazoa | EP00074 Homo sapiens 31139 | 1,60E-166 | 719 | NP 001343429.1 | protein transport protein Sec16B isoform 3 [Homo sapiens] | 719 | 0 | 1471,83 | 100 |
| EP00105 | Obazoa;Opisthokonta;Holozoa;Choanozoa;Metazoa | EP00105 Crassostrea gigas 42212 | 2,20E-166 | 993 | XP 019926360.1 | PREDICTED: protein transport protein Sec16A isoform X2 [Crassostrea gigas] | 993 | 0 | 2080,45 | 100 |
| EP00259 | Archaeplastida;Chloroplastida;Streptophyta;"core s | EP00259 Arabidopsis lyrata 14131 | 2,20E-166 | 1356 | XP 020866847.1 | protein transport protein SEC16A homolog isoform X2 [Arabidopsis lyrata subsp. lyrata] | 1356 | 0 | 2805,39 | 100 |
| EP00067 | Obazoa;Opisthokonta;Holozoa;Choanozoa;Metazoa | EP00067 Danio rerio 41535 | 2,50E-166 | 827 | XP 021327438.1 | protein transport protein Sec16B isoform X2 [Danio rerio] | 827 | 0 | 1722,98 | 100 |
| EP00023 | Amoebozoa;Evosea;Eumycetozoa;Dictyostelia;Dicty | EP00023 Dictyostelium discoideum 12290 | 2,80E-166 | 1992 | XP 646771.1 | hypothetical protein DDB G0269198 [Dictyostelium discoideum AX4] | 1992 | 0 | 3915,92 | 100 |
| EP00105 | Obazoa;Opisthokonta;Holozoa;Choanozoa;Metazoa | EP00105 Crassostrea gigas 42214 | 1,10E-165 | 945 | XP 019926362.1 | PREDICTED: protein transport protein Sec16A isoform X4 [Crassostrea gigas] | 945 | 0 | 1983,38 | 100 |
| EP00105 | Obazoa;Opisthokonta;Holozoa;Choanozoa;Metazoa | EP00105 Crassostrea gigas 20604 | 1,40E-165 | 995 | XP 011439430.2 | PREDICTED: protein transport protein Sec16A isoform X1 [Crassostrea gigas] | 995 | 0 | 2084,69 | 100 |
| EP00105 | Obazoa;Opisthokonta;Holozoa;Choanozoa;Metazoa | EP00105 Crassostrea gigas 42213 | 4,50E-165 | 975 | XP 019926361.1 | PREDICTED: protein transport protein Sec16A isoform X3 [Crassostrea gigas] | 975 | 0 | 2043,85 | 100 |
| EP00259 | Archaeplastida;Chloroplastida;Streptophyta;"core s | EP00259 Arabidopsis lyrata 1100 | 1,80E-163 | 1358 | XP 002865114.1 | protein transport protein SEC16B homolog isoform X1 [Arabidopsis lyrata subsp. lyrata] >ɛ | 1358 | 0 | 2822,73 | 100 |
| EP00259 | Archaeplastida;Chloroplastida;Streptophyta;"core s | EP00259 Arabidopsis lyrata 14150 | 1,80E-163 | 1358 | XP 002865114.1 | protein transport protein SEC16B homolog isoform X1 [Arabidopsis lyrata subsp. lyrata] >ɛ | 1358 | 0 | 2822,73 | 100 |
| EP00259 | Archaeplastida;Chloroplastida;Streptophyta;"core s | EP00259 Arabidopsis lyrata 14157 | 2,90E-163 | 1357 | XP 020866873.1 | protein transport protein SEC16B homolog isoform X2 [Arabidopsis lyrata subsp. lyrata] | 1357 | 0 | 2819,65 | 100 |
| EP00084 | Obazoa;Opisthokonta;Holozoa;Choanozoa;Metazoa | EP00084 Strigamia maritima 6860 | 7,80E-163 | 1797 | XP 022244251.1 | uncharacterized protein LOC106461550 isoform X3 [Limulus polyphemus] | 2244 | 0 | 630,943 | 52,82685512 |
| EP00131 | Obazoa;Opisthokonta;Nucletmycea;Fungi;"core Fur | EP00131 Spizellomyces punctatus 2826 | 1,10E-162 | 2211 | XP 016606650.1 | hypothetical protein SPPG 06291 [Spizellomyces punctatus DAOM BR117] | 2211 | 0 | 4528,01 | 100 |
| EP00064 | Obazoa;Opisthokonta;Holozoa;Choanozoa;Metazoa | EP00064 Thunnus orientalis 3613 | 2,00E-162 | 948 | XP 028443522.1 | protein transport protein Sec16B-like [Perca flavescens] | 941 | 0 | 1422,53 | 79,41810345 |
| EP00268 | Archaeplastida;Chloroplastida;Streptophyta;"core s | EP00268 Persea americana 9747 | 4,00E-162 | 1429 | XP 010258928.1 | PREDICTED: protein transport protein SEC16B homolog [Nelumbo nucifera] >gi\|72000941 | 1429 | 0 | 1569,67 | 64,93860846 |
| EP00268 | Archaeplastida;Chloroplastida;Streptophyta;"core s | EP00268 Persea americana 9749 | 4,00E-162 | 1429 | XP 010258928.1 | PREDICTED: protein transport protein SEC16B homolog [Nelumbo nucifera] >gi\|72000941 | 1429 | 0 | 1569,67 | 64,93860846 |
| EP00086 | Obazoa;Opisthokonta;Holozoa;Choanozoa;Metazoa | EP00086 Labidocera madurae 62016 | 6,40E-162 | 2035 | XP 023346730.1 | uncharacterized protein LOC111715609 isoform X1 [Eurytemora affinis] | 1957 | 0 | 1973,75 | 67,40775781 |
| EP00093 | Obazoa;Opisthokonta;Holozoa;Choanozoa;Metazoa | EP00093 Oithona nana 14117 | 2,20E-160 | 1029 | XP 014669562.1 | PREDICTED: protein transport protein Sec16A-like isoform X6 [Priapulus caudatus] | 1146 | 1,69E-131 | 436,417 | 47,46450304 |
| EP00094 | Obazoa;Opisthokonta;Holozoa;Choanozoa;Metazoa | EP00094 Tigriopus californicus 560 | 2,30E-160 | 1780 | XP 023346731.1 | protein transport protein Sec16-like isoform X2 [Eurytemora affinis] | 1941 | 1,20E-159 | 546,969 | 51,64179104 |
| EP00261 | Archaeplastida;Chloroplastida;Streptophyta;"core s | EP00261 Vitis vinifera 27501 | 2,60E-160 | 1400 | XP 010661318.1 | PREDICTED: protein transport protein SEC16B homolog [Vitis vinifera] >gi\|1104657609\|re | 1400 | 0 | 2900,15 | 100 |
| EP00261 | Archaeplastida;Chloroplastida;Streptophyta;"core s | EP00261 Vitis vinifera 40002 | 2,60E-160 | 1400 | XP 010661318.1 | PREDICTED: protein transport protein SEC16B homolog [Vitis vinifera] >gi\|1104657609\|re | 1400 | 0 | 2900,15 | 100 |
| EP00268 | Archaeplastida;Chloroplastida;Streptophyta;"core s | EP00268 Persea americana 9748 | 1,30E-159 | 1432 | XP 010258928.1 | PREDICTED: protein transport protein SEC16B homolog [Nelumbo nucifera] >gi\|72000941 | 1429 | 0 | 1543,87 | 64,99321574 |
| EP00268 | Archaeplastida;Chloroplastida;Streptophyta;"core s | EP00268 Persea americana 9750 | 1,30E-159 | 1432 | XP 010258928.1 | PREDICTED: protein transport protein SEC16B homolog [Nelumbo nucifera] >gi\|72000941 | 1429 | 0 | 1543,87 | 64,99321574 |
| EP00086 | Obazoa;Opisthokonta;Holozoa;Choanozoa;Metazoa | EP00086 Labidocera madurae 62015 | 1,60E-159 | 2019 | XP 023346731.1 | protein transport protein Sec16-like isoform X2 [Eurytemora affinis] | 1941 | 0 | 1956,41 | 67,49285033 |
| EP00087 | Obazoa;Opisthokonta;Holozoa;Choanozoa;Metazoa | EP00087 Eurytemora affinis 27274 | 2,70E-158 | 1957 | XP 023346730.1 | uncharacterized protein LOC111715609 isoform X1 [Eurytemora affinis] | 1957 | 0 | 4034,18 | 100 |
| EP00087 | Obazoa;Opisthokonta;Holozoa;Choanozoa;Metazoa | EP00087 Eurytemora affinis 27275 | 3,80E-158 | 1941 | XP 023346731.1 | protein transport protein Sec16-like isoform X2 [Eurytemora affinis] | 1941 | 0 | 4001,05 | 100 |
| EP00074 | Obazoa;Opisthokonta;Holozoa;Choanozoa;Metazoa | EP00074 Homo sapiens 67255 | 5,00E-158 | 2230 | XP 011517563.1 | protein transport protein Sec16A isoform X14 [Homo sapiens] | 2230 | 0 | 4484,48 | 100 |
| EP00258 | Archaeplastida;Chloroplastida;Streptophyta;"core s | EP00258 Populus trichocarpa 36109 | 5,20E-157 | 1405 | XP 024452128.1 | protein transport protein SEC16B homolog [Populus trichocarpa] >gi\|1375846436\|ref\|XP | 1405 | 0 | 2908,24 | 100 |
| EP00258 | Archaeplastida;Chloroplastida;Streptophyta;"core s | EP00258 Populus trichocarpa 36110 | 5,20E-157 | 1405 | XP 024452128.1 | protein transport protein SEC16B homolog [Populus trichocarpa] >gi\|1375846436\|ref\|XP | 1405 | 0 | 2908,24 | 100 |
| EP00105 | Obazoa;Opisthokonta;Holozoa;Choanozoa;Metazoa | EP00105 Crassostrea gigas 13368 | 2,50E-156 | 1436 | XP 011429675.1 | PREDICTED: uncharacterized protein LOC105329889 [Crassostrea gigas] | 1436 | 0 | 2995,68 | 100 |
| EP00258 | Archaeplastida;Chloroplastida;Streptophyta;"core s | EP00258 Populus trichocarpa 33743 | 4,50E-156 | 1412 | XP 024449762.1 | protein transport protein SEC16B homolog isoform X2 [Populus trichocarpa] | 1412 | 0 | 2918,64 | 100 |
| EP00258 | Archaeplastida;Chloroplastida;Streptophyta;"core s | EP00258 Populus trichocarpa 33742 | 5,80E-156 | 1414 | XP 024449761.1 | protein transport protein SEC16B homolog isoform X1 [Populus trichocarpa] | 1414 | 0 | 2922,88 | 100 |
| EP00095 | Obazoa;Opisthokonta;Holozoa;Choanozoa;Metazoa | EP00095 Tigriopus kingsejongensis 182 | 2,90E-155 | 1731 | XP 023346730.1 | uncharacterized protein LOC111715609 isoform X1 [Eurytemora affinis] | 1957 | 2,38E-159 | 545,428 | 50,57251908 |
| EP00095 | Obazoa;Opisthokonta;Holozoa;Choanozoa;Metazoa | EP00095 Tigriopus kingsejongensis 4258 | 2,90E-155 | 1731 | XP 023346730.1 | uncharacterized protein LOC111715609 isoform X1 [Eurytemora affinis] | 1957 | 2,38E-159 | 545,428 | 50,57251908 |
| EP00261 | Archaeplastida;Chloroplastida;Streptophyta;"core s | EP00261 Vitis vinifera 17045 | 4,20E-155 | 1423 | XP 010646525.1 | PREDICTED: protein transport protein SEC16A homolog [Vitis vinifera] | 1423 | 0 | 2949,84 | 100 |
| EP00256 | Archaeplastida;Chloroplastida;Streptophyta;"core s | EP00256 Glycine max 7772 | 3,90E-154 | 1404 | XP 003516666.1 | protein transport protein SEC16B homolog [Glycine max] | 1404 | 0 | 2905,16 | 100 |
| EP00088 | Obazoa;Opisthokonta;Holozoa;Choanozoa;Metazoa | EP00088 Acartia fossae 38520 | 1,10E-153 | 888 | XP 023346731.1 | protein transport protein Sec16-like isoform X2 [Eurytemora affinis] | 1941 | 0 | 587,03 | 72,55813953 |
| EP00088 | Obazoa;Opisthokonta;Holozoa;Choanozoa;Metazoa | EP00088 Acartia fossae 38521 | 1,10E-153 | 888 | XP 023346731.1 | protein transport protein Sec16-like isoform X2 [Eurytemora affinis] | 1941 | 0 | 587,03 | 72,55813953 |
| EP00088 | Obazoa;Opisthokonta;Holozoa;Choanozoa;Metazoa | EP00088 Acartia fossae 38522 | 1,10E-153 | 888 | XP 023346731.1 | protein transport protein Sec16-like isoform X2 [Eurytemora affinis] | 1941 | 0 | 587,03 | 72,55813953 |
| EP00088 | Obazoa;Opisthokonta;Holozoa;Choanozoa;Metazoa | EP00088 Acartia fossae 38523 | 1,10E-153 | 888 | XP 023346731.1 | protein transport protein Sec16-like isoform X2 [Eurytemora affinis] | 1941 | 0 | 587,03 | 72,55813953 |
| EP00088 | Obazoa;Opisthokonta;Holozoa;Choanozoa;Metazoa | EP00088 Acartia fossae 38524 | 1,10E-153 | 888 | XP 023346731.1 | protein transport protein Sec16-like isoform X2 [Eurytemora affinis] | 1941 | 0 | 587,03 | 72,55813953 |
| EP00088 | Obazoa;Opisthokonta;Holozoa;Choanozoa;Metazoa | EP00088 Acartia fossae 38525 | 1,10E-153 | 888 | XP 023346731.1 | protein transport protein Sec16-like isoform X2 [Eurytemora affinis] | 1941 | 0 | 587,03 | 72,55813953 |
| EP00096 | Obazoa;Opisthokonta;Holozoa;Choanozoa;Metazoa | EP00096 Caligus rogercresseyi 8847 | 7,60E-153 | 1116 | XP 023346731.1 | protein transport protein Sec16-like isoform X2 [Eurytemora affinis] | 1941 | 5,15E-132 | 452,981 | 49,12820513 |
| EP00256 | Archaeplastida;Chloroplastida;Streptophyta;"core s | EP00256 Glycine max 40939 | 4,70E-152 | 1412 | XP 006590566.1 | protein transport protein SEC16B homolog [Glycine max] >gi\|1590581767\|ref\|XP 02819: | 1412 | 0 | 2927,12 | 100 |
| EP00256 | Archaeplastida;Chloroplastida;Streptophyta;"core s | EP00256 Glycine max 19953 | 5,80E-152 | 1423 | XP 003538737.1 | protein transport protein SEC16B homolog [Glycine max] >gi\|1590582695\|ref\|XP 02819: | 1423 | 0 | 2951,77 | 100 |
| EP00074 | Obazoa;Opisthokonta;Holozoa;Choanozoa;Metazoa | EP00074 Homo sapiens 31138 | 9,70E-152 | 625 | NP 001343428.1 | protein transport protein Sec16B isoform 2 [Homo sapiens] | 625 | 0 | 1303,5 | 100 |
| EP00248 | Archaeplastida;Chloroplastida;Streptophyta;"core s | EP00248 Marchantia polymorpha 24255 | 1,60E-151 | 1733 | XP 019423445.1 | PREDICTED: protein transport protein SEC16B homolog [Lupinus angustifolius] >gi\|11174! | 1420 | 0 | 688,337 | 55,85464334 |
| EP00257 | Archaeplastida;Chloroplastida;Streptophyta;"core s | EP00257 Ricinus communis 27431 | 7,00E-151 | 1240 | XP 025014845.1 | protein transport protein SEC16B homolog [Ricinus communis] | 1240 | 0 | 2567,73 | 100 |
| EP00021 | Amoebozoa;Evosea;Eumycetozoa;Dictyostelia;Acyt | EP00021 Heterostelium album 5112 | 8,20E-151 | 1678 | XP 020431748.1 | hypothetical protein PPL 07486 [Heterostelium album PN500] | 1678 | 0 | 3358,93 | 100 |
| EP00256 | Archaeplastida;Chloroplastida;Streptophyta;"core s | EP00256 Glycine max 7771 | 1,20E-150 | 1424 | XP 003516665.1 | protein transport protein SEC16B homolog [Glycine max] | 1424 | 0 | 2945,61 | 100 |
| EP00251 | Archaeplastida;Chloroplastida;Streptophyta;"core s | EP00251 Ginkgo biloba 23000 | 2,50E-150 | 1651 | XP 006852622.1 | protein transport protein SEC16A homolog isoform X1 [Amborella trichopoda] | 1484 | 0 | 1088,56 | 57,73324984 |
| EP00268 | Archaeplastida;Chloroplastida;Streptophyta;"core s | EP00268 Persea americana 9752 | 6,70E-150 | 1226 | XP 010264304.1 | PREDICTED: protein transport protein SEC16A homolog isoform X1 [Nelumbo nucifera] | 1395 | 0 | 1353,19 | 66,66666667 |
| EP00268 | Archaeplastida;Chloroplastida;Streptophyta;"core s | EP00268 Persea americana 9751 | 3,10E-149 | 1235 | XP 010264304.1 | PREDICTED: protein transport protein SEC16A homolog isoform X1 [Nelumbo nucifera] | 1395 | 0 | 1361,67 | 66,30094044 |
| EP00264 | Archaeplastida;Chloroplastida;Streptophyta;"core s | EP00264 Hordeum vulgare 157067 | 3,60E-147 | 1033 | XP 020180505.1 | protein transport protein SEC16B homolog [Aegilops tauschii subsp. tauschii] | 1647 | 0 | 1944,86 | 93,41723136 |
| EP00022 | Amoebozoa;Evosea;Eumycetozoa;Dictyostelia;Acyt | EP00022 Cavenderia fasciculata 2800 | 1,30E-146 | 1778 | XP 004354463.1 | hypothetical protein DFA 10563 [Cavenderia fasciculata] | 1778 | 0 | 3585,42 | 100 |
| EP00123 | Obazoa;Opisthokonta;Holozoa;Ichthyosporea;Ichth | EP00123 Amoebidium parasiticum 6883 | 3,30E-146 | 993 | XP 004344939.1 | hypothetical protein ACA1 285550 [Acanthamoeba castellanii str. Neff] | 1515 | 1,37E-62 | 243,817 | 44,68085106 |
| EP00263 | Archaeplastida;Chloroplastida;Streptophyta;"core s | EP00263 Brachypodium distachyon 2663 | 5,50E-146 | 1554 | XP 003561234.1 | protein transport protein SEC16A homolog [Brachypodium distachyon] | 1554 | 0 | 3206,77 | 100 |
| EP00263 | Archaeplastida;Chloroplastida;Streptophyta;"core s | EP00263 Brachypodium distachyon 19486 | 7,30E-146 | 1509 | XP 010233662.1 | protein transport protein SEC16B homolog [Brachypodium distachyon] | 1509 | 0 | 3114,32 | 100 |
| EP00089 | Obazoa;Opisthokonta;Holozoa;Choanozoa;Metazoa | EP00089 Calanus finmarchicus 93268 | 1,00E-145 | 1300 | XP 023346731.1 | protein transport protein Sec16-like isoform X2 [Eurytemora affinis] | 1941 | 0 | 822,387 | 63,63636364 |
| EP00264 | Archaeplastida;Chloroplastida;Streptophyta;"core s | EP00264 Hordeum vulgare 157071 | 3,00E-145 | 1552 | XP 020180505.1 | protein transport protein SEC16B homolog [Aegilops tauschii subsp. tauschii] | 1647 | 0 | 2548,85 | 90,8045977 |
| EP00254 | Archaeplastida;Chloroplastida;Streptophyta;"core s | EP00254 Amborella trichopoda 7418 | 5,60E-145 | 1484 | XP 006852622.1 | protein transport protein SEC16A homolog isoform X1 [Amborella trichopoda] | 1484 | 0 | 3064,25 | 100 |
| EP00264 | Archaeplastida;Chloroplastida;Streptophyta;"core s | EP00264 Hordeum vulgare 41457 | 5,90E-145 | 1140 | XP 020151898.1 | protein transport protein SEC16A homolog [Aegilops tauschii subsp. tauschii] | 1490 | 0 | 1818,13 | 95,67456231 |
| EP00060 | Obazoa;Opisthokonta;Holozoa;Choanozoa;Metazoa | EP00060 Petromyzon marinus 19316 | 7,30E-145 | 2240 | XP 015222231.1 | PREDICTED: protein transport protein Sec16A isoform X1 [Lepisosteus oculatus] | 2483 | 0 | 713,375 | 66,45408163 |
| EP00057 | Obazoa;Opisthokonta;Holozoa;Choanozoa;Metazoa | EP00057 Strongylocentrotus purpuratus 29536 | 1,20E-144 | 1870 | XP 011684010.1 | PREDICTED: protein transport protein Sec16A isoform X2 [Strongylocentrotus purpuratus] | 1870 | 0 | 3801,91 | 100 |
| EP00264 | Archaeplastida;Chloroplastida;Streptophyta;"core s | EP00264 Hordeum vulgare 41454 | 1,40E-144 | 1140 | XP 020151898.1 | protein transport protein SEC16A homolog [Aegilops tauschii subsp. tauschii] | 1490 | 0 | 1780,76 | 95,69327731 |
| EP00264 | Archaeplastida;Chloroplastida;Streptophyta;"core s | EP00264 Hordeum vulgare 41458 | 1,70E-144 | 1125 | XP 020151898.1 | protein transport protein SEC16A homolog [Aegilops tauschii subsp. tauschii] | 1490 | 0 | 1805,8 | 95,64766839 |
| EP00057 | Obazoa;Opisthokonta;Holozoa;Choanozoa;Metazoa | EP00057 Strongylocentrotus purpuratus 3888 | 2,20E-144 | 1871 | XP 003727593.1 | PREDICTED: protein transport protein Sec16A isoform X1 [Strongylocentrotus purpuratus] | 1871 | 0 | 3803,06 | 100 |
| EP00130 | Obazoa;Opisthokonta;Nucletmycea;Fungi;"core Fur | EP00130 Batrachochytrium dendrobatidis 4985 | 3,40E-144 | 2643 | XP 006679860.1 | hypothetical protein BATDEDRAFT 25718 [Batrachochytrium dendrobatidis JAM81] | 2643 | 0 | 5449,4 | 100 |
| EP00031 | Obazoa;Apusomonadida;Thecamonadinae;Thecam | EP00031 Thecamonas trahens 10123 | 8,10E-144 | 2138 | XP 013762693.1 | hypothetical protein AMSG 00414 [Thecamonas trahens ATCC 50062] | 2138 | 0 | 4156,67 | 100 |
| EP00255 | Archaeplastida;Chloroplastida;Streptophyta;"core s | EP00255 Erythranthe guttata 17916 | 9,60E-144 | 1514 | XP 012845178.1 | PREDICTED: protein transport protein SEC16B homolog [Erythranthe guttata] >gi\|848890( | 1514 | 0 | 3123,95 | 100 |
| EP00255 | Archaeplastida;Chloroplastida;Streptophyta;"core s | EP00255 Erythranthe guttata 17917 | 9,60E-144 | 1514 | XP 012845178.1 | PREDICTED: protein transport protein SEC16B homolog [Erythranthe guttata] >gi\|848890( | 1514 | 0 | 3123,95 | 100 |
| EP00264 | Archaeplastida;Chloroplastida;Streptophyta;"core s | EP00264 Hordeum vulgare 41453 | 1,70E-143 | 1328 | XP 020151898.1 | protein transport protein SEC16A homolog [Aegilops tauschii subsp. tauschii] | 1490 | 0 | 2367,81 | 92,89985052 |
| EP00229 | Archaeplastida;Chloroplastida;Chlorophyta;Mamiel | EP00229 Micromonas commoda 9872 | 1,10E-142 | 1945 | XP 002509249.1 | predicted protein [Micromonas commoda] | 1945 | 0 | 3817,7 | 100 |
| EP00075 | Obazoa;Opisthokonta;Holozoa;Choanozoa;Metazoa | EP00075 Anas platyrhynchos 29384 | 1,80E-142 | 664 | XP 027318637.1 | protein transport protein Sec16B isoform X8 [Anas platyrhynchos] >gi\|1540448175\|ref\|X | 664 | 0 | 1353,96 | 100 |
| EP00075 | Obazoa;Opisthokonta;Holozoa;Choanozoa;Metazoa | EP00075 Anas platyrhynchos 29385 | 1,80E-142 | 664 | XP 027318637.1 | protein transport protein Sec16B isoform X8 [Anas platyrhynchos] >gi\|1540448175\|ref\|X | 664 | 0 | 1353,96 | 100 |
| EP00288 | Cryptista;Cryptophyceae;Cryptomonadales;"lineage | EP00288 Rhodomonas lens 10368 | 4,80E-142 | 677 | XP 020527880.1 | protein transport protein SEC16A homolog isoform X2 [Amborella trichopoda] | 1341 | 1,25E-67 | 251,906 | 42,81609195 |
| EP00249 | Archaeplastida;Chloroplastida;Streptophyta;"core s | EP00249 Psilotum nudum 22638 | 9,20E-142 | 1144 | XP 006852622.1 | protein transport protein SEC16A homolog isoform X1 [Amborella trichopoda] | 1484 | 0 | 709,523 | 54,97142857 |
| EP00264 | Archaeplastida;Chloroplastida;Streptophyta;"core s | EP00264 Hordeum vulgare 157070 | 2,00E-141 | 1582 | XP 020180505.1 | protein transport protein SEC16B homolog [Aegilops tauschii subsp. tauschii] | 1647 | 0 | 2565,03 | 91,06356199 |
| EP00145 | Obazoa;Opisthokonta;Nucletmycea;Fungi;"core Fur | EP00145 Schizosaccharomyces pombe 3423 | 2,40E-141 | 1995 | NP 594985.1 | putative multidomain vesicle coat component Sec16 [Schizosaccharomyces pombe] | 1995 | 0 | 4078,86 | 100 |
| EP00264 | Archaeplastida;Chloroplastida;Streptophyta;"core s | EP00264 Hordeum vulgare 41459 | 2,50E-141 | 1429 | XP 020151898.1 | protein transport protein SEC16A homolog [Aegilops tauschii subsp. tauschii] | 1490 | 0 | 2491,07 | 92,10342418 |
| EP00252 | Archaeplastida;Chloroplastida;Streptophyta;"core s | EP00252 Welwitschia mirabilis 9473 | 5,10E-141 | 1399 | XP 010264305.1 | PREDICTED: protein transport protein SEC16A homolog isoform X2 [Nelumbo nucifera] | 1393 | 0 | 771,155 | 55,41237113 |
| EP00252 | Archaeplastida;Chloroplastida;Streptophyta;"core s | EP00252 Welwitschia mirabilis 9475 | 1,90E-140 | 997 | XP 019423445.1 | PREDICTED: protein transport protein SEC16B homolog [Lupinus angustifolius] >gi\|11174! | 1420 | 0 | 736,102 | 56,00858369 |
| EP00530 | Sar;Stramenopiles;Gyrista;Ochrophyta;"BD clade";[ | EP00530 Phaeodactylum tricornutum 7488 | 2,60E-140 | 2083 | XP 002183614.1 | predicted protein [Phaeodactylum tricornutum CCAP 1055/1] | 2083 | 0 | 4234,48 | 100 |
| EP00267 | Archaeplastida;Chloroplastida;Streptophyta;"core s | EP00267 Zea mays 21331 | 5,80E-140 | 1478 | XP 008645146.1 | protein transport protein SEC16B homolog [Zea mays] | 1478 | 0 | 3055 | 100 |
| EP00252 | Archaeplastida;Chloroplastida;Streptophyta;"core s | EP00252 Welwitschia mirabilis 9474 | 1,10E-139 | 1751 | XP 010264305.1 | PREDICTED: protein transport protein SEC16A homolog isoform X2 [Nelumbo nucifera] | 1393 | 0 | 772,311 | 55,41237113 |
| EP00250 | Archaeplastida;Chloroplastida;Streptophyta;"core s | EP00250 Pteridium aquilinum 20169 | 1,40E-139 | 1632 | XP 006852622.1 | protein transport protein SEC16A homolog isoform X1 [Amborella trichopoda] | 1484 | 0 | 621,698 | 51,76817289 |
| EP00267 | Archaeplastida;Chloroplastida;Streptophyta;"core s | EP00267 Zea mays 38961 | 1,60E-139 | 1493 | NP 001348910.1 | uncharacterized protein LOC100383255 [Zea mays] | 1342 | 0 | 2787,29 | 100 |
| EP00099 | Obazoa;Opisthokonta;Holozoa;Choanozoa;Metazoa | EP00099 Drosophila melanogaster 1394 | 2,10E-139 | 2021 | NP 001096963.1 | secretory 16, isoform E [Drosophila melanogaster] | 2021 | 0 | 4117,38 | 100 |
| EP00250 | Archaeplastida;Chloroplastida;Streptophyta;"core s | EP00250 Pteridium aquilinum 21698 | 1,00E-138 | 1681 | XP 006852622.1 | protein transport protein SEC16A homolog isoform X1 [Amborella trichopoda] | 1484 | 0 | 680,633 | 50,07788162 |
| EP00264 | Archaeplastida;Chloroplastida;Streptophyta;"core s | EP00264 Hordeum vulgare 41455 | 1,20E-138 | 1274 | XP 020151898.1 | protein transport protein SEC16A homolog [Aegilops tauschii subsp. tauschii] | 1490 | 0 | 2184,84 | 91,21405751 |
| EP00024 | Amoebozoa;Evosea;Eumycetozoa;Dictyostelia;Dicty | EP00024 Dictyostelium purpureum 6047 | 1,70E-138 | 1854 | XP 003288899.1 | hypothetical protein DICPUDRAFT 98175 [Dictyostelium purpureum] | 1854 | 0 | 3693,28 | 100 |
| EP00254 | Archaeplastida;Chloroplastida;Streptophyta;"core s | EP00254 Amborella trichopoda 27042 | 2,10E-138 | 1341 | XP 020527880.1 | protein transport protein SEC16A homolog isoform X2 [Amborella trichopoda] | 1341 | 0 | 2777,66 | 100 |
| EP00266 | Archaeplastida;Chloroplastida;Streptophyta;"core s | EP00266 Sorghum bicolor 7869 | 2,30E-138 | 1479 | XP 002451351.1 | protein transport protein SEC16A homolog [Sorghum bicolor] | 1479 | 0 | 3053,08 | 100 |
| EP00059 | Obazoa;Opisthokonta;Holozoa;Choanozoa;Metazoa | EP00059 Ciona intestinalis 11360 | 2,30E-138 | 915 | XP 018669012.1 | protein transport protein Sec16A [Ciona intestinalis] | 915 | 0 | 1879,37 | 100 |
| EP00266 | Archaeplastida;Chloroplastida;Streptophyta;"core s | EP00266 Sorghum bicolor 31816 | 2,80E-138 | 1499 | XP 021314642.1 | protein transport protein SEC16A homolog isoform X2 [Sorghum bicolor] | 1499 | 0 | 3091,98 | 100 |
| EP00266 | Archaeplastida;Chloroplastida;Streptophyta;"core s | EP00266 Sorghum bicolor 31817 | 3,80E-138 | 1501 | XP 002451352.1 | protein transport protein SEC16A homolog isoform X1 [Sorghum bicolor] >gi\|1205951665 | 1501 | 0 | 3095,45 | 100 |
| EP00266 | Archaeplastida;Chloroplastida;Streptophyta;"core s | EP00266 Sorghum bicolor 7870 | 3,80E-138 | 1501 | XP 002451352.1 | protein transport protein SEC16A homolog isoform X1 [Sorghum bicolor] >gi\|1205951665 | 1501 | 0 | 3095,45 | 100 |
| EP00277 | Cryptista;Cryptophyceae;Cryptomonadales;"lineage | EP00277 Geminigera cryophila 22384 | 1,90E-137 | 1731 | XP 020527880.1 | protein transport protein SEC16A homolog isoform X2 [Amborella trichopoda] | 1341 | 1,08E-56 | 228,794 | 40,38997214 |
| EP00134 | Obazoa;Opisthokonta;Nucletmycea;Fungi;"core Fur | EP00134 Neurospora crassa 2112 | 1,10E-135 | 2084 | XP 011394599.1 | COPII coat assembly protein sec-16, variant 1 [Neurospora crassa OR74A] >gi\|758996814 | 2084 | 0 | 4210,22 | 100 |
| EP00134 | Obazoa;Opisthokonta;Nucletmycea;Fungi;"core Fur | EP00134 Neurospora crassa 2113 | 1,10E-135 | 2084 | XP 011394599.1 | COPII coat assembly protein sec-16, variant 1 [Neurospora crassa OR74A] >gi\|758996814 | 2084 | 0 | 4210,22 | 100 |
| EP00134 | Obazoa;Opisthokonta;Nucletmycea;Fungi;"core Fur | EP00134 Neurospora crassa 2114 | 1,10E-135 | 2084 | XP 011394599.1 | COPII coat assembly protein sec-16, variant 1 [Neurospora crassa OR74A] >gi\|758996814 | 2084 | 0 | 4210,22 | 100 |
| EP00276 | Archaeplastida;Glaucophyta;Gloeochaete;Gloeocha | EP00276 Gloeochaete wittrockiana 1783 | 2,80E-135 | 1443 | XP 022002241.1 | protein transport protein SEC16B homolog isoform X2 [Helianthus annuus] | 1329 | 2,55E-82 | 308,531 | 42,65463918 |
| EP00743 | Sar;Alveolata;Colponemidae;"MPE2-45 clade";strai | EP00743 Colponemidia sp Colp-15 2433 | 2,90E-134 | 810 | XP 020240850.1 | protein transport protein SEC16B homolog [Asparagus officinalis] | 1486 | 1,20E-58 | 228,409 | 51,22850123 |
| EP00574 | Sar;Stramenopiles;Gyrista;Ochrophyta;"BD clade";[ | EP00574 Skeletonema japonicum 187 | 5,10E-134 | 1800 | XP 002183614.1 | predicted protein [Phaeodactylum tricornutum CCAP 1055/1] | 2083 | 1,39E-112 | 409,453 | 47,63948498 |
| EP00027 | Amoebozoa;Evosea;Variosea;Filamoeba;Filamoeba | EP00027 Filamoeba nolandi 2639 | 1,80E-133 | 838 | XP 004354463.1 | hypothetical protein DFA 10563 [Cavenderia fasciculata] | 1778 | 8,01E-74 | 275,018 | 43,30042313 |
| EP00149 | Obazoa;Opisthokonta;Nucletmycea;Fungi;"core Fur | EP00149 Cryptococcus neoformans 425 | 3,50E-133 | 1998 | XP 012046565.1 | hypothetical protein CNAG 00148 [Cryptococcus neoformans var. grubii H99] | 1998 | 0 | 4005,68 | 100 |
| EP00260 | Archaeplastida;Chloroplastida;Streptophyta;"core s | EP00260 Arabidopsis thaliana 18309 | 4,50E-133 | 1061 | NP 199560.1 | RGPR-like protein [Arabidopsis thaliana] >gi\|1063736704\|ref\|NP 001318753.1\| RGPR-lik | 1361 | 0 | 2206,79 | 100 |
| EP00006 | Amoebozoa;Discosea;Centramoebia;"core centram | EP00006 Acanthamoeba castellanii 8457 | 5,70E-133 | 1515 | XP 004344939.1 | hypothetical protein ACA1 285550 [Acanthamoeba castellanii str. Neff] | 1515 | 0 | 3002,23 | 100 |
| EP00254 | Archaeplastida;Chloroplastida;Streptophyta;"core s | EP00254 Amborella trichopoda 22932 | 8,20E-133 | 1619 | XP 020523770.1 | LOW QUALITY PROTEIN: protein transport protein SEC16A homolog [Amborella trichopod | 1619 | 0 | 3350,07 | 100 |
| EP00269 | Archaeplastida;Chloroplastida;Streptophyta;"core s | EP00269 Selaginella moellendorffii 6455 | 1,30E-132 | 1190 | XP 002975264.1 | protein transport protein SEC16B homolog [Selaginella moellendorffii] | 1190 | 0 | 2457,17 | 100 |
| EP00267 | Archaeplastida;Chloroplastida;Streptophyta;"core s | EP00267 Zea mays 22917 | 1,40E-132 | 1821 | XP 008648102.1 | uncharacterized protein LOC100383190 isoform X2 [Zea mays] | 1821 | 0 | 3752,21 | 100 |
| EP00269 | Archaeplastida;Chloroplastida;Streptophyta;"core s | EP00269 Selaginella moellendorffii 11290 | 2,00E-132 | 1190 | XP 002987428.1 | protein transport protein SEC16B homolog [Selaginella moellendorffii] | 1190 | 0 | 2456,4 | 100 |
| EP00576 | Sar;Stramenopiles;Gyrista;Ochrophyta;"BD clade";[ | EP00576 Skeletonema menzelii 6755 | 2,30E-132 | 1866 | XP 002183614.1 | predicted protein [Phaeodactylum tricornutum CCAP 1055/1] | 2083 | 3,44E-118 | 427,557 | 52,00831025 |
| EP00260 | Archaeplastida;Chloroplastida;Streptophyta;"core s | EP00260 Arabidopsis thaliana 19676 | 2,80E-132 | 1054 | NP 001331519.1 | RGPR-like protein [Arabidopsis thaliana] | 1054 | 0 | 2189,85 | 100 |
| EP00575 | Sar;Stramenopiles;Gyrista;Ochrophyta;"BD clade";[ | EP00575 Skeletonema marinoi 38803 | 2,80E-132 | 1377 | XP 002183614.1 | predicted protein [Phaeodactylum tricornutum CCAP 1055/1] | 2083 | 1,03E-96 | 355,14 | 50 |
| EP00742 | Sar;Alveolata;Colponemidae;"MPE2-45 clade";strai | EP00742 Colponemidia sp Colp-10 7876 | 4,00E-132 | 772 | XP 004344939.1 | hypothetical protein ACA1 285550 [Acanthamoeba castellanii str. Neff] | 1515 | 4,23E-58 | 226,098 | 46,03481625 |
| EP00570 | Sar;Stramenopiles;Gyrista;Ochrophyta;"BD clade";[ | EP00570 Cyclotella meneghiniana 14282 | 4,50E-132 | 2728 | XP 002183614.1 | predicted protein [Phaeodactylum tricornutum CCAP 1055/1] | 2083 | 4,09E-102 | 379,407 | 52,72058824 |
| EP00551 | Sar;Stramenopiles;Gyrista;Ochrophyta;"BD clade";[ | EP00551 Chaetoceros affinis 13216 | 9,20E-132 | 2165 | XP 002183614.1 | predicted protein [Phaeodactylum tricornutum CCAP 1055/1] | 2083 | 9,50E-130 | 465,692 | 45,28301887 |
| EP00259 | Archaeplastida;Chloroplastida;Streptophyta;"core s | EP00259 Arabidopsis lyrata 14135 | 4,10E-131 | 1046 | XP 020866847.1 | protein transport protein SEC16A homolog isoform X2 [Arabidopsis lyrata subsp. lyrata] | 1356 | 0 | 2173,28 | 100 |
| EP00259 | Archaeplastida;Chloroplastida;Streptophyta;"core s | EP00259 Arabidopsis lyrata 14138 | 4,60E-131 | 1044 | XP 002865115.1 | protein transport protein SEC16A homolog isoform X1 [Arabidopsis lyrata subsp. lyrata] | 1358 | 0 | 2170,97 | 100 |
| EP00575 | Sar;Stramenopiles;Gyrista;Ochrophyta;"BD clade";[ | EP00575 Skeletonema marinoi 12924 | 4,90E-131 | 2138 | XP 002183614.1 | predicted protein [Phaeodactylum tricornutum CCAP 1055/1] | 2083 | 3,65E-108 | 397,897 | 49,84544049 |
| EP00575 | Sar;Stramenopiles;Gyrista;Ochrophyta;"BD clade";[ | EP00575 Skeletonema marinoi 12928 | 5,50E-131 | 2090 | XP 002183614.1 | predicted protein [Phaeodactylum tricornutum CCAP 1055/1] | 2083 | 5,91E-110 | 403,29 | 50,44247788 |

| | | | | | | | | | | |
|---|---|---|---|---|---|---|---|---|---|---|
| EP00089 | Obazoa;Opisthokonta;Holozoa;Choanozoa;Metazoa | EP00089 Calanus finmarchicus 93269 | 1,40E-130 | 845 | XP 023346731.1 | protein transport protein Sec16-like isoform X2 [Eurytemora affinis] | 1941 | 0 | 813,913 | 63,83219955 |
| EP00580 | Sar;Stramenopiles;Gyrista;Ochrophyta;"BD clade"; | EP00580 Thalassiosira gravida 19549 | 1,60E-130 | 1228 | XP 002183614.1 | predicted protein [Phaeodactylum tricornutum CCAP 1055/1] | 2083 | 4,91E-84 | 313,923 | 51,67785235 |
| EP00154 | Obazoa;Opisthokonta;Nucletmycea;Fungi;"core Fur | EP00154 Conidiobolus coronatus 6786 | 1,90E-130 | 1396 | XP 016275114.1 | COPII vesicle coat protein Sec16 [Rhodotorula toruloides NP11] | 2270 | 5,93E-67 | 262,307 | 48,11490126 |
| EP00276 | Archaeplastida;Glaucophyta;Gloeochaete;Gloeocha | EP00276 Gloeochaete wittrockiana 23328 | 3,70E-130 | 1453 | XP 022002241.1 | protein transport protein SEC16B homolog isoform X2 [Helianthus annuus] | 1329 | 1,47E-82 | 309,301 | 42,65463918 |
| EP00577 | Sar;Stramenopiles;Gyrista;Ochrophyta;"BD clade"; | EP00577 Skeletonema sp RCC1716 4364 | 1,10E-129 | 1373 | XP 002183614.1 | predicted protein [Phaeodactylum tricornutum CCAP 1055/1] | 2083 | 5,49E-98 | 358,992 | 49,04086739 |
| EP00266 | Archaeplastida;Chloroplastida;Streptophyta;"core s | EP00266 Sorghum bicolor 21428 | 2,40E-129 | 1891 | XP 021304254.1 | protein transport protein SEC16A homolog isoform X1 [Sorghum bicolor] | 1891 | 0 | 3894,35 | 100 |
| EP00569 | Sar;Stramenopiles;Gyrista;Ochrophyta;"BD clade"; | EP00569 Conticribra weissflogii 12916 | 3,20E-129 | 2390 | XP 002183614.1 | predicted protein [Phaeodactylum tricornutum CCAP 1055/1] | 2083 | 7,47E-117 | 426,402 | 51,91104934 |
| EP00577 | Sar;Stramenopiles;Gyrista;Ochrophyta;"BD clade"; | EP00577 Skeletonema sp RCC1716 4270 | 7,10E-129 | 1268 | XP 002183614.1 | predicted protein [Phaeodactylum tricornutum CCAP 1055/1] | 2083 | 1,72E-84 | 315,849 | 50,83955224 |
| EP00686 | Discoba;Heterolobosea;Tetramitia;Eutetramitia;"M | EP00686 Naegleria gruberi 15187 | 9,70E-129 | 1290 | XP 002683116.1 | RmlC-like cupin family protein [Naegleria gruberi] | 1290 | 0 | 2617,03 | 100 |
| EP00572 | Sar;Stramenopiles;Gyrista;Ochrophyta;"BD clade"; | EP00572 Skeletonema dohrnii 8548 | 1,00E-128 | 972 | XP 002183614.1 | predicted protein [Phaeodactylum tricornutum CCAP 1055/1] | 2083 | 3,68E-78 | 291,197 | 48,43260188 |
| EP00569 | Sar;Stramenopiles;Gyrista;Ochrophyta;"BD clade"; | EP00569 Conticribra weissflogii 1259 | 1,00E-128 | 2402 | XP 002183614.1 | predicted protein [Phaeodactylum tricornutum CCAP 1055/1] | 2083 | 4,99E-117 | 426,787 | 53,10344828 |
| EP00564 | Sar;Stramenopiles;Gyrista;Ochrophyta;"BD clade"; | EP00564 Ditylum brightwellii 14132 | 1,40E-128 | 1333 | XP 002183614.1 | predicted protein [Phaeodactylum tricornutum CCAP 1055/1] | 2083 | 8,75E-88 | 327,02 | 52,68817204 |
| EP00267 | Archaeplastida;Chloroplastida;Streptophyta;"core s | EP00267 Zea mays 22916 | 1,60E-128 | 1822 | NP 001348882.1 | uncharacterized protein LOC100383190 [Zea mays] | 1822 | 0 | 3754,14 | 100 |
| EP00119 | Obazoa;Opisthokonta;Holozoa;Choanozoa;Metazoa | EP00119 Amphimedon queenslandica 20158 | 2,40E-128 | 1105 | XP 019860577.1 | PREDICTED: protein transport protein Sec16A-like [Amphimedon queenslandica] | 1105 | 0 | 2260,72 | 100 |
| EP00118 | Obazoa;Opisthokonta;Holozoa;Choanozoa;Metazoa | EP00118 Oscarella pearsei 15406 | 7,90E-128 | 1816 | XP 022107909.1 | protein transport protein Sec16A-like isoform X3 [Acanthaster planci] | 2310 | 2,59E-91 | 342,813 | 45,04643963 |
| EP00577 | Sar;Stramenopiles;Gyrista;Ochrophyta;"BD clade"; | EP00577 Skeletonema sp RCC1716 3031 | 9,60E-128 | 938 | XP 002183614.1 | predicted protein [Phaeodactylum tricornutum CCAP 1055/1] | 2083 | 7,78E-74 | 277,33 | 47,20720721 |
| EP00266 | Archaeplastida;Chloroplastida;Streptophyta;"core s | EP00266 Sorghum bicolor 21429 | 1,10E-127 | 1890 | XP 021304255.1 | protein transport protein SEC16A homolog isoform X2 [Sorghum bicolor] | 1890 | 0 | 3892,43 | 100 |
| EP00578 | Sar;Stramenopiles;Gyrista;Ochrophyta;"BD clade"; | EP00578 Thalassiosira sp NH16 14626 | 1,50E-127 | 1390 | XP 002183614.1 | predicted protein [Phaeodactylum tricornutum CCAP 1055/1] | 2083 | 4,52E-90 | 334,724 | 48,8317757 |
| EP00577 | Sar;Stramenopiles;Gyrista;Ochrophyta;"BD clade"; | EP00577 Skeletonema sp RCC1716 5123 | 1,50E-127 | 2549 | XP 002183614.1 | predicted protein [Phaeodactylum tricornutum CCAP 1055/1] | 2083 | 5,24E-106 | 391,734 | 47,38438211 |
| EP00271 | Archaeplastida;Chloroplastida;Streptophyta;"core s | EP00271 Cylindrocystis brebissonii 3418 | 1,90E-127 | 2131 | XP 020527880.1 | protein transport protein SEC16A homolog isoform X2 [Amborella trichopoda] | 1341 | 1,83E-116 | 415,231 | 48,68004224 |
| EP00578 | Sar;Stramenopiles;Gyrista;Ochrophyta;"BD clade"; | EP00578 Thalassiosira sp NH16 8320 | 2,40E-127 | 1414 | XP 002183614.1 | predicted protein [Phaeodactylum tricornutum CCAP 1055/1] | 2083 | 5,43E-90 | 335,109 | 48,59484778 |
| EP00003 | CRuMs;Mantamonas;Mantamonas plastica;strain C | EP00003 Mantamonas plastica 22627 | 3,70E-127 | 972 | XP 004344939.1 | hypothetical protein ACA1 285550 [Acanthamoeba castellanii str. Neff] | 1515 | 4,37E-59 | 232,646 | 46,49227111 |
| EP00027 | Amoebozoa;Evosea;Variosea;Filamoeba;Filamoeba | EP00027 Filamoeba nolandi 11950 | 4,60E-127 | 888 | XP 004354463.1 | hypothetical protein DFA 10563 [Cavenderia fasciculata] | 1778 | 1,24E-79 | 293,508 | 42,63456091 |
| EP00584 | Sar;Stramenopiles;Gyrista;Ochrophyta;"BD clade"; | EP00584 Thalassiosira punctigera 26238 | 4,70E-127 | 2201 | XP 002183614.1 | predicted protein [Phaeodactylum tricornutum CCAP 1055/1] | 2083 | 2,79E-110 | 404,831 | 51,96787149 |
| EP00231 | Archaeplastida;Chloroplastida;Chlorophyta;Mamiel | EP00231 Micromonas pusilla 6752 | 5,60E-127 | 1814 | XP 003061507.1 | predicted protein [Micromonas pusilla CCMP1545] | 1814 | 0 | 3533,04 | 100 |
| EP00585 | Sar;Stramenopiles;Gyrista;Ochrophyta;"BD clade"; | EP00585 Thalassiosira rotula 4060 | 1,20E-126 | 1798 | XP 002183614.1 | predicted protein [Phaeodactylum tricornutum CCAP 1055/1] | 2083 | 2,44E-85 | 323,168 | 52,30024213 |
| EP00475 | Sar;Rhizaria;Endomyxa;"phytorhiza";Vampyrellida;' | EP00475 Leptophrys vorax 25199 | 1,50E-126 | 866 | XP 004344939.1 | hypothetical protein ACA1 285550 [Acanthamoeba castellanii str. Neff] | 1515 | 5,28E-53 | 212,231 | 46,0035524 |
| EP00475 | Sar;Rhizaria;Endomyxa;"phytorhiza";Vampyrellida;' | EP00475 Leptophrys vorax 25200 | 1,50E-126 | 866 | XP 004344939.1 | hypothetical protein ACA1 285550 [Acanthamoeba castellanii str. Neff] | 1515 | 5,28E-53 | 212,231 | 46,0035524 |
| EP00003 | CRuMs;Mantamonas;Mantamonas plastica;strain C | EP00003 Mantamonas plastica 22628 | 1,70E-126 | 917 | XP 004344939.1 | hypothetical protein ACA1 285550 [Acanthamoeba castellanii str. Neff] | 1515 | 8,94E-59 | 230,72 | 46,61117717 |
| EP00099 | Obazoa;Opisthokonta;Holozoa;Choanozoa;Metazoa | EP00099 Drosophila melanogaster 10847 | 2,80E-126 | 2137 | NP 001285157.1 | secretory 16, isoform H [Drosophila melanogaster] | 2137 | 0 | 4355,44 | 100 |
| EP00025 | Amoebozoa;Evosea;Eumycetozoa;Dictyostelia;Dicty | EP00025 Tieghemostelium lacteum 7869 | 1,10E-125 | 1912 | XP 646771.1 | hypothetical protein DDB G0269198 [Dictyostelium discoideum AX4] | 1992 | 5,31E-176 | 596,66 | 59,50155763 |
| EP00302 | Haptista;Haptophyta;Pavlovales;Diacronema;strain | EP00302 Diacronema sp CCMP2436 44594 | 1,60E-125 | 1496 | XP 004344939.1 | hypothetical protein ACA1 285550 [Acanthamoeba castellanii str. Neff] | 1515 | 2,28E-51 | 211,46 | 42,52631579 |
| EP00575 | Sar;Stramenopiles;Gyrista;Ochrophyta;"BD clade"; | EP00575 Skeletonema marinoi 3860 | 1,70E-125 | 2904 | XP 002183614.1 | predicted protein [Phaeodactylum tricornutum CCAP 1055/1] | 2083 | 1,06E-107 | 397,897 | 48,87556222 |
| EP00643 | Sar;Stramenopiles;Gyrista;Peronosporomycetes;"cc | EP00643 Hyaloperonospora arabidopsidis 7646 | 2,40E-125 | 2716 | XP 002908536.1 | COPII coat assembly protein SEC16, putative [Phytophthora infestans T30-4] | 2641 | 0 | 1250,73 | 62,88071066 |
| EP00148 | Obazoa;Opisthokonta;Nucletmycea;Fungi;"core Fur | EP00148 Cryptococcus gattii 4185 | 2,90E-125 | 1984 | XP 003191949.1 | Conserved hypothetical protein [Cryptococcus gattii WM276] | 2002 | 0 | 3530,73 | 95,45454545 |
| EP00072 | Obazoa;Opisthokonta;Holozoa;Choanozoa;Metazoa | EP00072 Mus musculus 66713 | 3,90E-125 | 565 | XP 017168476.1 | protein transport protein Sec16B isoform X4 [Mus musculus] | 565 | 0 | 1180,24 | 100 |
| EP00224 | Archaeplastida;Chloroplastida;Chlorophyta;Mamiel | EP00224 Ostreococcus tauri 6212 | 7,70E-125 | 1130 | XP 022839957.1 | COPII coat assembly protein, Sec16, partial [Ostreococcus tauri] | 1130 | 0 | 2290 | 100 |
| EP00512 | Sar;Stramenopiles;Bigyra;Sagenista;Labyrinthulomy | EP00512 Aurantiochytrium limacinum 11646 | 1,40E-124 | 1274 | XP 004344939.1 | hypothetical protein ACA1 285550 [Acanthamoeba castellanii str. Neff] | 1515 | 2,29E-48 | 201,06 | 52,50965251 |
| EP00090 | Obazoa;Opisthokonta;Holozoa;Choanozoa;Metazoa | EP00090 Calanus glacialis 314 | 1,80E-124 | 1812 | XP 023346731.1 | protein transport protein Sec16-like isoform X2 [Eurytemora affinis] | 1941 | 0 | 963,37 | 61,36195411 |
| EP00578 | Sar;Stramenopiles;Gyrista;Ochrophyta;"BD clade"; | EP00578 Thalassiosira sp NH16 7450 | 2,70E-124 | 1427 | XP 002183614.1 | predicted protein [Phaeodactylum tricornutum CCAP 1055/1] | 2083 | 2,10E-89 | 333,183 | 47,09371293 |
| EP00577 | Sar;Stramenopiles;Gyrista;Ochrophyta;"BD clade"; | EP00577 Skeletonema sp RCC1716 1513 | 5,10E-124 | 896 | XP 002183614.1 | predicted protein [Phaeodactylum tricornutum CCAP 1055/1] | 2083 | 1,86E-71 | 269,24 | 47,56756757 |
| EP00029 | Amoebozoa;Tubulinea;Echinamoebidia;Vermamoet | EP00029 Vermamoeba vermiformis 4110 | 7,60E-124 | 1658 | XP 004344939.1 | hypothetical protein ACA1 285550 [Acanthamoeba castellanii str. Neff] | 1515 | 1,05E-70 | 274,248 | 47,2693032 |
| EP00259 | Archaeplastida;Chloroplastida;Streptophyta;"core s | EP00259 Arabidopsis lyrata 19206 | 1,00E-123 | 1014 | XP 020871922.1 | LOW QUALITY PROTEIN: protein transport protein SEC16A homolog [Arabidopsis lyrata su | 1014 | 0 | 2098,94 | 100 |
| EP00081 | Obazoa;Opisthokonta;Holozoa;Choanozoa;Metazoa | EP00081 Caenorhabditis elegans 26016 | 2,60E-123 | 1632 | NP 509400.3 | Protein transport protein sec16 [Caenorhabditis elegans] | 1632 | 0 | 3383,19 | 100 |
| EP00090 | Obazoa;Opisthokonta;Holozoa;Choanozoa;Metazoa | EP00090 Calanus glacialis 315 | 2,70E-123 | 1808 | XP 023346731.1 | protein transport protein Sec16-like isoform X2 [Eurytemora affinis] | 1941 | 0 | 964,911 | 61,24721604 |
| EP00566 | Sar;Stramenopiles;Gyrista;Ochrophyta;"BD clade"; | EP00566 Odontella aurita 4283 | 3,20E-123 | 3089 | XP 002183614.1 | predicted protein [Phaeodactylum tricornutum CCAP 1055/1] | 2083 | 2,14E-133 | 479,945 | 51,51079137 |
| EP00013 | Amoebozoa;Discosea;Flabellinia;Stygamoeba;Styga | EP00013 Stygamoeba regulata 20863 | 4,50E-123 | 1396 | XP 004344939.1 | hypothetical protein ACA1 285550 [Acanthamoeba castellanii str. Neff] | 1515 | 1,64E-60 | 240,736 | 47,78012685 |
| EP00578 | Sar;Stramenopiles;Gyrista;Ochrophyta;"BD clade"; | EP00578 Thalassiosira sp NH16 21918 | 1,00E-122 | 964 | XP 002183614.1 | predicted protein [Phaeodactylum tricornutum CCAP 1055/1] | 2083 | 1,45E-70 | 268,085 | 53,49794239 |
| EP00560 | Sar;Stramenopiles;Gyrista;Ochrophyta;"BD clade"; | EP00560 Eucampia antarctica 4038 | 1,00E-122 | 1221 | XP 002183614.1 | predicted protein [Phaeodactylum tricornutum CCAP 1055/1] | 2083 | 2,74E-94 | 345,125 | 49,6350365 |
| EP00005 | Amoebozoa;Discosea;Centramoebia;"core centram | EP00005 Dracoamoeba jomungandri 4827 | 3,10E-122 | 1436 | XP 004344939.1 | hypothetical protein ACA1 285550 [Acanthamoeba castellanii str. Neff] | 1515 | 3,17E-92 | 340,117 | 45,22292994 |
| EP00099 | Obazoa;Opisthokonta;Holozoa;Choanozoa;Metazoa | EP00099 Drosophila melanogaster 10849 | 4,80E-122 | 2242 | NP 001285159.1 | secretory 16, isoform J [Drosophila melanogaster] | 2242 | 0 | 4577,31 | 100 |
| EP00667 | Discoba;Euglenozoa;Euglenida;Spirocuta;Euglenea; | EP00667 Euglena gracilis 4555 | 5,30E-122 | 664 | XP 004344939.1 | hypothetical protein ACA1 285550 [Acanthamoeba castellanii str. Neff] | 1515 | 8,75E-40 | 168,703 | 37,7672209 |
| EP00647 | Sar;Stramenopiles;Gyrista;Peronosporomycetes;"cc | EP00647 Phytophthora sojae 10217 | 5,40E-122 | 2808 | XP 009523981.1 | hypothetical protein PHYSODRAFT 262855 [Phytophthora sojae] | 2808 | 0 | 5603,87 | 100 |
| EP00306 | Haptista;Haptophyta;Pavlovales;Pavlova;strain CCN | EP00306 Pavlova sp CCMP459 16668 | 5,90E-122 | 1407 | XP 004344939.1 | hypothetical protein ACA1 285550 [Acanthamoeba castellanii str. Neff] | 1515 | 4,17E-59 | 236,113 | 47,2 |
| EP00646 | Sar;Stramenopiles;Gyrista;Peronosporomycetes;"cc | EP00646 Phytophthora ramorum 14933 | 7,30E-122 | 2462 | XP 002908536.1 | COPII coat assembly protein SEC16, putative [Phytophthora infestans T30-4] | 2641 | 0 | 2125,52 | 71,75112895 |
| EP00099 | Obazoa;Opisthokonta;Holozoa;Choanozoa;Metazoa | EP00099 Drosophila melanogaster 16955 | 5,80E-121 | 2250 | NP 572812.3 | secretory 16, isoform F [Drosophila melanogaster] | 2250 | 0 | 4594,26 | 100 |
| EP00319 | Haptista;Haptophyta;Prymnesiophyceae;Phaeocyst | EP00319 Phaeocystis globosa 14228 | 7,90E-121 | 634 | XP 002975264.1 | protein transport protein SEC16B homolog [Selaginella moellendorffii] | 1190 | 2,01E-45 | 184,882 | 40,41570439 |
| EP00579 | Sar;Stramenopiles;Gyrista;Ochrophyta;"BD clade"; | EP00579 Thalassiosira antarctica 25555 | 9,40E-121 | 2913 | XP 002183614.1 | predicted protein [Phaeodactylum tricornutum CCAP 1055/1] | 2083 | 9,77E-77 | 296,975 | 56,50080257 |
| EP00306 | Haptista;Haptophyta;Pavlovales;Pavlova;strain CCN | EP00306 Pavlova sp CCMP459 3062 | 3,50E-120 | 2342 | XP 004344939.1 | hypothetical protein ACA1 285550 [Acanthamoeba castellanii str. Neff] | 1515 | 5,00E-60 | 241,121 | 47,2 |
| EP00281 | Cryptista;Cryptophyceae;Cryptomonadales;"lineage | EP00281 Chroomonas sp CCMP1168 20830 | 3,70E-120 | 737 | XP 004344939.1 | hypothetical protein ACA1 285550 [Acanthamoeba castellanii str. Neff] | 1515 | 4,68E-58 | 225,328 | 42,51101322 |
| EP00147 | Obazoa;Opisthokonta;Nucletmycea;Fungi;"core Fur | EP00147 Laccaria bicolor 2250 | 5,10E-120 | 1596 | XP 001875397.1 | predicted protein [Laccaria bicolor S238N-H82] | 1596 | 0 | 3236,43 | 100 |
| EP00641 | Sar;Stramenopiles;Gyrista;Peronosporomycetes;"cc | EP00641 Albugo candida 2680 | 1,20E-119 | 2622 | XP 008890960.1 | hypothetical protein PPTG 01118 [Phytophthora parasitica INRA-310] | 1621 | 2,13E-169 | 579,326 | 64,24682396 |
| EP00641 | Sar;Stramenopiles;Gyrista;Peronosporomycetes;"cc | EP00641 Albugo candida 2677 | 2,70E-119 | 2605 | XP 008890960.1 | hypothetical protein PPTG 01118 [Phytophthora parasitica INRA-310] | 1621 | 1,39E-172 | 588,571 | 62,92134831 |
| EP00641 | Sar;Stramenopiles;Gyrista;Peronosporomycetes;"cc | EP00641 Albugo candida 2678 | 2,90E-119 | 2623 | XP 008890960.1 | hypothetical protein PPTG 01118 [Phytophthora parasitica INRA-310] | 1621 | 8,56E-168 | 574,704 | 64,3115942 |
| EP00514 | Sar;Stramenopiles;Bigyra;Sagenista;Labyrinthulomy | EP00514 Thraustochytrium sp LLF1b 10761 | 3,50E-119 | 2048 | XP 004344939.1 | hypothetical protein ACA1 285550 [Acanthamoeba castellanii str. Neff] | 1515 | 1,99E-48 | 202,986 | 47,90794979 |
| EP00641 | Sar;Stramenopiles;Gyrista;Peronosporomycetes;"cc | EP00641 Albugo candida 2679 | 4,20E-119 | 2606 | XP 008890960.1 | hypothetical protein PPTG 01118 [Phytophthora parasitica INRA-310] | 1621 | 5,58E-171 | 583,948 | 62,99065421 |
| EP00556 | Sar;Stramenopiles;Gyrista;Ochrophyta;"BD clade"; | EP00556 Chaetoceros neogracilis 6171 | 5,30E-119 | 2212 | XP 002183614.1 | predicted protein [Phaeodactylum tricornutum CCAP 1055/1] | 2083 | 4,49E-126 | 454,521 | 47,12871287 |
| EP00581 | Sar;Stramenopiles;Gyrista;Ochrophyta;"BD clade"; | EP00581 Thalassiosira minuscula 2189 | 6,00E-119 | 2866 | XP 002183614.1 | predicted protein [Phaeodactylum tricornutum CCAP 1055/1] | 2083 | 7,92E-122 | 442,965 | 50,16350556 |
| EP00578 | Sar;Stramenopiles;Gyrista;Ochrophyta;"BD clade"; | EP00578 Thalassiosira sp NH16 17141 | 6,70E-119 | 931 | XP 002183614.1 | predicted protein [Phaeodactylum tricornutum CCAP 1055/1] | 2083 | 3,63E-61 | 238,81 | 52,40384615 |
| EP00736 | Archaeplastida;"RR clade";Rhodelphis;Rhodelphis n | EP00736 Rhodelphis marinus 865 | 1,90E-118 | 1420 | XP 004344939.1 | hypothetical protein ACA1 285550 [Acanthamoeba castellanii str. Neff] | 1515 | 2,28E-49 | 204,912 | 44,36619718 |
| EP00462 | Sar;Rhizaria;Cercozoa;"crown cercozoa";Thecofilose | EP00462 Mataza sp D1 22271 | 2,70E-118 | 1234 | XP 004344939.1 | hypothetical protein ACA1 285550 [Acanthamoeba castellanii str. Neff] | 1515 | 3,65E-67 | 260,766 | 49,77924945 |
| EP00566 | Sar;Stramenopiles;Gyrista;Ochrophyta;"BD clade"; | EP00566 Odontella aurita 4341 | 5,00E-118 | 2260 | XP 002183614.1 | predicted protein [Phaeodactylum tricornutum CCAP 1055/1] | 2083 | 5,47E-130 | 467,233 | 52,2679693 |
| EP00089 | Obazoa;Opisthokonta;Holozoa;Choanozoa;Metazoa | EP00089 Calanus finmarchicus 93270 | 8,60E-118 | 748 | XP 023346731.1 | protein transport protein Sec16-like isoform X2 [Eurytemora affinis] | 1941 | 0 | 716,072 | 64,98103666 |
| EP00295 | Cryptista;Cryptophyceae;Goniomonas;"ma G clade" | EP00295 Goniomonas pacifica 8634 | 9,00E-118 | 980 | XP 020180505.1 | protein transport protein SEC16B homolog [Aegilops tauschii subsp. tauschii] | 1647 | 8,04E-51 | 207,223 | 40,75032342 |
| EP00554 | Sar;Stramenopiles;Gyrista;Ochrophyta;"BD clade"; | EP00554 Chaetoceros debilis 1952 | 1,00E-117 | 3457 | XP 002183614.1 | predicted protein [Phaeodactylum tricornutum CCAP 1055/1] | 2083 | 1,56E-123 | 449,129 | 47,58009543 |
| EP00125 | Obazoa;Opisthokonta;Holozoa;Ichthyosporea;Ichthy | EP00125 Sphaeroforma arctica 16493 | 3,90E-117 | 2420 | XP 014159568.1 | hypothetical protein SARC 02150 [Sphaeroforma arctica JP610] | 2420 | 0 | 4855,43 | 100 |
| EP00007 | Amoebozoa;Discosea;Flabellinia;"core flabellinids"; | EP00007 Cunea sp BSH-02190019 9886 | 4,90E-117 | 1697 | XP 004344939.1 | hypothetical protein ACA1 285550 [Acanthamoeba castellanii str. Neff] | 1515 | 4,41E-60 | 240,35 | 47,03389831 |
| EP00568 | Sar;Stramenopiles;Gyrista;Ochrophyta;"BD clade"; | EP00568 Trieres chinensis 14224 | 5,90E-117 | 1327 | XP 002183614.1 | predicted protein [Phaeodactylum tricornutum CCAP 1055/1] | 2083 | 4,25E-85 | 318,546 | 49,56949569 |
| EP00556 | Sar;Stramenopiles;Gyrista;Ochrophyta;"BD clade"; | EP00556 Chaetoceros neogracilis 20872 | 6,40E-117 | 1728 | XP 002183614.1 | predicted protein [Phaeodactylum tricornutum CCAP 1055/1] | 2083 | 1,43E-131 | 466,463 | 44,80910918 |
| EP00046 | Obazoa;Opisthokonta;Holozoa;Choanozoa;Choanof | EP00046 Monosiga brevicollis 5685 | 3,40E-116 | 2207 | XP 001747683.1 | hypothetical protein [Monosiga brevicollis MX1] | 2207 | 0 | 4491,03 | 100 |
| EP00223 | Archaeplastida;Chloroplastida;Chlorophyta;Mamiel | EP00223 Ostreococcus mediterraneus 5800 | 4,20E-116 | 1334 | XP 001420497.1 | predicted protein [Ostreococcus lucimarinus CCE9901] | 1174 | 0 | 866,3 | 58,42696629 |
| EP00092 | Obazoa;Opisthokonta;Holozoa;Choanozoa;Metazoa | EP00092 Neocalanus flemingeri 7473 | 8,70E-116 | 2334 | XP 023346731.1 | protein transport protein Sec16-like isoform X2 [Eurytemora affinis] | 1941 | 0 | 960,674 | 60,30418251 |
| EP00571 | Sar;Stramenopiles;Gyrista;Ochrophyta;"BD clade"; | EP00571 Detonula confervacea 14131 | 1,10E-115 | 2825 | XP 002183614.1 | predicted protein [Phaeodactylum tricornutum CCAP 1055/1] | 2083 | 1,61E-111 | 409,838 | 48,48684211 |
| EP00566 | Sar;Stramenopiles;Gyrista;Ochrophyta;"BD clade"; | EP00566 Odontella aurita 4282 | 1,20E-115 | 3091 | XP 002183614.1 | predicted protein [Phaeodactylum tricornutum CCAP 1055/1] | 2083 | 1,15E-133 | 480,715 | 51,36690647 |
| EP00741 | Archaeplastida;Glaucophyta;Cyanophora;Cyanopho | EP00741 Cyanophora paradoxa 7842 | 1,80E-115 | 2287 | XP 006852622.1 | protein transport protein SEC16A homolog isoform X1 [Amborella trichopoda] | 1484 | 1,56E-69 | 271,552 | 46,53465347 |
| EP00163 | Ancyromonadida;Fabomonas;Fabomonas tropica;st | EP00163 Fabomonas tropica 21794 | 2,30E-115 | 694 | XP 012756363.1 | hypothetical protein SAMD00019534 027760 [Acytostelium subglobosum LB1] | 1470 | 1,74E-56 | 219,55 | 40,55944056 |
| EP00671 | Discoba;Euglenozoa;Kinetoplastea;Metakinetoplast | EP00671 Bodo saltans 12290 | 3,30E-115 | 1477 | XP 814906.1 | hypothetical protein [Trypanosoma cruzi strain CL Brener] | 1482 | 1,75E-46 | 195,667 | 43,50282486 |
| EP00222 | Archaeplastida;Chloroplastida;Chlorophyta;Mamiel | EP00222 Ostreococcus lucimarinus 5140 | 3,60E-115 | 1174 | XP 001420497.1 | predicted protein [Ostreococcus lucimarinus CCE9901] | 1174 | 0 | 2371,27 | 100 |
| EP00081 | Obazoa;Opisthokonta;Holozoa;Choanozoa;Metazoa | EP00081 Caenorhabditis elegans 26015 | 3,80E-115 | 1614 | NP 509399.2 | Protein transport protein sec16 [Caenorhabditis elegans] | 1614 | 0 | 3346,98 | 100 |
| EP00081 | Obazoa;Opisthokonta;Holozoa;Choanozoa;Metazoa | EP00081 Caenorhabditis elegans 2445 | 8,10E-115 | 1609 | NP 001024557.1 | Protein transport protein sec16 [Caenorhabditis elegans] | 1609 | 0 | 3333,89 | 100 |
| EP00133 | Obazoa;Opisthokonta;Nucletmycea;Fungi;"core Fur | EP00133 Fusarium graminearum 8164 | 9,50E-115 | 1943 | XP 011323725.1 | hypothetical protein FGSG 05213 [Fusarium graminearum PH-1] | 1943 | 0 | 3927,1 | 100 |
| EP00264 | Archaeplastida;Chloroplastida;Streptophyta;"core s | EP00264 Hordeum vulgare 157066 | 1,90E-114 | 1157 | XP 020180505.1 | protein transport protein SEC16B homolog [Aegilops tauschii subsp. tauschii] | 1647 | 0 | 1838,16 | 91,12833764 |
| EP00146 | Obazoa;Opisthokonta;Nucletmycea;Fungi;"core Fur | EP00146 Coprinopsis cinerea 4306 | 2,50E-114 | 1628 | XP 001833253.2 | hypothetical protein CC1G 04232 [Coprinopsis cinerea okayama7#130] | 1628 | 0 | 3260,32 | 100 |
| EP00100 | Obazoa;Opisthokonta;Holozoa;Choanozoa;Metazoa | EP00100 Culex quinquefasciatus 16294 | 3,40E-114 | 1112 | XP 001868048.1 | conserved hypothetical protein [Culex quinquefasciatus] | 1112 | 0 | 2302,71 | 100 |
| EP00557 | Sar;Stramenopiles;Gyrista;Ochrophyta;"BD clade"; | EP00557 Chaetoceros sp GSL56 1656 | 3,50E-114 | 890 | XP 002183614.1 | predicted protein [Phaeodactylum tricornutum CCAP 1055/1] | 2083 | 2,86E-68 | 259,61 | 39,83050847 |
| EP00514 | Sar;Stramenopiles;Bigyra;Sagenista;Labyrinthulomy | EP00514 Thraustochytrium sp LLF1b 4331 | 7,30E-114 | 1894 | XP 004344939.1 | hypothetical protein ACA1 285550 [Acanthamoeba castellanii str. Neff] | 1515 | 1,75E-48 | 202,986 | 47,90794979 |
| EP00092 | Obazoa;Opisthokonta;Holozoa;Choanozoa;Metazoa | EP00092 Neocalanus flemingeri 7474 | 1,20E-113 | 1984 | XP 023346731.1 | protein transport protein Sec16-like isoform X2 [Eurytemora affinis] | 1941 | 0 | 958,748 | 60,22813688 |
| EP00557 | Sar;Stramenopiles;Gyrista;Ochrophyta;"BD clade"; | EP00557 Chaetoceros sp GSL56 5986 | 1,80E-113 | 904 | XP 002183614.1 | predicted protein [Phaeodactylum tricornutum CCAP 1055/1] | 2083 | 2,92E-67 | 256,914 | 41,20267261 |
| EP00092 | Obazoa;Opisthokonta;Holozoa;Choanozoa;Metazoa | EP00092 Neocalanus flemingeri 27971 | 3,10E-113 | 1885 | XP 023346731.1 | protein transport protein Sec16-like isoform X2 [Eurytemora affinis] | 1941 | 0 | 960,288 | 60,06051437 |
| EP00588 | Sar;Stramenopiles;Gyrista;Ochrophyta;"BD clade"; | EP00588 Corethron pennatum 27055 | 4,50E-113 | 1139 | XP 002183614.1 | predicted protein [Phaeodactylum tricornutum CCAP 1055/1] | 2083 | 1,78E-43 | 185,267 | 43,59605911 |
| EP00099 | Obazoa;Opisthokonta;Holozoa;Choanozoa;Metazoa | EP00099 Drosophila melanogaster 1395 | 1,40E-112 | 2221 | NP 001096964.1 | secretory 16, isoform D [Drosophila melanogaster] | 2221 | 0 | 4544,96 | 100 |
| EP00286 | Cryptista;Cryptophyceae;Cryptomonadales;"lineage | EP00286 Rhodomonas abbreviata 27402 | 1,50E-112 | 464 | XP 020180505.1 | protein transport protein SEC16B homolog [Aegilops tauschii subsp. tauschii] | 1647 | 3,38E-64 | 235,343 | 43,9516129 |
| EP00202 | Archaeplastida;Chloroplastida;Chlorophyta;"core ch | EP00202 Volvox carteri 8231 | 2,40E-112 | 3946 | XP 002953837.1 | hypothetical protein VOLCADRAFT 106114 [Volvox carteri f. nagariensis] | 3946 | 0 | 7765,22 | 100 |
| EP00546 | Sar;Stramenopiles;Gyrista;Ochrophyta;"BD clade"; | EP00546 Thalassionema frauenfeldii 16639 | 3,70E-112 | 2288 | XP 002183614.1 | predicted protein [Phaeodactylum tricornutum CCAP 1055/1] | 2083 | 2,56E-126 | 455,677 | 49,65089216 |
| EP00515 | Sar;Stramenopiles;Bigyra;Sagenista;Labyrinthulomy | EP00515 Schizochytrium aggregatum 20978 | 3,80E-112 | 2258 | XP 009523981.1 | hypothetical protein PHYSODRAFT 262855 [Phytophthora sojae] | 2808 | 3,91E-40 | 176,792 | 44,98269896 |
| EP00304 | Haptista;Haptophyta;Pavlovales;Pavlova;Pavlova gy | EP00304 Pavlova gyrans 14843 | 4,80E-112 | 1461 | XP 004344939.1 | hypothetical protein ACA1 285550 [Acanthamoeba castellanii str. Neff] | 1515 | 9,79E-53 | 215,698 | 41,29032258 |
| EP00289 | Cryptista;Cryptophyceae;Cryptomonadales;"lineage | EP00289 Rhodomonas salina 11823 | 8,90E-112 | 1134 | XP 005822601.1 | hypothetical protein GUITHDRAFT 158726 [Guillardia theta CCMP2712] | 305 | 6,17E-130 | 407,912 | 72,4137931 |
| EP00637 | Sar;Stramenopiles;Gyrista;Ochrophyta;Pinguiophyc | EP00637 Pinguiococcus pyrenoidosus 1658 | 1,20E-111 | 1192 | XP 008890960.1 | hypothetical protein PPTG 01118 [Phytophthora parasitica INRA-310] | 1621 | 5,87E-53 | 215,698 | 46,06240713 |
| EP00136 | Obazoa;Opisthokonta;Nucletmycea;Fungi;"core Fur | EP00136 Aspergillus niger 5887 | 1,80E-111 | 1835 | XP 001396692.2 | COPII coat assembly protein SEC16 [Aspergillus niger CBS 513.88] | 1835 | 0 | 3671,32 | 100 |
| EP00556 | Sar;Stramenopiles;Gyrista;Ochrophyta;"BD clade"; | EP00556 Chaetoceros neogracilis 1844 | 7,30E-111 | 1104 | XP 002183614.1 | predicted protein [Phaeodactylum tricornutum CCAP 1055/1] | 2083 | 1,76E-94 | 343,199 | 51,31004367 |
| EP00645 | Sar;Stramenopiles;Gyrista;Peronosporomycetes;"cc | EP00645 Phytophthora infestans 13838 | 9,80E-111 | 2641 | XP 002908536.1 | COPII coat assembly protein SEC16, putative [Phytophthora infestans T30-4] | 2641 | 0 | 5341,55 | 100 |
| EP00609 | Sar;Stramenopiles;Gyrista;Ochrophyta;"PX clade";P | EP00609 Ectocarpus siliculosus 15252 | 1,10E-110 | 3470 | XP 025416207.1 | pro-resilin-like [Sipha flava] | 740 | 2,40E-11 | 81,6481 | 48,94514768 |
| EP00587 | Sar;Stramenopiles;Gyrista;Ochrophyta;"BD clade"; | EP00587 Corethron hystrix 974 | 2,00E-110 | 1102 | XP 002183614.1 | predicted protein [Phaeodactylum tricornutum CCAP 1055/1] | 2083 | 9,09E-41 | 176,407 | 45,36817102 |
| EP00517 | Sar;Stramenopiles;Bigyra;Sagenista;Labyrinthulomy | EP00517 Thraustochytrida sp NY07348D 9537 | 2,20E-110 | 1970 | XP 030285773.1 | uncharacterized threonine-rich GPI-anchored glycoprotein PJ4664.02-like isoform X3 [Spa | 2016 | 4,82E-28 | 136,346 | 73,34851936 |
| EP00137 | Obazoa;Opisthokonta;Nucletmycea;Fungi;"core Fur | EP00137 Aspergillus oryzae 8179 | 2,70E-110 | 1826 | XP 001823030.1 | unnamed protein product [Aspergillus oryzae RIB40] | 1843 | 0 | 3641,66 | 99,89148128 |
| EP00587 | Sar;Stramenopiles;Gyrista;Ochrophyta;"BD clade"; | EP00587 Corethron hystrix 930 | 4,30E-110 | 1095 | XP 002183614.1 | predicted protein [Phaeodactylum tricornutum CCAP 1055/1] | 2083 | 1,38E-40 | 175,637 | 45,36817102 |
| EP00687 | Discoba;Heterolobosea;Tetramitia;Eutetramitia;"PS | EP00687 Percolomonas cosmopolitus 13867 | 4,70E-110 | 1260 | XP 004344939.1 | hypothetical protein ACA1 285550 [Acanthamoeba castellanii str. Neff] | 1515 | 3,08E-56 | 226,483 | 43,21223709 |
| EP00161 | Ancyromonadida;Ancyromonadidae;Ancyromonas; | EP00161 Ancyromonas sigmoides 15047 | 1,10E-109 | 865 | XP 004344939.1 | hypothetical protein ACA1 285550 [Acanthamoeba castellanii str. Neff] | 1515 | 1,63E-45 | 189,119 | 53,21969697 |
| EP00546 | Sar;Stramenopiles;Gyrista;Ochrophyta;"BD clade"; | EP00546 Thalassionema frauenfeldii 3683 | 1,30E-109 | 2242 | XP 002183614.1 | predicted protein [Phaeodactylum tricornutum CCAP 1055/1] | 2083 | 2,99E-127 | 458,373 | 49,8060512 |
| EP00092 | Obazoa;Opisthokonta;Holozoa;Choanozoa;Metazoa | EP00092 Neocalanus flemingeri 6266 | 1,60E-109 | 1751 | XP 023346731.1 | protein transport protein Sec16-like isoform X2 [Eurytemora affinis] | 1941 | 0 | 960,288 | 59,98492841 |
| EP00162 | Ancyromonadida;Ancyromonadidae;Nutomonas;Nu | EP00162 Nutomonas longa 10758 | 1,90E-109 | 543 | XP 004344939.1 | hypothetical protein ACA1 285550 [Acanthamoeba castellanii str. Neff] | 1515 | 5,96E-67 | 246,128 | 46,83544304 |
| EP00290 | Cryptista;Cryptophyceae;Cryptomonadales;Baffinel | EP00290 Baffinella sp CCMP2293 55407 | 3,20E-109 | 559 | XP 004344939.1 | hypothetical protein ACA1 285550 [Acanthamoeba castellanii str. Neff] | 1515 | 7,84E-56 | 214,157 | 40,3887689 |
| EP00513 | Sar;Stramenopiles;Bigyra;Sagenista;Labyrinthulomy | EP00513 Hondaea fermentalgiana 11061 | 9,40E-109 | 2387 | XP 004344939.1 | hypothetical protein ACA1 285550 [Acanthamoeba castellanii str. Neff] | 1515 | 3,15E-47 | 199,519 | 53,52380952 |
| EP00012 | Amoebozoa;Discosea;Flabellinia;"core flabellinids"; | EP00012 Vannella robusta 14462 | 9,80E-109 | 946 | XP 003288899.1 | hypothetical protein DICPUDRAFT 98175 [Dictyostelium purpureum] | 1854 | 1,32E-44 | 187,193 | 42,33687405 |
| EP00544 | Sar;Stramenopiles;Gyrista;Ochrophyta;"BD clade"; | EP00544 Staurosira sp CCMP2646 2551 | 1,20E-108 | 1995 | XP 002183614.1 | predicted protein [Phaeodactylum tricornutum CCAP 1055/1] | 2083 | 5,15E-162 | 559,681 | 50,7969508 |
| EP00642 | Sar;Stramenopiles;Gyrista;Peronosporomycetes;"cc | EP00642 Albugo laibachii 13666 | 1,60E-108 | 2659 | XP 008890960.1 | hypothetical protein PPTG 01118 [Phytophthora parasitica INRA-310] | 1621 | 1,98E-172 | 588,571 | 65,04854369 |
| EP00516 | Sar;Stramenopiles;Bigyra;Sagenista;Labyrinthulomy | EP00516 Thraustochytrida sp NY0313808BC1 1231 | 1,60E-108 | 1968 | XP 030285773.1 | uncharacterized threonine-rich GPI-anchored glycoprotein PJ4664.02-like isoform X3 [Spa | 2016 | 4,66E-28 | 136,346 | 73,34851936 |
| EP00594 | Sar;Stramenopiles;Gyrista;Ochrophyta;"BD clade"; | EP00594 Rhizosolenia setigera 10409 | 1,90E-108 | 1099 | XP 002183614.1 | predicted protein [Phaeodactylum tricornutum CCAP 1055/1] | 2083 | 4,83E-96 | 347,821 | 46,46840149 |
| EP00153 | Obazoa;Opisthokonta;Nucletmycea;Fungi;"core Fur | EP00153 Phycomyces blakesleeanus 11044 | 2,00E-108 | 1879 | XP 018294121.1 | hypothetical protein PHYBLDRAFT 75955 [Phycomyces blakesleeanus NRRL 1555(-)] | 1879 | 0 | 3876,25 | 100 |
| EP00594 | Sar;Stramenopiles;Gyrista;Ochrophyta;"BD clade"; | EP00594 Rhizosolenia setigera 5805 | 2,00E-108 | 1119 | XP 002183614.1 | predicted protein [Phaeodactylum tricornutum CCAP 1055/1] | 2083 | 4,47E-101 | 363,614 | 47,09236032 |
| EP00267 | Archaeplastida;Chloroplastida;Streptophyta;"core s | EP00267 Zea mays 38962 | 2,10E-108 | 1205 | XP 008677596.1 | protein transport protein SEC16B-like protein isoform X2 [Zea mays] | 1205 | 0 | 2506,09 | 100 |
| EP00594 | Sar;Stramenopiles;Gyrista;Ochrophyta;"BD clade"; | EP00594 Rhizosolenia setigera 12801 | 2,10E-108 | 1097 | XP 002183614.1 | predicted protein [Phaeodactylum tricornutum CCAP 1055/1] | 2083 | 4,79E-96 | 347,821 | 46,46840149 |
| EP00161 | Ancyromonadida;Ancyromonadidae;Ancyromonas; | EP00161 Ancyromonas sigmoides 15045 | 2,20E-108 | 792 | XP 004344939.1 | hypothetical protein ACA1 285550 [Acanthamoeba castellanii str. Neff] | 1515 | 1,62E-45 | 188,348 | 53,21969697 |
| EP00753 | Sar;Stramenopiles;Platysulcus;Platysulcus tardus;st | EP00753 Platysulcus tardus 4649 | 7,60E-108 | 1174 | XP 002967613.1 | galactokinase isoform X1 [Selaginella moellendorffii] | 505 | 1,50E-104 | 348,977 | 47,6861167 |
| EP00135 | Obazoa;Opisthokonta;Nucletmycea;Fungi;"core Fur | EP00135 Aspergillus nidulans 6611 | 1,00E-107 | 1947 | XP 664219.1 | hypothetical protein AN6615.2 [Aspergillus nidulans FGSC A4] | 1947 | 0 | 3917,47 | 100 |
| EP00655 | Sar;Stramenopiles;Gyrista;Peronosporomycetes;"cc | EP00655 Saprolegnia parasitica 13720 | 1,20E-107 | 2601 | XP 012207226.1 | hypothetical protein SPRG 12070 [Saprolegnia parasitica CBS 223.65] | 2601 | 0 | 5182,85 | 100 |
| EP00653 | Sar;Stramenopiles;Gyrista;Peronosporomycetes;"cc | EP00653 Aphanomyces invadans 10450 | 2,20E-107 | 2510 | XP 008871247.1 | hypothetical protein H310 07614 [Aphanomyces invadans] | 2510 | 0 | 5083,46 | 100 |
| EP00236 | Archaeplastida;Chloroplastida;Chlorophyta;Picocyst | EP00236 Picocystis salinarum 3261 | 4,90E-107 | 1470 | XP 028053810.1 | protein transport protein SEC16B homolog [Camellia sinensis] | 1469 | 3,82E-72 | 278,1 | 43,8410596 |
| EP00326 | Haptista;Haptophyta;Prymnesiophyceae;Prymnesia | EP00326 Haptolina ericina 42296 | 5,20E-107 | 1188 | XP 004344939.1 | hypothetical protein ACA1 285550 [Acanthamoeba castellanii str. Neff] | 1515 | 1,02E-42 | 182,57 | 46,13636364 |
| EP00144 | Obazoa;Opisthokonta;Nucletmycea;Fungi;"core Fur | EP00144 Saccharomyces cerevisiae 5369 | 5,60E-107 | 2195 | NP 015240.1 | Sec16p [Saccharomyces cerevisiae S288C] | 2195 | 0 | 4489,87 | 100 |
| EP00099 | Obazoa;Opisthokonta;Holozoa;Choanozoa;Metazoa | EP00099 Drosophila melanogaster 10848 | 9,70E-107 | 2326 | NP 001285158.1 | secretory 16, isoform I [Drosophila melanogaster] | 2326 | 0 | 4767,6 | 100 |
| EP00326 | Haptista;Haptophyta;Prymnesiophyceae;Prymnesia | EP00326 Haptolina ericina 14908 | 1,10E-106 | 1234 | XP 004344939.1 | hypothetical protein ACA1 285550 [Acanthamoeba castellanii str. Neff] | 1515 | 1,40E-42 | 182,185 | 46,13636364 |
| EP00211 | Archaeplastida;Chloroplastida;Chlorophyta;Chlorop | EP00211 Chloroparvula japonica 3564 | 1,20E-106 | 777 | XP 025811964.1 | protein transport protein SEC16A homolog isoform X1 [Panicum hallii] | 1891 | 3,40E-72 | 268,855 | 47,66763848 |
| EP00654 | Sar;Stramenopiles;Gyrista;Peronosporomycetes;"cc | EP00654 Saprolegnia diclina 5277 | 2,10E-106 | 2628 | XP 008608960.1 | hypothetical protein SDRG 05043 [Saprolegnia diclina VS20] | 2628 | 0 | 5237,93 | 100 |
| EP00586 | Sar;Stramenopiles;Gyrista;Ochrophyta;"BD clade"; | EP00586 Coscinodiscus wailesii 8664 | 2,50E-106 | 2997 | XP 002183614.1 | predicted protein [Phaeodactylum tricornutum CCAP 1055/1] | 2083 | 4,44E-93 | 350,517 | 49,09662215 |
| EP00051 | Obazoa;Opisthokonta;Holozoa;Choanozoa;Choanof | EP00051 Salpingoeca urceolata 25752 | 2,60E-106 | 1604 | XP 004344939.1 | hypothetical protein ACA1 285550 [Acanthamoeba castellanii str. Neff] | 1515 | 4,58E-40 | 174,866 | 41,4893617 |
| EP00129 | Obazoa;Opisthokonta;Nucletmycea;Fungi;"core Fur | EP00129 Allomyces macrogynus 8111 | 2,70E-106 | 2350 | XP 010760357.1 | hypothetical protein PADG 04750 [Paracoccidioides brasiliensis Pb18] | 1822 | 1,41E-39 | 174,481 | 44,12416851 |
| EP00559 | Sar;Stramenopiles;Gyrista;Ochrophyta;"BD clade"; | EP00559 Dactyliosolen fragilissimus 8136 | 3,20E-106 | 1076 | XP 002183614.1 | predicted protein [Phaeodactylum tricornutum CCAP 1055/1] | 2083 | 3,62E-82 | 305,449 | 50,31605563 |
| EP00326 | Haptista;Haptophyta;Prymnesiophyceae;Prymnesia | EP00326 Haptolina ericina 40544 | 3,90E-106 | 1212 | XP 004344939.1 | hypothetical protein ACA1 285550 [Acanthamoeba castellanii str. Neff] | 1515 | 1,31E-42 | 182,185 | 46,13636364 |
| EP00328 | Haptista;Haptophyta;Prymnesiophyceae;Prymnesia | EP00328 Prymnesium polylepis 59141 | 6,80E-106 | 903 | XP 004344939.1 | hypothetical protein ACA1 285550 [Acanthamoeba castellanii str. Neff] | 1515 | 6,64E-40 | 171,785 | 45,83333333 |
| EP00264 | Archaeplastida;Chloroplastida;Streptophyta;"core s | EP00264 Hordeum vulgare 157068 | 1,20E-105 | 1119 | XP 020180505.1 | protein transport protein SEC16B homolog [Aegilops tauschii subsp. tauschii] | 1647 | 0 | 1758,81 | 90,83629893 |
| EP00543 | Sar;Stramenopiles;Gyrista;Ochrophyta;"BD clade"; | EP00543 Licmophora paradoxa 8987 | 2,00E-105 | 1561 | XP 002183614.1 | predicted protein [Phaeodactylum tricornutum CCAP 1055/1] | 2083 | 7,03E-125 | 443,736 | 46,44852354 |
| EP00280 | Cryptista;Cryptophyceae;Cryptomonadales;"lineage | EP00280 Hanusia phi 19393 | 3,40E-105 | 420 | XP 027180549.1 | protein transport protein SEC16B homolog [Coffea eugenioides] | 1487 | 6,18E-48 | 186,422 | 40,24767802 |
| EP00012 | Amoebozoa;Discosea;Flabellinia;"core flabellinids"; | EP00012 Vannella robusta 1918 | 7,20E-105 | 1133 | XP 003288899.1 | hypothetical protein DICPUDRAFT 98175 [Dictyostelium purpureum] | 1854 | 2,14E-45 | 191,045 | 40 |
| EP00519 | Sar;Stramenopiles;Gyrista;Ochrophyta;"BD clade"; | EP00519 Triparma laevis 4888 | 1,30E-104 | 1070 | XP 002183614.1 | predicted protein [Phaeodactylum tricornutum CCAP 1055/1] | 2083 | 2,19E-50 | 206,838 | 43,94736842 |
| EP00563 | Sar;Stramenopiles;Gyrista;Ochrophyta;"BD clade"; | EP00563 Minutocellus polymorphus 6113 | 3,50E-104 | 2193 | XP 002183614.1 | predicted protein [Phaeodactylum tricornutum CCAP 1055/1] | 2083 | 1,48E-92 | 347,821 | 48,87152778 |
| EP00339 | Sar;Alveolata;Ciliophora;Intramacronucleata;"CON | EP00339 Tiarina fusus 25463 | 7,20E-104 | 1196 | XP 004344939.1 | hypothetical protein ACA1 285550 [Acanthamoeba castellanii str. Neff] | 1515 | 5,40E-44 | 186,808 | 39,77485929 |
| EP00650 | Sar;Stramenopiles;Gyrista;Peronosporomycetes;"cc | EP00650 Pythium arrhenomanes 918 | 8,30E-104 | 2175 | XP 002908536.1 | COPII coat assembly protein SEC16, putative [Phytophthora infestans T30-4] | 2641 | 0 | 661,374 | 70,29288703 |
| EP00549 | Sar;Stramenopiles;Gyrista;Ochrophyta;"BD clade"; | EP00549 Striatella unipunctata 21869 | 9,00E-104 | 1544 | XP 002183614.1 | predicted protein [Phaeodactylum tricornutum CCAP 1055/1] | 2083 | 1,38E-119 | 427,557 | 49,19908467 |
| EP00085 | Obazoa;Opisthokonta;Holozoa;Choanozoa;Metazoa | EP00085 Daphnia pulex 28804 | 1,10E-103 | 275 | XP 023234695.1 | protein transport protein Sec16A-like [Centruroides sculpturatus] | 764 | 3,10E-112 | 349,747 | 65,68265683 |
| EP00339 | Sar;Alveolata;Ciliophora;Intramacronucleata;"CON | EP00339 Tiarina fusus 25666 | 1,40E-103 | 1181 | XP 004344939.1 | hypothetical protein ACA1 285550 [Acanthamoeba castellanii str. Neff] | 1515 | 7,42E-44 | 186,037 | 39,77485929 |
| EP00549 | Sar;Stramenopiles;Gyrista;Ochrophyta;"BD clade"; | EP00549 Striatella unipunctata 21864 | 1,90E-103 | 1549 | XP 002183614.1 | predicted protein [Phaeodactylum tricornutum CCAP 1055/1] | 2083 | 2,55E-119 | 426,787 | 49,7723824 |
| EP00652 | Sar;Stramenopiles;Gyrista;Peronosporomycetes;"cc | EP00652 Aphanomyces astaci 10476 | 2,20E-103 | 2384 | XP 009831237.1 | hypothetical protein H257 07417 [Aphanomyces astaci] | 2384 | 0 | 4804,19 | 100 |
| EP00557 | Sar;Stramenopiles;Gyrista;Ochrophyta;"BD clade"; | EP00557 Chaetoceros sp GSL56 10718 | 3,90E-103 | 770 | XP 002183614.1 | predicted protein [Phaeodactylum tricornutum CCAP 1055/1] | 2083 | 1,80E-50 | 203,371 | 40,3785489 |
| EP00253 | Archaeplastida;Chloroplastida;Streptophyta;"core s | EP00253 Pinus taeda 23969 | 1,40E-102 | 733 | XP 010264304.1 | PREDICTED: protein transport protein SEC16A homolog isoform X1 [Nelumbo nucifera] | 1395 | 0 | 624,394 | 58 |

| | | | | | | | | | | |
|---|---|---|---|---|---|---|---|---|---|---|
| EP00054 | Obazoa;Opisthokonta;Holozoa;Choanozoa;Choanof | EP00054 Salpingoeca dolichothecata 27075 | 2,60E-102 | 1370 | XP 014769113.1 | PREDICTED: uncharacterized protein LOC106868398 isoform X4 [Octopus bimaculoides] | 2159 | 3,67E-43 | 184,882 | 46,20123203 |
| EP00002 | CRuMs;Collodictyonidae;Diphylleia;Diphylleia rotan | EP00002 Diphylleia rotans 22687 | 2,70E-102 | 1305 | XP 004344939.1 | hypothetical protein ACA1 285550 [Acanthamoeba castellanii str. Neff] | 1515 | 1,83E-47 | 198,364 | 34,38735178 |
| EP00123 | Obazoa;Opisthokonta;Holozoa;Ichthyosporea;Ichthy | EP00123 Amoebidium parasiticum 14287 | 4,40E-102 | 1111 | XP 004344939.1 | hypothetical protein ACA1 285550 [Acanthamoeba castellanii str. Neff] | 1515 | 1,41E-50 | 207,223 | 43,40425532 |
| EP00294 | Cryptista;Cryptophyceae;Goniomonas;"ma G clade" | EP00294 Goniomonas avonlea 15680 | 7,10E-102 | 587 | XP 020527880.1 | protein transport protein SEC16A homolog isoform X2 [Amborella trichopoda] | 1341 | 2,17E-45 | 183,726 | 40,37478705 |
| EP00590 | Sar;Stramenopiles;Gyrista;Ochrophyta;"BD clade";[ | EP00590 Aulacoseira subarctica 8993 | 1,60E-101 | 1924 | XP 002183614.1 | predicted protein [Phaeodactylum tricornutum CCAP 1055/1] | 2083 | 3,50E-84 | 319,701 | 47,00285442 |
| EP00002 | CRuMs;Collodictyonidae;Diphylleia;Diphylleia rotan | EP00002 Diphylleia rotans 22693 | 2,30E-101 | 1374 | XP 004344939.1 | hypothetical protein ACA1 285550 [Acanthamoeba castellanii str. Neff] | 1515 | 1,87E-47 | 198,364 | 34,38735178 |
| EP00548 | Sar;Stramenopiles;Gyrista;Ochrophyta;"BD clade";[ | EP00548 Thalassiothrix antarctica 4632 | 1,10E-100 | 1667 | XP 002183614.1 | predicted protein [Phaeodactylum tricornutum CCAP 1055/1] | 2083 | 2,31E-123 | 441,039 | 47,20734507 |
| EP00044 | Obazoa;Opisthokonta;Holozoa;Choanozoa;Choanof | EP00044 Salpingoeca rosetta 4597 | 1,90E-100 | 2149 | XP 004992030.1 | hypothetical protein PTSG 12463 [Salpingoeca rosetta] | 2149 | 0 | 4355,82 | 100 |
| EP00656 | Telonemia;Telonema;"Tsubtile clade";Telonema sul | EP00656 Telonema subtile 11507 | 2,70E-100 | 644 | XP 013728346.1 | protein transport protein SEC16A homolog [Brassica napus] | 1358 | 6,04E-39 | 165,622 | 47,14475432 |
| EP00555 | Sar;Stramenopiles;Gyrista;Ochrophyta;"BD clade";[ | EP00555 Chaetoceros dichaeta 260 | 7,20E-100 | 1417 | XP 002183614.1 | predicted protein [Phaeodactylum tricornutum CCAP 1055/1] | 2083 | 1,82E-92 | 342,813 | 49,33214604 |
| EP00300 | Haptista;Centroplasthelida;Pterocystida;Raphidista | EP00300 Choanocystis sp HF-7 18814 | 7,40E-100 | 715 | XP 003288899.1 | hypothetical protein DICPUDRAFT 98175 [Dictyostelium purpureum] | 1854 | 4,17E-32 | 145,591 | 38,79518072 |
| EP00026 | Amoebozoa;Evosea;Eumycetozoa;Myxogastria;Colu | EP00026 Physarum polycephalum 1296 | 1,10E-99 | 1167 | XP 646771.1 | hypothetical protein DDB G0269198 [Dictyostelium discoideum AX4] | 1992 | 1,33E-86 | 320,472 | 52,26337449 |
| EP00537 | Sar;Stramenopiles;Gyrista;Ochrophyta;"BD clade";[ | EP00537 Pseudo-nitzschia pungens 7448 | 1,40E-99 | 4056 | XP 002183614.1 | predicted protein [Phaeodactylum tricornutum CCAP 1055/1] | 2083 | 6,74E-123 | 447,588 | 49,80724749 |
| EP00537 | Sar;Stramenopiles;Gyrista;Ochrophyta;"BD clade";[ | EP00537 Pseudo-nitzschia pungens 19767 | 1,70E-99 | 2896 | XP 002183614.1 | predicted protein [Phaeodactylum tricornutum CCAP 1055/1] | 2083 | 1,76E-126 | 457,603 | 46,07768469 |
| EP00558 | Sar;Stramenopiles;Gyrista;Ochrophyta;"BD clade";[ | EP00558 Chaetoceros sp UNC1202 2446 | 1,80E-99 | 698 | XP 002183614.1 | predicted protein [Phaeodactylum tricornutum CCAP 1055/1] | 2083 | 7,86E-55 | 214,927 | 44,3902439 |
| EP00533 | Sar;Stramenopiles;Gyrista;Ochrophyta;"BD clade";[ | EP00533 Pseudo-nitzschia delicatissima 4842 | 4,50E-99 | 2797 | XP 002183614.1 | predicted protein [Phaeodactylum tricornutum CCAP 1055/1] | 2083 | 1,52E-123 | 448,358 | 49,88814318 |
| EP00339 | Sar;Alveolata;Ciliophora;Intramacronucleata;"CON: | EP00339 Tiarina fusus 25618 | 4,80E-99 | 1164 | XP 004344939.1 | hypothetical protein ACA1 285550 [Acanthamoeba castellanii str. Neff] | 1515 | 7,89E-44 | 186,037 | 39,77485929 |
| EP00562 | Sar;Stramenopiles;Gyrista;Ochrophyta;"BD clade";[ | EP00562 Extubocellulus spinifer 3562 | 6,40E-99 | 2252 | XP 002183614.1 | predicted protein [Phaeodactylum tricornutum CCAP 1055/1] | 2083 | 1,31E-99 | 370,548 | 49,67105263 |
| EP00761 | Discoba;Heterolobosea;Pharyngomonas;Pharyngon | EP00761 Pharyngomonas kirbyi 1530 | 6,50E-99 | 660 | XP 021631646.1 | protein transport protein SEC16B homolog isoform X1 [Manihot esculenta] >gi\|12162366 | 1430 | 1,69E-36 | 158,303 | 42,1978022 |
| EP00228 | Archaeplastida;Chloroplastida;Chlorophyta;Mamiel | EP00228 Micromonas bravo 1884 | 6,60E-99 | 1314 | XP 003061507.1 | predicted protein [Micromonas pusilla CCMP1545] | 1814 | 1,03E-84 | 317,005 | 57,4795082 |
| EP00471 | Sar;Rhizaria;Cercozoa;Chlorarachnea;"core chlorara | EP00471 Norrisiella sphaerica 12160 | 6,70E-99 | 1084 | XP 004344939.1 | hypothetical protein ACA1 285550 [Acanthamoeba castellanii str. Neff] | 1515 | 2,82E-48 | 199,904 | 46,88221709 |
| EP00531 | Sar;Stramenopiles;Gyrista;Ochrophyta;"BD clade";[ | EP00531 Pseudo-nitzschia arenysensis 8997 | 1,30E-98 | 2201 | XP 002183614.1 | predicted protein [Phaeodactylum tricornutum CCAP 1055/1] | 2083 | 3,23E-125 | 451,825 | 48,86807182 |
| EP00575 | Sar;Stramenopiles;Gyrista;Ochrophyta;"BD clade";[ | EP00575 Skeletonema marinoi 24745 | 1,50E-98 | 615 | XP 002183614.1 | predicted protein [Phaeodactylum tricornutum CCAP 1055/1] | 2083 | 2,18E-42 | 175,637 | 46,41509434 |
| EP00052 | Obazoa;Opisthokonta;Holozoa;Choanozoa;Choanof | EP00052 Salpingoeca macrocollata 10498 | 2,90E-98 | 981 | XP 004344939.1 | hypothetical protein ACA1 285550 [Acanthamoeba castellanii str. Neff] | 1515 | 2,46E-41 | 177,178 | 42,9280397 |
| EP00531 | Sar;Stramenopiles;Gyrista;Ochrophyta;"BD clade";[ | EP00531 Pseudo-nitzschia arenysensis 9033 | 3,70E-98 | 2402 | XP 002183614.1 | predicted protein [Phaeodactylum tricornutum CCAP 1055/1] | 2083 | 2,06E-125 | 453,366 | 48,79000781 |
| EP00472 | Sar;Rhizaria;Cercozoa;Chlorarachnea;"core chlorara | EP00472 Partenskyella glossopodia 276 | 4,50E-98 | 1412 | XP 004344939.1 | hypothetical protein ACA1 285550 [Acanthamoeba castellanii str. Neff] | 1515 | 5,07E-59 | 236,113 | 49,22178988 |
| EP00109 | Obazoa;Opisthokonta;Holozoa;Choanozoa;Metazoa | EP00109 Schistosoma mansoni 8929 | 6,10E-98 | 1407 | XP 018652912.1 | rgpr-related [Schistosoma mansoni] | 1407 | 0 | 2914,79 | 100 |
| EP00531 | Sar;Stramenopiles;Gyrista;Ochrophyta;"BD clade";[ | EP00531 Pseudo-nitzschia arenysensis 6735 | 6,30E-98 | 833 | XP 002183614.1 | predicted protein [Phaeodactylum tricornutum CCAP 1055/1] | 2083 | 3,15E-75 | 279,256 | 45,66115702 |
| EP00562 | Sar;Stramenopiles;Gyrista;Ochrophyta;"BD clade";[ | EP00562 Extubocellulus spinifer 31491 | 8,50E-98 | 1759 | XP 002183614.1 | predicted protein [Phaeodactylum tricornutum CCAP 1055/1] | 2083 | 8,49E-92 | 343,584 | 49,78540773 |
| EP00531 | Sar;Stramenopiles;Gyrista;Ochrophyta;"BD clade";[ | EP00531 Pseudo-nitzschia arenysensis 8924 | 9,20E-98 | 1940 | XP 002183614.1 | predicted protein [Phaeodactylum tricornutum CCAP 1055/1] | 2083 | 1,86E-126 | 453,751 | 48,86807182 |
| EP00469 | Sar;Rhizaria;Cercozoa;Chlorarachnea;"core chlorara | EP00469 Lotharella globosa 13128 | 9,70E-98 | 1082 | XP 004344939.1 | hypothetical protein ACA1 285550 [Acanthamoeba castellanii str. Neff] | 1515 | 3,97E-49 | 202,601 | 46,37681159 |
| EP00200 | Archaeplastida;Chloroplastida;Chlorophyta;"core ch | EP00200 Dunaliella tertiolecta 15083 | 1,30E-97 | 1592 | XP 012472009.1 | PREDICTED: protein transport protein SEC16B homolog isoform X1 [Gossypium raimondii] | 1391 | 7,42E-47 | 196,823 | 52,25806452 |
| EP00518 | Sar;Stramenopiles;Gyrista;Ochrophyta;"BD clade";E | EP00518 Triparma eleuthera 18045 | 1,60E-97 | 992 | XP 002183614.1 | predicted protein [Phaeodactylum tricornutum CCAP 1055/1] | 2083 | 8,23E-47 | 194,897 | 48,28973843 |
| EP00532 | Sar;Stramenopiles;Gyrista;Ochrophyta;"BD clade";[ | EP00532 Pseudo-nitzschia australis 6196 | 2,90E-97 | 1302 | XP 002183614.1 | predicted protein [Phaeodactylum tricornutum CCAP 1055/1] | 2083 | 6,99E-100 | 363,614 | 48,31704669 |
| EP00531 | Sar;Stramenopiles;Gyrista;Ochrophyta;"BD clade";[ | EP00531 Pseudo-nitzschia arenysensis 9022 | 3,10E-97 | 2435 | XP 002183614.1 | predicted protein [Phaeodactylum tricornutum CCAP 1055/1] | 2083 | 2,38E-125 | 453,366 | 48,79000781 |
| EP00534 | Sar;Stramenopiles;Gyrista;Ochrophyta;"BD clade";[ | EP00534 Pseudo-nitzschia fraudulenta 9194 | 3,40E-97 | 1485 | XP 002183614.1 | predicted protein [Phaeodactylum tricornutum CCAP 1055/1] | 2083 | 2,15E-109 | 395,586 | 52,69461078 |
| EP00233 | Archaeplastida;Chloroplastida;Palmophyllophyceae | EP00233 Prasinococcus capsulatus 4848 | 3,50E-97 | 1521 | XP 020240850.1 | protein transport protein SEC16B homolog [Asparagus officinalis] | 1486 | 2,12E-64 | 253,447 | 48,37905237 |
| EP00531 | Sar;Stramenopiles;Gyrista;Ochrophyta;"BD clade";[ | EP00531 Pseudo-nitzschia arenysensis 978 | 3,70E-97 | 970 | XP 002183614.1 | predicted protein [Phaeodactylum tricornutum CCAP 1055/1] | 2083 | 9,10E-80 | 296,204 | 46,16613419 |
| EP00434 | Sar;Alveolata;Myzozoa;"DP clade";Dinoflagellata;D | EP00434 Breviolum minutum 32947 | 5,20E-97 | 853 | XP 004344939.1 | hypothetical protein ACA1 285550 [Acanthamoeba castellanii str. Neff] | 1515 | 1,03E-17 | 100,523 | 42,29828851 |
| EP00265 | Archaeplastida;Chloroplastida;Streptophyta;"core s | EP00265 Oryza sativa 3438 | 5,80E-97 | 634 | XP 015626058.1 | protein transport protein SEC16B homolog [Oryza sativa Japonica Group] | 1514 | 0 | 1316,98 | 100 |
| EP00103 | Obazoa;Opisthokonta;Holozoa;Choanozoa;Metazoa | EP00103 Capitella teleta 4648 | 6,00E-97 | 2190 | XP 013096820.1 | PREDICTED: uncharacterized protein LOC106080064 isoform X8 [Biomphalaria glabrata] | 1837 | 2,05E-155 | 539,65 | 55,29801325 |
| EP00649 | Sar;Stramenopiles;Gyrista;Peronosporomycetes;"cc | EP00649 Plasmopara halstedii 13216 | 7,20E-97 | 2672 | XP 024584649.1 | Regucalcin gene promoter region-related protein (RGPR) [Plasmopara halstedii] | 2672 | 0 | 5494,09 | 100 |
| EP00531 | Sar;Stramenopiles;Gyrista;Ochrophyta;"BD clade";[ | EP00531 Pseudo-nitzschia arenysensis 16721 | 1,50E-96 | 1521 | XP 002183614.1 | predicted protein [Phaeodactylum tricornutum CCAP 1055/1] | 2083 | 3,35E-124 | 441,039 | 48,79000781 |
| EP00192 | Archaeplastida;Chloroplastida;Chlorophyta;"core ch | EP00192 Tetraselmis astigmatica 1544 | 1,50E-96 | 1718 | XP 024373338.1 | protein transport protein SEC16A homolog isoform X1 [Physcomitrella patens] >gi\|13738 | 1444 | 8,17E-62 | 245,743 | 42,30769231 |
| EP00132 | Obazoa;Opisthokonta;Nucletmycea;Fungi;"core Fur | EP00132 Gonapodya prolifera 2262 | 1,50E-96 | 1107 | XP 016606650.1 | hypothetical protein SPPG 06291 [Spizellomyces punctatus DAOM BR117] | 2211 | 2,05E-68 | 263,848 | 47,59358289 |
| EP00208 | Archaeplastida;Chloroplastida;Chlorophyta;"core ch | EP00208 Stichococcus sp RCC1054 8927 | 1,90E-96 | 2837 | XP 005649504.1 | hypothetical protein COCSUDRAFT 83652 [Coccomyxa subellipsoidea C-169] | 1231 | 1,40E-61 | 245,358 | 53,60824742 |
| EP00434 | Sar;Alveolata;Myzozoa;"DP clade";Dinoflagellata;D | EP00434 Breviolum minutum 32946 | 2,20E-96 | 844 | XP 004344939.1 | hypothetical protein ACA1 285550 [Acanthamoeba castellanii str. Neff] | 1515 | 9,86E-18 | 100,523 | 42,29828851 |
| EP00280 | Cryptista;Cryptophyceae;Cryptomonadales;"lineage | EP00280 Hanusia phi 17694 | 2,50E-96 | 332 | XP 023916411.1 | protein transport protein SEC16B homolog [Quercus suber] | 1465 | 1,09E-47 | 182,57 | 44,58204334 |
| EP00217 | Archaeplastida;Chloroplastida;Chlorophyta;Mamiel | EP00217 Crustomastix stigmatica 5691 | 2,80E-96 | 865 | XP 001420497.1 | predicted protein [Ostreococcus lucimarinus CCE9901] | 1174 | 9,79E-72 | 267,7 | 45,72425829 |
| EP00547 | Sar;Stramenopiles;Gyrista;Ochrophyta;"BD clade";[ | EP00547 Thalassionema nitzschioides 10426 | 2,90E-96 | 2523 | XP 002183614.1 | predicted protein [Phaeodactylum tricornutum CCAP 1055/1] | 2083 | 4,90E-129 | 465,307 | 44,99640029 |
| EP00269 | Archaeplastida;Chloroplastida;Streptophyta;"core s | EP00269 Selaginella moellendorffii 31690 | 3,40E-96 | 671 | XP 024532336.1 | protein transport protein SEC16B homolog isoform X1 [Selaginella moellendorffii] | 671 | 0 | 1387,09 | 100 |
| EP00269 | Archaeplastida;Chloroplastida;Streptophyta;"core s | EP00269 Selaginella moellendorffii 4945 | 3,40E-96 | 655 | XP 002971690.1 | protein transport protein SEC16B homolog isoform X2 [Selaginella moellendorffii] | 655 | 0 | 1352,04 | 100 |
| EP00213 | Archaeplastida;Chloroplastida;Chlorophyta;Chlorop | EP00213 Chloropicon mariensis 2649 | 4,20E-96 | 1345 | XP 024985174.1 | protein transport protein SEC16B homolog isoform X1 [Cynara cardunculus var. scolymus] | 1457 | 7,85E-58 | 231,876 | 47,07207207 |
| EP00269 | Archaeplastida;Chloroplastida;Streptophyta;"core s | EP00269 Selaginella moellendorffii 12732 | 4,20E-96 | 676 | XP 002990968.1 | protein transport protein SEC16A homolog [Selaginella moellendorffii] | 676 | 0 | 1417,14 | 100 |
| EP00327 | Haptista;Haptophyta;Prymnesiophyceae;Prymnesia | EP00327 Prymnesium parvum 41469 | 6,00E-96 | 2312 | XP 012756363.1 | hypothetical protein SAMD00019534 027760 [Acytostelium subglobosum LB1] | 1470 | 1,58E-34 | 157,532 | 34,88372093 |
| EP00050 | Obazoa;Opisthokonta;Holozoa;Choanozoa;Choanof | EP00050 Salpingoeca kvevrii 11299 | 9,70E-96 | 1978 | XP 004344939.1 | hypothetical protein ACA1 285550 [Acanthamoeba castellanii str. Neff] | 1515 | 2,79E-45 | 192,586 | 44,06779661 |
| EP00216 | Archaeplastida;Chloroplastida;Chlorophyta;Chlorop | EP00216 Chloropicon sp CCMP2111 5416 | 1,10E-95 | 1538 | XP 006852622.1 | protein transport protein SEC16A homolog isoform X1 [Amborella trichopoda] | 1484 | 1,17E-55 | 225,328 | 42,52336449 |
| EP00464 | Sar;Rhizaria;Cercozoa;Chlorarachnea;"core chlorara | EP00464 Amorphochlora amoebiformis 6292 | 1,20E-95 | 855 | XP 004344939.1 | hypothetical protein ACA1 285550 [Acanthamoeba castellanii str. Neff] | 1515 | 1,47E-38 | 167,162 | 43,23725055 |
| EP00613 | Sar;Stramenopiles;Gyrista;Ochrophyta;Dictyochoph | EP00613 Pedinella sp CCMP2098 26823 | 1,70E-95 | 1492 | XP 008232573.2 | PREDICTED: LOW QUALITY PROTEIN: protein transport protein SEC16B homolog [Prunus n | 1429 | 4,86E-25 | 125,561 | 49,18032787 |
| EP00327 | Haptista;Haptophyta;Prymnesiophyceae;Prymnesia | EP00327 Prymnesium parvum 14463 | 2,10E-95 | 1975 | XP 012756363.1 | hypothetical protein SAMD00019534 027760 [Acytostelium subglobosum LB1] | 1470 | 1,69E-32 | 150,599 | 37,31707317 |
| EP00269 | Archaeplastida;Chloroplastida;Streptophyta;"core s | EP00269 Selaginella moellendorffii 43073 | 6,00E-95 | 676 | XP 024543719.1 | protein transport protein SEC16A homolog [Selaginella moellendorffii] | 676 | 0 | 1417,52 | 100 |
| EP00129 | Obazoa;Opisthokonta;Nucletmycea;Fungi;"core Fur | EP00129 Allomyces macrogynus 9092 | 9,40E-95 | 2438 | XP 007690667.1 | hypothetical protein COCMIDRAFT 102534 [Bipolaris oryzae ATCC 44560] | 1799 | 9,52E-33 | 152,14 | 50,99667774 |
| EP00727 | Amoebozoa;Evosea;Archamoebea;Mastigamoebida | EP00727 Mastigamoeba balamuthi 6046 | 1,20E-94 | 1707 | XP 004354463.1 | hypothetical protein DFA 10563 [Cavenderia fasciculata] | 1778 | 7,83E-78 | 298,13 | 38,95216401 |
| EP00591 | Sar;Stramenopiles;Gyrista;Ochrophyta;"BD clade";[ | EP00591 Stephanopyxis turris 18413 | 2,70E-94 | 1906 | XP 002183614.1 | predicted protein [Phaeodactylum tricornutum CCAP 1055/1] | 2083 | 3,86E-70 | 274,248 | 49,65034965 |
| EP00755 | Discoba;Euglenozoa;Diplonemea;Diplonemidae;Sul | EP00755 Sulcionema specki 33431 | 3,90E-94 | 649 | ***no hit found*** | | | | | |
| EP00047 | Obazoa;Opisthokonta;Holozoa;Choanozoa;Choanof | EP00047 Mylnosiga fluctuans 20884 | 7,60E-94 | 1626 | XP 004344939.1 | hypothetical protein ACA1 285550 [Acanthamoeba castellanii str. Neff] | 1515 | 1,65E-40 | 176,407 | 44,39140811 |
| EP00523 | Sar;Stramenopiles;Gyrista;Ochrophyta;"BD clade";[ | EP00523 Amphiprora sp CCMP467 6926 | 7,70E-94 | 1400 | XP 002183614.1 | predicted protein [Phaeodactylum tricornutum CCAP 1055/1] | 2083 | 8,21E-105 | 380,563 | 49,73544974 |
| EP00658 | Telonemia;Telonema;strain P-2 | EP00658 Telonema sp P-2 54003 | 1,30E-93 | 816 | XP 021304255.1 | protein transport protein SEC16A homolog isoform X2 [Sorghum bicolor] | 1890 | 4,89E-39 | 168,703 | 47,50542299 |
| EP00151 | Obazoa;Opisthokonta;Nucletmycea;Fungi;"core Fur | EP00151 Rhizophagus irregularis 13061 | 1,70E-93 | 530 | XP 025177105.1 | Sec23-binding domain of Sec16-domain-containing protein, partial [Rhizophagus irregular | 530 | 0 | 1096,26 | 100 |
| EP00526 | Sar;Stramenopiles;Gyrista;Ochrophyta;"BD clade";[ | EP00526 Cylindrotheca closterium 1422 | 1,80E-93 | 1939 | XP 002183614.1 | predicted protein [Phaeodactylum tricornutum CCAP 1055/1] | 2083 | 3,44E-102 | 377,867 | 52,55813953 |
| EP00215 | Archaeplastida;Chloroplastida;Chlorophyta;Chlorop | EP00215 Chloropicon roscoffensis 5973 | 2,00E-93 | 1345 | XP 019423445.1 | PREDICTED: protein transport protein SEC16B homolog [Lupinus angustifolius] >gi\|11174 | 1420 | 7,84E-55 | 221,861 | 51,42857143 |
| EP00125 | Obazoa;Opisthokonta;Holozoa;Ichthyosporea;Ichthy | EP00125 Sphaeroforma arctica 16618 | 3,60E-93 | 1511 | XP 014159693.1 | hypothetical protein SARC 02054 [Sphaeroforma arctica JP610] | 1511 | 0 | 3095,45 | 100 |
| EP00299 | Haptista;Centroplasthelida;Pterocystida;Pterista;"P | EP00299 Pterocystis canadensis 17688 | 4,60E-93 | 747 | XP 012756363.1 | hypothetical protein SAMD00019534 027760 [Acytostelium subglobosum LB1] | 1470 | 1,70E-30 | 140,969 | 38,52040816 |
| EP00108 | Obazoa;Opisthokonta;Holozoa;Choanozoa;Metazoa | EP00108 Taenia solium 1338 | 8,30E-93 | 1262 | XP 024351479.1 | Protein transport protein Sec16A [Echinococcus granulosus] | 1265 | 0 | 1804,65 | 84,17266187 |
| EP00751 | Sar;Stramenopiles;Gyrista;Ochrophyta;"BD clade";[ | EP00751 Fragilariopsis kerguelensis 19268 | 1,30E-92 | 2878 | XP 002183614.1 | predicted protein [Phaeodactylum tricornutum CCAP 1055/1] | 2083 | 4,39E-93 | 350,517 | 52,21238938 |
| EP00269 | Archaeplastida;Chloroplastida;Streptophyta;"core s | EP00269 Selaginella moellendorffii 22256 | 1,50E-92 | 557 | XP 024522902.1 | protein transport protein SEC16A homolog, partial [Selaginella moellendorffii] | 557 | 0 | 1163,29 | 100 |
| EP00656 | Telonemia;Telonema;"Tsubtile clade";Telonema sul | EP00656 Telonema subtile 11513 | 1,60E-92 | 1190 | XP 013762693.1 | hypothetical protein AMSG 00414 [Thecamonas trahens ATCC 50062] | 2138 | 5,20E-45 | 190,274 | 48,17073171 |
| EP00117 | Obazoa;Opisthokonta;Holozoa;Choanozoa;Metazoa | EP00117 Sycon ciliatum 14688 | 2,50E-92 | 1056 | XP 026029230.1 | protein transport protein Sec16A isoform X1 [Astatotilapia calliptera] >gi\|1468945480\|re | 2285 | 1,89E-42 | 181,415 | 40,25157233 |
| EP00615 | Sar;Stramenopiles;Gyrista;Ochrophyta;Dictyochoph | EP00615 Pteridomonas danica 15579 | 2,60E-92 | 892 | XP 008890960.1 | hypothetical protein PPTG 01118 [Phytophthora parasitica INRA-310] | 1621 | 1,25E-37 | 164,851 | 40,625 |
| EP00039 | Obazoa;Opisthokonta;Holozoa;Choanozoa;Choanof | EP00039 Didymoeca costata 12426 | 4,00E-92 | 969 | XP 010661318.1 | PREDICTED: protein transport protein SEC16B homolog [Vitis vinifera] >gi\|1104657609\|re | 1400 | 6,66E-34 | 152,91 | 46,13733906 |
| EP00264 | Archaeplastida;Chloroplastida;Streptophyta;"core s | EP00264 Hordeum vulgare 157073 | 4,30E-92 | 588 | XP 020180505.1 | protein transport protein SEC16B homolog [Aegilops tauschii subsp. tauschii] | 1647 | 0 | 1072,77 | 93,03904924 |
| EP00102 | Obazoa;Opisthokonta;Holozoa;Choanozoa;Metazoa | EP00102 Acyrthosiphon pisum 7019 | 1,10E-91 | 1435 | XP 003241201.1 | protein transport protein Sec16B isoform X1 [Acyrthosiphon pisum] | 1435 | 0 | 2979,12 | 100 |
| EP00578 | Sar;Stramenopiles;Gyrista;Ochrophyta;"BD clade";[ | EP00578 Thalassiosira sp NH16 29979 | 1,40E-91 | 1757 | XP 002183614.1 | predicted protein [Phaeodactylum tricornutum CCAP 1055/1] | 2083 | 5,07E-88 | 331,643 | 48,03240741 |
| EP00698 | Malawimonadidae;Gefionella;Gefionella okellyi;stra | EP00698 Gefionella okellyi 3275 | 2,10E-91 | 1259 | XP 020240850.1 | protein transport protein SEC16B homolog [Asparagus officinalis] | 1486 | 1,92E-41 | 178,718 | 46,72657253 |
| EP00523 | Sar;Stramenopiles;Gyrista;Ochrophyta;"BD clade";[ | EP00523 Amphiprora sp CCMP467 16974 | 2,60E-91 | 2374 | XP 002183614.1 | predicted protein [Phaeodactylum tricornutum CCAP 1055/1] | 2083 | 2,22E-109 | 402,519 | 49,50842697 |
| EP00241 | Archaeplastida;Chloroplastida;Chlorophyta;Pyramir | EP00241 Pyramimonas parkeae 17541 | 2,80E-91 | 953 | XP 010542224.1 | PREDICTED: protein transport protein SEC16B homolog [Tarenaya hassleriana] | 1321 | 2,06E-75 | 281,182 | 48,61111111 |
| EP00751 | Sar;Stramenopiles;Gyrista;Ochrophyta;"BD clade";[ | EP00751 Fragilariopsis kerguelensis 13616 | 5,50E-91 | 2276 | XP 002183614.1 | predicted protein [Phaeodactylum tricornutum CCAP 1055/1] | 2083 | 7,71E-118 | 429,098 | 50,60606061 |
| EP00698 | Malawimonadidae;Gefionella;Gefionella okellyi;stra | EP00698 Gefionella okellyi 3278 | 6,10E-91 | 1362 | XP 009791559.1 | PREDICTED: uncharacterized protein LOC104238783 [Nicotiana sylvestris] | 1515 | 4,45E-41 | 177,563 | 48,375 |
| EP00192 | Archaeplastida;Chloroplastida;Chlorophyta;"core ch | EP00192 Tetraselmis astigmatica 2307 | 1,00E-90 | 997 | XP 024373338.1 | protein transport protein SEC16A homolog isoform X1 [Physcomitrella patens] >gi\|13738 | 1444 | 4,06E-60 | 236,113 | 42,64150943 |
| EP00004 | CRuMs;Rigifilida;Rigifila;Rigifila ramosa;strain CCAF | EP00004 Rigifila ramosa 22578 | 1,10E-90 | 1232 | XP 004344939.1 | hypothetical protein ACA1 285550 [Acanthamoeba castellanii str. Neff] | 1515 | 3,48E-42 | 181,03 | 45,30612245 |
| EP00002 | CRuMs;Collodictyonidae;Diphylleia;Diphylleia rotan | EP00002 Diphylleia rotans 22695 | 1,10E-90 | 1052 | XP 004344939.1 | hypothetical protein ACA1 285550 [Acanthamoeba castellanii str. Neff] | 1515 | 9,94E-48 | 197,978 | 35,75221239 |
| EP00152 | Obazoa;Opisthokonta;Nucletmycea;Fungi;"core Fur | EP00152 Rhizopus delemar 3508 | 1,60E-90 | 571 | XP 023461952.1 | hypothetical protein RHIMIDRAFT 247874 [Rhizopus microsporus ATCC 52813] | 612 | 0 | 666,766 | 75,52816901 |
| EP00002 | CRuMs;Collodictyonidae;Diphylleia;Diphylleia rotan | EP00002 Diphylleia rotans 22688 | 1,60E-90 | 1890 | XP 004344939.1 | hypothetical protein ACA1 285550 [Acanthamoeba castellanii str. Neff] | 1515 | 5,92E-47 | 197,978 | 35,53191489 |
| EP00265 | Archaeplastida;Chloroplastida;Streptophyta;"core s | EP00265 Oryza sativa 13139 | 1,80E-90 | 577 | XP 015643544.1 | protein transport protein SEC16B homolog isoform X1 [Oryza sativa Japonica Group] | 1483 | 0 | 1189,87 | 100 |
| EP00698 | Malawimonadidae;Gefionella;Gefionella okellyi;stra | EP00698 Gefionella okellyi 3272 | 2,50E-90 | 1321 | XP 009791559.1 | PREDICTED: uncharacterized protein LOC104238783 [Nicotiana sylvestris] | 1515 | 5,81E-41 | 177,178 | 48,5 |
| EP00464 | Sar;Rhizaria;Cercozoa;Chlorarachnea;"core chlorara | EP00464 Amorphochlora amoebiformis 9576 | 2,60E-90 | 1736 | XP 004344939.1 | hypothetical protein ACA1 285550 [Acanthamoeba castellanii str. Neff] | 1515 | 2,37E-40 | 176,022 | 42,85714286 |
| EP00124 | Obazoa;Opisthokonta;Holozoa;Ichthyosporea;Ichthy | EP00124 Ichthyophonus hoferi 1710 | 3,90E-90 | 1987 | XP 003288899.1 | hypothetical protein DICPUDRAFT 98175 [Dictyostelium purpureum] | 1854 | 2,82E-34 | 156,762 | 40 |
| EP00053 | Obazoa;Opisthokonta;Holozoa;Choanozoa;Choanof | EP00053 Salpingoeca punica 15774 | 4,00E-90 | 485 | XP 013096820.1 | PREDICTED: uncharacterized protein LOC106080064 isoform X8 [Biomphalaria glabrata] | 1837 | 9,33E-43 | 173,326 | 47,18309859 |
| EP00526 | Sar;Stramenopiles;Gyrista;Ochrophyta;"BD clade";[ | EP00526 Cylindrotheca closterium 4161 | 4,40E-90 | 4255 | XP 002183614.1 | predicted protein [Phaeodactylum tricornutum CCAP 1055/1] | 2083 | 9,48E-108 | 399,053 | 50,75757576 |
| EP00002 | CRuMs;Collodictyonidae;Diphylleia;Diphylleia rotan | EP00002 Diphylleia rotans 22691 | 4,90E-90 | 1890 | XP 004344939.1 | hypothetical protein ACA1 285550 [Acanthamoeba castellanii str. Neff] | 1515 | 6,39E-47 | 197,593 | 35,53191489 |
| EP00698 | Malawimonadidae;Gefionella;Gefionella okellyi;stra | EP00698 Gefionella okellyi 3273 | 8,30E-90 | 1367 | XP 020240850.1 | protein transport protein SEC16B homolog [Asparagus officinalis] | 1486 | 9,55E-42 | 179,874 | 46,72657253 |
| EP00002 | CRuMs;Collodictyonidae;Diphylleia;Diphylleia rotan | EP00002 Diphylleia rotans 22690 | 1,10E-89 | 1422 | XP 004344939.1 | hypothetical protein ACA1 285550 [Acanthamoeba castellanii str. Neff] | 1515 | 1,62E-46 | 195,667 | 35,75221239 |
| EP00469 | Sar;Rhizaria;Cercozoa;Chlorarachnea;"core chlorara | EP00469 Lotharella globosa 3753 | 1,30E-89 | 1869 | XP 004344939.1 | hypothetical protein ACA1 285550 [Acanthamoeba castellanii str. Neff] | 1515 | 7,16E-51 | 210,69 | 46,43584521 |
| EP00206 | Archaeplastida;Chloroplastida;Chlorophyta;"core ch | EP00206 Coccomyxa subellipsoidea 6955 | 3,40E-89 | 1231 | XP 005649504.1 | hypothetical protein COCSUDRAFT 83652 [Coccomyxa subellipsoidea C-169] | 1231 | 0 | 2426,74 | 100 |
| EP00036 | Obazoa;Opisthokonta;Holozoa;Choanozoa;Choanof | EP00036 Acanthoeca sp 10tr 15891 | 4,00E-89 | 1620 | XP 004344939.1 | hypothetical protein ACA1 285550 [Acanthamoeba castellanii str. Neff] | 1515 | 3,11E-34 | 155,992 | 43,30143541 |
| EP00246 | Archaeplastida;Chloroplastida;Streptophyta;"core s | EP00246 Nothoceros aenigmaticus 6214 | 4,70E-89 | 542 | XP 019423445.1 | PREDICTED: protein transport protein SEC16B homolog [Lupinus angustifolius] >gi\|11174 | 1420 | 4,87E-123 | 402,905 | 56,82819383 |
| EP00150 | Obazoa;Opisthokonta;Nucletmycea;Fungi;"core Fur | EP00150 Ustilago maydis 3112 | 4,90E-89 | 2222 | XP 011389100.1 | hypothetical protein UMAG 02736 [Ustilago maydis 521] | 2222 | 0 | 4487,18 | 100 |
| EP00594 | Sar;Stramenopiles;Gyrista;Ochrophyta;"BD clade";[ | EP00594 Rhizosolenia setigera 9414 | 6,20E-89 | 708 | XP 002183614.1 | predicted protein [Phaeodactylum tricornutum CCAP 1055/1] | 2083 | 5,64E-77 | 281,182 | 45,20547945 |
| EP00698 | Malawimonadidae;Gefionella;Gefionella okellyi;stra | EP00698 Gefionella okellyi 3277 | 1,20E-88 | 1381 | XP 020240850.1 | protein transport protein SEC16B homolog [Asparagus officinalis] | 1486 | 8,32E-42 | 180,259 | 46,72657253 |
| EP00464 | Sar;Rhizaria;Cercozoa;Chlorarachnea;"core chlorara | EP00464 Amorphochlora amoebiformis 9469 | 9,30E-88 | 2391 | XP 004344939.1 | hypothetical protein ACA1 285550 [Acanthamoeba castellanii str. Neff] | 1515 | 1,86E-40 | 177,178 | 42,39130435 |
| EP00037 | Obazoa;Opisthokonta;Holozoa;Choanozoa;Choanof | EP00037 Helgoeca nana 11082 | 9,50E-88 | 1614 | XP 004344939.1 | hypothetical protein ACA1 285550 [Acanthamoeba castellanii str. Neff] | 1515 | 4,17E-33 | 152,14 | 41,33333333 |
| EP00464 | Sar;Rhizaria;Cercozoa;Chlorarachnea;"core chlorara | EP00464 Amorphochlora amoebiformis 9554 | 1,40E-87 | 2316 | XP 004344939.1 | hypothetical protein ACA1 285550 [Acanthamoeba castellanii str. Neff] | 1515 | 1,35E-40 | 177,563 | 42,39130435 |
| EP00152 | Obazoa;Opisthokonta;Nucletmycea;Fungi;"core Fur | EP00152 Rhizopus delemar 5140 | 1,50E-87 | 2576 | XP 023470420.1 | actin-binding FH2 [Rhizopus microsporus ATCC 52813] | 1383 | 0 | 822,772 | 84,99061914 |
| EP00138 | Obazoa;Opisthokonta;Nucletmycea;Fungi;"core Fur | EP00138 Coccidioides posadasii 7105 | 4,30E-87 | 1295 | XP 003072104.1 | hypothetical protein CPC735 012770 [Coccidioides posadasii C735 delta SOWgp] | 1295 | 0 | 2647,85 | 100 |
| EP00535 | Sar;Stramenopiles;Gyrista;Ochrophyta;"BD clade";[ | EP00535 Pseudo-nitzschia heimii 2688 | 5,00E-87 | 1479 | XP 002183614.1 | predicted protein [Phaeodactylum tricornutum CCAP 1055/1] | 2083 | 1,52E-99 | 365,54 | 47,00260643 |
| EP00002 | CRuMs;Collodictyonidae;Diphylleia;Diphylleia rotan | EP00002 Diphylleia rotans 22694 | 6,40E-87 | 1771 | XP 004344939.1 | hypothetical protein ACA1 285550 [Acanthamoeba castellanii str. Neff] | 1515 | 1,25E-46 | 196,823 | 35,53191489 |
| EP00738 | Archaeplastida;Chloroplastida;Chlorophyta;Chlorop | EP00738 Chloropicon primus 7426 | 9,40E-87 | 1311 | XP 024010987.1 | protein transport protein SEC16B homolog [Eutrema salsugineum] | 1221 | 4,64E-61 | 241,121 | 47,72209567 |
| EP00036 | Obazoa;Opisthokonta;Holozoa;Choanozoa;Choanof | EP00036 Acanthoeca sp 10tr 2923 | 1,40E-86 | 1342 | XP 004344939.1 | hypothetical protein ACA1 285550 [Acanthamoeba castellanii str. Neff] | 1515 | 6,13E-33 | 151,369 | 41,59292035 |
| EP00751 | Sar;Stramenopiles;Gyrista;Ochrophyta;"BD clade";[ | EP00751 Fragilariopsis kerguelensis 47168 | 2,20E-86 | 2165 | XP 002183614.1 | predicted protein [Phaeodactylum tricornutum CCAP 1055/1] | 2083 | 8,97E-88 | 332,028 | 51,75 |
| EP00117 | Obazoa;Opisthokonta;Holozoa;Choanozoa;Metazoa | EP00117 Sycon ciliatum 12290 | 2,70E-86 | 1069 | XP 026029235.1 | protein transport protein Sec16A isoform X3 [Astatotilapia calliptera] | 2270 | 1,92E-42 | 181,415 | 40,25157233 |
| EP00230 | Archaeplastida;Chloroplastida;Chlorophyta;Mamiel | EP00230 Micromonas polaris 5144 | 5,10E-86 | 2012 | XP 002509249.1 | predicted protein [Micromonas commoda] | 1945 | 1,48E-91 | 343,584 | 57,64587525 |
| EP00687 | Discoba;Heterolobosea;Tetramitia;Eutetramitia;"PS | EP00687 Percolomonas cosmopolitus 5698 | 5,50E-86 | 1357 | XP 012756363.1 | hypothetical protein SAMD00019534 027760 [Acytostelium subglobosum LB1] | 1470 | 9,34E-42 | 179,874 | 43,90756303 |
| EP00618 | Sar;Stramenopiles;Gyrista;Ochrophyta;Dictyochoph | EP00618 Florenciella parvula 941 | 5,90E-86 | 998 | XP 008890960.1 | hypothetical protein PPTG 01118 [Phytophthora parasitica INRA-310] | 1621 | 4,15E-48 | 198,749 | 46,91780822 |
| EP00282 | Cryptista;Cryptophyceae;Cryptomonadales;"lineage | EP00282 Hemiselmis andersenii 37378 | 6,30E-86 | 651 | XP 004511687.1 | protein transport protein SEC16B homolog [Cicer arietinum] >gi\|1535901408\|ref\|XP 027 | 1424 | 4,26E-40 | 169,088 | 44,51097804 |
| EP00041 | Obazoa;Opisthokonta;Holozoa;Choanozoa;Choanof | EP00041 Stephanoeca diplocostata 35234 | 6,90E-86 | 1331 | XP 004344939.1 | hypothetical protein ACA1 285550 [Acanthamoeba castellanii str. Neff] | 1515 | 7,78E-33 | 150,984 | 40,10840108 |
| EP00648 | Sar;Stramenopiles;Gyrista;Peronosporomycetes;"cc | EP00648 Phytopythium vexans 4166 | 9,50E-86 | 2870 | XP 002908536.1 | COPII coat assembly protein SEC16, putative [Phytophthora infestans T30-4] | 2641 | 0 | 774,237 | 64,72457627 |
| EP00521 | Sar;Stramenopiles;Gyrista;Ochrophyta;"BD clade";[ | EP00521 Asterionellopsis glacialis 9836 | 9,60E-86 | 1453 | XP 002183614.1 | predicted protein [Phaeodactylum tricornutum CCAP 1055/1] | 2083 | 7,99E-111 | 399,438 | 46,67318982 |
| EP00587 | Sar;Stramenopiles;Gyrista;Ochrophyta;"BD clade";[ | EP00587 Corethron hystrix 19 | 2,20E-85 | 878 | XP 002183614.1 | predicted protein [Phaeodactylum tricornutum CCAP 1055/1] | 2083 | 1,41E-34 | 155,221 | 40,95238095 |
| EP00681 | Discoba;Euglenozoa;Kinetoplastea;Metakinetoplast | EP00681 Trypanosoma brucei 1122 | 7,80E-85 | 1387 | XP 011772368.1 | hypothetical protein, conserved [Trypanosoma brucei gambiense DAL972] | 1387 | 0 | 2812,71 | 100 |
| EP00542 | Sar;Stramenopiles;Gyrista;Ochrophyta;"BD clade";[ | EP00542 Grammatophora oceanica 11580 | 1,10E-84 | 1711 | XP 002183614.1 | predicted protein [Phaeodactylum tricornutum CCAP 1055/1] | 2083 | 3,60E-120 | 431,795 | 46,8892261 |
| EP00339 | Sar;Alveolata;Ciliophora;Intramacronucleata;"CON: | EP00339 Tiarina fusus 12555 | 1,20E-84 | 1867 | XP 002183614.1 | predicted protein [Phaeodactylum tricornutum CCAP 1055/1] | 2083 | 2,24E-77 | 297,745 | 44,65408805 |
| EP00442 | Sar;Alveolata;Myzozoa;"DP clade";Dinoflagellata;D | EP00442 Polarella glacialis 908 | 5,80E-84 | 1036 | XP 009223994.1 | hypothetical protein GGTG 07899 [Gaeumannomyces tritici R3-111a-1] | 1936 | 3,31E-18 | 102,449 | 43,5483871 |
| EP00553 | Sar;Stramenopiles;Gyrista;Ochrophyta;"BD clade";[ | EP00553 Chaetoceros curvisetus 10068 | 6,30E-84 | 595 | XP 002183614.1 | predicted protein [Phaeodactylum tricornutum CCAP 1055/1] | 2083 | 1,18E-44 | 182,185 | 45,39877301 |
| EP00522 | Sar;Stramenopiles;Gyrista;Ochrophyta;"BD clade";[ | EP00522 Amphiprora paludosa 2344 | 7,70E-84 | 2857 | XP 002183614.1 | predicted protein [Phaeodactylum tricornutum CCAP 1055/1] | 2083 | 3,50E-144 | 513,072 | 50,58179329 |
| EP00442 | Sar;Alveolata;Myzozoa;"DP clade";Dinoflagellata;D | EP00442 Polarella glacialis 828 | 9,70E-84 | 1043 | XP 009223994.1 | hypothetical protein GGTG 07899 [Gaeumannomyces tritici R3-111a-1] | 1936 | 1,69E-17 | 100,138 | 37,13355049 |
| EP00035 | Obazoa;Opisthokonta;Holozoa;Choanozoa;Choanof | EP00035 Acanthoeca spectabilis 22936 | 1,50E-83 | 1583 | XP 002109984.1 | hypothetical protein TRIADDRAFT 53450 [Trichoplax adhaerens] | 1642 | 2,25E-32 | 149,828 | 40,10840108 |
| EP00158 | Obazoa;Opisthokonta;Nucletmycea;Fungi;Aphelide | EP00158 Paraphelidium tribonemae 8368 | 2,20E-83 | 1236 | XP 002842484.1 | hypothetical protein [Tuber melanosporum Mel28] | 1599 | 2,22E-39 | 171,785 | 45,01108647 |
| EP00469 | Sar;Rhizaria;Cercozoa;Chlorarachnea;"core chlorara | EP00469 Lotharella globosa 6487 | 2,60E-83 | 2345 | XP 004344939.1 | hypothetical protein ACA1 285550 [Acanthamoeba castellanii str. Neff] | 1515 | 6,07E-54 | 221,09 | 46,32653061 |
| EP00339 | Sar;Alveolata;Ciliophora;Intramacronucleata;"CON: | EP00339 Tiarina fusus 12537 | 2,80E-83 | 2828 | XP 002183614.1 | predicted protein [Phaeodactylum tricornutum CCAP 1055/1] | 2083 | 2,98E-101 | 377,096 | 48,71611983 |
| EP00390 | Sar;Alveolata;Myzozoa;"AC clade";Colpodellida;Vitr | EP00390 Vitrella brassicaformis 22980 | 3,20E-83 | 1136 | XP 004344939.1 | hypothetical protein ACA1 285550 [Acanthamoeba castellanii str. Neff] | 1515 | 1,39E-27 | 133,265 | 40,36885246 |
| EP00442 | Sar;Alveolata;Myzozoa;"DP clade";Dinoflagellata;D | EP00442 Polarella glacialis 906 | 3,60E-83 | 1036 | XP 009223994.1 | hypothetical protein GGTG 07899 [Gaeumannomyces tritici R3-111a-1] | 1936 | 2,96E-18 | 102,834 | 43,5483871 |
| EP00048 | Obazoa;Opisthokonta;Holozoa;Choanozoa;Choanof | EP00048 Salpingoeca helianthica 13697 | 7,40E-83 | 555 | XP 004344939.1 | hypothetical protein ACA1 285550 [Acanthamoeba castellanii str. Neff] | 1515 | 1,52E-37 | 159,458 | 45,93908629 |
| EP00762 | Discoba;Jakobida;Andalucina;Andaluciidae;Andaluc | EP00762 Andalucia godoyi 3862 | 1,00E-82 | 1405 | XP 008871247.1 | hypothetical protein H310 07614 [Aphanomyces invadans] | 2510 | 2,06E-25 | 127,102 | 47,07207207 |
| EP00158 | Obazoa;Opisthokonta;Nucletmycea;Fungi;Aphelide | EP00158 Paraphelidium tribonemae 8367 | 1,10E-82 | 1236 | XP 002842484.1 | hypothetical protein [Tuber melanosporum Mel28] | 1599 | 1,21E-39 | 172,94 | 45,45454545 |
| EP00410 | Sar;Alveolata;Myzozoa;"DP clade";Dinoflagellata;D | EP00410 Alexandrium tamarense 65833 | 1,30E-82 | 1258 | XP 008890960.1 | hypothetical protein PPTG 01118 [Phytophthora parasitica INRA-310] | 1621 | 2,16E-20 | 110,153 | 43,70546318 |
| EP00760 | Discoba;Euglenozoa;Kinetoplastea;Prokinetoplastin | EP00760 Prokinetoplastina sp PhM-4 10195 | 1,30E-82 | 1087 | XP 029230831.1 | uncharacterized protein Tco025E 02121 [Trypanosoma conorhini] | 1464 | 8,16E-24 | 120,939 | 45,28301887 |
| EP00563 | Sar;Stramenopiles;Gyrista;Ochrophyta;"BD clade";[ | EP00563 Minutocellus polymorphus 13986 | 1,60E-82 | 1566 | XP 002183614.1 | predicted protein [Phaeodactylum tricornutum CCAP 1055/1] | 2083 | 2,68E-67 | 263,848 | 47,99154334 |
| EP00441 | Sar;Alveolata;Myzozoa;"DP clade";Dinoflagellata;D | EP00441 Pelagodinium beii 43776 | 4,60E-82 | 1185 | XP 009223994.1 | hypothetical protein GGTG 07899 [Gaeumannomyces tritici R3-111a-1] | 1936 | 1,41E-21 | 114,005 | 42,34527687 |
| EP00117 | Obazoa;Opisthokonta;Holozoa;Choanozoa;Metazoa | EP00117 Sycon ciliatum 45910 | 4,90E-82 | 1082 | XP 026029235.1 | protein transport protein Sec16A isoform X3 [Astatotilapia calliptera] | 2270 | 1,97E-42 | 181,415 | 40,25157233 |
| EP00760 | Discoba;Euglenozoa;Kinetoplastea;Prokinetoplastin | EP00760 Prokinetoplastina sp PhM-4 10193 | 8,70E-82 | 1063 | XP 029230831.1 | uncharacterized protein Tco025E 02121 [Trypanosoma conorhini] | 1464 | 9,82E-24 | 120,553 | 45,28301887 |
| EP00004 | CRuMs;Rigifilida;Rigifila;Rigifila ramosa;strain CCAF | EP00004 Rigifila ramosa 22575 | 7,00E-81 | 1042 | XP 004344939.1 | hypothetical protein ACA1 285550 [Acanthamoeba castellanii str. Neff] | 1515 | 2,70E-41 | 177,178 | 45,30612245 |
| EP00339 | Sar;Alveolata;Ciliophora;Intramacronucleata;"CON: | EP00339 Tiarina fusus 12649 | 1,60E-80 | 2441 | XP 002183614.1 | predicted protein [Phaeodactylum tricornutum CCAP 1055/1] | 2083 | 6,82E-102 | 378,252 | 48,68139701 |
| EP00418 | Sar;Alveolata;Myzozoa;"DP clade";Dinoflagellata;D | EP00418 Pyrodinium bahamense 43192 | 1,80E-80 | 827 | XP 002370242.1 | hypothetical protein TGME49 294730 [Toxoplasma gondii ME49] | 2142 | 5,64E-17 | 97,8265 | 39,44636678 |
| EP00328 | Haptista;Haptophyta;Prymnesiophyceae;Prymnesia | EP00328 Prymnesium polylepis 27136 | 2,50E-80 | 636 | XP 004344939.1 | hypothetical protein ACA1 285550 [Acanthamoeba castellanii str. Neff] | 1515 | 6,64E-41 | 171,4 | 47,5 |
| EP00770 | Metamonada;Preaxostyla;Oxymonadida;Monocerc | EP00770 Monocercomonoides exilis 652 | 2,80E-80 | 1906 | ***no hit found*** | | | | | |
| EP00439 | Sar;Alveolata;Myzozoa;"DP clade";Dinoflagellata;D | EP00439 Symbiodinium sp Y106 84093 | 3,60E-80 | 1480 | XP 028465267.1 | Pkinase-domain-containing protein [Sodiomyces alkalinus F11] | 395 | 6,99E-30 | 134,42 | 46,17737003 |
| EP00038 | Obazoa;Opisthokonta;Holozoa;Choanozoa;Choanof | EP00038 Savillea parva 20398 | 4,80E-80 | 1806 | XP 021349045.1 | uncharacterized protein LOC110447583 isoform X4 [Mizuhopecten yessoensis] | 2120 | 1,85E-33 | 154,066 | 35,65217391 |
| EP00439 | Sar;Alveolata;Myzozoa;"DP clade";Dinoflagellata;D | EP00439 Symbiodinium sp Y106 83819 | 5,30E-80 | 1484 | XP 028465267.1 | Pkinase-domain-containing protein [Sodiomyces alkalinus F11] | 395 | 6,95E-30 | 134,42 | 46,17737003 |
| EP00365 | Sar;Alveolata;Myzozoa;"AC clade";Apicomplexa;"cc | EP00365 Plasmodium falciparum 524 | 5,40E-80 | 1749 | XP 001347877.1 | conserved Plasmodium protein, unknown function [Plasmodium falciparum 3D7] | 1749 | 0 | 3400,91 | 100 |
| EP00234 | Archaeplastida;Chloroplastida;Palmophyllophyceae | EP00234 Prasinoderma coloniale 1870 | 6,70E-80 | 2008 | XP 003055453.1 | predicted protein [Micromonas pusilla CCMP1545] | 294 | 7,84E-50 | 189,889 | 50,76923077 |
| EP00443 | Sar;Alveolata;Myzozoa;"DP clade";Dinoflagellata;D | EP00443 Scrippsiella trochoidea 49072 | 7,60E-80 | 1079 | XP 012756363.1 | hypothetical protein SAMD00019534 027760 [Acytostelium subglobosum LB1] | 1470 | 8,70E-21 | 110,923 | 39,12087912 |
| EP00210 | Archaeplastida;Chloroplastida;Chlorophyta;"core ch | EP00210 Caulerpa lentillifera 2975 | 1,00E-79 | 940 | XP 024373338.1 | protein transport protein SEC16A homolog isoform X1 [Physcomitrella patens] >gi\|13738 | 1444 | 1,94E-45 | 189,504 | 39,81308411 |
| EP00339 | Sar;Alveolata;Ciliophora;Intramacronucleata;"CON: | EP00339 Tiarina fusus 12608 | 1,30E-79 | 2138 | XP 002183614.1 | predicted protein [Phaeodactylum tricornutum CCAP 1055/1] | 2083 | 4,84E-101 | 374,785 | 48,64479315 |
| EP00269 | Archaeplastida;Chloroplastida;Streptophyta;"core s | EP00269 Selaginella moellendorffii 22542 | 4,00E-79 | 381 | XP 024523188.1 | protein transport protein SEC16A homolog [Selaginella moellendorffii] | 381 | 0 | 790,415 | 100 |
| EP00529 | Sar;Stramenopiles;Gyrista;Ochrophyta;"BD clade";[ | EP00529 Nitzschia sp RCC80 8196 | 7,70E-79 | 1169 | XP 002183614.1 | predicted protein [Phaeodactylum tricornutum CCAP 1055/1] | 2083 | 2,18E-68 | 264,618 | 49,72617744 |

| | | | | | | | | | | |
|---|---|---|---|---|---|---|---|---|---|---|
| EP00752 | Sar;Stramenopiles;Gyrista;Ochrophyta;"PX clade";P | EP00752 Nemacystus decipiens 12780 | 1,10E-78 | 3926 | XP 005854093.1 | hypothetical protein NGA 0496400 [Nannochloropsis gaditana CCMP526] | 487 | 0 | 835,099 | 85,27472527 |
| EP00012 | Amoebozoa;Discosea;Flabellinia;"core flabellinids"; | EP00012 Vannella robusta 4126 | 1,10E-78 | 851 | XP 003288899.1 | hypothetical protein DICPUDRAFT 98175 [Dictyostelium purpureum] | 1854 | 5,52E-35 | 155,992 | 41,86046512 |
| EP00474 | Sar;Rhizaria;Endomyxa;"phytorhiza";Phytomyxea;" | EP00474 Spongospora subterranea 9655 | 2,10E-78 | 1008 | XP 004344939.1 | hypothetical protein ACA1 285550 [Acanthamoeba castellanii str. Neff] | 1515 | 3,19E-30 | 141,354 | 46,15384615 |
| EP00468 | Sar;Rhizaria;Cercozoa;Chlorarachnea;"core chlorara | EP00468 Gymnochlora sp CCMP2014 12696 | 3,20E-78 | 2259 | XP 004344939.1 | hypothetical protein ACA1 285550 [Acanthamoeba castellanii str. Neff] | 1515 | 8,96E-50 | 207,608 | 44,95798319 |
| EP00142 | Obazoa;Opisthokonta;Nucletmycea;Fungi;"core Fur | EP00142 Kluyveromyces lactis 3848 | 3,90E-78 | 2082 | XP 455029.1 | uncharacterized protein KLLA0 E23915g [Kluyveromyces lactis] | 2082 | 0 | 4228,71 | 100 |
| EP00755 | Discoba;Euglenozoa;Diplonemea;Diplonemidae;Sul | EP00755 Sulcionema specki 17032 | 7,50E-78 | 665 | XP 012207226.1 | hypothetical protein SPRG 12070 [Saprolegnia parasitica CBS 223.65] | 2601 | 4,83E-09 | 71,633 | 40,36697248 |
| EP00435 | Sar;Alveolata;Myzozoa;"DP clade";Dinoflagellata;D | EP00435 Cladocopium sp Y103 11019 | 8,50E-78 | 1008 | XP 016606650.1 | hypothetical protein SPPG 06291 [Spizellomyces punctatus DAOM BR117] | 2211 | 4,88E-19 | 105,145 | 38,91752577 |
| EP00529 | Sar;Stramenopiles;Gyrista;Ochrophyta;"BD clade";[ | EP00529 Nitzschia sp RCC80 6367 | 1,80E-77 | 1507 | XP 002183614.1 | predicted protein [Phaeodactylum tricornutum CCAP 1055/1] | 2083 | 3,16E-85 | 320,857 | 52,36309077 |
| EP00235 | Archaeplastida;Chloroplastida;Palmophyllophyceae | EP00235 Prasinoderma singulare 380 | 1,90E-77 | 1544 | XP 021977759.1 | protein transport protein SEC16B homolog isoform X2 [Helianthus annuus] | 1389 | 2,11E-40 | 175,637 | 44,26229508 |
| EP00529 | Sar;Stramenopiles;Gyrista;Ochrophyta;"BD clade";[ | EP00529 Nitzschia sp RCC80 9011 | 2,40E-77 | 2522 | XP 002183614.1 | predicted protein [Phaeodactylum tricornutum CCAP 1055/1] | 2083 | 3,56E-95 | 356,681 | 52,13675214 |
| EP00468 | Sar;Rhizaria;Cercozoa;Chlorarachnea;"core chlorara | EP00468 Gymnochlora sp CCMP2014 15293 | 3,60E-77 | 2279 | XP 004344939.1 | hypothetical protein ACA1 285550 [Acanthamoeba castellanii str. Neff] | 1515 | 1,07E-49 | 207,223 | 44,95798319 |
| EP00442 | Sar;Alveolata;Myzozoa;"DP clade";Dinoflagellata;D | EP00442 Polarella glacialis 1041 | 4,40E-77 | 1069 | XP 009223994.1 | hypothetical protein GGTG 07899 [Gaeumannomyces tritici R3-111a-1] | 1936 | 8,09E-18 | 101,293 | 37,25490196 |
| EP00435 | Sar;Alveolata;Myzozoa;"DP clade";Dinoflagellata;D | EP00435 Cladocopium sp Y103 9041 | 6,60E-77 | 891 | XP 016606650.1 | hypothetical protein SPPG 06291 [Spizellomyces punctatus DAOM BR117] | 2211 | 1,06E-18 | 103,99 | 38,97058824 |
| EP00111 | Obazoa;Opisthokonta;Holozoa;Choanozoa;Metazoa | EP00111 Clytia hemisphaerica 22257 | 6,70E-77 | 2217 | XP 012566805.1 | PREDICTED: uncharacterized protein LOC100197480 isoform X2 [Hydra vulgaris] | 2407 | 3,18E-134 | 481,871 | 49,3877551 |
| EP00644 | Sar;Stramenopiles;Gyrista;Peronosporomycetes;"cc | EP00644 Phytophthora capsici 6892 | 9,00E-77 | 679 | XP 002908536.1 | COPII coat assembly protein SEC16, putative [Phytophthora infestans T30-4] | 2641 | 2,07E-149 | 490,73 | 82,13457077 |
| EP00697 | Hemimastigophora;Spironemidae;Spironema;strain | EP00697 Spironema sp BW2 10718 | 1,70E-76 | 458 | XP 004344939.1 | hypothetical protein ACA1 285550 [Acanthamoeba castellanii str. Neff] | 1515 | 1,21E-45 | 181,03 | 52,79187817 |
| EP00429 | Sar;Alveolata;Myzozoa;"DP clade";Dinoflagellata;D | EP00429 Kryptoperidinium foliaceum 9246 | 3,00E-76 | 950 | XP 012756363.1 | hypothetical protein SAMD00019534 027760 [Acytostelium subglobosum LB1] | 1470 | 3,27E-24 | 121,709 | 36,11738149 |
| EP00419 | Sar;Alveolata;Myzozoa;"DP clade";Dinoflagellata;D | EP00419 Tripos fusus 53165 | 1,10E-75 | 895 | XP 027040890.1 | protein transport protein Sec16A-like [Pocillopora damicornis] | 2396 | 1,89E-21 | 112,849 | 38,13747228 |
| EP00331 | Sar;Alveolata;Ciliophora;Intramacronucleata;"CON: | EP00331 Platyophrya macrostoma 4967 | 1,40E-75 | 523 | XP 029239091.1 | uncharacterized protein TraAM80 04123 [Trypanosoma rangeli] | 1446 | 6,99E-37 | 156,762 | 45,28301887 |
| EP00298 | Haptista;Centroplasthelida;Panacanthocystida;Acar | EP00298 Acanthocystis sp HF-20 5480 | 1,80E-75 | 435 | XP 012756363.1 | hypothetical protein SAMD00019534 027760 [Acytostelium subglobosum LB1] | 1470 | 2,23E-32 | 141,354 | 37,09198813 |
| EP00448 | Sar;Alveolata;Myzozoa;"DP clade";Dinoflagellata;D | EP00448 Togula jolla 7089 | 2,40E-75 | 1080 | XP 004344939.1 | hypothetical protein ACA1 285550 [Acanthamoeba castellanii str. Neff] | 1515 | 3,11E-20 | 108,997 | 41,55555556 |
| EP00473 | Sar;Rhizaria;Endomyxa;"phytorhiza";Phytomyxea;" | EP00473 Plasmodiophora brassicae 7402 | 2,80E-75 | 921 | XP 021349048.1 | uncharacterized protein LOC110447583 isoform X7 [Mizuhopecten yessoensis] | 2085 | 7,43E-28 | 133,65 | 44,9168207 |
| EP00432 | Sar;Alveolata;Myzozoa;"DP clade";Dinoflagellata;D | EP00432 Prorocentrum cordatum 28595 | 4,60E-75 | 741 | XP 016606650.1 | hypothetical protein SPPG 06291 [Spizellomyces punctatus DAOM BR117] | 2211 | 9,90E-21 | 109,768 | 46,0815047 |
| EP00418 | Sar;Alveolata;Myzozoa;"DP clade";Dinoflagellata;D | EP00418 Pyrodinium bahamense 7490 | 4,90E-75 | 822 | XP 002370242.1 | hypothetical protein TGME49 294730 [Toxoplasma gondii ME49] | 2142 | 3,76E-17 | 98,5969 | 39,44636678 |
| EP00613 | Sar;Stramenopiles;Gyrista;Ochrophyta;Dictyochoph | EP00613 Pedinella sp CCMP2098 49134 | 6,90E-75 | 1121 | XP 012207226.1 | hypothetical protein SPRG 12070 [Saprolegnia parasitica CBS 223.65] | 2601 | 5,17E-22 | 115,161 | 47,6673428 |
| EP00753 | Sar;Stramenopiles;Platysulcus;Platysulcus tardus;st | EP00753 Platysulcus tardus 15410 | 7,00E-75 | 535 | XP 004344939.1 | hypothetical protein ACA1 285550 [Acanthamoeba castellanii str. Neff] | 1515 | 5,23E-36 | 154,451 | 46,71361502 |
| EP00124 | Obazoa;Opisthokonta;Holozoa;Ichthyosporea;Ichthy | EP00124 Ichthyophonus hoferi 5476 | 1,40E-74 | 455 | XP 004344939.1 | hypothetical protein ACA1 285550 [Acanthamoeba castellanii str. Neff] | 1515 | 1,23E-43 | 175,252 | 45,20123839 |
| EP00393 | Sar;Alveolata;Myzozoa;"DP clade";Dinoflagellata;"c | EP00393 Amoebophrya sp ex-Akashiwo-sanguinea 87135 | 1,50E-74 | 1270 | XP 004344939.1 | hypothetical protein ACA1 285550 [Acanthamoeba castellanii str. Neff] | 1515 | 9,44E-25 | 124,405 | 41,81415929 |
| EP00448 | Sar;Alveolata;Myzozoa;"DP clade";Dinoflagellata;D | EP00448 Togula jolla 35128 | 1,80E-74 | 1199 | XP 004344939.1 | hypothetical protein ACA1 285550 [Acanthamoeba castellanii str. Neff] | 1515 | 1,32E-20 | 110,538 | 41,55555556 |
| EP00418 | Sar;Alveolata;Myzozoa;"DP clade";Dinoflagellata;D | EP00418 Pyrodinium bahamense 47895 | 2,80E-74 | 829 | XP 002370242.1 | hypothetical protein TGME49 294730 [Toxoplasma gondii ME49] | 2142 | 3,04E-17 | 98,9821 | 39,44636678 |
| EP00672 | Discoba;Euglenozoa;Kinetoplastea;Metakinetoplast | EP00672 Neobodo designis 4672 | 3,30E-74 | 1499 | XP 029239091.1 | uncharacterized protein TraAM80 04123 [Trypanosoma rangeli] | 1446 | 9,73E-24 | 121,324 | 43,49206349 |
| EP00468 | Sar;Rhizaria;Cercozoa;Chlorarachnea;"core chlorara | EP00468 Gymnochlora sp CCMP2014 6308 | 7,80E-74 | 1926 | XP 004344939.1 | hypothetical protein ACA1 285550 [Acanthamoeba castellanii str. Neff] | 1515 | 1,99E-49 | 206,068 | 44,95798319 |
| EP00759 | Discoba;Euglenozoa;Kinetoplastea;Prokinetoplastin | EP00759 Prokinetoplastina sp PhF-6 30777 | 9,80E-74 | 798 | XP 008890960.1 | hypothetical protein PPTG 01118 [Phytophthora parasitica INRA-310] | 1621 | 1,99E-19 | 105,916 | 42,00426439 |
| EP00230 | Archaeplastida;Chloroplastida;Chlorophyta;Mamiel | EP00230 Micromonas polaris 435 | 1,30E-73 | 1498 | XP 002509249.1 | predicted protein [Micromonas commoda] | 1945 | 1,94E-80 | 305,449 | 57,79467681 |
| EP00468 | Sar;Rhizaria;Cercozoa;Chlorarachnea;"core chlorara | EP00468 Gymnochlora sp CCMP2014 2352 | 1,40E-72 | 2203 | XP 004344939.1 | hypothetical protein ACA1 285550 [Acanthamoeba castellanii str. Neff] | 1515 | 3,86E-50 | 208,764 | 45,16806723 |
| EP00658 | Telonemia;Telonema;strain P-2 | EP00658 Telonema sp P-2 54010 | 1,90E-72 | 527 | XP 021304254.1 | protein transport protein SEC16A homolog isoform X1 [Sorghum bicolor] | 1891 | 2,91E-33 | 146,362 | 47,41784038 |
| EP00450 | Sar;Alveolata;Myzozoa;"DP clade";Dinoflagellata;N | EP00450 Noctiluca scintillans 34040 | 2,30E-72 | 1175 | XP 004344939.1 | hypothetical protein ACA1 285550 [Acanthamoeba castellanii str. Neff] | 1515 | 7,59E-20 | 108,227 | 46,35761589 |
| EP00303 | Haptista;Haptophyta;Pavlovales;Exanthemachrysis; | EP00303 Exanthemachrysis gayraliae 14359 | 3,10E-72 | 619 | XP 025811964.1 | protein transport protein SEC16A homolog isoform X1 [Panicum hallii] | 1891 | 1,35E-23 | 117,857 | 37,94871795 |
| EP00283 | Cryptista;Cryptophyceae;Cryptomonadales;"lineage | EP00283 Hemiselmis rufescens 8734 | 7,30E-72 | 251 | XP 012845178.1 | PREDICTED: protein transport protein SEC16B homolog [Erythranthe guttata] >gi\|848890( | 1514 | 5,91E-39 | 154,451 | 47,58064516 |
| EP00184 | Archaeplastida;"RR clade";Rhodophyta;Proteorhod | EP00184 Porphyridium aerugineum 586 | 1,90E-71 | 1829 | XP 005711041.1 | unnamed protein product [Chondrus crispus] | 2546 | 1,72E-20 | 111,309 | 39,25925926 |
| EP00178 | Archaeplastida;"RR clade";Rhodophyta;Eurhodophy | EP00178 Gracilaria changii 19066 | 2,00E-71 | 2948 | XP 005711041.1 | unnamed protein product [Chondrus crispus] | 2546 | 0 | 672,929 | 49,39965695 |
| EP00040 | Obazoa;Opisthokonta;Holozoa;Choanozoa;Choanof | EP00040 Diaphanoeca grandis 30923 | 2,70E-71 | 407 | XP 019860577.1 | PREDICTED: protein transport protein Sec16A-like [Amphimedon queenslandica] | 1105 | 7,49E-27 | 124,02 | 43,12977099 |
| EP00754 | Discoba;Euglenozoa;Diplonemea;Diplonemidae;Rhy | EP00754 Rhynchopus humris 10960 | 3,10E-71 | 1147 | ***no hit found*** | | | | | |
| EP00368 | Sar;Alveolata;Myzozoa;"AC clade";Apicomplexa;"cc | EP00368 Plasmodium yoelii 5750 | 4,10E-71 | 1316 | XP 730982.2 | uncharacterized protein PY17X 0930300 [Plasmodium yoelii] | 1316 | 0 | 2612,41 | 100 |
| EP00503 | Sar;Stramenopiles;Bigyra;Opalozoa;Opalinata;Blast | EP00503 Blastocystis sp subtype4 3447 | 6,00E-71 | 1160 | XP 014528041.1 | hypothetical protein JH06 3685 [Blastocystis sp. subtype 4] | 1160 | 0 | 2328,9 | 100 |
| EP00730 | Obazoa;Opisthokonta;Holozoa;Choanozoa;Choanof | EP00730 Choanoeca flexa 4969 | 1,00E-70 | 1453 | XP 004992030.1 | hypothetical protein PTSG 12463 [Salpingoeca rosetta] | 2149 | 8,35E-21 | 112,079 | 36,55172414 |
| EP00529 | Sar;Stramenopiles;Gyrista;Ochrophyta;"BD clade";[ | EP00529 Nitzschia sp RCC80 41931 | 1,10E-70 | 1428 | XP 002183614.1 | predicted protein [Phaeodactylum tricornutum CCAP 1055/1] | 2083 | 2,04E-82 | 311,227 | 53,11004785 |
| EP00444 | Sar;Alveolata;Myzozoa;"DP clade";Dinoflagellata;D | EP00444 Apocalathium aciculiferum 33019 | 1,40E-70 | 980 | XP 012756363.1 | hypothetical protein SAMD00019534 027760 [Acytostelium subglobosum LB1] | 1470 | 1,93E-21 | 112,849 | 43,24894515 |
| EP00435 | Sar;Alveolata;Myzozoa;"DP clade";Dinoflagellata;D | EP00435 Cladocopium sp Y103 11257 | 2,50E-70 | 1039 | XP 016606650.1 | hypothetical protein SPPG 06291 [Spizellomyces punctatus DAOM BR117] | 2211 | 9,03E-19 | 104,375 | 38,91752577 |
| EP00300 | Haptista;Centroplasthelida;Pterocystida;Raphidista | EP00300 Choanocystis sp HF-7 18815 | 4,20E-70 | 495 | XP 646771.1 | hypothetical protein DDB G0269198 [Dictyostelium discoideum AX4] | 1992 | 5,56E-30 | 135,576 | 43,45679012 |
| EP00081 | Obazoa;Opisthokonta;Holozoa;Choanozoa;Metazoa | EP00081 Caenorhabditis elegans 18549 | 4,50E-70 | 1232 | NP 499022.1 | Protein transport protein Sec16 [Caenorhabditis elegans] | 1232 | 0 | 2544,61 | 100 |
| EP00311 | Haptista;Haptophyta;Prymnesiophyceae;Calcihapto | EP00311 Isochrysis galbana 10858 | 6,60E-70 | 2564 | XP 004344939.1 | hypothetical protein ACA1 285550 [Acanthamoeba castellanii str. Neff] | 1515 | 4,53E-31 | 146,747 | 42,89693593 |
| EP00269 | Archaeplastida;Chloroplastida;Streptophyta;"core s | EP00269 Selaginella moellendorffii 31614 | 1,30E-69 | 273 | XP 024532260.1 | protein transport protein SEC16B homolog isoform X2 [Selaginella moellendorffii] | 273 | 0 | 571,237 | 100 |
| EP00158 | Obazoa;Opisthokonta;Nucletmycea;Fungi;Aphelide | EP00158 Paraphelidium tribonemae 8366 | 1,30E-69 | 426 | XP 012946927.1 | PREDICTED: uncharacterized protein LOC101850917 isoform X3 [Aplysia californica] | 2276 | 1,23E-31 | 139,043 | 42,37918216 |
| EP00081 | Obazoa;Opisthokonta;Holozoa;Choanozoa;Metazoa | EP00081 Caenorhabditis elegans 8471 | 1,40E-69 | 1224 | NP 001293651.1 | Protein transport protein Sec16 [Caenorhabditis elegans] | 1224 | 0 | 2529,21 | 100 |
| EP00194 | Archaeplastida;Chloroplastida;Chlorophyta;"core ch | EP00194 Tetraselmis striata 37615 | 2,00E-68 | 783 | XP 020240850.1 | protein transport protein SEC16B homolog [Asparagus officinalis] | 1486 | 4,31E-59 | 229,565 | 44,82173175 |
| EP00446 | Sar;Alveolata;Myzozoa;"DP clade";Dinoflagellata;D | EP00446 Apocalathium sp SHHI-4 91619 | 4,20E-68 | 972 | XP 012756363.1 | hypothetical protein SAMD00019534 027760 [Acytostelium subglobosum LB1] | 1470 | 5,11E-21 | 111,309 | 42,39828694 |
| EP00189 | Archaeplastida;"RR clade";Rhodophyta;Proteorhod | EP00189 Rhodosorus marinus 912 | 2,20E-67 | 3361 | XP 005711041.1 | unnamed protein product [Chondrus crispus] | 2546 | 1,20E-37 | 169,088 | 42,27467811 |
| EP00592 | Sar;Stramenopiles;Gyrista;Ochrophyta;"BD clade";[ | EP00592 Proboscia alata 29354 | 2,30E-67 | 502 | XP 002183614.1 | predicted protein [Phaeodactylum tricornutum CCAP 1055/1] | 2083 | 5,24E-49 | 192,586 | 50,31055901 |
| EP00177 | Archaeplastida;"RR clade";Rhodophyta;Eurhodophy | EP00177 Eucheuma denticulatum 6069 | 3,20E-67 | 2012 | XP 005711041.1 | unnamed protein product [Chondrus crispus] | 2546 | 0 | 826,239 | 54,26774483 |
| EP00042 | Obazoa;Opisthokonta;Holozoa;Choanozoa;Choanof | EP00042 Codosiga hollandica 56371 | 4,60E-67 | 2325 | XP 021174898.1 | protein transport protein Sec16A isoform X2 [Fundulus heteroclitus] | 2341 | 1,27E-38 | 171,4 | 43,17617866 |
| EP00437 | Sar;Alveolata;Myzozoa;"DP clade";Dinoflagellata;D | EP00437 Effrenium voratum 24811 | 6,10E-67 | 687 | XP 016606650.1 | hypothetical protein SPPG 06291 [Spizellomyces punctatus DAOM BR117] | 2211 | 4,78E-20 | 107,457 | 44,83627204 |
| EP00674 | Discoba;Euglenozoa;Kinetoplastea;Metakinetoplast | EP00674 Leishmania major 7454 | 6,40E-67 | 1968 | XP 003722202.1 | conserved hypothetical protein [Leishmania major strain Friedlin] | 1968 | 0 | 3911,69 | 100 |
| EP00401 | Sar;Alveolata;Myzozoa;"DP clade";Dinoflagellata;D | EP00401 Gymnodinium catenatum 61934 | 7,60E-67 | 649 | XP 016606650.1 | hypothetical protein SPPG 06291 [Spizellomyces punctatus DAOM BR117] | 2211 | 7,40E-17 | 96,6709 | 38,50931677 |
| EP00043 | Obazoa;Opisthokonta;Holozoa;Choanozoa;Choanof | EP00043 Microstomoeca roanoka 16640 | 1,20E-66 | 1005 | XP 004992030.1 | hypothetical protein PTSG 12463 [Salpingoeca rosetta] | 2149 | 8,94E-140 | 473,396 | 63,86554622 |
| EP00325 | Haptista;Haptophyta;Prymnesiophyceae;Prymnesia | EP00325 Haptolina brevifila 18780 | 3,20E-66 | 965 | XP 004344939.1 | hypothetical protein ACA1 285550 [Acanthamoeba castellanii str. Neff] | 1515 | 1,09E-36 | 162,155 | 44,25287356 |
| EP00189 | Archaeplastida;"RR clade";Rhodophyta;Proteorhod | EP00189 Rhodosorus marinus 904 | 3,40E-66 | 3341 | XP 005711041.1 | unnamed protein product [Chondrus crispus] | 2546 | 1,00E-37 | 169,474 | 42,27467811 |
| EP00520 | Sar;Stramenopiles;Gyrista;Ochrophyta;"BD clade";E | EP00520 Triparma pacifica 17432 | 3,90E-66 | 1120 | XP 004344939.1 | hypothetical protein ACA1 285550 [Acanthamoeba castellanii str. Neff] | 1515 | 4,10E-32 | 147,902 | 44,89311164 |
| EP00139 | Obazoa;Opisthokonta;Nucletmycea;Fungi;"core Fur | EP00139 Candida albicans 4942 | 5,10E-66 | 2091 | XP 720651.2 | hypothetical protein CAALFM C503140CA [Candida albicans SC5314] | 2091 | 0 | 4235,64 | 100 |
| EP00437 | Sar;Alveolata;Myzozoa;"DP clade";Dinoflagellata;D | EP00437 Effrenium voratum 57964 | 5,20E-66 | 688 | XP 016606650.1 | hypothetical protein SPPG 06291 [Spizellomyces punctatus DAOM BR117] | 2211 | 4,64E-20 | 107,457 | 44,83627204 |
| EP00426 | Sar;Alveolata;Myzozoa;"DP clade";Dinoflagellata;D | EP00426 Karenia brevis 114147 | 1,10E-65 | 1033 | XP 002908536.1 | COPII coat assembly protein SEC16, putative [Phytophthora infestans T30-4] | 2641 | 5,23E-23 | 118,242 | 41,77693762 |
| EP00437 | Sar;Alveolata;Myzozoa;"DP clade";Dinoflagellata;D | EP00437 Effrenium voratum 52340 | 1,30E-65 | 521 | XP 016606650.1 | hypothetical protein SPPG 06291 [Spizellomyces punctatus DAOM BR117] | 2211 | 6,87E-18 | 98,9821 | 44,83627204 |
| EP00317 | Haptista;Haptophyta;Prymnesiophyceae;Phaeocyst | EP00317 Phaeocystis antarctica 10245 | 6,70E-65 | 392 | XP 004344939.1 | hypothetical protein ACA1 285550 [Acanthamoeba castellanii str. Neff] | 1515 | 7,86E-34 | 144,436 | 47,03557312 |
| EP00262 | Archaeplastida;Chloroplastida;Streptophyta;"core s | EP00262 Sarcandra glabra 5651 | 1,70E-64 | 523 | XP 010264305.1 | PREDICTED: protein transport protein SEC16A homolog isoform X2 [Nelumbo nucifera] | 1393 | 0 | 677,167 | 71,11111111 |
| EP00426 | Sar;Alveolata;Myzozoa;"DP clade";Dinoflagellata;D | EP00426 Karenia brevis 100499 | 2,40E-64 | 1041 | XP 002908536.1 | COPII coat assembly protein SEC16, putative [Phytophthora infestans T30-4] | 2641 | 1,44E-24 | 123,635 | 41,81523501 |
| EP00045 | Obazoa;Opisthokonta;Holozoa;Choanozoa;Choanof | EP00045 Choanoeca perplexa 17471 | 3,90E-64 | 1949 | XP 001747683.1 | hypothetical protein [Monosiga brevicollis MX1] | 2207 | 7,60E-94 | 351,288 | 43,07992203 |
| EP00269 | Archaeplastida;Chloroplastida;Streptophyta;"core s | EP00269 Selaginella moellendorffii 31613 | 1,60E-63 | 289 | XP 024532259.1 | protein transport protein SEC16B homolog isoform X1 [Selaginella moellendorffii] | 289 | 0 | 607,06 | 100 |
| EP00614 | Sar;Stramenopiles;Gyrista;Ochrophyta;Dictyochoph | EP00614 Pseudopedinella elastica 391 | 2,60E-63 | 866 | XP 008890960.1 | hypothetical protein PPTG 01118 [Phytophthora parasitica INRA-310] | 1621 | 1,10E-17 | 100,138 | 50,89108911 |
| EP00588 | Sar;Stramenopiles;Gyrista;Ochrophyta;"BD clade";[ | EP00588 Corethron pennatum 3672 | 2,70E-63 | 585 | XP 002183614.1 | predicted protein [Phaeodactylum tricornutum CCAP 1055/1] | 2083 | 1,61E-22 | 114,39 | 43,26241135 |
| EP00523 | Sar;Stramenopiles;Gyrista;Ochrophyta;"BD clade";[ | EP00523 Amphiprora sp CCMP467 13782 | 5,60E-63 | 1009 | XP 002183614.1 | predicted protein [Phaeodactylum tricornutum CCAP 1055/1] | 2083 | 3,07E-64 | 249,595 | 51,67130919 |
| EP00264 | Archaeplastida;Chloroplastida;Streptophyta;"core s | EP00264 Hordeum vulgare 41452 | 1,00E-62 | 914 | XP 020151898.1 | protein transport protein SEC16A homolog [Aegilops tauschii subsp. tauschii] | 1490 | 0 | 1478,38 | 88,18283166 |
| EP00387 | Sar;Alveolata;Myzozoa;"AC clade";Colpodellida;Chr | EP00387 Chromera velia 12698 | 1,10E-62 | 563 | XP 006358347.1 | PREDICTED: protein transport protein SEC16B homolog [Solanum tuberosum] | 1471 | 4,70E-18 | 99,7525 | 40 |
| EP00364 | Sar;Alveolata;Myzozoa;"AC clade";Apicomplexa;"cc | EP00364 Plasmodium chabaudi 4905 | 1,40E-62 | 1320 | XP 745504.2 | conserved Plasmodium protein, unknown function [Plasmodium chabaudi chabaudi] | 1320 | 0 | 2647,46 | 100 |
| EP00562 | Sar;Stramenopiles;Gyrista;Ochrophyta;"BD clade";[ | EP00562 Extubocellulus spinifer 24080 | 1,90E-62 | 708 | XP 002183614.1 | predicted protein [Phaeodactylum tricornutum CCAP 1055/1] | 2083 | 2,44E-51 | 204,527 | 49,49748744 |
| EP00402 | Sar;Alveolata;Myzozoa;"DP clade";Dinoflagellata;D | EP00402 Amphidinium carterae 4627 | 3,20E-62 | 608 | XP 003288899.1 | hypothetical protein DICPUDRAFT 98175 [Dictyostelium purpureum] | 1854 | 3,49E-16 | 94,3597 | 41,38702461 |
| EP00394 | Sar;Alveolata;Myzozoa;"DP clade";Dinoflagellata;"c | EP00394 Amoebophrya sp ex-Karlodinium-veneficum 4667 | 4,90E-62 | 532 | XP 008890960.1 | hypothetical protein PPTG 01118 [Phytophthora parasitica INRA-310] | 1621 | 1,57E-17 | 97,8265 | 41,82242991 |
| EP00744 | Sar;Alveolata;Colponemidae;Colponema;Colponem | EP00744 Colponema vietnamica 13382 | 1,80E-61 | 319 | XP 020431748.1 | hypothetical protein PPL 07486 [Heterostelium album PN500] | 1678 | 2,67E-35 | 146,362 | 46,84210526 |
| EP00705 | Metamonada;Parabasalia;Trichomonadida;Trichom | EP00705 Trichomonas vaginalis 55997 | 1,80E-61 | 2050 | XP 001580855.1 | hypothetical protein [Trichomonas vaginalis G3] | 2050 | 0 | 4021,08 | 100 |
| EP00141 | Obazoa;Opisthokonta;Nucletmycea;Fungi;"core Fur | EP00141 Yarrowia lipolytica 1312 | 3,20E-61 | 2183 | XP 500810.1 | YALI0B12694p [Yarrowia lipolytica CLIB122] | 2183 | 0 | 4368,92 | 100 |
| EP00506 | Sar;Stramenopiles;Bigyra;Sagenista;"MAST-04";stra | EP00506 MAST4 sp TOSAG23-1 7107 | 3,90E-61 | 1778 | XP 011312055.1 | PREDICTED: uncharacterized protein LOC105271924 [Fopius arisanus] | 2092 | 9,09E-19 | 105,531 | 37,5 |
| EP00126 | Obazoa;Opisthokonta;Holozoa;Ichthyosporea;Rhinc | EP00126 Sphaerothecum destruens 2098 | 5,70E-61 | 512 | XP 004344939.1 | hypothetical protein ACA1 285550 [Acanthamoeba castellanii str. Neff] | 1515 | 1,50E-15 | 91,2781 | 46,61016949 |
| EP00636 | Sar;Stramenopiles;Gyrista;Ochrophyta;Pinguiophyc | EP00636 Phaeomonas parva 17952 | 6,80E-61 | 315 | XP 002183614.1 | predicted protein [Phaeodactylum tricornutum CCAP 1055/1] | 2083 | 6,77E-29 | 127,487 | 37,22943723 |
| EP00592 | Sar;Stramenopiles;Gyrista;Ochrophyta;"BD clade";[ | EP00592 Proboscia alata 16020 | 1,30E-60 | 718 | XP 002183614.1 | predicted protein [Phaeodactylum tricornutum CCAP 1055/1] | 2083 | 1,33E-47 | 193,741 | 54,50901804 |
| EP00403 | Sar;Alveolata;Myzozoa;"DP clade";Dinoflagellata;D | EP00403 Amphidinium massartii 46355 | 1,60E-60 | 1121 | XP 011121253.1 | hypothetical protein AOL s00076g399 [Arthrobotrys oligospora ATCC 24927] | 1959 | 3,25E-20 | 109,383 | 42,15500945 |
| EP00750 | Sar;Stramenopiles;Bigyra;Opalozoa;Nanomonadea; | EP00750 Incisomonas marina 27668 | 1,90E-60 | 1176 | XP 008608960.1 | hypothetical protein SDRG 05043 [Saprolegnia diclina VS20] | 2628 | 7,36E-15 | 92,0485 | 52,734375 |
| EP00049 | Obazoa;Opisthokonta;Holozoa;Choanozoa;Choanof | EP00049 Salpingoeca infusionum 10020 | 2,90E-60 | 1676 | XP 004992030.1 | hypothetical protein PTSG 12463 [Salpingoeca rosetta] | 2149 | 1,11E-35 | 160,999 | 42,08144796 |
| EP00176 | Archaeplastida;"RR clade";Rhodophyta;Eurhodophy | EP00176 Chondrus crispus 1209 | 3,40E-60 | 2546 | XP 005711041.1 | unnamed protein product [Chondrus crispus] | 2546 | 0 | 5202,88 | 100 |
| EP00683 | Discoba;Euglenozoa;Kinetoplastea;Metakinetoplast | EP00683 Trypanosoma rangeli 6134 | 5,00E-60 | 1446 | XP 029239091.1 | uncharacterized protein TraAM80 04123 [Trypanosoma rangeli] | 1446 | 0 | 2928,27 | 100 |
| EP00376 | Sar;Alveolata;Myzozoa;"AC clade";Apicomplexa;"cc | EP00376 Eimeria tenella 5056 | 8,70E-60 | 5802 | XP 013232789.1 | hypothetical protein, conserved [Eimeria tenella] | 5802 | 0 | 11199,7 | 100 |
| EP00312 | Haptista;Haptophyta;Prymnesiophyceae;Calcihapto | EP00312 Isochrysis sp CCMP1244 39823 | 1,10E-59 | 439 | XP 004344939.1 | hypothetical protein ACA1 285550 [Acanthamoeba castellanii str. Neff] | 1515 | 1,29E-32 | 142,124 | 43,85964912 |
| EP00187 | Archaeplastida;"RR clade";Rhodophyta;Proteorhod | EP00187 Rhodella violacea 11202 | 1,90E-59 | 1349 | XP 005711041.1 | unnamed protein product [Chondrus crispus] | 2546 | 8,47E-31 | 144,436 | 40,8993576 |
| EP00607 | Sar;Stramenopiles;Gyrista;Ochrophyta;"CS clade";C | EP00607 Mallomonas sp CCMP3275 3098 | 2,10E-59 | 1255 | XP 009523981.1 | hypothetical protein PHYSODRAFT 262855 [Phytophthora sojae] | 2808 | 2,33E-10 | 77,411 | 49,8470948 |
| EP00603 | Sar;Stramenopiles;Gyrista;Ochrophyta;"CS clade";C | EP00603 Paraphysomonas imperforata 13536 | 4,80E-59 | 1644 | XP 009523981.1 | hypothetical protein PHYSODRAFT 262855 [Phytophthora sojae] | 2808 | 1,11E-17 | 102,064 | 34,71502591 |
| EP00315 | Haptista;Haptophyta;Prymnesiophyceae;Calcihapto | EP00315 Gephyrocapsa oceanica 35085 | 9,10E-59 | 428 | XP 004344939.1 | hypothetical protein ACA1 285550 [Acanthamoeba castellanii str. Neff] | 1515 | 2,56E-33 | 144,05 | 44,3148688 |
| EP00451 | Sar;Alveolata;Myzozoa;"DP clade";Dinoflagellata;O | EP00451 Oxyrrhis marina 46424 | 4,80E-58 | 857 | XP 003840368.1 | hypothetical protein LEMA P100200.1 [Leptosphaeria maculans JN3] | 1821 | 3,46E-10 | 75,8702 | 51,74825175 |
| EP00086 | Obazoa;Opisthokonta;Holozoa;Choanozoa;Metazoa | EP00086 Labidocera madurae 38613 | 6,20E-58 | 2426 | XP 020125811.1 | copii coat assembly protein sec16 [Diplodia corticola] | 2166 | 5,00E-19 | 107,071 | 44,32624113 |
| EP00086 | Obazoa;Opisthokonta;Holozoa;Choanozoa;Metazoa | EP00086 Labidocera madurae 38614 | 6,20E-58 | 2426 | XP 020125811.1 | copii coat assembly protein sec16 [Diplodia corticola] | 2166 | 5,00E-19 | 107,071 | 44,32624113 |
| EP00538 | Sar;Stramenopiles;Gyrista;Ochrophyta;"BD clade";[ | EP00538 Stauroneis constricta 12866 | 1,10E-57 | 1253 | XP 002183614.1 | predicted protein [Phaeodactylum tricornutum CCAP 1055/1] | 2083 | 1,76E-70 | 271,937 | 47,56606398 |
| EP00589 | Sar;Stramenopiles;Gyrista;Ochrophyta;"BD clade";[ | EP00589 Leptocylindrus danicus 26336 | 1,20E-57 | 1961 | XP 008890960.1 | hypothetical protein PPTG 01118 [Phytophthora parasitica INRA-310] | 1621 | 9,08E-28 | 135,191 | 44,64802314 |
| EP00536 | Sar;Stramenopiles;Gyrista;Ochrophyta;"BD clade";[ | EP00536 Pseudo-nitzschia multiseries 11689 | 2,10E-57 | 1613 | XP 002183614.1 | predicted protein [Phaeodactylum tricornutum CCAP 1055/1] | 2083 | 1,59E-100 | 370,163 | 50,42735043 |
| EP00157 | Obazoa;Opisthokonta;Nucletmycea;Fungi;"cryptom | EP00157 Rozella allomycis 3289 | 9,10E-57 | 1426 | ***no hit found*** | | | | | |
| EP00529 | Sar;Stramenopiles;Gyrista;Ochrophyta;"BD clade";[ | EP00529 Nitzschia sp RCC80 12717 | 2,00E-56 | 595 | XP 002183614.1 | predicted protein [Phaeodactylum tricornutum CCAP 1055/1] | 2083 | 2,17E-23 | 117,087 | 46,84684685 |
| EP00269 | Archaeplastida;Chloroplastida;Streptophyta;"core s | EP00269 Selaginella moellendorffii 16441 | 2,50E-56 | 431 | XP 024517087.1 | protein transport protein SEC16B homolog [Selaginella moellendorffii] >gi\|1376974737\|r | 431 | 0 | 901,353 | 100 |
| EP00269 | Archaeplastida;Chloroplastida;Streptophyta;"core s | EP00269 Selaginella moellendorffii 16442 | 2,50E-56 | 431 | XP 024517087.1 | protein transport protein SEC16B homolog [Selaginella moellendorffii] >gi\|1376974737\|r | 431 | 0 | 901,353 | 100 |
| EP00747 | Sar;Alveolata;Myzozoa;"DP clade";Dinoflagellata;D | EP00747 Dinoflagellata sp TGD 51025 | 2,70E-56 | 760 | XP 019922177.1 | PREDICTED: mucin-17 isoform X10 [Crassostrea gigas] | 6562 | 1,39E-10 | 77,0258 | 40,6557377 |
| EP00425 | Sar;Alveolata;Myzozoa;"DP clade";Dinoflagellata;D | EP00425 Heterocapsa triquetra 5354 | 2,80E-56 | 1059 | ***no hit found*** | | | | | |
| EP00620 | Sar;Stramenopiles;Gyrista;Ochrophyta;Dictyochoph | EP00620 Florenciella sp RCC1587 5374 | 2,90E-56 | 463 | XP 008890960.1 | hypothetical protein PPTG 01118 [Phytophthora parasitica INRA-310] | 1621 | 8,23E-35 | 149,443 | 43,22916667 |
| EP00739 | Archaeplastida;Chloroplastida;Chlorophyta;Mamiel | EP00739 Bathycoccus prasinos 6505 | 3,50E-56 | 1142 | XP 007509414.1 | unknown protein [Bathycoccus prasinos] | 1174 | 0 | 2269,2 | 98,99074853 |
| EP00167 | Archaeplastida;"RR clade";Rhodophyta;Cyanidiales; | EP00167 Galdieria sulphuraria 3310 | 7,60E-56 | 1395 | XP 005705546.1 | hypothetical protein Gasu 35950 [Galdieria sulphuraria] | 1395 | 0 | 2869,72 | 100 |
| EP00682 | Discoba;Euglenozoa;Kinetoplastea;Metakinetoplast | EP00682 Trypanosoma cruzi 12369 | 1,20E-55 | 1482 | XP 814906.1 | hypothetical protein [Trypanosoma cruzi strain CL Brener] | 1482 | 0 | 3006,47 | 100 |
| EP00179 | Archaeplastida;"RR clade";Rhodophyta;Proteorhod | EP00179 Madagascaria erythrocladioides 15049 | 1,50E-55 | 429 | XP 004354463.1 | hypothetical protein DFA 10563 [Cavenderia fasciculata] | 1778 | 4,31E-21 | 107,457 | 40,97222222 |
| EP00185 | Archaeplastida;"RR clade";Rhodophyta;Proteorhod | EP00185 Porphyridium purpureum 7149 | 2,40E-55 | 1475 | XP 005711041.1 | unnamed protein product [Chondrus crispus] | 2546 | 1,56E-15 | 94,7449 | 46,95259594 |
| EP00750 | Sar;Stramenopiles;Bigyra;Opalozoa;Nanomonadea; | EP00750 Incisomonas marina 27671 | 2,80E-55 | 1832 | XP 008608960.1 | hypothetical protein SDRG 05043 [Saprolegnia diclina VS20] | 2628 | 1,64E-15 | 95,1301 | 52,734375 |
| EP00658 | Telonemia;Telonema;strain P-2 | EP00658 Telonema sp P-2 54000 | 3,40E-55 | 398 | XP 018853918.1 | PREDICTED: protein transport protein SEC16A homolog isoform X2 [Juglans regia] | 1364 | 3,32E-24 | 115,931 | 45,93301435 |
| EP00406 | Sar;Alveolata;Myzozoa;"DP clade";Dinoflagellata;D | EP00406 Dinophysis acuminata 68854 | 6,10E-55 | 468 | XP 016606650.1 | hypothetical protein SPPG 06291 [Spizellomyces punctatus DAOM BR117] | 2211 | 6,01E-14 | 85,8853 | 44,48275862 |
| EP00363 | Sar;Alveolata;Myzozoa;"AC clade";Apicomplexa;"cc | EP00363 Plasmodium berghei 3548 | 6,30E-55 | 1331 | XP 676273.2 | conserved Plasmodium protein, unknown function [Plasmodium berghei ANKA] | 1331 | 0 | 2588,14 | 100 |
| EP00405 | Sar;Alveolata;Myzozoa;"DP clade";Dinoflagellata;D | EP00405 Crypthecodinium cohnii 56355 | 7,20E-55 | 535 | XP 012756363.1 | hypothetical protein SAMD00019534 027760 [Acytostelium subglobosum LB1] | 1470 | 1,41E-18 | 101,293 | 46,46226415 |
| EP00403 | Sar;Alveolata;Myzozoa;"DP clade";Dinoflagellata;D | EP00403 Amphidinium massartii 25259 | 8,00E-55 | 1304 | XP 011121253.1 | hypothetical protein AOL s00076g399 [Arthrobotrys oligospora ATCC 24927] | 1959 | 2,22E-20 | 110,153 | 42,15500945 |
| EP00747 | Sar;Alveolata;Myzozoa;"DP clade";Dinoflagellata;D | EP00747 Dinoflagellata sp TGD 51027 | 8,40E-55 | 808 | XP 019922177.1 | PREDICTED: mucin-17 isoform X10 [Crassostrea gigas] | 6562 | 5,83E-16 | 94,7449 | 43,89534884 |
| EP00186 | Archaeplastida;"RR clade";Rhodophyta;Proteorhod | EP00186 Timspurckia oligopyrenoides 1349 | 1,00E-54 | 1259 | XP 005711041.1 | unnamed protein product [Chondrus crispus] | 2546 | 2,51E-22 | 116,701 | 40,75630252 |
| EP00376 | Sar;Alveolata;Myzozoa;"AC clade";Apicomplexa;"cc | EP00376 Eimeria tenella 534 | 1,10E-54 | 5136 | XP 013228260.1 | hypothetical protein, conserved [Eimeria tenella] | 5136 | 0 | 9844,15 | 100 |
| EP00673 | Discoba;Euglenozoa;Kinetoplastea;Metakinetoplast | EP00673 Leishmania infantum 3733 | 1,20E-54 | 2158 | XP 001466649.2 | conserved hypothetical protein [Leishmania infantum JPCM5] | 2158 | 0 | 4265,3 | 100 |
| EP00444 | Sar;Alveolata;Myzozoa;"DP clade";Dinoflagellata;D | EP00444 Apocalathium aciculiferum 59300 | 1,90E-54 | 1002 | XP 004344939.1 | hypothetical protein ACA1 285550 [Acanthamoeba castellanii str. Neff] | 1515 | 2,34E-17 | 99,7525 | 38,70967742 |
| EP00140 | Obazoa;Opisthokonta;Nucletmycea;Fungi;"core Fur | EP00140 Debaryomyces hansenii 1570 | 3,80E-54 | 2130 | XP 457088.2 | DEHA2B02838p [Debaryomyces hansenii CBS767] | 2130 | 0 | 4350,82 | 100 |
| EP00509 | Sar;Stramenopiles;Bigyra;Sagenista;"MAST-04";stra | EP00509 MAST4 sp TOSAG41-1 4493 | 6,20E-54 | 1993 | XP 012756363.1 | hypothetical protein SAMD00019534 027760 [Acytostelium subglobosum LB1] | 1470 | 8,90E-20 | 108,997 | 47,2972973 |
| EP00245 | Archaeplastida;Chloroplastida;Streptophyta;"core s | EP00245 Coleochaete scutata 12722 | 6,40E-54 | 197 | XP 024523188.1 | protein transport protein SEC16A homolog [Selaginella moellendorffii] | 381 | 4,58E-70 | 226,868 | 63,80368098 |
| EP00682 | Discoba;Euglenozoa;Kinetoplastea;Metakinetoplast | EP00682 Trypanosoma cruzi 6928 | 1,50E-53 | 1482 | XP 809465.1 | hypothetical protein [Trypanosoma cruzi strain CL Brener] | 1482 | 0 | 3007,62 | 100 |
| EP00616 | Sar;Stramenopiles;Gyrista;Ochrophyta;Dictyochoph | EP00616 Rhizochromulina sp CCMP1243 18002 | 1,50E-53 | 586 | XP 008890960.1 | hypothetical protein PPTG 01118 [Phytophthora parasitica INRA-310] | 1621 | 3,55E-17 | 97,4413 | 39,9463807 |
| EP00194 | Archaeplastida;Chloroplastida;Chlorophyta;"core ch | EP00194 Tetraselmis striata 16865 | 1,60E-53 | 517 | XP 024373339.1 | protein transport protein SEC16B homolog isoform X2 [Physcomitrella patens] >gi\|13738! | 1264 | 4,84E-29 | 132,88 | 44,90566038 |
| EP00116 | Obazoa;Opisthokonta;Holozoa;Choanozoa;Metazoa | EP00116 Pleurobrachia bachei 1737 | 7,20E-53 | 861 | XP 029473752.1 | protein transport protein Sec16B-like isoform X3 [Rhinatrema bivittatum] | 988 | 8,29E-31 | 142,124 | 38,125 |
| EP00619 | Sar;Stramenopiles;Gyrista;Ochrophyta;Dictyochoph | EP00619 Florenciella sp RCC1007 6678 | 8,50E-53 | 358 | XP 002908536.1 | COPII coat assembly protein SEC16, putative [Phytophthora infestans T30-4] | 2641 | 1,83E-26 | 121,709 | 46,38157895 |
| EP00232 | Archaeplastida;Chloroplastida;Chlorophyta;Nephro | EP00232 Nephroselmis pyriformis 4987 | 9,80E-53 | 288 | XP 024373338.1 | protein transport protein SEC16A homolog isoform X1 [Physcomitrella patens] >gi\|13738! | 1444 | 1,57E-52 | 194,897 | 55,55555556 |
| EP00403 | Sar;Alveolata;Myzozoa;"DP clade";Dinoflagellata;D | EP00403 Amphidinium massartii 31865 | 1,00E-52 | 1107 | XP 011121253.1 | hypothetical protein AOL s00076g399 [Arthrobotrys oligospora ATCC 24927] | 1959 | 4,98E-20 | 108,612 | 42,15500945 |
| EP00518 | Sar;Stramenopiles;Gyrista;Ochrophyta;"BD clade";E | EP00518 Triparma eleuthera 168 | 1,20E-52 | 454 | XP 002183614.1 | predicted protein [Phaeodactylum tricornutum CCAP 1055/1] | 2083 | 1,33E-15 | 90,8929 | 47,59358289 |
| EP00189 | Archaeplastida;"RR clade";Rhodophyta;Proteorhod | EP00189 Rhodosorus marinus 8517 | 1,30E-52 | 1746 | XP 005711041.1 | unnamed protein product [Chondrus crispus] | 2546 | 5,51E-35 | 159,073 | 42,6124197 |
| EP00331 | Sar;Alveolata;Ciliophora;Intramacronucleata;"CON: | EP00331 Platyophrya macrostoma 199 | 1,50E-52 | 447 | XP 814906.1 | hypothetical protein [Trypanosoma cruzi strain CL Brener] | 1482 | 1,08E-21 | 109,383 | 51,03092784 |
| EP00056 | Obazoa;Opisthokonta;Holozoa;Choanozoa;Choanof | EP00056 Hartaetosiga gracilis 10177 | 2,40E-52 | 1228 | XP 004992030.1 | hypothetical protein PTSG 12463 [Salpingoeca rosetta] | 2149 | 1,51E-57 | 230,72 | 39,02912621 |
| EP00115 | Obazoa;Opisthokonta;Holozoa;Choanozoa;Metazoa | EP00115 Mnemiopsis leidyi 12869 | 2,90E-52 | 758 | XP 026152800.1 | protein transport protein Sec16A isoform X4 [Mastacembelus armatus] | 2176 | 7,39E-33 | 148,673 | 40,45801527 |
| EP00403 | Sar;Alveolata;Myzozoa;"DP clade";Dinoflagellata;D | EP00403 Amphidinium massartii 23952 | 6,60E-52 | 1293 | XP 011121253.1 | hypothetical protein AOL s00076g399 [Arthrobotrys oligospora ATCC 24927] | 1959 | 3,56E-19 | 106,301 | 42,15500945 |
| EP00756 | Discoba;Euglenozoa;Diplonemea;Hemistasiidae;Her | EP00756 Hemistasia phaeocysticola 61208 | 1,50E-51 | 634 | ***no hit found*** | | | | | |
| EP00143 | Obazoa;Opisthokonta;Nucletmycea;Fungi;"core Fur | EP00143 Candida glabrata 2324 | 4,70E-51 | 2238 | XP 447038.1 | uncharacterized protein CAGL0H05577g [[Candida] glabrata] | 2238 | 0 | 4568,07 | 100 |
| EP00602 | Sar;Stramenopiles;Gyrista;Ochrophyta;"CS clade";C | EP00602 Paraphysomonas bandaiensis 13521 | 5,30E-51 | 1490 | XP 002908536.1 | COPII coat assembly protein SEC16, putative [Phytophthora infestans T30-4] | 2641 | 3,62E-21 | 113,235 | 47,79771615 |
| EP00179 | Archaeplastida;"RR clade";Rhodophyta;Proteorhod | EP00179 Madagascaria erythrocladioides 14636 | 1,40E-50 | 1541 | XP 022845957.1 | protein transport protein SEC16B homolog [Olea europaea var. sylvestris] | 1414 | 1,22E-10 | 78,5666 | 48,23529412 |
| EP00750 | Sar;Stramenopiles;Bigyra;Opalozoa;Nanomonadea; | EP00750 Incisomonas marina 27667 | 1,70E-50 | 1692 | XP 008608960.1 | hypothetical protein SDRG 05043 [Saprolegnia diclina VS20] | 2628 | 2,44E-15 | 94,3597 | 52,734375 |
| EP00057 | Obazoa;Opisthokonta;Holozoa;Choanozoa;Metazoa | EP00057 Strongylocentrotus purpuratus 15881 | 1,10E-49 | 4112 | XP 011670355.1 | PREDICTED: mucin-17-like [Strongylocentrotus purpuratus] | 4112 | 0 | 7661,61 | 100 |
| EP00295 | Cryptista;Cryptophyceae;Goniomonas;"ma G clade" | EP00295 Goniomonas pacifica 12372 | 1,20E-49 | 486 | XP 020697806.1 | protein transport protein SEC16A homolog [Dendrobium catenatum] >gi\|1622115509\|ref | 1397 | 9,29E-21 | 107,457 | 41,59663866 |
| EP00749 | Sar;Stramenopiles;Bigyra;Opalozoa;Bicosoecida;"cc | EP00749 Cafeteria roenbergensis 8155 | 3,00E-49 | 791 | XP 002908536.1 | COPII coat assembly protein SEC16, putative [Phytophthora infestans T30-4] | 2641 | 7,74E-12 | 81,2629 | 38,8261851 |
| EP00756 | Discoba;Euglenozoa;Diplonemea;Hemistasiidae;Her | EP00756 Hemistasia phaeocysticola 52403 | 3,40E-49 | 409 | XP 023371948.1 | protein transport protein Sec16A isoform X1 [Otolemur garnettii] | 870 | 3,68E-08 | 67,0106 | 43,22033898 |
| EP00172 | Archaeplastida;"RR clade";Rhodophyta;Eurhodophy | EP00172 Hildenbrandia rubra 3868 | 5,60E-49 | 571 | XP 005711041.1 | unnamed protein product [Chondrus crispus] | 2546 | 2,26E-43 | 177,563 | 47,08249497 |
| EP00376 | Sar;Alveolata;Myzozoa;"AC clade";Apicomplexa;"cc | EP00376 Eimeria tenella 2514 | 9,30E-49 | 2958 | XP 013230241.1 | hypothetical protein, conserved [Eimeria tenella] | 2958 | 0 | 5676,29 | 100 |
| EP00427 | Sar;Alveolata;Myzozoa;"DP clade";Dinoflagellata;D | EP00427 Karlodinium veneficum 62429 | 1,00E-48 | 1255 | ***no hit found*** | | | | | |
| EP00573 | Sar;Stramenopiles;Gyrista;Ochrophyta;"BD clade";[ | EP00573 Skeletonema grethae 12880 | 2,00E-48 | 426 | XP 002183614.1 | predicted protein [Phaeodactylum tricornutum CCAP 1055/1] | 2083 | 2,81E-14 | 86,6557 | 66,42335766 |
| EP00393 | Sar;Alveolata;Myzozoa;"DP clade";Dinoflagellata;"c | EP00393 Amoebophrya sp ex-Akashiwo-sanguinea 8126 | 2,20E-48 | 1301 | XP 012756363.1 | hypothetical protein SAMD00019534 027760 [Acytostelium subglobosum LB1] | 1470 | 9,45E-20 | 107,842 | 47,78420039 |
| EP00675 | Discoba;Euglenozoa;Kinetoplastea;Metakinetoplast | EP00675 Leptomonas pyrrhocoris 7421 | 5,50E-48 | 1961 | XP 015658925.1 | hypothetical protein ABB37 04704 [Leptomonas pyrrhocoris] | 1961 | 0 | 3910,15 | 100 |
| EP00366 | Sar;Alveolata;Myzozoa;"AC clade";Apicomplexa;"cc | EP00366 Plasmodium knowlesi 1703 | 5,80E-48 | 1428 | XP 002259200.1 | hypothetical protein, conserved in Plasmodium species [Plasmodium knowlesi strain H] | 1428 | 0 | 2982,97 | 100 |

| | | | | | | | | | | |
|---|---|---|---|---|---|---|---|---|---|---|
| EP00670 | Discoba;Euglenozoa;Euglenida;Spirocuta;Euglenea;I | EP00670 Eutreptiella gymnastica 23706 | 6,90E-48 | 172 | XP 004344939.1 | hypothetical protein ACA1 285550 [Acanthamoeba castellanii str. Neff] | 1515 | 3,20E-23 | 105,916 | 38,69047619 |
| EP00676 | Discoba;Euglenozoa;Kinetoplastea;Metakinetoplast | EP00676 Leptomonas seymouri 5143 | 1,20E-47 | 2352 | XP 015658925.1 | hypothetical protein ABB37 04704 [Leptomonas pyrrhocoris] | 1961 | 0 | 1381,31 | 61,40149339 |
| EP00376 | Sar;Alveolata;Myzozoa;"AC clade";Apicomplexa;"cc | EP00376 Eimeria tenella 496 | 5,10E-47 | 4275 | XP 013228222.1 | hypothetical protein, conserved [Eimeria tenella] | 4275 | 0 | 8291,79 | 100 |
| EP00243 | Archaeplastida;Chloroplastida;Streptophyta;"core s | EP00243 Klebsormidium subtile 10664 | 5,90E-47 | 133 | XP 010692158.1 | PREDICTED: protein transport protein SEC16B homolog [Beta vulgaris subsp. vulgaris] >gi\| | 1373 | 2,10E-52 | 187,578 | 65,41353383 |
| EP00321 | Haptista;Haptophyta;Prymnesiophyceae;Phaeocyst | EP00321 Phaeocystis sp CCMP2710 16445 | 9,50E-47 | 343 | XP 004344939.1 | hypothetical protein ACA1 285550 [Acanthamoeba castellanii str. Neff] | 1515 | 1,37E-24 | 115,931 | 45,92274678 |
| EP00392 | Sar;Alveolata;Myzozoa;"DP clade";Dinoflagellata;"c | EP00392 Amoebophrya sp AT5 13438 | 2,70E-46 | 1079 | ***no hit found*** | | | | | |
| EP00739 | Archaeplastida;Chloroplastida;Chlorophyta;Mamiel | EP00739 Bathycoccus prasinos 7666 | 1,10E-45 | 544 | XP 007509414.1 | unknown protein [Bathycoccus prasinos] | 1174 | 0 | 815,453 | 84 |
| EP00244 | Archaeplastida;Chloroplastida;Streptophyta;"core s | EP00244 Chara vulgaris 8503 | 2,50E-45 | 118 | XP 019709693.1 | protein transport protein SEC16A homolog isoform X2 [Elaeis guineensis] | 1410 | 5,44E-52 | 185,652 | 65,51724138 |
| EP00589 | Sar;Stramenopiles;Gyrista;Ochrophyta;"BD clade";[ | EP00589 Leptocylindrus danicus 11600 | 3,90E-45 | 1188 | XP 002183614.1 | predicted protein [Phaeodactylum tricornutum CCAP 1055/1] | 2083 | 1,50E-22 | 117,087 | 46,11344538 |
| EP00397 | Sar;Alveolata;Myzozoa;"DP clade";Dinoflagellata;"c | EP00397 Hematodinium sp SG-2012 13097 | 7,90E-45 | 1030 | XP 015839572.1 | PREDICTED: protein transport protein Sec16A isoform X4 [Tribolium castaneum] | 1354 | 6,91E-19 | 104,76 | 46,61458333 |
| EP00455 | Sar;Rhizaria;"eorhiza";Aquavolonida;"NC10 C lineag | EP00455 Lapot gusevi 53636 | 1,10E-44 | 346 | XP 029230831.1 | uncharacterized protein Tco025E 02121 [Trypanosoma conorhini] | 1464 | 9,20E-16 | 89,7373 | 66,66666667 |
| EP00744 | Sar;Alveolata;Colponemidae;Colponema;Colponem | EP00744 Colponema vietnamica 590 | 2,80E-43 | 1227 | ***no hit found*** | | | | | |
| EP00600 | Sar;Stramenopiles;Gyrista;Ochrophyta;"CS clade";C | EP00600 Ochromonadales sp CCMP1393 9370 | 3,10E-43 | 1952 | ***no hit found*** | | | | | |
| EP00639 | Sar;Stramenopiles;Gyrista;Ochrophyta;Raphidophy | EP00639 Heterosigma akashiwo 9558 | 1,10E-42 | 196 | XP 002908536.1 | COPII coat assembly protein SEC16, putative [Phytophthora infestans T30-4] | 2641 | 2,22E-22 | 104,375 | 37,8238342 |
| EP00171 | Archaeplastida;"RR clade";Rhodophyta;Eurhodophy | EP00171 Calliarthron tuberculosum 14677 | 1,50E-42 | 621 | XP 005711041.1 | unnamed protein product [Chondrus crispus] | 2546 | 1,30E-79 | 286,189 | 47,76470588 |
| EP00367 | Sar;Alveolata;Myzozoa;"AC clade";Apicomplexa;"cc | EP00367 Plasmodium vivax 3305 | 2,00E-42 | 1454 | XP 001615317.1 | hypothetical protein, conserved [Plasmodium vivax] | 1454 | 0 | 3019,18 | 100 |
| EP00376 | Sar;Alveolata;Myzozoa;"AC clade";Apicomplexa;"cc | EP00376 Eimeria tenella 2225 | 2,50E-42 | 3489 | XP 013229952.1 | hypothetical protein, conserved, partial [Eimeria tenella] | 3489 | 0 | 6681,27 | 100 |
| EP00467 | Sar;Rhizaria;Cercozoa;Chlorarachnea;"core chlorara | EP00467 Chlorarachnion reptans 2385 | 2,60E-42 | 306 | XP 004344939.1 | hypothetical protein ACA1 285550 [Acanthamoeba castellanii str. Neff] | 1515 | 1,47E-25 | 117,472 | 59,55882353 |
| EP00055 | Obazoa;Opisthokonta;Holozoa;Choanozoa;Choanof | EP00055 Hartaetosiga balthica 14439 | 3,80E-42 | 496 | XP 004992030.1 | hypothetical protein PTSG 12463 [Salpingoeca rosetta] | 2149 | 4,57E-57 | 216,083 | 40,78549849 |
| EP00454 | Sar;Alveolata;Myzozoa;"DP clade";Perkinsea;Perkin | EP00454 Perkinsus marinus 22478 | 6,40E-42 | 1430 | XP 002787764.1 | hypothetical protein Pmar PMAR012541 [Perkinsus marinus ATCC 50983] | 1430 | 0 | 2859,71 | 100 |
| EP00765 | Metamonada;"BF clade";Fornicata;Carpediemonas; | EP00765 Carpediemonas membranifera 10921 | 1,70E-41 | 806 | XP 001688188.1 | AGAP011214-PA, partial [Anopheles gambiae str. PEST] | 750 | 4,18E-10 | 75,0998 | 41,66666667 |
| EP00599 | Sar;Stramenopiles;Gyrista;Ochrophyta;"CS clade";C | EP00599 Poterioochromonas sp BG-1 3126 | 2,20E-41 | 2252 | XP 008608960.1 | hypothetical protein SDRG 05043 [Saprolegnia diclina VS20] | 2628 | 1,44E-21 | 115,546 | 52,41090147 |
| EP00742 | Sar;Alveolata;Colponemidae;"MPE2-45 clade";strai | EP00742 Colponemidia sp Colp-10 18014 | 3,00E-41 | 817 | XP 003872434.1 | conserved hypothetical protein [Leishmania mexicana MHOM/GT/2001/U1103] | 1955 | 1,91E-08 | 70,0922 | 49,14089347 |
| EP00629 | Sar;Stramenopiles;Gyrista;Ochrophyta;Pelagophyce | EP00629 Pelagomonadales sp RCC1024 5869 | 3,40E-41 | 808 | XP 008347457.3 | protein transport protein SEC16B homolog [Malus domestica] | 1424 | 6,99E-08 | 68,1662 | 41,17647059 |
| EP00057 | Obazoa;Opisthokonta;Holozoa;Choanozoa;Metazoa | EP00057 Strongylocentrotus purpuratus 28935 | 8,80E-41 | 344 | XP 011683409.1 | PREDICTED: protein transport protein Sec16B-like [Strongylocentrotus purpuratus] | 344 | 0 | 702,59 | 100 |
| EP00539 | Sar;Stramenopiles;Gyrista;Ochrophyta;"BD clade";[ | EP00539 Tryblionella compressa 8893 | 9,60E-41 | 491 | XP 002183614.1 | predicted protein [Phaeodactylum tricornutum CCAP 1055/1] | 2083 | 3,84E-17 | 96,2857 | 46,01769912 |
| EP00617 | Sar;Stramenopiles;Gyrista;Ochrophyta;Dictyochoph | EP00617 Octactis speculum 1252 | 3,40E-40 | 352 | XP 002908536.1 | COPII coat assembly protein SEC16, putative [Phytophthora infestans T30-4] | 2641 | 3,62E-13 | 82,0333 | 34,4 |
| EP00599 | Sar;Stramenopiles;Gyrista;Ochrophyta;"CS clade";C | EP00599 Poterioochromonas sp BG-1 9571 | 4,90E-40 | 2197 | XP 008890960.1 | hypothetical protein PPTG 01118 [Phytophthora parasitica INRA-310] | 1621 | 3,63E-22 | 117,087 | 56,13207547 |
| EP00668 | Discoba;Euglenozoa;Euglenida;Spirocuta;Euglenea;I | EP00668 Euglena longa 44894 | 1,60E-39 | 153 | XP 004344939.1 | hypothetical protein ACA1 285550 [Acanthamoeba castellanii str. Neff] | 1515 | 7,49E-15 | 80,8777 | 34,58646617 |
| EP00376 | Sar;Alveolata;Myzozoa;"AC clade";Apicomplexa;"cc | EP00376 Eimeria tenella 830 | 1,70E-39 | 9860 | XP 013228556.1 | hypothetical protein, conserved [Eimeria tenella] | 9860 | 0 | 19804,3 | 100 |
| EP00599 | Sar;Stramenopiles;Gyrista;Ochrophyta;"CS clade";C | EP00599 Poterioochromonas sp BG-1 17857 | 1,80E-39 | 2252 | XP 008890960.1 | hypothetical protein PPTG 01118 [Phytophthora parasitica INRA-310] | 1621 | 1,35E-22 | 118,627 | 56,13207547 |
| EP00126 | Obazoa;Opisthokonta;Holozoa;Ichthyosporea;Rhinc | EP00126 Sphaerothecum destruens 9912 | 3,10E-39 | 864 | XP 004344939.1 | hypothetical protein ACA1 285550 [Acanthamoeba castellanii str. Neff] | 1515 | 1,10E-18 | 103,605 | 47,22222222 |
| EP00740 | Archaeplastida;Chloroplastida;Chlorophyta;Mamiel | EP00740 Mantoniella antarctica 11683 | 3,70E-39 | 163 | XP 002509249.1 | predicted protein [Micromonas commoda] | 1945 | 1,51E-69 | 238,039 | 73,88535032 |
| EP00376 | Sar;Alveolata;Myzozoa;"AC clade";Apicomplexa;"cc | EP00376 Eimeria tenella 7161 | 4,10E-39 | 3109 | XP 013234898.1 | hypothetical protein, conserved [Eimeria tenella] | 3109 | 0 | 5995,23 | 100 |
| EP00462 | Sar;Rhizaria;Cercozoa;"crown cercozoa";Thecofilose | EP00462 Mataza sp D1 22328 | 4,70E-39 | 528 | ***no hit found*** | | | | | |
| EP00291 | Cryptista;Cryptophyceae;Cryptomonadales;Crypton | EP00291 Cryptomonas curvata 22702 | 5,90E-39 | 118 | XP 009415458.1 | PREDICTED: protein transport protein SEC16B homolog isoform X1 [Musa acuminata subsı | 1421 | 1,33E-22 | 101,679 | 52,98507463 |
| EP00376 | Sar;Alveolata;Myzozoa;"AC clade";Apicomplexa;"cc | EP00376 Eimeria tenella 3864 | 6,60E-39 | 3068 | XP 013231597.1 | hypothetical protein ETH 00006040 [Eimeria tenella] | 3068 | 0 | 5964,03 | 100 |
| EP00126 | Obazoa;Opisthokonta;Holozoa;Ichthyosporea;Rhinc | EP00126 Sphaerothecum destruens 9913 | 1,70E-38 | 894 | XP 004344939.1 | hypothetical protein ACA1 285550 [Acanthamoeba castellanii str. Neff] | 1515 | 1,21E-18 | 103,605 | 47,22222222 |
| EP00750 | Sar;Stramenopiles;Bigyra;Opalozoa;Nanomonadea; | EP00750 Incisomonas marina 27669 | 1,90E-38 | 439 | XP 016851775.1 | PREDICTED: protein transport protein Sec16B [Anolis carolinensis] | 990 | 1,08E-08 | 68,9366 | 43,36283186 |
| EP00407 | Sar;Alveolata;Myzozoa;"DP clade";Dinoflagellata;D | EP00407 Lingulodinium polyedra 14127 | 2,20E-38 | 255 | XP 009523981.1 | hypothetical protein PHYSODRAFT 262855 [Phytophthora sojae] | 2808 | 3,08E-12 | 77,0258 | 46,78899083 |
| EP00378 | Sar;Alveolata;Myzozoa;"AC clade";Apicomplexa;"cc | EP00378 Neospora caninum 172 | 5,10E-38 | 2103 | XP 003879709.1 | conserved hypothetical protein [Neospora caninum Liverpool] | 2103 | 0 | 4062,69 | 100 |
| EP00020 | Amoebozoa;Evosea;Cutosea;Sapocribrum;Sapocribr | EP00020 Sapocribrum chincoteaguense 10058 | 7,70E-38 | 1571 | WP 148340744.1 | DUF945 family protein [Aquicella siphonis] | 689 | 2,29E-32 | 147,132 | 66,20689655 |
| EP00061 | Obazoa;Opisthokonta;Holozoa;Choanozoa;Metazoa | EP00061 Rhincodon typus 17205 | 1,70E-37 | 217 | XP 020378983.1 | LOW QUALITY PROTEIN: protein transport protein Sec16A-like [Rhincodon typus] | 2296 | 4,66E-109 | 353,984 | 94,50980392 |
| EP00088 | Obazoa;Opisthokonta;Holozoa;Choanozoa;Metazoa | EP00088 Acartia fossae 58075 | 2,00E-37 | 188 | XP 023346731.1 | protein transport protein Sec16-like isoform X2 [Eurytemora affinis] | 1941 | 1,97E-89 | 296,204 | 78,88888889 |
| EP00376 | Sar;Alveolata;Myzozoa;"AC clade";Apicomplexa;"cc | EP00376 Eimeria tenella 1013 | 2,20E-37 | 7937 | XP 013228739.1 | Midasin, related [Eimeria tenella] | 7937 | 0 | 15819 | 100 |
| EP00658 | Telonemia;Telonema;strain P-2 | EP00658 Telonema sp P-2 79807 | 4,40E-37 | 430 | XP 814906.1 | hypothetical protein [Trypanosoma cruzi strain CL Brener] | 1482 | 4,44E-15 | 88,9669 | 47,01754386 |
| EP00376 | Sar;Alveolata;Myzozoa;"AC clade";Apicomplexa;"cc | EP00376 Eimeria tenella 5967 | 6,60E-37 | 1958 | XP 013233700.1 | sybindin-like family domain-containing protein, putative [Eimeria tenella] | 1958 | 0 | 3860,84 | 100 |
| EP00706 | Metamonada;Parabasalia;Tritrichomonas;Tritrichor | EP00706 Tritrichomonas foetus 10885 | 3,60E-36 | 1534 | XP 001580855.1 | hypothetical protein [Trichomonas vaginalis G3] | 2050 | 1,39E-66 | 261,536 | 42,76595745 |
| EP00377 | Sar;Alveolata;Myzozoa;"AC clade";Apicomplexa;"cc | EP00377 Hammondia hammondi 5186 | 4,40E-36 | 1845 | XP 002370242.1 | hypothetical protein TGME49 294730 [Toxoplasma gondii ME49] | 2142 | 0 | 1549,64 | 90,37267081 |
| EP00061 | Obazoa;Opisthokonta;Holozoa;Choanozoa;Metazoa | EP00061 Rhincodon typus 19065 | 5,50E-36 | 1486 | XP 020378983.1 | LOW QUALITY PROTEIN: protein transport protein Sec16A-like [Rhincodon typus] | 2296 | 0 | 2862,4 | 98,06701031 |
| EP00267 | Archaeplastida;Chloroplastida;Streptophyta;"core s | EP00267 Zea mays 15586 | 1,70E-35 | 485 | NP 001348910.1 | uncharacterized protein LOC100383255 [Zea mays] | 1342 | 0 | 993,416 | 100 |
| EP00020 | Amoebozoa;Evosea;Cutosea;Sapocribrum;Sapocribr | EP00020 Sapocribrum chincoteaguense 7526 | 1,80E-35 | 1441 | WP 148340744.1 | DUF945 family protein [Aquicella siphonis] | 689 | 8,24E-36 | 157,532 | 63,23529412 |
| EP00677 | Discoba;Euglenozoa;Kinetoplastea;Metakinetoplast | EP00677 Phytomonas sp EM1 114 | 3,20E-35 | 1841 | XP 010700735.1 | hypothetical protein LPMP 291270 [Leishmania panamensis] | 2222 | 6,64E-84 | 318,931 | 52,64735265 |
| EP00598 | Sar;Stramenopiles;Gyrista;Ochrophyta;"CS clade";C | EP00598 Pedospumella elongata 6432 | 1,40E-34 | 2515 | ***no hit found*** | | | | | |
| EP00169 | Archaeplastida;"RR clade";Rhodophyta;Eurhodophy | EP00169 Porphyra umbilicalis 2135 | 2,00E-34 | 3158 | ***no hit found*** | | | | | |
| EP00613 | Sar;Stramenopiles;Gyrista;Ochrophyta;Dictyochoph | EP00613 Pedinella sp CCMP2098 12851 | 2,90E-34 | 324 | XP 026401446.1 | protein transport protein SEC16A homolog isoform X1 [Papaver somniferum] | 1558 | 9,30E-17 | 92,0485 | 48,25581395 |
| EP00020 | Amoebozoa;Evosea;Cutosea;Sapocribrum;Sapocribr | EP00020 Sapocribrum chincoteaguense 9135 | 3,20E-34 | 1446 | WP 148340744.1 | DUF945 family protein [Aquicella siphonis] | 689 | 4,93E-33 | 149,058 | 64,74820144 |
| EP00376 | Sar;Alveolata;Myzozoa;"AC clade";Apicomplexa;"cc | EP00376 Eimeria tenella 5172 | 3,20E-34 | 2903 | XP 013232905.1 | tRNA-splicing endonuclease positive effector, putative [Eimeria tenella] | 2903 | 0 | 5788,38 | 100 |
| EP00063 | Obazoa;Opisthokonta;Holozoa;Choanozoa;Metazoa | EP00063 Salmo salar 49762 | 3,60E-34 | 190 | XP 014024597.1 | PREDICTED: protein transport protein Sec16B-like [Salmo salar] | 190 | 5,48E-137 | 389,423 | 100 |
| EP00316 | Haptista;Haptophyta;Prymnesiophyceae;Calcihapto | EP00316 Scyphosphaera apsteinii 8626 | 4,50E-34 | 445 | XP 003840368.1 | hypothetical protein LEMA P100200.1 [Leptosphaeria maculans JN3] | 1821 | 7,86E-13 | 82,4185 | 54,14847162 |
| EP00466 | Sar;Rhizaria;Cercozoa;Chlorarachnea;"core chlorara | EP00466 Bigelowiella natans 4593 | 9,30E-34 | 131 | XP 024391503.1 | protein transport protein SEC16A homolog isoform X1 [Physcomitrella patens] >gi\|13739: | 1457 | 2,49E-25 | 110,153 | 57,66423358 |
| EP00421 | Sar;Alveolata;Myzozoa;"DP clade";Dinoflagellata;D | EP00421 Protoceratium reticulatum 28031 | 1,10E-33 | 155 | XP 009523981.1 | hypothetical protein PHYSODRAFT 262855 [Phytophthora sojae] | 2808 | 4,28E-12 | 73,1738 | 47,70642202 |
| EP00630 | Sar;Stramenopiles;Gyrista;Ochrophyta;Pelagophyce | EP00630 Chrysocystis fragilis 6297 | 1,20E-33 | 1727 | XP 002908536.1 | COPII coat assembly protein SEC16, putative [Phytophthora infestans T30-4] | 2641 | 4,64E-11 | 80,1073 | 36,19631902 |
| EP00127 | Obazoa;Opisthokonta;Holozoa;Pluriformea;Corallo | EP00127 Corallochytrium limacisporum 5087 | 1,90E-33 | 1715 | WP 089609777.1 | hypothetical protein [Dehalobacterium formicoaceticum] | 2198 | 2,24E-52 | 215,698 | 50,42979943 |
| EP00044 | Obazoa;Opisthokonta;Holozoa;Choanozoa;Choanof | EP00044 Salpingoeca rosetta 4269 | 7,30E-33 | 7577 | XP 004991702.1 | lankamycin synthase [Salpingoeca rosetta] | 7577 | 0 | 15401,4 | 100 |
| EP00237 | Archaeplastida;Chloroplastida;Chlorophyta;Pycnocc | EP00237 Pycnococcus provasolii 831 | 1,10E-32 | 519 | XP 010530052.1 | PREDICTED: protein transport protein SEC16B homolog [Tarenaya hassleriana] | 1377 | 1,48E-11 | 78,9518 | 47,40740741 |
| EP00455 | Sar;Rhizaria;"eorhiza";Aquavolonida;"NC10 C lineag | EP00455 Lapot gusevi 26800 | 1,40E-32 | 129 | XP 011772368.1 | hypothetical protein, conserved [Trypanosoma brucei gambiense DAL972] | 1387 | 1,11E-27 | 116,701 | 47,40740741 |
| EP00599 | Sar;Stramenopiles;Gyrista;Ochrophyta;"CS clade";C | EP00599 Poterioochromonas sp BG-1 13900 | 4,30E-32 | 1514 | XP 008608960.1 | hypothetical protein SDRG 05043 [Saprolegnia diclina VS20] | 2628 | 8,75E-21 | 112,079 | 50,86580087 |
| EP00612 | Sar;Stramenopiles;Gyrista;Ochrophyta;"PX clade";X | EP00612 Vaucheria litorea 1681 | 8,70E-32 | 461 | XP 017641979.1 | PREDICTED: protein transport protein SEC16B homolog [Gossypium arboreum] | 1386 | 2,10E-10 | 74,7146 | 45,12471655 |
| EP00308 | Haptista;Haptophyta;Prymnesiophyceae;Calcihapto | EP00308 Calcidiscus leptoporus 9913 | 9,70E-32 | 231 | XP 006646765.2 | PREDICTED: protein transport protein SEC16A homolog [Oryza brachyantha] | 1514 | 2,87E-17 | 90,8929 | 51,79487179 |
| EP00376 | Sar;Alveolata;Myzozoa;"AC clade";Apicomplexa;"cc | EP00376 Eimeria tenella 7344 | 2,80E-31 | 1515 | XP 013235083.1 | hypothetical protein ETH 00020910, partial [Eimeria tenella] | 1515 | 0 | 2897,84 | 100 |
| EP00599 | Sar;Stramenopiles;Gyrista;Ochrophyta;"CS clade";C | EP00599 Poterioochromonas sp BG-1 2144 | 1,30E-30 | 1056 | XP 009523981.1 | hypothetical protein PHYSODRAFT 262855 [Phytophthora sojae] | 2808 | 3,28E-12 | 83,1889 | 39,49044586 |
| EP00269 | Archaeplastida;Chloroplastida;Streptophyta;"core s | EP00269 Selaginella moellendorffii 35435 | 1,50E-30 | 119 | XP 024536081.1 | protein transport protein SEC16B homolog [Selaginella moellendorffii] | 119 | 9,62E-85 | 251,521 | 100 |
| EP00262 | Archaeplastida;Chloroplastida;Streptophyta;"core s | EP00262 Sarcandra glabra 5848 | 2,00E-30 | 167 | XP 010258928.1 | PREDICTED: protein transport protein SEC16B homolog [Nelumbo nucifera] >gi\|72000941 | 1429 | 1,92E-77 | 260,766 | 81,17647059 |
| EP00428 | Sar;Alveolata;Myzozoa;"DP clade";Dinoflagellata;D | EP00428 Durinskia baltica 71905 | 3,10E-30 | 161 | XP 002183614.1 | predicted protein [Phaeodactylum tricornutum CCAP 1055/1] | 2083 | 1,47E-16 | 86,2705 | 46,04316547 |
| EP00044 | Obazoa;Opisthokonta;Holozoa;Choanozoa;Choanof | EP00044 Salpingoeca rosetta 548 | 4,90E-30 | 2342 | XP 004987981.1 | hypothetical protein PTSG 11062 [Salpingoeca rosetta] | 2342 | 0 | 4790,33 | 100 |
| EP00746 | Sar;Alveolata;Myzozoa;"DP clade";Dinoflagellata;"I | EP00746 Dinoflagellata sp MGD 144431 | 5,20E-30 | 826 | ***no hit found*** | | | | | |
| EP00462 | Sar;Rhizaria;Cercozoa;"crown cercozoa";Thecofilose | EP00462 Mataza sp D1 7206 | 1,00E-29 | 909 | XP 023392200.1 | protein transport protein Sec16A isoform X2 [Pteropus vampyrus] | 2046 | 2,37E-08 | 70,0922 | 41,60583942 |
| EP00658 | Telonemia;Telonema;strain P-2 | EP00658 Telonema sp P-2 79806 | 1,80E-29 | 367 | XP 814906.1 | hypothetical protein [Trypanosoma cruzi strain CL Brener] | 1482 | 9,64E-12 | 77,7962 | 48,16326531 |
| EP00191 | Archaeplastida;Chloroplastida;Chlorophyta;"core ch | EP00191 Tetraselmis sp GSL018 8752 | 2,00E-29 | 562 | XP 024373338.1 | protein transport protein SEC16A homolog isoform X1 [Physcomitrella patens] >gi\|13738! | 1444 | 2,27E-26 | 125,561 | 44,23529412 |
| EP00599 | Sar;Stramenopiles;Gyrista;Ochrophyta;"CS clade";C | EP00599 Poterioochromonas sp BG-1 6636 | 6,40E-29 | 1785 | XP 008890960.1 | hypothetical protein PPTG 01118 [Phytophthora parasitica INRA-310] | 1621 | 1,78E-12 | 84,7297 | 54,02684564 |
| EP00376 | Sar;Alveolata;Myzozoa;"AC clade";Apicomplexa;"cc | EP00376 Eimeria tenella 7392 | 7,80E-29 | 1535 | XP 013235131.1 | hypothetical protein, conserved [Eimeria tenella] | 1535 | 0 | 2928,27 | 100 |
| EP00112 | Obazoa;Opisthokonta;Holozoa;Choanozoa;Metazoa | EP00112 Aurelia sp Birch-Aquarium-sp1 19266 | 1,80E-28 | 3380 | XP 028515408.1 | uncharacterized protein LOC110240827 [Exaiptasia pallida] | 11210 | 0 | 880,552 | 46,34146341 |
| EP00577 | Sar;Stramenopiles;Gyrista;Ochrophyta;"BD clade";[ | EP00577 Skeletonema sp RCC1716 2936 | 1,20E-27 | 939 | XP 002290923.1 | predicted protein [Thalassiosira pseudonana CCMP1335] | 1041 | 0 | 1293,87 | 74,34002112 |
| EP00160 | Obazoa;Opisthokonta;Nucletmycea;Rotosphaerida; | EP00160 Parvularia atlantis 20090 | 1,80E-27 | 270 | XP 017235227.1 | PREDICTED: protein transport protein SEC16A homolog [Daucus carota subsp. sativus] >gi | 1558 | 8,73E-07 | 60,8474 | 44,02985075 |
| EP00286 | Cryptista;Cryptophyceae;Cryptomonadales;"lineage | EP00286 Rhodomonas abbreviata 4251 | 2,60E-27 | 163 | XP 003061507.1 | predicted protein [Micromonas pusilla CCMP1545] | 1814 | 5,70E-15 | 81,6481 | 43,30708661 |
| EP00297 | Cryptista;Palpitomonas;Palpitomonas bilix;strain N | EP00297 Palpitomonas bilix 9239 | 4,20E-27 | 520 | XP 030469312.1 | protein transport protein SEC16A homolog isoform X2 [Syzygium oleosum] | 1444 | 2,72E-06 | 62,003 | 42,79835391 |
| EP00638 | Sar;Stramenopiles;Gyrista;Ochrophyta;Raphidophy | EP00638 Chattonella subsalsa 6320 | 4,60E-27 | 392 | XP 024584649.1 | Regucalcin gene promoter region-related protein (RGPR) [Plasmopara halstedii] | 2672 | 4,92E-17 | 94,3597 | 56,91489362 |
| EP00455 | Sar;Rhizaria;"eorhiza";Aquavolonida;"NC10 C lineag | EP00455 Lapot gusevi 17584 | 1,70E-26 | 123 | XP 004344939.1 | hypothetical protein ACA1 285550 [Acanthamoeba castellanii str. Neff] | 1515 | 9,82E-17 | 85,1149 | 51,42857143 |
| EP00376 | Sar;Alveolata;Myzozoa;"AC clade";Apicomplexa;"cc | EP00376 Eimeria tenella 4774 | 1,90E-26 | 1904 | XP 013232507.1 | hypothetical protein, conserved [Eimeria tenella] | 1904 | 0 | 3761,84 | 100 |
| EP00246 | Archaeplastida;Chloroplastida;Streptophyta;"core s | EP00246 Nothoceros aenigmaticus 10860 | 2,40E-26 | 200 | XP 028751820.1 | LOW QUALITY PROTEIN: protein transport protein SEC16A homolog [Prosopis alba] | 1393 | 1,90E-46 | 173,711 | 52,84974093 |
| EP00402 | Sar;Alveolata;Myzozoa;"DP clade";Dinoflagellata;D | EP00402 Amphidinium carterae 5886 | 2,80E-26 | 761 | XP 018001640.1 | COPII coat assembly protein sec16 [Phialophora attae] | 1858 | 1,89E-15 | 92,8189 | 42,20779221 |
| EP00569 | Sar;Stramenopiles;Gyrista;Ochrophyta;"BD clade";[ | EP00569 Conticribra weissflogii 8970 | 3,40E-26 | 911 | XP 002290923.1 | predicted protein [Thalassiosira pseudonana CCMP1335] | 1041 | 0 | 1142,49 | 73,31812999 |
| EP00376 | Sar;Alveolata;Myzozoa;"AC clade";Apicomplexa;"cc | EP00376 Eimeria tenella 1312 | 3,40E-26 | 2805 | XP 013229038.1 | zinc finger (C3HC4 RING finger) protein, putative [Eimeria tenella] | 2805 | 0 | 5314,97 | 100 |
| EP00575 | Sar;Stramenopiles;Gyrista;Ochrophyta;"BD clade";[ | EP00575 Skeletonema marinoi 29751 | 3,90E-26 | 1060 | XP 002290923.1 | predicted protein [Thalassiosira pseudonana CCMP1335] | 1041 | 0 | 1378,62 | 74,30939227 |
| EP00575 | Sar;Stramenopiles;Gyrista;Ochrophyta;"BD clade";[ | EP00575 Skeletonema marinoi 16834 | 4,10E-26 | 1098 | XP 002290923.1 | predicted protein [Thalassiosira pseudonana CCMP1335] | 1041 | 0 | 1429,85 | 73,96396396 |
| EP00579 | Sar;Stramenopiles;Gyrista;Ochrophyta;"BD clade";[ | EP00579 Thalassiosira antarctica 907 | 8,00E-26 | 1129 | XP 002290923.1 | predicted protein [Thalassiosira pseudonana CCMP1335] | 1041 | 0 | 1126,31 | 78,4217016 |
| EP00577 | Sar;Stramenopiles;Gyrista;Ochrophyta;"BD clade";[ | EP00577 Skeletonema sp RCC1716 2090 | 1,00E-25 | 1092 | XP 002290923.1 | predicted protein [Thalassiosira pseudonana CCMP1335] | 1041 | 0 | 1449,11 | 74,16137806 |
| EP00572 | Sar;Stramenopiles;Gyrista;Ochrophyta;"BD clade";[ | EP00572 Skeletonema dohrnii 1732 | 1,40E-25 | 1090 | XP 002290923.1 | predicted protein [Thalassiosira pseudonana CCMP1335] | 1041 | 0 | 1442,56 | 74,11444142 |
| EP00577 | Sar;Stramenopiles;Gyrista;Ochrophyta;"BD clade";[ | EP00577 Skeletonema sp RCC1716 2443 | 1,60E-25 | 1094 | XP 002290923.1 | predicted protein [Thalassiosira pseudonana CCMP1335] | 1041 | 0 | 1446,8 | 74,11764706 |
| EP00376 | Sar;Alveolata;Myzozoa;"AC clade";Apicomplexa;"cc | EP00376 Eimeria tenella 2049 | 2,30E-25 | 1711 | XP 013229775.1 | Niemann-Pick type C1 disease protein/ patched like cholesterol transporter of the SecD fa | 1711 | 0 | 3367,02 | 100 |
| EP00577 | Sar;Stramenopiles;Gyrista;Ochrophyta;"BD clade";[ | EP00577 Skeletonema sp RCC1716 5839 | 2,30E-25 | 779 | XP 002290923.1 | predicted protein [Thalassiosira pseudonana CCMP1335] | 1041 | 0 | 732,635 | 78,6036036 |
| EP00269 | Archaeplastida;Chloroplastida;Streptophyta;"core s | EP00269 Selaginella moellendorffii 35313 | 2,40E-25 | 230 | XP 024535959.1 | protein transport protein SEC16B homolog isoform X2 [Selaginella moellendorffii] | 230 | 1,54E-171 | 479,945 | 100 |
| EP00376 | Sar;Alveolata;Myzozoa;"AC clade";Apicomplexa;"cc | EP00376 Eimeria tenella 4818 | 2,40E-25 | 1848 | XP 013232551.1 | hypothetical protein, conserved [Eimeria tenella] | 1848 | 0 | 3575,41 | 100 |
| EP00267 | Archaeplastida;Chloroplastida;Streptophyta;"core s | EP00267 Zea mays 15533 | 2,90E-25 | 409 | NP 001348882.1 | uncharacterized protein LOC100383190 [Zea mays] | 1822 | 0 | 825,854 | 100 |
| EP00404 | Sar;Alveolata;Myzozoa;"DP clade";Dinoflagellata;D | EP00404 Azadinium spinosum 13883 | 7,20E-25 | 138 | XP 009523981.1 | hypothetical protein PHYSODRAFT 262855 [Phytophthora sojae] | 2808 | 1,45E-08 | 62,3882 | 45,34883721 |
| EP00585 | Sar;Stramenopiles;Gyrista;Ochrophyta;"BD clade";[ | EP00585 Thalassiosira rotula 6207 | 8,60E-25 | 1115 | XP 002290923.1 | predicted protein [Thalassiosira pseudonana CCMP1335] | 1041 | 0 | 1115,91 | 75,65543071 |
| EP00376 | Sar;Alveolata;Myzozoa;"AC clade";Apicomplexa;"cc | EP00376 Eimeria tenella 1012 | 9,10E-25 | 1882 | XP 013228738.1 | hypothetical protein, conserved [Eimeria tenella] | 1882 | 0 | 3657,46 | 100 |
| EP00541 | Sar;Stramenopiles;Gyrista;Ochrophyta;"BD clade";[ | EP00541 Cyclophora tenuis 3238 | 1,20E-24 | 494 | XP 002183614.1 | predicted protein [Phaeodactylum tricornutum CCAP 1055/1] | 2083 | 5,29E-31 | 138,658 | 62,35294118 |
| EP00376 | Sar;Alveolata;Myzozoa;"AC clade";Apicomplexa;"cc | EP00376 Eimeria tenella 4772 | 1,20E-24 | 2391 | XP 013232505.1 | hypothetical protein, conserved [Eimeria tenella] | 2391 | 0 | 4559,21 | 100 |
| EP00126 | Obazoa;Opisthokonta;Holozoa;Ichthyosporea;Rhinc | EP00126 Sphaerothecum destruens 12717 | 1,40E-24 | 99 | XP 004344939.1 | hypothetical protein ACA1 285550 [Acanthamoeba castellanii str. Neff] | 1515 | 1,03E-16 | 83,9593 | 39,39393939 |
| EP00151 | Obazoa;Opisthokonta;Nucletmycea;Fungi;"core Fur | EP00151 Rhizophagus irregularis 13060 | 1,70E-24 | 480 | XP 025177104.1 | hypothetical protein GLOIN 2v127817 [Rhizophagus irregularis DAOM 181602=DAOM 19 | 480 | 0 | 978,778 | 100 |
| EP00376 | Sar;Alveolata;Myzozoa;"AC clade";Apicomplexa;"cc | EP00376 Eimeria tenella 1764 | 2,10E-24 | 2467 | XP 013229490.1 | hypothetical protein, conserved [Eimeria tenella] | 2467 | 0 | 4714,44 | 100 |
| EP00376 | Sar;Alveolata;Myzozoa;"AC clade";Apicomplexa;"cc | EP00376 Eimeria tenella 6435 | 2,30E-24 | 2223 | XP 013234168.1 | Possible conserved eukaryotic alpha beta hydrolase, related [Eimeria tenella] | 2223 | 0 | 4364,68 | 100 |
| EP00571 | Sar;Stramenopiles;Gyrista;Ochrophyta;"BD clade";[ | EP00571 Detonula confervacea 2408 | 2,30E-24 | 1132 | XP 002290923.1 | predicted protein [Thalassiosira pseudonana CCMP1335] | 1041 | 0 | 1134,4 | 76,89440994 |
| EP00264 | Archaeplastida;Chloroplastida;Streptophyta;"core s | EP00264 Hordeum vulgare 57091 | 2,70E-24 | 866 | XP 020187483.1 | protein transport protein SEC31 homolog B isoform X1 [Aegilops tauschii subsp. tauschii] | 1135 | 0 | 1621,29 | 94,57900807 |
| EP00500 | Sar;Stramenopiles;Bigyra;Opalozoa;Bicosoecida;str | EP00500 Bicosoecida sp ms1 11052 | 2,80E-24 | 99 | XP 004344939.1 | hypothetical protein ACA1 285550 [Acanthamoeba castellanii str. Neff] | 1515 | 5,15E-13 | 73,559 | 44,44444444 |
| EP00264 | Archaeplastida;Chloroplastida;Streptophyta;"core s | EP00264 Hordeum vulgare 57085 | 2,90E-24 | 889 | XP 020187484.1 | protein transport protein SEC31 homolog B isoform X2 [Aegilops tauschii subsp. tauschii] | 1134 | 0 | 1626,3 | 94,38073394 |
| EP00264 | Archaeplastida;Chloroplastida;Streptophyta;"core s | EP00264 Hordeum vulgare 57070 | 2,90E-24 | 907 | XP 020187484.1 | protein transport protein SEC31 homolog B isoform X2 [Aegilops tauschii subsp. tauschii] | 1134 | 0 | 1707,96 | 94,82378855 |
| EP00264 | Archaeplastida;Chloroplastida;Streptophyta;"core s | EP00264 Hordeum vulgare 57084 | 2,90E-24 | 907 | XP 020187484.1 | protein transport protein SEC31 homolog B isoform X2 [Aegilops tauschii subsp. tauschii] | 1134 | 0 | 1707,96 | 94,82378855 |
| EP00264 | Archaeplastida;Chloroplastida;Streptophyta;"core s | EP00264 Hordeum vulgare 57087 | 2,90E-24 | 907 | XP 020187484.1 | protein transport protein SEC31 homolog B isoform X2 [Aegilops tauschii subsp. tauschii] | 1134 | 0 | 1707,96 | 94,82378855 |
| EP00264 | Archaeplastida;Chloroplastida;Streptophyta;"core s | EP00264 Hordeum vulgare 57093 | 2,90E-24 | 907 | XP 020187484.1 | protein transport protein SEC31 homolog B isoform X2 [Aegilops tauschii subsp. tauschii] | 1134 | 0 | 1707,96 | 94,82378855 |
| EP00376 | Sar;Alveolata;Myzozoa;"AC clade";Apicomplexa;"cc | EP00376 Eimeria tenella 6198 | 3,00E-24 | 3847 | XP 013233931.1 | hypothetical protein, conserved [Eimeria tenella] | 3847 | 0 | 7634,64 | 100 |
| EP00264 | Archaeplastida;Chloroplastida;Streptophyta;"core s | EP00264 Hordeum vulgare 57094 | 3,00E-24 | 1030 | XP 020187483.1 | protein transport protein SEC31 homolog B isoform X1 [Aegilops tauschii subsp. tauschii] | 1135 | 0 | 1942,55 | 95,49718574 |
| EP00264 | Archaeplastida;Chloroplastida;Streptophyta;"core s | EP00264 Hordeum vulgare 57072 | 3,20E-24 | 941 | XP 020187483.1 | protein transport protein SEC31 homolog B isoform X1 [Aegilops tauschii subsp. tauschii] | 1135 | 0 | 1773,44 | 94,79277365 |
| EP00264 | Archaeplastida;Chloroplastida;Streptophyta;"core s | EP00264 Hordeum vulgare 57096 | 3,20E-24 | 941 | XP 020187483.1 | protein transport protein SEC31 homolog B isoform X1 [Aegilops tauschii subsp. tauschii] | 1135 | 0 | 1768,82 | 94,98933902 |
| EP00264 | Archaeplastida;Chloroplastida;Streptophyta;"core s | EP00264 Hordeum vulgare 57081 | 3,20E-24 | 942 | XP 020187483.1 | protein transport protein SEC31 homolog B isoform X1 [Aegilops tauschii subsp. tauschii] | 1135 | 0 | 1771,52 | 94,98933902 |
| EP00264 | Archaeplastida;Chloroplastida;Streptophyta;"core s | EP00264 Hordeum vulgare 57086 | 3,70E-24 | 1011 | XP 020187483.1 | protein transport protein SEC31 homolog B isoform X1 [Aegilops tauschii subsp. tauschii] | 1135 | 0 | 1923,67 | 95,82929195 |
| EP00264 | Archaeplastida;Chloroplastida;Streptophyta;"core s | EP00264 Hordeum vulgare 57083 | 3,80E-24 | 1032 | XP 020187483.1 | protein transport protein SEC31 homolog B isoform X1 [Aegilops tauschii subsp. tauschii] | 1135 | 0 | 1967,97 | 95,35333979 |
| EP00264 | Archaeplastida;Chloroplastida;Streptophyta;"core s | EP00264 Hordeum vulgare 57074 | 4,10E-24 | 1060 | XP 020187483.1 | protein transport protein SEC31 homolog B isoform X1 [Aegilops tauschii subsp. tauschii] | 1135 | 0 | 1945,24 | 95,10166359 |
| EP00264 | Archaeplastida;Chloroplastida;Streptophyta;"core s | EP00264 Hordeum vulgare 57078 | 4,30E-24 | 1088 | XP 020187483.1 | protein transport protein SEC31 homolog B isoform X1 [Aegilops tauschii subsp. tauschii] | 1135 | 0 | 2011,11 | 94,9168207 |
| EP00376 | Sar;Alveolata;Myzozoa;"AC clade";Apicomplexa;"cc | EP00376 Eimeria tenella 1197 | 5,40E-24 | 2080 | XP 013228923.1 | phosphatidylinositol 3-kinase, putative [Eimeria tenella] | 2080 | 0 | 4182,87 | 100 |
| EP00574 | Sar;Stramenopiles;Gyrista;Ochrophyta;"BD clade";[ | EP00574 Skeletonema japonicum 9291 | 9,70E-24 | 1113 | XP 002290923.1 | predicted protein [Thalassiosira pseudonana CCMP1335] | 1041 | 0 | 1397,49 | 72,89211242 |
| EP00262 | Archaeplastida;Chloroplastida;Streptophyta;"core s | EP00262 Sarcandra glabra 5849 | 1,20E-23 | 116 | XP 003516666.1 | protein transport protein SEC16B homolog [Glycine max] | 1404 | 3,79E-54 | 191,815 | 76,72413793 |
| EP00746 | Sar;Alveolata;Myzozoa;"DP clade";Dinoflagellata;"I | EP00746 Dinoflagellata sp MGD 56450 | 1,50E-23 | 209 | XP 009523981.1 | hypothetical protein PHYSODRAFT 262855 [Phytophthora sojae] | 2808 | 1,05E-09 | 68,1662 | 43,44262295 |
| EP00376 | Sar;Alveolata;Myzozoa;"AC clade";Apicomplexa;"cc | EP00376 Eimeria tenella 2964 | 1,70E-23 | 3901 | XP 013230694.1 | hypothetical protein, conserved [Eimeria tenella] | 3901 | 0 | 7644,27 | 100 |
| EP00576 | Sar;Stramenopiles;Gyrista;Ochrophyta;"BD clade";[ | EP00576 Skeletonema menzelii 4606 | 1,80E-23 | 1082 | XP 002290923.1 | predicted protein [Thalassiosira pseudonana CCMP1335] | 1041 | 0 | 1439,09 | 73,1261426 |
| EP00263 | Archaeplastida;Chloroplastida;Streptophyta;"core s | EP00263 Brachypodium distachyon 3407 | 1,80E-23 | 1121 | XP 003562538.1 | protein transport protein SEC31 homolog B [Brachypodium distachyon] | 1121 | 0 | 2313,88 | 100 |
| EP00173 | Archaeplastida;"RR clade";Rhodophyta;Eurhodophy | EP00173 Palmaria palmata 2590 | 1,90E-23 | 403 | XP 005711041.1 | unnamed protein product [Chondrus crispus] | 2546 | 4,73E-15 | 88,5817 | 37,07482993 |
| EP00437 | Sar;Alveolata;Myzozoa;"DP clade";Dinoflagellata;D | EP00437 Effrenium voratum 57552 | 1,90E-23 | 189 | XP 016606650.1 | hypothetical protein SPPG 06291 [Spizellomyces punctatus DAOM BR117] | 2211 | 3,55E-09 | 65,855 | 36,31578947 |
| EP00019 | Amoebozoa;Evosea;Cutosea;Armaparvus;Armaparv | EP00019 Armaparvus languidus 1385 | 2,10E-23 | 110 | XP 004344939.1 | hypothetical protein ACA1 285550 [Acanthamoeba castellanii str. Neff] | 1515 | 1,23E-22 | 101,293 | 45,87155963 |
| EP00256 | Archaeplastida;Chloroplastida;Streptophyta;"core s | EP00256 Glycine max 20553 | 2,40E-23 | 1113 | XP 003539884.2 | protein transport protein SEC31 homolog B [Glycine max] | 1113 | 0 | 2300,4 | 100 |
| EP00264 | Archaeplastida;Chloroplastida;Streptophyta;"core s | EP00264 Hordeum vulgare 57095 | 3,50E-23 | 899 | XP 020187484.1 | protein transport protein SEC31 homolog B isoform X2 [Aegilops tauschii subsp. tauschii] | 1134 | 0 | 1602,03 | 94,77777778 |
| EP00264 | Archaeplastida;Chloroplastida;Streptophyta;"core s | EP00264 Hordeum vulgare 57082 | 3,50E-23 | 897 | XP 020187483.1 | protein transport protein SEC31 homolog B isoform X1 [Aegilops tauschii subsp. tauschii] | 1135 | 0 | 1606,65 | 94,76614699 |
| EP00264 | Archaeplastida;Chloroplastida;Streptophyta;"core s | EP00264 Hordeum vulgare 57080 | 3,70E-23 | 920 | XP 020187484.1 | protein transport protein SEC31 homolog B isoform X2 [Aegilops tauschii subsp. tauschii] | 1134 | 0 | 1610,89 | 94,57364341 |
| EP00264 | Archaeplastida;Chloroplastida;Streptophyta;"core s | EP00264 Hordeum vulgare 57077 | 3,90E-23 | 938 | XP 020187484.1 | protein transport protein SEC31 homolog B isoform X2 [Aegilops tauschii subsp. tauschii] | 1134 | 0 | 1691,4 | 94,99467519 |
| EP00264 | Archaeplastida;Chloroplastida;Streptophyta;"core s | EP00264 Hordeum vulgare 57097 | 3,90E-23 | 938 | XP 020187484.1 | protein transport protein SEC31 homolog B isoform X2 [Aegilops tauschii subsp. tauschii] | 1134 | 0 | 1691,4 | 94,99467519 |
| EP00284 | Cryptista;Cryptophyceae;Cryptomonadales;"lineage | EP00284 Hemiselmis tepida 12874 | 4,80E-23 | 80 | XP 002975264.1 | protein transport protein SEC16B homolog [Selaginella moellendorffii] | 1190 | 9,01E-13 | 71,633 | 47,36842105 |
| EP00264 | Archaeplastida;Chloroplastida;Streptophyta;"core s | EP00264 Hordeum vulgare 57089 | 4,90E-23 | 1063 | XP 020187483.1 | protein transport protein SEC31 homolog B isoform X1 [Aegilops tauschii subsp. tauschii] | 1135 | 0 | 1952,56 | 95,4887218 |
| EP00160 | Obazoa;Opisthokonta;Nucletmycea;Rotosphaerida; | EP00160 Parvularia atlantis 21828 | 5,50E-23 | 189 | XP 002624527.1 | COPII coat assembly protein SEC16 [Blastomyces gilchristii SLH14081] | 1776 | 1,13E-11 | 73,1738 | 44,05594406 |
| EP00170 | Archaeplastida;"RR clade";Rhodophyta;Eurhodophy | EP00170 Pyropia yezoensis 7489 | 8,40E-23 | 789 | ***no hit found*** | | | | | |
| EP00265 | Archaeplastida;Chloroplastida;Streptophyta;"core s | EP00265 Oryza sativa 16563 | 1,10E-22 | 1127 | XP 015646666.1 | protein transport protein SEC31 homolog B [Oryza sativa Japonica Group] | 1127 | 0 | 2329,29 | 100 |
| EP00589 | Sar;Stramenopiles;Gyrista;Ochrophyta;"BD clade";[ | EP00589 Leptocylindrus danicus 13574 | 1,10E-22 | 1052 | XP 002290923.1 | predicted protein [Thalassiosira pseudonana CCMP1335] | 1041 | 0 | 914,835 | 52,39410681 |
| EP00376 | Sar;Alveolata;Myzozoa;"AC clade";Apicomplexa;"cc | EP00376 Eimeria tenella 2810 | 1,20E-22 | 3306 | XP 013230538.1 | leucine rich repeat containing protein / ankyrin repeat containing protein, putative [Eimer | 3306 | 0 | 6433,59 | 100 |
| EP00023 | Amoebozoa;Evosea;Eumycetozoa;Dictyostelia;Dicty | EP00023 Dictyostelium discoideum 6370 | 1,20E-22 | 4582 | XP 640504.5 | protein kinase, Atypical group [Dictyostelium discoideum AX4] | 4582 | 0 | 9348,01 | 100 |
| EP00054 | Obazoa;Opisthokonta;Holozoa;Choanozoa;Choanof | EP00054 Salpingoeca dolichothecata 27048 | 1,30E-22 | 1714 | XP 004994749.1 | hypothetical protein PTSG 12199 [Salpingoeca rosetta] | 796 | 2,46E-07 | 67,3958 | 37,74509804 |
| EP00573 | Sar;Stramenopiles;Gyrista;Ochrophyta;"BD clade";[ | EP00573 Skeletonema grethae 7300 | 1,40E-22 | 1092 | XP 002290923.1 | predicted protein [Thalassiosira pseudonana CCMP1335] | 1041 | 0 | 1410,97 | 72,71906052 |
| EP00537 | Sar;Stramenopiles;Gyrista;Ochrophyta;"BD clade";[ | EP00537 Pseudo-nitzschia pungens 6117 | 1,60E-22 | 1029 | XP 002290923.1 | predicted protein [Thalassiosira pseudonana CCMP1335] | 1041 | 0 | 929,858 | 51,89753321 |
| EP00529 | Sar;Stramenopiles;Gyrista;Ochrophyta;"BD clade";[ | EP00529 Nitzschia sp RCC80 14341 | 1,90E-22 | 718 | XP 002290923.1 | predicted protein [Thalassiosira pseudonana CCMP1335] | 1041 | 0 | 588,956 | 48,10810811 |
| EP00376 | Sar;Alveolata;Myzozoa;"AC clade";Apicomplexa;"cc | EP00376 Eimeria tenella 436 | 2,20E-22 | 1750 | XP 013228162.1 | hypothetical protein, conserved [Eimeria tenella] | 1750 | 0 | 3339,28 | 100 |
| EP00454 | Sar;Alveolata;Myzozoa;"DP clade";Perkinsea;Perkin | EP00454 Perkinsus marinus 12682 | 2,50E-22 | 197 | XP 002777968.1 | hypothetical protein Pmar PMAR023945 [Perkinsus marinus ATCC 50983] | 197 | 9,37E-143 | 404,445 | 100 |
| EP00581 | Sar;Stramenopiles;Gyrista;Ochrophyta;"BD clade";[ | EP00581 Thalassiosira minuscula 9890 | 2,80E-22 | 1118 | XP 002290923.1 | predicted protein [Thalassiosira pseudonana CCMP1335] | 1041 | 0 | 1104,35 | 75,06361323 |
| EP00578 | Sar;Stramenopiles;Gyrista;Ochrophyta;"BD clade";[ | EP00578 Thalassiosira sp NH16 6546 | 2,90E-22 | 1086 | XP 002290923.1 | predicted protein [Thalassiosira pseudonana CCMP1335] | 1041 | 0 | 1431 | 74,17883212 |
| EP00578 | Sar;Stramenopiles;Gyrista;Ochrophyta;"BD clade";[ | EP00578 Thalassiosira sp NH16 8747 | 2,90E-22 | 1086 | XP 002290923.1 | predicted protein [Thalassiosira pseudonana CCMP1335] | 1041 | 0 | 1431 | 74,17883212 |
| EP00767 | Metamonada;"BF clade";Fornicata;Chilomastix;Chil | EP00767 Chilomastix cuspidata 2650 | 5,40E-22 | 779 | ***no hit found*** | | | | | |

| | | | | | | | | | | |
|---|---|---|---|---|---|---|---|---|---|---|
| EP00736 | Archaeplastida;"RR clade";Rhodelphis;Rhodelphis n | EP00736 Rhodelphis marinus 9419 | 7,90E-22 | 209 | XP 019860577.1 | PREDICTED: protein transport protein Sec16A-like [Amphimedon queenslandica] | 1105 | 1,03E-07 | 62,3882 | 36,875 |
| EP00233 | Archaeplastida;Chloroplastida;Palmophyllophyceae | EP00233 Prasinococcus capsulatus 8292 | 8,50E-22 | 1173 | XP 028797788.1 | protein transport protein SEC31 homolog B-like [Prosopis alba] >gi\|1624055540\|ref\|XP ( | 1107 | 0 | 689,108 | 51,16751269 |
| EP00376 | Sar;Alveolata;Myzozoa;"AC clade";Apicomplexa;"cc | EP00376 Eimeria tenella 2334 | 9,40E-22 | 1786 | XP 013230061.1 | importin alpha, putative [Eimeria tenella] | 1786 | 0 | 3493,75 | 100 |
| EP00022 | Amoebozoa;Evosea;Eumycetozoa;Dictyostelia;Acyt | EP00022 Cavenderia fasciculata 3380 | 1,20E-21 | 1147 | XP 004355165.1 | WD40 repeat-containing protein [Cavenderia fasciculata] | 1147 | 0 | 2356,25 | 100 |
| EP00529 | Sar;Stramenopiles;Gyrista;Ochrophyta;"BD clade";[ | EP00529 Nitzschia sp RCC80 16201 | 1,30E-21 | 669 | XP 002290923.1 | predicted protein [Thalassiosira pseudonana CCMP1335] | 1041 | 3,38E-180 | 549,28 | 48,45814978 |
| EP00286 | Cryptista;Cryptophyceae;Cryptomonadales;"lineage | EP00286 Rhodomonas abbreviata 25336 | 1,30E-21 | 447 | ***no hit found*** | | | | | |
| EP00301 | Haptista;Centroplasthelida;Pterocystida;Raphidista | EP00301 Raphidiophrys heterophryoidea 16156 | 1,40E-21 | 175 | XP 020431748.1 | hypothetical protein PPL 07486 [Heterostelium album PN500] | 1678 | 2,27E-07 | 60,077 | 40,18691589 |
| EP00612 | Sar;Stramenopiles;Gyrista;Ochrophyta;"PX clade";X | EP00612 Vaucheria litorea 6404 | 1,50E-21 | 458 | XP 024584649.1 | Regucalcin gene promoter region-related protein (RGPR) [Plasmopara halstedii] | 2672 | 1,24E-15 | 91,2781 | 64,81481481 |
| EP00583 | Sar;Stramenopiles;Gyrista;Ochrophyta;"BD clade";[ | EP00583 Thalassiosira pseudonana 4989 | 2,10E-21 | 1041 | XP 002290923.1 | predicted protein [Thalassiosira pseudonana CCMP1335] | 1041 | 0 | 2156,72 | 100 |
| EP00770 | Metamonada;Preaxostyla;Oxymonadida;Monocerc | EP00770 Monocercomonoides exilis 13625 | 2,40E-21 | 4007 | XP 026291516.1 | obscurin-like isoform X4 [Frankliniella occidentalis] | 6099 | 2,76E-09 | 75,8702 | 39,23076923 |
| EP00738 | Archaeplastida;Chloroplastida;Chlorophyta;Chlorop | EP00738 Chloropicon primus 7215 | 2,60E-21 | 1137 | XP 006837403.2 | protein transport protein SEC31 homolog B isoform X1 [Amborella trichopoda] | 1155 | 0 | 741,88 | 50,91210614 |
| EP00021 | Amoebozoa;Evosea;Eumycetozoa;Dictyostelia;Acyt | EP00021 Heterostelium album 375 | 3,00E-21 | 836 | XP 020427011.1 | WD40 repeat-containing protein [Heterostelium album PN500] | 836 | 0 | 1731,46 | 100 |
| EP00536 | Sar;Stramenopiles;Gyrista;Ochrophyta;"BD clade";[ | EP00536 Pseudo-nitzschia multiseries 3977 | 3,20E-21 | 1036 | XP 002290923.1 | predicted protein [Thalassiosira pseudonana CCMP1335] | 1041 | 0 | 934,48 | 53,32708529 |
| EP00529 | Sar;Stramenopiles;Gyrista;Ochrophyta;"BD clade";[ | EP00529 Nitzschia sp RCC80 10748 | 3,50E-21 | 638 | XP 002290923.1 | predicted protein [Thalassiosira pseudonana CCMP1335] | 1041 | 8,61E-162 | 500,36 | 46,95121951 |
| EP00260 | Archaeplastida;Chloroplastida;Streptophyta;"core s | EP00260 Arabidopsis thaliana 9055 | 3,50E-21 | 963 | NP 001320897.1 | Transducin/WD40 repeat-like superfamily protein [Arabidopsis thaliana] >gi\|1063684652 | 963 | 0 | 2005,33 | 100 |
| EP00260 | Archaeplastida;Chloroplastida;Streptophyta;"core s | EP00260 Arabidopsis thaliana 9056 | 3,50E-21 | 963 | NP 001320897.1 | Transducin/WD40 repeat-like superfamily protein [Arabidopsis thaliana] >gi\|1063684652 | 963 | 0 | 2005,33 | 100 |
| EP00445 | Sar;Alveolata;Myzozoa;"DP clade";Dinoflagellata;D | EP00445 Apocalathium hangoei 40645 | 4,10E-21 | 710 | ***no hit found*** | | | | | |
| EP00529 | Sar;Stramenopiles;Gyrista;Ochrophyta;"BD clade";[ | EP00529 Nitzschia sp RCC80 6078 | 4,20E-21 | 1008 | XP 002290923.1 | predicted protein [Thalassiosira pseudonana CCMP1335] | 1041 | 0 | 838,951 | 48,64864865 |
| EP00529 | Sar;Stramenopiles;Gyrista;Ochrophyta;"BD clade";[ | EP00529 Nitzschia sp RCC80 5931 | 4,40E-21 | 1039 | XP 002290923.1 | predicted protein [Thalassiosira pseudonana CCMP1335] | 1041 | 0 | 885,174 | 49,48162111 |
| EP00258 | Archaeplastida;Chloroplastida;Streptophyta;"core s | EP00258 Populus trichocarpa 8084 | 4,60E-21 | 1135 | XP 002313327.2 | protein transport protein SEC31 homolog B [Populus trichocarpa] | 1135 | 0 | 2344,7 | 100 |
| EP00213 | Archaeplastida;Chloroplastida;Chlorophyta;Chlorop | EP00213 Chloropicon mariensis 530 | 4,70E-21 | 1098 | XP 006837403.2 | protein transport protein SEC31 homolog B isoform X1 [Amborella trichopoda] | 1155 | 0 | 705,671 | 51,98961938 |
| EP00376 | Sar;Alveolata;Myzozoa;"AC clade";Apicomplexa;"cc | EP00376 Eimeria tenella 1042 | 6,60E-21 | 1763 | XP 013228768.1 | SOH1 domain-containing protein, putative [Eimeria tenella] | 1763 | 0 | 3271,1 | 100 |
| EP00613 | Sar;Stramenopiles;Gyrista;Ochrophyta;Dictyochoph | EP00613 Pedinella sp CCMP2098 21931 | 7,40E-21 | 2033 | ***no hit found*** | | | | | |
| EP00260 | Archaeplastida;Chloroplastida;Streptophyta;"core s | EP00260 Arabidopsis thaliana 22312 | 8,90E-21 | 969 | NP 173317.2 | Transducin/WD40 repeat-like superfamily protein [Arabidopsis thaliana] | 969 | 0 | 2017,66 | 100 |
| EP00245 | Archaeplastida;Chloroplastida;Streptophyta;"core s | EP00245 Coleochaete scutata 5205 | 9,30E-21 | 1133 | XP 024379159.1 | protein transport protein SEC31 homolog B-like isoform X2 [Physcomitrella patens] | 1166 | 0 | 1063,91 | 58,20770519 |
| EP00376 | Sar;Alveolata;Myzozoa;"AC clade";Apicomplexa;"cc | EP00376 Eimeria tenella 225 | 1,10E-20 | 1602 | XP 013227951.1 | Condensin subunit, related [Eimeria tenella] | 1602 | 0 | 3044,6 | 100 |
| EP00215 | Archaeplastida;Chloroplastida;Chlorophyta;Chlorop | EP00215 Chloropicon roscoffensis 11657 | 1,20E-20 | 1158 | XP 020519257.1 | protein transport protein SEC31 homolog B isoform X2 [Amborella trichopoda] | 1154 | 0 | 750,74 | 53,54785479 |
| EP00245 | Archaeplastida;Chloroplastida;Streptophyta;"core s | EP00245 Coleochaete scutata 5204 | 1,30E-20 | 1125 | XP 024379159.1 | protein transport protein SEC31 homolog B-like isoform X2 [Physcomitrella patens] | 1166 | 0 | 1067,76 | 58,24915825 |
| EP00376 | Sar;Alveolata;Myzozoa;"AC clade";Apicomplexa;"cc | EP00376 Eimeria tenella 5940 | 1,60E-20 | 2110 | XP 013233673.1 | hypothetical protein, conserved [Eimeria tenella] | 2110 | 0 | 4180,56 | 100 |
| EP00376 | Sar;Alveolata;Myzozoa;"AC clade";Apicomplexa;"cc | EP00376 Eimeria tenella 3566 | 1,70E-20 | 2135 | XP 013231297.1 | protein kinase (incomplete catalytic triad), putative [Eimeria tenella] | 2135 | 0 | 4332,33 | 100 |
| EP00283 | Cryptista;Cryptophyceae;Cryptomonadales;"lineage | EP00283 Hemiselmis rufescens 1040 | 1,70E-20 | 369 | ***no hit found*** | | | | | |
| EP00668 | Discoba;Euglenozoa;Euglenida;Spirocuta;Euglenea;I | EP00668 Euglena longa 39649 | 2,10E-20 | 115 | XP 008608960.1 | hypothetical protein SDRG 05043 [Saprolegnia diclina VS20] | 2628 | 5,35E-08 | 59,6918 | 45,96774194 |
| EP00529 | Sar;Stramenopiles;Gyrista;Ochrophyta;"BD clade";[ | EP00529 Nitzschia sp RCC80 6158 | 2,30E-20 | 1008 | XP 002290923.1 | predicted protein [Thalassiosira pseudonana CCMP1335] | 1041 | 0 | 835,869 | 48,64864865 |
| EP00089 | Obazoa;Opisthokonta;Holozoa;Choanozoa;Metazoa | EP00089 Calanus finmarchicus 93274 | 2,40E-20 | 287 | XP 023346731.1 | protein transport protein Sec16-like isoform X2 [Eurytemora affinis] | 1941 | 3,04E-31 | 133,65 | 59,22330097 |
| EP00089 | Obazoa;Opisthokonta;Holozoa;Choanozoa;Metazoa | EP00089 Calanus finmarchicus 93280 | 2,40E-20 | 287 | XP 023346731.1 | protein transport protein Sec16-like isoform X2 [Eurytemora affinis] | 1941 | 3,04E-31 | 133,65 | 59,22330097 |
| EP00521 | Sar;Stramenopiles;Gyrista;Ochrophyta;"BD clade";[ | EP00521 Asterionellopsis glacialis 9074 | 3,00E-20 | 1056 | XP 002290923.1 | predicted protein [Thalassiosira pseudonana CCMP1335] | 1041 | 0 | 1042,72 | 59,5505618 |
| EP00217 | Archaeplastida;Chloroplastida;Chlorophyta;Mamiel | EP00217 Crustomastix stigmatica 518 | 3,20E-20 | 1048 | XP 001420597.1 | predicted protein, partial [Ostreococcus lucimarinus CCE9901] | 1077 | 0 | 830,861 | 58,18399044 |
| EP00212 | Archaeplastida;Chloroplastida;Chlorophyta;Chlorop | EP00212 Chloropicon laureae 8145 | 3,70E-20 | 1177 | XP 006837403.2 | protein transport protein SEC31 homolog B isoform X1 [Amborella trichopoda] | 1155 | 0 | 672,159 | 51,23966942 |
| EP00267 | Archaeplastida;Chloroplastida;Streptophyta;"core s | EP00267 Zea mays 33490 | 3,80E-20 | 1128 | XP 008667827.1 | uncharacterized protein LOC100279874 isoform X1 [Zea mays] | 1128 | 0 | 2331,21 | 100 |
| EP00267 | Archaeplastida;Chloroplastida;Streptophyta;"core s | EP00267 Zea mays 8600 | 3,80E-20 | 1129 | NP 001146299.1 | uncharacterized protein LOC100279874 [Zea mays] | 1129 | 0 | 2333,14 | 100 |
| EP00123 | Obazoa;Opisthokonta;Holozoa;Ichthyosporea;Ichth | EP00123 Amoebidium parasiticum 16335 | 4,00E-20 | 150 | XP 011270238.1 | hypothetical protein, variant [Capsaspora owczarzaki ATCC 30864] | 1951 | 1,91E-07 | 59,6918 | 66,66666667 |
| EP00573 | Sar;Stramenopiles;Gyrista;Ochrophyta;"BD clade";[ | EP00573 Skeletonema grethae 10705 | 4,10E-20 | 128 | XP 002183614.1 | predicted protein [Phaeodactylum tricornutum CCAP 1055/1] | 2083 | 6,71E-12 | 71,633 | 41,02564103 |
| EP00269 | Archaeplastida;Chloroplastida;Streptophyta;"core s | EP00269 Selaginella moellendorffii 35312 | 4,30E-20 | 246 | XP 024535958.1 | protein transport protein SEC16B homolog isoform X1 [Selaginella moellendorffii] | 246 | 0 | 514,227 | 100 |
| EP00023 | Amoebozoa;Evosea;Eumycetozoa;Dictyostelia;Dicty | EP00023 Dictyostelium discoideum 8371 | 4,90E-20 | 2625 | XP 642625.1 | hypothetical protein DDB G0277419 [Dictyostelium discoideum AX4] | 2625 | 0 | 5239,47 | 100 |
| EP00266 | Archaeplastida;Chloroplastida;Streptophyta;"core s | EP00266 Sorghum bicolor 27605 | 4,90E-20 | 1127 | XP 021310431.1 | protein transport protein SEC31 homolog B isoform X2 [Sorghum bicolor] | 1127 | 0 | 2325,05 | 100 |
| EP00266 | Archaeplastida;Chloroplastida;Streptophyta;"core s | EP00266 Sorghum bicolor 27604 | 4,90E-20 | 1128 | XP 021310430.1 | protein transport protein SEC31 homolog B isoform X1 [Sorghum bicolor] | 1128 | 0 | 2326,21 | 100 |
| EP00254 | Archaeplastida;Chloroplastida;Streptophyta;"core s | EP00254 Amborella trichopoda 2356 | 5,20E-20 | 1155 | XP 006837403.2 | protein transport protein SEC31 homolog B isoform X1 [Amborella trichopoda] | 1155 | 0 | 2385,14 | 100 |
| EP00254 | Archaeplastida;Chloroplastida;Streptophyta;"core s | EP00254 Amborella trichopoda 18419 | 5,30E-20 | 1154 | XP 020519257.1 | protein transport protein SEC31 homolog B isoform X2 [Amborella trichopoda] | 1154 | 0 | 2383,22 | 100 |
| EP00297 | Cryptista;Palpitomonas;Palpitomonas bilix;strain N | EP00297 Palpitomonas bilix 1177 | 5,70E-20 | 1112 | XP 005831033.1 | secretory protein Sec31 [Guillardia theta CCMP2712] | 1081 | 3,95E-130 | 433,335 | 46,68246445 |
| EP00297 | Cryptista;Palpitomonas;Palpitomonas bilix;strain N | EP00297 Palpitomonas bilix 1286 | 5,80E-20 | 1123 | XP 005831033.1 | secretory protein Sec31 [Guillardia theta CCMP2712] | 1081 | 3,95E-139 | 457,603 | 46,8339307 |
| EP00267 | Archaeplastida;Chloroplastida;Streptophyta;"core s | EP00267 Zea mays 25822 | 5,80E-20 | 1129 | XP 008653419.1 | protein transport protein SEC31 homolog B isoform X2 [Zea mays] | 1129 | 0 | 2328,13 | 100 |
| EP00267 | Archaeplastida;Chloroplastida;Streptophyta;"core s | EP00267 Zea mays 25821 | 5,80E-20 | 1130 | XP 008653418.1 | protein transport protein SEC31 homolog B isoform X1 [Zea mays] | 1130 | 0 | 2330,06 | 100 |
| EP00297 | Cryptista;Palpitomonas;Palpitomonas bilix;strain N | EP00297 Palpitomonas bilix 1369 | 5,90E-20 | 1135 | XP 005831033.1 | secretory protein Sec31 [Guillardia theta CCMP2712] | 1081 | 1,21E-139 | 459,529 | 46,56398104 |
| EP00269 | Archaeplastida;Chloroplastida;Streptophyta;"core s | EP00269 Selaginella moellendorffii 35315 | 6,40E-20 | 228 | XP 024535961.1 | protein transport protein SEC16B homolog isoform X3 [Selaginella moellendorffii] | 228 | 2,31E-170 | 476,863 | 100 |
| EP00535 | Sar;Stramenopiles;Gyrista;Ochrophyta;"BD clade";[ | EP00535 Pseudo-nitzschia heimii 9026 | 6,80E-20 | 1037 | XP 002290923.1 | predicted protein [Thalassiosira pseudonana CCMP1335] | 1041 | 0 | 875,93 | 51,14503817 |
| EP00567 | Sar;Stramenopiles;Gyrista;Ochrophyta;"BD clade";[ | EP00567 Triceratium dubium 13754 | 7,40E-20 | 108 | XP 002183614.1 | predicted protein [Phaeodactylum tricornutum CCAP 1055/1] | 2083 | 1,34E-16 | 83,9593 | 43,92523364 |
| EP00570 | Sar;Stramenopiles;Gyrista;Ochrophyta;"BD clade";[ | EP00570 Cyclotella meneghiniana 9440 | 8,40E-20 | 1073 | XP 002290923.1 | predicted protein [Thalassiosira pseudonana CCMP1335] | 1041 | 0 | 1323,92 | 68,20702403 |
| EP00245 | Archaeplastida;Chloroplastida;Streptophyta;"core s | EP00245 Coleochaete scutata 17159 | 9,00E-20 | 190 | XP 016436112.1 | PREDICTED: protein transport protein SEC16B homolog [Nicotiana tabacum] | 289 | 2,61E-21 | 98,2117 | 49,72375691 |
| EP00658 | Telonemia;Telonema;strain P-2 | EP00658 Telonema sp P-2 54001 | 1,00E-19 | 275 | ***no hit found*** | | | | | |
| EP00533 | Sar;Stramenopiles;Gyrista;Ochrophyta;"BD clade";[ | EP00533 Pseudo-nitzschia delicatissima 282 | 1,10E-19 | 1027 | XP 002290923.1 | predicted protein [Thalassiosira pseudonana CCMP1335] | 1041 | 0 | 892,493 | 53,37132004 |
| EP00533 | Sar;Stramenopiles;Gyrista;Ochrophyta;"BD clade";[ | EP00533 Pseudo-nitzschia delicatissima 303 | 1,30E-19 | 1037 | XP 002290923.1 | predicted protein [Thalassiosira pseudonana CCMP1335] | 1041 | 0 | 930,243 | 52,64650284 |
| EP00089 | Obazoa;Opisthokonta;Holozoa;Choanozoa;Metazoa | EP00089 Calanus finmarchicus 93275 | 1,40E-19 | 287 | XP 023346731.1 | protein transport protein Sec16-like isoform X2 [Eurytemora affinis] | 1941 | 2,95E-31 | 133,65 | 59,22330097 |
| EP00269 | Archaeplastida;Chloroplastida;Streptophyta;"core s | EP00269 Selaginella moellendorffii 35316 | 1,50E-19 | 205 | XP 024535962.1 | protein transport protein SEC16A homolog isoform X4 [Selaginella moellendorffii] >gi\|137 | 205 | 4,62E-145 | 410,994 | 100 |
| EP00269 | Archaeplastida;Chloroplastida;Streptophyta;"core s | EP00269 Selaginella moellendorffii 35317 | 1,50E-19 | 205 | XP 024535962.1 | protein transport protein SEC16A homolog isoform X4 [Selaginella moellendorffii] >gi\|137 | 205 | 4,62E-145 | 410,994 | 100 |
| EP00637 | Sar;Stramenopiles;Gyrista;Ochrophyta;Pinguiophyc | EP00637 Pinguiococcus pyrenoidosus 9326 | 1,80E-19 | 1070 | XP 009038384.1 | hypothetical protein AURANDRAFT 28308, partial [Aureococcus anophagefferens] | 728 | 2,96E-133 | 430,254 | 47,66214178 |
| EP00603 | Sar;Stramenopiles;Gyrista;Ochrophyta;"CS clade";C | EP00603 Paraphysomonas imperforata 15328 | 1,90E-19 | 1092 | XP 009515715.1 | hypothetical protein PHYSODRAFT 322106 [Phytophthora sojae] | 1262 | 0 | 628,246 | 51,59386068 |
| EP00089 | Obazoa;Opisthokonta;Holozoa;Choanozoa;Metazoa | EP00089 Calanus finmarchicus 93273 | 1,90E-19 | 552 | XP 023346731.1 | protein transport protein Sec16-like isoform X2 [Eurytemora affinis] | 1941 | 2,65E-32 | 143,665 | 51,64835165 |
| EP00027 | Amoebozoa;Evosea;Variosea;Filamoeba;Filamoeba | EP00027 Filamoeba nolandi 14028 | 2,00E-19 | 1039 | XP 012754771.1 | hypothetical protein SAMD00019534 058260 [Acytostelium subglobosum LB1] | 1179 | 0 | 605,905 | 49,28057554 |
| EP00027 | Amoebozoa;Evosea;Variosea;Filamoeba;Filamoeba | EP00027 Filamoeba nolandi 17434 | 2,00E-19 | 1084 | XP 012754771.1 | hypothetical protein SAMD00019534 058260 [Acytostelium subglobosum LB1] | 1179 | 0 | 650,203 | 48,99159664 |
| EP00223 | Archaeplastida;Chloroplastida;Chlorophyta;Mamiel | EP00223 Ostreococcus mediterraneus 6592 | 2,00E-19 | 623 | XP 001418383.1 | predicted protein [Ostreococcus lucimarinus CCE9901] | 592 | 9,85E-139 | 426,402 | 58,50234009 |
| EP00530 | Sar;Stramenopiles;Gyrista;Ochrophyta;"BD clade";[ | EP00530 Phaeodactylum tricornutum 5026 | 2,60E-19 | 1133 | XP 002181151.1 | predicted protein [Phaeodactylum tricornutum CCAP 1055/1] | 1133 | 0 | 2337,76 | 100 |
| EP00428 | Sar;Alveolata;Myzozoa;"DP clade";Dinoflagellata;D | EP00428 Durinskia baltica 38493 | 2,70E-19 | 199 | XP 021977758.1 | protein transport protein SEC16B homolog isoform X1 [Helianthus annuus] | 1391 | 1,92E-06 | 58,151 | 34,68208092 |
| EP00379 | Sar;Alveolata;Myzozoa;"AC clade";Apicomplexa;"cc | EP00379 Toxoplasma gondii 3361 | 2,70E-19 | 2142 | XP 002370242.1 | hypothetical protein TGME49 294730 [Toxoplasma gondii ME49] | 2142 | 0 | 4143,58 | 100 |
| EP00616 | Sar;Stramenopiles;Gyrista;Ochrophyta;Dictyochoph | EP00616 Rhizochromulina sp CCMP1243 17886 | 2,80E-19 | 1091 | XP 009038384.1 | hypothetical protein AURANDRAFT 28308, partial [Aureococcus anophagefferens] | 728 | 0 | 670,233 | 60 |
| EP00027 | Amoebozoa;Evosea;Variosea;Filamoeba;Filamoeba | EP00027 Filamoeba nolandi 1027 | 2,80E-19 | 1120 | XP 012754771.1 | hypothetical protein SAMD00019534 058260 [Acytostelium subglobosum LB1] | 1179 | 0 | 699,123 | 49,14004914 |
| EP00029 | Amoebozoa;Tubulinea;Echinamoebidia;Vermamoet | EP00029 Vermamoeba vermiformis 14674 | 2,80E-19 | 1164 | XP 012754771.1 | hypothetical protein SAMD00019534 058260 [Acytostelium subglobosum LB1] | 1179 | 0 | 673,315 | 46,73650282 |
| EP00163 | Ancyromonadida;Fabomonas;Fabomonas tropica;st | EP00163 Fabomonas tropica 31182 | 3,80E-19 | 277 | XP 020866873.1 | protein transport protein SEC16B homolog isoform X2 [Arabidopsis lyrata subsp. lyrata] | 1357 | 3,14E-10 | 71,633 | 55,74324324 |
| EP00160 | Obazoa;Opisthokonta;Nucletmycea;Rotosphaerida; | EP00160 Parvularia atlantis 21825 | 4,40E-19 | 112 | XP 011429675.1 | PREDICTED: uncharacterized protein LOC105329889 [Crassostrea gigas] | 1436 | 1,74E-08 | 61,2326 | 34,7826087 |
| EP00593 | Sar;Stramenopiles;Gyrista;Ochrophyta;"BD clade";[ | EP00593 Proboscia inermis 5656 | 4,40E-19 | 699 | XP 002183614.1 | predicted protein [Phaeodactylum tricornutum CCAP 1055/1] | 2083 | 9,33E-19 | 103,219 | 62,83783784 |
| EP00528 | Sar;Stramenopiles;Gyrista;Ochrophyta;"BD clade";[ | EP00528 Fragilariopsis cylindrus 9894 | 4,60E-19 | 1060 | XP 002181151.1 | predicted protein [Phaeodactylum tricornutum CCAP 1055/1] | 1133 | 0 | 947,962 | 56,22280244 |
| EP00520 | Sar;Stramenopiles;Gyrista;Ochrophyta;"BD clade";E | EP00520 Triparma pacifica 6064 | 4,60E-19 | 889 | XP 001209741.1 | conserved hypothetical protein [Aspergillus terreus NIH2624] | 1806 | 5,41E-10 | 75,485 | 48,89867841 |
| EP00614 | Sar;Stramenopiles;Gyrista;Ochrophyta;Dictyochoph | EP00614 Pseudopedinella elastica 13777 | 5,60E-19 | 1144 | XP 009038384.1 | hypothetical protein AURANDRAFT 28308, partial [Aureococcus anophagefferens] | 728 | 0 | 614,379 | 58,86990802 |
| EP00164 | Ancoracysta;Ancoracysta twista;strain TD-1 | EP00164 Ancoracysta twista 37197 | 7,20E-19 | 133 | XP 004344939.1 | hypothetical protein ACA1 285550 [Acanthamoeba castellanii str. Neff] | 1515 | 6,49E-10 | 65,855 | 41,12149533 |
| EP00740 | Archaeplastida;Chloroplastida;Chlorophyta;Mamiel | EP00740 Mantoniella antarctica 9500 | 7,40E-19 | 207 | XP 002500425.1 | predicted protein [Micromonas commoda] | 1395 | 5,68E-19 | 95,1301 | 61,88340807 |
| EP00023 | Amoebozoa;Evosea;Eumycetozoa;Dictyostelia;Dicty | EP00023 Dictyostelium discoideum 3868 | 7,70E-19 | 4246 | XP 637397.1 | vacuolar protein sorting-associated protein 13 family protein [Dictyostelium discoideum A | 4246 | 0 | 8599,57 | 100 |
| EP00258 | Archaeplastida;Chloroplastida;Streptophyta;"core s | EP00258 Populus trichocarpa 20684 | 9,40E-19 | 1138 | XP 024436703.1 | protein transport protein SEC31 homolog B [Populus trichocarpa] | 1138 | 0 | 2353,17 | 100 |
| EP00580 | Sar;Stramenopiles;Gyrista;Ochrophyta;"BD clade";[ | EP00580 Thalassiosira gravida 5176 | 1,00E-18 | 1130 | XP 002290923.1 | predicted protein [Thalassiosira pseudonana CCMP1335] | 1041 | 0 | 1382,08 | 74,05452946 |
| EP00257 | Archaeplastida;Chloroplastida;Streptophyta;"core s | EP00257 Ricinus communis 20787 | 1,00E-18 | 894 | XP 015580116.2 | protein transport protein SEC31 homolog B [Ricinus communis] | 894 | 0 | 1850,48 | 100 |
| EP00584 | Sar;Stramenopiles;Gyrista;Ochrophyta;"BD clade";[ | EP00584 Thalassiosira punctigera 2653 | 1,20E-18 | 1276 | XP 002290923.1 | predicted protein [Thalassiosira pseudonana CCMP1335] | 1041 | 0 | 1074,31 | 73,50746269 |
| EP00195 | Archaeplastida;Chloroplastida;Chlorophyta;"core ch | EP00195 Chlamydomonas chlamydogama 12889 | 1,30E-18 | 1245 | XP 001697039.1 | COP-II coat subunit [Chlamydomonas reinhardtii] | 1313 | 0 | 1023,46 | 64,79028698 |
| EP00203 | Archaeplastida;Chloroplastida;Chlorophyta;"core ch | EP00203 Chlorella variabilis 9656 | 1,50E-18 | 3297 | XP 005852202.1 | hypothetical protein CHLNCDRAFT 133426 [Chlorella variabilis] | 3297 | 0 | 6368,49 | 100 |
| EP00751 | Sar;Stramenopiles;Gyrista;Ochrophyta;"BD clade";[ | EP00751 Fragilariopsis kerguelensis 1392 | 1,70E-18 | 1069 | XP 002181151.1 | predicted protein [Phaeodactylum tricornutum CCAP 1055/1] | 1133 | 0 | 898,271 | 56,59246575 |
| EP00256 | Archaeplastida;Chloroplastida;Streptophyta;"core s | EP00256 Glycine max 17560 | 1,80E-18 | 1118 | XP 003534381.1 | protein transport protein SEC31 homolog B [Glycine max] >gi\|1590556237\|ref\|XP 02824 | 1118 | 0 | 2311,18 | 100 |
| EP00376 | Sar;Alveolata;Myzozoa;"AC clade";Apicomplexa;"cc | EP00376 Eimeria tenella 479 | 2,60E-18 | 1356 | XP 013228205.1 | hypothetical protein, conserved [Eimeria tenella] | 1356 | 0 | 2629,74 | 100 |
| EP00620 | Sar;Stramenopiles;Gyrista;Ochrophyta;Dictyochoph | EP00620 Florenciella sp RCC1587 6688 | 2,70E-18 | 350 | ***no hit found*** | | | | | |
| EP00264 | Archaeplastida;Chloroplastida;Streptophyta;"core s | EP00264 Hordeum vulgare 57071 | 2,80E-18 | 615 | XP 020187483.1 | protein transport protein SEC31 homolog B isoform X1 [Aegilops tauschii subsp. tauschii] | 1135 | 0 | 1214,91 | 97,15719064 |
| EP00299 | Haptista;Centroplasthelida;Pterocystida;Pterista;"P | EP00299 Pterocystis canadensis 17689 | 3,00E-18 | 286 | XP 018914533.1 | PREDICTED: sporozoite surface protein 2-like isoform X2 [Bemisia tabaci] | 349 | 2,06E-06 | 59,3066 | 51,5625 |
| EP00467 | Sar;Rhizaria;Cercozoa;Chlorarachnea;"core chlorara | EP00467 Chlorarachnion reptans 201 | 3,00E-18 | 617 | ***no hit found*** | | | | | |
| EP00603 | Sar;Stramenopiles;Gyrista;Ochrophyta;"CS clade";C | EP00603 Paraphysomonas imperforata 8215 | 3,10E-18 | 1109 | XP 009515715.1 | hypothetical protein PHYSODRAFT 322106 [Phytophthora sojae] | 1262 | 0 | 622,083 | 53,12855518 |
| EP00293 | Cryptista;Cryptophyceae;Cryptomonadales;Proteon | EP00293 Proteomonas sulcata 11960 | 3,50E-18 | 181 | ***no hit found*** | | | | | |
| EP00297 | Cryptista;Palpitomonas;Palpitomonas bilix;strain N | EP00297 Palpitomonas bilix 12676 | 3,50E-18 | 301 | XP 019254119.1 | PREDICTED: protein transport protein SEC16A homolog [Nicotiana attenuata] | 1521 | 2,25E-06 | 60,077 | 38,96713615 |
| EP00376 | Sar;Alveolata;Myzozoa;"AC clade";Apicomplexa;"cc | EP00376 Eimeria tenella 617 | 3,70E-18 | 1271 | XP 013228343.1 | beta-tubulin cofactor d, putative [Eimeria tenella] | 1271 | 0 | 2412,49 | 100 |
| EP00376 | Sar;Alveolata;Myzozoa;"AC clade";Apicomplexa;"cc | EP00376 Eimeria tenella 8216 | 4,10E-18 | 2453 | XP 013235960.1 | hypothetical protein, conserved [Eimeria tenella] | 2453 | 0 | 4832,7 | 100 |
| EP00044 | Obazoa;Opisthokonta;Holozoa;Choanozoa;Choanof | EP00044 Salpingoeca rosetta 1367 | 4,20E-18 | 1985 | XP 004988800.1 | hypothetical protein PTSG 10421 [Salpingoeca rosetta] | 1985 | 0 | 3898,21 | 100 |
| EP00162 | Ancyromonadida;Ancyromonadidae;Nutomonas;Nu | EP00162 Nutomonas longa 4907 | 4,70E-18 | 228 | XP 010494986.1 | PREDICTED: protein transport protein SEC16A homolog isoform X2 [Camelina sativa] | 1382 | 1,80E-10 | 70,8626 | 49,08256881 |
| EP00249 | Archaeplastida;Chloroplastida;Streptophyta;"core s | EP00249 Psilotum nudum 22019 | 4,70E-18 | 805 | XP 006482944.1 | protein transport protein SEC31 homolog B isoform X1 [Citrus sinensis] >gi\|1350271342\|r | 1120 | 0 | 652,899 | 57,10814095 |
| EP00445 | Sar;Alveolata;Myzozoa;"DP clade";Dinoflagellata;D | EP00445 Apocalathium hangoei 63884 | 4,90E-18 | 1167 | XP 013251586.1 | hypothetical protein, conserved [Eimeria acervulina] | 851 | 2,15E-19 | 105,916 | 40,60150376 |
| EP00455 | Sar;Rhizaria;"eorhiza";Aquavolonida;"NC10 C lineag | EP00455 Lapot gusevi 53634 | 5,40E-18 | 266 | XP 029230831.1 | uncharacterized protein Tco025E 02121 [Trypanosoma conorhini] | 1464 | 2,00E-16 | 89,3521 | 66,66666667 |
| EP00650 | Sar;Stramenopiles;Gyrista;Peronosporomycetes;"cc | EP00650 Pythium arrhenomanes 39 | 5,70E-18 | 1149 | XP 009515715.1 | hypothetical protein PHYSODRAFT 322106 [Phytophthora sojae] | 1262 | 0 | 1523,84 | 74,98051442 |
| EP00222 | Archaeplastida;Chloroplastida;Chlorophyta;Mamiel | EP00222 Ostreococcus lucimarinus 5240 | 5,80E-18 | 1077 | XP 001420597.1 | predicted protein, partial [Ostreococcus lucimarinus CCE9901] | 1077 | 0 | 2195,62 | 100 |
| EP00643 | Sar;Stramenopiles;Gyrista;Peronosporomycetes;"cc | EP00643 Hyaloperonospora arabidopsidis 7137 | 6,10E-18 | 797 | XP 009515715.1 | hypothetical protein PHYSODRAFT 322106 [Phytophthora sojae] | 1262 | 0 | 1408,28 | 87,38853503 |
| EP00020 | Amoebozoa;Evosea;Cutosea;Sapocribrum;Sapocribr | EP00020 Sapocribrum chincoteaguense 4061 | 6,20E-18 | 444 | ***no hit found*** | | | | | |
| EP00192 | Archaeplastida;Chloroplastida;Chlorophyta;"core ch | EP00192 Tetraselmis astigmatica 495 | 6,20E-18 | 1222 | XP 022152713.1 | protein transport protein SEC31 homolog B [Momordica charantia] | 1114 | 0 | 820,461 | 55,71895425 |
| EP00192 | Archaeplastida;Chloroplastida;Chlorophyta;"core ch | EP00192 Tetraselmis astigmatica 724 | 6,30E-18 | 1221 | XP 022152713.1 | protein transport protein SEC31 homolog B [Momordica charantia] | 1114 | 0 | 778,859 | 54,53802126 |
| EP00532 | Sar;Stramenopiles;Gyrista;Ochrophyta;"BD clade";[ | EP00532 Pseudo-nitzschia australis 180 | 6,30E-18 | 1048 | XP 002290923.1 | predicted protein [Thalassiosira pseudonana CCMP1335] | 1041 | 0 | 927,932 | 50,66666667 |
| EP00249 | Archaeplastida;Chloroplastida;Streptophyta;"core s | EP00249 Psilotum nudum 22020 | 7,30E-18 | 1167 | XP 015881568.1 | protein transport protein SEC31 homolog B isoform X1 [Ziziphus jujuba] | 1154 | 0 | 1118,61 | 62,54098361 |
| EP00540 | Sar;Stramenopiles;Gyrista;Ochrophyta;"BD clade";[ | EP00540 Astrosyne radiata 9643 | 7,40E-18 | 727 | XP 002181151.1 | predicted protein [Phaeodactylum tricornutum CCAP 1055/1] | 1133 | 0 | 566,229 | 55,79896907 |
| EP00513 | Sar;Stramenopiles;Bigyra;Sagenista;Labyrinthulomy | EP00513 Hondaea fermentalgiana 1036 | 8,70E-18 | 1322 | XP 009515715.1 | hypothetical protein PHYSODRAFT 322106 [Phytophthora sojae] | 1262 | 6,82E-143 | 477,633 | 54,30944964 |
| EP00134 | Obazoa;Opisthokonta;Nucletmycea;Fungi;"core Fur | EP00134 Neurospora crassa 8942 | 1,00E-17 | 1256 | XP 963510.1 | protein transporter sec-31 [Neurospora crassa OR74A] | 1256 | 0 | 2548,08 | 100 |
| EP00083 | Obazoa;Opisthokonta;Holozoa;Choanozoa;Metazoa | EP00083 Stegodyphus mimosarum 16877 | 1,00E-17 | 1223 | XP 021001566.1 | protein transport protein Sec31A-like [Parasteatoda tepidariorum] | 1219 | 0 | 1770,75 | 76,98476343 |
| EP00646 | Sar;Stramenopiles;Gyrista;Peronosporomycetes;"cc | EP00646 Phytophthora ramorum 12371 | 1,10E-17 | 1328 | XP 009515715.1 | hypothetical protein PHYSODRAFT 322106 [Phytophthora sojae] | 1262 | 0 | 2228,75 | 92,49011858 |
| EP00554 | Sar;Stramenopiles;Gyrista;Ochrophyta;"BD clade";[ | EP00554 Chaetoceros debilis 15116 | 1,10E-17 | 1087 | XP 002290923.1 | predicted protein [Thalassiosira pseudonana CCMP1335] | 1041 | 0 | 829,706 | 59,14948454 |
| EP00269 | Archaeplastida;Chloroplastida;Streptophyta;"core s | EP00269 Selaginella moellendorffii 31691 | 1,10E-17 | 284 | XP 024532337.1 | protein transport protein SEC16A homolog [Selaginella moellendorffii] | 284 | 0 | 561,607 | 100 |
| EP00648 | Sar;Stramenopiles;Gyrista;Peronosporomycetes;"cc | EP00648 Phytopythium vexans 1193 | 1,20E-17 | 1259 | XP 009515715.1 | hypothetical protein PHYSODRAFT 322106 [Phytophthora sojae] | 1262 | 0 | 1755,73 | 77,80355762 |
| EP00240 | Archaeplastida;Chloroplastida;Chlorophyta;Pyramir | EP00240 Pyramimonas obovata 6635 | 1,20E-17 | 162 | XP 007155661.1 | hypothetical protein PHAVU 003G220900g [Phaseolus vulgaris] | 1379 | 3,52E-08 | 62,003 | 66,42857143 |
| EP00428 | Sar;Alveolata;Myzozoa;"DP clade";Dinoflagellata;D | EP00428 Durinskia baltica 79388 | 1,20E-17 | 158 | XP 002183614.1 | predicted protein [Phaeodactylum tricornutum CCAP 1055/1] | 2083 | 1,47E-07 | 60,077 | 43,68932039 |
| EP00613 | Sar;Stramenopiles;Gyrista;Ochrophyta;Dictyochoph | EP00613 Pedinella sp CCMP2098 56696 | 1,30E-17 | 487 | ***no hit found*** | | | | | |
| EP00259 | Archaeplastida;Chloroplastida;Streptophyta;"core s | EP00259 Arabidopsis lyrata 29045 | 1,50E-17 | 1102 | XP 020881761.1 | protein transport protein SEC31 homolog B isoform X2 [Arabidopsis lyrata subsp. lyrata] | 1102 | 0 | 2286,15 | 100 |
| EP00739 | Archaeplastida;Chloroplastida;Chlorophyta;Mamiel | EP00739 Bathycoccus prasinos 3475 | 1,70E-17 | 1386 | XP 007512259.1 | predicted protein [Bathycoccus prasinos] | 1096 | 0 | 2241,08 | 99,8173516 |
| EP00259 | Archaeplastida;Chloroplastida;Streptophyta;"core s | EP00259 Arabidopsis lyrata 6541 | 1,90E-17 | 1104 | XP 002878518.1 | protein transport protein SEC31 homolog B isoform X1 [Arabidopsis lyrata subsp. lyrata] | 1104 | 0 | 2291,92 | 100 |
| EP00647 | Sar;Stramenopiles;Gyrista;Peronosporomycetes;"cc | EP00647 Phytophthora sojae 1951 | 2,00E-17 | 1262 | XP 009515715.1 | hypothetical protein PHYSODRAFT 322106 [Phytophthora sojae] | 1262 | 0 | 2581,59 | 100 |
| EP00228 | Archaeplastida;Chloroplastida;Chlorophyta;Mamiel | EP00228 Micromonas bravo 5246 | 2,40E-17 | 400 | XP 002500425.1 | predicted protein [Micromonas commoda] | 1395 | 1,27E-26 | 123,25 | 61,78861789 |
| EP00186 | Archaeplastida;"RR clade";Rhodophyta;Proteorhod | EP00186 Timspurckia oligopyrenoides 231 | 2,40E-17 | 306 | XP 030500564.1 | protein transport protein SEC16B homolog [Cannabis sativa] >gi\|1731704580\|ref\|XP 03( | 1470 | 4,67E-07 | 62,3882 | 55,17241379 |
| EP00251 | Archaeplastida;Chloroplastida;Streptophyta;"core s | EP00251 Ginkgo biloba 34254 | 2,40E-17 | 1160 | XP 006837403.2 | protein transport protein SEC31 homolog B isoform X1 [Amborella trichopoda] | 1155 | 0 | 1486,09 | 68,51535836 |
| EP00264 | Archaeplastida;Chloroplastida;Streptophyta;"core s | EP00264 Hordeum vulgare 57098 | 2,50E-17 | 545 | XP 020187483.1 | protein transport protein SEC31 homolog B isoform X1 [Aegilops tauschii subsp. tauschii] | 1135 | 0 | 1084,32 | 96,8164794 |
| EP00252 | Archaeplastida;Chloroplastida;Streptophyta;"core s | EP00252 Welwitschia mirabilis 10265 | 2,70E-17 | 1077 | XP 015881568.1 | protein transport protein SEC31 homolog B isoform X1 [Ziziphus jujuba] | 1154 | 0 | 1140,18 | 61,2244898 |
| EP00260 | Archaeplastida;Chloroplastida;Streptophyta;"core s | EP00260 Arabidopsis thaliana 31898 | 3,50E-17 | 1102 | NP 191905.3 | transducin family protein / WD-40 repeat family protein [Arabidopsis thaliana] | 1102 | 0 | 2285,76 | 100 |
| EP00086 | Obazoa;Opisthokonta;Holozoa;Choanozoa;Metazoa | EP00086 Labidocera madurae 25886 | 3,60E-17 | 1555 | XP 023341345.1 | AT-rich interactive domain-containing protein 2-like isoform X3 [Eurytemora affinis] | 1573 | 0 | 1161,36 | 82,68456376 |
| EP00182 | Archaeplastida;"RR clade";Rhodophyta;Proteorhod | EP00182 Erythrolobus australicus 2245 | 3,60E-17 | 392 | XP 016436112.1 | PREDICTED: protein transport protein SEC16B homolog [Nicotiana tabacum] | 289 | 8,08E-13 | 78,9518 | 59,3220339 |
| EP00260 | Archaeplastida;Chloroplastida;Streptophyta;"core s | EP00260 Arabidopsis thaliana 46528 | 3,70E-17 | 1104 | NP 851024.1 | transducin family protein / WD-40 repeat family protein [Arabidopsis thaliana] | 1104 | 0 | 2289,61 | 100 |
| EP00044 | Obazoa;Opisthokonta;Holozoa;Choanozoa;Choanof | EP00044 Salpingoeca rosetta 2726 | 5,20E-17 | 4709 | XP 004990159.1 | beige/BEACH domain-containing protein [Salpingoeca rosetta] | 4709 | 0 | 9599,93 | 100 |
| EP00525 | Sar;Stramenopiles;Gyrista;Ochrophyta;"BD clade";[ | EP00525 Craspedostauros australis 2246 | 5,20E-17 | 88 | XP 002183614.1 | predicted protein [Phaeodactylum tricornutum CCAP 1055/1] | 2083 | 1,70E-14 | 77,0258 | 46,51162791 |
| EP00232 | Archaeplastida;Chloroplastida;Chlorophyta;Nephro | EP00232 Nephroselmis pyriformis 24447 | 5,30E-17 | 981 | XP 001420597.1 | predicted protein, partial [Ostreococcus lucimarinus CCE9901] | 1077 | 0 | 798,119 | 58,67768595 |
| EP00644 | Sar;Stramenopiles;Gyrista;Peronosporomycetes;"cc | EP00644 Phytophthora capsici 209 | 5,90E-17 | 1252 | XP 009515715.1 | hypothetical protein PHYSODRAFT 322106 [Phytophthora sojae] | 1262 | 0 | 2175,59 | 90,19762846 |
| EP00044 | Obazoa;Opisthokonta;Holozoa;Choanozoa;Choanof | EP00044 Salpingoeca rosetta 6989 | 5,90E-17 | 1335 | XP 004994422.1 | hypothetical protein PTSG 04335 [Salpingoeca rosetta] | 1335 | 0 | 2705,63 | 100 |
| EP00269 | Archaeplastida;Chloroplastida;Streptophyta;"core s | EP00269 Selaginella moellendorffii 35434 | 6,40E-17 | 149 | XP 024536080.1 | protein transport protein SEC16A homolog [Selaginella moellendorffii] | 149 | 5,94E-102 | 297,745 | 100 |
| EP00761 | Discoba;Heterolobosea;Pharyngomonas;Pharyngon | EP00761 Pharyngomonas kirbyi 14333 | 6,80E-17 | 756 | XP 005831033.1 | secretory protein Sec31 [Guillardia theta CCMP2712] | 1081 | 4,28E-131 | 425,631 | 46,59498208 |
| EP00376 | Sar;Alveolata;Myzozoa;"AC clade";Apicomplexa;"cc | EP00376 Eimeria tenella 3756 | 8,60E-17 | 1844 | XP 013231487.1 | hypothetical protein, conserved [Eimeria tenella] | 1844 | 0 | 3588,51 | 100 |
| EP00200 | Archaeplastida;Chloroplastida;Chlorophyta;"core ch | EP00200 Dunaliella tertiolecta 13460 | 9,10E-17 | 1890 | ***no hit found*** | | | | | |
| EP00440 | Sar;Alveolata;Myzozoa;"DP clade";Dinoflagellata;D | EP00440 Ansanella granifera 48298 | 1,00E-16 | 745 | ***no hit found*** | | | | | |
| EP00559 | Sar;Stramenopiles;Gyrista;Ochrophyta;"BD clade";[ | EP00559 Dactyliosolen fragilissimus 6456 | 1,10E-16 | 1106 | XP 002290923.1 | predicted protein [Thalassiosira pseudonana CCMP1335] | 1041 | 0 | 1056,2 | 58,5125448 |
| EP00248 | Archaeplastida;Chloroplastida;Streptophyta;"core s | EP00248 Marchantia polymorpha 2687 | 1,20E-16 | 1165 | XP 024379159.1 | protein transport protein SEC31 homolog B-like isoform X2 [Physcomitrella patens] | 1166 | 0 | 1108,98 | 59,21601334 |
| EP00639 | Sar;Stramenopiles;Gyrista;Ochrophyta;Raphidophy | EP00639 Heterosigma akashiwo 9070 | 1,20E-16 | 725 | XP 009515715.1 | hypothetical protein PHYSODRAFT 322106 [Phytophthora sojae] | 1262 | 0 | 623,239 | 54,06091371 |
| EP00736 | Archaeplastida;"RR clade";Rhodelphis;Rhodelphis n | EP00736 Rhodelphis marinus 9603 | 1,20E-16 | 1097 | XP 010552745.1 | PREDICTED: protein transport protein SEC31 homolog B-like isoform X1 [Tarenaya hassleri | 1114 | 5,63E-160 | 513,072 | 43,59415305 |
| EP00538 | Sar;Stramenopiles;Gyrista;Ochrophyta;"BD clade";[ | EP00538 Stauroneis constricta 306 | 1,40E-16 | 1053 | XP 002290923.1 | predicted protein [Thalassiosira pseudonana CCMP1335] | 1041 | 0 | 1021,54 | 54,62184874 |
| EP00649 | Sar;Stramenopiles;Gyrista;Peronosporomycetes;"cc | EP00649 Plasmopara halstedii 181 | 1,50E-16 | 1104 | XP 024571614.1 | Vesicle coat complex COPII, subunit SEC31 [Plasmopara halstedii] | 1104 | 0 | 2274,59 | 100 |
| EP00376 | Sar;Alveolata;Myzozoa;"AC clade";Apicomplexa;"cc | EP00376 Eimeria tenella 2916 | 1,50E-16 | 2133 | XP 013230646.1 | hypothetical protein, conserved [Eimeria tenella] | 2133 | 0 | 4331,94 | 100 |
| EP00645 | Sar;Stramenopiles;Gyrista;Peronosporomycetes;"cc | EP00645 Phytophthora infestans 11954 | 1,60E-16 | 1264 | XP 002906652.1 | protein transporter Sec31A, putative [Phytophthora infestans T30-4] | 1264 | 0 | 2591,99 | 100 |
| EP00023 | Amoebozoa;Evosea;Eumycetozoa;Dictyostelia;Dicty | EP00023 Dictyostelium discoideum 11933 | 1,70E-16 | 1355 | XP 646396.1 | WD40 repeat-containing protein [Dictyostelium discoideum AX4] | 1355 | 0 | 2789,99 | 100 |
| EP00512 | Sar;Stramenopiles;Bigyra;Sagenista;Labyrinthulomy | EP00512 Aurantiochytrium limacinum 6408 | 1,80E-16 | 497 | ***no hit found*** | | | | | |
| EP00376 | Sar;Alveolata;Myzozoa;"AC clade";Apicomplexa;"cc | EP00376 Eimeria tenella 2518 | 1,80E-16 | 5838 | XP 013230245.1 | UBA/TS-N domain-containing protein, putative [Eimeria tenella] | 5838 | 0 | 11712 | 100 |
| EP00339 | Sar;Alveolata;Ciliophora;Intramacronucleata;"CON: | EP00339 Tiarina fusus 498 | 1,80E-16 | 1049 | XP 002290923.1 | predicted protein [Thalassiosira pseudonana CCMP1335] | 1041 | 0 | 1000,35 | 54,13533835 |
| EP00523 | Sar;Stramenopiles;Gyrista;Ochrophyta;"BD clade";[ | EP00523 Amphiprora sp CCMP467 2780 | 1,90E-16 | 1162 | XP 002181151.1 | predicted protein [Phaeodactylum tricornutum CCAP 1055/1] | 1133 | 0 | 914,45 | 55,2852349 |
| EP00602 | Sar;Stramenopiles;Gyrista;Ochrophyta;"CS clade";C | EP00602 Paraphysomonas bandaiensis 4721 | 1,90E-16 | 1052 | XP 002290923.1 | predicted protein [Thalassiosira pseudonana CCMP1335] | 1041 | 0 | 694,501 | 45,72490706 |
| EP00090 | Obazoa;Opisthokonta;Holozoa;Choanozoa;Metazoa | EP00090 Calanus glacialis 45931 | 2,00E-16 | 406 | XP 023346731.1 | protein transport protein Sec16-like isoform X2 [Eurytemora affinis] | 1941 | 1,60E-31 | 137,887 | 51,32978723 |

| | | | | | | | | | | |
|---|---|---|---|---|---|---|---|---|---|---|
| EP00478 | Sar;Rhizaria;Endomyxa;"zoorhiza";Filoreta;"Ftenera | EP00478 Filoreta tenera 2146 | 2,10E-16 | 241 | ***no hit found*** | | | | | |
| EP00074 | Obazoa;Opisthokonta;Holozoa;Choanozoa;Metazoa | EP00074 Homo sapiens 93406 | 2,40E-16 | 1546 | XP 016870853.1 | protein transport protein Sec16A isoform X18 [Homo sapiens] | 1546 | 0 | 3093,91 | 100 |
| EP00237 | Archaeplastida;Chloroplastida;Chlorophyta;Pycnocc | EP00237 Pycnococcus provasolii 8538 | 2,50E-16 | 154 | XP 010530052.1 | PREDICTED: protein transport protein SEC16B homolog [Tarenaya hassleriana] | 1377 | 3,05E-08 | 62,003 | 47,32142857 |
| EP00376 | Sar;Alveolata;Myzozoa;"AC clade";Apicomplexa;"cc | EP00376 Eimeria tenella 1246 | 2,80E-16 | 3206 | XP 013228972.1 | hypothetical protein, conserved [Eimeria tenella] | 3206 | 0 | 6345,38 | 100 |
| EP00676 | Discoba;Euglenozoa;Kinetoplastea;Metakinetoplast | EP00676 Leptomonas seymouri 6387 | 2,80E-16 | 1285 | XP 015653768.1 | hypothetical protein ABB37 08623 [Leptomonas pyrrhocoris] | 1298 | 0 | 1245,34 | 69,77443609 |
| EP00255 | Archaeplastida;Chloroplastida;Streptophyta;"core s | EP00255 Erythranthe guttata 22432 | 2,80E-16 | 1129 | XP 012849694.1 | PREDICTED: protein transport protein SEC31 homolog B [Erythranthe guttata] | 1129 | 0 | 2330,83 | 100 |
| EP00556 | Sar;Stramenopiles;Gyrista;Ochrophyta;"BD clade";[ | EP00556 Chaetoceros neogracilis 17689 | 3,00E-16 | 1151 | XP 002290923.1 | predicted protein [Thalassiosira pseudonana CCMP1335] | 1041 | 0 | 796,579 | 59,57446809 |
| EP00245 | Archaeplastida;Chloroplastida;Streptophyta;"core s | EP00245 Coleochaete scutata 12723 | 3,10E-16 | 141 | XP 002987428.1 | protein transport protein SEC16B homolog [Selaginella moellendorffii] | 1190 | 5,45E-31 | 126,716 | 54,03225806 |
| EP00687 | Discoba;Heterolobosea;Tetramitia;Eutetramitia;"PS | EP00687 Percolomonas cosmopolitus 13695 | 3,60E-16 | 1108 | XP 012754771.1 | hypothetical protein SAMD00019534 058260 [Acytostelium subglobosum LB1] | 1179 | 2,75E-113 | 389,808 | 47,27646454 |
| EP00627 | Sar;Stramenopiles;Gyrista;Ochrophyta;Pelagophyce | EP00627 Pelagomonas calceolata 12009 | 3,70E-16 | 969 | XP 009038384.1 | hypothetical protein AURANDRAFT 28308, partial [Aureococcus anophagefferens] | 728 | 0 | 876,315 | 72,27586207 |
| EP00027 | Amoebozoa;Evosea;Variosea;Filamoeba;Filamoeba | EP00027 Filamoeba nolandi 6312 | 3,70E-16 | 213 | ***no hit found*** | | | | | |
| EP00157 | Obazoa;Opisthokonta;Nucletmycea;Fungi;"cryptom | EP00157 Rozella allomycis 3290 | 3,90E-16 | 361 | XP 010542224.1 | PREDICTED: protein transport protein SEC16B homolog [Tarenaya hassleriana] | 1321 | 1,31E-07 | 64,6994 | 43,93939394 |
| EP00627 | Sar;Stramenopiles;Gyrista;Ochrophyta;Pelagophyce | EP00627 Pelagomonas calceolata 26403 | 4,00E-16 | 980 | XP 009038384.1 | hypothetical protein AURANDRAFT 28308, partial [Aureococcus anophagefferens] | 728 | 0 | 874,004 | 72,27586207 |
| EP00241 | Archaeplastida;Chloroplastida;Chlorophyta;Pyramir | EP00241 Pyramimonas parkeae 4035 | 4,10E-16 | 662 | XP 009348578.1 | PREDICTED: protein transport protein SEC16B homolog [Pyrus x bretschneideri] >gi\|69444 | 1417 | 1,29E-10 | 76,6406 | 60,64516129 |
| EP00455 | Sar;Rhizaria;"eorhiza";Aquavolonida;"NC10 C lineag | EP00455 Lapot gusevi 53635 | 4,90E-16 | 197 | XP 029230831.1 | uncharacterized protein Tco025E 02121 [Trypanosoma conorhini] | 1464 | 1,04E-16 | 88,1965 | 66,66666667 |
| EP00200 | Archaeplastida;Chloroplastida;Chlorophyta;"core ch | EP00200 Dunaliella tertiolecta 17338 | 5,00E-16 | 1095 | XP 001689665.1 | hypothetical protein CHLREDRAFT 146684 [Chlamydomonas reinhardtii] | 1145 | 2,61E-21 | 112,464 | 49,20634921 |
| EP00247 | Archaeplastida;Chloroplastida;Streptophyta;"core s | EP00247 Physcomitrella patens 22908 | 5,00E-16 | 1166 | XP 024379159.1 | protein transport protein SEC31 homolog B-like isoform X2 [Physcomitrella patens] | 1166 | 0 | 2388,61 | 100 |
| EP00089 | Obazoa;Opisthokonta;Holozoa;Choanozoa;Metazoa | EP00089 Calanus finmarchicus 93271 | 5,10E-16 | 458 | XP 023346731.1 | protein transport protein Sec16-like isoform X2 [Eurytemora affinis] | 1941 | 1,80E-32 | 142,124 | 51,64835165 |
| EP00089 | Obazoa;Opisthokonta;Holozoa;Choanozoa;Metazoa | EP00089 Calanus finmarchicus 93272 | 5,10E-16 | 458 | XP 023346731.1 | protein transport protein Sec16-like isoform X2 [Eurytemora affinis] | 1941 | 1,80E-32 | 142,124 | 51,64835165 |
| EP00564 | Sar;Stramenopiles;Gyrista;Ochrophyta;"BD clade";[ | EP00564 Ditylum brightwellii 7747 | 5,20E-16 | 1036 | XP 002290923.1 | predicted protein [Thalassiosira pseudonana CCMP1335] | 1041 | 0 | 1124 | 60,15037594 |
| EP00588 | Sar;Stramenopiles;Gyrista;Ochrophyta;"BD clade";[ | EP00588 Corethron pennatum 7886 | 5,80E-16 | 603 | XP 002908536.1 | COPII coat assembly protein SEC16, putative [Phytophthora infestans T30-4] | 2641 | 2,46E-10 | 75,485 | 48,54368932 |
| EP00326 | Haptista;Haptophyta;Prymnesiophyceae;Prymnesia | EP00326 Haptolina ericina 35715 | 6,10E-16 | 435 | XP 021848631.1 | protein transport protein SEC16B homolog [Spinacia oleracea] | 1425 | 3,79E-06 | 60,8474 | 40,60324826 |
| EP00566 | Sar;Stramenopiles;Gyrista;Ochrophyta;"BD clade";[ | EP00566 Odontella aurita 4216 | 6,20E-16 | 1158 | XP 002290923.1 | predicted protein [Thalassiosira pseudonana CCMP1335] | 1041 | 0 | 1123,23 | 64,82112436 |
| EP00633 | Sar;Stramenopiles;Gyrista;Ochrophyta;Pelagophyce | EP00633 Aureoumbra lagunensis 2734 | 6,80E-16 | 1155 | XP 002290923.1 | predicted protein [Thalassiosira pseudonana CCMP1335] | 1041 | 0 | 578,556 | 52,14105793 |
| EP00534 | Sar;Stramenopiles;Gyrista;Ochrophyta;"BD clade";[ | EP00534 Pseudo-nitzschia fraudulenta 863 | 7,00E-16 | 984 | XP 002290923.1 | predicted protein [Thalassiosira pseudonana CCMP1335] | 1041 | 0 | 868,611 | 53,28031809 |
| EP00376 | Sar;Alveolata;Myzozoa;"AC clade";Apicomplexa;"cc | EP00376 Eimeria tenella 4081 | 8,10E-16 | 1570 | XP 013231814.1 | hypothetical protein, conserved [Eimeria tenella] | 1570 | 0 | 3034,97 | 100 |
| EP00376 | Sar;Alveolata;Myzozoa;"AC clade";Apicomplexa;"cc | EP00376 Eimeria tenella 169 | 8,70E-16 | 1649 | XP 013227895.1 | hypothetical protein, conserved [Eimeria tenella] | 1649 | 0 | 3172,1 | 100 |
| EP00633 | Sar;Stramenopiles;Gyrista;Ochrophyta;Pelagophyce | EP00633 Aureoumbra lagunensis 11226 | 8,90E-16 | 1175 | XP 002290923.1 | predicted protein [Thalassiosira pseudonana CCMP1335] | 1041 | 7,88E-180 | 565,459 | 52,64900662 |
| EP00289 | Cryptista;Cryptophyceae;Cryptomonadales;"lineage | EP00289 Rhodomonas salina 3346 | 9,70E-16 | 86 | XP 020523770.1 | LOW QUALITY PROTEIN: protein transport protein SEC16A homolog [Amborella trichopod | 1619 | 1,91E-08 | 59,6918 | 53,44827586 |
| EP00194 | Archaeplastida;Chloroplastida;Chlorophyta;"core ch | EP00194 Tetraselmis striata 13005 | 1,00E-15 | 1234 | XP 006482945.1 | protein transport protein SEC31 homolog B isoform X2 [Citrus sinensis] >gi\|1350271346\| | 1117 | 0 | 842,803 | 53,36099585 |
| EP00303 | Haptista;Haptophyta;Pavlovales;Exanthemachrysis; | EP00303 Exanthemachrysis gayraliae 9922 | 1,00E-15 | 61 | XP 002683116.1 | RmlC-like cupin family protein [Naegleria gruberi] | 1290 | 9,15E-06 | 50,8322 | 46,55172414 |
| EP00556 | Sar;Stramenopiles;Gyrista;Ochrophyta;"BD clade";[ | EP00556 Chaetoceros neogracilis 2277 | 1,00E-15 | 714 | XP 002181151.1 | predicted protein [Phaeodactylum tricornutum CCAP 1055/1] | 1133 | 2,76E-145 | 462,611 | 52,59165613 |
| EP00194 | Archaeplastida;Chloroplastida;Chlorophyta;"core ch | EP00194 Tetraselmis striata 36357 | 1,10E-15 | 1272 | XP 028119649.1 | protein transport protein SEC31 homolog B-like [Camellia sinensis] | 1118 | 0 | 813,527 | 55,50277118 |
| EP00359 | Sar;Alveolata;Ciliophora;Postciliodesmatophora;He | EP00359 Climacostomum virens 9049 | 1,10E-15 | 504 | ***no hit found*** | | | | | |
| EP00617 | Sar;Stramenopiles;Gyrista;Ochrophyta;Dictyochoph | EP00617 Octactis speculum 27664 | 1,40E-15 | 288 | XP 008890960.1 | hypothetical protein PPTG 01118 [Phytophthora parasitica INRA-310] | 1621 | 1,47E-06 | 60,4622 | 54,7826087 |
| EP00203 | Archaeplastida;Chloroplastida;Chlorophyta;"core ch | EP00203 Chlorella variabilis 1235 | 1,40E-15 | 1805 | XP 005843781.1 | hypothetical protein CHLNCDRAFT 139911 [Chlorella variabilis] | 1805 | 0 | 3615,47 | 100 |
| EP00019 | Amoebozoa;Evosea;Cutosea;Armaparvus;Armaparv | EP00019 Armaparvus languidus 1393 | 1,40E-15 | 379 | ***no hit found*** | | | | | |
| EP00602 | Sar;Stramenopiles;Gyrista;Ochrophyta;"CS clade";C | EP00602 Paraphysomonas bandaiensis 5046 | 1,40E-15 | 1052 | XP 002290923.1 | predicted protein [Thalassiosira pseudonana CCMP1335] | 1041 | 0 | 683,715 | 48,19944598 |
| EP00641 | Sar;Stramenopiles;Gyrista;Peronosporomycetes;"cc | EP00641 Albugo candida 10096 | 1,50E-15 | 1181 | XP 009515715.1 | hypothetical protein PHYSODRAFT 322106 [Phytophthora sojae] | 1262 | 0 | 1399,03 | 65,96078431 |
| EP00550 | Sar;Stramenopiles;Gyrista;Ochrophyta;"BD clade";[ | EP00550 Attheya septentrionalis 8923 | 1,70E-15 | 1057 | XP 002290923.1 | predicted protein [Thalassiosira pseudonana CCMP1335] | 1041 | 0 | 1097,03 | 59,74264706 |
| EP00670 | Discoba;Euglenozoa;Euglenida;Spirocuta;Euglenea;I | EP00670 Eutreptiella gymnastica 7877 | 1,70E-15 | 1032 | XP 020585787.1 | protein transport protein SEC31 homolog B isoform X1 [Phalaenopsis equestris] | 1127 | 1,77E-175 | 552,362 | 45,44634806 |
| EP00376 | Sar;Alveolata;Myzozoa;"AC clade";Apicomplexa;"cc | EP00376 Eimeria tenella 101 | 2,00E-15 | 2879 | XP 013227827.1 | hypothetical protein ETH 00004910 [Eimeria tenella] | 2879 | 0 | 5583,84 | 100 |
| EP00591 | Sar;Stramenopiles;Gyrista;Ochrophyta;"BD clade";[ | EP00591 Stephanopyxis turris 9575 | 2,10E-15 | 1054 | XP 002290923.1 | predicted protein [Thalassiosira pseudonana CCMP1335] | 1041 | 0 | 1083,55 | 58,08550186 |
| EP00200 | Archaeplastida;Chloroplastida;Chlorophyta;"core ch | EP00200 Dunaliella tertiolecta 2506 | 2,10E-15 | 1277 | XP 001697039.1 | COP-II coat subunit [Chlamydomonas reinhardtii] | 1313 | 0 | 969,918 | 64,4198895 |
| EP00200 | Archaeplastida;Chloroplastida;Chlorophyta;"core ch | EP00200 Dunaliella tertiolecta 19755 | 2,20E-15 | 1304 | XP 001697039.1 | COP-II coat subunit [Chlamydomonas reinhardtii] | 1313 | 0 | 970,689 | 64,4198895 |
| EP00468 | Sar;Rhizaria;Cercozoa;Chlorarachnea;"core chlorara | EP00468 Gymnochlora sp CCMP2014 7452 | 2,20E-15 | 1441 | XP 005831033.1 | secretory protein Sec31 [Guillardia theta CCMP2712] | 1081 | 7,13E-110 | 384,415 | 44,81012658 |
| EP00639 | Sar;Stramenopiles;Gyrista;Ochrophyta;Raphidophy | EP00639 Heterosigma akashiwo 1561 | 2,30E-15 | 940 | XP 009515715.1 | hypothetical protein PHYSODRAFT 322106 [Phytophthora sojae] | 1262 | 0 | 658,292 | 52,14285714 |
| EP00511 | Sar;Stramenopiles;Bigyra;Sagenista;Labyrinthulomy | EP00511 Aplanochytrium stocchinoi 486 | 2,40E-15 | 380 | ***no hit found*** | | | | | |
| EP00200 | Archaeplastida;Chloroplastida;Chlorophyta;"core ch | EP00200 Dunaliella tertiolecta 8576 | 2,60E-15 | 1330 | XP 001697039.1 | COP-II coat subunit [Chlamydomonas reinhardtii] | 1313 | 0 | 971,074 | 64,4198895 |
| EP00011 | Amoebozoa;Discosea;Flabellinia;"core flabellinids"; | EP00011 Vannella sp DIVA3-517-6-12 11286 | 2,70E-15 | 366 | ***no hit found*** | | | | | |
| EP00670 | Discoba;Euglenozoa;Euglenida;Spirocuta;Euglenea;I | EP00670 Eutreptiella gymnastica 13723 | 2,80E-15 | 1056 | XP 020585787.1 | protein transport protein SEC31 homolog B isoform X1 [Phalaenopsis equestris] | 1127 | 1,56E-170 | 540,036 | 46,05146406 |
| EP00197 | Archaeplastida;Chloroplastida;Chlorophyta;"core ch | EP00197 Chlamydomonas leiostraca 9517 | 2,80E-15 | 1163 | XP 001697039.1 | COP-II coat subunit [Chlamydomonas reinhardtii] | 1313 | 0 | 1019,22 | 67,41321389 |
| EP00593 | Sar;Stramenopiles;Gyrista;Ochrophyta;"BD clade";[ | EP00593 Proboscia inermis 10168 | 2,80E-15 | 621 | XP 002290923.1 | predicted protein [Thalassiosira pseudonana CCMP1335] | 1041 | 4,68E-172 | 526,554 | 58,29383886 |
| EP00202 | Archaeplastida;Chloroplastida;Chlorophyta;"core ch | EP00202 Volvox carteri 10433 | 2,80E-15 | 1711 | XP 002956039.1 | hypothetical protein VOLCADRAFT 107038 [Volvox carteri f. nagariensis] | 1711 | 0 | 3364,32 | 100 |
| EP00218 | Archaeplastida;Chloroplastida;Chlorophyta;Mamiel | EP00218 Dolichomastix tenuilepis 6061 | 2,90E-15 | 305 | XP 022002240.1 | protein transport protein SEC16B homolog isoform X1 [Helianthus annuus] | 1331 | 3,59E-17 | 92,8189 | 56,42458101 |
| EP00007 | Amoebozoa;Discosea;Flabellinia;"core flabellinids"; | EP00007 Cunea sp BSH-02190019 9876 | 2,90E-15 | 5930 | XP 002895837.1 | Hect-domain containing peotein, putative [Phytophthora infestans T30-4] | 2300 | 8,66E-63 | 253,062 | 47,28971963 |
| EP00543 | Sar;Stramenopiles;Gyrista;Ochrophyta;"BD clade";[ | EP00543 Licmophora paradoxa 10351 | 3,00E-15 | 986 | XP 002181151.1 | predicted protein [Phaeodactylum tricornutum CCAP 1055/1] | 1133 | 0 | 852,818 | 56,35359116 |
| EP00615 | Sar;Stramenopiles;Gyrista;Ochrophyta;Dictyochoph | EP00615 Pteridomonas danica 11892 | 3,30E-15 | 1181 | XP 009515715.1 | hypothetical protein PHYSODRAFT 322106 [Phytophthora sojae] | 1262 | 0 | 613,609 | 49,28176796 |
| EP00696 | Hemimastigophora;Spironemidae;Hemimastix;Hem | EP00696 Hemimastix kukwesjijk 14916 | 3,30E-15 | 967 | XP 005831033.1 | secretory protein Sec31 [Guillardia theta CCMP2712] | 1081 | 8,76E-119 | 399,053 | 48,7544484 |
| EP00297 | Cryptista;Palpitomonas;Palpitomonas bilix;strain N | EP00297 Palpitomonas bilix 17362 | 3,30E-15 | 181 | ***no hit found*** | | | | | |
| EP00120 | Obazoa;Opisthokonta;Holozoa;Filasterea;Capsaspoı | EP00120 Capsaspora owczarzaki 8232 | 3,40E-15 | 2093 | XP 011270413.1 | hypothetical protein CAOG 08783 [Capsaspora owczarzaki ATCC 30864] | 2093 | 0 | 4261,45 | 100 |
| EP00224 | Archaeplastida;Chloroplastida;Chlorophyta;Mamiel | EP00224 Ostreococcus tauri 6302 | 3,50E-15 | 1080 | XP 022840047.1 | Helicase, C-terminal [Ostreococcus tauri] | 1080 | 0 | 2227,98 | 100 |
| EP00179 | Archaeplastida;"RR clade";Rhodophyta;Proteorhod | EP00179 Madagascaria erythrocladioides 27208 | 3,70E-15 | 435 | XP 003288054.1 | hypothetical protein DICPUDRAFT 152246 [Dictyostelium purpureum] | 1228 | 4,55E-46 | 181,415 | 43,22344322 |
| EP00546 | Sar;Stramenopiles;Gyrista;Ochrophyta;"BD clade";[ | EP00546 Thalassionema frauenfeldii 12639 | 3,90E-15 | 1002 | XP 002290923.1 | predicted protein [Thalassiosira pseudonana CCMP1335] | 1041 | 0 | 938,332 | 54,13891532 |
| EP00044 | Obazoa;Opisthokonta;Holozoa;Choanozoa;Choanof | EP00044 Salpingoeca rosetta 5652 | 3,90E-15 | 1720 | XP 004993085.1 | hypothetical protein PTSG 12412 [Salpingoeca rosetta] | 1720 | 0 | 3362,39 | 100 |
| EP00044 | Obazoa;Opisthokonta;Holozoa;Choanozoa;Choanof | EP00044 Salpingoeca rosetta 862 | 4,00E-15 | 5597 | XP 004988295.1 | hypothetical protein PTSG 13118 [Salpingoeca rosetta] | 5597 | 0 | 11502 | 100 |
| EP00582 | Sar;Stramenopiles;Gyrista;Ochrophyta;"BD clade";[ | EP00582 Thalassiosira oceanica 4901 | 4,10E-15 | 376 | XP 002290923.1 | predicted protein [Thalassiosira pseudonana CCMP1335] | 1041 | 1,13E-122 | 388,267 | 71,31782946 |
| EP00768 | Metamonada;"BF clade";Fornicata;Dysnectes;Dysn | EP00768 Dysnectes brevis 1393 | 5,10E-15 | 371 | XP 025362829.1 | hypothetical protein BDZ90DRAFT 231966 [Jaminaea rosea] | 2332 | 2,17E-07 | 64,3142 | 41,86046512 |
| EP00202 | Archaeplastida;Chloroplastida;Chlorophyta;"core ch | EP00202 Volvox carteri 4184 | 5,20E-15 | 3835 | XP 002949790.1 | hypothetical protein VOLCADRAFT 90172 [Volvox carteri f. nagariensis] | 3835 | 0 | 7476,71 | 100 |
| EP00321 | Haptista;Haptophyta;Prymnesiophyceae;Phaeocyst | EP00321 Phaeocystis sp CCMP2710 7200 | 5,40E-15 | 246 | XP 010150971.1 | PREDICTED: protein FAM184A-like [Eurypyga helias] | 287 | 3,02E-23 | 104,76 | 42,32804233 |
| EP00762 | Discoba;Jakobida;Andalucina;Andaluciidae;Andaluc | EP00762 Andalucia godoyi 3033 | 5,90E-15 | 1036 | XP 006482944.1 | protein transport protein SEC31 homolog B isoform X1 [Citrus sinensis] >gi\|1350271342\| | 1120 | 4,11E-143 | 466,848 | 47,90059983 |
| EP00163 | Ancyromonadida;Fabomonas;Fabomonas tropica;st | EP00163 Fabomonas tropica 33726 | 6,00E-15 | 1069 | XP 017701229.2 | protein transport protein SEC31 homolog B-like [Phoenix dactylifera] | 1044 | 0 | 628,246 | 51,22807018 |
| EP00232 | Archaeplastida;Chloroplastida;Chlorophyta;Nephro | EP00232 Nephroselmis pyriformis 10601 | 6,40E-15 | 166 | XP 002987428.1 | protein transport protein SEC16B homolog [Selaginella moellendorffii] | 1190 | 6,04E-10 | 67,3958 | 43,19526627 |
| EP00023 | Amoebozoa;Evosea;Eumycetozoa;Dictyostelia;Dicty | EP00023 Dictyostelium discoideum 4963 | 6,70E-15 | 802 | XP 638542.1 | hypothetical protein DDB G0284465 [Dictyostelium discoideum AX4] | 802 | 0 | 1556,19 | 100 |
| EP00300 | Haptista;Centroplasthelida;Pterocystida;Raphidista | EP00300 Choanocystis sp HF-7 18813 | 6,80E-15 | 233 | ***no hit found*** | | | | | |
| EP00035 | Obazoa;Opisthokonta;Holozoa;Choanozoa;Choanof | EP00035 Acanthoeca spectabilis 38929 | 7,00E-15 | 1081 | XP 026477575.1 | uncharacterized protein LOC113383530 [Ctenocephalides felis] | 638 | 1,15E-11 | 80,4925 | 39,28571429 |
| EP00540 | Sar;Stramenopiles;Gyrista;Ochrophyta;"BD clade";[ | EP00540 Astrosyne radiata 9999 | 7,30E-15 | 741 | XP 002181151.1 | predicted protein [Phaeodactylum tricornutum CCAP 1055/1] | 1133 | 0 | 622,468 | 55,86734694 |
| EP00376 | Sar;Alveolata;Myzozoa;"AC clade";Apicomplexa;"cc | EP00376 Eimeria tenella 5585 | 7,40E-15 | 2184 | XP 013233318.1 | hypothetical protein, conserved [Eimeria tenella] | 2184 | 0 | 4115,84 | 100 |
| EP00247 | Archaeplastida;Chloroplastida;Streptophyta;"core s | EP00247 Physcomitrella patens 7824 | 7,40E-15 | 1167 | XP 024364075.1 | protein transport protein SEC31 homolog B-like isoform X2 [Physcomitrella patens] | 1167 | 0 | 2386,3 | 100 |
| EP00594 | Sar;Stramenopiles;Gyrista;Ochrophyta;"BD clade";[ | EP00594 Rhizosolenia setigera 4213 | 7,50E-15 | 1108 | XP 002290923.1 | predicted protein [Thalassiosira pseudonana CCMP1335] | 1041 | 0 | 994,186 | 56,68161435 |
| EP00578 | Sar;Stramenopiles;Gyrista;Ochrophyta;"BD clade";[ | EP00578 Thalassiosira sp NH16 17595 | 7,80E-15 | 328 | XP 002183614.1 | predicted protein [Phaeodactylum tricornutum CCAP 1055/1] | 2083 | 3,06E-09 | 69,707 | 37,5 |
| EP00223 | Archaeplastida;Chloroplastida;Chlorophyta;Mamiel | EP00223 Ostreococcus mediterraneus 1505 | 8,00E-15 | 1238 | XP 001420597.1 | predicted protein, partial [Ostreococcus lucimarinus CCE9901] | 1077 | 0 | 1226,08 | 82,69754768 |
| EP00168 | Archaeplastida;"RR clade";Rhodophyta;Eurhodophy | EP00168 Porphyra purpurea 4978 | 8,30E-15 | 541 | XP 005711041.1 | unnamed protein product [Chondrus crispus] | 2546 | 3,74E-09 | 71,2478 | 39,00414938 |
| EP00231 | Archaeplastida;Chloroplastida;Chlorophyta;Mamiel | EP00231 Micromonas pusilla 211 | 8,60E-15 | 1138 | XP 003054966.1 | predicted protein [Micromonas pusilla CCMP1545] | 1138 | 0 | 2301,94 | 100 |
| EP00260 | Archaeplastida;Chloroplastida;Streptophyta;"core s | EP00260 Arabidopsis thaliana 5598 | 8,60E-15 | 1094 | NP 001190171.1 | transducin family protein / WD-40 repeat family protein [Arabidopsis thaliana] | 1094 | 0 | 2268,43 | 100 |
| EP00376 | Sar;Alveolata;Myzozoa;"AC clade";Apicomplexa;"cc | EP00376 Eimeria tenella 3070 | 8,70E-15 | 930 | XP 013230800.1 | hypothetical protein, conserved [Eimeria tenella] | 930 | 0 | 1776,91 | 100 |
| EP00024 | Amoebozoa;Evosea;Eumycetozoa;Dictyostelia;Dicty | EP00024 Dictyostelium purpureum 5202 | 9,50E-15 | 1228 | XP 003288054.1 | hypothetical protein DICPUDRAFT 152246 [Dictyostelium purpureum] | 1228 | 0 | 2531,52 | 100 |
| EP00376 | Sar;Alveolata;Myzozoa;"AC clade";Apicomplexa;"cc | EP00376 Eimeria tenella 6764 | 1,10E-14 | 973 | XP 013234499.1 | hypothetical protein ETH 00009770, partial [Eimeria tenella] | 973 | 0 | 1788,85 | 100 |
| EP00019 | Amoebozoa;Evosea;Cutosea;Armaparvus;Armaparv | EP00019 Armaparvus languidus 10698 | 1,30E-14 | 353 | ***no hit found*** | | | | | |
| EP00280 | Cryptista;Cryptophyceae;Cryptomonadales;"lineage | EP00280 Hanusia phi 571 | 1,40E-14 | 433 | ***no hit found*** | | | | | |
| EP00211 | Archaeplastida;Chloroplastida;Chlorophyta;Chlorop | EP00211 Chloroparvula japonica 131 | 1,50E-14 | 1095 | XP 024379159.1 | protein transport protein SEC31 homolog B-like isoform X2 [Physcomitrella patens] | 1166 | 0 | 791,956 | 51,28424658 |
| EP00605 | Sar;Stramenopiles;Gyrista;Ochrophyta;"CS clade";C | EP00605 Chrysophyceae sp TOSAG23-4 2180 | 1,70E-14 | 1035 | XP 002290923.1 | predicted protein [Thalassiosira pseudonana CCMP1335] | 1041 | 0 | 617,846 | 46,55647383 |
| EP00738 | Archaeplastida;Chloroplastida;Chlorophyta;Chlorop | EP00738 Chloropicon primus 5642 | 1,70E-14 | 647 | XP 019445058.1 | PREDICTED: protein transport protein SEC16A homolog isoform X2 [Lupinus angustifolius] | 1422 | 6,84E-06 | 61,2326 | 46,83544304 |
| EP00376 | Sar;Alveolata;Myzozoa;"AC clade";Apicomplexa;"cc | EP00376 Eimeria tenella 7920 | 1,70E-14 | 2214 | XP 013235662.1 | hypothetical protein ETH 00017500 [Eimeria tenella] | 2214 | 0 | 4264,53 | 100 |
| EP00445 | Sar;Alveolata;Myzozoa;"DP clade";Dinoflagellata;D | EP00445 Apocalathium hangoei 85317 | 1,80E-14 | 660 | ***no hit found*** | | | | | |
| EP00199 | Archaeplastida;Chloroplastida;Chlorophyta;"core ch | EP00199 Chlamydomonas sp CCMP681 1514 | 1,80E-14 | 929 | XP 002948107.1 | hypothetical protein VOLCADRAFT 120559 [Volvox carteri f. nagariensis] | 1302 | 1,27E-129 | 431,795 | 59,51742627 |
| EP00467 | Sar;Rhizaria;Cercozoa;Chlorarachnea;"core chlorara | EP00467 Chlorarachnion reptans 17407 | 2,20E-14 | 197 | ***no hit found*** | | | | | |
| EP00125 | Obazoa;Opisthokonta;Holozoa;Ichthyosporea;Ichthy | EP00125 Sphaeroforma arctica 16619 | 2,20E-14 | 486 | XP 014159694.1 | hypothetical protein SARC 02055 [Sphaeroforma arctica JP610] | 486 | 0 | 995,342 | 100 |
| EP00103 | Obazoa;Opisthokonta;Holozoa;Choanozoa;Metazoa | EP00103 Capitella teleta 21379 | 2,30E-14 | 1165 | XP 013384747.1 | protein transport protein Sec31A-like [Lingula anatina] | 1860 | 0 | 1219,14 | 61,5 |
| EP00566 | Sar;Stramenopiles;Gyrista;Ochrophyta;"BD clade";[ | EP00566 Odontella aurita 2062 | 2,40E-14 | 1078 | XP 002290923.1 | predicted protein [Thalassiosira pseudonana CCMP1335] | 1041 | 0 | 1161,36 | 62,10045662 |
| EP00566 | Sar;Stramenopiles;Gyrista;Ochrophyta;"BD clade";[ | EP00566 Odontella aurita 36466 | 2,40E-14 | 1078 | XP 002290923.1 | predicted protein [Thalassiosira pseudonana CCMP1335] | 1041 | 0 | 1161,36 | 62,10045662 |
| EP00536 | Sar;Stramenopiles;Gyrista;Ochrophyta;"BD clade";[ | EP00536 Pseudo-nitzschia multiseries 11688 | 2,40E-14 | 431 | XP 002183614.1 | predicted protein [Phaeodactylum tricornutum CCAP 1055/1] | 2083 | 3,77E-12 | 80,1073 | 53,48837209 |
| EP00159 | Obazoa;Opisthokonta;Nucletmycea;Rotosphaerida; | EP00159 Fonticula alba 2644 | 2,50E-14 | 3536 | XP 009494760.1 | hypothetical protein H696 02574 [Fonticula alba] | 3536 | 0 | 7050,68 | 100 |
| EP00553 | Sar;Stramenopiles;Gyrista;Ochrophyta;"BD clade";[ | EP00553 Chaetoceros curvisetus 625 | 2,50E-14 | 614 | XP 002181151.1 | predicted protein [Phaeodactylum tricornutum CCAP 1055/1] | 1133 | 1,36E-126 | 409,838 | 52,64847512 |
| EP00376 | Sar;Alveolata;Myzozoa;"AC clade";Apicomplexa;"cc | EP00376 Eimeria tenella 3065 | 2,50E-14 | 3159 | XP 013230795.1 | DNA polymerase, putative [Eimeria tenella] | 3159 | 0 | 6267,18 | 100 |
| EP00264 | Archaeplastida;Chloroplastida;Streptophyta;"core s | EP00264 Hordeum vulgare 57073 | 2,70E-14 | 354 | XP 020187484.1 | protein transport protein SEC31 homolog B isoform X2 [Aegilops tauschii subsp. tauschii] | 1134 | 0 | 687,567 | 95,48022599 |
| EP00232 | Archaeplastida;Chloroplastida;Chlorophyta;Nephro | EP00232 Nephroselmis pyriformis 21148 | 2,90E-14 | 339 | XP 013713824.1 | protein transport protein SEC16A homolog isoform X2 [Brassica napus] | 1333 | 4,76E-08 | 65,855 | 54,42176871 |
| EP00566 | Sar;Stramenopiles;Gyrista;Ochrophyta;"BD clade";[ | EP00566 Odontella aurita 2098 | 2,90E-14 | 1078 | XP 002290923.1 | predicted protein [Thalassiosira pseudonana CCMP1335] | 1041 | 0 | 1162,13 | 62,22627737 |
| EP00557 | Sar;Stramenopiles;Gyrista;Ochrophyta;"BD clade";[ | EP00557 Chaetoceros sp GSL56 11507 | 3,20E-14 | 1118 | XP 002290923.1 | predicted protein [Thalassiosira pseudonana CCMP1335] | 1041 | 0 | 810,446 | 55,74193548 |
| EP00133 | Obazoa;Opisthokonta;Nucletmycea;Fungi;"core Fur | EP00133 Fusarium graminearum 9354 | 3,30E-14 | 1294 | XP 011324915.1 | hypothetical protein FGSG 06265 [Fusarium graminearum PH-1] | 1294 | 0 | 2629,74 | 100 |
| EP00019 | Amoebozoa;Evosea;Cutosea;Armaparvus;Armaparv | EP00019 Armaparvus languidus 4367 | 3,30E-14 | 854 | XP 003288054.1 | hypothetical protein DICPUDRAFT 152246 [Dictyostelium purpureum] | 1228 | 2,90E-169 | 533,102 | 49,21171171 |
| EP00019 | Amoebozoa;Evosea;Cutosea;Armaparvus;Armaparv | EP00019 Armaparvus languidus 6296 | 3,40E-14 | 195 | ***no hit found*** | | | | | |
| EP00545 | Sar;Stramenopiles;Gyrista;Ochrophyta;"BD clade";[ | EP00545 Synedropsis sp CCMP1620 273 | 3,50E-14 | 969 | XP 002176596.1 | predicted protein [Phaeodactylum tricornutum CCAP 1055/1] | 752 | 2,56E-92 | 319,701 | 58,51979346 |
| EP00234 | Archaeplastida;Chloroplastida;Palmophyllophyceae | EP00234 Prasinoderma coloniale 5727 | 3,50E-14 | 1185 | XP 020519257.1 | protein transport protein SEC31 homolog B isoform X2 [Amborella trichopoda] | 1154 | 0 | 766,918 | 50,41118421 |
| EP00593 | Sar;Stramenopiles;Gyrista;Ochrophyta;"BD clade";[ | EP00593 Proboscia inermis 12009 | 3,60E-14 | 80 | XP 002183614.1 | predicted protein [Phaeodactylum tricornutum CCAP 1055/1] | 2083 | 2,61E-13 | 73,1738 | 54,92957746 |
| EP00152 | Obazoa;Opisthokonta;Nucletmycea;Fungi;"core Fur | EP00152 Rhizopus delemar 1905 | 4,10E-14 | 673 | XP 023467476.1 | WD40 repeat-like protein [Rhizopus microsporus ATCC 52813] | 720 | 0 | 666,766 | 60,81081081 |
| EP00019 | Amoebozoa;Evosea;Cutosea;Armaparvus;Armaparv | EP00019 Armaparvus languidus 3887 | 4,20E-14 | 1134 | XP 012754771.1 | hypothetical protein SAMD00019534 058260 [Acytostelium subglobosum LB1] | 1179 | 4,31E-174 | 553,518 | 44,3012884 |
| EP00545 | Sar;Stramenopiles;Gyrista;Ochrophyta;"BD clade";[ | EP00545 Synedropsis sp CCMP1620 6889 | 4,20E-14 | 708 | XP 002181151.1 | predicted protein [Phaeodactylum tricornutum CCAP 1055/1] | 1133 | 0 | 614,764 | 56,84754522 |
| EP00325 | Haptista;Haptophyta;Prymnesiophyceae;Prymnesia | EP00325 Haptolina brevifila 108 | 4,30E-14 | 223 | ***no hit found*** | | | | | |
| EP00376 | Sar;Alveolata;Myzozoa;"AC clade";Apicomplexa;"cc | EP00376 Eimeria tenella 2755 | 4,70E-14 | 940 | XP 013230483.1 | hypothetical protein, conserved [Eimeria tenella] | 940 | 0 | 1812,74 | 100 |
| EP00231 | Archaeplastida;Chloroplastida;Chlorophyta;Mamiel | EP00231 Micromonas pusilla 4392 | 4,80E-14 | 1002 | XP 003059147.1 | predicted protein [Micromonas pusilla CCMP1545] | 1002 | 0 | 1922,13 | 100 |
| EP00376 | Sar;Alveolata;Myzozoa;"AC clade";Apicomplexa;"cc | EP00376 Eimeria tenella 4063 | 5,00E-14 | 854 | XP 013231796.1 | hypothetical protein, conserved [Eimeria tenella] | 854 | 0 | 1597,02 | 100 |
| EP00556 | Sar;Stramenopiles;Gyrista;Ochrophyta;"BD clade";[ | EP00556 Chaetoceros neogracilis 7877 | 5,00E-14 | 1050 | XP 002295126.1 | small GTPase, partial [Thalassiosira pseudonana CCMP1335] | 223 | 2,57E-30 | 130,183 | 44,90740741 |
| EP00675 | Discoba;Euglenozoa;Kinetoplastea;Metakinetoplast | EP00675 Leptomonas pyrrhocoris 2264 | 5,00E-14 | 1298 | XP 015653768.1 | hypothetical protein ABB37 08623 [Leptomonas pyrrhocoris] | 1298 | 0 | 2633,6 | 100 |
| EP00594 | Sar;Stramenopiles;Gyrista;Ochrophyta;"BD clade";[ | EP00594 Rhizosolenia setigera 12073 | 5,70E-14 | 757 | XP 002290923.1 | predicted protein [Thalassiosira pseudonana CCMP1335] | 1041 | 0 | 630,172 | 56,51612903 |
| EP00376 | Sar;Alveolata;Myzozoa;"AC clade";Apicomplexa;"cc | EP00376 Eimeria tenella 5607 | 5,70E-14 | 4739 | XP 013233340.1 | hypothetical protein, conserved [Eimeria tenella] | 4739 | 0 | 9559,48 | 100 |
| EP00612 | Sar;Stramenopiles;Gyrista;Ochrophyta;"PX clade";X | EP00612 Vaucheria litorea 6457 | 5,70E-14 | 1148 | XP 002290923.1 | predicted protein [Thalassiosira pseudonana CCMP1335] | 1041 | 0 | 645,966 | 53,14505777 |
| EP00567 | Sar;Stramenopiles;Gyrista;Ochrophyta;"BD clade";[ | EP00567 Triceratium dubium 14749 | 6,20E-14 | 71 | XP 002183614.1 | predicted protein [Phaeodactylum tricornutum CCAP 1055/1] | 2083 | 4,82E-10 | 63,5438 | 49,05660377 |
| EP00390 | Sar;Alveolata;Myzozoa;"AC clade";Colpodellida;Vitr | EP00390 Vitrella brassicaformis 2818 | 6,40E-14 | 1948 | XP 014529636.1 | hypothetical protein JH06 0320 [Blastocystis sp. subtype 4] | 318 | 1,91E-10 | 75,485 | 51,72413793 |
| EP00044 | Obazoa;Opisthokonta;Holozoa;Choanozoa;Choanof | EP00044 Salpingoeca rosetta 10581 | 6,70E-14 | 3529 | XP 004998014.1 | hypothetical protein PTSG 11826 [Salpingoeca rosetta] | 3529 | 0 | 7217,86 | 100 |
| EP00566 | Sar;Stramenopiles;Gyrista;Ochrophyta;"BD clade";[ | EP00566 Odontella aurita 2147 | 6,80E-14 | 1078 | XP 002290923.1 | predicted protein [Thalassiosira pseudonana CCMP1335] | 1041 | 0 | 1160,21 | 62,00913242 |
| EP00268 | Archaeplastida;Chloroplastida;Streptophyta;"core s | EP00268 Persea americana 17114 | 6,80E-14 | 485 | XP 010279523.1 | PREDICTED: protein transport protein SEC31 homolog B-like [Nelumbo nucifera] | 1129 | 0 | 639,032 | 66,10526316 |
| EP00019 | Amoebozoa;Evosea;Cutosea;Armaparvus;Armaparv | EP00019 Armaparvus languidus 3956 | 8,00E-14 | 1118 | XP 012754771.1 | hypothetical protein SAMD00019534 058260 [Acytostelium subglobosum LB1] | 1179 | 5,59E-174 | 552,362 | 45,05928854 |
| EP00319 | Haptista;Haptophyta;Prymnesiophyceae;Phaeocyst | EP00319 Phaeocystis globosa 9684 | 9,80E-14 | 387 | XP 005831033.1 | secretory protein Sec31 [Guillardia theta CCMP2712] | 1081 | 4,27E-72 | 253,832 | 49,69512195 |
| EP00376 | Sar;Alveolata;Myzozoa;"AC clade";Apicomplexa;"cc | EP00376 Eimeria tenella 5720 | 1,00E-13 | 2313 | XP 013233453.1 | hypothetical protein, conserved [Eimeria tenella] | 2313 | 0 | 4439,03 | 100 |
| EP00376 | Sar;Alveolata;Myzozoa;"AC clade";Apicomplexa;"cc | EP00376 Eimeria tenella 4815 | 1,30E-13 | 3151 | XP 013232548.1 | hypothetical protein, conserved [Eimeria tenella] | 3151 | 0 | 6094,23 | 100 |
| EP00494 | Sar;Rhizaria;Retaria;Radiolaria;Acantharea;Astrolor | EP00494 Astrolonche serrata 19923 | 1,40E-13 | 134 | XP 020180505.1 | protein transport protein SEC16B homolog [Aegilops tauschii subsp. tauschii] | 1647 | 4,34E-33 | 132,494 | 91,42857143 |
| EP00242 | Archaeplastida;Chloroplastida;Chlorophyta;Pyramir | EP00242 Pyramimonas sp CCMP2087 13024 | 1,40E-13 | 231 | XP 024379159.1 | protein transport protein SEC31 homolog B-like isoform X2 [Physcomitrella patens] | 1166 | 5,49E-57 | 205,297 | 53,33333333 |
| EP00560 | Sar;Stramenopiles;Gyrista;Ochrophyta;"BD clade";[ | EP00560 Eucampia antarctica 5826 | 1,60E-13 | 1037 | XP 002290923.1 | predicted protein [Thalassiosira pseudonana CCMP1335] | 1041 | 0 | 832,402 | 65,00711238 |
| EP00247 | Archaeplastida;Chloroplastida;Streptophyta;"core s | EP00247 Physcomitrella patens 7814 | 1,60E-13 | 1195 | XP 024364065.1 | protein transport protein SEC31 homolog B-like isoform X1 [Physcomitrella patens] | 1195 | 0 | 2444,46 | 100 |
| EP00628 | Sar;Stramenopiles;Gyrista;Ochrophyta;Pelagophyce | EP00628 Pelagophyceae sp CCMP2097 12126 | 1,70E-13 | 1223 | XP 009038384.1 | hypothetical protein AURANDRAFT 28308, partial [Aureococcus anophagefferens] | 728 | 0 | 815,068 | 67,97297297 |
| EP00007 | Amoebozoa;Discosea;Flabellinia;"core flabellinids"; | EP00007 Cunea sp BSH-02190019 2153 | 1,70E-13 | 5547 | XP 022071969.1 | probable proline--tRNA ligase, mitochondrial [Acanthochromis polyacanthus] | 483 | 2,56E-103 | 352,829 | 52,38095238 |
| EP00376 | Sar;Alveolata;Myzozoa;"AC clade";Apicomplexa;"cc | EP00376 Eimeria tenella 8062 | 1,80E-13 | 1297 | XP 013235804.1 | hypothetical protein ETH 00003945 [Eimeria tenella] | 1297 | 0 | 2441,38 | 100 |
| EP00770 | Metamonada;Preaxostyla;Oxymonadida;Monocerc | EP00770 Monocercomonoides exilis 2464 | 2,00E-13 | 4038 | XP 008879363.1 | hypothetical protein H310 13588 [Aphanomyces invadans] | 1853 | 3,03E-34 | 157,918 | 50,71283096 |
| EP00051 | Obazoa;Opisthokonta;Holozoa;Choanozoa;Choanof | EP00051 Salpingoeca urceolata 16636 | 2,00E-13 | 891 | XP 001634737.1 | predicted protein [Nematostella vectensis] | 1219 | 0 | 575,859 | 49,6009122 |
| EP00551 | Sar;Stramenopiles;Gyrista;Ochrophyta;"BD clade";[ | EP00551 Chaetoceros affinis 12722 | 2,00E-13 | 1062 | XP 002290923.1 | predicted protein [Thalassiosira pseudonana CCMP1335] | 1041 | 0 | 937,176 | 51,78236398 |
| EP00284 | Cryptista;Cryptophyceae;Cryptomonadales;"lineage | EP00284 Hemiselmis tepida 5388 | 2,20E-13 | 66 | XP 026401447.1 | protein transport protein SEC16A homolog isoform X2 [Papaver somniferum] | 1553 | 1,99E-07 | 55,8398 | 45,90163934 |
| EP00547 | Sar;Stramenopiles;Gyrista;Ochrophyta;"BD clade";[ | EP00547 Thalassionema nitzschioides 4476 | 2,20E-13 | 1002 | XP 002290923.1 | predicted protein [Thalassiosira pseudonana CCMP1335] | 1041 | 0 | 948,347 | 54,51950523 |
| EP00472 | Sar;Rhizaria;Cercozoa;Chlorarachnea;"core chlorara | EP00472 Partenskyella glossopodia 5994 | 2,40E-13 | 744 | XP 024188898.1 | uncharacterized protein LOC112193100 [Rosa chinensis] | 278 | 1,58E-29 | 129,028 | 48,57142857 |
| EP00299 | Haptista;Centroplasthelida;Pterocystida;Pterista;"P | EP00299 Pterocystis canadensis 17687 | 2,50E-13 | 236 | XP 018914532.1 | PREDICTED: eisosome protein sle1-like isoform X1 [Bemisia tabaci] | 400 | 1,88E-06 | 58,9214 | 51,5625 |
| EP00299 | Haptista;Centroplasthelida;Pterocystida;Pterista;"P | EP00299 Pterocystis canadensis 17690 | 2,50E-13 | 236 | XP 018914532.1 | PREDICTED: eisosome protein sle1-like isoform X1 [Bemisia tabaci] | 400 | 1,88E-06 | 58,9214 | 51,5625 |
| EP00088 | Obazoa;Opisthokonta;Holozoa;Choanozoa;Metazoa | EP00088 Acartia fossae 24576 | 2,60E-13 | 107 | XP 023346731.1 | protein transport protein Sec16-like isoform X2 [Eurytemora affinis] | 1941 | 9,76E-15 | 78,5666 | 73,33333333 |
| EP00739 | Archaeplastida;Chloroplastida;Chlorophyta;Mamiel | EP00739 Bathycoccus prasinos 5129 | 2,80E-13 | 926 | XP 007513334.1 | predicted protein [Bathycoccus prasinos] | 854 | 0 | 1708,35 | 99,29988331 |
| EP00270 | Archaeplastida;Chloroplastida;Streptophyta;"core s | EP00270 Netrium digitus 1582 | 2,80E-13 | 273 | XP 017234870.1 | PREDICTED: protein transport protein SEC16B homolog [Daucus carota subsp. sativus] | 1421 | 2,17E-10 | 72,0182 | 53,42465753 |
| EP00115 | Obazoa;Opisthokonta;Holozoa;Choanozoa;Metazoa | EP00115 Mnemiopsis leidyi 1067 | 2,90E-13 | 1106 | XP 030420897.1 | protein transport protein Sec31A isoform X6 [Gopherus evgoodei] | 1225 | 0 | 757,288 | 50,64516129 |
| EP00468 | Sar;Rhizaria;Cercozoa;Chlorarachnea;"core chlorara | EP00468 Gymnochlora sp CCMP2014 2814 | 2,90E-13 | 924 | ***no hit found*** | | | | | |
| EP00010 | Amoebozoa;Discosea;Flabellinia;"core flabellinids"; | EP00010 Vexillifera sp DIVA3-564-2 2422 | 3,20E-13 | 882 | XP 007887153.1 | PREDICTED: protein transport protein Sec31A isoform X5 [Callorhinchus milii] | 1153 | 1,83E-90 | 320,857 | 42,60614934 |
| EP00642 | Sar;Stramenopiles;Gyrista;Peronosporomycetes;"cc | EP00642 Albugo laibachii 7578 | 3,20E-13 | 1186 | XP 009515715.1 | hypothetical protein PHYSODRAFT 322106 [Phytophthora sojae] | 1262 | 0 | 1403,65 | 66,25194401 |
| EP00617 | Sar;Stramenopiles;Gyrista;Ochrophyta;Dictyochoph | EP00617 Octactis speculum 24864 | 3,20E-13 | 1150 | XP 002290923.1 | predicted protein [Thalassiosira pseudonana CCMP1335] | 1041 | 0 | 662,144 | 52,57819104 |
| EP00585 | Sar;Stramenopiles;Gyrista;Ochrophyta;"BD clade";[ | EP00585 Thalassiosira rotula 22289 | 3,20E-13 | 538 | XP 002181151.1 | predicted protein [Phaeodactylum tricornutum CCAP 1055/1] | 1133 | 6,31E-118 | 384,03 | 52,7972028 |
| EP00581 | Sar;Stramenopiles;Gyrista;Ochrophyta;"BD clade";[ | EP00581 Thalassiosira minuscula 4567 | 3,30E-13 | 1037 | XP 002290314.1 | predicted protein [Thalassiosira pseudonana CCMP1335] | 772 | 1,81E-153 | 483,797 | 71,7037037 |
| EP00592 | Sar;Stramenopiles;Gyrista;Ochrophyta;"BD clade";[ | EP00592 Proboscia alata 7733 | 3,30E-13 | 1062 | XP 002290923.1 | predicted protein [Thalassiosira pseudonana CCMP1335] | 1041 | 0 | 993,03 | 58,04066543 |
| EP00184 | Archaeplastida;"RR clade";Rhodophyta;Proteorhod | EP00184 Porphyridium aerugineum 5040 | 3,50E-13 | 659 | ***no hit found*** | | | | | |
| EP00376 | Sar;Alveolata;Myzozoa;"AC clade";Apicomplexa;"cc | EP00376 Eimeria tenella 5304 | 3,90E-13 | 1232 | XP 013233037.1 | hypothetical protein ETH 00000205 [Eimeria tenella] | 1232 | 0 | 2315,42 | 100 |
| EP00376 | Sar;Alveolata;Myzozoa;"AC clade";Apicomplexa;"cc | EP00376 Eimeria tenella 128 | 3,90E-13 | 1395 | XP 013227854.1 | WD-40 repeat protein, putative [Eimeria tenella] | 1395 | 0 | 2710,25 | 100 |

| | | | | | | | | | | |
|---|---|---|---|---|---|---|---|---|---|---|
| EP00608 | Sar;Stramenopiles;Gyrista;Ochrophyta;"CS clade";S | EP00608 Synchroma pusillum 7855 | 4,30E-13 | 418 | XP 002290923.1 | predicted protein [Thalassiosira pseudonana CCMP1335] | 1041 | 1,07E-89 | 302,368 | 49,30747922 |
| EP00541 | Sar;Stramenopiles;Gyrista;Ochrophyta;"BD clade";D | EP00541 Cyclophora tenuis 5712 | 4,30E-13 | 453 | XP 002183614.1 | predicted protein [Phaeodactylum tricornutum CCAP 1055/1] | 2083 | 1,81E-31 | 139,043 | 61,30952381 |
| EP00053 | Obazoa;Opisthokonta;Holozoa;Choanozoa;Choanof | EP00053 Salpingoeca punica 616 | 4,50E-13 | 768 | XP 002602930.1 | hypothetical protein BRAFLDRAFT 62485 [Branchiostoma floridae] | 1113 | 0 | 568,54 | 48,2278481 |
| EP00013 | Amoebozoa;Discosea;Flabellinia;Stygamoeba;Styga | EP00013 Stygamoeba regulata 19222 | 5,00E-13 | 860 | XP 005835801.1 | hypothetical protein GUITHDRAFT 151698 [Guillardia theta CCMP2712] | 367 | 7,84E-32 | 138,658 | 46,62576687 |
| EP00462 | Sar;Rhizaria;Cercozoa;"crown cercozoa";Thecofilose | EP00462 Mataza sp D1 26369 | 6,60E-13 | 89 | XP 013892991.1 | hypothetical protein MNEG 13990 [Monoraphidium neglectum] | 818 | 4,06E-08 | 58,9214 | 56,06060606 |
| EP00291 | Cryptista;Cryptophyceae;Cryptomonadales;Crypton | EP00291 Cryptomonas curvata 3481 | 6,60E-13 | 68 | XP 011312055.1 | PREDICTED: uncharacterized protein LOC105271924 [Fopius arisanus] | 2092 | 3,72E-07 | 55,0694 | 48,14814815 |
| EP00314 | Haptista;Haptophyta;Prymnesiophyceae;Calcihapto | EP00314 Emiliania huxleyi 34360 | 6,70E-13 | 4317 | XP 005790421.1 | hypothetical protein EMIHUDRAFT 97896 [Emiliania huxleyi CCMP1516] | 4317 | 0 | 8548,33 | 100 |
| EP00198 | Archaeplastida;Chloroplastida;Chlorophyta;"core ch | EP00198 Chlamydomonas reinhardtii 7702 | 6,80E-13 | 1313 | XP 001697039.1 | COP-II coat subunit [Chlamydomonas reinhardtii] | 1313 | 0 | 2672,5 | 100 |
| EP00536 | Sar;Stramenopiles;Gyrista;Ochrophyta;"BD clade";D | EP00536 Pseudo-nitzschia multiseries 4156 | 7,00E-13 | 1755 | ***no hit found*** | | | | | |
| EP00116 | Obazoa;Opisthokonta;Holozoa;Choanozoa;Metazoa | EP00116 Pleurobrachia bachei 4994 | 7,00E-13 | 406 | XP 020600558.1 | serine/threonine-protein kinase pim-2-like [Orbicella faveolata] | 426 | 8,08E-28 | 124,02 | 39,03345725 |
| EP00429 | Sar;Alveolata;Myzozoa;"DP clade";Dinoflagellata;D | EP00429 Kryptoperidinium foliaceum 63774 | 7,40E-13 | 867 | XP 002773527.1 | conserved hypothetical protein [Perkinsus marinus ATCC 50983] | 1211 | 0 | 645,966 | 50,05931198 |
| EP00372 | Sar;Alveolata;Myzozoa;"AC clade";Apicomplexa;"cc | EP00372 Theileria equi 4938 | 7,60E-13 | 957 | XP 004833510.1 | conserved hypothetical protein [Theileria equi strain WA] | 957 | 0 | 1971,82 | 100 |
| EP00376 | Sar;Alveolata;Myzozoa;"AC clade";Apicomplexa;"cc | EP00376 Eimeria tenella 3311 | 7,70E-13 | 1202 | XP 013231042.1 | thiF family domain-containing protein, putative [Eimeria tenella] | 1202 | 0 | 2420,19 | 100 |
| EP00054 | Obazoa;Opisthokonta;Holozoa;Choanozoa;Choanof | EP00054 Salpingoeca dolichothecata 15646 | 9,30E-13 | 906 | XP 007887151.1 | PREDICTED: protein transport protein Sec31A isoform X3 [Callorhinchus milii] | 1236 | 0 | 588,186 | 51,65816327 |
| EP00230 | Archaeplastida;Chloroplastida;Chlorophyta;Mamiel | EP00230 Micromonas polaris 4503 | 9,40E-13 | 1259 | XP 002500425.1 | predicted protein [Micromonas commoda] | 1395 | 1,06E-35 | 159,844 | 59,65665236 |
| EP00188 | Archaeplastida;"RR clade";Rhodophyta;Proteorhod | EP00188 Purpureofilum apyrenoidigerum 6041 | 9,60E-13 | 125 | XP 023635307.1 | protein transport protein SEC16A homolog isoform X2 [Capsella rubella] | 1387 | 4,53E-07 | 57,3806 | 38,88888889 |
| EP00512 | Sar;Stramenopiles;Bigyra;Sagenista;Labyrinthulomy | EP00512 Aurantiochytrium limacinum 5049 | 9,60E-13 | 964 | XP 006482944.1 | protein transport protein SEC31 homolog B isoform X1 [Citrus sinensis] >gi\|1350271342\|r | 1120 | 6,28E-51 | 206,453 | 43,52557127 |
| EP00590 | Sar;Stramenopiles;Gyrista;Ochrophyta;"BD clade";D | EP00590 Aulacoseira subarctica 3442 | 1,00E-12 | 1069 | XP 002290923.1 | predicted protein [Thalassiosira pseudonana CCMP1335] | 1041 | 0 | 940,643 | 56,43115942 |
| EP00271 | Archaeplastida;Chloroplastida;Streptophyta;"core s | EP00271 Cylindrocystis brebissonii 6221 | 1,10E-12 | 1272 | XP 028119649.1 | protein transport protein SEC31 homolog B-like [Camellia sinensis] | 1118 | 0 | 786,178 | 54,10216718 |
| EP00026 | Amoebozoa;Evosea;Eumycetozoa;Myxogastria;Colu | EP00026 Physarum polycephalum 2399 | 1,20E-12 | 914 | XP 005831033.1 | secretory protein Sec31 [Guillardia theta CCMP2712] | 1081 | 0 | 586,26 | 51,66297118 |
| EP00555 | Sar;Stramenopiles;Gyrista;Ochrophyta;"BD clade";D | EP00555 Chaetoceros dichaeta 26 | 1,20E-12 | 923 | XP 002181151.1 | predicted protein [Phaeodactylum tricornutum CCAP 1055/1] | 1133 | 0 | 721,85 | 55,72289157 |
| EP00706 | Metamonada;Parabasalia;Tritrichomonas;Tritrichor | EP00706 Tritrichomonas foetus 2069 | 1,30E-12 | 920 | XP 001580855.1 | hypothetical protein [Trichomonas vaginalis G3] | 2050 | 1,94E-22 | 115,931 | 38,97849462 |
| EP00051 | Obazoa;Opisthokonta;Holozoa;Choanozoa;Choanof | EP00051 Salpingoeca urceolata 25201 | 1,30E-12 | 633 | XP 001748484.1 | hypothetical protein [Monosiga brevicollis MX1] | 442 | 1,08E-28 | 129,798 | 41,89189189 |
| EP00247 | Archaeplastida;Chloroplastida;Streptophyta;"core s | EP00247 Physcomitrella patens 22899 | 1,30E-12 | 1179 | XP 024379150.1 | protein transport protein SEC31 homolog B-like isoform X1 [Physcomitrella patens] | 1179 | 0 | 2414,03 | 100 |
| EP00446 | Sar;Alveolata;Myzozoa;"DP clade";Dinoflagellata;D | EP00446 Apocalathium sp SHHI-4 79387 | 1,40E-12 | 753 | XP 002773527.1 | conserved hypothetical protein [Perkinsus marinus ATCC 50983] | 1211 | 0 | 604,749 | 51,83312263 |
| EP00376 | Sar;Alveolata;Myzozoa;"AC clade";Apicomplexa;"cc | EP00376 Eimeria tenella 5021 | 1,40E-12 | 3666 | XP 013232754.1 | hypothetical protein, conserved [Eimeria tenella] | 3666 | 0 | 7486,34 | 100 |
| EP00462 | Sar;Rhizaria;Cercozoa;"crown cercozoa";Thecofilose | EP00462 Mataza sp D1 24759 | 1,50E-12 | 1253 | XP 020585788.1 | protein transport protein SEC31 homolog B isoform X2 [Phalaenopsis equestris] | 1125 | 1,08E-124 | 423,32 | 43,47826087 |
| EP00512 | Sar;Stramenopiles;Bigyra;Sagenista;Labyrinthulomy | EP00512 Aurantiochytrium limacinum 2158 | 1,50E-12 | 1312 | XP 008907364.1 | hypothetical protein, variant 2 [Phytophthora parasitica INRA-310] | 1175 | 6,27E-97 | 347,821 | 51,96428571 |
| EP00191 | Archaeplastida;Chloroplastida;Chlorophyta;"core ch | EP00191 Tetraselmis sp GSL018 11699 | 1,60E-12 | 1134 | XP 006837403.2 | protein transport protein SEC31 homolog B isoform X1 [Amborella trichopoda] | 1155 | 0 | 804,668 | 48,98989899 |
| EP00558 | Sar;Stramenopiles;Gyrista;Ochrophyta;"BD clade";D | EP00558 Chaetoceros sp UNC1202 13907 | 1,70E-12 | 150 | XP 002290923.1 | predicted protein [Thalassiosira pseudonana CCMP1335] | 1041 | 9,18E-59 | 206,068 | 65,54054054 |
| EP00586 | Sar;Stramenopiles;Gyrista;Ochrophyta;"BD clade";D | EP00586 Coscinodiscus wailesii 2928 | 1,70E-12 | 784 | XP 002290923.1 | predicted protein [Thalassiosira pseudonana CCMP1335] | 1041 | 0 | 634,41 | 55,72519084 |
| EP00522 | Sar;Stramenopiles;Gyrista;Ochrophyta;"BD clade";D | EP00522 Amphiprora paludosa 11499 | 1,80E-12 | 820 | XP 002181151.1 | predicted protein [Phaeodactylum tricornutum CCAP 1055/1] | 1133 | 0 | 649,818 | 56,14906832 |
| EP00183 | Archaeplastida;"RR clade";Rhodophyta;Proteorhod | EP00183 Erythrolobus madagascarensis 203 | 1,80E-12 | 192 | ***no hit found*** | | | | | |
| EP00636 | Sar;Stramenopiles;Gyrista;Ochrophyta;Pinguiophyc | EP00636 Phaeomonas parva 6666 | 1,90E-12 | 253 | XP 009515715.1 | hypothetical protein PHYSODRAFT 322106 [Phytophthora sojae] | 1262 | 1,73E-44 | 170,244 | 47,75510204 |
| EP00656 | Telonemia;Telonema;"Tsubtile clade";Telonema sul | EP00656 Telonema subtile 32752 | 2,00E-12 | 115 | ***no hit found*** | | | | | |
| EP00376 | Sar;Alveolata;Myzozoa;"AC clade";Apicomplexa;"cc | EP00376 Eimeria tenella 1817 | 2,00E-12 | 2290 | XP 013229543.1 | hypothetical protein, conserved [Eimeria tenella] | 2290 | 0 | 4504,51 | 100 |
| EP00301 | Haptista;Centroplasthelida;Pterocystida;Raphidista | EP00301 Raphidiophrys heterophryoidea 16108 | 2,10E-12 | 238 | ***no hit found*** | | | | | |
| EP00638 | Sar;Stramenopiles;Gyrista;Ochrophyta;Raphidophy | EP00638 Chattonella subsalsa 17267 | 2,30E-12 | 1170 | XP 009515715.1 | hypothetical protein PHYSODRAFT 322106 [Phytophthora sojae] | 1262 | 0 | 674,47 | 51,22494432 |
| EP00704 | Metamonada;"BF clade";Fornicata;Kipferlia;Kipferli | EP00704 Kipferlia bialata 1703 | 2,40E-12 | 2093 | XP 026291516.1 | obscurin-like isoform X4 [Frankliniella occidentalis] | 6099 | 5,26E-26 | 130,183 | 39,28571429 |
| EP00376 | Sar;Alveolata;Myzozoa;"AC clade";Apicomplexa;"cc | EP00376 Eimeria tenella 8121 | 2,80E-12 | 1116 | XP 013235864.1 | hypothetical protein ETH 00028115 [Eimeria tenella] | 1116 | 0 | 2130,91 | 100 |
| EP00566 | Sar;Stramenopiles;Gyrista;Ochrophyta;"BD clade";D | EP00566 Odontella aurita 1249 | 2,80E-12 | 552 | XP 002290923.1 | predicted protein [Thalassiosira pseudonana CCMP1335] | 1041 | 4,60E-175 | 531,561 | 62,72401434 |
| EP00474 | Sar;Rhizaria;Endomyxa;"phytorhiza";Phytomyxea;"( | EP00474 Spongospora subterranea 5006 | 2,80E-12 | 1630 | ***no hit found*** | | | | | |
| EP00299 | Haptista;Centroplasthelida;Pterocystida;Pterista;"P | EP00299 Pterocystis canadensis 10864 | 2,90E-12 | 924 | XP 020585787.1 | protein transport protein SEC31 homolog B isoform X1 [Phalaenopsis equestris] | 1127 | 2,24E-128 | 424,476 | 44,42988204 |
| EP00412 | Sar;Alveolata;Myzozoa;"DP clade";Dinoflagellata;D | EP00412 Alexandrium andersonii 10197 | 3,00E-12 | 77 | ***no hit found*** | | | | | |
| EP00229 | Archaeplastida;Chloroplastida;Chlorophyta;Mamiel | EP00229 Micromonas commoda 1051 | 3,10E-12 | 1395 | XP 002500425.1 | predicted protein [Micromonas commoda] | 1395 | 0 | 2732,21 | 100 |
| EP00457 | Sar;Rhizaria;Cercozoa;"crown cercozoa";Imbricatea | EP00457 Paulinella chromatophora 689 | 3,10E-12 | 1239 | XP 028397312.1 | protein transport protein Sec31A-like [Dendronephthya gigantea] | 1203 | 2,17E-94 | 340,117 | 43,87678437 |
| EP00687 | Discoba;Heterolobosea;Tetramitia;Eutetramitia;"PS | EP00687 Percolomonas cosmopolitus 4664 | 3,10E-12 | 645 | XP 012386130.1 | protein transport protein Sec31A isoform X17 [Dasypus novemcinctus] | 1066 | 2,72E-33 | 147,902 | 42,14501511 |
| EP00638 | Sar;Stramenopiles;Gyrista;Ochrophyta;Raphidophy | EP00638 Chattonella subsalsa 9075 | 3,10E-12 | 1105 | XP 009515715.1 | hypothetical protein PHYSODRAFT 322106 [Phytophthora sojae] | 1262 | 0 | 595,504 | 52,21843003 |
| EP00667 | Discoba;Euglenozoa;Euglenida;Spirocuta;Euglenea;I | EP00667 Euglena gracilis 1138 | 3,20E-12 | 884 | XP 030046480.1 | protein transport protein Sec31A isoform X6 [Microcaecilia unicolor] | 1139 | 5,43E-115 | 387,882 | 47,05882353 |
| EP00244 | Archaeplastida;Chloroplastida;Streptophyta;"core s | EP00244 Chara vulgaris 8504 | 3,40E-12 | 117 | XP 011070128.1 | protein transport protein SEC16B homolog [Sesamum indicum] | 1458 | 1,61E-22 | 101,293 | 53,33333333 |
| EP00222 | Archaeplastida;Chloroplastida;Chlorophyta;Mamiel | EP00222 Ostreococcus lucimarinus 3026 | 3,40E-12 | 592 | XP 001418383.1 | predicted protein [Ostreococcus lucimarinus CCE9901] | 592 | 0 | 1187,94 | 100 |
| EP00082 | Obazoa;Opisthokonta;Holozoa;Choanozoa;Metazoa | EP00082 Ixodes scapularis 3454 | 3,40E-12 | 347 | XP 029846214.1 | uncharacterized protein LOC8036454 isoform X1 [Ixodes scapularis] | 2152 | 0 | 700,664 | 99,71181556 |
| EP00200 | Archaeplastida;Chloroplastida;Chlorophyta;"core ch | EP00200 Dunaliella tertiolecta 4903 | 3,40E-12 | 1044 | XP 001689665.1 | hypothetical protein CHLREDRAFT 146684 [Chlamydomonas reinhardtii] | 1145 | 1,16E-30 | 142,51 | 37,16381418 |
| EP00242 | Archaeplastida;Chloroplastida;Chlorophyta;Pyramir | EP00242 Pyramimonas sp CCMP2087 12936 | 3,60E-12 | 578 | XP 020225819.1 | protein transport protein SEC16B homolog [Cajanus cajan] | 1326 | 1,39E-14 | 88,9669 | 53,125 |
| EP00120 | Obazoa;Opisthokonta;Holozoa;Filasterea;Capsaspoi | EP00120 Capsaspora owczarzaki 5841 | 3,70E-12 | 1962 | XP 004363971.1 | hypothetical protein CAOG 03132 [Capsaspora owczarzaki ATCC 30864] | 1962 | 0 | 3889,73 | 100 |
| EP00044 | Obazoa;Opisthokonta;Holozoa;Choanozoa;Choanof | EP00044 Salpingoeca rosetta 6364 | 3,70E-12 | 1098 | XP 004993797.1 | hypothetical protein PTSG 12276 [Salpingoeca rosetta] | 1098 | 0 | 2224,51 | 100 |
| EP00241 | Archaeplastida;Chloroplastida;Chlorophyta;Pyramir | EP00241 Pyramimonas parkeae 18086 | 3,80E-12 | 357 | XP 002987428.1 | protein transport protein SEC16B homolog [Selaginella moellendorffii] | 1190 | 4,04E-09 | 69,3218 | 45,91836735 |
| EP00376 | Sar;Alveolata;Myzozoa;"AC clade";Apicomplexa;"cc | EP00376 Eimeria tenella 4300 | 3,90E-12 | 2087 | XP 013232033.1 | hypothetical protein, conserved [Eimeria tenella] | 2087 | 0 | 4122,01 | 100 |
| EP00638 | Sar;Stramenopiles;Gyrista;Ochrophyta;Raphidophy | EP00638 Chattonella subsalsa 9040 | 3,90E-12 | 1150 | XP 009515715.1 | hypothetical protein PHYSODRAFT 322106 [Phytophthora sojae] | 1262 | 0 | 675,241 | 49,82935154 |
| EP00013 | Amoebozoa;Discosea;Flabellinia;Stygamoeba;Styga | EP00013 Stygamoeba regulata 29253 | 3,90E-12 | 1051 | XP 004989501.1 | hypothetical protein PTSG 09244 [Salpingoeca rosetta] | 1883 | 6,75E-21 | 111,309 | 100 |
| EP00022 | Amoebozoa;Evosea;Eumycetozoa;Dictyostelia;Acyto | EP00022 Cavenderia fasciculata 11948 | 4,80E-12 | 1807 | XP 004367225.1 | putative mediator complex subunit 12 [Cavenderia fasciculata] | 1807 | 0 | 3655,92 | 100 |
| EP00550 | Sar;Stramenopiles;Gyrista;Ochrophyta;"BD clade";D | EP00550 Attheya septentrionalis 11877 | 4,80E-12 | 69 | XP 002183614.1 | predicted protein [Phaeodactylum tricornutum CCAP 1055/1] | 2083 | 1,90E-09 | 61,6178 | 40,32258065 |
| EP00562 | Sar;Stramenopiles;Gyrista;Ochrophyta;"BD clade";D | EP00562 Extubocellulus spinifer 32585 | 5,20E-12 | 931 | XP 002290923.1 | predicted protein [Thalassiosira pseudonana CCMP1335] | 1041 | 0 | 892,493 | 60,71044133 |
| EP00754 | Discoba;Euglenozoa;Diplonemea;Diplonemidae;Rhy | EP00754 Rhynchopus humris 38061 | 5,30E-12 | 424 | ***no hit found*** | | | | | |
| EP00206 | Archaeplastida;Chloroplastida;Chlorophyta;"core ch | EP00206 Coccomyxa subellipsoidea 846 | 5,50E-12 | 1272 | XP 005643395.1 | hypothetical protein COCSUDRAFT 49081 [Coccomyxa subellipsoidea C-169] | 1272 | 0 | 2584,67 | 100 |
| EP00201 | Archaeplastida;Chloroplastida;Chlorophyta;"core ch | EP00201 Polytomella parva 20681 | 6,00E-12 | 1740 | ***no hit found*** | | | | | |
| EP00120 | Obazoa;Opisthokonta;Holozoa;Filasterea;Capsaspoi | EP00120 Capsaspora owczarzaki 8057 | 6,00E-12 | 1951 | XP 011270238.1 | hypothetical protein, variant [Capsaspora owczarzaki ATCC 30864] | 1951 | 0 | 3867,77 | 100 |
| EP00236 | Archaeplastida;Chloroplastida;Chlorophyta;Picocyst | EP00236 Picocystis salinarum 4318 | 6,10E-12 | 606 | XP 018455948.1 | PREDICTED: protein transport protein SEC16A homolog isoform X3 [Raphanus sativus] | 1363 | 1,33E-06 | 63,5438 | 50,96774194 |
| EP00316 | Haptista;Haptophyta;Prymnesiophyceae;Calcihapto | EP00316 Scyphosphaera apsteinii 8887 | 6,20E-12 | 464 | ***no hit found*** | | | | | |
| EP00632 | Sar;Stramenopiles;Gyrista;Ochrophyta;Pelagophyce | EP00632 Arachnochrysis sp CCMP2950 12220 | 6,20E-12 | 1006 | ***no hit found*** | | | | | |
| EP00562 | Sar;Stramenopiles;Gyrista;Ochrophyta;"BD clade";D | EP00562 Extubocellulus spinifer 1573 | 6,30E-12 | 995 | XP 002290923.1 | predicted protein [Thalassiosira pseudonana CCMP1335] | 1041 | 0 | 929,858 | 59,11764706 |
| EP00548 | Sar;Stramenopiles;Gyrista;Ochrophyta;"BD clade";D | EP00548 Thalassiothrix antarctica 20837 | 6,80E-12 | 1017 | XP 002181151.1 | predicted protein [Phaeodactylum tricornutum CCAP 1055/1] | 1133 | 0 | 816,224 | 55,31167691 |
| EP00317 | Haptista;Haptophyta;Prymnesiophyceae;Phaeocyst | EP00317 Phaeocystis antarctica 47446 | 6,80E-12 | 869 | XP 005788197.1 | secretory protein 31 [Emiliania huxleyi CCMP1516] | 1166 | 0 | 848,581 | 62,06896552 |
| EP00200 | Archaeplastida;Chloroplastida;Chlorophyta;"core ch | EP00200 Dunaliella tertiolecta 14603 | 7,50E-12 | 1301 | XP 002954917.1 | hypothetical protein VOLCADRAFT 118887 [Volvox carteri f. nagariensis] | 4212 | 2,66E-24 | 123,25 | 54,03225806 |
| EP00466 | Sar;Rhizaria;Cercozoa;Chlorarachnea;"core chlorara | EP00466 Bigelowiella natans 19194 | 7,60E-12 | 1116 | XP 021688424.1 | protein transport protein SEC31 homolog B-like [Hevea brasiliensis] | 1130 | 2,39E-106 | 370,163 | 42,82786885 |
| EP00562 | Sar;Stramenopiles;Gyrista;Ochrophyta;"BD clade";D | EP00562 Extubocellulus spinifer 14407 | 7,80E-12 | 1082 | XP 002290923.1 | predicted protein [Thalassiosira pseudonana CCMP1335] | 1041 | 0 | 1057,74 | 58,99543379 |
| EP00374 | Sar;Alveolata;Myzozoa;"AC clade";Apicomplexa;"cc | EP00374 Theileria orientalis 2513 | 8,40E-12 | 211 | XP 009691345.1 | conserved hypothetical protein [Theileria orientalis strain Shintoku] | 211 | 6,33E-154 | 434,106 | 100 |
| EP00514 | Sar;Stramenopiles;Bigyra;Sagenista;Labyrinthulomy | EP00514 Thraustochytrium sp LLF1b 5282 | 8,50E-12 | 1312 | XP 021624622.1 | protein transport protein SEC31 homolog B-like [Manihot esculenta] | 1130 | 3,54E-106 | 373,244 | 45,37815126 |
| EP00568 | Sar;Stramenopiles;Gyrista;Ochrophyta;"BD clade";D | EP00568 Trieres chinensis 11945 | 8,60E-12 | 1016 | XP 002290923.1 | predicted protein [Thalassiosira pseudonana CCMP1335] | 1041 | 8,33E-163 | 516,153 | 54,94699647 |
| EP00390 | Sar;Alveolata;Myzozoa;"AC clade";Colpodellida;Vitr | EP00390 Vitrella brassicaformis 2135 | 8,80E-12 | 1182 | XP 002140912.1 | hypothetical protein [Cryptosporidium muris RN66] | 1225 | 0 | 728,783 | 51,23266564 |
| EP00544 | Sar;Stramenopiles;Gyrista;Ochrophyta;"BD clade";D | EP00544 Staurosira sp CCMP2646 4285 | 9,90E-12 | 1036 | XP 002290923.1 | predicted protein [Thalassiosira pseudonana CCMP1335] | 1041 | 0 | 1029,24 | 57,63033175 |
| EP00514 | Sar;Stramenopiles;Bigyra;Sagenista;Labyrinthulomy | EP00514 Thraustochytrium sp LLF1b 11695 | 1,00E-11 | 1334 | XP 021624622.1 | protein transport protein SEC31 homolog B-like [Manihot esculenta] | 1130 | 5,76E-107 | 375,941 | 46,4312547 |
| EP00010 | Amoebozoa;Discosea;Flabellinia;"core flabellinids"; | EP00010 Vexillifera sp DIVA3-564-2 8190 | 1,10E-11 | 259 | ***no hit found*** | | | | | |
| EP00100 | Obazoa;Opisthokonta;Holozoa;Choanozoa;Metazoa | EP00100 Culex quinquefasciatus 8062 | 1,20E-11 | 225 | XP 001850120.1 | conserved hypothetical protein [Culex quinquefasciatus] | 225 | 9,14E-166 | 465,307 | 100 |
| EP00215 | Archaeplastida;Chloroplastida;Chlorophyta;Chlorop | EP00215 Chloropicon roscoffensis 6332 | 1,20E-11 | 627 | XP 020225819.1 | protein transport protein SEC16B homolog [Cajanus cajan] | 1326 | 7,94E-08 | 67,3958 | 47,22222222 |
| EP00751 | Sar;Stramenopiles;Gyrista;Ochrophyta;"BD clade";D | EP00751 Fragilariopsis kerguelensis 43370 | 1,20E-11 | 980 | XP 002290400.1 | ankyrin domain-containing protein, partial [Thalassiosira pseudonana CCMP1335] | 139 | 1,55E-24 | 110,538 | 47,44525547 |
| EP00066 | Obazoa;Opisthokonta;Holozoa;Choanozoa;Metazoa | EP00066 Takifugu rubripes 10880 | 1,30E-11 | 478 | XP 029693435.1 | protein transport protein Sec31A-like isoform X3 [Takifugu rubripes] | 1192 | 0 | 993,416 | 100 |
| EP00628 | Sar;Stramenopiles;Gyrista;Ochrophyta;Pelagophyce | EP00628 Pelagophyceae sp CCMP2097 35359 | 1,40E-11 | 429 | XP 002908536.1 | COPII coat assembly protein SEC16, putative [Phytophthora infestans T30-4] | 2641 | 5,03E-06 | 60,8474 | 47,59615385 |
| EP00235 | Archaeplastida;Chloroplastida;Palmophyllophyceae | EP00235 Prasinoderma singulare 229 | 1,40E-11 | 1129 | XP 020519257.1 | protein transport protein SEC31 homolog B isoform X2 [Amborella trichopoda] | 1154 | 0 | 716,072 | 49,95836803 |
| EP00376 | Sar;Alveolata;Myzozoa;"AC clade";Apicomplexa;"cc | EP00376 Eimeria tenella 6074 | 1,50E-11 | 952 | XP 013233807.1 | hypothetical protein, conserved [Eimeria tenella] | 952 | 0 | 1827,37 | 100 |
| EP00588 | Sar;Stramenopiles;Gyrista;Ochrophyta;"BD clade";D | EP00588 Corethron pennatum 8634 | 1,60E-11 | 1069 | XP 002290923.1 | predicted protein [Thalassiosira pseudonana CCMP1335] | 1041 | 0 | 682,174 | 53,45060893 |
| EP00059 | Obazoa;Opisthokonta;Holozoa;Choanozoa;Metazoa | EP00059 Ciona intestinalis 21002 | 1,60E-11 | 1163 | XP 026696656.1 | protein transport protein Sec31A isoform X7 [Ciona intestinalis] | 1163 | 0 | 2422,51 | 100 |
| EP00239 | Archaeplastida;Chloroplastida;Chlorophyta;Pyramir | EP00239 Pterosperma sp CCMP1384 2158 | 1,70E-11 | 1127 | XP 024379150.1 | protein transport protein SEC31 homolog B-like isoform X1 [Physcomitrella patens] | 1179 | 0 | 858,981 | 53,08641975 |
| EP00191 | Archaeplastida;Chloroplastida;Chlorophyta;"core ch | EP00191 Tetraselmis sp GSL018 6438 | 1,80E-11 | 1799 | ***no hit found*** | | | | | |
| EP00059 | Obazoa;Opisthokonta;Holozoa;Choanozoa;Metazoa | EP00059 Ciona intestinalis 21001 | 1,80E-11 | 1188 | XP 026696655.1 | protein transport protein Sec31A isoform X6 [Ciona intestinalis] | 1188 | 0 | 2475,66 | 100 |
| EP00059 | Obazoa;Opisthokonta;Holozoa;Choanozoa;Metazoa | EP00059 Ciona intestinalis 21000 | 1,80E-11 | 1190 | XP 026696654.1 | protein transport protein Sec31A isoform X5 [Ciona intestinalis] | 1190 | 0 | 2477,59 | 100 |
| EP00110 | Obazoa;Opisthokonta;Holozoa;Choanozoa;Metazoa | EP00110 Nematostella vectensis 17336 | 1,80E-11 | 1219 | XP 001634737.1 | predicted protein [Nematostella vectensis] | 1219 | 0 | 2526,12 | 100 |
| EP00059 | Obazoa;Opisthokonta;Holozoa;Choanozoa;Metazoa | EP00059 Ciona intestinalis 20997 | 1,80E-11 | 1213 | XP 026696651.1 | protein transport protein Sec31A isoform X2 [Ciona intestinalis] | 1213 | 0 | 2525,74 | 100 |
| EP00059 | Obazoa;Opisthokonta;Holozoa;Choanozoa;Metazoa | EP00059 Ciona intestinalis 20996 | 1,80E-11 | 1215 | XP 026696650.1 | protein transport protein Sec31A isoform X1 [Ciona intestinalis] | 1215 | 0 | 2529,98 | 100 |
| EP00376 | Sar;Alveolata;Myzozoa;"AC clade";Apicomplexa;"cc | EP00376 Eimeria tenella 8426 | 1,90E-11 | 1075 | XP 013236170.1 | hypothetical protein, conserved, partial [Eimeria tenella] | 1075 | 0 | 2090,08 | 100 |
| EP00405 | Sar;Alveolata;Myzozoa;"DP clade";Dinoflagellata;D | EP00405 Crypthecodinium cohnii 40919 | 1,90E-11 | 927 | ***no hit found*** | | | | | |
| EP00761 | Discoba;Heterolobosea;Pharyngomonas;Pharyngon | EP00761 Pharyngomonas kirbyi 9669 | 2,00E-11 | 266 | ***no hit found*** | | | | | |
| EP00249 | Archaeplastida;Chloroplastida;Streptophyta;"core s | EP00249 Psilotum nudum 31423 | 2,00E-11 | 93 | XP 024373338.1 | protein transport protein SEC16A homolog isoform X1 [Physcomitrella patens] >gi\|13738! | 1444 | 2,11E-21 | 97,0561 | 60,63829787 |
| EP00592 | Sar;Stramenopiles;Gyrista;Ochrophyta;"BD clade";D | EP00592 Proboscia alata 14944 | 2,10E-11 | 424 | XP 002183614.1 | predicted protein [Phaeodactylum tricornutum CCAP 1055/1] | 2083 | 2,17E-18 | 98,9821 | 60,94674556 |
| EP00376 | Sar;Alveolata;Myzozoa;"AC clade";Apicomplexa;"cc | EP00376 Eimeria tenella 320 | 2,20E-11 | 996 | XP 013228046.1 | hypothetical protein, conserved [Eimeria tenella] | 996 | 0 | 1949,48 | 100 |
| EP00089 | Obazoa;Opisthokonta;Holozoa;Choanozoa;Metazoa | EP00089 Calanus finmarchicus 23840 | 2,20E-11 | 522 | XP 023336929.1 | protein transport protein Sec31A-like [Eurytemora affinis] | 1195 | 0 | 593,578 | 64,61538462 |
| EP00104 | Obazoa;Opisthokonta;Holozoa;Choanozoa;Metazoa | EP00104 Lingula anatina 5383 | 2,30E-11 | 1860 | XP 013384747.1 | protein transport protein Sec31A-like [Lingula anatina] | 1860 | 0 | 3853,52 | 100 |
| EP00740 | Archaeplastida;Chloroplastida;Chlorophyta;Mamiel | EP00740 Mantoniella antarctica 6861 | 2,40E-11 | 203 | XP 002509249.1 | predicted protein [Micromonas commoda] | 1945 | 2,71E-18 | 92,8189 | 66,12903226 |
| EP00298 | Haptista;Centroplasthelida;Panacanthocystida;Acar | EP00298 Acanthocystis sp HF-20 23915 | 2,40E-11 | 162 | ***no hit found*** | | | | | |
| EP00387 | Sar;Alveolata;Myzozoa;"AC clade";Colpodellida;Chr | EP00387 Chromera velia 2723 | 2,40E-11 | 364 | ***no hit found*** | | | | | |
| EP00451 | Sar;Alveolata;Myzozoa;"DP clade";Dinoflagellata;O | EP00451 Oxyrrhis marina 27106 | 2,50E-11 | 218 | ***no hit found*** | | | | | |
| EP00520 | Sar;Stramenopiles;Gyrista;Ochrophyta;"BD clade";E | EP00520 Triparma pacifica 7477 | 2,50E-11 | 235 | XP 002290923.1 | predicted protein [Thalassiosira pseudonana CCMP1335] | 1041 | 3,07E-71 | 244,973 | 56,55737705 |
| EP00112 | Obazoa;Opisthokonta;Holozoa;Choanozoa;Metazoa | EP00112 Aurelia sp Birch-Aquarium-sp1 11275 | 2,50E-11 | 1162 | XP 028397312.1 | protein transport protein Sec31A-like [Dendronephthya gigantea] | 1203 | 0 | 1103,58 | 58,07754443 |
| EP00755 | Discoba;Euglenozoa;Diplonemea;Diplonemidae;Sul | EP00755 Sulcionema specki 21184 | 2,60E-11 | 755 | XP 014545106.1 | Protein transport protein sec31, partial [Metarhizium brunneum ARSEF 3297] | 1248 | 1,13E-57 | 224,172 | 44,70246734 |
| EP00232 | Archaeplastida;Chloroplastida;Chlorophyta;Nephro | EP00232 Nephroselmis pyriformis 2097 | 2,80E-11 | 106 | XP 018457010.1 | PREDICTED: protein transport protein SEC16A homolog isoform X3 [Raphanus sativus] | 1080 | 6,06E-10 | 65,0846 | 47,56097561 |
| EP00089 | Obazoa;Opisthokonta;Holozoa;Choanozoa;Metazoa | EP00089 Calanus finmarchicus 23823 | 2,90E-11 | 781 | XP 023336929.1 | protein transport protein Sec31A-like [Eurytemora affinis] | 1195 | 0 | 951,429 | 66,20428752 |
| EP00655 | Sar;Stramenopiles;Gyrista;Peronosporomycetes;"cc | EP00655 Saprolegnia parasitica 12465 | 3,20E-11 | 1190 | XP 012205971.1 | hypothetical protein SPRG 11633 [Saprolegnia parasitica CBS 223.65] | 1190 | 0 | 2427,51 | 100 |
| EP00443 | Sar;Alveolata;Myzozoa;"DP clade";Dinoflagellata;D | EP00443 Scrippsiella trochoidea 89701 | 3,20E-11 | 1111 | ***no hit found*** | | | | | |
| EP00635 | Sar;Stramenopiles;Gyrista;Ochrophyta;Pelagophyce | EP00635 Sarcinochrysidales sp CCMP3193 4767 | 3,30E-11 | 875 | XP 009535381.1 | hypothetical protein PHYSODRAFT 525781 [Phytophthora sojae] | 716 | 2,11E-16 | 95,5153 | 50 |
| EP00269 | Archaeplastida;Chloroplastida;Streptophyta;"core s | EP00269 Selaginella moellendorffii 35424 | 3,30E-11 | 377 | XP 024536070.1 | protein transport protein SEC16A homolog [Selaginella moellendorffii] >gi\|1376936154\|r | 377 | 0 | 756,133 | 100 |
| EP00269 | Archaeplastida;Chloroplastida;Streptophyta;"core s | EP00269 Selaginella moellendorffii 35425 | 3,30E-11 | 377 | XP 024536070.1 | protein transport protein SEC16A homolog [Selaginella moellendorffii] >gi\|1376936154\|r | 377 | 0 | 756,133 | 100 |
| EP00269 | Archaeplastida;Chloroplastida;Streptophyta;"core s | EP00269 Selaginella moellendorffii 35426 | 3,30E-11 | 377 | XP 024536070.1 | protein transport protein SEC16A homolog [Selaginella moellendorffii] >gi\|1376936154\|r | 377 | 0 | 756,133 | 100 |
| EP00269 | Archaeplastida;Chloroplastida;Streptophyta;"core s | EP00269 Selaginella moellendorffii 35427 | 3,30E-11 | 377 | XP 024536070.1 | protein transport protein SEC16A homolog [Selaginella moellendorffii] >gi\|1376936154\|r | 377 | 0 | 756,133 | 100 |
| EP00020 | Amoebozoa;Evosea;Cutosea;Sapocribrum;Sapocribr | EP00020 Sapocribrum chincoteaguense 1821 | 3,70E-11 | 1041 | XP 012754771.1 | hypothetical protein SAMD00019534 058260 [Acytostelium subglobosum LB1] | 1179 | 9,85E-134 | 443,736 | 43,26375712 |
| EP00020 | Amoebozoa;Evosea;Cutosea;Sapocribrum;Sapocribr | EP00020 Sapocribrum chincoteaguense 14133 | 3,90E-11 | 1000 | XP 012754771.1 | hypothetical protein SAMD00019534 058260 [Acytostelium subglobosum LB1] | 1179 | 4,12E-132 | 437,958 | 44,26540284 |
| EP00191 | Archaeplastida;Chloroplastida;Chlorophyta;"core ch | EP00191 Tetraselmis sp GSL018 9715 | 3,90E-11 | 1795 | ***no hit found*** | | | | | |
| EP00236 | Archaeplastida;Chloroplastida;Chlorophyta;Picocyst | EP00236 Picocystis salinarum 474 | 3,90E-11 | 1401 | XP 006837403.2 | protein transport protein SEC31 homolog B isoform X1 [Amborella trichopoda] | 1155 | 0 | 810,446 | 51,91163976 |
| EP00043 | Obazoa;Opisthokonta;Holozoa;Choanozoa;Choanof | EP00043 Microstomoeca roanoka 16639 | 4,20E-11 | 841 | XP 004992030.1 | hypothetical protein PTSG 12463 [Salpingoeca rosetta] | 2149 | 1,52E-110 | 383,645 | 58,86287625 |
| EP00089 | Obazoa;Opisthokonta;Holozoa;Choanozoa;Metazoa | EP00089 Calanus finmarchicus 23843 | 4,30E-11 | 502 | XP 023336929.1 | protein transport protein Sec31A-like [Eurytemora affinis] | 1195 | 0 | 586,26 | 65,0390625 |
| EP00668 | Discoba;Euglenozoa;Euglenida;Spirocuta;Euglenea;I | EP00668 Euglena longa 6628 | 4,30E-11 | 1096 | XP 009391540.1 | PREDICTED: protein transport protein SEC31 homolog B-like [Musa acuminata subsp. mala | 1123 | 2,19E-151 | 490,73 | 46,55319149 |
| EP00241 | Archaeplastida;Chloroplastida;Chlorophyta;Pyramir | EP00241 Pyramimonas parkeae 12586 | 4,40E-11 | 268 | XP 002975264.1 | protein transport protein SEC16B homolog [Selaginella moellendorffii] | 1190 | 8,48E-08 | 63,929 | 46,11398964 |
| EP00002 | CRuMs;Collodictyonidae;Diphylleia;Diphylleia rotan | EP00002 Diphylleia rotans 14283 | 4,50E-11 | 1070 | XP 005831033.1 | secretory protein Sec31 [Guillardia theta CCMP2712] | 1081 | 0 | 586,26 | 46,48711944 |
| EP00089 | Obazoa;Opisthokonta;Holozoa;Choanozoa;Metazoa | EP00089 Calanus finmarchicus 23824 | 4,50E-11 | 761 | XP 023336929.1 | protein transport protein Sec31A-like [Eurytemora affinis] | 1195 | 0 | 936,021 | 66,36481242 |
| EP00100 | Obazoa;Opisthokonta;Holozoa;Choanozoa;Metazoa | EP00100 Culex quinquefasciatus 8061 | 4,80E-11 | 315 | XP 001850119.1 | conserved hypothetical protein [Culex quinquefasciatus] | 315 | 0 | 612,838 | 100 |
| EP00279 | Cryptista;Cryptophyceae;Cryptomonadales;"lineage | EP00279 Guillardia theta 13314 | 4,80E-11 | 1081 | XP 005831033.1 | secretory protein Sec31 [Guillardia theta CCMP2712] | 1081 | 0 | 2230,68 | 100 |
| EP00002 | CRuMs;Collodictyonidae;Diphylleia;Diphylleia rotan | EP00002 Diphylleia rotans 14282 | 5,10E-11 | 1138 | XP 005831033.1 | secretory protein Sec31 [Guillardia theta CCMP2712] | 1081 | 0 | 587,415 | 46,48711944 |
| EP00002 | CRuMs;Collodictyonidae;Diphylleia;Diphylleia rotan | EP00002 Diphylleia rotans 14284 | 5,10E-11 | 1138 | XP 005831033.1 | secretory protein Sec31 [Guillardia theta CCMP2712] | 1081 | 0 | 587,415 | 46,48711944 |
| EP00089 | Obazoa;Opisthokonta;Holozoa;Choanozoa;Metazoa | EP00089 Calanus finmarchicus 23835 | 5,30E-11 | 713 | XP 023336929.1 | protein transport protein Sec31A-like [Eurytemora affinis] | 1195 | 0 | 898,656 | 68,3029453 |
| EP00131 | Obazoa;Opisthokonta;Nucletmycea;Fungi;"core Fur | EP00131 Spizellomyces punctatus 8981 | 5,60E-11 | 1378 | XP 016612805.1 | hypothetical protein SPPG 00469 [Spizellomyces punctatus DAOM BR117] | 1378 | 0 | 2814,64 | 100 |
| EP00567 | Sar;Stramenopiles;Gyrista;Ochrophyta;"BD clade";D | EP00567 Triceratium dubium 1392 | 5,70E-11 | 126 | XP 002183614.1 | predicted protein [Phaeodactylum tricornutum CCAP 1055/1] | 2083 | 9,58E-14 | 76,6406 | 51,85185185 |
| EP00058 | Obazoa;Opisthokonta;Holozoa;Choanozoa;Metazoa | EP00058 Branchiostoma floridae 17442 | 5,70E-11 | 1113 | XP 002602930.1 | hypothetical protein BRAFLDRAFT 62485 [Branchiostoma floridae] | 1113 | 0 | 2301,17 | 100 |
| EP00115 | Obazoa;Opisthokonta;Holozoa;Choanozoa;Metazoa | EP00115 Mnemiopsis leidyi 12871 | 5,80E-11 | 1554 | XP 002740116.2 | PREDICTED: centrosomal protein KIAA1731 homolog [Saccoglossus kowalevskii] | 1757 | 9,07E-17 | 98,5969 | 46,83544304 |
| EP00197 | Archaeplastida;Chloroplastida;Chlorophyta;"core ch | EP00197 Chlamydomonas leiostraca 6878 | 5,80E-11 | 1267 | XP 002956039.1 | hypothetical protein VOLCADRAFT 107038 [Volvox carteri f. nagariensis] | 1711 | 5,19E-16 | 95,9005 | 46,58119658 |
| EP00037 | Obazoa;Opisthokonta;Holozoa;Choanozoa;Choanof | EP00037 Helgoeca nana 30341 | 6,80E-11 | 1618 | ***no hit found*** | | | | | |
| EP00078 | Obazoa;Opisthokonta;Holozoa;Choanozoa;Metazoa | EP00078 Anolis carolinensis 29154 | 6,90E-11 | 1224 | XP 016849264.1 | PREDICTED: LOW QUALITY PROTEIN: protein transport protein Sec31A [Anolis carolinensis | 1224 | 0 | 2519,19 | 100 |
| EP00033 | Obazoa;Breviatea;Pygsuia;Pygsuia biforma;strain P( | EP00033 Pygsuia biforma 1563 | 7,10E-11 | 963 | XP 020427011.1 | WD40 repeat-containing protein [Heterostelium album PN500] | 836 | 8,60E-135 | 434,876 | 43,62573099 |
| EP00184 | Archaeplastida;"RR clade";Rhodophyta;Proteorhod | EP00184 Porphyridium aerugineum 2778 | 7,30E-11 | 1325 | XP 016449557.1 | PREDICTED: LOW QUALITY PROTEIN: protein transport protein SEC31 homolog B-like [Nic | 1130 | 1,58E-123 | 421,394 | 48,41149773 |
| EP00301 | Haptista;Centroplasthelida;Pterocystida;Raphidista | EP00301 Raphidiophrys heterophryoidea 26410 | 7,40E-11 | 1024 | XP 004355165.1 | WD40 repeat-containing protein [Cavenderia fasciculata] | 1147 | 7,00E-148 | 479,945 | 48,11320755 |
| EP00372 | Sar;Alveolata;Myzozoa;"AC clade";Apicomplexa;"cc | EP00372 Theileria equi 4179 | 8,40E-11 | 184 | XP 004832751.1 | conserved hypothetical protein [Theileria equi strain WA] | 184 | 1,49E-130 | 372,859 | 100 |
| EP00385 | Sar;Alveolata;Myzozoa;"AC clade";Apicomplexa;Cry | EP00385 Cryptosporidium parvum 3686 | 8,50E-11 | 610 | XP 628577.1 | hypothetical protein [Cryptosporidium parvum Iowa II] | 610 | 0 | 1250,34 | 100 |
| EP00376 | Sar;Alveolata;Myzozoa;"AC clade";Apicomplexa;"cc | EP00376 Eimeria tenella 4254 | 8,60E-11 | 1103 | XP 013231987.1 | hypothetical protein, conserved [Eimeria tenella] | 1103 | 0 | 2133,99 | 100 |
| EP00520 | Sar;Stramenopiles;Gyrista;Ochrophyta;"BD clade";E | EP00520 Triparma pacifica 9358 | 8,80E-11 | 721 | XP 002290923.1 | predicted protein [Thalassiosira pseudonana CCMP1335] | 1041 | 5,39E-175 | 537,724 | 47,55244755 |
| EP00206 | Archaeplastida;Chloroplastida;Chlorophyta;"core ch | EP00206 Coccomyxa subellipsoidea 1011 | 8,80E-11 | 1799 | XP 005643560.1 | hypothetical protein COCSUDRAFT 48975 [Coccomyxa subellipsoidea C-169] | 1799 | 0 | 3538,43 | 100 |
| EP00189 | Archaeplastida;"RR clade";Rhodophyta;Proteorhod | EP00189 Rhodosorus marinus 8750 | 9,00E-11 | 1884 | XP 025602404.1 | mucin-16-like [Athalia rosae] | 811 | 4,13E-30 | 141,354 | 77,09251101 |
| EP00601 | Sar;Stramenopiles;Gyrista;Ochrophyta;"CS clade";C | EP00601 Ochromonadales sp CCMP2298 2047 | 9,30E-11 | 1091 | XP 002906652.1 | protein transporter Sec31A, putative [Phytophthora infestans T30-4] | 1264 | 0 | 654,825 | 50,88967972 |
| EP00427 | Sar;Alveolata;Myzozoa;"DP clade";Dinoflagellata;D | EP00427 Karlodinium veneficum 29353 | 1,00E-10 | 683 | ***no hit found*** | | | | | |
| EP00089 | Obazoa;Opisthokonta;Holozoa;Choanozoa;Metazoa | EP00089 Calanus finmarchicus 23836 | 1,10E-10 | 693 | XP 023336929.1 | protein transport protein Sec31A-like [Eurytemora affinis] | 1195 | 0 | 890,952 | 68,65248227 |
| EP00376 | Sar;Alveolata;Myzozoa;"AC clade";Apicomplexa;"cc | EP00376 Eimeria tenella 5980 | 1,10E-10 | 1253 | XP 013233713.1 | diacylglycerol kinase, putative [Eimeria tenella] | 1253 | 0 | 2558,87 | 100 |
| EP00020 | Amoebozoa;Evosea;Cutosea;Sapocribrum;Sapocribr | EP00020 Sapocribrum chincoteaguense 9081 | 1,10E-10 | 414 | WP 148340744.1 | DUF945 family protein [Aquicella siphonis] | 689 | 1,41E-27 | 125,946 | 59,16666667 |
| EP00550 | Sar;Stramenopiles;Gyrista;Ochrophyta;"BD clade";D | EP00550 Attheya septentrionalis 8873 | 1,20E-10 | 213 | XP 002183614.1 | predicted protein [Phaeodactylum tricornutum CCAP 1055/1] | 2083 | 1,66E-10 | 70,4774 | 42,59259259 |
| EP00686 | Discoba;Heterolobosea;Tetramitia;Eutetramitia;"M | EP00686 Naegleria gruberi 5775 | 1,20E-10 | 1934 | XP 002673704.1 | predicted protein [Naegleria gruberi] | 1934 | 0 | 4000,67 | 100 |
| EP00347 | Sar;Alveolata;Ciliophora;Intramacronucleata;"SAL"; | EP00347 Sterkiella histriomuscorum 1717 | 1,30E-10 | 1288 | XP 002773527.1 | conserved hypothetical protein [Perkinsus marinus ATCC 50983] | 1211 | 1,28E-93 | 338,576 | 43,05396096 |
| EP00589 | Sar;Stramenopiles;Gyrista;Ochrophyta;"BD clade";D | EP00589 Leptocylindrus danicus 11724 | 1,30E-10 | 1100 | XP 002290923.1 | predicted protein [Thalassiosira pseudonana CCMP1335] | 1041 | 0 | 825,854 | 57,83410138 |
| EP00443 | Sar;Alveolata;Myzozoa;"DP clade";Dinoflagellata;D | EP00443 Scrippsiella trochoidea 20878 | 1,40E-10 | 3575 | XP 013897136.1 | hypothetical protein MNEG 9845 [Monoraphidium neglectum] | 406 | 3,34E-19 | 104,375 | 37,95620438 |
| EP00438 | Sar;Alveolata;Myzozoa;"DP clade";Dinoflagellata;D | EP00438 Fugacium kawagutii 36520 | 1,40E-10 | 623 | XP 013756677.1 | hypothetical protein AMSG 12049 [Thecamonas trahens ATCC 50062] | 1953 | 1,14E-42 | 176,792 | 55,96707819 |

| | | | | | | | | | | |
|---|---|---|---|---|---|---|---|---|---|---|
| EP00216 | Archaeplastida;Chloroplastida;Chlorophyta;Chlorop | EP00216 Chloropicon sp CCMP2111 3717 | 1,50E-10 | 1165 | XP 020519257.1 | protein transport protein SEC31 homolog B isoform X2 [Amborella trichopoda] | 1154 | 0 | 736,873 | 49,54507858 |
| EP00302 | Haptista;Haptophyta;Pavlovales;Diacronema;strain | EP00302 Diacronema sp CCMP2436 33560 | 1,50E-10 | 1253 | XP 005788197.1 | secretory protein 31 [Emiliania huxleyi CCMP1516] | 1166 | 0 | 619,772 | 52,70092227 |
| EP00376 | Sar;Alveolata;Myzozoa;"AC clade";Apicomplexa;"cc | EP00376 Eimeria tenella 3448 | 1,50E-10 | 1371 | XP 013231179.1 | hypothetical protein, conserved [Eimeria tenella] | 1371 | 0 | 2687,14 | 100 |
| EP00067 | Obazoa;Opisthokonta;Holozoa;Choanozoa;Metazoa | EP00067 Danio rerio 16426 | 1,80E-10 | 1171 | XP 001921581.3 | protein transport protein Sec31A isoform X3 [Danio rerio] | 1171 | 0 | 2401,71 | 100 |
| EP00067 | Obazoa;Opisthokonta;Holozoa;Choanozoa;Metazoa | EP00067 Danio rerio 24911 | 1,80E-10 | 1174 | XP 005173060.1 | protein transport protein Sec31A isoform X1 [Danio rerio] >gi\|528496920\|ref\|XP 00517 | 1174 | 0 | 2407,1 | 100 |
| EP00067 | Obazoa;Opisthokonta;Holozoa;Choanozoa;Metazoa | EP00067 Danio rerio 24912 | 1,80E-10 | 1174 | XP 005173060.1 | protein transport protein Sec31A isoform X1 [Danio rerio] >gi\|528496920\|ref\|XP 00517 | 1174 | 0 | 2407,1 | 100 |
| EP00428 | Sar;Alveolata;Myzozoa;"DP clade";Dinoflagellata;D | EP00428 Durinskia baltica 42789 | 1,90E-10 | 85 | ***no hit found*** | | | | | |
| EP00019 | Amoebozoa;Evosea;Cutosea;Armaparvus;Armaparv | EP00019 Armaparvus languidus 1559 | 1,90E-10 | 1144 | XP 012754771.1 | hypothetical protein SAMD00019534 058260 [Acytostelium subglobosum LB1] | 1179 | 1,69E-170 | 544,273 | 45,55125725 |
| EP00303 | Haptista;Haptophyta;Pavlovales;Exanthemachrysis; | EP00303 Exanthemachrysis gayraliae 14080 | 2,00E-10 | 433 | XP 008480553.3 | uncharacterized protein DDB G0290685-like [Diaphorina citri] | 1958 | 1,77E-09 | 71,633 | 39,453125 |
| EP00376 | Sar;Alveolata;Myzozoa;"AC clade";Apicomplexa;"cc | EP00376 Eimeria tenella 3338 | 2,10E-10 | 1193 | XP 013231069.1 | zinc finger (C3HC4 type RING finger) protein, putative, partial [Eimeria tenella] | 1193 | 0 | 2358,95 | 100 |
| EP00418 | Sar;Alveolata;Myzozoa;"DP clade";Dinoflagellata;D | EP00418 Pyrodinium bahamense 28772 | 2,10E-10 | 119 | ***no hit found*** | | | | | |
| EP00191 | Archaeplastida;Chloroplastida;Chlorophyta;"core ch | EP00191 Tetraselmis sp GSL018 23398 | 2,30E-10 | 1802 | ***no hit found*** | | | | | |
| EP00505 | Sar;Stramenopiles;Bigyra;Sagenista;"MAST-04";stra | EP00505 MAST4 sp SCG-Rhode-Island 2166 | 2,40E-10 | 2220 | ***no hit found*** | | | | | |
| EP00015 | Amoebozoa;Evosea;Archamoebea;Entamoebidae;E | EP00015 Entamoeba histolytica 8154 | 2,50E-10 | 112 | XP 657594.1 | hypothetical protein EHI 111070 [Entamoeba histolytica HM-1:IMSS] >gi\|672801350\|ref | 112 | 1,22E-74 | 225,713 | 100 |
| EP00016 | Amoebozoa;Evosea;Archamoebea;Entamoebidae;E | EP00016 Entamoeba nuttalli 1103 | 2,50E-10 | 112 | XP 657594.1 | hypothetical protein EHI 111070 [Entamoeba histolytica HM-1:IMSS] >gi\|672801350\|ref | 112 | 1,22E-74 | 225,713 | 100 |
| EP00406 | Sar;Alveolata;Myzozoa;"DP clade";Dinoflagellata;D | EP00406 Dinophysis acuminata 50073 | 2,60E-10 | 63 | XP 022812224.1 | conserved Plasmodium protein, unknown function [Plasmodium yoelii] | 1494 | 1,89E-06 | 53,1434 | 57,14285714 |
| EP00563 | Sar;Stramenopiles;Gyrista;Ochrophyta;"BD clade";[ | EP00563 Minutocellus polymorphus 15576 | 2,70E-10 | 1079 | XP 002290923.1 | predicted protein [Thalassiosira pseudonana CCMP1335] | 1041 | 0 | 1043,88 | 55,92286501 |
| EP00022 | Amoebozoa;Evosea;Eumycetozoa;Dictyostelia;Acyt | EP00022 Cavenderia fasciculata 6185 | 2,80E-10 | 1033 | XP 004358674.1 | hypothetical protein DFA 00689 [Cavenderia fasciculata] | 1033 | 0 | 2040,39 | 100 |
| EP00052 | Obazoa;Opisthokonta;Holozoa;Choanozoa;Choanof | EP00052 Salpingoeca macrocollata 22630 | 2,80E-10 | 763 | XP 009060088.1 | hypothetical protein LOTGIDRAFT 106381 [Lottia gigantea] | 914 | 4,92E-167 | 514,998 | 44,94680851 |
| EP00308 | Haptista;Haptophyta;Prymnesiophyceae;Calcihapto | EP00308 Calcidiscus leptoporus 24147 | 2,90E-10 | 1138 | XP 005788197.1 | secretory protein 31 [Emiliania huxleyi CCMP1516] | 1166 | 0 | 876,7 | 63,03162486 |
| EP00425 | Sar;Alveolata;Myzozoa;"DP clade";Dinoflagellata;D | EP00425 Heterocapsa triquetra 25847 | 2,90E-10 | 113 | ***no hit found*** | | | | | |
| EP00203 | Archaeplastida;Chloroplastida;Chlorophyta;"core ch | EP00203 Chlorella variabilis 8701 | 2,90E-10 | 1216 | XP 005851247.1 | hypothetical protein CHLNCDRAFT 137973 [Chlorella variabilis] | 1216 | 0 | 2368,58 | 100 |
| EP00368 | Sar;Alveolata;Myzozoa;"AC clade";Apicomplexa;"cc | EP00368 Plasmodium yoelii 1562 | 3,60E-10 | 1494 | XP 022812224.1 | conserved Plasmodium protein, unknown function [Plasmodium yoelii] | 1494 | 0 | 2847 | 100 |
| EP00444 | Sar;Alveolata;Myzozoa;"DP clade";Dinoflagellata;D | EP00444 Apocalathium aciculiferum 35181 | 3,60E-10 | 1202 | XP 002773527.1 | conserved hypothetical protein [Perkinsus marinus ATCC 50983] | 1211 | 0 | 642,884 | 49,39320388 |
| EP00262 | Archaeplastida;Chloroplastida;Streptophyta;"core s | EP00262 Sarcandra glabra 4271 | 3,70E-10 | 114 | XP 021814369.1 | protein transport protein SEC16B homolog [Prunus avium] | 1419 | 2,04E-51 | 183,726 | 78,07017544 |
| EP00023 | Amoebozoa;Evosea;Eumycetozoa;Dictyostelia;Dicty | EP00023 Dictyostelium discoideum 3935 | 3,70E-10 | 5222 | XP 637470.1 | hypothetical protein DDB G0286931 [Dictyostelium discoideum AX4] | 5222 | 0 | 10546,4 | 100 |
| EP00480 | Sar;Rhizaria;Retaria;Foraminifera;"monothalamid c | EP00480 Reticulomyxa filosa 26163 | 3,90E-10 | 710 | ***no hit found*** | | | | | |
| EP00598 | Sar;Stramenopiles;Gyrista;Ochrophyta;"CS clade";C | EP00598 Pedospumella elongata 3301 | 3,90E-10 | 1146 | XP 009515715.1 | hypothetical protein PHYSODRAFT 322106 [Phytophthora sojae] | 1262 | 0 | 649,047 | 49,05882353 |
| EP00440 | Sar;Alveolata;Myzozoa;"DP clade";Dinoflagellata;D | EP00440 Ansanella granifera 63599 | 3,90E-10 | 175 | ***no hit found*** | | | | | |
| EP00376 | Sar;Alveolata;Myzozoa;"AC clade";Apicomplexa;"cc | EP00376 Eimeria tenella 4813 | 4,00E-10 | 1631 | XP 013232546.1 | hypothetical protein ETH 00011470 [Eimeria tenella] | 1631 | 0 | 3221,41 | 100 |
| EP00419 | Sar;Alveolata;Myzozoa;"DP clade";Dinoflagellata;D | EP00419 Tripos fusus 63703 | 4,10E-10 | 113 | XP 022812224.1 | conserved Plasmodium protein, unknown function [Plasmodium yoelii] | 1494 | 9,04E-07 | 56,225 | 57,44680851 |
| EP00201 | Archaeplastida;Chloroplastida;Chlorophyta;"core ch | EP00201 Polytomella parva 598 | 4,10E-10 | 971 | XP 001697039.1 | COP-II coat subunit [Chlamydomonas reinhardtii] | 1313 | 0 | 800,816 | 64,22018349 |
| EP00200 | Archaeplastida;Chloroplastida;Chlorophyta;"core ch | EP00200 Dunaliella tertiolecta 11842 | 4,20E-10 | 1637 | ***no hit found*** | | | | | |
| EP00429 | Sar;Alveolata;Myzozoa;"DP clade";Dinoflagellata;D | EP00429 Kryptoperidinium foliaceum 1683 | 4,30E-10 | 998 | XP 002181151.1 | predicted protein [Phaeodactylum tricornutum CCAP 1055/1] | 1133 | 0 | 791,956 | 53,55191257 |
| EP00276 | Archaeplastida;Glaucophyta;Gloeochaete;Gloeocha | EP00276 Gloeochaete wittrockiana 12242 | 4,40E-10 | 3167 | XP 024926767.1 | E3 ubiquitin-protein ligase UPL1-like [Ziziphus jujuba] | 3746 | 0 | 766,918 | 45,98708487 |
| EP00405 | Sar;Alveolata;Myzozoa;"DP clade";Dinoflagellata;D | EP00405 Crypthecodinium cohnii 9801 | 4,50E-10 | 981 | ***no hit found*** | | | | | |
| EP00106 | Obazoa;Opisthokonta;Holozoa;Choanozoa;Metazoa | EP00106 Octopus bimaculoides 1774 | 4,50E-10 | 1231 | XP 014769216.1 | PREDICTED: protein transport protein Sec31A-like isoform X1 [Octopus bimaculoides] | 1231 | 0 | 2539,61 | 100 |
| EP00106 | Obazoa;Opisthokonta;Holozoa;Choanozoa;Metazoa | EP00106 Octopus bimaculoides 1778 | 4,70E-10 | 1220 | XP 014769220.1 | PREDICTED: protein transport protein Sec31A-like isoform X2 [Octopus bimaculoides] | 1220 | 0 | 2519,58 | 100 |
| EP00531 | Sar;Stramenopiles;Gyrista;Ochrophyta;"BD clade";[ | EP00531 Pseudo-nitzschia arenysensis 18197 | 4,70E-10 | 583 | XP 002177153.1 | predicted protein [Phaeodactylum tricornutum CCAP 1055/1] | 495 | 1,31E-26 | 124,405 | 72,85714286 |
| EP00191 | Archaeplastida;Chloroplastida;Chlorophyta;"core ch | EP00191 Tetraselmis sp GSL018 21 | 5,00E-10 | 656 | ***no hit found*** | | | | | |
| EP00014 | Amoebozoa;Evosea;Archamoebea;Entamoebidae;E | EP00014 Entamoeba dispar 7546 | 5,10E-10 | 112 | XP 001734782.1 | hypothetical protein [Entamoeba dispar SAW760] | 112 | 3,09E-74 | 224,557 | 100 |
| EP00561 | Sar;Stramenopiles;Gyrista;Ochrophyta;"BD clade";[ | EP00561 Arcocellulus cornucervis 6616 | 5,10E-10 | 838 | XP 002290923.1 | predicted protein [Thalassiosira pseudonana CCMP1335] | 1041 | 0 | 792,341 | 56 |
| EP00341 | Sar;Alveolata;Ciliophora;Intramacronucleata;"SAL"; | EP00341 Mesodinium pulex 10076 | 5,40E-10 | 183 | XP 002290923.1 | predicted protein [Thalassiosira pseudonana CCMP1335] | 1041 | 3,12E-50 | 183,726 | 52,43902439 |
| EP00063 | Obazoa;Opisthokonta;Holozoa;Choanozoa;Metazoa | EP00063 Salmo salar 57746 | 5,40E-10 | 271 | XP 014032581.1 | PREDICTED: uncharacterized protein LOC106588272 [Salmo salar] | 271 | 0 | 554,673 | 100 |
| EP00445 | Sar;Alveolata;Myzozoa;"DP clade";Dinoflagellata;D | EP00445 Apocalathium hangoei 95215 | 5,50E-10 | 1284 | XP 002773527.1 | conserved hypothetical protein [Perkinsus marinus ATCC 50983] | 1211 | 0 | 648,277 | 49,44785276 |
| EP00376 | Sar;Alveolata;Myzozoa;"AC clade";Apicomplexa;"cc | EP00376 Eimeria tenella 5554 | 5,80E-10 | 1062 | XP 013233287.1 | adaptin, putative, partial [Eimeria tenella] | 1062 | 0 | 2078,91 | 100 |
| EP00441 | Sar;Alveolata;Myzozoa;"DP clade";Dinoflagellata;D | EP00441 Pelagodinium beii 36985 | 6,10E-10 | 370 | ***no hit found*** | | | | | |
| EP00364 | Sar;Alveolata;Myzozoa;"AC clade";Apicomplexa;"cc | EP00364 Plasmodium chabaudi 2510 | 6,20E-10 | 1485 | XP 016655365.1 | conserved Plasmodium protein, unknown function [Plasmodium chabaudi chabaudi] | 1485 | 0 | 2876,66 | 100 |
| EP00473 | Sar;Rhizaria;Endomyxa;"phytorhiza";Phytomyxea;"( | EP00473 Plasmodiophora brassicae 4429 | 6,60E-10 | 1515 | XP 012262776.1 | major antigen-like isoform X1 [Athalia rosae] | 1710 | 8,84E-11 | 78,9518 | 39,07734057 |
| EP00357 | Sar;Alveolata;Ciliophora;Intramacronucleata;Protoc | EP00357 Protocruzia adherens 15826 | 6,70E-10 | 2336 | ***no hit found*** | | | | | |
| EP00157 | Obazoa;Opisthokonta;Nucletmycea;Fungi;"cryptom | EP00157 Rozella allomycis 1178 | 6,80E-10 | 1122 | XP 022104560.1 | protein transport protein Sec31A-like isoform X1 [Acanthaster planci] | 1228 | 5,33E-137 | 455,677 | 42,15530903 |
| EP00469 | Sar;Rhizaria;Cercozoa;Chlorarachnea;"core chlorara | EP00469 Lotharella globosa 8156 | 6,90E-10 | 1450 | XP 021372939.1 | macrophage mannose receptor 1-like [Mizuhopecten yessoensis] | 716 | 3,23E-27 | 130,954 | 40,42553191 |
| EP00101 | Obazoa;Opisthokonta;Holozoa;Choanozoa;Metazoa | EP00101 Apis mellifera 5311 | 7,00E-10 | 1247 | XP 006563951.2 | protein transport protein Sec31A [Apis mellifera] | 1247 | 0 | 2571,58 | 100 |
| EP00445 | Sar;Alveolata;Myzozoa;"DP clade";Dinoflagellata;D | EP00445 Apocalathium hangoei 41275 | 7,00E-10 | 1327 | XP 002773527.1 | conserved hypothetical protein [Perkinsus marinus ATCC 50983] | 1211 | 0 | 648,662 | 49,44785276 |
| EP00051 | Obazoa;Opisthokonta;Holozoa;Choanozoa;Choanof | EP00051 Salpingoeca urceolata 20382 | 7,00E-10 | 995 | XP 012556979.1 | PREDICTED: histone-lysine N-methyltransferase SETDB1-like [Hydra vulgaris] >gi\|8282064 | 1371 | 1,55E-44 | 187,193 | 42,94670846 |
| EP00740 | Archaeplastida;Chloroplastida;Chlorophyta;Mamiel | EP00740 Mantoniella antarctica 4844 | 7,10E-10 | 581 | XP 002503975.1 | predicted protein [Micromonas commoda] | 1399 | 1,62E-119 | 394,815 | 73,15789474 |
| EP00060 | Obazoa;Opisthokonta;Holozoa;Choanozoa;Metazoa | EP00060 Petromyzon marinus 43825 | 7,20E-10 | 608 | XP 008943825.1 | PREDICTED: circadian locomoter output cycles protein kaput, partial [Merops nubicus] | 483 | 7,51E-109 | 345,125 | 57,83783784 |
| EP00017 | Amoebozoa;Evosea;Archamoebea;Entamoebidae;E | EP00017 Entamoeba invadens 6625 | 7,40E-10 | 113 | XP 004256770.1 | hypothetical protein EIN 403130 [Entamoeba invadens IP1] | 113 | 6,18E-76 | 228,794 | 100 |
| EP00021 | Amoebozoa;Evosea;Eumycetozoa;Dictyostelia;Acyt | EP00021 Heterostelium album 6746 | 7,90E-10 | 1472 | XP 020433382.1 | putative mediator complex subunit 12 [Heterostelium album PN500] | 1472 | 0 | 3031,51 | 100 |
| EP00735 | Archaeplastida;"RR clade";Rhodelphis;Rhodelphis li | EP00735 Rhodelphis limneticus 4984 | 8,10E-10 | 1147 | XP 020585787.1 | protein transport protein SEC31 homolog B isoform X1 [Phalaenopsis equestris] | 1127 | 0 | 679,478 | 52,78232406 |
| EP00738 | Archaeplastida;Chloroplastida;Chlorophyta;Chlorop | EP00738 Chloropicon primus 2904 | 8,20E-10 | 718 | XP 005643560.1 | hypothetical protein COCSUDRAFT 48975 [Coccomyxa subellipsoidea C-169] | 1799 | 2,87E-07 | 65,855 | 51,26582278 |
| EP00429 | Sar;Alveolata;Myzozoa;"DP clade";Dinoflagellata;D | EP00429 Kryptoperidinium foliaceum 15684 | 8,30E-10 | 246 | XP 002183614.1 | predicted protein [Phaeodactylum tricornutum CCAP 1055/1] | 2083 | 2,47E-20 | 100,523 | 52,30769231 |
| EP00201 | Archaeplastida;Chloroplastida;Chlorophyta;"core ch | EP00201 Polytomella parva 15534 | 8,60E-10 | 356 | XP 001697039.1 | COP-II coat subunit [Chlamydomonas reinhardtii] | 1313 | 8,34E-61 | 221,476 | 58,84244373 |
| EP00376 | Sar;Alveolata;Myzozoa;"AC clade";Apicomplexa;"cc | EP00376 Eimeria tenella 4057 | 8,70E-10 | 982 | XP 013231790.1 | hypothetical protein ETH 00014925, partial [Eimeria tenella] | 982 | 0 | 1835,08 | 100 |
| EP00280 | Cryptista;Cryptophyceae;Cryptomonadales;"lineage | EP00280 Hanusia phi 15016 | 8,80E-10 | 973 | XP 005831033.1 | secretory protein Sec31 [Guillardia theta CCMP2712] | 1081 | 0 | 1642,86 | 93,99563319 |
| EP00120 | Obazoa;Opisthokonta;Holozoa;Filasterea;Capsaspo | EP00120 Capsaspora owczarzaki 536 | 8,90E-10 | 1308 | XP 004343028.1 | SEC31B protein [Capsaspora owczarzaki ATCC 30864] | 1308 | 0 | 2628,2 | 100 |
| EP00376 | Sar;Alveolata;Myzozoa;"AC clade";Apicomplexa;"cc | EP00376 Eimeria tenella 7335 | 8,90E-10 | 1155 | XP 013235074.1 | hypothetical protein, conserved, partial [Eimeria tenella] | 1155 | 0 | 2259,18 | 100 |
| EP00655 | Sar;Stramenopiles;Gyrista;Peronosporomycetes;"cc | EP00655 Saprolegnia parasitica 1785 | 9,10E-10 | 1021 | XP 012195291.1 | hypothetical protein SPRG 19095 [Saprolegnia parasitica CBS 223.65] | 1021 | 0 | 2097,78 | 100 |
| EP00401 | Sar;Alveolata;Myzozoa;"DP clade";Dinoflagellata;D | EP00401 Gymnodinium catenatum 30172 | 9,40E-10 | 464 | ***no hit found*** | | | | | |
| EP00376 | Sar;Alveolata;Myzozoa;"AC clade";Apicomplexa;"cc | EP00376 Eimeria tenella 295 | 9,80E-10 | 1234 | XP 013228021.1 | arginyl-tRNA synthetase, putative [Eimeria tenella] | 1234 | 0 | 2429,82 | 100 |
| EP00363 | Sar;Alveolata;Myzozoa;"AC clade";Apicomplexa;"cc | EP00363 Plasmodium berghei 1098 | 1,00E-09 | 1474 | XP 022712940.1 | conserved Plasmodium protein, unknown function [Plasmodium berghei ANKA] | 1474 | 0 | 2811,17 | 100 |
| EP00119 | Obazoa;Opisthokonta;Holozoa;Choanozoa;Metazoa | EP00119 Amphimedon queenslandica 20159 | 1,00E-09 | 683 | XP 019860578.1 | PREDICTED: uncharacterized protein LOC105315103 [Amphimedon queenslandica] | 683 | 0 | 1377,07 | 100 |
| EP00223 | Archaeplastida;Chloroplastida;Chlorophyta;Mamiel | EP00223 Ostreococcus mediterraneus 10002 | 1,00E-09 | 1708 | XP 022840888.1 | Serine/threonine-protein kinase, active site [Ostreococcus tauri] | 955 | 0 | 944,88 | 67,06706707 |
| EP00210 | Archaeplastida;Chloroplastida;Chlorophyta;"core ch | EP00210 Caulerpa lentillifera 8679 | 1,00E-09 | 916 | ***no hit found*** | | | | | |
| EP00161 | Ancyromonadida;Ancyromonadidae;Ancyromonas;/ | EP00161 Ancyromonas sigmoides 15051 | 1,00E-09 | 135 | XP 017235227.1 | PREDICTED: protein transport protein SEC16A homolog [Daucus carota subsp. sativus] >gi | 1558 | 1,12E-06 | 56,9954 | 47,2972973 |
| EP00376 | Sar;Alveolata;Myzozoa;"AC clade";Apicomplexa;"cc | EP00376 Eimeria tenella 1071 | 1,10E-09 | 1572 | XP 013228797.1 | hypothetical protein, conserved [Eimeria tenella] | 1572 | 0 | 2986,44 | 100 |
| EP00751 | Sar;Stramenopiles;Gyrista;Ochrophyta;"BD clade";[ | EP00751 Fragilariopsis kerguelensis 49921 | 1,10E-09 | 485 | XP 002183614.1 | predicted protein [Phaeodactylum tricornutum CCAP 1055/1] | 2083 | 2,08E-12 | 81,2629 | 46,36363636 |
| EP00062 | Obazoa;Opisthokonta;Holozoa;Choanozoa;Metazoa | EP00062 Callorhinchus milii 4896 | 1,20E-09 | 1236 | XP 007887151.1 | PREDICTED: protein transport protein Sec31A isoform X3 [Callorhinchus milii] | 1236 | 0 | 2548,85 | 100 |
| EP00383 | Sar;Alveolata;Myzozoa;"AC clade";Apicomplexa;Cry | EP00383 Cryptosporidium hominis 3742 | 1,30E-09 | 608 | XP 668554.1 | hypothetical protein [Cryptosporidium hominis TU502] | 608 | 0 | 1245,34 | 100 |
| EP00650 | Sar;Stramenopiles;Gyrista;Peronosporomycetes;"cc | EP00650 Pythium arrhenomanes 917 | 1,30E-09 | 202 | XP 002908536.1 | COPII coat assembly protein SEC16, putative [Phytophthora infestans T30-4] | 2641 | 2,03E-39 | 153,68 | 74,19354839 |
| EP00185 | Archaeplastida;"RR clade";Rhodophyta;Proteorhod | EP00185 Porphyridium purpureum 8872 | 1,30E-09 | 2500 | ***no hit found*** | | | | | |
| EP00658 | Telonemia;Telonema;strain P-2 | EP00658 Telonema sp P-2 14615 | 1,30E-09 | 464 | WP 141668442.1 | DUF4868 domain-containing protein [Clostridioides difficile] | 193 | 1,64E-07 | 62,003 | 82,35294118 |
| EP00746 | Sar;Alveolata;Myzozoa;"DP clade";Dinoflagellata;"I | EP00746 Dinoflagellata sp MGD 19809 | 1,30E-09 | 700 | XP 002773527.1 | conserved hypothetical protein [Perkinsus marinus ATCC 50983] | 1211 | 0 | 605,905 | 52,94117647 |
| EP00428 | Sar;Alveolata;Myzozoa;"DP clade";Dinoflagellata;D | EP00428 Durinskia baltica 60558 | 1,30E-09 | 1187 | XP 009515715.1 | hypothetical protein PHYSODRAFT 322106 [Phytophthora sojae] | 1262 | 0 | 646,736 | 50,64478312 |
| EP00051 | Obazoa;Opisthokonta;Holozoa;Choanozoa;Choanof | EP00051 Salpingoeca urceolata 20383 | 1,30E-09 | 1055 | XP 012556979.1 | PREDICTED: histone-lysine N-methyltransferase SETDB1-like [Hydra vulgaris] >gi\|8282064 | 1371 | 5,54E-60 | 236,113 | 44,95867769 |
| EP00440 | Sar;Alveolata;Myzozoa;"DP clade";Dinoflagellata;D | EP00440 Ansanella granifera 49184 | 1,40E-09 | 876 | XP 002773527.1 | conserved hypothetical protein [Perkinsus marinus ATCC 50983] | 1211 | 0 | 614,764 | 48,41463415 |
| EP00654 | Sar;Stramenopiles;Gyrista;Peronosporomycetes;"cc | EP00654 Saprolegnia diclina 6829 | 1,50E-09 | 1189 | XP 008610512.1 | hypothetical protein SDRG 06509 [Saprolegnia diclina VS20] | 1189 | 0 | 2417,5 | 100 |
| EP00402 | Sar;Alveolata;Myzozoa;"DP clade";Dinoflagellata;D | EP00402 Amphidinium carterae 8724 | 1,50E-09 | 754 | XP 004356402.1 | hypothetical protein DFA 08919 [Cavenderia fasciculata] | 1450 | 8,07E-45 | 185,267 | 71,86147186 |
| EP00208 | Archaeplastida;Chloroplastida;Chlorophyta;"core ch | EP00208 Stichococcus sp RCC1054 8433 | 1,50E-09 | 1138 | XP 001697039.1 | COP-II coat subunit [Chlamydomonas reinhardtii] | 1313 | 4,11E-173 | 554,673 | 49,56236324 |
| EP00044 | Obazoa;Opisthokonta;Holozoa;Choanozoa;Choanof | EP00044 Salpingoeca rosetta 8725 | 1,60E-09 | 2827 | XP 004996158.1 | hypothetical protein PTSG 02659 [Salpingoeca rosetta] | 2827 | 0 | 5744,08 | 100 |
| EP00044 | Obazoa;Opisthokonta;Holozoa;Choanozoa;Choanof | EP00044 Salpingoeca rosetta 11281 | 1,70E-09 | 2050 | XP 004998714.1 | hypothetical protein PTSG 00162 [Salpingoeca rosetta] | 2050 | 0 | 4126,24 | 100 |
| EP00208 | Archaeplastida;Chloroplastida;Chlorophyta;"core ch | EP00208 Stichococcus sp RCC1054 8287 | 1,70E-09 | 1222 | XP 005846158.1 | hypothetical protein CHLNCDRAFT 58390 [Chlorella variabilis] | 98 | 1,54E-19 | 95,1301 | 57,89473684 |
| EP00528 | Sar;Stramenopiles;Gyrista;Ochrophyta;"BD clade";[ | EP00528 Fragilariopsis cylindrus 1558 | 1,70E-09 | 931 | ***no hit found*** | | | | | |
| EP00095 | Obazoa;Opisthokonta;Holozoa;Choanozoa;Metazoa | EP00095 Tigriopus kingsejongensis 8051 | 1,70E-09 | 1161 | XP 013780411.2 | protein transport protein Sec31A-like [Limulus polyphemus] | 1162 | 0 | 774,622 | 57,4002574 |
| EP00089 | Obazoa;Opisthokonta;Holozoa;Choanozoa;Metazoa | EP00089 Calanus finmarchicus 53472 | 1,70E-09 | 110 | XP 023346731.1 | protein transport protein Sec16-like isoform X2 [Eurytemora affinis] | 1941 | 5,02E-21 | 96,6709 | 58,65384615 |
| EP00005 | Amoebozoa;Discosea;Centramoebia;"core centram | EP00005 Dracoamoeba jomungandri 9333 | 1,80E-09 | 1409 | XP 024379159.1 | protein transport protein SEC31 homolog B-like isoform X2 [Physcomitrella patens] | 1166 | 1,95E-138 | 464,922 | 45,37131231 |
| EP00289 | Cryptista;Cryptophyceae;Cryptomonadales;"lineage | EP00289 Rhodomonas salina 14312 | 1,80E-09 | 2191 | XP 013417931.2 | fibrillin-1 [Lingula anatina] | 1757 | 3,89E-101 | 374,015 | 57,20720721 |
| EP00023 | Amoebozoa;Evosea;Eumycetozoa;Dictyostelia;Dicty | EP00023 Dictyostelium discoideum 6076 | 1,90E-09 | 3080 | XP 640187.1 | involucrin repeat-containing protein [Dictyostelium discoideum AX4] | 3080 | 0 | 6148,54 | 100 |
| EP00426 | Sar;Alveolata;Myzozoa;"DP clade";Dinoflagellata;D | EP00426 Karenia brevis 123932 | 1,90E-09 | 101 | ***no hit found*** | | | | | |
| EP00191 | Archaeplastida;Chloroplastida;Chlorophyta;"core ch | EP00191 Tetraselmis sp GSL018 21804 | 1,90E-09 | 436 | XP 024373338.1 | protein transport protein SEC16A homolog isoform X1 [Physcomitrella patens] >gi\|13738 | 1444 | 4,03E-14 | 86,2705 | 45,25862069 |
| EP00575 | Sar;Stramenopiles;Gyrista;Ochrophyta;"BD clade";[ | EP00575 Skeletonema marinoi 38793 | 1,90E-09 | 1101 | XP 002296320.1 | predicted protein [Thalassiosira pseudonana CCMP1335] | 1033 | 3,53E-38 | 166,777 | 46,18181818 |
| EP00626 | Sar;Stramenopiles;Gyrista;Ochrophyta;Pelagophyce | EP00626 Pelagococcus subviridis 3112 | 2,00E-09 | 269 | XP 009038384.1 | hypothetical protein AURANDRAFT 28308, partial [Aureococcus anophagefferens] | 728 | 5,98E-61 | 214,542 | 75,16339869 |
| EP00044 | Obazoa;Opisthokonta;Holozoa;Choanozoa;Choanof | EP00044 Salpingoeca rosetta 3454 | 2,00E-09 | 1670 | XP 004990887.1 | hypothetical protein PTSG 07388 [Salpingoeca rosetta] | 1670 | 0 | 3340,82 | 100 |
| EP00589 | Sar;Stramenopiles;Gyrista;Ochrophyta;"BD clade";[ | EP00589 Leptocylindrus danicus 12501 | 2,00E-09 | 319 | XP 024584649.1 | Regucalcin gene promoter region-related protein (RGPR) [Plasmopara halstedii] | 2672 | 5,79E-12 | 77,7962 | 54,30107527 |
| EP00166 | Archaeplastida;"RR clade";Rhodophyta;Cyanidiales; | EP00166 Cyanidium caldarium 2562 | 2,10E-09 | 327 | ***no hit found*** | | | | | |
| EP00474 | Sar;Rhizaria;Endomyxa;"phytorhiza";Phytomyxea;"( | EP00474 Spongospora subterranea 8637 | 2,10E-09 | 1691 | XP 005831033.1 | secretory protein Sec31 [Guillardia theta CCMP2712] | 1081 | 6,18E-115 | 402,134 | 42,51870324 |
| EP00654 | Sar;Stramenopiles;Gyrista;Peronosporomycetes;"cc | EP00654 Saprolegnia diclina 8243 | 2,20E-09 | 1006 | XP 008611926.1 | hypothetical protein SDRG 07848 [Saprolegnia diclina VS20] | 1006 | 0 | 2066,58 | 100 |
| EP00077 | Obazoa;Opisthokonta;Holozoa;Choanozoa;Metazoa | EP00077 Pelodiscus sinensis 18401 | 2,20E-09 | 1215 | XP 014426335.1 | LOW QUALITY PROTEIN: protein transport protein Sec31A [Pelodiscus sinensis] | 1215 | 0 | 2509,95 | 100 |
| EP00567 | Sar;Stramenopiles;Gyrista;Ochrophyta;"BD clade";[ | EP00567 Triceratium dubium 1627 | 2,20E-09 | 950 | XP 002287238.1 | predicted protein [Thalassiosira pseudonana CCMP1335] | 997 | 7,52E-86 | 306,99 | 54,71976401 |
| EP00675 | Discoba;Euglenozoa;Kinetoplastea;Metakinetoplast | EP00675 Leptomonas pyrrhocoris 1307 | 2,30E-09 | 2387 | XP 015652811.1 | putative ubiquitin-protein ligase [Leptomonas pyrrhocoris] | 2387 | 0 | 4831,54 | 100 |
| EP00376 | Sar;Alveolata;Myzozoa;"AC clade";Apicomplexa;"cc | EP00376 Eimeria tenella 162 | 2,30E-09 | 3419 | XP 013227888.1 | hypothetical protein ETH 00005215 [Eimeria tenella] | 3419 | 0 | 6584,97 | 100 |
| EP00512 | Sar;Stramenopiles;Bigyra;Sagenista;Labyrinthulomy | EP00512 Aurantiochytrium limacinum 9445 | 2,30E-09 | 1257 | XP 009515715.1 | hypothetical protein PHYSODRAFT 322106 [Phytophthora sojae] | 1262 | 4,34E-140 | 468,389 | 51,18790497 |
| EP00044 | Obazoa;Opisthokonta;Holozoa;Choanozoa;Choanof | EP00044 Salpingoeca rosetta 641 | 2,30E-09 | 2720 | XP 004988074.1 | Ptpn18 protein [Salpingoeca rosetta] | 2720 | 0 | 5502,56 | 100 |
| EP00184 | Archaeplastida;"RR clade";Rhodophyta;Proteorhod | EP00184 Porphyridium aerugineum 801 | 2,40E-09 | 673 | XP 021631664.1 | protein transport protein SEC16B homolog isoform X2 [Manihot esculenta] | 1399 | 4,42E-09 | 71,633 | 55,06756757 |
| EP00376 | Sar;Alveolata;Myzozoa;"AC clade";Apicomplexa;"cc | EP00376 Eimeria tenella 4724 | 2,40E-09 | 1152 | XP 013232457.1 | hypothetical protein, conserved [Eimeria tenella] | 1152 | 0 | 2183,68 | 100 |
| EP00376 | Sar;Alveolata;Myzozoa;"AC clade";Apicomplexa;"cc | EP00376 Eimeria tenella 5715 | 2,50E-09 | 1374 | XP 013233448.1 | hypothetical protein, conserved [Eimeria tenella] | 1374 | 0 | 2648,23 | 100 |
| EP00063 | Obazoa;Opisthokonta;Holozoa;Choanozoa;Metazoa | EP00063 Salmo salar 78169 | 2,60E-09 | 1220 | XP 014053004.1 | PREDICTED: ubiquitin carboxyl-terminal hydrolase 28-like isoform X2 [Salmo salar] | 1220 | 0 | 2542,3 | 100 |
| EP00070 | Obazoa;Opisthokonta;Holozoa;Choanozoa;Metazoa | EP00070 Physeter macrocephalus 40897 | 2,70E-09 | 1237 | XP 028347791.1 | protein transport protein Sec31A isoform X5 [Physeter catodon] | 1237 | 0 | 2550,01 | 100 |
| EP00070 | Obazoa;Opisthokonta;Holozoa;Choanozoa;Metazoa | EP00070 Physeter macrocephalus 40895 | 2,80E-09 | 1250 | XP 028347789.1 | protein transport protein Sec31A isoform X3 [Physeter catodon] | 1250 | 0 | 2578,51 | 100 |
| EP00070 | Obazoa;Opisthokonta;Holozoa;Choanozoa;Metazoa | EP00070 Physeter macrocephalus 14064 | 2,80E-09 | 1252 | XP 023971132.1 | protein transport protein Sec31A isoform X2 [Physeter catodon] | 1252 | 0 | 2581,21 | 100 |
| EP00190 | Archaeplastida;"RR clade";Rhodophyta;Proteorhod | EP00190 Stylonematophyceae sp CCMP1999 5670 | 2,80E-09 | 94 | XP 006646765.2 | PREDICTED: protein transport protein SEC16A homolog [Oryza brachyantha] | 1514 | 1,07E-08 | 60,8474 | 40 |
| EP00041 | Obazoa;Opisthokonta;Holozoa;Choanozoa;Choanof | EP00041 Stephanoeca diplocostata 38873 | 2,80E-09 | 1098 | ***no hit found*** | | | | | |
| EP00070 | Obazoa;Opisthokonta;Holozoa;Choanozoa;Metazoa | EP00070 Physeter macrocephalus 14061 | 2,90E-09 | 1265 | XP 023971129.1 | protein transport protein Sec31A isoform X1 [Physeter catodon] >gi\|1348534269\|ref\|XP | 1265 | 0 | 2610,1 | 100 |
| EP00070 | Obazoa;Opisthokonta;Holozoa;Choanozoa;Metazoa | EP00070 Physeter macrocephalus 14062 | 2,90E-09 | 1265 | XP 023971129.1 | protein transport protein Sec31A isoform X1 [Physeter catodon] >gi\|1348534269\|ref\|XP | 1265 | 0 | 2610,1 | 100 |
| EP00070 | Obazoa;Opisthokonta;Holozoa;Choanozoa;Metazoa | EP00070 Physeter macrocephalus 14063 | 2,90E-09 | 1265 | XP 023971129.1 | protein transport protein Sec31A isoform X1 [Physeter catodon] >gi\|1348534269\|ref\|XP | 1265 | 0 | 2610,1 | 100 |
| EP00070 | Obazoa;Opisthokonta;Holozoa;Choanozoa;Metazoa | EP00070 Physeter macrocephalus 40894 | 2,90E-09 | 1265 | XP 023971129.1 | protein transport protein Sec31A isoform X1 [Physeter catodon] >gi\|1348534269\|ref\|XP | 1265 | 0 | 2610,1 | 100 |
| EP00062 | Obazoa;Opisthokonta;Holozoa;Choanozoa;Metazoa | EP00062 Callorhinchus milii 4893 | 2,90E-09 | 1173 | XP 007887152.1 | PREDICTED: protein transport protein Sec31A isoform X4 [Callorhinchus milii] | 1173 | 0 | 2421,74 | 100 |
| EP00376 | Sar;Alveolata;Myzozoa;"AC clade";Apicomplexa;"cc | EP00376 Eimeria tenella 368 | 3,10E-09 | 2473 | XP 013228094.1 | NEK kinase, putative [Eimeria tenella] | 2473 | 0 | 5044,56 | 100 |
| EP00089 | Obazoa;Opisthokonta;Holozoa;Choanozoa;Metazoa | EP00089 Calanus finmarchicus 23844 | 3,20E-09 | 492 | XP 023336929.1 | protein transport protein Sec31A-like [Eurytemora affinis] | 1195 | 0 | 578,556 | 66,53225806 |
| EP00062 | Obazoa;Opisthokonta;Holozoa;Choanozoa;Metazoa | EP00062 Callorhinchus milii 4894 | 3,20E-09 | 1248 | XP 007887150.1 | PREDICTED: protein transport protein Sec31A isoform X2 [Callorhinchus milii] | 1248 | 0 | 2571,58 | 100 |
| EP00062 | Obazoa;Opisthokonta;Holozoa;Choanozoa;Metazoa | EP00062 Callorhinchus milii 4892 | 3,30E-09 | 1261 | XP 007887149.1 | PREDICTED: protein transport protein Sec31A isoform X1 [Callorhinchus milii] | 1261 | 0 | 2600,47 | 100 |
| EP00652 | Sar;Stramenopiles;Gyrista;Peronosporomycetes;"cc | EP00652 Aphanomyces astaci 4330 | 3,30E-09 | 751 | XP 009825091.1 | hypothetical protein, variant [Aphanomyces astaci] | 751 | 0 | 1504,58 | 100 |
| EP00732 | Obazoa;Opisthokonta;Nucletmycea;Rotosphaerida; | EP00732 Lithocolla globosa 751 | 3,40E-09 | 1073 | XP 025022721.1 | protein transport protein Sec31A isoform X7 [Python bivittatus] | 1137 | 7,54E-163 | 520,776 | 45,30059272 |
| EP00136 | Obazoa;Opisthokonta;Nucletmycea;Fungi;"core Fur | EP00136 Aspergillus niger 7777 | 3,50E-09 | 1259 | XP 001399310.1 | protein transport protein sec31 [Aspergillus niger CBS 513.88] >gi\|1419017715\|ref\|XP 0 | 1259 | 0 | 2575,05 | 100 |
| EP00286 | Cryptista;Cryptophyceae;Cryptomonadales;"lineage | EP00286 Rhodomonas abbreviata 21035 | 3,50E-09 | 340 | XP 005831033.1 | secretory protein Sec31 [Guillardia theta CCMP2712] | 1081 | 5,24E-150 | 458,759 | 74,24749164 |
| EP00611 | Sar;Stramenopiles;Gyrista;Ochrophyta;"PX clade";X | EP00611 Tribonema gayanum 13758 | 3,60E-09 | 104 | XP 012195291.1 | hypothetical protein SPRG 19095 [Saprolegnia parasitica CBS 223.65] | 1021 | 9,39E-36 | 138,658 | 62,13592233 |
| EP00060 | Obazoa;Opisthokonta;Holozoa;Choanozoa;Metazoa | EP00060 Petromyzon marinus 40316 | 3,60E-09 | 707 | XP 014340496.1 | PREDICTED: protein transport protein Sec31B [Latimeria chalumnae] | 1243 | 0 | 978,393 | 70,05494505 |
| EP00317 | Haptista;Haptophyta;Prymnesiophyceae;Phaeocyst | EP00317 Phaeocystis antarctica 25598 | 3,70E-09 | 932 | WP 123259744.1 | cell envelope integrity protein TolA [Bacillus circulans] | 983 | 2,03E-14 | 89,7373 | 61,01694915 |
| EP00443 | Sar;Alveolata;Myzozoa;"DP clade";Dinoflagellata;D | EP00443 Scrippsiella trochoidea 42410 | 3,70E-09 | 1186 | ***no hit found*** | | | | | |
| EP00432 | Sar;Alveolata;Myzozoa;"DP clade";Dinoflagellata;D | EP00432 Prorocentrum cordatum 105776 | 3,70E-09 | 230 | XP 003881576.1 | putative plectin [Neospora caninum Liverpool] | 2378 | 1,04E-06 | 60,077 | 51,92307692 |
| EP00200 | Archaeplastida;Chloroplastida;Chlorophyta;"core ch | EP00200 Dunaliella tertiolecta 9255 | 3,90E-09 | 1379 | XP 011400305.1 | Alpha-L-arabinofuranosidase 1 [Auxenochlorella protothecoides] | 716 | 1,86E-85 | 305,449 | 47,81783681 |
| EP00172 | Archaeplastida;"RR clade";Rhodophyta;Eurhodophy | EP00172 Hildenbrandia rubra 174 | 4,20E-09 | 1215 | XP 005716292.1 | unnamed protein product [Chondrus crispus] | 1246 | 0 | 617,846 | 43,80853278 |
| EP00200 | Archaeplastida;Chloroplastida;Chlorophyta;"core ch | EP00200 Dunaliella tertiolecta 17714 | 4,30E-09 | 1249 | XP 002954917.1 | hypothetical protein VOLCADRAFT 118887 [Volvox carteri f. nagariensis] | 4212 | 4,00E-32 | 149,058 | 42,88888889 |
| EP00200 | Archaeplastida;Chloroplastida;Chlorophyta;"core ch | EP00200 Dunaliella tertiolecta 14803 | 4,40E-09 | 1534 | XP 002956610.1 | hypothetical protein VOLCADRAFT 67166 [Volvox carteri f. nagariensis] | 656 | 4,47E-105 | 360,533 | 73,64016736 |
| EP00290 | Cryptista;Cryptophyceae;Cryptomonadales;Baffinel | EP00290 Baffinella sp CCMP2293 57086 | 4,50E-09 | 712 | ***no hit found*** | | | | | |
| EP00562 | Sar;Stramenopiles;Gyrista;Ochrophyta;"BD clade";[ | EP00562 Extubocellulus spinifer 25162 | 4,50E-09 | 909 | ***no hit found*** | | | | | |
| EP00411 | Sar;Alveolata;Myzozoa;"DP clade";Dinoflagellata;D | EP00411 Alexandrium monilatum 100826 | 4,80E-09 | 66 | ***no hit found*** | | | | | |
| EP00652 | Sar;Stramenopiles;Gyrista;Peronosporomycetes;"cc | EP00652 Aphanomyces astaci 4329 | 4,90E-09 | 994 | XP 009825090.1 | hypothetical protein H257 02942 [Aphanomyces astaci] | 994 | 0 | 1994,16 | 100 |
| EP00004 | CRuMs;Rigifilida;Rigifila;Rigifila ramosa;strain CCAF | EP00004 Rigifila ramosa 18181 | 4,90E-09 | 1100 | XP 011432043.1 | PREDICTED: protein transport protein Sec31A isoform X2 [Crassostrea gigas] | 1191 | 1,74E-172 | 548,51 | 44,9382716 |
| EP00510 | Sar;Stramenopiles;Bigyra;Sagenista;Labyrinthulomy | EP00510 Aplanochytrium kerguelense 124 | 4,90E-09 | 124 | ***no hit found*** | | | | | |
| EP00405 | Sar;Alveolata;Myzozoa;"DP clade";Dinoflagellata;D | EP00405 Crypthecodinium cohnii 18218 | 4,90E-09 | 1250 | XP 002773527.1 | conserved hypothetical protein [Perkinsus marinus ATCC 50983] | 1211 | 0 | 625,55 | 47,60736196 |
| EP00370 | Sar;Alveolata;Myzozoa;"AC clade";Apicomplexa;"cc | EP00370 Babesia bigemina 4448 | 5,00E-09 | 138 | XP 012770305.1 | -COPII coat assembly protein sec16 [Babesia bigemina] | 138 | 2,37E-98 | 287,73 | 100 |
| EP00289 | Cryptista;Cryptophyceae;Cryptomonadales;"lineage | EP00289 Rhodomonas salina 9244 | 5,20E-09 | 1751 | XP 019623052.1 | PREDICTED: fibrillin-2-like [Branchiostoma belcheri] | 1067 | 2,90E-174 | 565,844 | 57,7874818 |
| EP00376 | Sar;Alveolata;Myzozoa;"AC clade";Apicomplexa;"cc | EP00376 Eimeria tenella 848 | 5,30E-09 | 623 | XP 013228574.1 | hypothetical protein, conserved [Eimeria tenella] | 623 | 0 | 1226,46 | 100 |
| EP00499 | Sar;Stramenopiles;Bigyra;Opalozoa;Bicosoecida;"cc | EP00499 Cafeteria sp CaronLabIsolate 2291 | 5,30E-09 | 281 | ***no hit found*** | | | | | |
| EP00261 | Archaeplastida;Chloroplastida;Streptophyta;"core s | EP00261 Vitis vinifera 17797 | 5,40E-09 | 167 | XP 010647605.2 | PREDICTED: protein transport protein SEC16B homolog [Vitis vinifera] | 167 | 5,43E-120 | 344,739 | 100 |
| EP00752 | Sar;Stramenopiles;Gyrista;Ochrophyta;"PX clade";P | EP00752 Nemacystus decipiens 5079 | 5,50E-09 | 992 | ***no hit found*** | | | | | |
| EP00751 | Sar;Stramenopiles;Gyrista;Ochrophyta;"BD clade";[ | EP00751 Fragilariopsis kerguelensis 43960 | 5,50E-09 | 863 | XP 002290400.1 | ankyrin domain-containing protein, partial [Thalassiosira pseudonana CCMP1335] | 139 | 9,86E-25 | 110,923 | 47,44525547 |
| EP00305 | Haptista;Haptophyta;Pavlovales;Diacronema;Diacr | EP00305 Diacronema lutheri 2524 | 5,60E-09 | 163 | XP 005831033.1 | secretory protein Sec31 [Guillardia theta CCMP2712] | 1081 | 1,14E-34 | 138,272 | 48,7012987 |
| EP00025 | Amoebozoa;Evosea;Eumycetozoa;Dictyostelia;Dicty | EP00025 Tieghemostelium lacteum 9592 | 5,60E-09 | 1177 | XP 003288054.1 | hypothetical protein DICPUDRAFT 152246 [Dictyostelium purpureum] | 1228 | 0 | 1027,31 | 58,81889764 |
| EP00445 | Sar;Alveolata;Myzozoa;"DP clade";Dinoflagellata;D | EP00445 Apocalathium hangoei 41459 | 5,90E-09 | 861 | ***no hit found*** | | | | | |
| EP00545 | Sar;Stramenopiles;Gyrista;Ochrophyta;"BD clade";[ | EP00545 Synedropsis sp CCMP1620 5490 | 5,90E-09 | 245 | XP 002183614.1 | predicted protein [Phaeodactylum tricornutum CCAP 1055/1] | 2083 | 4,30E-17 | 90,8929 | 53,10734463 |
| EP00653 | Sar;Stramenopiles;Gyrista;Peronosporomycetes;"cc | EP00653 Aphanomyces invadans 5120 | 6,00E-09 | 1040 | XP 008865917.1 | hypothetical protein H310 03725 [Aphanomyces invadans] | 1040 | 0 | 2101,63 | 100 |
| EP00070 | Obazoa;Opisthokonta;Holozoa;Choanozoa;Metazoa | EP00070 Physeter macrocephalus 40896 | 6,40E-09 | 1239 | XP 028347790.1 | protein transport protein Sec31A isoform X4 [Physeter catodon] | 1239 | 0 | 2555,79 | 100 |
| EP00044 | Obazoa;Opisthokonta;Holozoa;Choanozoa;Choanof | EP00044 Salpingoeca rosetta 866 | 6,50E-09 | 2214 | XP 004988299.1 | TKL protein kinase [Salpingoeca rosetta] | 2214 | 0 | 4534,94 | 100 |

| | | | | | | | | | | | |
|---|---|---|---|---|---|---|---|---|---|---|---|
| EP00609 | Sar;Stramenopiles;Gyrista;Ochrophyta;"PX clade";P | EP00609 | Ectocarpus siliculosus 12535 | 6,80E-09 | 942 | XP 002290923.1 | predicted protein [Thalassiosira pseudonana CCMP1335] | 1041 | 4,26E-114 | 384,8 | 48,12785388 |
| EP00656 | Telonemia;Telonema;"Tsubtile clade";Telonema sul | EP00656 | Telonema subtile 42034 | 6,90E-09 | 157 | ***no hit found*** | | | | | |
| EP00023 | Amoebozoa;Evosea;Eumycetozoa;Dictyostelia;Dicty | EP00023 | Dictyostelium discoideum 2185 | 6,90E-09 | 5650 | XP 635620.1 | hypothetical protein DDB G0290641 [Dictyostelium discoideum AX4] | 5650 | 0 | 11452,7 | 100 |
| EP00752 | Sar;Stramenopiles;Gyrista;Ochrophyta;"PX clade";P | EP00752 | Nemacystus decipiens 7249 | 7,20E-09 | 1196 | XP 009515715.1 | hypothetical protein PHYSODRAFT 322106 [Phytophthora sojae] | 1262 | 4,65E-164 | 530,791 | 53,39578454 |
| EP00500 | Sar;Stramenopiles;Bigyra;Opalozoa;Bicosoecida;stra | EP00500 | Bicosoecida sp ms1 4083 | 7,40E-09 | 962 | XP 009515715.1 | hypothetical protein PHYSODRAFT 322106 [Phytophthora sojae] | 1262 | 2,37E-170 | 540,421 | 46,635183 |
| EP00381 | Sar;Alveolata;Myzozoa;"AC clade";Apicomplexa;"gr | EP00381 | Gregarina niphandrodes 2706 | 7,50E-09 | 984 | XP 011131136.1 | hypothetical protein GNI 100640 [Gregarina niphandrodes] | 984 | 0 | 2029,22 | 100 |
| EP00531 | Sar;Stramenopiles;Gyrista;Ochrophyta;"BD clade";[ | EP00531 | Pseudo-nitzschia arenysensis 18131 | 7,60E-09 | 573 | XP 002177153.1 | predicted protein [Phaeodactylum tricornutum CCAP 1055/1] | 495 | 1,28E-26 | 124,02 | 72,85714286 |
| EP00482 | Sar;Rhizaria;Retaria;Foraminifera;Globothalamea;R | EP00482 | Globobulimina sp turgida-auriculata 121634 | 7,70E-09 | 228 | ***no hit found*** | | | | | |
| EP00376 | Sar;Alveolata;Myzozoa;"AC clade";Apicomplexa;"cc | EP00376 | Eimeria tenella 1431 | 7,80E-09 | 1081 | XP 013229157.1 | hypothetical protein, conserved [Eimeria tenella] | 1081 | 0 | 2127,44 | 100 |
| EP00089 | Obazoa;Opisthokonta;Holozoa;Choanozoa;Metazoa | EP00089 | Calanus finmarchicus 23837 | 7,80E-09 | 683 | XP 023336929.1 | protein transport protein Sec31A-like [Eurytemora affinis] | 1195 | 0 | 884,019 | 69,81132075 |
| EP00244 | Archaeplastida;Chloroplastida;Streptophyta;"core s | EP00244 | Chara vulgaris 8505 | 8,10E-09 | 210 | XP 010542224.1 | PREDICTED: protein transport protein SEC16B homolog [Tarenaya hassleriana] | 1321 | 4,12E-22 | 103,99 | 45,04504505 |
| EP00066 | Obazoa;Opisthokonta;Holozoa;Choanozoa;Metazoa | EP00066 | Takifugu rubripes 13766 | 8,10E-09 | 1329 | XP 011603032.2 | protein transport protein Sec31A isoform X2 [Takifugu rubripes] | 1329 | 0 | 2720,65 | 99,92475546 |
| EP00751 | Sar;Stramenopiles;Gyrista;Ochrophyta;"BD clade";[ | EP00751 | Fragilariopsis kerguelensis 13481 | 8,40E-09 | 1238 | XP 002177413.1 | predicted protein [Phaeodactylum tricornutum CCAP 1055/1] | 1046 | 3,49E-16 | 95,9005 | 48,54368932 |
| EP00581 | Sar;Stramenopiles;Gyrista;Ochrophyta;"BD clade";[ | EP00581 | Thalassiosira minuscula 10060 | 8,40E-09 | 473 | XP 002286037.1 | predicted protein [Thalassiosira pseudonana CCMP1335] | 328 | 3,91E-38 | 151,369 | 56,52173913 |
| EP00519 | Sar;Stramenopiles;Gyrista;Ochrophyta;"BD clade";E | EP00519 | Triparma laevis 12752 | 8,60E-09 | 303 | XP 002290923.1 | predicted protein [Thalassiosira pseudonana CCMP1335] | 1041 | 2,10E-68 | 239,965 | 54,13533835 |
| EP00327 | Haptista;Haptophyta;Prymnesiophyceae;Prymnesia | EP00327 | Prymnesium parvum 36788 | 8,70E-09 | 1057 | XP 005788197.1 | secretory protein 31 [Emiliania huxleyi CCMP1516] | 1166 | 0 | 825,469 | 59,47136564 |
| EP00044 | Obazoa;Opisthokonta;Holozoa;Choanozoa;Choanof | EP00044 | Salpingoeca rosetta 7506 | 8,70E-09 | 1220 | XP 004994939.1 | hypothetical protein PTSG 04042 [Salpingoeca rosetta] | 1220 | 0 | 2491,46 | 100 |
| EP00770 | Metamonada;Preaxostyla;Oxymonadida;Monocerc | EP00770 | Monocercomonoides exilis 6700 | 8,70E-09 | 2893 | ***no hit found*** | | | | | |
| EP00376 | Sar;Alveolata;Myzozoa;"AC clade";Apicomplexa;"cc | EP00376 | Eimeria tenella 1179 | 8,80E-09 | 882 | XP 013228905.1 | Chromosome II, complete genome, related [Eimeria tenella] | 882 | 0 | 1757,65 | 100 |
| EP00376 | Sar;Alveolata;Myzozoa;"AC clade";Apicomplexa;"cc | EP00376 | Eimeria tenella 523 | 9,00E-09 | 1147 | XP 013228249.1 | hypothetical protein, conserved [Eimeria tenella] | 1147 | 0 | 2244,93 | 100 |
| EP00760 | Discoba;Euglenozoa;Kinetoplastea;Prokinetoplastin | EP00760 | Prokinetoplastina sp PhM-4 23022 | 9,20E-09 | 969 | XP 001634737.1 | predicted protein [Nematostella vectensis] | 1219 | 4,03E-114 | 389,423 | 46,68008048 |
| EP00376 | Sar;Alveolata;Myzozoa;"AC clade";Apicomplexa;"cc | EP00376 | Eimeria tenella 3191 | 9,90E-09 | 1260 | XP 013230922.1 | hypothetical protein, conserved [Eimeria tenella] | 1260 | 0 | 2452,17 | 100 |
| EP00592 | Sar;Stramenopiles;Gyrista;Ochrophyta;"BD clade";[ | EP00592 | Proboscia alata 27301 | 1,00E-08 | 409 | XP 002183614.1 | predicted protein [Phaeodactylum tricornutum CCAP 1055/1] | 2083 | 1,76E-16 | 93,2041 | 60,13986014 |
| EP00376 | Sar;Alveolata;Myzozoa;"AC clade";Apicomplexa;"cc | EP00376 | Eimeria tenella 1620 | 1,00E-08 | 1423 | XP 013229346.1 | hypothetical protein, conserved [Eimeria tenella] | 1423 | 0 | 2841,6 | 100 |
| EP00529 | Sar;Stramenopiles;Gyrista;Ochrophyta;"BD clade";[ | EP00529 | Nitzschia sp RCC80 5956 | 1,00E-08 | 1012 | XP 002176599.1 | predicted protein [Phaeodactylum tricornutum CCAP 1055/1] | 1181 | 0 | 790,03 | 58,54779412 |
| EP00656 | Telonemia;Telonema;"Tsubtile clade";Telonema sul | EP00656 | Telonema subtile 42033 | 1,10E-08 | 145 | ***no hit found*** | | | | | |
| EP00749 | Sar;Stramenopiles;Bigyra;Opalozoa;Bicosoecida;"cc | EP00749 | Cafeteria roenbergensis 8382 | 1,10E-08 | 1653 | XP 022662203.1 | S-formylglutathione hydrolase-like [Varroa destructor] >gi\|1253756348\|ref\|XP 0226622( | 283 | 5,64E-83 | 284,648 | 57,25806452 |
| EP00089 | Obazoa;Opisthokonta;Holozoa;Choanozoa;Metazoa | EP00089 | Calanus finmarchicus 23845 | 1,10E-08 | 484 | XP 023336929.1 | protein transport protein Sec31A-like [Eurytemora affinis] | 1195 | 0 | 578,556 | 66,2601626 |
| EP00262 | Archaeplastida;Chloroplastida;Streptophyta;"core s | EP00262 | Sarcandra glabra 5652 | 1,10E-08 | 181 | XP 010264305.1 | PREDICTED: protein transport protein SEC16A homolog isoform X2 [Nelumbo nucifera] | 1393 | 2,49E-74 | 252,677 | 70,16574586 |
| EP00192 | Archaeplastida;Chloroplastida;Chlorophyta;"core ch | EP00192 | Tetraselmis astigmatica 1344 | 1,10E-08 | 2062 | ***no hit found*** | | | | | |
| EP00454 | Sar;Alveolata;Myzozoa;"DP clade";Perkinsea;Perkin | EP00454 | Perkinsus marinus 8557 | 1,10E-08 | 1575 | XP 002773822.1 | conserved hypothetical protein [Perkinsus marinus ATCC 50983] | 1575 | 0 | 3247,99 | 100 |
| EP00055 | Obazoa;Opisthokonta;Holozoa;Choanozoa;Choanof | EP00055 | Hartaetosiga balthica 14434 | 1,10E-08 | 254 | XP 004992030.1 | hypothetical protein PTSG 12463 [Salpingoeca rosetta] | 2149 | 4,70E-11 | 73,559 | 47,61904762 |
| EP00294 | Cryptista;Cryptophyceae;Goniomonas;"ma G clade" | EP00294 | Goniomonas avonlea 15459 | 1,10E-08 | 1091 | XP 005831033.1 | secretory protein Sec31 [Guillardia theta CCMP2712] | 1081 | 0 | 767,689 | 55,99505562 |
| EP00224 | Archaeplastida;Chloroplastida;Chlorophyta;Mamiel | EP00224 | Ostreococcus tauri 1691 | 1,20E-08 | 617 | XP 003079704.1 | Uso1/p115-like vesicle tethering protein,C-terminal [Ostreococcus tauri] | 617 | 0 | 1230,7 | 100 |
| EP00658 | Telonemia;Telonema;strain P-2 | EP00658 | Telonema sp P-2 14616 | 1,30E-08 | 324 | ***no hit found*** | | | | | |
| EP00348 | Sar;Alveolata;Ciliophora;Intramacronucleata;"SAL"; | EP00348 | Stylonychia lemnae 14086 | 1,30E-08 | 1350 | XP 002773527.1 | conserved hypothetical protein [Perkinsus marinus ATCC 50983] | 1211 | 3,70E-98 | 352,829 | 43,24324324 |
| EP00233 | Archaeplastida;Chloroplastida;Palmophyllophyceae | EP00233 | Prasinococcus capsulatus 2147 | 1,30E-08 | 1143 | XP 021844781.1 | golgin candidate 6 [Spinacia oleracea] | 920 | 1,26E-10 | 77,7962 | 47,68211921 |
| EP00185 | Archaeplastida;"RR clade";Rhodophyta;Proteorhod | EP00185 | Porphyridium purpureum 2160 | 1,30E-08 | 1313 | XP 020696568.1 | protein transport protein SEC31 homolog B [Dendrobium catenatum] | 1130 | 4,47E-99 | 353,214 | 48,90829694 |
| EP00404 | Sar;Alveolata;Myzozoa;"DP clade";Dinoflagellata;D | EP00404 | Azadinium spinosum 55158 | 1,40E-08 | 91 | XP 001615328.1 | liver stage antigen, putative [Plasmodium vivax] | 1507 | 6,43E-08 | 58,5362 | 47,36842105 |
| EP00758 | Discoba;Euglenozoa;Kinetoplastea;Metakinetoplast | EP00758 | Cryptobia borreli 1232 | 1,40E-08 | 420 | ***no hit found*** | | | | | |
| EP00242 | Archaeplastida;Chloroplastida;Chlorophyta;Pyramir | EP00242 | Pyramimonas sp CCMP2087 16974 | 1,40E-08 | 61 | XP 012845178.1 | PREDICTED: protein transport protein SEC16B homolog [Erythranthe guttata] >gi\|848890( | 1514 | 3,02E-08 | 57,7658 | 57,69230769 |
| EP00566 | Sar;Stramenopiles;Gyrista;Ochrophyta;"BD clade";[ | EP00566 | Odontella aurita 14322 | 1,40E-08 | 536 | ***no hit found*** | | | | | |
| EP00754 | Discoba;Euglenozoa;Diplonemea;Diplonemidae;Rhy | EP00754 | Rhynchopus humris 25154 | 1,50E-08 | 917 | ***no hit found*** | | | | | |
| EP00154 | Obazoa;Opisthokonta;Nucletmycea;Fungi;"core Fur | EP00154 | Conidiobolus coronatus 4769 | 1,50E-08 | 1281 | XP 021881067.1 | hypothetical protein BCR41DRAFT 396657 [Lobosporangium transversale] | 1406 | 0 | 691,034 | 52,51215559 |
| EP00076 | Obazoa;Opisthokonta;Holozoa;Choanozoa;Metazoa | EP00076 | Gallus gallus 13294 | 1,50E-08 | 1358 | XP 015131740.2 | protein transport protein Sec31A isoform X6 [Gallus gallus] | 1358 | 0 | 2794,61 | 100 |
| EP00458 | Sar;Rhizaria;Cercozoa;"crown cercozoa";Imbricatea | EP00458 | Abollifer prolabens 3273 | 1,60E-08 | 125 | ***no hit found*** | | | | | |
| EP00750 | Sar;Stramenopiles;Bigyra;Opalozoa;Nanomonadea; | EP00750 | Incisomonas marina 9284 | 1,60E-08 | 2638 | ***no hit found*** | | | | | |
| EP00217 | Archaeplastida;Chloroplastida;Chlorophyta;Mamiel | EP00217 | Crustomastix stigmatica 9069 | 1,60E-08 | 92 | XP 018455952.1 | PREDICTED: protein transport protein SEC16A homolog isoform X2 [Raphanus sativus] | 465 | 4,58E-07 | 56,225 | 53,7037037 |
| EP00456 | Sar;Rhizaria;Cercozoa;"crown cercozoa";Imbricatea | EP00456 | Euglypha rotunda 55925 | 1,70E-08 | 133 | ***no hit found*** | | | | | |
| EP00407 | Sar;Alveolata;Myzozoa;"DP clade";Dinoflagellata;D | EP00407 | Lingulodinium polyedra 131865 | 1,80E-08 | 1189 | ***no hit found*** | | | | | |
| EP00427 | Sar;Alveolata;Myzozoa;"DP clade";Dinoflagellata;D | EP00427 | Karlodinium veneficum 41561 | 1,80E-08 | 707 | ***no hit found*** | | | | | |
| EP00165 | Archaeplastida;"RR clade";Rhodophyta;Cyanidiales; | EP00165 | Cyanidioschyzon merolae 600 | 1,80E-08 | 2319 | XP 005535363.1 | hypothetical protein CYME CMC155C [Cyanidioschyzon merolae strain 10D] | 2319 | 0 | 4679,01 | 100 |
| EP00162 | Ancyromonadida;Ancyromonadidae;Nutomonas;Nu | EP00162 | Nutomonas longa 17191 | 1,90E-08 | 379 | XP 024987452.1 | protein transport protein SEC31 homolog B-like [Cynara cardunculus var. scolymus] | 1096 | 3,72E-46 | 179,874 | 39,02439024 |
| EP00376 | Sar;Alveolata;Myzozoa;"AC clade";Apicomplexa;"cc | EP00376 | Eimeria tenella 1136 | 1,90E-08 | 1042 | XP 013228862.1 | hypothetical protein, conserved [Eimeria tenella] | 1042 | 0 | 1917,12 | 100 |
| EP00049 | Obazoa;Opisthokonta;Holozoa;Choanozoa;Choanof | EP00049 | Salpingoeca infusionum 10021 | 1,90E-08 | 217 | XP 004992030.1 | hypothetical protein PTSG 12463 [Salpingoeca rosetta] | 2149 | 1,84E-09 | 67,781 | 52,5 |
| EP00653 | Sar;Stramenopiles;Gyrista;Peronosporomycetes;"co | EP00653 | Aphanomyces invadans 12744 | 2,00E-08 | 1217 | XP 008873541.1 | hypothetical protein H310 09285 [Aphanomyces invadans] | 1217 | 0 | 2493 | 100 |
| EP00120 | Obazoa;Opisthokonta;Holozoa;Filasterea;Capsaspoi | EP00120 | Capsaspora owczarzaki 7152 | 2,10E-08 | 2831 | XP 004365400.1 | hypothetical protein CAOG 00529 [Capsaspora owczarzaki ATCC 30864] | 2831 | 0 | 5725,59 | 100 |
| EP00639 | Sar;Stramenopiles;Gyrista;Ochrophyta;Raphidophy | EP00639 | Heterosigma akashiwo 34232 | 2,30E-08 | 277 | XP 024584649.1 | Regucalcin gene promoter region-related protein (RGPR) [Plasmopara halstedii] | 2672 | 6,04E-08 | 64,6994 | 45,67901235 |
| EP00376 | Sar;Alveolata;Myzozoa;"AC clade";Apicomplexa;"cc | EP00376 | Eimeria tenella 4234 | 2,30E-08 | 666 | XP 013231967.1 | hypothetical protein, conserved, partial [Eimeria tenella] | 666 | 0 | 1228 | 100 |
| EP00379 | Sar;Alveolata;Myzozoa;"AC clade";Apicomplexa;"cc | EP00379 | Toxoplasma gondii 7562 | 2,50E-08 | 2315 | XP 018637885.1 | surface antigen repeat-containing protein [Toxoplasma gondii ME49] | 2315 | 0 | 4590,79 | 100 |
| EP00613 | Sar;Stramenopiles;Gyrista;Ochrophyta;Dictyochoph | EP00613 | Pedinella sp CCMP2098 49256 | 2,50E-08 | 399 | XP 009515715.1 | hypothetical protein PHYSODRAFT 322106 [Phytophthora sojae] | 1262 | 1,71E-72 | 256,529 | 53,77358491 |
| EP00202 | Archaeplastida;Chloroplastida;Chlorophyta;"core ch | EP00202 | Volvox carteri 2501 | 2,50E-08 | 1302 | XP 002948107.1 | hypothetical protein VOLCADRAFT 120559 [Volvox carteri f. nagariensis] | 1302 | 0 | 2675,97 | 100 |
| EP00597 | Sar;Stramenopiles;Gyrista;Ochrophyta;"CS clade";C | EP00597 | Dinobryon sp UTEXLB2267 17383 | 2,50E-08 | 2313 | XP 005824228.1 | hypothetical protein GUITHDRAFT 165547 [Guillardia theta CCMP2712] | 1743 | 0 | 947,192 | 47,41863075 |
| EP00537 | Sar;Stramenopiles;Gyrista;Ochrophyta;"BD clade";[ | EP00537 | Pseudo-nitzschia pungens 10216 | 2,50E-08 | 714 | XP 002180114.1 | predicted protein [Phaeodactylum tricornutum CCAP 1055/1] | 441 | 1,45E-53 | 201,06 | 65,35947712 |
| EP00376 | Sar;Alveolata;Myzozoa;"AC clade";Apicomplexa;"cc | EP00376 | Eimeria tenella 8172 | 2,60E-08 | 1233 | XP 013235915.1 | hypothetical protein, conserved [Eimeria tenella] | 1233 | 0 | 2363,96 | 100 |
| EP00312 | Haptista;Haptophyta;Prymnesiophyceae;Calcihapto | EP00312 | Isochrysis sp CCMP1244 16808 | 2,60E-08 | 309 | ***no hit found*** | | | | | |
| EP00373 | Sar;Alveolata;Myzozoa;"AC clade";Apicomplexa;"cc | EP00373 | Theileria annulata 3447 | 2,60E-08 | 322 | XP 955204.1 | hypothetical protein TA04945 [Theileria annulata] | 322 | 0 | 670,233 | 100 |
| EP00089 | Obazoa;Opisthokonta;Holozoa;Choanozoa;Metazoa | EP00089 | Calanus finmarchicus 23838 | 2,70E-08 | 675 | XP 023336929.1 | protein transport protein Sec31A-like [Eurytemora affinis] | 1195 | 0 | 884,019 | 69,6350365 |
| EP00403 | Sar;Alveolata;Myzozoa;"DP clade";Dinoflagellata;D | EP00403 | Amphidinium massartii 7220 | 2,70E-08 | 983 | XP 002766585.1 | calcium-dependent protein kinase, putative [Perkinsus marinus ATCC 50983] | 331 | 1,63E-82 | 280,026 | 54,18060201 |
| EP00539 | Sar;Stramenopiles;Gyrista;Ochrophyta;"BD clade";[ | EP00539 | Tryblionella compressa 4876 | 2,80E-08 | 244 | XP 002183614.1 | predicted protein [Phaeodactylum tricornutum CCAP 1055/1] | 2083 | 1,09E-16 | 89,7373 | 57,71812081 |
| EP00426 | Sar;Alveolata;Myzozoa;"DP clade";Dinoflagellata;D | EP00426 | Karenia brevis 77842 | 2,90E-08 | 1151 | XP 002773527.1 | conserved hypothetical protein [Perkinsus marinus ATCC 50983] | 1211 | 0 | 629,402 | 50,3649635 |
| EP00613 | Sar;Stramenopiles;Gyrista;Ochrophyta;Dictyochoph | EP00613 | Pedinella sp CCMP2098 60745 | 2,90E-08 | 1123 | XP 005856379.1 | t-complex 11 like isoform cra b [Nannochloropsis gaditana CCMP526] | 976 | 9,20E-47 | 193,741 | 45,67901235 |
| EP00301 | Haptista;Centroplasthelida;Pterocystida;Raphidista | EP00301 | Raphidiophrys heterophryoidea 20150 | 2,90E-08 | 258 | ***no hit found*** | | | | | |
| EP00743 | Sar;Alveolata;Colponemidae;"MPE2-45 clade";strai | EP00743 | Colponemidia sp Colp-15 6695 | 3,00E-08 | 836 | XP 002773527.1 | conserved hypothetical protein [Perkinsus marinus ATCC 50983] | 1211 | 0 | 608,601 | 51,71568627 |
| EP00192 | Archaeplastida;Chloroplastida;Chlorophyta;"core ch | EP00192 | Tetraselmis astigmatica 1584 | 3,10E-08 | 2052 | ***no hit found*** | | | | | |
| EP00364 | Sar;Alveolata;Myzozoa;"AC clade";Apicomplexa;"cc | EP00364 | Plasmodium chabaudi 356 | 3,10E-08 | 1394 | XP 016653211.1 | protein transport protein SEC31, putative [Plasmodium chabaudi chabaudi] | 1394 | 0 | 2854,7 | 100 |
| EP00618 | Sar;Stramenopiles;Gyrista;Ochrophyta;Dictyochoph | EP00618 | Florenciella parvula 31829 | 3,20E-08 | 675 | XP 002289284.1 | probable eh domain-containing protein, partial [Thalassiosira pseudonana CCMP1335] | 382 | 0 | 538,11 | 69,13580247 |
| EP00099 | Obazoa;Opisthokonta;Holozoa;Choanozoa;Metazoa | EP00099 | Drosophila melanogaster 1345 | 3,20E-08 | 1209 | NP 001096900.3 | uncharacterized protein Dmel CG9650, isoform L [Drosophila melanogaster] | 1213 | 0 | 2473,74 | 100 |
| EP00751 | Sar;Stramenopiles;Gyrista;Ochrophyta;"BD clade";[ | EP00751 | Fragilariopsis kerguelensis 12022 | 3,20E-08 | 963 | ***no hit found*** | | | | | |
| EP00269 | Archaeplastida;Chloroplastida;Streptophyta;"core s | EP00269 | Selaginella moellendorffii 21884 | 3,30E-08 | 291 | XP 024522530.1 | pentatricopeptide repeat-containing protein At4g02750-like [Selaginella moellendorffii] | 291 | 0 | 603,979 | 100 |
| EP00595 | Sar;Stramenopiles;Gyrista;Ochrophyta;"CS clade";C | EP00595 | Chromulina nebulosa 2146 | 3,30E-08 | 354 | XP 002908536.1 | COPII coat assembly protein SEC16, putative [Phytophthora infestans T30-4] | 2641 | 8,91E-08 | 65,4698 | 48,1865285 |
| EP00145 | Obazoa;Opisthokonta;Nucletmycea;Fungi;"core Fur | EP00145 | Schizosaccharomyces pombe 3951 | 3,30E-08 | 1224 | NP 595582.1 | putative COPII-coated vesicle-associated protein Sec31 [Schizosaccharomyces pombe] | 1224 | 0 | 2503,4 | 100 |
| EP00118 | Obazoa;Opisthokonta;Holozoa;Choanozoa;Metazoa | EP00118 | Oscarella pearsei 25957 | 3,40E-08 | 1165 | XP 028397312.1 | protein transport protein Sec31A-like [Dendronephthya gigantea] | 1203 | 0 | 959,133 | 55,48387097 |
| EP00235 | Archaeplastida;Chloroplastida;Palmophyllophyceae | EP00235 | Prasinoderma singulare 307 | 3,50E-08 | 1714 | ***no hit found*** | | | | | |
| EP00613 | Sar;Stramenopiles;Gyrista;Ochrophyta;Dictyochoph | EP00613 | Pedinella sp CCMP2098 34992 | 3,50E-08 | 473 | XP 009515715.1 | hypothetical protein PHYSODRAFT 322106 [Phytophthora sojae] | 1262 | 4,35E-80 | 280,796 | 52,43128964 |
| EP00076 | Obazoa;Opisthokonta;Holozoa;Choanozoa;Metazoa | EP00076 | Gallus gallus 13300 | 3,60E-08 | 1214 | XP 015131746.1 | protein transport protein Sec31A isoform X11 [Gallus gallus] | 1214 | 0 | 2500,32 | 100 |
| EP00557 | Sar;Stramenopiles;Gyrista;Ochrophyta;"BD clade";[ | EP00557 | Chaetoceros sp GSL56 9427 | 3,60E-08 | 210 | XP 002183614.1 | predicted protein [Phaeodactylum tricornutum CCAP 1055/1] | 2083 | 9,92E-16 | 85,8853 | 48,52941176 |
| EP00076 | Obazoa;Opisthokonta;Holozoa;Choanozoa;Metazoa | EP00076 | Gallus gallus 13299 | 3,60E-08 | 1226 | XP 015131745.1 | protein transport protein Sec31A isoform X10 [Gallus gallus] | 1226 | 0 | 2524,97 | 100 |
| EP00076 | Obazoa;Opisthokonta;Holozoa;Choanozoa;Metazoa | EP00076 | Gallus gallus 13298 | 3,60E-08 | 1229 | XP 015131744.1 | protein transport protein Sec31A isoform X9 [Gallus gallus] | 1229 | 0 | 2530,36 | 100 |
| EP00451 | Sar;Alveolata;Myzozoa;"DP clade";Dinoflagellata;O | EP00451 | Oxyrrhis marina 12112 | 3,70E-08 | 411 | ***no hit found*** | | | | | |
| EP00076 | Obazoa;Opisthokonta;Holozoa;Choanozoa;Metazoa | EP00076 | Gallus gallus 13291 | 3,70E-08 | 1260 | XP 015131737.1 | protein transport protein Sec31A isoform X3 [Gallus gallus] | 1260 | 0 | 2595,85 | 100 |
| EP00076 | Obazoa;Opisthokonta;Holozoa;Choanozoa;Metazoa | EP00076 | Gallus gallus 13297 | 3,70E-08 | 1242 | XP 015131743.1 | protein transport protein Sec31A isoform X8 [Gallus gallus] | 1242 | 0 | 2558,1 | 100 |
| EP00269 | Archaeplastida;Chloroplastida;Streptophyta;"core s | EP00269 | Selaginella moellendorffii 35446 | 3,70E-08 | 179 | XP 024536092.1 | uncharacterized protein LOC112348194 [Selaginella moellendorffii] | 179 | 1,99E-129 | 369,392 | 100 |
| EP00076 | Obazoa;Opisthokonta;Holozoa;Choanozoa;Metazoa | EP00076 | Gallus gallus 1721 | 3,70E-08 | 1227 | NP 001026312.1 | protein transport protein Sec31A [Gallus gallus] | 1227 | 0 | 2529,21 | 100 |
| EP00216 | Archaeplastida;Chloroplastida;Chlorophyta;Chlorop | EP00216 | Chloropicon sp CCMP2111 5846 | 3,80E-08 | 632 | ***no hit found*** | | | | | |
| EP00284 | Cryptista;Cryptophyceae;Cryptomonadales;"lineage | EP00284 | Hemiselmis tepida 15242 | 3,90E-08 | 258 | XP 005831033.1 | secretory protein Sec31 [Guillardia theta CCMP2712] | 1081 | 1,43E-108 | 346,665 | 64,0625 |
| EP00076 | Obazoa;Opisthokonta;Holozoa;Choanozoa;Metazoa | EP00076 | Gallus gallus 13301 | 3,90E-08 | 1273 | XP 015131747.1 | protein transport protein Sec31A isoform X12 [Gallus gallus] >gi\|971393222\|ref\|XP 0151 | 1273 | 0 | 2620,5 | 100 |
| EP00076 | Obazoa;Opisthokonta;Holozoa;Choanozoa;Metazoa | EP00076 | Gallus gallus 13302 | 3,90E-08 | 1273 | XP 015131747.1 | protein transport protein Sec31A isoform X12 [Gallus gallus] >gi\|971393222\|ref\|XP 0151 | 1273 | 0 | 2620,5 | 100 |
| EP00076 | Obazoa;Opisthokonta;Holozoa;Choanozoa;Metazoa | EP00076 | Gallus gallus 13303 | 3,90E-08 | 1273 | XP 015131747.1 | protein transport protein Sec31A isoform X12 [Gallus gallus] >gi\|971393222\|ref\|XP 0151 | 1273 | 0 | 2620,5 | 100 |
| EP00076 | Obazoa;Opisthokonta;Holozoa;Choanozoa;Metazoa | EP00076 | Gallus gallus 39874 | 3,90E-08 | 1273 | XP 015131747.1 | protein transport protein Sec31A isoform X12 [Gallus gallus] >gi\|971393222\|ref\|XP 0151 | 1273 | 0 | 2620,5 | 100 |
| EP00210 | Archaeplastida;Chloroplastida;Chlorophyta;"core ch | EP00210 | Caulerpa lentillifera 1256 | 3,90E-08 | 964 | ***no hit found*** | | | | | |
| EP00611 | Sar;Stramenopiles;Gyrista;Ochrophyta;"PX clade";X | EP00611 | Tribonema gayanum 19171 | 4,00E-08 | 622 | XP 014152139.1 | hypothetical protein SARC 09322 [Sphaeroforma arctica JP610] | 856 | 2,96E-27 | 128,642 | 39,48717949 |
| EP00201 | Archaeplastida;Chloroplastida;Chlorophyta;"core ch | EP00201 | Polytomella parva 13367 | 4,00E-08 | 168 | ***no hit found*** | | | | | |
| EP00044 | Obazoa;Opisthokonta;Holozoa;Choanozoa;Choanof | EP00044 | Salpingoeca rosetta 140 | 4,10E-08 | 1162 | XP 004987573.1 | hypothetical protein PTSG 11422, partial [Salpingoeca rosetta] | 1162 | 0 | 2385,53 | 100 |
| EP00475 | Sar;Rhizaria;Endomyxa;"phytorhiza";Vampyrellida;' | EP00475 | Leptophrys vorax 6557 | 4,10E-08 | 338 | XP 017219748.1 | PREDICTED: protein transport protein SEC16B homolog [Daucus carota subsp. sativus] >gi\| | 1358 | 5,51E-15 | 87,0409 | 53,05164319 |
| EP00376 | Sar;Alveolata;Myzozoa;"AC clade";Apicomplexa;"cc | EP00376 | Eimeria tenella 678 | 4,20E-08 | 1466 | XP 013228404.1 | hypothetical protein, conserved [Eimeria tenella] | 1466 | 0 | 2854,7 | 100 |
| EP00076 | Obazoa;Opisthokonta;Holozoa;Choanozoa;Metazoa | EP00076 | Gallus gallus 13295 | 4,30E-08 | 1356 | XP 015131741.2 | protein transport protein Sec31A isoform X7 [Gallus gallus] | 1356 | 0 | 2793,45 | 100 |
| EP00294 | Cryptista;Cryptophyceae;Goniomonas;"ma G clade" | EP00294 | Goniomonas avonlea 15335 | 4,30E-08 | 1142 | XP 005831033.1 | secretory protein Sec31 [Guillardia theta CCMP2712] | 1081 | 0 | 800,816 | 56,21370499 |
| EP00213 | Archaeplastida;Chloroplastida;Chlorophyta;Chlorop | EP00213 | Chloropicon mariensis 7378 | 4,40E-08 | 817 | ***no hit found*** | | | | | |
| EP00066 | Obazoa;Opisthokonta;Holozoa;Choanozoa;Metazoa | EP00066 | Takifugu rubripes 13767 | 4,40E-08 | 1295 | XP 029693527.1 | protein transport protein Sec31A isoform X3 [Takifugu rubripes] | 1312 | 0 | 2641,68 | 99,92378049 |
| EP00076 | Obazoa;Opisthokonta;Holozoa;Choanozoa;Metazoa | EP00076 | Gallus gallus 13293 | 4,40E-08 | 1368 | XP 015131739.2 | protein transport protein Sec31A isoform X5 [Gallus gallus] | 1368 | 0 | 2813,87 | 100 |
| EP00076 | Obazoa;Opisthokonta;Holozoa;Choanozoa;Metazoa | EP00076 | Gallus gallus 13292 | 4,40E-08 | 1369 | XP 015131738.2 | protein transport protein Sec31A isoform X4 [Gallus gallus] | 1369 | 0 | 2815,41 | 100 |
| EP00076 | Obazoa;Opisthokonta;Holozoa;Choanozoa;Metazoa | EP00076 | Gallus gallus 13290 | 4,50E-08 | 1381 | XP 015131736.2 | protein transport protein Sec31A isoform X2 [Gallus gallus] | 1381 | 0 | 2840,83 | 100 |
| EP00076 | Obazoa;Opisthokonta;Holozoa;Choanozoa;Metazoa | EP00076 | Gallus gallus 13289 | 4,50E-08 | 1384 | XP 015131735.2 | protein transport protein Sec31A isoform X1 [Gallus gallus] | 1384 | 0 | 2847 | 100 |
| EP00089 | Obazoa;Opisthokonta;Holozoa;Choanozoa;Metazoa | EP00089 | Calanus finmarchicus 23846 | 4,80E-08 | 478 | XP 023336929.1 | protein transport protein Sec31A-like [Eurytemora affinis] | 1195 | 0 | 573,933 | 66,39344262 |
| EP00376 | Sar;Alveolata;Myzozoa;"AC clade";Apicomplexa;"cc | EP00376 | Eimeria tenella 4662 | 5,00E-08 | 1143 | XP 013232395.1 | hypothetical protein, conserved [Eimeria tenella] | 1143 | 0 | 2144,01 | 100 |
| EP00285 | Cryptista;Cryptophyceae;Cryptomonadales;"lineage | EP00285 | Hemiselmis virescens 13376 | 5,10E-08 | 517 | XP 005831033.1 | secretory protein Sec31 [Guillardia theta CCMP2712] | 1081 | 8,65E-151 | 468,389 | 57,77351248 |
| EP00706 | Metamonada;Parabasalia;Tritrichomonas;Tritrichor | EP00706 | Tritrichomonas foetus 17602 | 5,10E-08 | 746 | XP 001307212.1 | hypothetical protein [Trichomonas vaginalis G3] | 1164 | 1,94E-95 | 331,257 | 58,43270869 |
| EP00099 | Obazoa;Opisthokonta;Holozoa;Choanozoa;Metazoa | EP00099 | Drosophila melanogaster 1344 | 5,10E-08 | 1209 | NP 001096899.2 | uncharacterized protein Dmel CG9650, isoform G [Drosophila melanogaster] | 1209 | 0 | 2474,12 | 100 |
| EP00339 | Sar;Alveolata;Ciliophora;Intramacronucleata;"CON3 | EP00339 | Tiarina fusus 847 | 5,10E-08 | 987 | XP 025172421.1 | hypothetical protein GLOIN 2v1665522 [Rhizophagus irregularis DAOM 181602=DAOM 1 | 225 | 2,39E-39 | 155,992 | 46,19883041 |
| EP00367 | Sar;Alveolata;Myzozoa;"AC clade";Apicomplexa;"cc | EP00367 | Plasmodium vivax 3316 | 5,20E-08 | 1507 | XP 001615328.1 | liver stage antigen, putative [Plasmodium vivax] | 1507 | 0 | 3030,74 | 100 |
| EP00044 | Obazoa;Opisthokonta;Holozoa;Choanozoa;Choanof | EP00044 | Salpingoeca rosetta 3906 | 5,20E-08 | 1557 | XP 004991339.1 | hypothetical protein PTSG 07542 [Salpingoeca rosetta] | 1557 | 0 | 3123,18 | 100 |
| EP00200 | Archaeplastida;Chloroplastida;Chlorophyta;"core ch | EP00200 | Dunaliella tertiolecta 10243 | 5,20E-08 | 1422 | ***no hit found*** | | | | | |
| EP00191 | Archaeplastida;Chloroplastida;Chlorophyta;"core ch | EP00191 | Tetraselmis sp GSL018 16632 | 5,20E-08 | 1789 | ***no hit found*** | | | | | |
| EP00376 | Sar;Alveolata;Myzozoa;"AC clade";Apicomplexa;"cc | EP00376 | Eimeria tenella 8096 | 5,30E-08 | 668 | XP 013235839.1 | hypothetical protein, conserved, partial [Eimeria tenella] | 668 | 0 | 1301,19 | 100 |
| EP00618 | Sar;Stramenopiles;Gyrista;Ochrophyta;Dictyochoph | EP00618 | Florenciella parvula 12733 | 5,40E-08 | 840 | XP 009038384.1 | hypothetical protein AURANDRAFT 28308, partial [Aureococcus anophagefferens] | 728 | 0 | 678,707 | 60,13422819 |
| EP00475 | Sar;Rhizaria;Endomyxa;"phytorhiza";Vampyrellida;' | EP00475 | Leptophrys vorax 2693 | 5,50E-08 | 627 | ***no hit found*** | | | | | |
| EP00475 | Sar;Rhizaria;Endomyxa;"phytorhiza";Vampyrellida;' | EP00475 | Leptophrys vorax 2694 | 5,50E-08 | 627 | ***no hit found*** | | | | | |
| EP00475 | Sar;Rhizaria;Endomyxa;"phytorhiza";Vampyrellida;' | EP00475 | Leptophrys vorax 2695 | 5,50E-08 | 627 | ***no hit found*** | | | | | |
| EP00390 | Sar;Alveolata;Myzozoa;"AC clade";Colpodellida;Vitr | EP00390 | Vitrella brassicaformis 19563 | 5,60E-08 | 1821 | XP 007515401.1 | predicted protein [Bathycoccus prasinos] | 1306 | 1,79E-68 | 266,544 | 44,96350365 |
| EP00011 | Amoebozoa;Discosea;Flabellinia;"core flabellinids"; | EP00011 | Vannella sp DIVA3-517-6-12 12930 | 5,70E-08 | 1098 | XP 006482944.1 | protein transport protein SEC31 homolog B isoform X1 [Citrus sinensis] >gi\|1350271342\|r | 1120 | 9,31E-136 | 449,129 | 43,66471735 |
| EP00024 | Amoebozoa;Evosea;Eumycetozoa;Dictyostelia;Dicty | EP00024 | Dictyostelium purpureum 6792 | 5,90E-08 | 632 | XP 003289644.1 | hypothetical protein DICPUDRAFT 98447 [Dictyostelium purpureum] | 632 | 0 | 1277,69 | 100 |
| EP00405 | Sar;Alveolata;Myzozoa;"DP clade";Dinoflagellata;D | EP00405 | Crypthecodinium cohnii 50629 | 6,00E-08 | 710 | ***no hit found*** | | | | | |
| EP00580 | Sar;Stramenopiles;Gyrista;Ochrophyta;"BD clade";[ | EP00580 | Thalassiosira gravida 20787 | 6,50E-08 | 588 | XP 002296732.1 | hypothetical protein THAPSDRAFT 269874 [Thalassiosira pseudonana CCMP1335] | 277 | 5,22E-110 | 340,117 | 77,91798107 |
| EP00183 | Archaeplastida;"RR clade";Rhodophyta;Proteorhod | EP00183 | Erythrolobus madagascarensis 5823 | 7,00E-08 | 1250 | XP 022567395.1 | protein transport protein SEC31 homolog B isoform X1 [Brassica napus] | 1110 | 1,08E-120 | 411,764 | 45,96078431 |
| EP00376 | Sar;Alveolata;Myzozoa;"AC clade";Apicomplexa;"cc | EP00376 | Eimeria tenella 3817 | 7,10E-08 | 936 | XP 013231549.1 | hypothetical protein ETH 00041785, partial [Eimeria tenella] | 936 | 0 | 1745,71 | 100 |
| EP00315 | Haptista;Haptophyta;Prymnesiophyceae;Calcihapto | EP00315 | Gephyrocapsa oceanica 27691 | 7,10E-08 | 399 | ***no hit found*** | | | | | |
| EP00282 | Cryptista;Cryptophyceae;Cryptomonadales;"lineage | EP00282 | Hemiselmis andersenii 30750 | 7,10E-08 | 99 | ***no hit found*** | | | | | |
| EP00405 | Sar;Alveolata;Myzozoa;"DP clade";Dinoflagellata;D | EP00405 | Crypthecodinium cohnii 11073 | 7,20E-08 | 996 | ***no hit found*** | | | | | |
| EP00449 | Sar;Alveolata;Myzozoa;"DP clade";Dinoflagellata;D | EP00449 | Zooxanthella nutricula 58004 | 7,50E-08 | 812 | XP 002773527.1 | conserved hypothetical protein [Perkinsus marinus ATCC 50983] | 1211 | 0 | 638,647 | 50,24096386 |
| EP00630 | Sar;Stramenopiles;Gyrista;Ochrophyta;Pelagophyce | EP00630 | Chrysocystis fragilis 2945 | 7,60E-08 | 1060 | XP 002181151.1 | predicted protein [Phaeodactylum tricornutum CCAP 1055/1] | 1133 | 4,18E-66 | 253,832 | 44,56692913 |
| EP00333 | Sar;Alveolata;Ciliophora;Intramacronucleata;"CON3 | EP00333 | Tetrahymena thermophila 25593 | 7,60E-08 | 1488 | XP 012655749.1 | hypothetical protein TTHERM 000755929 [Tetrahymena thermophila SB210] | 1488 | 0 | 2912,09 | 100 |
| EP00405 | Sar;Alveolata;Myzozoa;"DP clade";Dinoflagellata;D | EP00405 | Crypthecodinium cohnii 52758 | 7,90E-08 | 748 | XP 002768122.1 | Carbon catabolite derepressing protein kinase, putative [Perkinsus marinus ATCC 50983] | 773 | 7,28E-147 | 457,603 | 69,91869919 |
| EP00280 | Cryptista;Cryptophyceae;Cryptomonadales;"lineage | EP00280 | Hanusia phi 4192 | 7,90E-08 | 666 | XP 005831033.1 | secretory protein Sec31 [Guillardia theta CCMP2712] | 1081 | 0 | 1018,84 | 91,58576052 |
| EP00770 | Metamonada;Preaxostyla;Oxymonadida;Monocerc | EP00770 | Monocercomonoides exilis 14011 | 8,20E-08 | 4620 | XP 024360346.1 | protein ILITYHIA-like [Physcomitrella patens] | 2614 | 4,26E-51 | 214,157 | 51,88216039 |
| EP00106 | Obazoa;Opisthokonta;Holozoa;Choanozoa;Metazoa | EP00106 | Octopus bimaculoides 1708 | 8,30E-08 | 1032 | XP 014769150.1 | PREDICTED: uncharacterized protein LOC106868418 [Octopus bimaculoides] | 1032 | 0 | 2151,71 | 100 |
| EP00445 | Sar;Alveolata;Myzozoa;"DP clade";Dinoflagellata;D | EP00445 | Apocalathium hangoei 46942 | 8,40E-08 | 159 | XP 007375323.1 | hypothetical protein SPAPADRAFT 61142 [Spathaspora passalidarum NRRL Y-27907] | 312 | 9,85E-07 | 57,3806 | 50,61728395 |
| EP00044 | Obazoa;Opisthokonta;Holozoa;Choanozoa;Choanof | EP00044 | Salpingoeca rosetta 305 | 8,50E-08 | 577 | XP 004987738.1 | hypothetical protein PTSG 11239 [Salpingoeca rosetta] | 577 | 0 | 1176,77 | 100 |
| EP00613 | Sar;Stramenopiles;Gyrista;Ochrophyta;Dictyochoph | EP00613 | Pedinella sp CCMP2098 17389 | 8,50E-08 | 982 | XP 009038384.1 | hypothetical protein AURANDRAFT 28308, partial [Aureococcus anophagefferens] | 728 | 5,19E-45 | 185,267 | 60,38961039 |
| EP00044 | Obazoa;Opisthokonta;Holozoa;Choanozoa;Choanof | EP00044 | Salpingoeca rosetta 7876 | 8,90E-08 | 2008 | XP 004995309.1 | hypothetical protein PTSG 03578 [Salpingoeca rosetta] | 2008 | 0 | 4106,21 | 100 |
| EP00448 | Sar;Alveolata;Myzozoa;"DP clade";Dinoflagellata;D | EP00448 | Togula jolla 809 | 8,90E-08 | 1118 | XP 002773527.1 | conserved hypothetical protein [Perkinsus marinus ATCC 50983] | 1211 | 0 | 664,84 | 51,12426036 |
| EP00366 | Sar;Alveolata;Myzozoa;"AC clade";Apicomplexa;"cc | EP00366 | Plasmodium knowlesi 1715 | 8,90E-08 | 1400 | XP 002259212.1 | hypothetical protein, conserved in Plasmodium species [Plasmodium knowlesi strain H] | 1400 | 0 | 2798,85 | 100 |
| EP00127 | Obazoa;Opisthokonta;Holozoa;Pluriformea;Corallo | EP00127 | Corallochytrium limacisporum 2296 | 9,00E-08 | 1210 | XP 028397312.1 | protein transport protein Sec31A-like [Dendronephthya gigantea] | 1203 | 0 | 608,601 | 46,86774942 |
| EP00073 | Obazoa;Opisthokonta;Holozoa;Choanozoa;Metazoa | EP00073 | Rattus norvegicus 15041 | 9,30E-08 | 1249 | NP 148981.1 | protein transport protein Sec31A [Rattus norvegicus] | 1249 | 0 | 2568,11 | 100 |
| EP00280 | Cryptista;Cryptophyceae;Cryptomonadales;"lineage | EP00280 | Hanusia phi 2459 | 9,40E-08 | 980 | XP 005831033.1 | secretory protein Sec31 [Guillardia theta CCMP2712] | 1081 | 0 | 1619,75 | 94,10480349 |
| EP00371 | Sar;Alveolata;Myzozoa;"AC clade";Apicomplexa;"cc | EP00371 | Babesia bovis 469 | 9,60E-08 | 136 | XP 001609116.1 | hypothetical protein [Babesia bovis T2Bo] | 136 | 2,45E-95 | 279,641 | 100 |
| EP00105 | Obazoa;Opisthokonta;Holozoa;Choanozoa;Metazoa | EP00105 | Crassostrea gigas 40449 | 9,70E-08 | 3047 | XP 019924597.1 | PREDICTED: probable chitinase 3 [Crassostrea gigas] | 3047 | 0 | 6369,65 | 100 |
| EP00120 | Obazoa;Opisthokonta;Holozoa;Filasterea;Capsaspoi | EP00120 | Capsaspora owczarzaki 2299 | 1,10E-07 | 898 | XP 004345696.1 | hypothetical protein CAOG 06106 [Capsaspora owczarzaki ATCC 30864] | 898 | 0 | 1822,75 | 100 |
| EP00013 | Amoebozoa;Discosea;Flabellinia;Stygamoeba;Styga | EP00013 | Stygamoeba regulata 15152 | 1,10E-07 | 1179 | XP 020165258.1 | ultraviolet-B receptor UVR8-like [Aegilops tauschii subsp. tauschii] | 442 | 2,89E-34 | 148,288 | 42,82238443 |
| EP00376 | Sar;Alveolata;Myzozoa;"AC clade";Apicomplexa;"cc | EP00376 | Eimeria tenella 4215 | 1,20E-07 | 1538 | XP 013231948.1 | hypothetical protein, conserved [Eimeria tenella] | 1538 | 0 | 3028,42 | 100 |
| EP00089 | Obazoa;Opisthokonta;Holozoa;Choanozoa;Metazoa | EP00089 | Calanus finmarchicus 23839 | 1,20E-07 | 669 | XP 023336929.1 | protein transport protein Sec31A-like [Eurytemora affinis] | 1195 | 0 | 878,626 | 69,75036711 |
| EP00099 | Obazoa;Opisthokonta;Holozoa;Choanozoa;Metazoa | EP00099 | Drosophila melanogaster 16620 | 1,20E-07 | 1140 | NP 727173.2 | uncharacterized protein Dmel CG9650, isoform I [Drosophila melanogaster] | 1207 | 0 | 2326,59 | 100 |
| EP00693 | Discoba;Jakobida;Histionina;Jakoba;Jakoba libera | EP00693 | Jakoba libera 1755 | 1,30E-07 | 217 | ***no hit found*** | | | | | |
| EP00129 | Obazoa;Opisthokonta;Nucletmycea;Fungi;"core Fur | EP00129 | Allomyces macrogynus 9915 | 1,30E-07 | 856 | XP 021881067.1 | hypothetical protein BCR41DRAFT 396657 [Lobosporangium transversale] | 1406 | 1,59E-113 | 387,882 | 47,83748362 |
| EP00376 | Sar;Alveolata;Myzozoa;"AC clade";Apicomplexa;"cc | EP00376 | Eimeria tenella 6378 | 1,30E-07 | 1457 | XP 013234111.1 | hypothetical protein, conserved [Eimeria tenella] | 1457 | 0 | 2862,4 | 100 |

| | | | | | | | | | | |
|---|---|---|---|---|---|---|---|---|---|---|
| EP00114 | Obazoa;Opisthokonta;Holozoa;Choanozoa;Metazoa | EP00114 Trichoplax sp H2 11650 | 1,30E-07 | 1114 | XP 002115837.1 | hypothetical protein TRIADDRAFT 64202 [Trichoplax adhaerens] | 978 | 0 | 1781,92 | 96,1682243 |
| EP00229 | Archaeplastida;Chloroplastida;Chlorophyta;Mamiel | EP00229 Micromonas commoda 4601 | 1,30E-07 | 1399 | XP 002503975.1 | predicted protein [Micromonas commoda] | 1399 | 0 | 2806,55 | 100 |
| EP00625 | Sar;Stramenopiles;Gyrista;Ochrophyta;Pelagophyce | EP00625 Aureococcus anophagefferens 6302 | 1,40E-07 | 728 | XP 009038384.1 | hypothetical protein AURANDRAFT 28308, partial [Aureococcus anophagefferens] | 728 | 0 | 1460,28 | 100 |
| EP00421 | Sar;Alveolata;Myzozoa;"DP clade";Dinoflagellata;D | EP00421 Protoceratium reticulatum 58032 | 1,40E-07 | 1162 | XP 002773527.1 | conserved hypothetical protein [Perkinsus marinus ATCC 50983] | 1211 | 0 | 750,355 | 49,95996797 |
| EP00160 | Obazoa;Opisthokonta;Nucletmycea;Rotosphaerida; | EP00160 Parvularia atlantis 21827 | 1,40E-07 | 103 | ***no hit found*** | | | | | |
| EP00377 | Sar;Alveolata;Myzozoa;"AC clade";Apicomplexa;"cc | EP00377 Hammondia hammondi 2390 | 1,40E-07 | 2301 | XP 018637885.1 | surface antigen repeat-containing protein [Toxoplasma gondii ME49] | 2315 | 0 | 3549,99 | 87,33850129 |
| EP00421 | Sar;Alveolata;Myzozoa;"DP clade";Dinoflagellata;D | EP00421 Protoceratium reticulatum 54119 | 1,40E-07 | 1176 | XP 002773527.1 | conserved hypothetical protein [Perkinsus marinus ATCC 50983] | 1211 | 0 | 752,281 | 48,43373494 |
| EP00237 | Archaeplastida;Chloroplastida;Chlorophyta;Pycnocc | EP00237 Pycnococcus provasolii 126 | 1,50E-07 | 1237 | XP 006837403.2 | protein transport protein SEC31 homolog B isoform X1 [Amborella trichopoda] | 1155 | 0 | 748,814 | 53,52564103 |
| EP00073 | Obazoa;Opisthokonta;Holozoa;Choanozoa;Metazoa | EP00073 Rattus norvegicus 30005 | 1,50E-07 | 1236 | XP 006250741.1 | PREDICTED: protein transport protein Sec31A isoform X3 [Rattus norvegicus] | 1236 | 0 | 2542,3 | 100 |
| EP00073 | Obazoa;Opisthokonta;Holozoa;Choanozoa;Metazoa | EP00073 Rattus norvegicus 30009 | 1,50E-07 | 1218 | XP 006250745.1 | PREDICTED: protein transport protein Sec31A isoform X7 [Rattus norvegicus] | 1218 | 0 | 2505,71 | 100 |
| EP00099 | Obazoa;Opisthokonta;Holozoa;Choanozoa;Metazoa | EP00099 Drosophila melanogaster 26490 | 1,50E-07 | 1207 | NP 727173.2 | uncharacterized protein Dmel CG9650, isoform I [Drosophila melanogaster] | 1207 | 0 | 2467,96 | 100 |
| EP00384 | Sar;Alveolata;Myzozoa;"AC clade";Apicomplexa;Cry | EP00384 Cryptosporidium muris 843 | 1,60E-07 | 1300 | XP 002139888.1 | hypothetical protein [Cryptosporidium muris RN66] | 1300 | 0 | 2624,74 | 100 |
| EP00073 | Obazoa;Opisthokonta;Holozoa;Choanozoa;Metazoa | EP00073 Rattus norvegicus 30007 | 1,60E-07 | 1231 | XP 006250743.1 | PREDICTED: protein transport protein Sec31A isoform X5 [Rattus norvegicus] | 1231 | 0 | 2532,29 | 100 |
| EP00073 | Obazoa;Opisthokonta;Holozoa;Choanozoa;Metazoa | EP00073 Rattus norvegicus 30002 | 1,60E-07 | 1262 | XP 006250738.1 | PREDICTED: protein transport protein Sec31A isoform X1 [Rattus norvegicus] >gi\|5643828 | 1262 | 0 | 2595,08 | 100 |
| EP00073 | Obazoa;Opisthokonta;Holozoa;Choanozoa;Metazoa | EP00073 Rattus norvegicus 30003 | 1,60E-07 | 1262 | XP 006250738.1 | PREDICTED: protein transport protein Sec31A isoform X1 [Rattus norvegicus] >gi\|5643828 | 1262 | 0 | 2595,08 | 100 |
| EP00503 | Sar;Stramenopiles;Bigyra;Opalozoa;Opalinata;Blast | EP00503 Blastocystis sp subtype4 5543 | 1,70E-07 | 857 | XP 014530137.1 | hypothetical protein JH06 2352 [Blastocystis sp. subtype 4] | 857 | 0 | 1760,35 | 100 |
| EP00403 | Sar;Alveolata;Myzozoa;"DP clade";Dinoflagellata;D | EP00403 Amphidinium massartii 42823 | 1,70E-07 | 78 | ***no hit found*** | | | | | |
| EP00678 | Discoba;Euglenozoa;Kinetoplastea;Metakinetoplast | EP00678 Phytomonas sp Hart1 2464 | 1,70E-07 | 1116 | XP 015655531.1 | putative protein transport protein Sec31 [Leptomonas pyrrhocoris] | 1178 | 0 | 804,668 | 53,34900118 |
| EP00044 | Obazoa;Opisthokonta;Holozoa;Choanozoa;Choanof | EP00044 Salpingoeca rosetta 10136 | 1,80E-07 | 1226 | XP 004997569.1 | TKL/MLK/MLK protein kinase [Salpingoeca rosetta] | 1226 | 0 | 2510,33 | 100 |
| EP00237 | Archaeplastida;Chloroplastida;Chlorophyta;Pycnocc | EP00237 Pycnococcus provasolii 1576 | 1,80E-07 | 353 | XP 023755508.1 | protein transport protein SEC16B homolog isoform X1 [Lactuca sativa] >gi\|1340423345\|r | 1236 | 2,90E-07 | 63,5438 | 56,47668394 |
| EP00376 | Sar;Alveolata;Myzozoa;"AC clade";Apicomplexa;"cc | EP00376 Eimeria tenella 3296 | 1,80E-07 | 1756 | XP 013231027.1 | plectin, putative [Eimeria tenella] | 1756 | 0 | 3513,39 | 100 |
| EP00482 | Sar;Rhizaria;Retaria;Foraminifera;Globothalamea;R | EP00482 Globobulimina sp turgida-auriculata 2763 | 1,90E-07 | 1330 | ***no hit found*** | | | | | |
| EP00044 | Obazoa;Opisthokonta;Holozoa;Choanozoa;Choanof | EP00044 Salpingoeca rosetta 5541 | 1,90E-07 | 764 | XP 004992974.1 | hypothetical protein PTSG 06078 [Salpingoeca rosetta] | 764 | 0 | 1559,66 | 100 |
| EP00388 | Sar;Alveolata;Myzozoa;"AC clade";Colpodellida;Col | EP00388 Colpodella angusta 8234 | 1,90E-07 | 108 | XP 024752873.1 | SEC16 protein [Trichoderma citrinoviride] | 1850 | 6,38E-06 | 53,5286 | 60,6741573 |
| EP00698 | Malawimonadidae;Gefionella;Gefionella okellyi;stra | EP00698 Gefionella okellyi 4125 | 2,00E-07 | 1141 | XP 006482944.1 | protein transport protein SEC31 homolog B isoform X1 [Citrus sinensis] >gi\|1350271342\|r | 1120 | 0 | 581,252 | 47,41235392 |
| EP00698 | Malawimonadidae;Gefionella;Gefionella okellyi;stra | EP00698 Gefionella okellyi 4126 | 2,00E-07 | 1141 | XP 006482944.1 | protein transport protein SEC31 homolog B isoform X1 [Citrus sinensis] >gi\|1350271342\|r | 1120 | 0 | 581,252 | 47,41235392 |
| EP00339 | Sar;Alveolata;Ciliophora;Intramacronucleata;"CON | EP00339 Tiarina fusus 25600 | 2,00E-07 | 1111 | XP 020427011.1 | WD40 repeat-containing protein [Heterostelium album PN500] | 836 | 1,60E-140 | 454,136 | 42,75534442 |
| EP00620 | Sar;Stramenopiles;Gyrista;Ochrophyta;Dictyochoph | EP00620 Florenciella sp RCC1587 2595 | 2,10E-07 | 1071 | XP 009038384.1 | hypothetical protein AURANDRAFT 28308, partial [Aureococcus anophagefferens] | 728 | 0 | 665,226 | 59,65909091 |
| EP00044 | Obazoa;Opisthokonta;Holozoa;Choanozoa;Choanof | EP00044 Salpingoeca rosetta 5384 | 2,10E-07 | 682 | XP 004992817.1 | Glrp1 protein [Salpingoeca rosetta] | 682 | 0 | 1363,59 | 100 |
| EP00013 | Amoebozoa;Discosea;Flabellinia;Stygamoeba;Styga | EP00013 Stygamoeba regulata 11474 | 2,10E-07 | 1170 | XP 019200437.1 | PREDICTED: ultraviolet-B receptor UVR8-like [Ipomoea nil] >gi\|1109251305\|ref\|XP 0192 | 442 | 1,43E-34 | 149,058 | 41,64456233 |
| EP00198 | Archaeplastida;Chloroplastida;Chlorophyta;"core ch | EP00198 Chlamydomonas reinhardtii 11769 | 2,20E-07 | 1430 | XP 001701106.1 | predicted protein [Chlamydomonas reinhardtii] | 1430 | 0 | 2834,67 | 100 |
| EP00385 | Sar;Alveolata;Myzozoa;"AC clade";Apicomplexa;Cry | EP00385 Cryptosporidium parvum 2170 | 2,20E-07 | 1205 | XP 627059.1 | coiled coil protein [Cryptosporidium parvum Iowa II] | 1205 | 0 | 2407,87 | 100 |
| EP00017 | Amoebozoa;Evosea;Archamoebea;Entamoebidae;E | EP00017 Entamoeba invadens 7518 | 2,30E-07 | 103 | XP 004257663.1 | hypothetical protein EIN 359850 [Entamoeba invadens IP1] | 103 | 6,05E-68 | 207,994 | 100 |
| EP00429 | Sar;Alveolata;Myzozoa;"DP clade";Dinoflagellata;D | EP00429 Kryptoperidinium foliaceum 74534 | 2,30E-07 | 838 | XP 002773527.1 | conserved hypothetical protein [Perkinsus marinus ATCC 50983] | 1211 | 8,83E-77 | 281,952 | 60,37735849 |
| EP00019 | Amoebozoa;Evosea;Cutosea;Armaparvus;Armaparv | EP00019 Armaparvus languidus 2439 | 2,30E-07 | 314 | ***no hit found*** | | | | | |
| EP00308 | Haptista;Haptophyta;Prymnesiophyceae;Calcihapto | EP00308 Calcidiscus leptoporus 9042 | 2,40E-07 | 444 | ***no hit found*** | | | | | |
| EP00328 | Haptista;Haptophyta;Prymnesiophyceae;Prymnesia | EP00328 Prymnesium polylepis 23092 | 2,40E-07 | 219 | ***no hit found*** | | | | | |
| EP00615 | Sar;Stramenopiles;Gyrista;Ochrophyta;Dictyochoph | EP00615 Pteridomonas danica 10108 | 2,40E-07 | 650 | XP 022105118.1 | replication protein A 70 kDa DNA-binding subunit-like isoform X3 [Acanthaster planci] | 625 | 2,88E-118 | 375,555 | 47,81954887 |
| EP00321 | Haptista;Haptophyta;Prymnesiophyceae;Phaeocyst | EP00321 Phaeocystis sp CCMP2710 3402 | 2,40E-07 | 281 | XP 005788197.1 | secretory protein 31 [Emiliania huxleyi CCMP1516] | 1166 | 1,13E-74 | 257,299 | 61,41975309 |
| EP00317 | Haptista;Haptophyta;Prymnesiophyceae;Phaeocyst | EP00317 Phaeocystis antarctica 36702 | 2,60E-07 | 141 | ***no hit found*** | | | | | |
| EP00279 | Cryptista;Cryptophyceae;Cryptomonadales;"lineage | EP00279 Guillardia theta 509 | 2,60E-07 | 649 | XP 005818228.1 | hypothetical protein GUITHDRAFT 122555 [Guillardia theta CCMP2712] | 649 | 0 | 1338,55 | 100 |
| EP00129 | Obazoa;Opisthokonta;Nucletmycea;Fungi;"core Fur | EP00129 Allomyces macrogynus 13604 | 2,60E-07 | 1270 | XP 016612805.1 | hypothetical protein SPPG 00469 [Spizellomyces punctatus DAOM BR117] | 1378 | 2,11E-174 | 563,533 | 49,8210451 |
| EP00057 | Obazoa;Opisthokonta;Holozoa;Choanozoa;Metazoa | EP00057 Strongylocentrotus purpuratus 12990 | 2,60E-07 | 990 | XP 011667464.1 | PREDICTED: mucin-2-like [Strongylocentrotus purpuratus] | 990 | 0 | 1828,91 | 100 |
| EP00090 | Obazoa;Opisthokonta;Holozoa;Choanozoa;Metazoa | EP00090 Calanus glacialis 44162 | 2,60E-07 | 882 | XP 023336929.1 | protein transport protein Sec31A-like [Eurytemora affinis] | 1195 | 0 | 975,696 | 64,83870968 |
| EP00620 | Sar;Stramenopiles;Gyrista;Ochrophyta;Dictyochoph | EP00620 Florenciella sp RCC1587 9968 | 2,80E-07 | 643 | XP 002289284.1 | probable eh domain-containing protein, partial [Thalassiosira pseudonana CCMP1335] | 382 | 0 | 551,592 | 66,66666667 |
| EP00038 | Obazoa;Opisthokonta;Holozoa;Choanozoa;Choanof | EP00038 Savillea parva 129 | 2,80E-07 | 1230 | XP 005175847.2 | PREDICTED: mucin-17-like [Musca domestica] | 1113 | 5,60E-25 | 124,79 | 61,20689655 |
| EP00376 | Sar;Alveolata;Myzozoa;"AC clade";Apicomplexa;"cc | EP00376 Eimeria tenella 754 | 2,90E-07 | 648 | XP 013228480.1 | hypothetical protein, conserved [Eimeria tenella] | 648 | 0 | 1227,62 | 100 |
| EP00365 | Sar;Alveolata;Myzozoa;"AC clade";Apicomplexa;"cc | EP00365 Plasmodium falciparum 536 | 2,90E-07 | 1544 | XP 001347889.2 | conserved Plasmodium protein, unknown function [Plasmodium falciparum 3D7] | 1544 | 0 | 2929,04 | 100 |
| EP00376 | Sar;Alveolata;Myzozoa;"AC clade";Apicomplexa;"cc | EP00376 Eimeria tenella 407 | 3,00E-07 | 2372 | XP 013228133.1 | hypothetical protein, conserved [Eimeria tenella] | 2372 | 0 | 4655,12 | 100 |
| EP00068 | Obazoa;Opisthokonta;Holozoa;Choanozoa;Metazoa | EP00068 Latimeria chalumnae 13755 | 3,20E-07 | 1170 | XP 006006621.1 | PREDICTED: protein transport protein Sec31A [Latimeria chalumnae] | 1170 | 0 | 2410,95 | 100 |
| EP00295 | Cryptista;Cryptophyceae;Goniomonas;"ma G clade" | EP00295 Goniomonas pacifica 33487 | 3,40E-07 | 340 | XP 005831033.1 | secretory protein Sec31 [Guillardia theta CCMP2712] | 1081 | 2,73E-57 | 210,305 | 48,04270463 |
| EP00376 | Sar;Alveolata;Myzozoa;"AC clade";Apicomplexa;"cc | EP00376 Eimeria tenella 2467 | 3,40E-07 | 522 | XP 013230194.1 | hypothetical protein, conserved, partial [Eimeria tenella] | 522 | 0 | 976,467 | 100 |
| EP00203 | Archaeplastida;Chloroplastida;Chlorophyta;"core ch | EP00203 Chlorella variabilis 9650 | 3,40E-07 | 563 | XP 005852196.1 | hypothetical protein CHLNCDRAFT 133410 [Chlorella variabilis] | 563 | 0 | 1089,33 | 100 |
| EP00376 | Sar;Alveolata;Myzozoa;"AC clade";Apicomplexa;"cc | EP00376 Eimeria tenella 5581 | 3,60E-07 | 2240 | XP 013233314.1 | hypothetical protein ETH 00033395 [Eimeria tenella] | 2240 | 0 | 4362,37 | 100 |
| EP00752 | Sar;Stramenopiles;Gyrista;Ochrophyta;"PX clade";P | EP00752 Nemacystus decipiens 8879 | 3,70E-07 | 1318 | ***no hit found*** | | | | | |
| EP00397 | Sar;Alveolata;Myzozoa;"DP clade";Dinoflagellata;"c | EP00397 Hematodinium sp SG-2012 2351 | 3,80E-07 | 1041 | XP 002140912.1 | hypothetical protein [Cryptosporidium muris RN66] | 1225 | 5,45E-148 | 482,641 | 47,58064516 |
| EP00376 | Sar;Alveolata;Myzozoa;"AC clade";Apicomplexa;"cc | EP00376 Eimeria tenella 2339 | 3,90E-07 | 1836 | XP 013230066.1 | hypothetical protein ETH 00035195, partial [Eimeria tenella] | 1836 | 0 | 3659 | 100 |
| EP00553 | Sar;Stramenopiles;Gyrista;Ochrophyta;"BD clade";[ | EP00553 Chaetoceros curvisetus 10956 | 4,00E-07 | 98 | ***no hit found*** | | | | | |
| EP00228 | Archaeplastida;Chloroplastida;Chlorophyta;Mamiel | EP00228 Micromonas bravo 1698 | 4,00E-07 | 941 | XP 002503975.1 | predicted protein [Micromonas commoda] | 1399 | 0 | 1231,08 | 79,64135021 |
| EP00316 | Haptista;Haptophyta;Prymnesiophyceae;Calcihapto | EP00316 Scyphosphaera apsteinii 22964 | 4,00E-07 | 1083 | XP 005788197.1 | secretory protein 31 [Emiliania huxleyi CCMP1516] | 1166 | 0 | 867,84 | 62,36323851 |
| EP00283 | Cryptista;Cryptophyceae;Cryptomonadales;"lineage | EP00283 Hemiselmis rufescens 16197 | 4,10E-07 | 365 | XP 005831033.1 | secretory protein Sec31 [Guillardia theta CCMP2712] | 1081 | 1,23E-118 | 377,867 | 64,26116838 |
| EP00383 | Sar;Alveolata;Myzozoa;"AC clade";Apicomplexa;Cry | EP00383 Cryptosporidium hominis 1548 | 4,20E-07 | 1206 | XP 666359.1 | T10G3.5 [Cryptosporidium hominis TU502] | 1206 | 0 | 2415,57 | 100 |
| EP00445 | Sar;Alveolata;Myzozoa;"DP clade";Dinoflagellata;D | EP00445 Apocalathium hangoei 92790 | 4,80E-07 | 830 | XP 004995411.1 | hypothetical protein PTSG 03685 [Salpingoeca rosetta] | 254 | 3,60E-08 | 66,2402 | 96,07843137 |
| EP00293 | Cryptista;Cryptophyceae;Cryptomonadales;Proteon | EP00293 Proteomonas sulcata 13253 | 4,90E-07 | 621 | XP 005831033.1 | secretory protein Sec31 [Guillardia theta CCMP2712] | 1081 | 1,07E-149 | 469,544 | 74,09836066 |
| EP00376 | Sar;Alveolata;Myzozoa;"AC clade";Apicomplexa;"cc | EP00376 Eimeria tenella 1946 | 4,90E-07 | 1201 | XP 013229672.1 | ERCC4 domain-containing protein, related [Eimeria tenella] | 1201 | 0 | 2311,18 | 100 |
| EP00376 | Sar;Alveolata;Myzozoa;"AC clade";Apicomplexa;"cc | EP00376 Eimeria tenella 1225 | 5,00E-07 | 1743 | XP 013228951.1 | DNA-dependent RNA polymerase, putative [Eimeria tenella] | 1743 | 0 | 3466,4 | 100 |
| EP00019 | Amoebozoa;Evosea;Cutosea;Armaparvus;Armaparv | EP00019 Armaparvus languidus 1656 | 5,00E-07 | 299 | ***no hit found*** | | | | | |
| EP00636 | Sar;Stramenopiles;Gyrista;Ochrophyta;Pinguiophyc | EP00636 Phaeomonas parva 18559 | 5,10E-07 | 130 | XP 024584649.1 | Regucalcin gene promoter region-related protein (RGPR) [Plasmopara halstedii] | 2672 | 3,93E-08 | 60,8474 | 64,0776699 |
| EP00405 | Sar;Alveolata;Myzozoa;"DP clade";Dinoflagellata;D | EP00405 Crypthecodinium cohnii 33886 | 5,20E-07 | 956 | ***no hit found*** | | | | | |
| EP00670 | Discoba;Euglenozoa;Euglenida;Spirocuta;Euglenea;I | EP00670 Eutreptiella gymnastica 12268 | 5,20E-07 | 683 | ***no hit found*** | | | | | |
| EP00446 | Sar;Alveolata;Myzozoa;"DP clade";Dinoflagellata;D | EP00446 Apocalathium sp SHHI-4 11787 | 5,40E-07 | 230 | ***no hit found*** | | | | | |
| EP00515 | Sar;Stramenopiles;Bigyra;Sagenista;Labyrinthulomy | EP00515 Schizochytrium aggregatum 11338 | 5,50E-07 | 814 | XP 008865383.1 | hypothetical protein H310 03344 [Aphanomyces invadans] | 664 | 9,53E-18 | 99,3673 | 37,86666667 |
| EP00411 | Sar;Alveolata;Myzozoa;"DP clade";Dinoflagellata;D | EP00411 Alexandrium monilatum 1482 | 6,00E-07 | 184 | ***no hit found*** | | | | | |
| EP00482 | Sar;Rhizaria;Retaria;Foraminifera;Globothalamea;R | EP00482 Globobulimina sp turgida-auriculata 1740 | 6,00E-07 | 1538 | XP 023243444.1 | transmembrane protease serine 9-like [Centruroides sculpturatus] | 864 | 6,22E-17 | 98,5969 | 51,33333333 |
| EP00376 | Sar;Alveolata;Myzozoa;"AC clade";Apicomplexa;"cc | EP00376 Eimeria tenella 674 | 6,10E-07 | 885 | XP 013228400.1 | hypothetical protein, conserved [Eimeria tenella] | 885 | 0 | 1716,82 | 100 |
| EP00561 | Sar;Stramenopiles;Gyrista;Ochrophyta;"BD clade";[ | EP00561 Arcocellulus cornucervis 8765 | 6,10E-07 | 710 | XP 002286990.1 | predicted protein [Thalassiosira pseudonana CCMP1335] | 913 | 9,09E-94 | 320,857 | 54,9898167 |
| EP00478 | Sar;Rhizaria;Endomyxa;"zoorhiza";Filoreta;"Ftenera | EP00478 Filoreta tenera 8111 | 6,40E-07 | 176 | ***no hit found*** | | | | | |
| EP00619 | Sar;Stramenopiles;Gyrista;Ochrophyta;Dictyochoph | EP00619 Florenciella sp RCC1007 14439 | 6,40E-07 | 189 | XP 002183614.1 | predicted protein [Phaeodactylum tricornutum CCAP 1055/1] | 2083 | 7,87E-08 | 62,003 | 56,56565657 |
| EP00230 | Archaeplastida;Chloroplastida;Chlorophyta;Mamiel | EP00230 Micromonas polaris 4890 | 6,60E-07 | 1634 | XP 002503975.1 | predicted protein [Micromonas commoda] | 1399 | 0 | 1208,36 | 77,37603306 |
| EP00376 | Sar;Alveolata;Myzozoa;"AC clade";Apicomplexa;"cc | EP00376 Eimeria tenella 4032 | 6,60E-07 | 834 | XP 013231765.1 | hypothetical protein, conserved [Eimeria tenella] | 834 | 0 | 1608,58 | 100 |
| EP00520 | Sar;Stramenopiles;Gyrista;Ochrophyta;"BD clade";E | EP00520 Triparma pacifica 2314 | 6,80E-07 | 117 | XP 024584649.1 | Regucalcin gene promoter region-related protein (RGPR) [Plasmopara halstedii] | 2672 | 1,92E-08 | 61,2326 | 66,66666667 |
| EP00105 | Obazoa;Opisthokonta;Holozoa;Choanozoa;Metazoa | EP00105 Crassostrea gigas 15136 | 6,90E-07 | 1191 | XP 011432043.1 | PREDICTED: protein transport protein Sec31A isoform X2 [Crassostrea gigas] | 1191 | 0 | 2471,04 | 100 |
| EP00073 | Obazoa;Opisthokonta;Holozoa;Choanozoa;Metazoa | EP00073 Rattus norvegicus 30011 | 7,10E-07 | 1179 | XP 006250747.1 | PREDICTED: protein transport protein Sec31A isoform X9 [Rattus norvegicus] | 1179 | 0 | 2427,13 | 100 |
| EP00449 | Sar;Alveolata;Myzozoa;"DP clade";Dinoflagellata;D | EP00449 Zooxanthella nutricula 22667 | 7,10E-07 | 81 | XP 001615328.1 | liver stage antigen, putative [Plasmodium vivax] | 1507 | 1,19E-07 | 57,3806 | 62,5 |
| EP00105 | Obazoa;Opisthokonta;Holozoa;Choanozoa;Metazoa | EP00105 Crassostrea gigas 15135 | 7,20E-07 | 1211 | XP 011432042.1 | PREDICTED: protein transport protein Sec31A isoform X1 [Crassostrea gigas] | 1211 | 0 | 2512,26 | 100 |
| EP00024 | Amoebozoa;Evosea;Eumycetozoa;Dictyostelia;Dicty | EP00024 Dictyostelium purpureum 4182 | 7,20E-07 | 4177 | XP 003287034.1 | hypothetical protein DICPUDRAFT 151088 [Dictyostelium purpureum] | 4177 | 0 | 8499,8 | 100 |
| EP00227 | Archaeplastida;Chloroplastida;Chlorophyta;Mamiel | EP00227 Mantoniella beaufortii 4558 | 7,30E-07 | 929 | XP 002503975.1 | predicted protein [Micromonas commoda] | 1399 | 0 | 1168,68 | 81,49425287 |
| EP00073 | Obazoa;Opisthokonta;Holozoa;Choanozoa;Metazoa | EP00073 Rattus norvegicus 30008 | 7,40E-07 | 1223 | XP 006250744.1 | PREDICTED: protein transport protein Sec31A isoform X6 [Rattus norvegicus] | 1223 | 0 | 2515,34 | 100 |
| EP00107 | Obazoa;Opisthokonta;Holozoa;Choanozoa;Metazoa | EP00107 Lottia gigantea 17981 | 7,60E-07 | 914 | XP 009060088.1 | hypothetical protein LOTGIDRAFT 106381 [Lottia gigantea] | 914 | 0 | 1900,56 | 100 |
| EP00321 | Haptista;Haptophyta;Prymnesiophyceae;Phaeocyst | EP00321 Phaeocystis sp CCMP2710 64 | 7,60E-07 | 774 | XP 005778453.1 | hypothetical protein EMIHUDRAFT 469065 [Emiliania huxleyi CCMP1516] | 711 | 6,31E-134 | 422,935 | 55,88865096 |
| EP00637 | Sar;Stramenopiles;Gyrista;Ochrophyta;Pinguiophyc | EP00637 Pinguiococcus pyrenoidosus 6667 | 7,70E-07 | 1524 | XP 008609468.1 | hypothetical protein SDRG 05527 [Saprolegnia diclina VS20] | 1030 | 1,06E-99 | 355,525 | 49,45736434 |
| EP00139 | Obazoa;Opisthokonta;Nucletmycea;Fungi;"core Fur | EP00139 Candida albicans 4057 | 8,10E-07 | 1265 | XP 718807.2 | Pga63p [Candida albicans SC5314] | 1265 | 0 | 2566,19 | 100 |
| EP00021 | Amoebozoa;Evosea;Eumycetozoa;Dictyostelia;Acyto | EP00021 Heterostelium album 8117 | 8,30E-07 | 602 | XP 020434753.1 | DNAJ heat shock N-terminal domain-containing protein [Heterostelium album PN500] | 602 | 0 | 1229,93 | 100 |
| EP00402 | Sar;Alveolata;Myzozoa;"DP clade";Dinoflagellata;D | EP00402 Amphidinium carterae 1750 | 8,30E-07 | 930 | XP 020391963.1 | CCR4-NOT transcription complex subunit 3 isoform X2 [Rhincodon typus] | 526 | 3,39E-44 | 179,104 | 45,56962025 |
| EP00437 | Sar;Alveolata;Myzozoa;"DP clade";Dinoflagellata;D | EP00437 Effrenium voratum 18820 | 8,30E-07 | 338 | XP 004988696.1 | hypothetical protein PTSG 10731 [Salpingoeca rosetta] | 481 | 1,80E-08 | 66,6254 | 34,44444444 |
| EP00201 | Archaeplastida;Chloroplastida;Chlorophyta;"core ch | EP00201 Polytomella parva 15733 | 8,50E-07 | 546 | ***no hit found*** | | | | | |
| EP00376 | Sar;Alveolata;Myzozoa;"AC clade";Apicomplexa;"cc | EP00376 Eimeria tenella 3517 | 8,50E-07 | 1762 | XP 013231248.1 | hypothetical protein, conserved [Eimeria tenella] | 1762 | 0 | 3435,58 | 100 |
| EP00697 | Hemimastigophora;Spironemidae;Spironema;strain | EP00697 Spironema sp BW2 9046 | 8,50E-07 | 118 | ***no hit found*** | | | | | |
| EP00282 | Cryptista;Cryptophyceae;Cryptomonadales;"lineage | EP00282 Hemiselmis andersenii 30804 | 8,50E-07 | 1101 | XP 005831033.1 | secretory protein Sec31 [Guillardia theta CCMP2712] | 1081 | 0 | 1082,78 | 65,2688172 |
| EP00123 | Obazoa;Opisthokonta;Holozoa;Ichthyosporea;Ichthy | EP00123 Amoebidium parasiticum 17318 | 8,60E-07 | 894 | XP 028661194.1 | protein transport protein Sec31A isoform X9 [Erpetoichthys calabaricus] | 1094 | 0 | 614,379 | 50,21978022 |
| EP00440 | Sar;Alveolata;Myzozoa;"DP clade";Dinoflagellata;D | EP00440 Ansanella granifera 30666 | 8,70E-07 | 295 | XP 002773527.1 | conserved hypothetical protein [Perkinsus marinus ATCC 50983] | 1211 | 9,20E-64 | 227,254 | 49,8381877 |
| EP00023 | Amoebozoa;Evosea;Eumycetozoa;Dictyostelia;Dicty | EP00023 Dictyostelium discoideum 5933 | 8,70E-07 | 1398 | XP 640040.1 | hypothetical protein DDB G0282421 [Dictyostelium discoideum AX4] | 1398 | 0 | 2749,54 | 100 |
| EP00282 | Cryptista;Cryptophyceae;Cryptomonadales;"lineage | EP00282 Hemiselmis andersenii 15963 | 8,70E-07 | 1173 | XP 005831033.1 | secretory protein Sec31 [Guillardia theta CCMP2712] | 1081 | 0 | 1039,25 | 64,00437637 |
| EP00119 | Obazoa;Opisthokonta;Holozoa;Choanozoa;Metazoa | EP00119 Amphimedon queenslandica 22332 | 8,90E-07 | 433 | XP 019862751.1 | PREDICTED: protein transport protein Sec16B-like, partial [Amphimedon queenslandica] | 433 | 0 | 864,374 | 100 |
| EP00376 | Sar;Alveolata;Myzozoa;"AC clade";Apicomplexa;"cc | EP00376 Eimeria tenella 6284 | 8,90E-07 | 1074 | XP 013234017.1 | hypothetical protein, conserved [Eimeria tenella] | 1074 | 0 | 2109,34 | 100 |
| EP00237 | Archaeplastida;Chloroplastida;Chlorophyta;Pycnocc | EP00237 Pycnococcus provasolii 13372 | 9,00E-07 | 1013 | ***no hit found*** | | | | | |
| EP00754 | Discoba;Euglenozoa;Diplonemea;Diplonemidae;Rhy | EP00754 Rhynchopus humris 19015 | 9,20E-07 | 503 | XP 022095045.1 | ubiquitin carboxyl-terminal hydrolase MINDY-2-like isoform X2 [Acanthaster planci] | 453 | 3,38E-61 | 216,853 | 47,11111111 |
| EP00516 | Sar;Stramenopiles;Bigyra;Sagenista;Labyrinthulomy | EP00516 Thraustochytrida sp NY0313808BC1 2098 | 9,50E-07 | 1331 | XP 008873541.1 | hypothetical protein H310 09285 [Aphanomyces invadans] | 1217 | 1,90E-156 | 513,072 | 47,5433526 |
| EP00517 | Sar;Stramenopiles;Bigyra;Sagenista;Labyrinthulomy | EP00517 Thraustochytrida sp NY07348D 9631 | 9,50E-07 | 1331 | XP 008873541.1 | hypothetical protein H310 09285 [Aphanomyces invadans] | 1217 | 1,90E-156 | 513,072 | 47,5433526 |
| EP00023 | Amoebozoa;Evosea;Eumycetozoa;Dictyostelia;Dicty | EP00023 Dictyostelium discoideum 5978 | 9,50E-07 | 1046 | XP 640086.1 | hypothetical protein DDB G0282501 [Dictyostelium discoideum AX4] | 1046 | 0 | 2026,52 | 100 |
| EP00200 | Archaeplastida;Chloroplastida;Chlorophyta;"core ch | EP00200 Dunaliella tertiolecta 14038 | 9,50E-07 | 2466 | XP 019193568.1 | PREDICTED: uncharacterized protein LOC109187727 [Ipomoea nil] | 2400 | 7,35E-117 | 427,943 | 44,83430799 |
| EP00120 | Obazoa;Opisthokonta;Holozoa;Filasterea;Capsaspoi | EP00120 Capsaspora owczarzaki 5164 | 9,60E-07 | 1609 | XP 004349938.2 | hypothetical protein CAOG 01418 [Capsaspora owczarzaki ATCC 30864] | 1609 | 0 | 3270,72 | 100 |
| EP00348 | Sar;Alveolata;Ciliophora;Intramacronucleata;"SAL"; | EP00348 Stylonychia lemnae 8354 | 9,90E-07 | 795 | ***no hit found*** | | | | | |
| EP00668 | Discoba;Euglenozoa;Euglenida;Spirocuta;Euglenea;I | EP00668 Euglena longa 48486 | 1,00E-06 | 945 | XP 024373656.1 | uncharacterized protein LOC112281406 isoform X3 [Physcomitrella patens] | 1251 | 1,21E-10 | 77,411 | 40,12605042 |
| EP00754 | Discoba;Euglenozoa;Diplonemea;Diplonemidae;Rhy | EP00754 Rhynchopus humris 19017 | 1,10E-06 | 514 | XP 022095045.1 | ubiquitin carboxyl-terminal hydrolase MINDY-2-like isoform X2 [Acanthaster planci] | 453 | 4,67E-61 | 216,853 | 47,11111111 |
| EP00628 | Sar;Stramenopiles;Gyrista;Ochrophyta;Pelagophyce | EP00628 Pelagophyceae sp CCMP2097 21520 | 1,10E-06 | 569 | XP 002290923.1 | predicted protein [Thalassiosira pseudonana CCMP1335] | 1041 | 6,38E-95 | 322,013 | 50,91514143 |
| EP00376 | Sar;Alveolata;Myzozoa;"AC clade";Apicomplexa;"cc | EP00376 Eimeria tenella 889 | 1,10E-06 | 1585 | XP 013228615.1 | hypothetical protein, conserved [Eimeria tenella] | 1585 | 0 | 2967,95 | 100 |
| EP00554 | Sar;Stramenopiles;Gyrista;Ochrophyta;"BD clade";[ | EP00554 Chaetoceros debilis 13426 | 1,10E-06 | 623 | XP 002294075.1 | predicted protein [Thalassiosira pseudonana CCMP1335] | 636 | 8,37E-27 | 126,716 | 42,6056338 |
| EP00245 | Archaeplastida;Chloroplastida;Streptophyta;"core s | EP00245 Coleochaete scutata 12724 | 1,20E-06 | 239 | XP 024373339.1 | protein transport protein SEC16B homolog isoform X2 [Physcomitrella patens] >gi\|137389 | 1264 | 3,01E-09 | 67,781 | 46,37681159 |
| EP00233 | Archaeplastida;Chloroplastida;Palmophyllophyceae | EP00233 Prasinococcus capsulatus 4787 | 1,20E-06 | 1383 | XP 002952699.1 | hypothetical protein VOLCADRAFT 36748 [Volvox carteri f. nagariensis] | 78 | 1,55E-17 | 88,5817 | 55,26315789 |
| EP00285 | Cryptista;Cryptophyceae;Cryptomonadales;"lineage | EP00285 Hemiselmis virescens 4053 | 1,20E-06 | 440 | ***no hit found*** | | | | | |
| EP00085 | Obazoa;Opisthokonta;Holozoa;Choanozoa;Metazoa | EP00085 Daphnia pulex 28805 | 1,20E-06 | 97 | ***no hit found*** | | | | | |
| EP00443 | Sar;Alveolata;Myzozoa;"DP clade";Dinoflagellata;D | EP00443 Scrippsiella trochoidea 72630 | 1,30E-06 | 213 | ***no hit found*** | | | | | |
| EP00060 | Obazoa;Opisthokonta;Holozoa;Choanozoa;Metazoa | EP00060 Petromyzon marinus 17883 | 1,30E-06 | 1435 | XP 007903785.1 | PREDICTED: paraspeckle component 1 isoform X2 [Callorhinchus milii] | 568 | 5,20E-167 | 522,316 | 69,13875598 |
| EP00132 | Obazoa;Opisthokonta;Nucletmycea;Fungi;"core Fur | EP00132 Gonapodya prolifera 8778 | 1,30E-06 | 1276 | XP 016612805.1 | hypothetical protein SPPG 00469 [Spizellomyces punctatus DAOM BR117] | 1378 | 0 | 890,182 | 51,69616519 |
| EP00601 | Sar;Stramenopiles;Gyrista;Ochrophyta;"CS clade";C | EP00601 Ochromonadales sp CCMP2298 3071 | 1,30E-06 | 1273 | XP 012194097.1 | hypothetical protein SPRG 18883 [Saprolegnia parasitica CBS 223.65] | 685 | 2,10E-57 | 222,631 | 62,83783784 |
| EP00044 | Obazoa;Opisthokonta;Holozoa;Choanozoa;Choanof | EP00044 Salpingoeca rosetta 10559 | 1,30E-06 | 6003 | XP 004997992.1 | MlnB [Salpingoeca rosetta] | 6003 | 0 | 12257,8 | 100 |
| EP00290 | Cryptista;Cryptophyceae;Cryptomonadales;Baffinel | EP00290 Baffinella sp CCMP2293 44408 | 1,40E-06 | 955 | XP 005838119.1 | hypothetical protein GUITHDRAFT 150997 [Guillardia theta CCMP2712] | 207 | 1,74E-51 | 189,889 | 62,61261261 |
| EP00675 | Discoba;Euglenozoa;Kinetoplastea;Metakinetoplast | EP00675 Leptomonas pyrrhocoris 4027 | 1,40E-06 | 1178 | XP 015655531.1 | putative protein transport protein Sec31 [Leptomonas pyrrhocoris] | 1178 | 0 | 2412,88 | 100 |
| EP00142 | Obazoa;Opisthokonta;Nucletmycea;Fungi;"core Fur | EP00142 Kluyveromyces lactis 4117 | 1,40E-06 | 1231 | XP 455314.1 | uncharacterized protein KLLA0 F05159g [Kluyveromyces lactis] | 1231 | 0 | 2494,15 | 100 |
| EP00023 | Amoebozoa;Evosea;Eumycetozoa;Dictyostelia;Dicty | EP00023 Dictyostelium discoideum 12142 | 1,40E-06 | 687 | XP 646613.1 | hypothetical protein DDB G0270200 [Dictyostelium discoideum AX4] | 687 | 0 | 1395,18 | 100 |
| EP00367 | Sar;Alveolata;Myzozoa;"AC clade";Apicomplexa;"cc | EP00367 Plasmodium vivax 894 | 1,50E-06 | 1438 | XP 001612905.1 | hypothetical protein, conserved [Plasmodium vivax] | 1438 | 0 | 2978,73 | 100 |
| EP00434 | Sar;Alveolata;Myzozoa;"DP clade";Dinoflagellata;D | EP00434 Breviolum minutum 45957 | 1,50E-06 | 245 | XP 002773527.1 | conserved hypothetical protein [Perkinsus marinus ATCC 50983] | 1211 | 9,70E-72 | 247,669 | 55,18672199 |
| EP00044 | Obazoa;Opisthokonta;Holozoa;Choanozoa;Choanof | EP00044 Salpingoeca rosetta 9102 | 1,50E-06 | 548 | XP 004996535.1 | excision repair cross-complementing rodent repair deficiency [Salpingoeca rosetta] | 548 | 0 | 1113,98 | 100 |
| EP00098 | Obazoa;Opisthokonta;Holozoa;Choanozoa;Metazoa | EP00098 Tribolium castaneum 3041 | 1,60E-06 | 2075 | XP 008192913.1 | PREDICTED: uncharacterized protein LOC658865 isoform X2 [Tribolium castaneum] | 2075 | 0 | 4253,36 | 100 |
| EP00298 | Haptista;Centroplasthelida;Panacanthocystida;Acar | EP00298 Acanthocystis sp HF-20 9321 | 1,60E-06 | 380 | WP 131338334.1 | hypothetical protein [Kribbella soli] | 1332 | 4,80E-17 | 93,9745 | 47,09302326 |
| EP00007 | Amoebozoa;Discosea;Flabellinia;"core flabellinids"; | EP00007 Cunea sp BSH-02190019 10096 | 1,60E-06 | 1835 | XP 019963803.1 | PREDICTED: 5'-3' exoribonuclease 1 [Paralichthys olivaceus] | 1717 | 7,71E-173 | 580,867 | 49,89414255 |
| EP00608 | Sar;Stramenopiles;Gyrista;Ochrophyta;"CS clade";S | EP00608 Synchroma pusillum 4432 | 1,60E-06 | 61 | ***no hit found*** | | | | | |
| EP00203 | Archaeplastida;Chloroplastida;Chlorophyta;"core ch | EP00203 Chlorella variabilis 3630 | 1,60E-06 | 2798 | XP 005846176.1 | hypothetical protein CHLNCDRAFT 136183 [Chlorella variabilis] | 2798 | 0 | 5467,12 | 100 |
| EP00358 | Sar;Alveolata;Ciliophora;Postciliodesmatophora;He | EP00358 Blepharisma japonicum 7308 | 1,70E-06 | 1123 | XP 001440954.1 | hypothetical protein [Paramecium tetraurelia strain d4-2] | 988 | 3,75E-85 | 308,145 | 40,32496307 |
| EP00639 | Sar;Stramenopiles;Gyrista;Ochrophyta;Raphidophy | EP00639 Heterosigma akashiwo 6152 | 1,70E-06 | 175 | XP 002908536.1 | COPII coat assembly protein SEC16, putative [Phytophthora infestans T30-4] | 2641 | 1,49E-09 | 66,6254 | 49,25373134 |
| EP00008 | Amoebozoa;Discosea;Flabellinia;"core flabellinids"; | EP00008 Paramoeba atlantica 3417 | 1,80E-06 | 82 | ***no hit found*** | | | | | |
| EP00007 | Amoebozoa;Discosea;Flabellinia;"core flabellinids"; | EP00007 Cunea sp BSH-02190019 8557 | 1,80E-06 | 923 | XP 004358077.1 | CUE domain containing protein [Acanthamoeba castellanii str. Neff] | 777 | 7,94E-26 | 125,946 | 43,10344828 |
| EP00376 | Sar;Alveolata;Myzozoa;"AC clade";Apicomplexa;"cc | EP00376 Eimeria tenella 8125 | 1,80E-06 | 735 | XP 013235868.1 | hypothetical protein ETH 00010180 [Eimeria tenella] | 735 | 0 | 1430,62 | 100 |
| EP00050 | Obazoa;Opisthokonta;Holozoa;Choanozoa;Choanof | EP00050 Salpingoeca kvevrii 11404 | 1,80E-06 | 885 | XP 002602930.1 | hypothetical protein BRAFLDRAFT 62485 [Branchiostoma floridae] | 1113 | 0 | 581,252 | 48,13384813 |
| EP00743 | Sar;Alveolata;Colponemidae;"MPE2-45 clade";strai | EP00743 Colponemidia sp Colp-15 7110 | 1,80E-06 | 977 | ***no hit found*** | | | | | |
| EP00257 | Archaeplastida;Chloroplastida;Streptophyta;"core s | EP00257 Ricinus communis 3478 | 1,90E-06 | 911 | XP 002515595.2 | transcriptional corepressor LEUNIG isoform X2 [Ricinus communis] | 911 | 0 | 1866,28 | 100 |
| EP00044 | Obazoa;Opisthokonta;Holozoa;Choanozoa;Choanof | EP00044 Salpingoeca rosetta 283 | 1,90E-06 | 801 | XP 004987716.1 | hypothetical protein PTSG 11219 [Salpingoeca rosetta] | 801 | 0 | 1639,01 | 100 |
| EP00740 | Archaeplastida;Chloroplastida;Chlorophyta;Mamiel | EP00740 Mantoniella antarctica 17302 | 1,90E-06 | 1164 | XP 029645372.1 | putative uncharacterized protein DDB G0294196 [Octopus vulgaris] | 262 | 8,74E-20 | 100,908 | 46,26865672 |
| EP00299 | Haptista;Centroplasthelida;Pterocystida;Pterista;"P | EP00299 Pterocystis canadensis 8172 | 1,90E-06 | 506 | XP 014785182.1 | PREDICTED: presenilin-1-like isoform X3 [Octopus bimaculoides] | 507 | 3,91E-80 | 268,085 | 46,03174603 |
| EP00446 | Sar;Alveolata;Myzozoa;"DP clade";Dinoflagellata;D | EP00446 Apocalathium sp SHHI-4 67462 | 2,00E-06 | 111 | ***no hit found*** | | | | | |
| EP00376 | Sar;Alveolata;Myzozoa;"AC clade";Apicomplexa;"cc | EP00376 Eimeria tenella 813 | 2,00E-06 | 1721 | XP 013228539.1 | hypothetical protein, conserved [Eimeria tenella] | 1721 | 0 | 3458,31 | 100 |
| EP00407 | Sar;Alveolata;Myzozoa;"DP clade";Dinoflagellata;D | EP00407 Lingulodinium polyedra 5539 | 2,00E-06 | 112 | ***no hit found*** | | | | | |
| EP00044 | Obazoa;Opisthokonta;Holozoa;Choanozoa;Choanof | EP00044 Salpingoeca rosetta 3426 | 2,10E-06 | 4739 | XP 004990859.1 | plasminogen [Salpingoeca rosetta] | 4739 | 0 | 9504,79 | 100 |
| EP00124 | Obazoa;Opisthokonta;Holozoa;Ichthyosporea;Ichthy | EP00124 Ichthyophonus hoferi 2852 | 2,20E-06 | 1134 | XP 011674242.1 | PREDICTED: putative exonuclease GOR-like protein [Strongylocentrotus purpuratus] | 215 | 5,10E-53 | 195,282 | 47 |
| EP00746 | Sar;Alveolata;Myzozoa;"DP clade";Dinoflagellata;"I | EP00746 Dinoflagellata sp MGD 42848 | 2,30E-06 | 303 | ***no hit found*** | | | | | |
| EP00376 | Sar;Alveolata;Myzozoa;"AC clade";Apicomplexa;"cc | EP00376 Eimeria tenella 1944 | 2,40E-06 | 1035 | XP 013229670.1 | lysyl-tRNA synthetase, putative [Eimeria tenella] | 1035 | 0 | 2042,7 | 100 |
| EP00407 | Sar;Alveolata;Myzozoa;"DP clade";Dinoflagellata;D | EP00407 Lingulodinium polyedra 38793 | 2,40E-06 | 549 | ***no hit found*** | | | | | |
| EP00179 | Archaeplastida;"RR clade";Rhodophyta;Proteorhod | EP00179 Madagascaria erythrocladioides 14904 | 2,50E-06 | 400 | XP 005852426.1 | threonine synthase, partial [Nannochloropsis gaditana CCMP526] | 401 | 2,63E-38 | 152,14 | 55,37190083 |
| EP00770 | Metamonada;Preaxostyla;Oxymonadida;Monocerc | EP00770 Monocercomonoides exilis 15470 | 2,50E-06 | 3387 | ***no hit found*** | | | | | |
| EP00391 | Sar;Alveolata;Myzozoa;"DP clade";Dinoflagellata;"c | EP00391 Amoebophrya sp Ameob2 260 | 2,60E-06 | 1700 | XP 028527165.1 | protein transport protein SEC31, putative [Plasmodium gallinaceum] | 1329 | 4,27E-49 | 204,142 | 42,03389831 |
| EP00376 | Sar;Alveolata;Myzozoa;"AC clade";Apicomplexa;"cc | EP00376 Eimeria tenella 3899 | 2,60E-06 | 879 | XP 013231632.1 | hypothetical protein, conserved [Eimeria tenella] | 879 | 0 | 1728,38 | 100 |
| EP00053 | Obazoa;Opisthokonta;Holozoa;Choanozoa;Choanof | EP00053 Salpingoeca punica 9134 | 2,60E-06 | 834 | XP 001744209.1 | hypothetical protein, partial [Monosiga brevicollis MX1] | 3611 | 3,46E-19 | 105,531 | 49,55223881 |
| EP00575 | Sar;Stramenopiles;Gyrista;Ochrophyta;"BD clade";[ | EP00575 Skeletonema marinoi 21588 | 2,60E-06 | 1224 | XP 002296320.1 | predicted protein [Thalassiosira pseudonana CCMP1335] | 1033 | 7,39E-38 | 166,007 | 45,72490706 |
| EP00638 | Sar;Stramenopiles;Gyrista;Ochrophyta;Raphidophy | EP00638 Chattonella subsalsa 286 | 2,60E-06 | 952 | XP 008860831.1 | hypothetical protein H310 00028 [Aphanomyces invadans] | 707 | 6,30E-52 | 205,297 | 42,53393665 |

| | | | | | | | | | | |
|---|---|---|---|---|---|---|---|---|---|---|
| EP00044 | Obazoa;Opisthokonta;Holozoa;Choanozoa;Choanof | EP00044 Salpingoeca rosetta 10436 | 2,70E-06 | 3953 | XP 004997869.1 | hypothetical protein PTSG 01888 [Salpingoeca rosetta] | 3953 | 0 | 8151,96 | 100 |
| EP00531 | Sar;Stramenopiles;Gyrista;Ochrophyta;"BD clade";[ | EP00531 Pseudo-nitzschia arenysensis 615 | 2,70E-06 | 1058 | XP 002180851.1 | predicted protein, partial [Phaeodactylum tricornutum CCAP 1055/1] | 258 | 3,58E-138 | 425,631 | 88,96103896 |
| EP00339 | Sar;Alveolata;Ciliophora;Intramacronucleata;"CON: | EP00339 Tiarina fusus 8590 | 2,80E-06 | 893 | XP 020427011.1 | WD40 repeat-containing protein [Heterostelium album PN500] | 836 | 1,29E-124 | 405,986 | 43,58353511 |
| EP00302 | Haptista;Haptophyta;Pavlovales;Diacronema;strain | EP00302 Diacronema sp CCMP2436 34915 | 2,80E-06 | 448 | ***no hit found*** | | | | | |
| EP00017 | Amoebozoa;Evosea;Archamoebea;Entamoebidae;E | EP00017 Entamoeba invadens 4726 | 2,80E-06 | 888 | XP 004254871.1 | hypothetical protein EIN 222530 [Entamoeba invadens IP1] | 888 | 0 | 1832,38 | 100 |
| EP00196 | Archaeplastida;Chloroplastida;Chlorophyta;"core ch | EP00196 Chlamydomonas euryale 17850 | 2,90E-06 | 768 | ***no hit found*** | | | | | |
| EP00652 | Sar;Stramenopiles;Gyrista;Peronosporomycetes;"cc | EP00652 Aphanomyces astaci 9134 | 2,90E-06 | 1221 | XP 009829895.1 | hypothetical protein H257 06393 [Aphanomyces astaci] | 1221 | 0 | 2483,75 | 100 |
| EP00445 | Sar;Alveolata;Myzozoa;"DP clade";Dinoflagellata;D | EP00445 Apocalathium hangoei 93672 | 2,90E-06 | 476 | ***no hit found*** | | | | | |
| EP00770 | Metamonada;Preaxostyla;Oxymonadida;Monocerc | EP00770 Monocercomonoides exilis 16076 | 2,90E-06 | 1363 | XP 019928790.1 | PREDICTED: transcriptional repressor NF-X1 [Crassostrea gigas] >gi\|1139829209\|ref\|XP ( | 1347 | 4,83E-32 | 148,288 | 40 |
| EP00621 | Sar;Stramenopiles;Gyrista;Ochrophyta;Dictyochoph | EP00621 Florenciella sp RCC1693 4610 | 3,00E-06 | 569 | XP 002181151.1 | predicted protein [Phaeodactylum tricornutum CCAP 1055/1] | 1133 | 1,43E-54 | 210,305 | 45,16129032 |
| EP00727 | Amoebozoa;Evosea;Archamoebea;Mastigamoebida | EP00727 Mastigamoeba balamuthi 6837 | 3,10E-06 | 1235 | XP 020313548.1 | protein transport protein Sec31A isoform X4 [Oncorhynchus kisutch] | 1429 | 5,62E-116 | 404,831 | 45,80384226 |
| EP00102 | Obazoa;Opisthokonta;Holozoa;Choanozoa;Metazoa | EP00102 Acyrthosiphon pisum 3646 | 3,10E-06 | 557 | XP 001946376.2 | nucleolin isoform X2 [Acyrthosiphon pisum] | 557 | 0 | 1068,53 | 100 |
| EP00195 | Archaeplastida;Chloroplastida;Chlorophyta;"core ch | EP00195 Chlamydomonas chlamydogama 11445 | 3,20E-06 | 1238 | XP 002956039.1 | hypothetical protein VOLCADRAFT 107038 [Volvox carteri f. nagariensis] | 1711 | 4,87E-14 | 89,3521 | 58,65384615 |
| EP00445 | Sar;Alveolata;Myzozoa;"DP clade";Dinoflagellata;D | EP00445 Apocalathium hangoei 34368 | 3,20E-06 | 764 | ***no hit found*** | | | | | |
| EP00531 | Sar;Stramenopiles;Gyrista;Ochrophyta;"BD clade";[ | EP00531 Pseudo-nitzschia arenysensis 579 | 3,30E-06 | 1060 | XP 002180765.1 | predicted protein, partial [Phaeodactylum tricornutum CCAP 1055/1] | 165 | 2,18E-32 | 134,035 | 49,15254237 |
| EP00533 | Sar;Stramenopiles;Gyrista;Ochrophyta;"BD clade";[ | EP00533 Pseudo-nitzschia delicatissima 9458 | 3,40E-06 | 1137 | XP 002286990.1 | predicted protein [Thalassiosira pseudonana CCMP1335] | 913 | 1,62E-91 | 324,709 | 59,75609756 |
| EP00063 | Obazoa;Opisthokonta;Holozoa;Choanozoa;Metazoa | EP00063 Salmo salar 7736 | 3,40E-06 | 1230 | XP 013982571.1 | PREDICTED: protein transport protein Sec31A-like isoform X6 [Salmo salar] | 1230 | 0 | 2517,27 | 100 |
| EP00674 | Discoba;Euglenozoa;Kinetoplastea;Metakinetoplast | EP00674 Leishmania major 810 | 3,40E-06 | 1158 | XP 001681582.1 | putative protein transport protein Sec31 [Leishmania major strain Friedlin] | 1158 | 0 | 2354,33 | 100 |
| EP00325 | Haptista;Haptophyta;Prymnesiophyceae;Prymnesia | EP00325 Haptolina brevifila 6421 | 3,50E-06 | 373 | XP 005788197.1 | secretory protein 31 [Emiliania huxleyi CCMP1516] | 1166 | 2,10E-65 | 234,958 | 64,02266289 |
| EP00421 | Sar;Alveolata;Myzozoa;"DP clade";Dinoflagellata;D | EP00421 Protoceratium reticulatum 39100 | 3,50E-06 | 123 | ***no hit found*** | | | | | |
| EP00426 | Sar;Alveolata;Myzozoa;"DP clade";Dinoflagellata;D | EP00426 Karenia brevis 34913 | 3,60E-06 | 474 | WP 144226726.1 | YSIRK-type signal peptide-containing protein [Lactobacillus reuteri] | 1432 | 1,70E-19 | 103,219 | 45,80152672 |
| EP00674 | Discoba;Euglenozoa;Kinetoplastea;Metakinetoplast | EP00674 Leishmania major 1037 | 3,60E-06 | 475 | XP 001681811.1 | putative lectin [Leishmania major strain Friedlin] | 475 | 0 | 975,311 | 100 |
| EP00376 | Sar;Alveolata;Myzozoa;"AC clade";Apicomplexa;"cc | EP00376 Eimeria tenella 6246 | 3,60E-06 | 1808 | XP 013233979.1 | intracellular protease, putative [Eimeria tenella] | 1808 | 0 | 3546,9 | 100 |
| EP00547 | Sar;Stramenopiles;Gyrista;Ochrophyta;"BD clade";[ | EP00547 Thalassionema nitzschioides 9253 | 3,60E-06 | 908 | XP 002185914.1 | NF-X1-like transcription factor [Phaeodactylum tricornutum CCAP 1055/1] | 906 | 7,23E-24 | 119,783 | 54,08805031 |
| EP00437 | Sar;Alveolata;Myzozoa;"DP clade";Dinoflagellata;D | EP00437 Effrenium voratum 16390 | 3,70E-06 | 1148 | XP 002140912.1 | hypothetical protein [Cryptosporidium muris RN66] | 1225 | 1,20E-122 | 417,542 | 47,51552795 |
| EP00429 | Sar;Alveolata;Myzozoa;"DP clade";Dinoflagellata;D | EP00429 Kryptoperidinium foliaceum 67091 | 3,70E-06 | 762 | ***no hit found*** | | | | | |
| EP00200 | Archaeplastida;Chloroplastida;Chlorophyta;"core ch | EP00200 Dunaliella tertiolecta 8579 | 3,80E-06 | 993 | XP 002946679.1 | hypothetical protein VOLCADRAFT 86877 [Volvox carteri f. nagariensis] | 969 | 5,46E-110 | 373,629 | 51,49812734 |
| EP00218 | Archaeplastida;Chloroplastida;Chlorophyta;Mamiel | EP00218 Dolichomastix tenuilepis 1864 | 3,80E-06 | 108 | XP 002987428.1 | protein transport protein SEC16B homolog [Selaginella moellendorffii] | 1190 | 2,22E-06 | 55,0694 | 50 |
| EP00334 | Sar;Alveolata;Ciliophora;Intramacronucleata;"CON: | EP00334 Paramecium tetraurelia 13204 | 3,80E-06 | 729 | XP 001436193.1 | hypothetical protein [Paramecium tetraurelia strain d4-2] | 729 | 0 | 1453,73 | 100 |
| EP00376 | Sar;Alveolata;Myzozoa;"AC clade";Apicomplexa;"cc | EP00376 Eimeria tenella 2372 | 3,80E-06 | 743 | XP 013230099.1 | hypothetical protein, conserved [Eimeria tenella] | 743 | 0 | 1438,32 | 100 |
| EP00376 | Sar;Alveolata;Myzozoa;"AC clade";Apicomplexa;"cc | EP00376 Eimeria tenella 4391 | 3,90E-06 | 1540 | XP 013232124.1 | hypothetical protein, conserved [Eimeria tenella] | 1540 | 0 | 2998,38 | 100 |
| EP00279 | Cryptista;Cryptophyceae;Cryptomonadales;"lineage | EP00279 Guillardia theta 11771 | 4,10E-06 | 115 | XP 005829490.1 | hypothetical protein GUITHDRAFT 153547 [Guillardia theta CCMP2712] | 115 | 2,36E-73 | 222,631 | 100 |
| EP00317 | Haptista;Haptophyta;Prymnesiophyceae;Phaeocyst | EP00317 Phaeocystis antarctica 49993 | 4,10E-06 | 823 | XP 005778453.1 | hypothetical protein EMIHUDRAFT 469065 [Emiliania huxleyi CCMP1516] | 711 | 4,25E-132 | 419,853 | 55,26315789 |
| EP00013 | Amoebozoa;Discosea;Flabellinia;Stygamoeba;Styga | EP00013 Stygamoeba regulata 10991 | 4,20E-06 | 670 | XP 004335566.1 | hypothetical protein ACA1 020160, partial [Acanthamoeba castellanii str. Neff] | 853 | 1,73E-45 | 184,496 | 45,65916399 |
| EP00742 | Sar;Alveolata;Colponemidae;"MPE2-45 clade";strai | EP00742 Colponemidia sp Colp-10 7804 | 4,40E-06 | 812 | XP 002773527.1 | conserved hypothetical protein [Perkinsus marinus ATCC 50983] | 1211 | 0 | 611,298 | 51,23152709 |
| EP00092 | Obazoa;Opisthokonta;Holozoa;Choanozoa;Metazoa | EP00092 Neocalanus flemingeri 12535 | 4,50E-06 | 825 | XP 023336929.1 | protein transport protein Sec31A-like [Eurytemora affinis] | 1195 | 4,66E-111 | 376,711 | 49,75369458 |
| EP00024 | Amoebozoa;Evosea;Eumycetozoa;Dictyostelia;Dicty | EP00024 Dictyostelium purpureum 12179 | 4,60E-06 | 234 | XP 003295031.1 | hypothetical protein DICPUDRAFT 160153 [Dictyostelium purpureum] | 234 | 1,89E-164 | 462,611 | 100 |
| EP00059 | Obazoa;Opisthokonta;Holozoa;Choanozoa;Metazoa | EP00059 Ciona intestinalis 519 | 4,60E-06 | 1205 | NP 001071923.1 | zinc finger protein [Ciona intestinalis] | 1205 | 0 | 2518,42 | 100 |
| EP00044 | Obazoa;Opisthokonta;Holozoa;Choanozoa;Choanof | EP00044 Salpingoeca rosetta 6913 | 4,60E-06 | 1812 | XP 004994346.1 | hypothetical protein PTSG 05031 [Salpingoeca rosetta] | 1812 | 0 | 3691,35 | 100 |
| EP00135 | Obazoa;Opisthokonta;Nucletmycea;Fungi;"core Fur | EP00135 Aspergillus nidulans 6253 | 4,80E-06 | 1275 | XP 663861.1 | hypothetical protein AN6257.2 [Aspergillus nidulans FGSC A4] | 1275 | 0 | 2597 | 100 |
| EP00696 | Hemimastigophora;Spironemidae;Hemimastix;Hem | EP00696 Hemimastix kukwesjijk 13575 | 4,80E-06 | 537 | XP 024401968.1 | origin of replication complex subunit 2-like isoform X1 [Physcomitrella patens] | 418 | 1,92E-63 | 222,631 | 47,67801858 |
| EP00457 | Sar;Rhizaria;Cercozoa;"crown cercozoa";Imbricatea | EP00457 Paulinella chromatophora 1275 | 4,80E-06 | 935 | WP 083253388.1 | HTTM domain-containing protein [Algibacter aquaticus] | 527 | 1,68E-28 | 132,109 | 41,69960474 |
| EP00607 | Sar;Stramenopiles;Gyrista;Ochrophyta;"CS clade";C | EP00607 Mallomonas sp CCMP3275 5434 | 5,00E-06 | 1125 | XP 009515715.1 | hypothetical protein PHYSODRAFT 322106 [Phytophthora sojae] | 1262 | 0 | 632,484 | 49,65831435 |
| EP00005 | Amoebozoa;Discosea;Centramoebia;"core centram | EP00005 Dracoamoeba jomungandri 9845 | 5,20E-06 | 677 | XP 002597290.1 | hypothetical protein BRAFLDRAFT 203599 [Branchiostoma floridae] | 431 | 9,20E-57 | 208,764 | 45,25139665 |
| EP00658 | Telonemia;Telonema;strain P-2 | EP00658 Telonema sp P-2 14614 | 5,20E-06 | 299 | WP 141668271.1 | hypothetical protein, partial [Clostridioides difficile] | 106 | 3,65E-08 | 60,8474 | 53,96825397 |
| EP00059 | Obazoa;Opisthokonta;Holozoa;Choanozoa;Metazoa | EP00059 Ciona intestinalis 11998 | 5,20E-06 | 1201 | XP 018670239.1 | zinc finger protein isoform X1 [Ciona intestinalis] | 1201 | 0 | 2510,33 | 100 |
| EP00439 | Sar;Alveolata;Myzozoa;"DP clade";Dinoflagellata;D | EP00439 Symbiodinium sp Y106 63626 | 5,20E-06 | 714 | XP 021967544.1 | glycosyltransferase 25 family member-like [Folsomia candida] | 585 | 6,53E-23 | 114,775 | 44,9197861 |
| EP00054 | Obazoa;Opisthokonta;Holozoa;Choanozoa;Choanof | EP00054 Salpingoeca dolichothecata 27078 | 5,30E-06 | 311 | ***no hit found*** | | | | | |
| EP00376 | Sar;Alveolata;Myzozoa;"AC clade";Apicomplexa;"cc | EP00376 Eimeria tenella 6878 | 5,30E-06 | 716 | XP 013234614.1 | Wee kinase, putative, partial [Eimeria tenella] | 716 | 0 | 1378,62 | 100 |
| EP00022 | Amoebozoa;Evosea;Eumycetozoa;Dictyostelia;Acyt | EP00022 Cavenderia fasciculata 10980 | 5,40E-06 | 979 | XP 004366257.1 | hypothetical protein DFA 03847 [Cavenderia fasciculata] | 979 | 0 | 1991,85 | 100 |
| EP00376 | Sar;Alveolata;Myzozoa;"AC clade";Apicomplexa;"cc | EP00376 Eimeria tenella 2368 | 5,50E-06 | 3065 | XP 013230095.1 | hypothetical protein, conserved [Eimeria tenella] | 3065 | 0 | 6042,61 | 100 |
| EP00457 | Sar;Rhizaria;Cercozoa;"crown cercozoa";Imbricatea | EP00457 Paulinella chromatophora 937 | 5,80E-06 | 991 | WP 083253388.1 | HTTM domain-containing protein [Algibacter aquaticus] | 527 | 3,77E-28 | 131,339 | 41,69960474 |
| EP00044 | Obazoa;Opisthokonta;Holozoa;Choanozoa;Choanof | EP00044 Salpingoeca rosetta 2165 | 5,80E-06 | 1061 | XP 004989598.1 | hypothetical protein PTSG 09339 [Salpingoeca rosetta] | 1061 | 0 | 2185,99 | 100 |
| EP00244 | Archaeplastida;Chloroplastida;Streptophyta;"core s | EP00244 Chara vulgaris 1374 | 5,90E-06 | 121 | XP 024379150.1 | protein transport protein SEC31 homolog B-like isoform X1 [Physcomitrella patens] | 1179 | 8,43E-43 | 159,458 | 58,62068966 |
| EP00065 | Obazoa;Opisthokonta;Holozoa;Choanozoa;Metazoa | EP00065 Mola mola 22605 | 6,00E-06 | 239 | XP 022614901.1 | protein transport protein Sec16B [Seriola dumerili] | 828 | 5,43E-45 | 170,629 | 69,50672646 |
| EP00376 | Sar;Alveolata;Myzozoa;"AC clade";Apicomplexa;"cc | EP00376 Eimeria tenella 1301 | 6,00E-06 | 736 | XP 013229027.1 | hypothetical protein, conserved [Eimeria tenella] | 736 | 0 | 1407,89 | 100 |
| EP00531 | Sar;Stramenopiles;Gyrista;Ochrophyta;"BD clade";[ | EP00531 Pseudo-nitzschia arenysensis 631 | 6,20E-06 | 1069 | XP 002180765.1 | predicted protein, partial [Phaeodactylum tricornutum CCAP 1055/1] | 165 | 2,47E-32 | 134,035 | 49,15254237 |
| EP00237 | Archaeplastida;Chloroplastida;Chlorophyta;Pycnocc | EP00237 Pycnococcus provasolii 12569 | 6,20E-06 | 766 | WP 083296208.1 | endonuclease/exonuclease/phosphatase family protein [Duganella sp. HH101] | 317 | 2,32E-32 | 138,658 | 48,47457627 |
| EP00537 | Sar;Stramenopiles;Gyrista;Ochrophyta;"BD clade";[ | EP00537 Pseudo-nitzschia pungens 12536 | 6,30E-06 | 1164 | XP 002180639.1 | predicted protein [Phaeodactylum tricornutum CCAP 1055/1] | 460 | 5,42E-20 | 105,531 | 42,51968504 |
| EP00357 | Sar;Alveolata;Ciliophora;Intramacronucleata;Protoc | EP00357 Protocruzia adherens 16673 | 6,40E-06 | 699 | XP 002957446.1 | hypothetical protein VOLCADRAFT 98490 [Volvox carteri f. nagariensis] | 4518 | 3,26E-06 | 62,7734 | 31,39534884 |
| EP00597 | Sar;Stramenopiles;Gyrista;Ochrophyta;"CS clade";C | EP00597 Dinobryon sp UTEXLB2267 19409 | 6,70E-06 | 225 | XP 024584649.1 | Regucalcin gene promoter region-related protein (RGPR) [Plasmopara halstedii] | 2672 | 5,99E-09 | 66,2402 | 59,25925926 |
| EP00384 | Sar;Alveolata;Myzozoa;"AC clade";Apicomplexa;Cry | EP00384 Cryptosporidium muris 2064 | 6,80E-06 | 593 | XP 002141109.1 | hypothetical protein [Cryptosporidium muris RN66] | 593 | 0 | 1217,22 | 100 |
| EP00298 | Haptista;Centroplasthelida;Panacanthocystida;Acar | EP00298 Acanthocystis sp HF-20 22518 | 6,80E-06 | 754 | XP 005831033.1 | secretory protein Sec31 [Guillardia theta CCMP2712] | 1081 | 4,55E-124 | 406,757 | 44,59627329 |
| EP00326 | Haptista;Haptophyta;Prymnesiophyceae;Prymnesia | EP00326 Haptolina ericina 28609 | 7,10E-06 | 359 | XP 005788197.1 | secretory protein 31 [Emiliania huxleyi CCMP1516] | 1166 | 4,11E-59 | 216,468 | 59,88857939 |
| EP00104 | Obazoa;Opisthokonta;Holozoa;Choanozoa;Metazoa | EP00104 Lingula anatina 26106 | 7,10E-06 | 1378 | XP 013409116.1 | uncharacterized protein LOC106172777 [Lingula anatina] >gi\|919054328\|ref\|XP 013409 | 1378 | 0 | 2866,64 | 100 |
| EP00104 | Obazoa;Opisthokonta;Holozoa;Choanozoa;Metazoa | EP00104 Lingula anatina 26107 | 7,10E-06 | 1378 | XP 013409116.1 | uncharacterized protein LOC106172777 [Lingula anatina] >gi\|919054328\|ref\|XP 013409 | 1378 | 0 | 2866,64 | 100 |
| EP00104 | Obazoa;Opisthokonta;Holozoa;Choanozoa;Metazoa | EP00104 Lingula anatina 26108 | 7,10E-06 | 1378 | XP 013409116.1 | uncharacterized protein LOC106172777 [Lingula anatina] >gi\|919054328\|ref\|XP 013409 | 1378 | 0 | 2866,64 | 100 |
| EP00104 | Obazoa;Opisthokonta;Holozoa;Choanozoa;Metazoa | EP00104 Lingula anatina 26109 | 7,10E-06 | 1378 | XP 013409116.1 | uncharacterized protein LOC106172777 [Lingula anatina] >gi\|919054328\|ref\|XP 013409 | 1378 | 0 | 2866,64 | 100 |
| EP00104 | Obazoa;Opisthokonta;Holozoa;Choanozoa;Metazoa | EP00104 Lingula anatina 26110 | 7,10E-06 | 1378 | XP 013409116.1 | uncharacterized protein LOC106172777 [Lingula anatina] >gi\|919054328\|ref\|XP 013409 | 1378 | 0 | 2866,64 | 100 |
| EP00104 | Obazoa;Opisthokonta;Holozoa;Choanozoa;Metazoa | EP00104 Lingula anatina 26111 | 7,10E-06 | 1378 | XP 013409116.1 | uncharacterized protein LOC106172777 [Lingula anatina] >gi\|919054328\|ref\|XP 013409 | 1378 | 0 | 2866,64 | 100 |
| EP00104 | Obazoa;Opisthokonta;Holozoa;Choanozoa;Metazoa | EP00104 Lingula anatina 26112 | 7,10E-06 | 1378 | XP 013409116.1 | uncharacterized protein LOC106172777 [Lingula anatina] >gi\|919054328\|ref\|XP 013409 | 1378 | 0 | 2866,64 | 100 |
| EP00104 | Obazoa;Opisthokonta;Holozoa;Choanozoa;Metazoa | EP00104 Lingula anatina 26113 | 7,10E-06 | 1378 | XP 013409116.1 | uncharacterized protein LOC106172777 [Lingula anatina] >gi\|919054328\|ref\|XP 013409 | 1378 | 0 | 2866,64 | 100 |
| EP00104 | Obazoa;Opisthokonta;Holozoa;Choanozoa;Metazoa | EP00104 Lingula anatina 26114 | 7,10E-06 | 1378 | XP 013409116.1 | uncharacterized protein LOC106172777 [Lingula anatina] >gi\|919054328\|ref\|XP 013409 | 1378 | 0 | 2866,64 | 100 |
| EP00104 | Obazoa;Opisthokonta;Holozoa;Choanozoa;Metazoa | EP00104 Lingula anatina 26115 | 7,10E-06 | 1378 | XP 013409116.1 | uncharacterized protein LOC106172777 [Lingula anatina] >gi\|919054328\|ref\|XP 013409 | 1378 | 0 | 2866,64 | 100 |
| EP00376 | Sar;Alveolata;Myzozoa;"AC clade";Apicomplexa;"cc | EP00376 Eimeria tenella 761 | 7,20E-06 | 900 | XP 013228487.1 | hypothetical protein, conserved [Eimeria tenella] | 900 | 0 | 1712,97 | 100 |
| EP00490 | Sar;Rhizaria;Retaria;Foraminifera;Tubothalamea;M | EP00490 Sorites sp Unknown 17237 | 7,30E-06 | 394 | XP 002773527.1 | conserved hypothetical protein [Perkinsus marinus ATCC 50983] | 1211 | 1,77E-80 | 278,485 | 57,32899023 |
| EP00597 | Sar;Stramenopiles;Gyrista;Ochrophyta;"CS clade";C | EP00597 Dinobryon sp UTEXLB2267 6232 | 7,80E-06 | 1127 | XP 002290923.1 | predicted protein [Thalassiosira pseudonana CCMP1335] | 1041 | 0 | 672,929 | 47,90940767 |
| EP00237 | Archaeplastida;Chloroplastida;Chlorophyta;Pycnocc | EP00237 Pycnococcus provasolii 469 | 7,90E-06 | 553 | ***no hit found*** | | | | | |
| EP00200 | Archaeplastida;Chloroplastida;Chlorophyta;"core ch | EP00200 Dunaliella tertiolecta 15056 | 8,10E-06 | 1222 | WP 015169633.1 | tRNA lysidine(34) synthetase TilS [Synechococcus sp. PCC 7502] | 318 | 2,88E-40 | 162,54 | 56,33802817 |
| EP00025 | Amoebozoa;Evosea;Eumycetozoa;Dictyostelia;Dicty | EP00025 Tieghemostelium lacteum 8036 | 8,20E-06 | 478 | XP 012751089.1 | hypothetical protein SAMD00019534 092210 [Acytostelium subglobosum LB1] | 362 | 5,55E-110 | 339,347 | 59,16666667 |
| EP00418 | Sar;Alveolata;Myzozoa;"DP clade";Dinoflagellata;D | EP00418 Pyrodinium bahamense 95786 | 8,30E-06 | 479 | ***no hit found*** | | | | | |
| EP00369 | Sar;Alveolata;Myzozoa;"AC clade";Apicomplexa;"cc | EP00369 Babesia microti 2390 | 8,40E-06 | 498 | XP 021337669.1 | conserved Plasmodium protein, unknown function [Babesia microti strain RI] | 498 | 0 | 1023,08 | 100 |
| EP00002 | CRuMs;Collodictyonidae;Diphylleia;Diphylleia rotan | EP00002 Diphylleia rotans 14281 | 8,60E-06 | 665 | XP 012567035.1 | PREDICTED: protein transport protein Sec31A-like isoform X1 [Hydra vulgaris] | 1175 | 5,87E-56 | 216,853 | 40,16260163 |
| EP00051 | Obazoa;Opisthokonta;Holozoa;Choanozoa;Choanof | EP00051 Salpingoeca urceolata 664 | 8,70E-06 | 852 | XP 010769212.1 | PREDICTED: WD repeat-containing protein 47-like isoform X2 [Notothenia coriiceps] | 881 | 5,52E-74 | 270,011 | 44,26580922 |
| EP00200 | Archaeplastida;Chloroplastida;Chlorophyta;"core ch | EP00200 Dunaliella tertiolecta 18052 | 8,80E-06 | 552 | XP 029483841.1 | protein phosphatase methylesterase 1-like isoform X1 [Oncorhynchus nerka] | 447 | 1,49E-61 | 219,164 | 47,96573876 |
| EP00376 | Sar;Alveolata;Myzozoa;"AC clade";Apicomplexa;"cc | EP00376 Eimeria tenella 3428 | 8,80E-06 | 1962 | XP 013231159.1 | hypothetical protein, conserved [Eimeria tenella] | 1962 | 0 | 3725,64 | 100 |
| EP00438 | Sar;Alveolata;Myzozoa;"DP clade";Dinoflagellata;D | EP00438 Fugacium kawagutii 23184 | 8,90E-06 | 210 | XP 002773527.1 | conserved hypothetical protein [Perkinsus marinus ATCC 50983] | 1211 | 4,33E-46 | 172,94 | 48,95833333 |
| EP00021 | Amoebozoa;Evosea;Eumycetozoa;Dictyostelia;Acyt | EP00021 Heterostelium album 8661 | 9,20E-06 | 1195 | XP 020435297.1 | hypothetical protein PPL 03970 [Heterostelium album PN500] | 1195 | 0 | 2414,42 | 100 |
| EP00601 | Sar;Stramenopiles;Gyrista;Ochrophyta;"CS clade";C | EP00601 Ochromonadales sp CCMP2298 34516 | 9,30E-06 | 1559 | XP 002183894.1 | Mer3 meiotic cross-over helicase [Phaeodactylum tricornutum CCAP 1055/1] | 1710 | 1,34E-14 | 91,6633 | 43,23432343 |
| EP00376 | Sar;Alveolata;Myzozoa;"AC clade";Apicomplexa;"cc | EP00376 Eimeria tenella 3862 | 9,90E-06 | 2234 | XP 013231595.1 | hypothetical protein ETH 00006030 [Eimeria tenella] | 2234 | 0 | 4435,17 | 100 |
| EP00405 | Sar;Alveolata;Myzozoa;"DP clade";Dinoflagellata;D | EP00405 Crypthecodinium cohnii 42610 | 1,00E-05 | 387 | YP 009052475.1 | pp1ab [Ball python nidovirus 1] | 8108 | 5,36E-12 | 78,9518 | 43,1372549 |
| EP00289 | Cryptista;Cryptophyceae;Cryptomonadales;"lineage | EP00289 Rhodomonas salina 11612 | 1,00E-05 | 706 | XP 005831033.1 | secretory protein Sec31 [Guillardia theta CCMP2712] | 1081 | 1,35E-143 | 456,447 | 71,12299465 |
| EP00444 | Sar;Alveolata;Myzozoa;"DP clade";Dinoflagellata;D | EP00444 Apocalathium aciculiferum 54101 | 1,00E-05 | 730 | XP 002773527.1 | conserved hypothetical protein [Perkinsus marinus ATCC 50983] | 1211 | 1,72E-75 | 275,404 | 59,20577617 |
| EP00200 | Archaeplastida;Chloroplastida;Chlorophyta;"core ch | EP00200 Dunaliella tertiolecta 10264 | 1,10E-05 | 2675 | XP 001703665.1 | predicted protein, partial [Chlamydomonas reinhardtii] | 1776 | 9,93E-50 | 207,994 | 51,2516469 |
| EP00281 | Cryptista;Cryptophyceae;Cryptomonadales;"lineage | EP00281 Chroomonas sp CCMP1168 21003 | 1,10E-05 | 755 | XP 005831033.1 | secretory protein Sec31 [Guillardia theta CCMP2712] | 1081 | 1,16E-117 | 389,808 | 65,65656566 |
| EP00024 | Amoebozoa;Evosea;Eumycetozoa;Dictyostelia;Dicty | EP00024 Dictyostelium purpureum 1948 | 1,10E-05 | 321 | XP 003284800.1 | hypothetical protein DICPUDRAFT 75738 [Dictyostelium purpureum] | 321 | 0 | 642,499 | 100 |
| EP00187 | Archaeplastida;"RR clade";Rhodophyta;Proteorhod | EP00187 Rhodella violacea 10787 | 1,10E-05 | 1029 | XP 004303090.1 | PREDICTED: protein transport protein SEC31 homolog B isoform X2 [Fragaria vesca subsp. | 1111 | 3,01E-86 | 311,612 | 41,65085389 |
| EP00331 | Sar;Alveolata;Ciliophora;Intramacronucleata;"CON: | EP00331 Platyophrya macrostoma 20544 | 1,10E-05 | 1759 | XP 028883513.1 | putative regulator of transcription factor TFIID [Trypanosoma theileri] | 2342 | 0 | 1113,21 | 48,01306478 |
| EP00429 | Sar;Alveolata;Myzozoa;"DP clade";Dinoflagellata;D | EP00429 Kryptoperidinium foliaceum 133133 | 1,10E-05 | 161 | XP 002183614.1 | predicted protein [Phaeodactylum tricornutum CCAP 1055/1] | 2083 | 6,23E-23 | 104,375 | 61,64383562 |
| EP00152 | Obazoa;Opisthokonta;Nucletmycea;Fungi;"core Fur | EP00152 Rhizopus delemar 15023 | 1,20E-05 | 916 | XP 023468011.1 | hypothetical protein RHIMIDRAFT 236286 [Rhizopus microsporus ATCC 52813] | 1104 | 0 | 1400,57 | 84,30962343 |
| EP00141 | Obazoa;Opisthokonta;Nucletmycea;Fungi;"core Fur | EP00141 Yarrowia lipolytica 4982 | 1,20E-05 | 1184 | XP 504598.1 | YALI0E30635p [Yarrowia lipolytica CLIB122] | 1184 | 0 | 2399,78 | 100 |
| EP00218 | Archaeplastida;Chloroplastida;Chlorophyta;Mamiel | EP00218 Dolichomastix tenuilepis 812 | 1,20E-05 | 1026 | XP 001420597.1 | predicted protein, partial [Ostreococcus lucimarinus CCE9901] | 1077 | 0 | 788,104 | 57,4566474 |
| EP00300 | Haptista;Centroplasthelida;Pterocystida;Raphidista | EP00300 Choanocystis sp HF-7 23847 | 1,20E-05 | 715 | XP 005831033.1 | secretory protein Sec31 [Guillardia theta CCMP2712] | 1081 | 1,15E-131 | 425,631 | 42,74406332 |
| EP00376 | Sar;Alveolata;Myzozoa;"AC clade";Apicomplexa;"cc | EP00376 Eimeria tenella 5683 | 1,20E-05 | 1642 | XP 013233416.1 | hypothetical protein, conserved [Eimeria tenella] | 1642 | 0 | 3266,48 | 100 |
| EP00023 | Amoebozoa;Evosea;Eumycetozoa;Dictyostelia;Dicty | EP00023 Dictyostelium discoideum 8117 | 1,20E-05 | 541 | XP 642368.1 | ARF/SAR superfamily protein [Dictyostelium discoideum AX4] | 541 | 0 | 1063,91 | 100 |
| EP00183 | Archaeplastida;"RR clade";Rhodophyta;Proteorhod | EP00183 Erythrolobus madagascarensis 2340 | 1,20E-05 | 1674 | ***no hit found*** | | | | | |
| EP00037 | Obazoa;Opisthokonta;Holozoa;Choanozoa;Choanof | EP00037 Helgoeca nana 11773 | 1,20E-05 | 1185 | XP 001745056.1 | hypothetical protein [Monosiga brevicollis MX1] | 787 | 5,97E-35 | 155,606 | 49,13151365 |
| EP00407 | Sar;Alveolata;Myzozoa;"DP clade";Dinoflagellata;D | EP00407 Lingulodinium polyedra 106540 | 1,20E-05 | 490 | ***no hit found*** | | | | | |
| EP00059 | Obazoa;Opisthokonta;Holozoa;Choanozoa;Metazoa | EP00059 Ciona intestinalis 17578 | 1,30E-05 | 1170 | XP 026693232.1 | zinc finger protein isoform X3 [Ciona intestinalis] | 1170 | 0 | 2447,93 | 100 |
| EP00286 | Cryptista;Cryptophyceae;Cryptomonadales;"lineage | EP00286 Rhodomonas abbreviata 15221 | 1,30E-05 | 404 | XP 005838423.1 | hypothetical protein GUITHDRAFT 161523 [Guillardia theta CCMP2712] | 986 | 2,51E-17 | 95,1301 | 41,06463878 |
| EP00672 | Discoba;Euglenozoa;Kinetoplastea;Metakinetoplast | EP00672 Neobodo designis 329 | 1,30E-05 | 902 | XP 028882558.1 | putative protein transport protein Sec31 [Trypanosoma theileri] | 1294 | 2,35E-124 | 416,772 | 40,90287278 |
| EP00120 | Obazoa;Opisthokonta;Holozoa;Filasterea;Capsaspoi | EP00120 Capsaspora owczarzaki 1332 | 1,30E-05 | 2915 | XP 004344217.2 | SNF2 family helicase [Capsaspora owczarzaki ATCC 30864] | 2915 | 0 | 5933,21 | 100 |
| EP00376 | Sar;Alveolata;Myzozoa;"AC clade";Apicomplexa;"cc | EP00376 Eimeria tenella 7127 | 1,40E-05 | 1701 | XP 013234864.1 | cation-transporting ATPase, putative [Eimeria tenella] | 1701 | 0 | 3385,12 | 100 |
| EP00376 | Sar;Alveolata;Myzozoa;"AC clade";Apicomplexa;"cc | EP00376 Eimeria tenella 5602 | 1,40E-05 | 632 | XP 013233335.1 | hypothetical protein, conserved [Eimeria tenella] | 632 | 0 | 1268,06 | 100 |
| EP00321 | Haptista;Haptophyta;Prymnesiophyceae;Phaeocyst | EP00321 Phaeocystis sp CCMP2710 5383 | 1,40E-05 | 85 | ***no hit found*** | | | | | |
| EP00581 | Sar;Stramenopiles;Gyrista;Ochrophyta;"BD clade";[ | EP00581 Thalassiosira minuscula 19346 | 1,50E-05 | 1714 | XP 002294375.1 | predicted protein [Thalassiosira pseudonana CCMP1335] | 1576 | 6,35E-117 | 417,542 | 63,99437412 |
| EP00676 | Discoba;Euglenozoa;Kinetoplastea;Metakinetoplast | EP00676 Leptomonas seymouri 655 | 1,50E-05 | 1198 | XP 015655531.1 | putative protein transport protein Sec31 [Leptomonas pyrrhocoris] | 1178 | 0 | 1650,18 | 80,52588332 |
| EP00405 | Sar;Alveolata;Myzozoa;"DP clade";Dinoflagellata;D | EP00405 Crypthecodinium cohnii 41706 | 1,50E-05 | 540 | ***no hit found*** | | | | | |
| EP00257 | Archaeplastida;Chloroplastida;Streptophyta;"core s | EP00257 Ricinus communis 14771 | 1,50E-05 | 920 | XP 015572528.1 | transcriptional corepressor LEUNIG isoform X1 [Ricinus communis] | 920 | 0 | 1884,77 | 100 |
| EP00279 | Cryptista;Cryptophyceae;Cryptomonadales;"lineage | EP00279 Guillardia theta 14783 | 1,50E-05 | 603 | XP 005832502.1 | hypothetical protein GUITHDRAFT 108396 [Guillardia theta CCMP2712] | 603 | 0 | 1252,65 | 100 |
| EP00120 | Obazoa;Opisthokonta;Holozoa;Filasterea;Capsaspoi | EP00120 Capsaspora owczarzaki 3320 | 1,50E-05 | 2415 | XP 004347402.1 | hypothetical protein CAOG 04655 [Capsaspora owczarzaki ATCC 30864] | 2415 | 0 | 4887,01 | 100 |
| EP00578 | Sar;Stramenopiles;Gyrista;Ochrophyta;"BD clade";[ | EP00578 Thalassiosira sp NH16 2656 | 1,60E-05 | 486 | ***no hit found*** | | | | | |
| EP00357 | Sar;Alveolata;Ciliophora;Intramacronucleata;Protoc | EP00357 Protocruzia adherens 522 | 1,60E-05 | 1015 | ***no hit found*** | | | | | |
| EP00556 | Sar;Stramenopiles;Gyrista;Ochrophyta;"BD clade";[ | EP00556 Chaetoceros neogracilis 13252 | 1,60E-05 | 676 | XP 002289734.1 | predicted protein [Thalassiosira pseudonana CCMP1335] | 627 | 6,78E-12 | 80,1073 | 43,88489209 |
| EP00138 | Obazoa;Opisthokonta;Nucletmycea;Fungi;"core Fur | EP00138 Coccidioides posadasii 6257 | 1,70E-05 | 1237 | XP 003071256.1 | WD domain, G-beta repeat containing protein [Coccidioides posadasii C735 delta SOWgp] | 1237 | 0 | 2531,52 | 100 |
| EP00469 | Sar;Rhizaria;Cercozoa;Chlorarachnea;"core chlorara | EP00469 Lotharella globosa 14922 | 1,80E-05 | 900 | XP 004335566.1 | hypothetical protein ACA1 020160, partial [Acanthamoeba castellanii str. Neff] | 853 | 3,60E-33 | 149,443 | 50,40871935 |
| EP00595 | Sar;Stramenopiles;Gyrista;Ochrophyta;"CS clade";C | EP00595 Chromulina nebulosa 931 | 1,80E-05 | 773 | XP 008860830.1 | hypothetical protein H310 00027 [Aphanomyces invadans] | 256 | 5,99E-14 | 83,1889 | 38,9380531 |
| EP00044 | Obazoa;Opisthokonta;Holozoa;Choanozoa;Choanof | EP00044 Salpingoeca rosetta 9964 | 1,80E-05 | 749 | XP 004997397.1 | hypothetical protein PTSG 01422 [Salpingoeca rosetta] | 749 | 0 | 1529,23 | 100 |
| EP00633 | Sar;Stramenopiles;Gyrista;Ochrophyta;Pelagophyce | EP00633 Aureoumbra lagunensis 7037 | 1,80E-05 | 614 | WP 011677359.1 | hypothetical protein [Oenococcus oeni] | 430 | 1,01E-24 | 117,857 | 98,68421053 |
| EP00499 | Sar;Stramenopiles;Bigyra;Opalozoa;Bicosoecida;"cc | EP00499 Cafeteria sp CaronLabIsolate 11987 | 1,90E-05 | 60 | ***no hit found*** | | | | | |
| EP00011 | Amoebozoa;Discosea;Flabellinia;"core flabellinids"; | EP00011 Vannella sp DIVA3-517-6-12 13189 | 1,90E-05 | 1278 | XP 011265577.2 | arf-GAP domain and FG repeat-containing protein 1 isoform X1 [Camponotus floridanus] | 411 | 7,15E-33 | 143,665 | 41,21863799 |
| EP00137 | Obazoa;Opisthokonta;Nucletmycea;Fungi;"core Fur | EP00137 Aspergillus oryzae 6755 | 1,90E-05 | 1273 | XP 023091002.1 | unnamed protein product [Aspergillus oryzae RIB40] | 1291 | 0 | 2576,59 | 100 |
| EP00376 | Sar;Alveolata;Myzozoa;"AC clade";Apicomplexa;"cc | EP00376 Eimeria tenella 230 | 1,90E-05 | 854 | XP 013227956.1 | hypothetical protein, conserved [Eimeria tenella] | 854 | 0 | 1643,25 | 100 |
| EP00445 | Sar;Alveolata;Myzozoa;"DP clade";Dinoflagellata;D | EP00445 Apocalathium hangoei 28066 | 2,00E-05 | 876 | ***no hit found*** | | | | | |
| EP00584 | Sar;Stramenopiles;Gyrista;Ochrophyta;"BD clade";[ | EP00584 Thalassiosira punctigera 27612 | 2,00E-05 | 890 | XP 002286037.1 | predicted protein [Thalassiosira pseudonana CCMP1335] | 328 | 3,16E-26 | 121,324 | 49,68553459 |
| EP00656 | Telonemia;Telonema;"Tsubtile clade";Telonema sul | EP00656 Telonema subtile 18587 | 2,00E-05 | 1113 | XP 002313327.2 | protein transport protein SEC31 homolog B [Populus trichocarpa] | 1135 | 1,13E-134 | 446,817 | 43,84027188 |
| EP00428 | Sar;Alveolata;Myzozoa;"DP clade";Dinoflagellata;D | EP00428 Durinskia baltica 19654 | 2,00E-05 | 1696 | WP 032847157.1 | Ig-like domain-containing protein [Mannheimia granulomatis] | 2035 | 2,63E-10 | 77,7962 | 48,1865285 |
| EP00648 | Sar;Stramenopiles;Gyrista;Peronosporomycetes;"cc | EP00648 Phytopythium vexans 2072 | 2,10E-05 | 2129 | XP 008908864.1 | hypothetical protein PPTG 13179 [Phytophthora parasitica INRA-310] | 2114 | 0 | 2264,19 | 72,37778832 |
| EP00445 | Sar;Alveolata;Myzozoa;"DP clade";Dinoflagellata;D | EP00445 Apocalathium hangoei 92227 | 2,10E-05 | 733 | ***no hit found*** | | | | | |
| EP00656 | Telonemia;Telonema;"Tsubtile clade";Telonema sul | EP00656 Telonema subtile 18586 | 2,10E-05 | 1168 | XP 002313327.2 | protein transport protein SEC31 homolog B [Populus trichocarpa] | 1135 | 5,36E-148 | 484,182 | 43,48184818 |
| EP00405 | Sar;Alveolata;Myzozoa;"DP clade";Dinoflagellata;D | EP00405 Crypthecodinium cohnii 38573 | 2,10E-05 | 990 | ***no hit found*** | | | | | |
| EP00556 | Sar;Stramenopiles;Gyrista;Ochrophyta;"BD clade";[ | EP00556 Chaetoceros neogracilis 2663 | 2,10E-05 | 645 | XP 002289734.1 | predicted protein [Thalassiosira pseudonana CCMP1335] | 627 | 1,02E-11 | 79,337 | 43,88489209 |
| EP00541 | Sar;Stramenopiles;Gyrista;Ochrophyta;"BD clade";[ | EP00541 Cyclophora tenuis 456 | 2,10E-05 | 535 | XP 002181151.1 | predicted protein [Phaeodactylum tricornutum CCAP 1055/1] | 1133 | 1,19E-138 | 438,728 | 54,56171735 |
| EP00443 | Sar;Alveolata;Myzozoa;"DP clade";Dinoflagellata;D | EP00443 Scrippsiella trochoidea 62623 | 2,20E-05 | 880 | ***no hit found*** | | | | | |
| EP00331 | Sar;Alveolata;Ciliophora;Intramacronucleata;"CON: | EP00331 Platyophrya macrostoma 11581 | 2,20E-05 | 863 | XP 029229941.1 | transcriptional regulatory protein NOT1 [Trypanosoma conorhini] | 1202 | 1,17E-157 | 501,901 | 47,78869779 |
| EP00445 | Sar;Alveolata;Myzozoa;"DP clade";Dinoflagellata;D | EP00445 Apocalathium hangoei 43384 | 2,20E-05 | 1495 | XP 014299070.1 | PREDICTED: mucin-17 [Microplitis demolitor] | 5803 | 9,25E-26 | 128,642 | 96,77419355 |
| EP00593 | Sar;Stramenopiles;Gyrista;Ochrophyta;"BD clade";[ | EP00593 Proboscia inermis 8677 | 2,30E-05 | 64 | XP 002183614.1 | predicted protein [Phaeodactylum tricornutum CCAP 1055/1] | 2083 | 3,06E-08 | 58,151 | 66,66666667 |
| EP00616 | Sar;Stramenopiles;Gyrista;Ochrophyta;Dictyochoph | EP00616 Rhizochromulina sp CCMP1243 17949 | 2,30E-05 | 1262 | XP 024577888.1 | sec14 cytosolic [Plasmopara halstedii] | 645 | 2,61E-28 | 133,65 | 37,13080169 |
| EP00770 | Metamonada;Preaxostyla;Oxymonadida;Monocerc | EP00770 Monocercomonoides exilis 8739 | 2,30E-05 | 908 | XP 029906089.1 | ubiquitin carboxyl-terminal hydrolase 13 isoform X5 [Myripristis murdjan] >gi\|170878620 | 766 | 3,83E-20 | 107,457 | 52,65017668 |
| EP00634 | Sar;Stramenopiles;Gyrista;Ochrophyta;Pelagophyce | EP00634 Sargassococcus sp CCMP2135 15996 | 2,30E-05 | 834 | XP 009040700.1 | hypothetical protein AURANDRAFT 67110 [Aureococcus anophagefferens] | 578 | 9,09E-51 | 198,364 | 58,68263473 |
| EP00301 | Haptista;Centroplasthelida;Pterocystida;Raphidista | EP00301 Raphidiophrys heterophryoidea 5969 | 2,40E-05 | 738 | XP 028727297.1 | protein NLRC5 [Peromyscus leucopus] | 1839 | 7,41E-07 | 64,6994 | 39,79591837 |
| EP00521 | Sar;Stramenopiles;Gyrista;Ochrophyta;"BD clade";[ | EP00521 Asterionellopsis glacialis 21615 | 2,40E-05 | 443 | XP 002183614.1 | predicted protein [Phaeodactylum tricornutum CCAP 1055/1] | 2083 | 7,35E-28 | 128,257 | 66,25 |
| EP00023 | Amoebozoa;Evosea;Eumycetozoa;Dictyostelia;Dicty | EP00023 Dictyostelium discoideum 4765 | 2,40E-05 | 837 | XP 638336.1 | anaphase promoting complex subunit 4 [Dictyostelium discoideum AX4] | 837 | 0 | 1683,69 | 100 |
| EP00529 | Sar;Stramenopiles;Gyrista;Ochrophyta;"BD clade";[ | EP00529 Nitzschia sp RCC80 17957 | 2,40E-05 | 829 | XP 002180574.1 | predicted protein [Phaeodactylum tricornutum CCAP 1055/1] | 915 | 1,76E-117 | 387,497 | 56,01217656 |
| EP00160 | Obazoa;Opisthokonta;Nucletmycea;Rotosphaerida; | EP00160 Parvularia atlantis 21826 | 2,40E-05 | 167 | ***no hit found*** | | | | | |
| EP00147 | Obazoa;Opisthokonta;Nucletmycea;Fungi;"core Fur | EP00147 Laccaria bicolor 16641 | 2,40E-05 | 1437 | XP 001889788.1 | protein transport protein sec31 [Laccaria bicolor S238N-H82] | 1437 | 0 | 2910,17 | 100 |
| EP00559 | Sar;Stramenopiles;Gyrista;Ochrophyta;"BD clade";[ | EP00559 Dactyliosolen fragilissimus 8072 | 2,50E-05 | 554 | ***no hit found*** | | | | | |
| EP00241 | Archaeplastida;Chloroplastida;Chlorophyta;Pyramir | EP00241 Pyramimonas parkeae 14657 | 2,50E-05 | 1183 | XP 006837403.2 | protein transport protein SEC31 homolog B isoform X1 [Amborella trichopoda] | 1155 | 0 | 794,267 | 52,51677852 |
| EP00548 | Sar;Stramenopiles;Gyrista;Ochrophyta;"BD clade";[ | EP00548 Thalassiothrix antarctica 14912 | 2,50E-05 | 2322 | XP 009514630.1 | hypothetical protein PHYSODRAFT 467355 [Phytophthora sojae] | 2377 | 0 | 1135,55 | 46,43637954 |
| EP00044 | Obazoa;Opisthokonta;Holozoa;Choanozoa;Choanof | EP00044 Salpingoeca rosetta 954 | 2,60E-05 | 7071 | XP 004988387.1 | hypothetical protein PTSG 10337 [Salpingoeca rosetta] | 7071 | 0 | 14527,4 | 100 |
| EP00429 | Sar;Alveolata;Myzozoa;"DP clade";Dinoflagellata;D | EP00429 Kryptoperidinium foliaceum 46838 | 2,60E-05 | 73 | XP 021878887.1 | Sec23-binding domain of Sec16-domain-containing protein [Lobosporangium transversale | 1291 | 8,13E-06 | 51,6026 | 49,27536232 |
| EP00202 | Archaeplastida;Chloroplastida;Chlorophyta;"core ch | EP00202 Volvox carteri 9898 | 2,60E-05 | 2925 | XP 002955504.1 | hypothetical protein VOLCADRAFT 96436 [Volvox carteri f. nagariensis] | 2925 | 0 | 5787,61 | 100 |
| EP00591 | Sar;Stramenopiles;Gyrista;Ochrophyta;"BD clade";[ | EP00591 Stephanopyxis turris 13807 | 2,60E-05 | 674 | XP 008907363.1 | hypothetical protein, variant 1 [Phytophthora parasitica INRA-310] | 1206 | 4,47E-54 | 211,46 | 41,92634561 |
| EP00528 | Sar;Stramenopiles;Gyrista;Ochrophyta;"BD clade";[ | EP00528 Fragilariopsis cylindrus 12344 | 2,70E-05 | 1353 | ***no hit found*** | | | | | |
| EP00414 | Sar;Alveolata;Myzozoa;"DP clade";Dinoflagellata;D | EP00414 Alexandrium minutum 4683 | 2,70E-05 | 207 | XP 002771093.1 | hypothetical protein Pmar PMAR000901 [Perkinsus marinus ATCC 50983] | 173 | 1,04E-08 | 62,3882 | 41,61490683 |

| | | | | | | | | | | |
|---|---|---|---|---|---|---|---|---|---|---|
| EP00482 | Sar;Rhizaria;Retaria;Foraminifera;Globothalamea;R | EP00482 Globobulimina sp turgida-auriculata 56163 | 2,70E-05 | 428 | XP 010694716.1 | PREDICTED: protein transport protein SEC31 homolog B isoform X2 [Beta vulgaris subsp. v | 1146 | 7,33E-38 | 157,147 | 46,13259669 |
| EP00658 | Telonemia;Telonema;strain P-2 | EP00658 Telonema sp P-2 73618 | 2,80E-05 | 175 | WP 141668293.1 | hypothetical protein [Clostridioides difficile] | 62 | 7,99E-10 | 62,003 | 92,85714286 |
| EP00099 | Obazoa;Opisthokonta;Holozoa;Choanozoa;Metazoa | EP00099 Drosophila melanogaster 26459 | 2,80E-05 | 700 | NP 727103.1 | Cdc7 kinase, isoform A [Drosophila melanogaster] | 700 | 0 | 1462,97 | 100 |
| EP00023 | Amoebozoa;Evosea;Eumycetozoa;Dictyostelia;Dicty | EP00023 Dictyostelium discoideum 10187 | 2,80E-05 | 1916 | XP 644514.1 | hypothetical protein DDB G0273795 [Dictyostelium discoideum AX4] >gi\|66822763\|ref\|) | 1916 | 0 | 3809,61 | 100 |
| EP00023 | Amoebozoa;Evosea;Eumycetozoa;Dictyostelia;Dicty | EP00023 Dictyostelium discoideum 10398 | 2,80E-05 | 1916 | XP 644514.1 | hypothetical protein DDB G0273795 [Dictyostelium discoideum AX4] >gi\|66822763\|ref\|) | 1916 | 0 | 3809,61 | 100 |
| EP00376 | Sar;Alveolata;Myzozoa;"AC clade";Apicomplexa;"cc | EP00376 Eimeria tenella 1231 | 2,80E-05 | 1150 | XP 013228957.1 | Poly(A) ribonuclease subunit, related [Eimeria tenella] | 1150 | 0 | 2239,92 | 100 |
| EP00218 | Archaeplastida;Chloroplastida;Chlorophyta;Mamiel | EP00218 Dolichomastix tenuilepis 14312 | 2,90E-05 | 385 | ***no hit found*** | | | | | |
| EP00578 | Sar;Stramenopiles;Gyrista;Ochrophyta;"BD clade";[ | EP00578 Thalassiosira sp NH16 10859 | 2,90E-05 | 484 | ***no hit found*** | | | | | |
| EP00124 | Obazoa;Opisthokonta;Holozoa;Ichthyosporea;Ichthy | EP00124 Ichthyophonus hoferi 2597 | 2,90E-05 | 946 | XP 022104643.1 | protein transport protein Sec31A-like isoform X2 [Acanthaster planci] | 1192 | 7,06E-147 | 476,093 | 49,47267498 |
| EP00289 | Cryptista;Cryptophyceae;Cryptomonadales;"lineage | EP00289 Rhodomonas salina 18165 | 3,00E-05 | 702 | XP 005831033.1 | secretory protein Sec31 [Guillardia theta CCMP2712] | 1081 | 3,47E-143 | 455,292 | 67,63925729 |
| EP00044 | Obazoa;Opisthokonta;Holozoa;Choanozoa;Choanof | EP00044 Salpingoeca rosetta 8538 | 3,00E-05 | 1704 | XP 004995971.1 | hypothetical protein PTSG 12015 [Salpingoeca rosetta] | 1704 | 0 | 3501,45 | 100 |
| EP00408 | Sar;Alveolata;Myzozoa;"DP clade";Dinoflagellata;D | EP00408 Alexandrium catenella 65505 | 3,10E-05 | 752 | YP 009052475.1 | pp1ab [Ball python nidovirus 1] | 8108 | 1,98E-09 | 73,1738 | 51,57894737 |
| EP00755 | Discoba;Euglenozoa;Diplonemea;Diplonemidae;Sul | EP00755 Sulcionema specki 4398 | 3,10E-05 | 917 | XP 002675650.1 | predicted protein [Naegleria gruberi] | 422 | 1,74E-10 | 75,485 | 32,11009174 |
| EP00443 | Sar;Alveolata;Myzozoa;"DP clade";Dinoflagellata;D | EP00443 Scrippsiella trochoidea 42828 | 3,10E-05 | 1383 | XP 018984480.1 | hypothetical protein BABINDRAFT 37888 [Babjeviella inositovora NRRL Y-12698] | 561 | 2,12E-54 | 211,075 | 41,88034188 |
| EP00013 | Amoebozoa;Discosea;Flabellinia;Stygamoeba;Styga | EP00013 Stygamoeba regulata 23694 | 3,10E-05 | 959 | ***no hit found*** | | | | | |
| EP00529 | Sar;Stramenopiles;Gyrista;Ochrophyta;"BD clade";[ | EP00529 Nitzschia sp RCC80 31591 | 3,20E-05 | 362 | ***no hit found*** | | | | | |
| EP00429 | Sar;Alveolata;Myzozoa;"DP clade";Dinoflagellata;D | EP00429 Kryptoperidinium foliaceum 77568 | 3,20E-05 | 705 | XP 002773527.1 | conserved hypothetical protein [Perkinsus marinus ATCC 50983] | 1211 | 0 | 584,719 | 53,09491059 |
| EP00556 | Sar;Stramenopiles;Gyrista;Ochrophyta;"BD clade";[ | EP00556 Chaetoceros neogracilis 13781 | 3,20E-05 | 792 | XP 003084120.2 | K Homology domain, type 1 [Ostreococcus tauri] | 416 | 1,04E-16 | 94,3597 | 37,7245509 |
| EP00752 | Sar;Stramenopiles;Gyrista;Ochrophyta;"PX clade";P | EP00752 Nemacystus decipiens 4101 | 3,20E-05 | 1608 | XP 008870269.1 | GTP-binding protein LepA [Aphanomyces invadans] | 646 | 0 | 639,032 | 58,27922078 |
| EP00235 | Archaeplastida;Chloroplastida;Palmophyllophyceae | EP00235 Prasinoderma singulare 541 | 3,20E-05 | 990 | XP 002948555.1 | hypothetical protein VOLCADRAFT 88764 [Volvox carteri f. nagariensis] | 1482 | 2,21E-12 | 83,1889 | 49,05660377 |
| EP00120 | Obazoa;Opisthokonta;Holozoa;Filasterea;Capsaspor | EP00120 Capsaspora owczarzaki 6572 | 3,30E-05 | 2317 | XP 004364776.1 | hypothetical protein CAOG 01908 [Capsaspora owczarzaki ATCC 30864] | 2317 | 0 | 4787,25 | 100 |
| EP00405 | Sar;Alveolata;Myzozoa;"DP clade";Dinoflagellata;D | EP00405 Crypthecodinium cohnii 58023 | 3,30E-05 | 465 | WP 117275230.1 | DUF4116 domain-containing protein [Mycoplasma edwardii] | 690 | 3,69E-30 | 134,806 | 42,5 |
| EP00044 | Obazoa;Opisthokonta;Holozoa;Choanozoa;Choanof | EP00044 Salpingoeca rosetta 3451 | 3,40E-05 | 1172 | XP 004990884.1 | hypothetical protein PTSG 07385 [Salpingoeca rosetta] | 1172 | 0 | 2404,4 | 100 |
| EP00601 | Sar;Stramenopiles;Gyrista;Ochrophyta;"CS clade";C | EP00601 Ochromonadales sp CCMP2298 24370 | 3,50E-05 | 494 | WP 144226726.1 | YSIRK-type signal peptide-containing protein [Lactobacillus reuteri] | 1432 | 3,04E-40 | 166,007 | 64,63414634 |
| EP00099 | Obazoa;Opisthokonta;Holozoa;Choanozoa;Metazoa | EP00099 Drosophila melanogaster 10642 | 3,50E-05 | 699 | NP 001284950.1 | Cdc7 kinase, isoform B [Drosophila melanogaster] | 699 | 0 | 1459,89 | 100 |
| EP00358 | Sar;Alveolata;Ciliophora;Postciliodesmatophora;He | EP00358 Blepharisma japonicum 4220 | 3,50E-05 | 157 | ***no hit found*** | | | | | |
| EP00009 | Amoebozoa;Discosea;Flabellinia;"core flabellinids"; | EP00009 Paramoeba aestuarina 6759 | 3,50E-05 | 53 | ***no hit found*** | | | | | |
| EP00484 | Sar;Rhizaria;Retaria;Foraminifera;Globothalamea;R | EP00484 Ammonia sp Unknown 20149 | 3,50E-05 | 371 | XP 018637002.1 | Rab1 protein [Toxoplasma gondii ME49] | 216 | 4,76E-73 | 235,728 | 64,13043478 |
| EP00405 | Sar;Alveolata;Myzozoa;"DP clade";Dinoflagellata;D | EP00405 Crypthecodinium cohnii 10398 | 3,50E-05 | 1907 | ***no hit found*** | | | | | |
| EP00256 | Archaeplastida;Chloroplastida;Streptophyta;"core s | EP00256 Glycine max 44495 | 3,70E-05 | 897 | XP 006596119.1 | transcriptional corepressor LEUNIG isoform X2 [Glycine max] >gi\|1590600323\|ref\|XP 02 | 897 | 0 | 1848,56 | 100 |
| EP00425 | Sar;Alveolata;Myzozoa;"DP clade";Dinoflagellata;D | EP00425 Heterocapsa triquetra 22555 | 3,70E-05 | 659 | XP 002784138.1 | hypothetical protein Pmar PMAR003392 [Perkinsus marinus ATCC 50983] | 292 | 8,19E-29 | 127,102 | 46,5648855 |
| EP00376 | Sar;Alveolata;Myzozoa;"AC clade";Apicomplexa;"cc | EP00376 Eimeria tenella 4367 | 3,90E-05 | 1361 | XP 013232100.1 | endonuclease/exonuclease/phosphatase domain-containing protein, putative, partial [Eim | 1361 | 0 | 2711,41 | 100 |
| EP00301 | Haptista;Centroplasthelida;Pterocystida;Raphidista | EP00301 Raphidiophrys heterophryoidea 5971 | 3,90E-05 | 731 | XP 028727297.1 | protein NLRC5 [Peromyscus leucopus] | 1839 | 8,36E-07 | 64,6994 | 39,79591837 |
| EP00022 | Amoebozoa;Evosea;Eumycetozoa;Dictyostelia;Acyto | EP00022 Cavenderia fasciculata 6841 | 3,90E-05 | 734 | XP 004359330.1 | putative homeobox transcription factor [Cavenderia fasciculata] | 734 | 0 | 1515,36 | 100 |
| EP00052 | Obazoa;Opisthokonta;Holozoa;Choanozoa;Choanof | EP00052 Salpingoeca macrocollata 7687 | 3,90E-05 | 607 | XP 019856761.1 | PREDICTED: cap-specific mRNA (nucleoside-2'-O-)-methyltransferase 2-like isoform X1 [An | 939 | 2,52E-08 | 68,9366 | 33,73493976 |
| EP00160 | Obazoa;Opisthokonta;Nucletmycea;Rotosphaerida; | EP00160 Parvularia atlantis 20091 | 3,90E-05 | 99 | ***no hit found*** | | | | | |
| EP00674 | Discoba;Euglenozoa;Kinetoplastea;Metakinetoplast | EP00674 Leishmania major 4476 | 4,00E-05 | 1295 | XP 001685264.1 | kinetoplast-associated protein-like protein [Leishmania major strain Friedlin] | 1295 | 0 | 2450,24 | 100 |
| EP00099 | Obazoa;Opisthokonta;Holozoa;Choanozoa;Metazoa | EP00099 Drosophila melanogaster 19171 | 4,10E-05 | 1240 | NP 610414.1 | secretory 31, isoform A [Drosophila melanogaster] | 1240 | 0 | 2558,48 | 100 |
| EP00613 | Sar;Stramenopiles;Gyrista;Ochrophyta;Dictyochoph | EP00613 Pedinella sp CCMP2098 67614 | 4,10E-05 | 437 | XP 005819637.1 | hypothetical protein GUITHDRAFT 166656 [Guillardia theta CCMP2712] | 1207 | 2,94E-37 | 155,606 | 52,60115607 |
| EP00134 | Obazoa;Opisthokonta;Nucletmycea;Fungi;"core Fur | EP00134 Neurospora crassa 9357 | 4,10E-05 | 1452 | XP 964067.2 | hypothetical protein NCU09807 [Neurospora crassa OR74A] | 1452 | 0 | 3009,16 | 100 |
| EP00404 | Sar;Alveolata;Myzozoa;"DP clade";Dinoflagellata;D | EP00404 Azadinium spinosum 29382 | 4,10E-05 | 691 | XP 002773527.1 | conserved hypothetical protein [Perkinsus marinus ATCC 50983] | 1211 | 9,37E-175 | 540,806 | 57,73955774 |
| EP00752 | Sar;Stramenopiles;Gyrista;Ochrophyta;"PX clade";P | EP00752 Nemacystus decipiens 4100 | 4,20E-05 | 1609 | XP 008870269.1 | GTP-binding protein LepA [Aphanomyces invadans] | 646 | 0 | 639,032 | 58,27922078 |
| EP00203 | Archaeplastida;Chloroplastida;Chlorophyta;"core ch | EP00203 Chlorella variabilis 7690 | 4,20E-05 | 959 | XP 005850236.1 | hypothetical protein CHLNCDRAFT 141882 [Chlorella variabilis] | 959 | 0 | 1941,78 | 100 |
| EP00570 | Sar;Stramenopiles;Gyrista;Ochrophyta;"BD clade";[ | EP00570 Cyclotella meneghiniana 1317 | 4,30E-05 | 1344 | XP 002296589.1 | predicted protein [Thalassiosira pseudonana CCMP1335] | 2042 | 2,14E-19 | 107,071 | 57,92079208 |
| EP00200 | Archaeplastida;Chloroplastida;Chlorophyta;"core ch | EP00200 Dunaliella tertiolecta 19386 | 4,30E-05 | 990 | ***no hit found*** | | | | | |
| EP00648 | Sar;Stramenopiles;Gyrista;Peronosporomycetes;"cc | EP00648 Phytopythium vexans 10654 | 4,30E-05 | 1156 | XP 009520239.1 | hypothetical protein PHYSODRAFT 326004 [Phytophthora sojae] | 2070 | 0 | 1075,08 | 60,6557377 |
| EP00536 | Sar;Stramenopiles;Gyrista;Ochrophyta;"BD clade";[ | EP00536 Pseudo-nitzschia multiseries 10613 | 4,30E-05 | 678 | ***no hit found*** | | | | | |
| EP00132 | Obazoa;Opisthokonta;Nucletmycea;Fungi;"core Fur | EP00132 Gonapodya prolifera 2261 | 4,40E-05 | 1759 | ***no hit found*** | | | | | |
| EP00764 | Metamonada;"BF clade";Fornicata;Aduncisulcus;Ad | EP00764 Aduncisulcus paluster 5127 | 4,40E-05 | 745 | ***no hit found*** | | | | | |
| EP00110 | Obazoa;Opisthokonta;Holozoa;Choanozoa;Metazoa | EP00110 Nematostella vectensis 783 | 4,50E-05 | 222 | XP 001618184.1 | hypothetical protein NEMVEDRAFT v1g225411 [Nematostella vectensis] | 222 | 9,08E-163 | 457,218 | 100 |
| EP00148 | Obazoa;Opisthokonta;Nucletmycea;Fungi;"core Fur | EP00148 Cryptococcus gattii 3088 | 4,50E-05 | 1433 | XP 003196195.1 | Protein transport protein SEC31, putative [Cryptococcus gattii WM276] | 1433 | 0 | 2876,27 | 98,74389393 |
| EP00620 | Sar;Stramenopiles;Gyrista;Ochrophyta;Dictyochoph | EP00620 Florenciella sp RCC1587 7335 | 4,50E-05 | 1455 | XP 002186176.1 | predicted protein, partial [Phaeodactylum tricornutum CCAP 1055/1] | 225 | 2,56E-44 | 171,014 | 44,10480349 |
| EP00120 | Obazoa;Opisthokonta;Holozoa;Filasterea;Capsaspor | EP00120 Capsaspora owczarzaki 7887 | 4,60E-05 | 1860 | XP 011270068.1 | hypothetical protein CAOG 08486 [Capsaspora owczarzaki ATCC 30864] | 1860 | 0 | 3818,47 | 100 |
| EP00578 | Sar;Stramenopiles;Gyrista;Ochrophyta;"BD clade";[ | EP00578 Thalassiosira sp NH16 3042 | 4,60E-05 | 482 | XP 002295385.1 | hypothetical protein THAPS 264495, partial [Thalassiosira pseudonana CCMP1335] | 87 | 4,29E-14 | 77,7962 | 47,82608696 |
| EP00052 | Obazoa;Opisthokonta;Holozoa;Choanozoa;Choanof | EP00052 Salpingoeca macrocollata 23033 | 4,60E-05 | 125 | ***no hit found*** | | | | | |
| EP00523 | Sar;Stramenopiles;Gyrista;Ochrophyta;"BD clade";[ | EP00523 Amphiprora sp CCMP467 6621 | 4,80E-05 | 1106 | ***no hit found*** | | | | | |
| EP00044 | Obazoa;Opisthokonta;Holozoa;Choanozoa;Choanof | EP00044 Salpingoeca rosetta 3958 | 4,90E-05 | 901 | XP 004991391.1 | hypothetical protein PTSG 12612 [Salpingoeca rosetta] | 901 | 0 | 1833,15 | 100 |
| EP00240 | Archaeplastida;Chloroplastida;Chlorophyta;Pyramir | EP00240 Pyramimonas obovata 15070 | 4,90E-05 | 187 | XP 020107946.1 | protein transport protein SEC31 homolog B [Ananas comosus] | 1119 | 1,70E-45 | 170,244 | 48,29545455 |
| EP00041 | Obazoa;Opisthokonta;Holozoa;Choanozoa;Choanof | EP00041 Stephanoeca diplocostata 23037 | 5,00E-05 | 1039 | XP 007887151.1 | PREDICTED: protein transport protein Sec31A isoform X3 [Callorhinchus milii] | 1236 | 3,34E-52 | 211,46 | 40,27586207 |
| EP00531 | Sar;Stramenopiles;Gyrista;Ochrophyta;"BD clade";[ | EP00531 Pseudo-nitzschia arenysensis 551 | 5,00E-05 | 1057 | XP 002180851.1 | predicted protein, partial [Phaeodactylum tricornutum CCAP 1055/1] | 258 | 3,50E-138 | 425,631 | 88,92508143 |
| EP00125 | Obazoa;Opisthokonta;Holozoa;Ichthyosporea;Ichthy | EP00125 Sphaeroforma arctica 11457 | 5,30E-05 | 893 | XP 014154532.1 | hypothetical protein SARC 07021 [Sphaeroforma arctica JP610] | 893 | 0 | 1829,68 | 100 |
| EP00606 | Sar;Stramenopiles;Gyrista;Ochrophyta;"CS clade";C | EP00606 Chrysophyceae sp TOSAG23-5 1532 | 5,30E-05 | 678 | XP 009040700.1 | hypothetical protein AURANDRAFT 67110 [Aureococcus anophagefferens] | 578 | 1,28E-14 | 88,5817 | 37,1541502 |
| EP00403 | Sar;Alveolata;Myzozoa;"DP clade";Dinoflagellata;D | EP00403 Amphidinium massartii 20296 | 5,40E-05 | 726 | XP 005822890.1 | hypothetical protein GUITHDRAFT 55424, partial [Guillardia theta CCMP2712] | 373 | 1,76E-11 | 77,7962 | 34,24657534 |
| EP00200 | Archaeplastida;Chloroplastida;Chlorophyta;"core ch | EP00200 Dunaliella tertiolecta 17123 | 5,40E-05 | 1664 | ***no hit found*** | | | | | |
| EP00628 | Sar;Stramenopiles;Gyrista;Ochrophyta;Pelagophyce | EP00628 Pelagophyceae sp CCMP2097 11280 | 5,50E-05 | 672 | ***no hit found*** | | | | | |
| EP00376 | Sar;Alveolata;Myzozoa;"AC clade";Apicomplexa;"cc | EP00376 Eimeria tenella 6210 | 5,60E-05 | 1714 | XP 013233943.1 | hypothetical protein, conserved [Eimeria tenella] | 1714 | 0 | 3345,06 | 100 |
| EP00194 | Archaeplastida;Chloroplastida;Chlorophyta;"core ch | EP00194 Tetraselmis striata 22521 | 5,60E-05 | 85 | ***no hit found*** | | | | | |
| EP00401 | Sar;Alveolata;Myzozoa;"DP clade";Dinoflagellata;D | EP00401 Gymnodinium catenatum 8803 | 5,70E-05 | 291 | XP 002773527.1 | conserved hypothetical protein [Perkinsus marinus ATCC 50983] | 1211 | 5,07E-75 | 258,84 | 55,73248408 |
| EP00009 | Amoebozoa;Discosea;Flabellinia;"core flabellinids"; | EP00009 Paramoeba aestuarina 1351 | 5,70E-05 | 454 | XP 014775718.1 | PREDICTED: FK506-binding protein 15-like isoform X3 [Octopus bimaculoides] | 1017 | 1,92E-57 | 214,542 | 48,89867841 |
| EP00670 | Discoba;Euglenozoa;Euglenida;Spirocuta;Euglenea;I | EP00670 Eutreptiella gymnastica 15776 | 5,70E-05 | 1810 | XP 017251332.1 | PREDICTED: E3 ubiquitin-protein ligase UPL1 isoform X1 [Daucus carota subsp. sativus] | 3775 | 6,53E-31 | 145,976 | 39,50276243 |
| EP00592 | Sar;Stramenopiles;Gyrista;Ochrophyta;"BD clade";[ | EP00592 Proboscia alata 29414 | 5,70E-05 | 804 | XP 002183614.1 | predicted protein [Phaeodactylum tricornutum CCAP 1055/1] | 2083 | 4,56E-17 | 98,2117 | 48,86211513 |
| EP00200 | Archaeplastida;Chloroplastida;Chlorophyta;"core ch | EP00200 Dunaliella tertiolecta 14748 | 5,70E-05 | 433 | XP 013894390.1 | hypothetical protein MNEG 12590 [Monoraphidium neglectum] | 328 | 1,28E-128 | 383,645 | 74,44089457 |
| EP00306 | Haptista;Haptophyta;Pavlovales;Pavlova;strain CCN | EP00306 Pavlova sp CCMP459 22062 | 5,80E-05 | 1269 | XP 002313327.2 | protein transport protein SEC31 homolog B [Populus trichocarpa] | 1135 | 0 | 651,744 | 48,51239669 |
| EP00038 | Obazoa;Opisthokonta;Holozoa;Choanozoa;Choanof | EP00038 Savillea parva 12450 | 5,80E-05 | 993 | XP 009081252.1 | PREDICTED: protein transport protein Sec31A isoform X3 [Acanthisitta chloris] | 1175 | 5,13E-31 | 143,665 | 46,25984252 |
| EP00040 | Obazoa;Opisthokonta;Holozoa;Choanozoa;Choanof | EP00040 Diaphanoeca grandis 24388 | 5,90E-05 | 173 | ***no hit found*** | | | | | |
| EP00256 | Archaeplastida;Chloroplastida;Streptophyta;"core s | EP00256 Glycine max 23050 | 6,00E-05 | 902 | XP 003544622.1 | transcriptional corepressor LEUNIG isoform X1 [Glycine max] >gi\|1590600321\|ref\|XP 02 | 902 | 0 | 1858,57 | 100 |
| EP00770 | Metamonada;Preaxostyla;Oxymonadida;Monocerc | EP00770 Monocercomonoides exilis 15433 | 6,10E-05 | 3034 | XP 002042100.1 | GM10100 [Drosophila sechellia] | 503 | 4,92E-108 | 366,311 | 45,21912351 |
| EP00466 | Sar;Rhizaria;Cercozoa;Chlorarachnea;"core chlorara | EP00466 Bigelowiella natans 4592 | 6,10E-05 | 329 | ***no hit found*** | | | | | |
| EP00227 | Archaeplastida;Chloroplastida;Chlorophyta;Mamiel | EP00227 Mantoniella beaufortii 3169 | 6,10E-05 | 828 | ***no hit found*** | | | | | |
| EP00472 | Sar;Rhizaria;Cercozoa;Chlorarachnea;"core chlorara | EP00472 Partenskyella glossopodia 6724 | 6,20E-05 | 755 | ***no hit found*** | | | | | |
| EP00282 | Cryptista;Cryptophyceae;Cryptomonadales;"lineage | EP00282 Hemiselmis andersenii 64 | 6,30E-05 | 1079 | XP 005831033.1 | secretory protein Sec31 [Guillardia theta CCMP2712] | 1081 | 0 | 1063,14 | 63,90728477 |
| EP00044 | Obazoa;Opisthokonta;Holozoa;Choanozoa;Choanof | EP00044 Salpingoeca rosetta 5380 | 6,50E-05 | 1544 | XP 004992813.1 | hypothetical protein PTSG 05920 [Salpingoeca rosetta] | 1544 | 0 | 3142,44 | 100 |
| EP00022 | Amoebozoa;Evosea;Eumycetozoa;Dictyostelia;Acyto | EP00022 Cavenderia fasciculata 7296 | 6,50E-05 | 2967 | XP 004359786.1 | putative E3 ubiquitin-protein ligase [Cavenderia fasciculata] | 2967 | 0 | 6065,34 | 100 |
| EP00008 | Amoebozoa;Discosea;Flabellinia;"core flabellinids"; | EP00008 Paramoeba atlantica 3031 | 6,50E-05 | 1058 | WP 068670387.1 | hypothetical protein [Thermosulfurimonas dismutans] | 327 | 1,06E-13 | 84,3445 | 42,30769231 |
| EP00751 | Sar;Stramenopiles;Gyrista;Ochrophyta;"BD clade";[ | EP00751 Fragilariopsis kerguelensis 12611 | 6,50E-05 | 1280 | XP 002186077.1 | predicted protein [Phaeodactylum tricornutum CCAP 1055/1] | 771 | 1,69E-47 | 194,512 | 55,49132948 |
| EP00194 | Archaeplastida;Chloroplastida;Chlorophyta;"core ch | EP00194 Tetraselmis striata 30049 | 6,70E-05 | 296 | ***no hit found*** | | | | | |
| EP00004 | CRuMs;Rigifilida;Rigifila;Rigifila ramosa;strain CCAF | EP00004 Rigifila ramosa 12234 | 6,80E-05 | 4005 | XP 012187456.1 | hypothetical protein PHSY 001436 [Pseudozyma hubeiensis SY62] | 578 | 4,41E-168 | 542,732 | 56,54648956 |
| EP00004 | CRuMs;Rigifilida;Rigifila;Rigifila ramosa;strain CCAF | EP00004 Rigifila ramosa 12237 | 6,80E-05 | 4005 | XP 012187456.1 | hypothetical protein PHSY 001436 [Pseudozyma hubeiensis SY62] | 578 | 4,41E-168 | 542,732 | 56,54648956 |
| EP00044 | Obazoa;Opisthokonta;Holozoa;Choanozoa;Choanof | EP00044 Salpingoeca rosetta 1460 | 6,90E-05 | 1031 | XP 004988893.1 | hypothetical protein PTSG 13061 [Salpingoeca rosetta] | 1031 | 0 | 2111,65 | 100 |
| EP00045 | Obazoa;Opisthokonta;Holozoa;Choanozoa;Choanof | EP00045 Choanoeca perplexa 10460 | 6,90E-05 | 609 | XP 004994760.1 | hypothetical protein PTSG 04670 [Salpingoeca rosetta] | 1237 | 5,91E-13 | 83,9593 | 54,27631579 |
| EP00203 | Archaeplastida;Chloroplastida;Chlorophyta;"core ch | EP00203 Chlorella variabilis 1145 | 7,00E-05 | 2766 | XP 005843691.1 | hypothetical protein CHLNCDRAFT 55047 [Chlorella variabilis] | 2766 | 0 | 5162,82 | 100 |
| EP00683 | Discoba;Euglenozoa;Kinetoplastea;Metakinetoplast | EP00683 Trypanosoma rangeli 7231 | 7,10E-05 | 1379 | XP 029240188.1 | putative protein transport protein Sec31 [Trypanosoma rangeli] | 1379 | 0 | 2784,98 | 100 |
| EP00754 | Discoba;Euglenozoa;Diplonemea;Diplonemidae;Rhy | EP00754 Rhynchopus humris 39362 | 7,10E-05 | 511 | ***no hit found*** | | | | | |
| EP00521 | Sar;Stramenopiles;Gyrista;Ochrophyta;"BD clade";[ | EP00521 Asterionellopsis glacialis 12423 | 7,20E-05 | 471 | XP 002178720.1 | predicted protein [Phaeodactylum tricornutum CCAP 1055/1] | 452 | 1,40E-77 | 258,84 | 49,66666667 |
| EP00125 | Obazoa;Opisthokonta;Holozoa;Ichthyosporea;Ichthy | EP00125 Sphaeroforma arctica 11724 | 7,20E-05 | 2195 | XP 014154799.1 | hypothetical protein SARC 06765 [Sphaeroforma arctica JP610] | 2195 | 0 | 4489,87 | 100 |
| EP00044 | Obazoa;Opisthokonta;Holozoa;Choanozoa;Choanof | EP00044 Salpingoeca rosetta 6363 | 7,20E-05 | 1063 | XP 004993796.1 | hypothetical protein PTSG 05218 [Salpingoeca rosetta] | 1063 | 0 | 2144,78 | 100 |
| EP00376 | Sar;Alveolata;Myzozoa;"AC clade";Apicomplexa;"cc | EP00376 Eimeria tenella 420 | 7,30E-05 | 3496 | XP 013228146.1 | Protein EMB-4, isoform d, related [Eimeria tenella] | 3496 | 0 | 6935,12 | 100 |
| EP00443 | Sar;Alveolata;Myzozoa;"DP clade";Dinoflagellata;D | EP00443 Scrippsiella trochoidea 14801 | 7,40E-05 | 806 | XP 025109932.1 | CAP-Gly domain-containing linker protein 3-like isoform X3 [Pomacea canaliculata] | 662 | 3,04E-42 | 174,866 | 42,74406332 |
| EP00435 | Sar;Alveolata;Myzozoa;"DP clade";Dinoflagellata;D | EP00435 Cladocopium sp Y103 11363 | 7,40E-05 | 491 | XP 028394615.1 | procollagen galactosyltransferase 1-like [Dendronephthya gigantea] | 585 | 5,22E-17 | 95,1301 | 47,27272727 |
| EP00376 | Sar;Alveolata;Myzozoa;"AC clade";Apicomplexa;"cc | EP00376 Eimeria tenella 4362 | 7,50E-05 | 786 | XP 013232095.1 | zinc finger (C3HC4 type) / FHA domain-containing protein, putative [Eimeria tenella] | 786 | 0 | 1516,9 | 100 |
| EP00046 | Obazoa;Opisthokonta;Holozoa;Choanozoa;Choanof | EP00046 Monosiga brevicollis 6743 | 7,60E-05 | 961 | XP 001748741.1 | hypothetical protein [Monosiga brevicollis MX1] | 961 | 0 | 1984,15 | 100 |
| EP00428 | Sar;Alveolata;Myzozoa;"DP clade";Dinoflagellata;D | EP00428 Durinskia baltica 2745 | 7,60E-05 | 580 | ***no hit found*** | | | | | |
| EP00633 | Sar;Stramenopiles;Gyrista;Ochrophyta;Pelagophyce | EP00633 Aureoumbra lagunensis 5593 | 7,70E-05 | 838 | XP 009040700.1 | hypothetical protein AURANDRAFT 67110 [Aureococcus anophagefferens] | 578 | 6,02E-56 | 213,001 | 52,85714286 |
| EP00376 | Sar;Alveolata;Myzozoa;"AC clade";Apicomplexa;"cc | EP00376 Eimeria tenella 6577 | 7,70E-05 | 888 | XP 013234311.1 | hypothetical protein ETH 00009765, partial [Eimeria tenella] | 888 | 0 | 1678,3 | 100 |
| EP00007 | Amoebozoa;Discosea;Flabellinia;"core flabellinids"; | EP00007 Cunea sp BSH-02190019 4867 | 7,80E-05 | 959 | ***no hit found*** | | | | | |
| EP00053 | Obazoa;Opisthokonta;Holozoa;Choanozoa;Choanof | EP00053 Salpingoeca punica 14056 | 7,80E-05 | 911 | ***no hit found*** | | | | | |
| EP00125 | Obazoa;Opisthokonta;Holozoa;Ichthyosporea;Ichthy | EP00125 Sphaeroforma arctica 11723 | 7,80E-05 | 2096 | XP 014154799.1 | hypothetical protein SARC 06765 [Sphaeroforma arctica JP610] | 2195 | 0 | 4271,08 | 100 |
| EP00023 | Amoebozoa;Evosea;Eumycetozoa;Dictyostelia;Dicty | EP00023 Dictyostelium discoideum 6597 | 7,80E-05 | 602 | XP 640741.1 | hypothetical protein DDB G0281351 [Dictyostelium discoideum AX4] | 602 | 0 | 1217,99 | 100 |
| EP00044 | Obazoa;Opisthokonta;Holozoa;Choanozoa;Choanof | EP00044 Salpingoeca rosetta 7706 | 8,00E-05 | 582 | XP 004995139.1 | hypothetical protein PTSG 04243 [Salpingoeca rosetta] | 582 | 0 | 1185,24 | 100 |
| EP00401 | Sar;Alveolata;Myzozoa;"DP clade";Dinoflagellata;D | EP00401 Gymnodinium catenatum 63075 | 8,10E-05 | 633 | ***no hit found*** | | | | | |
| EP00319 | Haptista;Haptophyta;Prymnesiophyceae;Phaeocyst | EP00319 Phaeocystis globosa 34135 | 8,20E-05 | 324 | XP 005788197.1 | secretory protein 31 [Emiliania huxleyi CCMP1516] | 1166 | 1,99E-78 | 269,626 | 63,15789474 |
| EP00376 | Sar;Alveolata;Myzozoa;"AC clade";Apicomplexa;"cc | EP00376 Eimeria tenella 7100 | 8,20E-05 | 686 | XP 013234836.1 | hypothetical protein, conserved, partial [Eimeria tenella] | 686 | 0 | 1301,19 | 100 |
| EP00152 | Obazoa;Opisthokonta;Nucletmycea;Fungi;"core Fur | EP00152 Rhizopus delemar 6076 | 8,20E-05 | 1015 | XP 023468011.1 | hypothetical protein RHIMIDRAFT 236286 [Rhizopus microsporus ATCC 52813] | 1104 | 0 | 1634,77 | 86,55913978 |
| EP00289 | Cryptista;Cryptophyceae;Cryptomonadales;"lineage | EP00289 Rhodomonas salina 11341 | 8,30E-05 | 695 | XP 005831033.1 | secretory protein Sec31 [Guillardia theta CCMP2712] | 1081 | 1,38E-143 | 456,062 | 71,12299465 |
| EP00200 | Archaeplastida;Chloroplastida;Chlorophyta;"core ch | EP00200 Dunaliella tertiolecta 10415 | 8,30E-05 | 2337 | XP 002951522.1 | hypothetical protein VOLCADRAFT 92108 [Volvox carteri f. nagariensis] | 2115 | 4,77E-77 | 297,745 | 52,86783042 |
| EP00613 | Sar;Stramenopiles;Gyrista;Ochrophyta;Dictyochoph | EP00613 Pedinella sp CCMP2098 48938 | 8,60E-05 | 606 | XP 009042371.1 | hypothetical protein AURANDRAFT 68438, partial [Aureococcus anophagefferens] | 817 | 2,50E-15 | 90,8929 | 45,21963824 |
| EP00376 | Sar;Alveolata;Myzozoa;"AC clade";Apicomplexa;"cc | EP00376 Eimeria tenella 7307 | 8,60E-05 | 1095 | XP 013235046.1 | DEHA2A05324p, related, related [Eimeria tenella] | 1095 | 0 | 2139,77 | 100 |
| EP00106 | Obazoa;Opisthokonta;Holozoa;Choanozoa;Metazoa | EP00106 Octopus bimaculoides 19576 | 8,70E-05 | 1170 | XP 014787018.1 | PREDICTED: ubiquitin carboxyl-terminal hydrolase 25-like isoform X3 [Octopus bimaculoid | 1170 | 0 | 2445,62 | 100 |
| EP00044 | Obazoa;Opisthokonta;Holozoa;Choanozoa;Choanof | EP00044 Salpingoeca rosetta 10848 | 8,80E-05 | 990 | XP 004998281.1 | hypothetical protein PTSG 00812 [Salpingoeca rosetta] | 990 | 0 | 2023,05 | 100 |
| EP00023 | Amoebozoa;Evosea;Eumycetozoa;Dictyostelia;Dicty | EP00023 Dictyostelium discoideum 1251 | 8,90E-05 | 1527 | XP 629538.1 | hypothetical protein DDB G0292556 [Dictyostelium discoideum AX4] | 1527 | 0 | 3029,58 | 100 |
| EP00289 | Cryptista;Cryptophyceae;Cryptomonadales;"lineage | EP00289 Rhodomonas salina 3501 | 9,30E-05 | 373 | XP 005831033.1 | secretory protein Sec31 [Guillardia theta CCMP2712] | 1081 | 6,54E-136 | 423,705 | 73,91304348 |
| EP00521 | Sar;Stramenopiles;Gyrista;Ochrophyta;"BD clade";[ | EP00521 Asterionellopsis glacialis 15695 | 9,40E-05 | 841 | XP 002181573.1 | predicted protein [Phaeodactylum tricornutum CCAP 1055/1] | 829 | 2,07E-118 | 388,267 | 45,95959596 |
| EP00578 | Sar;Stramenopiles;Gyrista;Ochrophyta;"BD clade";[ | EP00578 Thalassiosira sp NH16 24178 | 9,50E-05 | 480 | XP 002295385.1 | hypothetical protein THAPS 264495, partial [Thalassiosira pseudonana CCMP1335] | 87 | 4,27E-14 | 77,7962 | 47,82608696 |
| EP00237 | Archaeplastida;Chloroplastida;Chlorophyta;Pycnocc | EP00237 Pycnococcus provasolii 1387 | 9,60E-05 | 919 | XP 021993887.1 | zinc finger protein BRUTUS-like [Helianthus annuus] | 1195 | 4,59E-76 | 281,567 | 56,9124424 |
| EP00241 | Archaeplastida;Chloroplastida;Chlorophyta;Pyramir | EP00241 Pyramimonas parkeae 4376 | 9,60E-05 | 897 | XP 001450301.1 | hypothetical protein [Paramecium tetraurelia strain d4-2] | 504 | 1,34E-15 | 92,0485 | 73 |
| EP00239 | Archaeplastida;Chloroplastida;Chlorophyta;Pyramir | EP00239 Pterosperma sp CCMP1384 8413 | 9,80E-05 | 908 | XP 024363375.1 | vacuolar-sorting protein BRO1-like [Physcomitrella patens] | 861 | 6,97E-173 | 533,102 | 45,72158365 |
| EP00309 | Haptista;Haptophyta;Prymnesiophyceae;Calcihapto | EP00309 Chrysotila carterae 18593 | 9,80E-05 | 711 | XP 004346254.2 | TBC1 domain family protein [Capsaspora owczarzaki ATCC 30864] | 1005 | 2,25E-29 | 136,346 | 42,85714286 |
| EP00751 | Sar;Stramenopiles;Gyrista;Ochrophyta;"BD clade";[ | EP00751 Fragilariopsis kerguelensis 795 | 9,80E-05 | 1493 | ***no hit found*** | | | | | |
| EP00402 | Sar;Alveolata;Myzozoa;"DP clade";Dinoflagellata;D | EP00402 Amphidinium carterae 30733 | 9,80E-05 | 656 | XP 004356402.1 | hypothetical protein DFA 08919 [Cavenderia fasciculata] | 1450 | 9,41E-25 | 121,709 | 61,47308782 |
| EP00376 | Sar;Alveolata;Myzozoa;"AC clade";Apicomplexa;"cc | EP00376 Eimeria tenella 5096 | 9,90E-05 | 1816 | XP 013232829.1 | hypothetical protein, conserved [Eimeria tenella] | 1816 | 0 | 3590,82 | 100 |
| EP00418 | Sar;Alveolata;Myzozoa;"DP clade";Dinoflagellata;D | EP00418 Pyrodinium bahamense 36300 | 0,0001 | 890 | ***no hit found*** | | | | | |
| EP00309 | Haptista;Haptophyta;Prymnesiophyceae;Calcihapto | EP00309 Chrysotila carterae 29809 | 0,0001 | 1097 | XP 005788197.1 | secretory protein 31 [Emiliania huxleyi CCMP1516] | 1166 | 0 | 883,634 | 61,31078224 |
| EP00441 | Sar;Alveolata;Myzozoa;"DP clade";Dinoflagellata;D | EP00441 Pelagodinium beii 35279 | 0,0001 | 240 | XP 002771093.1 | hypothetical protein Pmar PMAR000901 [Perkinsus marinus ATCC 50983] | 173 | 4,91E-06 | 55,8398 | 49,16666667 |
| EP00601 | Sar;Stramenopiles;Gyrista;Ochrophyta;"CS clade";C | EP00601 Ochromonadales sp CCMP2298 28564 | 0,0001 | 1286 | XP 002183894.1 | Mer3 meiotic cross-over helicase [Phaeodactylum tricornutum CCAP 1055/1] | 1710 | 7,78E-14 | 88,5817 | 42,28187919 |
| EP00198 | Archaeplastida;Chloroplastida;Chlorophyta;"core ch | EP00198 Chlamydomonas reinhardtii 13965 | 0,0001 | 1367 | XP 001703302.1 | predicted protein [Chlamydomonas reinhardtii] | 1367 | 0 | 2669,42 | 100 |
| EP00256 | Archaeplastida;Chloroplastida;Streptophyta;"core s | EP00256 Glycine max 33050 | 0,0001 | 907 | XP 006578171.1 | transcriptional corepressor LEUNIG isoform X2 [Glycine max] | 907 | 0 | 1858,96 | 100 |
| EP00036 | Obazoa;Opisthokonta;Holozoa;Choanozoa;Choanof | EP00036 Acanthoeca sp 10tr 4057 | 0,00011 | 685 | XP 010181369.1 | PREDICTED: protein transport protein Sec31A [Mesitornis unicolor] | 989 | 2,75E-41 | 172,94 | 43,30042313 |
| EP00196 | Archaeplastida;Chloroplastida;Chlorophyta;"core ch | EP00196 Chlamydomonas euryale 18287 | 0,00011 | 1484 | XP 002956958.1 | hypothetical protein VOLCADRAFT 67693 [Volvox carteri f. nagariensis] | 283 | 5,25E-53 | 198,364 | 53,30882353 |
| EP00532 | Sar;Stramenopiles;Gyrista;Ochrophyta;"BD clade";[ | EP00532 Pseudo-nitzschia australis 20775 | 0,00011 | 749 | XP 002184843.1 | predicted protein [Phaeodactylum tricornutum CCAP 1055/1] | 696 | 0 | 728,398 | 66,22516556 |
| EP00434 | Sar;Alveolata;Myzozoa;"DP clade";Dinoflagellata;D | EP00434 Breviolum minutum 30679 | 0,00011 | 1235 | XP 025179275.1 | putative vacuolar protein sorting-associated protein [Rhizophagus irregularis DAOM 1816 | 577 | 5,42E-166 | 514,998 | 51,19453925 |
| EP00011 | Amoebozoa;Discosea;Flabellinia;"core flabellinids"; | EP00011 Vannella sp DIVA3-517-6-12 6741 | 0,00011 | 1707 | XP 004338529.1 | RhoGEF domain containing protein [Acanthamoeba castellanii str. Neff] | 670 | 1,49E-38 | 166,392 | 36,75496689 |
| EP00054 | Obazoa;Opisthokonta;Holozoa;Choanozoa;Choanof | EP00054 Salpingoeca dolichothecata 15541 | 0,00011 | 829 | ***no hit found*** | | | | | |
| EP00139 | Obazoa;Opisthokonta;Nucletmycea;Fungi;"core Fur | EP00139 Candida albicans 5339 | 0,00011 | 506 | XP 721805.2 | putative glucan endo-1\3-beta-D-glucosidase [Candida albicans SC5314] | 506 | 0 | 994,186 | 100 |
| EP00759 | Discoba;Euglenozoa;Kinetoplastea;Prokinetoplastin | EP00759 Prokinetoplastina sp PhF-6 4471 | 0,00011 | 934 | XP 004355165.1 | WD40 repeat-containing protein [Cavenderia fasciculata] | 1147 | 3,30E-114 | 387,497 | 46,18181818 |
| EP00401 | Sar;Alveolata;Myzozoa;"DP clade";Dinoflagellata;D | EP00401 Gymnodinium catenatum 61742 | 0,00011 | 644 | ***no hit found*** | | | | | |
| EP00484 | Sar;Rhizaria;Retaria;Foraminifera;Globothalamea;R | EP00484 Ammonia sp Unknown 23607 | 0,00011 | 795 | ***no hit found*** | | | | | |
| EP00593 | Sar;Stramenopiles;Gyrista;Ochrophyta;"BD clade";[ | EP00593 Proboscia inermis 11550 | 0,00011 | 158 | ***no hit found*** | | | | | |
| EP00730 | Obazoa;Opisthokonta;Holozoa;Choanozoa;Choanof | EP00730 Choanoeca flexa 4743 | 0,00012 | 422 | XP 004489519.1 | glycine-rich RNA-binding protein 4, mitochondrial-like [Cicer arietinum] | 265 | 9,66E-14 | 81,6481 | 46,34146341 |
| EP00401 | Sar;Alveolata;Myzozoa;"DP clade";Dinoflagellata;D | EP00401 Gymnodinium catenatum 61623 | 0,00012 | 647 | ***no hit found*** | | | | | |
| EP00376 | Sar;Alveolata;Myzozoa;"AC clade";Apicomplexa;"cc | EP00376 Eimeria tenella 4415 | 0,00012 | 994 | XP 013232148.1 | hypothetical protein, conserved [Eimeria tenella] | 994 | 0 | 1927,14 | 100 |
| EP00078 | Obazoa;Opisthokonta;Holozoa;Choanozoa;Metazoa | EP00078 Anolis carolinensis 28642 | 0,00012 | 3001 | XP 016848752.1 | PREDICTED: mucin-2 [Anolis carolinensis] | 3001 | 0 | 6108,48 | 100 |
| EP00256 | Archaeplastida;Chloroplastida;Streptophyta;"core s | EP00256 Glycine max 23051 | 0,00012 | 893 | XP 003544623.1 | transcriptional corepressor LEUNIG isoform X3 [Glycine max] >gi\|1590600325\|ref\|XP 02 | 893 | 0 | 1840,85 | 100 |
| EP00412 | Sar;Alveolata;Myzozoa;"DP clade";Dinoflagellata;D | EP00412 Alexandrium andersonii 26257 | 0,00012 | 265 | ***no hit found*** | | | | | |
| EP00122 | Obazoa;Opisthokonta;Holozoa;Ichthyosporea;Ichthy | EP00122 Pirum gemmata 21427 | 0,00013 | 392 | WP 116174136.1 | AAA family ATPase [Kutzneria buriramensis] | 1626 | 3,42E-08 | 67,0106 | 67,42424242 |
| EP00256 | Archaeplastida;Chloroplastida;Streptophyta;"core s | EP00256 Glycine max 11062 | 0,00013 | 912 | XP 003522633.1 | transcriptional corepressor LEUNIG isoform X1 [Glycine max] | 912 | 0 | 1868,59 | 100 |
| EP00618 | Sar;Stramenopiles;Gyrista;Ochrophyta;Dictyochoph | EP00618 Florenciella parvula 17602 | 0,00013 | 726 | XP 008610288.1 | PEK/GCN2 protein kinase [Saprolegnia diclina VS20] | 1395 | 9,72E-14 | 87,0409 | 41,6918429 |
| EP00161 | Ancyromonadida;Ancyromonadidae;Ancyromonas;/ | EP00161 Ancyromonas sigmoides 22971 | 0,00013 | 775 | XP 005831033.1 | secretory protein Sec31 [Guillardia theta CCMP2712] | 1081 | 8,79E-37 | 160,614 | 46,83098592 |
| EP00376 | Sar;Alveolata;Myzozoa;"AC clade";Apicomplexa;"cc | EP00376 Eimeria tenella 976 | 0,00013 | 669 | XP 013228702.1 | transmembrane domain-containing protein, putative [Eimeria tenella] | 669 | 0 | 1280,39 | 100 |
| EP00673 | Discoba;Euglenozoa;Kinetoplastea;Metakinetoplast | EP00673 Leishmania infantum 1300 | 0,00013 | 448 | XP 001464086.1 | putative lectin [Leishmania infantum JPCM5] | 448 | 0 | 929,087 | 100 |
| EP00166 | Archaeplastida;"RR clade";Rhodophyta;Cyanidiales; | EP00166 Cyanidium caldarium 4105 | 0,00013 | 948 | XP 005538478.1 | vesicle coat complex COPII, subunit Sec31 [Cyanidioschyzon merolae strain 10D] | 1307 | 1,00E-176 | 557,755 | 46,58753709 |
| EP00537 | Sar;Stramenopiles;Gyrista;Ochrophyta;"BD clade";[ | EP00537 Pseudo-nitzschia pungens 2665 | 0,00014 | 479 | XP 002179384.1 | predicted protein [Phaeodactylum tricornutum CCAP 1055/1] | 352 | 1,09E-144 | 427,557 | 63,76404494 |
| EP00166 | Archaeplastida;"RR clade";Rhodophyta;Cyanidiales; | EP00166 Cyanidium caldarium 6307 | 0,00014 | 1324 | XP 005535367.1 | probable mitochondrial 2-oxoglutarate/malate carrier protein [Cyanidioschyzon merolae | 317 | 6,01E-28 | 126,716 | 60,78431373 |
| EP00096 | Obazoa;Opisthokonta;Holozoa;Choanozoa;Metazoa | EP00096 Caligus rogercresseyi 16768 | 0,00015 | 711 | XP 022127448.1 | tudor domain-containing protein 3 [Pieris rapae] | 684 | 2,74E-47 | 188,734 | 47,15909091 |
| EP00457 | Sar;Rhizaria;Cercozoa;"crown cercozoa";Imbricatea | EP00457 Paulinella chromatophora 3334 | 0,00015 | 723 | XP 002643350.1 | Hypothetical protein CBG15947 [Caenorhabditis briggsae] | 370 | 3,96E-19 | 100,908 | 41,47465438 |
| EP00634 | Sar;Stramenopiles;Gyrista;Ochrophyta;Pelagophyce | EP00634 Sargassococcus sp CCMP2135 3346 | 0,00015 | 436 | XP 009041823.1 | hypothetical protein AURANDRAFT 67970 [Aureococcus anophagefferens] | 1796 | 2,35E-43 | 173,711 | 45,07042254 |

| | | | | | | | | | | |
|---|---|---|---|---|---|---|---|---|---|---|
| EP00078 | Obazoa;Opisthokonta;Holozoa;Choanozoa;Metazoa | EP00078 Anolis carolinensis 23811 | 0,00015 | 2438 | XP 008121353.1 | PREDICTED: zinc finger protein 638, partial [Anolis carolinensis] | 2438 | 0 | 4877 | 100 |
| EP00158 | Obazoa;Opisthokonta;Nucletmycea;Fungi;Aphelidea | EP00158 Paraphelidium tribonemae 9299 | 0,00015 | 1218 | XP 021881067.1 | hypothetical protein BCR41DRAFT 396657 [Lobosporangium transversale] | 1406 | 0 | 593,193 | 50,0394633 |
| EP00290 | Cryptista;Cryptophyceae;Cryptomonadales;Baffinel | EP00290 Baffinella sp CCMP2293 12787 | 0,00015 | 1171 | XP 005831033.1 | secretory protein Sec31 [Guillardia theta CCMP2712] | 1081 | 0 | 1072 | 70,19527235 |
| EP00023 | Amoebozoa;Evosea;Eumycetozoa;Dictyostelia;Dicty | EP00023 Dictyostelium discoideum 3487 | 0,00015 | 158 | XP 636990.1 | hypothetical protein DDB G0287915 [Dictyostelium discoideum AX4] | 158 | 1,90E-111 | 322,398 | 100 |
| EP00556 | Sar;Stramenopiles;Gyrista;Ochrophyta;"BD clade";C | EP00556 Chaetoceros neogracilis 19693 | 0,00015 | 433 | XP 023932950.1 | inositol phosphoceramide mannosyltransferase 2-like [Lingula anatina] | 363 | 2,34E-30 | 130,183 | 43,90243902 |
| EP00698 | Malawimonadidae;Gefionella;Gefionella okellyi;stra | EP00698 Gefionella okellyi 19338 | 0,00015 | 441 | ***no hit found*** | | | | | |
| EP00464 | Sar;Rhizaria;Cercozoa;Chlorarachnea;"core chlorara | EP00464 Amorphochlora amoebiformis 9983 | 0,00015 | 772 | ***no hit found*** | | | | | |
| EP00376 | Sar;Alveolata;Myzozoa;"AC clade";Apicomplexa;"cc | EP00376 Eimeria tenella 8220 | 0,00016 | 1584 | XP 013235964.1 | hypothetical protein, conserved [Eimeria tenella] | 1584 | 0 | 3082,74 | 100 |
| EP00193 | Archaeplastida;Chloroplastida;Chlorophyta;"core ch | EP00193 Tetraselmis chuii 3963 | 0,00016 | 1158 | XP 024379150.1 | protein transport protein SEC31 homolog B-like isoform X1 [Physcomitrella patens] | 1179 | 0 | 831,632 | 52,11382114 |
| EP00600 | Sar;Stramenopiles;Gyrista;Ochrophyta;"CS clade";C | EP00600 Ochromonadales sp CCMP1393 16644 | 0,00016 | 2124 | XP 008907364.1 | hypothetical protein, variant 2 [Phytophthora parasitica INRA-310] | 1175 | 0 | 609,757 | 48,60956618 |
| EP00311 | Haptista;Haptophyta;Prymnesiophyceae;Calcihapto | EP00311 Isochrysis galbana 19552 | 0,00016 | 1121 | XP 005788197.1 | secretory protein 31 [Emiliania huxleyi CCMP1516] | 1166 | 0 | 1082,4 | 69,8284561 |
| EP00435 | Sar;Alveolata;Myzozoa;"DP clade";Dinoflagellata;D | EP00435 Cladocopium sp Y103 11321 | 0,00016 | 473 | XP 028394615.1 | procollagen galactosyltransferase 1-like [Dendronephthya gigantea] | 585 | 4,00E-17 | 95,1301 | 47,27272727 |
| EP00600 | Sar;Stramenopiles;Gyrista;Ochrophyta;"CS clade";C | EP00600 Ochromonadales sp CCMP1393 9380 | 0,00017 | 934 | WP 120752464.1 | hypothetical protein [Aquimarina sp. BL5] | 3354 | 3,54E-38 | 167,162 | 51,32075472 |
| EP00401 | Sar;Alveolata;Myzozoa;"DP clade";Dinoflagellata;D | EP00401 Gymnodinium catenatum 24480 | 0,00017 | 859 | ***no hit found*** | | | | | |
| EP00445 | Sar;Alveolata;Myzozoa;"DP clade";Dinoflagellata;D | EP00445 Apocalathium hangoei 42927 | 0,00017 | 506 | ***no hit found*** | | | | | |
| EP00708 | Metamonada;Preaxostyla;Paratrimastigidae;Paratr | EP00708 Paratrimastix pyriformis 2000 | 0,00017 | 1017 | XP 016681984.1 | PREDICTED: protein transport protein SEC31 homolog B-like isoform X1 [Gossypium hirsut | 1112 | 4,49E-126 | 420,624 | 47,61459308 |
| EP00515 | Sar;Stramenopiles;Bigyra;Sagenista;Labyrinthulom | EP00515 Schizochytrium aggregatum 21299 | 0,00017 | 369 | XP 004357047.1 | small MutS related family protein [Cavenderia fasciculata] | 329 | 4,37E-20 | 100,138 | 46,12244898 |
| EP00153 | Obazoa;Opisthokonta;Nucletmycea;Fungi;"core Fur | EP00153 Phycomyces blakesleeanus 5727 | 0,00017 | 1116 | XP 018288804.1 | hypothetical protein PHYBLDRAFT 64156 [Phycomyces blakesleeanus NRRL 1555(-)] | 1116 | 0 | 2322,35 | 100 |
| EP00024 | Amoebozoa;Evosea;Eumycetozoa;Dictyostelia;Dicty | EP00024 Dictyostelium purpureum 11831 | 0,00017 | 694 | XP 003294683.1 | hypothetical protein DICPUDRAFT 85137 [Dictyostelium purpureum] | 694 | 0 | 1389,02 | 100 |
| EP00317 | Haptista;Haptophyta;Prymnesiophyceae;Phaeocyst | EP00317 Phaeocystis antarctica 247 | 0,00017 | 657 | ***no hit found*** | | | | | |
| EP00404 | Sar;Alveolata;Myzozoa;"DP clade";Dinoflagellata;D | EP00404 Azadinium spinosum 48855 | 0,00017 | 171 | ***no hit found*** | | | | | |
| EP00706 | Metamonada;Parabasalia;Tritrichomonas;Tritrichor | EP00706 Tritrichomonas foetus 9675 | 0,00018 | 1439 | XP 001311898.1 | Clan CA, family C19, ubiquitin hydrolase-like cysteine peptidase [Trichomonas vaginalis G3 | 1201 | 2,06E-90 | 330,872 | 39,83193277 |
| EP00577 | Sar;Stramenopiles;Gyrista;Ochrophyta;"BD clade";C | EP00577 Skeletonema sp RCC1716 2047 | 0,00018 | 1081 | XP 002289786.1 | predicted protein [Thalassiosira pseudonana CCMP1335] | 1191 | 0 | 872,848 | 66,45398616 |
| EP00011 | Amoebozoa;Discosea;Flabellinia;"core flabellinids"; | EP00011 Vannella sp DIVA3-517-6-12 6866 | 0,00018 | 758 | XP 004368323.1 | dynamin domain containing protein [Acanthamoeba castellanii str. Neff] | 1263 | 0 | 823,928 | 58,48329049 |
| EP00504 | Sar;Stramenopiles;Bigyra;Opalozoa;Opalinata;Blast | EP00504 Blastocystis sp subtype7 4633 | 0,00018 | 870 | XP 012898216.1 | uncharacterized protein [Blastocystis hominis] | 870 | 0 | 1786,54 | 100 |
| EP00044 | Obazoa;Opisthokonta;Holozoa;Choanozoa;Choanof | EP00044 Salpingoeca rosetta 6728 | 0,00018 | 1250 | XP 004994161.1 | hypothetical protein PTSG 04843 [Salpingoeca rosetta] | 1250 | 0 | 2563,87 | 100 |
| EP00393 | Sar;Alveolata;Myzozoa;"DP clade";Dinoflagellata;"c | EP00393 Amoebophrya sp ex-Akashiwo-sanguinea 57143 | 0,00018 | 938 | XP 007931629.1 | hypothetical protein MYCFIDRAFT 168463 [Pseudocercospora fijiensis CIRAD86] | 702 | 5,55E-52 | 205,297 | 47,91288566 |
| EP00201 | Archaeplastida;Chloroplastida;Chlorophyta;"core ch | EP00201 Polytomella parva 6668 | 0,00018 | 551 | XP 002946119.1 | hypothetical protein VOLCADRAFT 115694 [Volvox carteri f. nagariensis] | 577 | 1,26E-60 | 219,935 | 58,92857143 |
| EP00437 | Sar;Alveolata;Myzozoa;"DP clade";Dinoflagellata;D | EP00437 Effrenium voratum 15315 | 0,00018 | 225 | XP 002772264.1 | hypothetical protein Pmar PMAR023065 [Perkinsus marinus ATCC 50983] | 173 | 7,14E-08 | 60,4622 | 48,125 |
| EP00322 | Haptista;Haptophyta;Prymnesiophyceae;Prymnesia | EP00322 Chrysochromulina rotalis 27437 | 0,00018 | 576 | ***no hit found*** | | | | | |
| EP00044 | Obazoa;Opisthokonta;Holozoa;Choanozoa;Choanof | EP00044 Salpingoeca rosetta 5763 | 0,00019 | 1260 | XP 004993196.1 | ADAM-17 protein [Salpingoeca rosetta] | 1260 | 0 | 2602,01 | 100 |
| EP00334 | Sar;Alveolata;Ciliophora;Intramacronucleata;"CON | EP00334 Paramecium tetraurelia 14339 | 0,00019 | 869 | XP 001437328.1 | hypothetical protein [Paramecium tetraurelia strain d4-2] | 869 | 0 | 1749,56 | 100 |
| EP00445 | Sar;Alveolata;Myzozoa;"DP clade";Dinoflagellata;D | EP00445 Apocalathium hangoei 42164 | 0,00019 | 801 | XP 005818320.1 | hypothetical protein GUITHDRAFT 122458 [Guillardia theta CCMP2712] | 189 | 6,10E-10 | 70,0922 | 98,27586207 |
| EP00286 | Cryptista;Cryptophyceae;Cryptomonadales;"lineage | EP00286 Rhodomonas abbreviata 26319 | 0,00019 | 599 | ***no hit found*** | | | | | |
| EP00444 | Sar;Alveolata;Myzozoa;"DP clade";Dinoflagellata;D | EP00444 Apocalathium aciculiferum 46171 | 0,00019 | 86 | ***no hit found*** | | | | | |
| EP00301 | Haptista;Centroplasthelida;Pterocystida;Raphidista | EP00301 Raphidiophrys heterophryoidea 8014 | 0,00019 | 1069 | XP 028843481.1 | xylosyltransferase 2 [Denticeps clupeoides] | 874 | 2,20E-23 | 118,627 | 47,50733138 |
| EP00418 | Sar;Alveolata;Myzozoa;"DP clade";Dinoflagellata;D | EP00418 Pyrodinium bahamense 48699 | 0,00019 | 1618 | XP 002781537.1 | calcium-dependent protein kinase, putative [Perkinsus marinus ATCC 50983] | 990 | 6,49E-87 | 318,161 | 50,82987552 |
| EP00237 | Archaeplastida;Chloroplastida;Chlorophyta;Pycnoc | EP00237 Pycnococcus provasolii 124 | 0,00019 | 1262 | XP 005649785.1 | hypothetical protein COCSUDRAFT 65166 [Coccomyxa subellipsoidea C-169] | 856 | 1,96E-22 | 115,931 | 47,6635514 |
| EP00376 | Sar;Alveolata;Myzozoa;"AC clade";Apicomplexa;"cc | EP00376 Eimeria tenella 4838 | 0,0002 | 2498 | XP 013232571.1 | hypothetical protein, conserved [Eimeria tenella] | 2498 | 0 | 4988,71 | 100 |
| EP00462 | Sar;Rhizaria;Cercozoa;"crown cercozoa";Thecofilos | EP00462 Mataza sp D1 21777 | 0,00021 | 520 | XP 024529345.1 | methyltransferase-like protein 14 homolog [Selaginella moellendorffii] | 420 | 8,11E-114 | 352,829 | 64,78405316 |
| EP00382 | Sar;Alveolata;Myzozoa;"AC clade";Apicomplexa;"gr | EP00382 Lankesteria abbotti 1889 | 0,00021 | 463 | ***no hit found*** | | | | | |
| EP00101 | Obazoa;Opisthokonta;Holozoa;Choanozoa;Metazoa | EP00101 Apis mellifera 9988 | 0,00021 | 3122 | XP 016767280.2 | ankyrin repeat domain-containing protein 17 isoform X3 [Apis mellifera] | 3122 | 0 | 6454 | 100 |
| EP00079 | Obazoa;Opisthokonta;Holozoa;Choanozoa;Metazoa | EP00079 Xenopus tropicalis 6634 | 0,00021 | 659 | NP 001121427.1 | splicing factor, proline- and glutamine-rich [Xenopus tropicalis] | 659 | 0 | 1337,01 | 100 |
| EP00044 | Obazoa;Opisthokonta;Holozoa;Choanozoa;Choanof | EP00044 Salpingoeca rosetta 2377 | 0,00022 | 970 | XP 004989810.1 | AGC/RSK/MSK protein kinase [Salpingoeca rosetta] | 970 | 0 | 2005,33 | 100 |
| EP00042 | Obazoa;Opisthokonta;Holozoa;Choanozoa;Choanof | EP00042 Codosiga hollandica 40361 | 0,00022 | 409 | XP 004994691.1 | hypothetical protein PTSG 04597 [Salpingoeca rosetta] | 286 | 1,12E-45 | 168,703 | 56,875 |
| EP00179 | Archaeplastida;"RR clade";Rhodophyta;Proteorhod | EP00179 Madagascaria erythrocladioides 27920 | 0,00022 | 963 | ***no hit found*** | | | | | |
| EP00572 | Sar;Stramenopiles;Gyrista;Ochrophyta;"BD clade";C | EP00572 Skeletonema dohrnii 11534 | 0,00022 | 1149 | XP 003080029.1 | Putative 5-3 exonuclease [Ostreococcus tauri] | 1057 | 0 | 574,318 | 53,32640333 |
| EP00312 | Haptista;Haptophyta;Prymnesiophyceae;Calcihapto | EP00312 Isochrysis sp CCMP1244 28316 | 0,00022 | 718 | XP 005788197.1 | secretory protein 31 [Emiliania huxleyi CCMP1516] | 1166 | 0 | 1417,52 | 99,86052999 |
| EP00670 | Discoba;Euglenozoa;Euglenida;Spirocuta;Euglenea;I | EP00670 Eutreptiella gymnastica 8171 | 0,00022 | 2203 | XP 011650480.1 | PREDICTED: E3 ubiquitin-protein ligase UPL1 isoform X4 [Cucumis sativus] | 3783 | 2,26E-54 | 223,787 | 37,76418242 |
| EP00751 | Sar;Stramenopiles;Gyrista;Ochrophyta;"BD clade";C | EP00751 Fragilariopsis kerguelensis 30799 | 0,00022 | 810 | ***no hit found*** | | | | | |
| EP00008 | Amoebozoa;Discosea;Flabellinia;"core flabellinids"; | EP00008 Paramoeba atlantica 869 | 0,00022 | 1893 | XP 024385481.1 | protein MOR1-like isoform X2 [Physcomitrella patens] | 1998 | 2,89E-121 | 436,802 | 39,48256468 |
| EP00376 | Sar;Alveolata;Myzozoa;"AC clade";Apicomplexa;"cc | EP00376 Eimeria tenella 4962 | 0,00023 | 1799 | XP 013232695.1 | hypothetical protein ETH 00023690 [Eimeria tenella] | 1799 | 0 | 3556,92 | 100 |
| EP00443 | Sar;Alveolata;Myzozoa;"DP clade";Dinoflagellata;D | EP00443 Scrippsiella trochoidea 96889 | 0,00023 | 553 | XP 002768122.1 | Carbon catabolite derepressing protein kinase, putative [Perkinsus marinus ATCC 50983] | 773 | 7,98E-155 | 471,085 | 66,15384615 |
| EP00435 | Sar;Alveolata;Myzozoa;"DP clade";Dinoflagellata;D | EP00435 Cladocopium sp Y103 24560 | 0,00023 | 498 | XP 023340916.1 | myb-like protein X isoform X1 [Eurytemora affinis] | 590 | 6,51E-12 | 79,337 | 56,02409639 |
| EP00451 | Sar;Alveolata;Myzozoa;"DP clade";Dinoflagellata;O | EP00451 Oxyrrhis marina 10055 | 0,00023 | 558 | ***no hit found*** | | | | | |
| EP00378 | Sar;Alveolata;Myzozoa;"AC clade";Apicomplexa;"cc | EP00378 Neospora caninum 2038 | 0,00023 | 2378 | XP 003881576.1 | putative plectin [Neospora caninum Liverpool] | 2378 | 0 | 4682,09 | 100 |
| EP00290 | Cryptista;Cryptophyceae;Cryptomonadales;Baffinel | EP00290 Baffinella sp CCMP2293 36248 | 0,00023 | 1188 | XP 005831033.1 | secretory protein Sec31 [Guillardia theta CCMP2712] | 1081 | 0 | 1021,92 | 69,51391299 |
| EP00007 | Amoebozoa;Discosea;Flabellinia;"core flabellinids"; | EP00007 Cunea sp BSH-02190019 5503 | 0,00023 | 1128 | XP 005835801.1 | hypothetical protein GUITHDRAFT 151698 [Guillardia theta CCMP2712] | 367 | 8,07E-32 | 139,043 | 46,62576687 |
| EP00200 | Archaeplastida;Chloroplastida;Chlorophyta;"core ch | EP00200 Dunaliella tertiolecta 14328 | 0,00023 | 1355 | ***no hit found*** | | | | | |
| EP00516 | Sar;Stramenopiles;Bigyra;Sagenista;Labyrinthulom | EP00516 Thraustochytrida sp NY0313808BC1 11062 | 0,00024 | 1319 | XP 008907364.1 | hypothetical protein, variant 2 [Phytophthora parasitica INRA-310] | 1175 | 4,39E-151 | 497,278 | 49,46695096 |
| EP00322 | Haptista;Haptophyta;Prymnesiophyceae;Prymnesia | EP00322 Chrysochromulina rotalis 970 | 0,00024 | 529 | ***no hit found*** | | | | | |
| EP00434 | Sar;Alveolata;Myzozoa;"DP clade";Dinoflagellata;D | EP00434 Breviolum minutum 30678 | 0,00024 | 683 | XP 002764392.1 | homeobox-containing protein, putative [Perkinsus marinus ATCC 50983] | 310 | 2,91E-48 | 182,185 | 64,70588235 |
| EP00256 | Archaeplastida;Chloroplastida;Streptophyta;"core s | EP00256 Glycine max 11063 | 0,00024 | 903 | XP 003522634.1 | transcriptional corepressor LEUNIG isoform X3 [Glycine max] | 903 | 0 | 1852,03 | 100 |
| EP00686 | Discoba;Heterolobosea;Tetramitia;Eutetramitia;"M | EP00686 Naegleria gruberi 7943 | 0,00024 | 473 | XP 002675872.1 | wiscott-aldrich syndrome protein [Naegleria gruberi] | 473 | 0 | 951,429 | 100 |
| EP00084 | Obazoa;Opisthokonta;Holozoa;Choanozoa;Metazoa | EP00084 Strigamia maritima 3081 | 0,00024 | 1233 | XP 013780411.2 | protein transport protein Sec31A-like [Limulus polyphemus] | 1162 | 0 | 1085,09 | 61,5503876 |
| EP00027 | Amoebozoa;Evosea;Variosea;Filamoeba;Filamoeba | EP00027 Filamoeba nolandi 18986 | 0,00024 | 432 | ***no hit found*** | | | | | |
| EP00756 | Discoba;Euglenozoa;Diplonemea;Hemistasiidae;Her | EP00756 Hemistasia phaeocysticola 11920 | 0,00025 | 717 | XP 020608340.1 | dipeptidyl peptidase 9-like [Orbicella faveolata] | 861 | 1,66E-83 | 292,352 | 52,43128964 |
| EP00756 | Discoba;Euglenozoa;Diplonemea;Hemistasiidae;Her | EP00756 Hemistasia phaeocysticola 11921 | 0,00025 | 717 | XP 020608340.1 | dipeptidyl peptidase 9-like [Orbicella faveolata] | 861 | 1,66E-83 | 292,352 | 52,43128964 |
| EP00756 | Discoba;Euglenozoa;Diplonemea;Hemistasiidae;Her | EP00756 Hemistasia phaeocysticola 11922 | 0,00025 | 717 | XP 020608340.1 | dipeptidyl peptidase 9-like [Orbicella faveolata] | 861 | 1,66E-83 | 292,352 | 52,43128964 |
| EP00393 | Sar;Alveolata;Myzozoa;"DP clade";Dinoflagellata;"c | EP00393 Amoebophrya sp ex-Akashiwo-sanguinea 26332 | 0,00025 | 702 | XP 002788366.1 | hypothetical protein, conserved [Perkinsus marinus ATCC 50983] | 413 | 1,88E-53 | 199,519 | 52,97450425 |
| EP00023 | Amoebozoa;Evosea;Eumycetozoa;Dictyostelia;Dicty | EP00023 Dictyostelium discoideum 3841 | 0,00025 | 1524 | XP 637368.1 | DUF1683 family protein [Dictyostelium discoideum AX4] | 1524 | 0 | 3088,9 | 100 |
| EP00003 | CRuMs;Mantamonas;Mantamonas plastica;strain C | EP00003 Mantamonas plastica 27868 | 0,00026 | 1034 | XP 005831033.1 | secretory protein Sec31 [Guillardia theta CCMP2712] | 1081 | 1,06E-139 | 456,833 | 50,20689655 |
| EP00003 | CRuMs;Mantamonas;Mantamonas plastica;strain C | EP00003 Mantamonas plastica 27870 | 0,00026 | 1034 | XP 005831033.1 | secretory protein Sec31 [Guillardia theta CCMP2712] | 1081 | 1,06E-139 | 456,833 | 50,20689655 |
| EP00050 | Obazoa;Opisthokonta;Holozoa;Choanozoa;Choanof | EP00050 Salpingoeca kvevrii 5300 | 0,00026 | 781 | XP 029637693.1 | uncharacterized protein LOC115212971 [Octopus vulgaris] | 339 | 8,96E-15 | 87,4261 | 45,34883721 |
| EP00437 | Sar;Alveolata;Myzozoa;"DP clade";Dinoflagellata;D | EP00437 Effrenium voratum 23612 | 0,00026 | 681 | WP 131928543.1 | Ig-like domain-containing protein [Bibersteinia trehalosi] | 2280 | 7,67E-12 | 80,8777 | 34,56221198 |
| EP00116 | Obazoa;Opisthokonta;Holozoa;Choanozoa;Metazoa | EP00116 Pleurobrachia bachei 1600 | 0,00026 | 902 | XP 020375845.1 | chromodomain-helicase-DNA-binding protein 3-like, partial [Rhincodon typus] | 644 | 6,09E-69 | 251,521 | 50,13333333 |
| EP00562 | Sar;Stramenopiles;Gyrista;Ochrophyta;"BD clade";C | EP00562 Extubocellulus spinifer 20498 | 0,00026 | 1221 | XP 003080029.1 | Putative 5-3 exonuclease [Ostreococcus tauri] | 1057 | 0 | 599,742 | 54,24836601 |
| EP00682 | Discoba;Euglenozoa;Kinetoplastea;Metakinetoplast | EP00682 Trypanosoma cruzi 5195 | 0,00026 | 550 | XP 807732.1 | lectin [Trypanosoma cruzi strain CL Brener] | 550 | 0 | 1107,82 | 100 |
| EP00429 | Sar;Alveolata;Myzozoa;"DP clade";Dinoflagellata;D | EP00429 Kryptoperidinium foliaceum 35204 | 0,00026 | 508 | ***no hit found*** | | | | | |
| EP00096 | Obazoa;Opisthokonta;Holozoa;Choanozoa;Metazoa | EP00096 Caligus rogercresseyi 16769 | 0,00026 | 711 | XP 022127448.1 | tudor domain-containing protein 3 [Pieris rapae] | 684 | 3,40E-48 | 191,43 | 47,30113636 |
| EP00770 | Metamonada;Preaxostyla;Oxymonadida;Monocerc | EP00770 Monocercomonoides exilis 6814 | 0,00027 | 1743 | ***no hit found*** | | | | | |
| EP00628 | Sar;Stramenopiles;Gyrista;Ochrophyta;Pelagophyce | EP00628 Pelagophyceae sp CCMP2097 27761 | 0,00027 | 623 | XP 009042925.1 | hypothetical protein AURANDRAFT 68941, partial [Aureococcus anophagefferens] | 265 | 2,89E-21 | 104,375 | 57,59493671 |
| EP00445 | Sar;Alveolata;Myzozoa;"DP clade";Dinoflagellata;D | EP00445 Apocalathium hangoei 41691 | 0,00027 | 654 | YP 009052475.1 | pp1ab [Ball python nidovirus 1] | 8108 | 2,30E-07 | 66,2402 | 33,66336634 |
| EP00294 | Cryptista;Cryptophyceae;Goniomonas;"ma G clade" | EP00294 Goniomonas avonlea 8168 | 0,00028 | 715 | WP 062778875.1 | DNA translocase FtsK [Kluyvera intermedia] | 1374 | 3,42E-13 | 85,1149 | 70,44025157 |
| EP00122 | Obazoa;Opisthokonta;Holozoa;Ichthyosporea;Ichth | EP00122 Pirum gemmata 18191 | 0,00028 | 664 | XP 014045659.1 | PREDICTED: nesprin-2-like isoform X2 [Salmo salar] | 8751 | 4,59E-08 | 68,5514 | 58,2781457 |
| EP00443 | Sar;Alveolata;Myzozoa;"DP clade";Dinoflagellata;D | EP00443 Scrippsiella trochoidea 42692 | 0,00028 | 1080 | ***no hit found*** | | | | | |
| EP00575 | Sar;Stramenopiles;Gyrista;Ochrophyta;"BD clade";C | EP00575 Skeletonema marinoi 35695 | 0,00028 | 1209 | XP 002289786.1 | predicted protein [Thalassiosira pseudonana CCMP1335] | 1191 | 0 | 995,727 | 65,49145299 |
| EP00436 | Sar;Alveolata;Myzozoa;"DP clade";Dinoflagellata;D | EP00436 Durusdinium trenchii 23498 | 0,00028 | 331 | ***no hit found*** | | | | | |
| EP00044 | Obazoa;Opisthokonta;Holozoa;Choanozoa;Choanof | EP00044 Salpingoeca rosetta 5683 | 0,00028 | 4245 | XP 004993116.1 | hypothetical protein PTSG 06227 [Salpingoeca rosetta] | 4245 | 0 | 8616,52 | 100 |
| EP00132 | Obazoa;Opisthokonta;Nucletmycea;Fungi;"core Fur | EP00132 Gonapodya prolifera 1770 | 0,00028 | 551 | ***no hit found*** | | | | | |
| EP00036 | Obazoa;Opisthokonta;Holozoa;Choanozoa;Choanof | EP00036 Acanthoeca sp 10tr 15327 | 0,00028 | 3485 | WP 118443790.1 | carbohydrate-binding protein [Clostridium sp. AF37-5] | 1193 | 7,39E-60 | 239,965 | 46,5034965 |
| EP00426 | Sar;Alveolata;Myzozoa;"DP clade";Dinoflagellata;D | EP00426 Karenia brevis 59937 | 0,00028 | 944 | XP 008908976.1 | hypothetical protein PPTG 13264 [Phytophthora parasitica INRA-310] | 619 | 6,70E-46 | 186,037 | 47,40882917 |
| EP00451 | Sar;Alveolata;Myzozoa;"DP clade";Dinoflagellata;O | EP00451 Oxyrrhis marina 38063 | 0,00028 | 509 | XP 004987934.1 | hypothetical protein PTSG 11010 [Salpingoeca rosetta] | 397 | 1,08E-08 | 68,5514 | 98,14814815 |
| EP00475 | Sar;Rhizaria;Endomyxa;"phytorhiza";Vampyrellida; | EP00475 Leptophrys vorax 36768 | 0,00028 | 1099 | NP 001320897.1 | Transducin/WD40 repeat-like superfamily protein [Arabidopsis thaliana] >gi\|1063684652 | 963 | 1,15E-37 | 164,851 | 39,18495298 |
| EP00003 | CRuMs;Mantamonas;Mantamonas plastica;strain C | EP00003 Mantamonas plastica 27867 | 0,00029 | 1184 | XP 005831033.1 | secretory protein Sec31 [Guillardia theta CCMP2712] | 1081 | 0 | 608,216 | 50,80091533 |
| EP00003 | CRuMs;Mantamonas;Mantamonas plastica;strain C | EP00003 Mantamonas plastica 27871 | 0,00029 | 1184 | XP 005831033.1 | secretory protein Sec31 [Guillardia theta CCMP2712] | 1081 | 0 | 608,216 | 50,80091533 |
| EP00613 | Sar;Stramenopiles;Gyrista;Ochrophyta;Dictyochoph | EP00613 Pedinella sp CCMP2098 13268 | 0,00029 | 884 | XP 025195602.1 | histone-lysine N-methyltransferase 2C-like [Melanaphis sacchari] | 5193 | 6,17E-25 | 124,405 | 97,33333333 |
| EP00359 | Sar;Alveolata;Ciliophora;Postciliodesmatophora;He | EP00359 Climacostomum virens 6826 | 0,00029 | 631 | ***no hit found*** | | | | | |
| EP00040 | Obazoa;Opisthokonta;Holozoa;Choanozoa;Choanof | EP00040 Diaphanoeca grandis 30915 | 0,00029 | 322 | ***no hit found*** | | | | | |
| EP00200 | Archaeplastida;Chloroplastida;Chlorophyta;"core ch | EP00200 Dunaliella tertiolecta 9215 | 0,00029 | 954 | XP 002951275.1 | hypothetical protein VOLCADRAFT 91788 [Volvox carteri f. nagariensis] | 1243 | 5,86E-51 | 206,838 | 55,11363636 |
| EP00376 | Sar;Alveolata;Myzozoa;"AC clade";Apicomplexa;"cc | EP00376 Eimeria tenella 3927 | 0,00029 | 467 | XP 013231660.1 | selenocysteine lyase, putative [Eimeria tenella] | 467 | 0 | 929,087 | 100 |
| EP00445 | Sar;Alveolata;Myzozoa;"DP clade";Dinoflagellata;D | EP00445 Apocalathium hangoei 42763 | 0,0003 | 793 | XP 640040.1 | hypothetical protein DDB G0282421 [Dictyostelium discoideum AX4] | 1398 | 3,13E-19 | 105,145 | 87,5 |
| EP00752 | Sar;Stramenopiles;Gyrista;Ochrophyta;"PX clade";P | EP00752 Nemacystus decipiens 16386 | 0,0003 | 750 | XP 010528722.1 | PREDICTED: general negative regulator of transcription subunit 3-like isoform X2 [Tarenay | 823 | 5,67E-122 | 394,43 | 48,40236686 |
| EP00376 | Sar;Alveolata;Myzozoa;"AC clade";Apicomplexa;"cc | EP00376 Eimeria tenella 1751 | 0,0003 | 952 | XP 013229477.1 | hypothetical protein, conserved [Eimeria tenella] | 952 | 0 | 1845,86 | 100 |
| EP00387 | Sar;Alveolata;Myzozoa;"AC clade";Colpodellida;Chr | EP00387 Chromera velia 17160 | 0,00031 | 413 | XP 018638621.1 | poly(ADP-ribose) polymerase and DNA-Ligase Zn-finger region domain-containing protein | 392 | 5,62E-13 | 80,8777 | 33,81924198 |
| EP00201 | Archaeplastida;Chloroplastida;Chlorophyta;"core ch | EP00201 Polytomella parva 11440 | 0,00031 | 848 | ***no hit found*** | | | | | |
| EP00442 | Sar;Alveolata;Myzozoa;"DP clade";Dinoflagellata;D | EP00442 Polarella glacialis 1438 | 0,00031 | 932 | XP 019180959.1 | PREDICTED: ubiquitin carboxyl-terminal hydrolase 2-like [Ipomoea nil] | 1013 | 2,65E-19 | 105,531 | 51,95530726 |
| EP00044 | Obazoa;Opisthokonta;Holozoa;Choanozoa;Choanof | EP00044 Salpingoeca rosetta 9132 | 0,00031 | 1592 | XP 004995565.1 | hypothetical protein PTSG 11834 [Salpingoeca rosetta] | 1592 | 0 | 3311,93 | 100 |
| EP00185 | Archaeplastida;"RR clade";Rhodophyta;Proteorhod | EP00185 Porphyridium purpureum 1959 | 0,00032 | 455 | XP 005707237.1 | hypothetical protein Gasu 19590 [Galdieria sulphuraria] | 286 | 6,84E-24 | 110,923 | 45,8041958 |
| EP00101 | Obazoa;Opisthokonta;Holozoa;Choanozoa;Metazoa | EP00101 Apis mellifera 19529 | 0,00032 | 813 | XP 026301097.1 | alpha-protein kinase 1 isoform X1 [Apis mellifera] | 813 | 0 | 1639,01 | 100 |
| EP00194 | Archaeplastida;Chloroplastida;Chlorophyta;"core ch | EP00194 Tetraselmis striata 2141 | 0,00033 | 1784 | XP 005644893.1 | hypothetical protein COCSUDRAFT 83549 [Coccomyxa subellipsoidea C-169] | 3057 | 3,41E-78 | 300,827 | 50,42527339 |
| EP00023 | Amoebozoa;Evosea;Eumycetozoa;Dictyostelia;Dicty | EP00023 Dictyostelium discoideum 3134 | 0,00033 | 1697 | XP 636616.1 | sterol glucosyltransferase [Dictyostelium discoideum AX4] | 1697 | 0 | 3462,16 | 100 |
| EP00044 | Obazoa;Opisthokonta;Holozoa;Choanozoa;Choanof | EP00044 Salpingoeca rosetta 6866 | 0,00034 | 1000 | XP 004994299.1 | hypothetical protein PTSG 12234 [Salpingoeca rosetta] | 1000 | 0 | 2070,05 | 100 |
| EP00589 | Sar;Stramenopiles;Gyrista;Ochrophyta;"BD clade";C | EP00589 Leptocylindrus danicus 11872 | 0,00034 | 604 | ***no hit found*** | | | | | |
| EP00290 | Cryptista;Cryptophyceae;Cryptomonadales;Baffinel | EP00290 Baffinella sp CCMP2293 53963 | 0,00034 | 531 | XP 005830393.1 | hypothetical protein GUITHDRAFT 140464 [Guillardia theta CCMP2712] | 914 | 3,74E-51 | 198,364 | 50,26595745 |
| EP00025 | Amoebozoa;Evosea;Eumycetozoa;Dictyostelia;Dicty | EP00025 Tieghemostelium lacteum 4652 | 0,00034 | 331 | XP 003293362.1 | hypothetical protein DICPUDRAFT 83923 [Dictyostelium purpureum] | 376 | 2,92E-15 | 86,2705 | 35,41666667 |
| EP00104 | Obazoa;Opisthokonta;Holozoa;Choanozoa;Metazoa | EP00104 Lingula anatina 36664 | 0,00035 | 622 | XP 013421201.1 | chitinase-like protein 4 [Lingula anatina] | 622 | 0 | 1297,72 | 100 |
| EP00023 | Amoebozoa;Evosea;Eumycetozoa;Dictyostelia;Dicty | EP00023 Dictyostelium discoideum 3792 | 0,00036 | 1898 | XP 637315.1 | ubiquitin-protein ligase domain-containing protein [Dictyostelium discoideum AX4] | 1898 | 0 | 3848,13 | 100 |
| EP00024 | Amoebozoa;Evosea;Eumycetozoa;Dictyostelia;Dicty | EP00024 Dictyostelium purpureum 9037 | 0,00036 | 1304 | XP 003291889.1 | hypothetical protein DICPUDRAFT 99174 [Dictyostelium purpureum] | 1304 | 0 | 2620,88 | 100 |
| EP00425 | Sar;Alveolata;Myzozoa;"DP clade";Dinoflagellata;D | EP00425 Heterocapsa triquetra 4858 | 0,00036 | 640 | ***no hit found*** | | | | | |
| EP00645 | Sar;Stramenopiles;Gyrista;Peronosporomycetes;"cc | EP00645 Phytophthora infestans 3484 | 0,00036 | 839 | XP 002898182.1 | conserved hypothetical protein [Phytophthora infestans T30-4] | 839 | 0 | 1604,73 | 100 |
| EP00473 | Sar;Rhizaria;Endomyxa;"phytorhiza";Phytomyxea;"( | EP00473 Plasmodiophora brassicae 2440 | 0,00036 | 488 | XP 004342616.1 | hypothetical protein CAOG 08015 [Capsaspora owczarzaki ATCC 30864] | 355 | 6,39E-48 | 178,718 | 34,91620112 |
| EP00512 | Sar;Stramenopiles;Bigyra;Sagenista;Labyrinthulom | EP00512 Aurantiochytrium limacinum 13940 | 0,00037 | 1239 | ***no hit found*** | | | | | |
| EP00334 | Sar;Alveolata;Ciliophora;Intramacronucleata;"CON | EP00334 Paramecium tetraurelia 29803 | 0,00037 | 327 | XP 001452811.1 | hypothetical protein [Paramecium tetraurelia strain d4-2] | 327 | 0 | 649,432 | 100 |
| EP00751 | Sar;Stramenopiles;Gyrista;Ochrophyta;"BD clade";C | EP00751 Fragilariopsis kerguelensis 28840 | 0,00038 | 903 | XP 002290400.1 | ankyrin domain-containing protein, partial [Thalassiosira pseudonana CCMP1335] | 139 | 1,11E-24 | 110,923 | 47,44525547 |
| EP00435 | Sar;Alveolata;Myzozoa;"DP clade";Dinoflagellata;D | EP00435 Cladocopium sp Y103 75561 | 0,00038 | 1774 | WP 133138619.1 | hypothetical protein [Legionella sp. 2055-AUS-E] | 2195 | 3,04E-10 | 77,411 | 41,53846154 |
| EP00730 | Obazoa;Opisthokonta;Holozoa;Choanozoa;Choanof | EP00730 Choanoeca flexa 3934 | 0,00039 | 797 | XP 001746649.1 | hypothetical protein [Monosiga brevicollis MX1] | 1562 | 5,75E-121 | 408,683 | 56,65024631 |
| EP00628 | Sar;Stramenopiles;Gyrista;Ochrophyta;Pelagophyce | EP00628 Pelagophyceae sp CCMP2097 18807 | 0,00039 | 1191 | XP 009038384.1 | hypothetical protein AURANDRAFT 28308, partial [Aureococcus anophagefferens] | 728 | 0 | 785,408 | 66,21253406 |
| EP00484 | Sar;Rhizaria;Retaria;Foraminifera;Globothalamea;R | EP00484 Ammonia sp Unknown 2258 | 0,00039 | 585 | WP 129326415.1 | YSIRK-type signal peptide-containing protein [Streptococcus oralis] | 1923 | 2,67E-19 | 103,99 | 61,84971098 |
| EP00532 | Sar;Stramenopiles;Gyrista;Ochrophyta;"BD clade";C | EP00532 Pseudo-nitzschia australis 17721 | 0,00039 | 841 | XP 002180574.1 | predicted protein [Phaeodactylum tricornutum CCAP 1055/1] | 915 | 2,13E-120 | 395,586 | 56,26740947 |
| EP00099 | Obazoa;Opisthokonta;Holozoa;Choanozoa;Metazoa | EP00099 Drosophila melanogaster 25366 | 0,00039 | 1241 | NP 724716.1 | secretory 31, isoform B [Drosophila melanogaster] | 1241 | 0 | 2560,41 | 100 |
| EP00441 | Sar;Alveolata;Myzozoa;"DP clade";Dinoflagellata;D | EP00441 Pelagodinium beii 472 | 0,00039 | 674 | ***no hit found*** | | | | | |
| EP00674 | Discoba;Euglenozoa;Kinetoplastea;Metakinetoplast | EP00674 Leishmania major 2050 | 0,00039 | 1321 | XP 001682826.1 | conserved hypothetical protein [Leishmania major strain Friedlin] | 1321 | 0 | 2692,53 | 100 |
| EP00196 | Archaeplastida;Chloroplastida;Chlorophyta;"core ch | EP00196 Chlamydomonas euryale 18939 | 0,0004 | 986 | YP 009052475.1 | pp1ab [Ball python nidovirus 1] | 8108 | 1,46E-07 | 67,781 | 35,32338308 |
| EP00024 | Amoebozoa;Evosea;Eumycetozoa;Dictyostelia;Dicty | EP00024 Dictyostelium purpureum 4795 | 0,0004 | 711 | XP 003287647.1 | hypothetical protein DICPUDRAFT 97778 [Dictyostelium purpureum] | 711 | 0 | 1432,54 | 100 |
| EP00484 | Sar;Rhizaria;Retaria;Foraminifera;Globothalamea;R | EP00484 Ammonia sp Unknown 1069 | 0,0004 | 893 | XP 019864687.1 | PREDICTED: LOW QUALITY PROTEIN: programmed cell death 6-interacting protein [Aethin | 839 | 5,27E-65 | 244,588 | 34,05221339 |
| EP00376 | Sar;Alveolata;Myzozoa;"AC clade";Apicomplexa;"cc | EP00376 Eimeria tenella 7174 | 0,00041 | 809 | XP 013234912.1 | hypothetical protein, conserved [Eimeria tenella] | 809 | 0 | 1622,45 | 100 |
| EP00044 | Obazoa;Opisthokonta;Holozoa;Choanozoa;Choanof | EP00044 Salpingoeca rosetta 2519 | 0,00041 | 288 | XP 004989952.1 | hypothetical protein PTSG 09522 [Salpingoeca rosetta] | 288 | 0 | 569,696 | 100 |
| EP00613 | Sar;Stramenopiles;Gyrista;Ochrophyta;Dictyochoph | EP00613 Pedinella sp CCMP2098 65103 | 0,00041 | 522 | ***no hit found*** | | | | | |
| EP00405 | Sar;Alveolata;Myzozoa;"DP clade";Dinoflagellata;D | EP00405 Crypthecodinium cohnii 42139 | 0,00041 | 916 | XP 008610876.1 | anaphase-promoting complex subunit 3 [Saprolegnia diclina VS20] | 597 | 5,94E-99 | 331,643 | 45,95469256 |
| EP00408 | Sar;Alveolata;Myzozoa;"DP clade";Dinoflagellata;D | EP00408 Alexandrium catenella 2125 | 0,00042 | 594 | ***no hit found*** | | | | | |
| EP00159 | Obazoa;Opisthokonta;Nucletmycea;Rotosphaerida; | EP00159 Fonticula alba 5355 | 0,00042 | 2756 | XP 009497471.1 | hypothetical protein H696 05349 [Fonticula alba] | 2756 | 0 | 5182,08 | 100 |
| EP00393 | Sar;Alveolata;Myzozoa;"DP clade";Dinoflagellata;"c | EP00393 Amoebophrya sp ex-Akashiwo-sanguinea 105073 | 0,00043 | 264 | XP 002772264.1 | hypothetical protein Pmar PMAR023065 [Perkinsus marinus ATCC 50983] | 173 | 1,83E-07 | 60,077 | 31,3559322 |
| EP00437 | Sar;Alveolata;Myzozoa;"DP clade";Dinoflagellata;D | EP00437 Effrenium voratum 55345 | 0,00043 | 90 | ***no hit found*** | | | | | |
| EP00428 | Sar;Alveolata;Myzozoa;"DP clade";Dinoflagellata;D | EP00428 Durinskia baltica 13331 | 0,00044 | 892 | XP 002779439.1 | replication protein A 70kDa, putative [Perkinsus marinus ATCC 50983] | 788 | 5,15E-55 | 214,927 | 39,95535714 |
| EP00697 | Hemimastigophora;Spironemidae;Spironema;strain | EP00697 Spironema sp BW2 15250 | 0,00044 | 546 | XP 643680.1 | hypothetical protein DDB G0275359 [Dictyostelium discoideum AX4] | 462 | 1,28E-60 | 216,853 | 36,99788584 |
| EP00200 | Archaeplastida;Chloroplastida;Chlorophyta;"core ch | EP00200 Dunaliella tertiolecta 14295 | 0,00045 | 380 | ***no hit found*** | | | | | |
| EP00044 | Obazoa;Opisthokonta;Holozoa;Choanozoa;Choanof | EP00044 Salpingoeca rosetta 846 | 0,00045 | 1126 | XP 004988279.1 | hypothetical protein PTSG 13115 [Salpingoeca rosetta] | 1126 | 0 | 2277,67 | 100 |
| EP00445 | Sar;Alveolata;Myzozoa;"DP clade";Dinoflagellata;D | EP00445 Apocalathium hangoei 93376 | 0,00046 | 504 | XP 015519886.1 | PREDICTED: uncharacterized protein LOC107224373 [Neodiprion lecontei] | 671 | 2,74E-12 | 80,4925 | 96,07843137 |
| EP00532 | Sar;Stramenopiles;Gyrista;Ochrophyta;"BD clade";C | EP00532 Pseudo-nitzschia australis 8431 | 0,00046 | 557 | XP 002287857.1 | predicted protein [Thalassiosira pseudonana CCMP1335] | 500 | 2,44E-51 | 193,356 | 55,50458716 |
| EP00731 | Obazoa;Opisthokonta;Holozoa;Choanozoa;Metazoa | EP00731 Ephydatia muelleri 13340 | 0,00046 | 473 | XP 002947210.1 | hypothetical protein VOLCADRAFT 120422 [Volvox carteri f. nagariensis] | 1283 | 1,79E-18 | 100,138 | 65,68627451 |
| EP00057 | Obazoa;Opisthokonta;Holozoa;Choanozoa;Metazoa | EP00057 Strongylocentrotus purpuratus 12992 | 0,00046 | 1003 | XP 011667466.1 | PREDICTED: mucin-22-like [Strongylocentrotus purpuratus] | 1003 | 0 | 1715,66 | 100 |
| EP00099 | Obazoa;Opisthokonta;Holozoa;Choanozoa;Metazoa | EP00099 Drosophila melanogaster 27028 | 0,00047 | 1517 | NP 728554.1 | ubiquitin specific protease 10, isoform A [Drosophila melanogaster] >gi\|442629288\|ref\|N | 1517 | 0 | 3062,32 | 100 |
| EP00099 | Obazoa;Opisthokonta;Holozoa;Choanozoa;Metazoa | EP00099 Drosophila melanogaster 8512 | 0,00047 | 1517 | NP 728554.1 | ubiquitin specific protease 10, isoform A [Drosophila melanogaster] >gi\|442629288\|ref\|N | 1517 | 0 | 3062,32 | 100 |
| EP00101 | Obazoa;Opisthokonta;Holozoa;Choanozoa;Metazoa | EP00101 Apis mellifera 4611 | 0,00047 | 2947 | XP 006562479.2 | zinc finger homeobox protein 4 isoform X3 [Apis mellifera] | 2947 | 0 | 6072,66 | 100 |
| EP00024 | Amoebozoa;Evosea;Eumycetozoa;Dictyostelia;Dicty | EP00024 Dictyostelium purpureum 1222 | 0,00047 | 374 | XP 003284074.1 | hypothetical protein DICPUDRAFT 75027 [Dictyostelium purpureum] | 374 | 0 | 769,229 | 100 |
| EP00563 | Sar;Stramenopiles;Gyrista;Ochrophyta;"BD clade";C | EP00563 Minutocellus polymorphus 5356 | 0,00047 | 68 | ***no hit found*** | | | | | |
| EP00419 | Sar;Alveolata;Myzozoa;"DP clade";Dinoflagellata;D | EP00419 Tripos fusus 52243 | 0,00048 | 616 | XP 004988696.1 | hypothetical protein PTSG 10731 [Salpingoeca rosetta] | 481 | 3,67E-10 | 73,9442 | 33,70786517 |
| EP00200 | Archaeplastida;Chloroplastida;Chlorophyta;"core ch | EP00200 Dunaliella tertiolecta 11020 | 0,00048 | 2298 | ***no hit found*** | | | | | |
| EP00480 | Sar;Rhizaria;Retaria;Foraminifera;"monothalamid c | EP00480 Reticulomyxa filosa 31996 | 0,00049 | 493 | XP 015253920.1 | PREDICTED: snurportin-1 [Cyprinodon variegatus] >gi\|974109575\|ref\|XP 015253921.1\| I | 371 | 6,00E-35 | 144,436 | 47,71929825 |
| EP00044 | Obazoa;Opisthokonta;Holozoa;Choanozoa;Choanof | EP00044 Salpingoeca rosetta 7793 | 0,00049 | 1379 | XP 004995226.1 | hypothetical protein PTSG 03494 [Salpingoeca rosetta] | 1379 | 0 | 2766,1 | 100 |
| EP00429 | Sar;Alveolata;Myzozoa;"DP clade";Dinoflagellata;D | EP00429 Kryptoperidinium foliaceum 61183 | 0,00049 | 305 | ***no hit found*** | | | | | |
| EP00500 | Sar;Stramenopiles;Bigyra;Opalozoa;Bicosoecida;str | EP00500 Bicosoecida sp ms1 8688 | 0,00049 | 53 | ***no hit found*** | | | | | |
| EP00044 | Obazoa;Opisthokonta;Holozoa;Choanozoa;Choanof | EP00044 Salpingoeca rosetta 7730 | 0,00049 | 572 | XP 004995163.1 | hypothetical protein PTSG 12144 [Salpingoeca rosetta] | 572 | 0 | 1166,76 | 100 |
| EP00287 | Cryptista;Cryptophyceae;Cryptomonadales;"lineage | EP00287 Rhodomonas sp CCMP768 28463 | 0,00049 | 584 | XP 005823393.1 | hypothetical protein GUITHDRAFT 89990 [Guillardia theta CCMP2712] | 295 | 7,54E-153 | 449,899 | 77,41935484 |

| | | | | | | | | | | |
|---|---|---|---|---|---|---|---|---|---|---|
| EP00618 | Sar;Stramenopiles;Gyrista;Ochrophyta;Dictyochoph | EP00618 Florenciella parvula 39274 | 0,0005 | 133 | ***no hit found*** | | | | | |
| EP00435 | Sar;Alveolata;Myzozoa;"DP clade";Dinoflagellata;D | EP00435 Cladocopium sp Y103 52111 | 0,0005 | 382 | ***no hit found*** | | | | | |
| EP00201 | Archaeplastida;Chloroplastida;Chlorophyta;"core ch | EP00201 Polytomella parva 19074 | 0,0005 | 615 | XP 002946119.1 | hypothetical protein VOLCADRAFT 115694 [Volvox carteri f. nagariensis] | 577 | 5,99E-60 | 219,935 | 58,92857143 |
| EP00120 | Obazoa;Opisthokonta;Holozoa;Filasterea;Capsaspor | EP00120 Capsaspora owczarzaki 5005 | 0,0005 | 2246 | XP 004349759.1 | thyroid hormone receptor interactor 12 [Capsaspora owczarzaki ATCC 30864] | 2246 | 0 | 4506,05 | 100 |
| EP00106 | Obazoa;Opisthokonta;Holozoa;Choanozoa;Metazoa | EP00106 Octopus bimaculoides 19580 | 0,0005 | 1116 | XP 014787022.1 | PREDICTED: ubiquitin carboxyl-terminal hydrolase 25-like isoform X6 [Octopus bimaculoid | 1116 | 0 | 2332,37 | 100 |
| EP00331 | Sar;Alveolata;Ciliophora;Intramacronucleata;"CON: | EP00331 Platyophrya macrostoma 15793 | 0,00051 | 422 | XP 009607598.1 | PREDICTED: ubiquitin receptor RAD23c-like isoform X2 [Nicotiana tomentosiformis] >gi\|1( | 365 | 7,06E-64 | 218,779 | 53,125 |
| EP00331 | Sar;Alveolata;Ciliophora;Intramacronucleata;"CON: | EP00331 Platyophrya macrostoma 16825 | 0,00051 | 422 | XP 009607598.1 | PREDICTED: ubiquitin receptor RAD23c-like isoform X2 [Nicotiana tomentosiformis] >gi\|1( | 365 | 7,06E-64 | 218,779 | 53,125 |
| EP00751 | Sar;Stramenopiles;Gyrista;Ochrophyta;"BD clade";[ | EP00751 Fragilariopsis kerguelensis 35663 | 0,00051 | 624 | XP 002185439.1 | predicted protein [Phaeodactylum tricornutum CCAP 1055/1] | 698 | 3,24E-11 | 77,7962 | 48,38709677 |
| EP00201 | Archaeplastida;Chloroplastida;Chlorophyta;"core ch | EP00201 Polytomella parva 5630 | 0,00054 | 508 | XP 002946119.1 | hypothetical protein VOLCADRAFT 115694 [Volvox carteri f. nagariensis] | 577 | 2,86E-61 | 220,32 | 58,92857143 |
| EP00309 | Haptista;Haptophyta;Prymnesiophyceae;Calcihapto | EP00309 Chrysotila carterae 17361 | 0,00054 | 819 | XP 005791037.1 | hypothetical protein EMIHUDRAFT 224393 [Emiliania huxleyi CCMP1516] | 384 | 4,13E-21 | 107,071 | 53,99239544 |
| EP00302 | Haptista;Haptophyta;Pavlovales;Diacronema;strain | EP00302 Diacronema sp CCMP2436 44026 | 0,00054 | 433 | XP 002784418.1 | conserved hypothetical protein [Perkinsus marinus ATCC 50983] | 384 | 8,78E-30 | 129,413 | 41,96078431 |
| EP00451 | Sar;Alveolata;Myzozoa;"DP clade";Dinoflagellata;O | EP00451 Oxyrrhis marina 40470 | 0,00055 | 834 | WP 096699322.1 | LPXTG cell wall anchor domain-containing protein [Brochothrix thermosphacta] | 525 | 2,03E-30 | 137,502 | 54,4 |
| EP00682 | Discoba;Euglenozoa;Kinetoplastea;Metakinetoplast | EP00682 Trypanosoma cruzi 17676 | 0,00055 | 1232 | XP 820213.1 | protein transport protein sec31 [Trypanosoma cruzi strain CL Brener] | 1232 | 0 | 2531,9 | 100 |
| EP00266 | Archaeplastida;Chloroplastida;Streptophyta;"core s | EP00266 Sorghum bicolor 9088 | 0,00055 | 782 | XP 002453383.1 | protein STRUBBELIG-RECEPTOR FAMILY 3 isoform X1 [Sorghum bicolor] | 782 | 0 | 1578,15 | 100 |
| EP00102 | Obazoa;Opisthokonta;Holozoa;Choanozoa;Metazoa | EP00102 Acyrthosiphon pisum 11196 | 0,00056 | 2197 | XP 008179841.2 | uncharacterized protein LOC100165959 isoform X1 [Acyrthosiphon pisum] | 2197 | 0 | 4421,31 | 100 |
| EP00574 | Sar;Stramenopiles;Gyrista;Ochrophyta;"BD clade";[ | EP00574 Skeletonema japonicum 5510 | 0,00056 | 1004 | XP 002289786.1 | predicted protein [Thalassiosira pseudonana CCMP1335] | 1191 | 0 | 810,061 | 64,32160804 |
| EP00597 | Sar;Stramenopiles;Gyrista;Ochrophyta;"CS clade";C | EP00597 Dinobryon sp UTEXLB2267 10890 | 0,00056 | 165 | ***no hit found*** | | | | | |
| EP00044 | Obazoa;Opisthokonta;Holozoa;Choanozoa;Choanof | EP00044 Salpingoeca rosetta 138 | 0,00058 | 1885 | XP 004987571.1 | WD-40 repeat protein [Salpingoeca rosetta] | 1885 | 0 | 3904,37 | 100 |
| EP00435 | Sar;Alveolata;Myzozoa;"DP clade";Dinoflagellata;D | EP00435 Cladocopium sp Y103 10739 | 0,00058 | 603 | XP 023148314.1 | procollagen galactosyltransferase 2-like [Amphiprion ocellaris] | 481 | 4,41E-17 | 95,5153 | 45,13274336 |
| EP00146 | Obazoa;Opisthokonta;Nucletmycea;Fungi;"core Fur | EP00146 Coprinopsis cinerea 4081 | 0,00059 | 977 | XP 001832994.2 | hypothetical protein CC1G 01056 [Coprinopsis cinerea okayama7#130] | 977 | 0 | 2034,61 | 100 |
| EP00754 | Discoba;Euglenozoa;Diplonemea;Diplonemidae;Rhy | EP00754 Rhynchopus humris 20361 | 0,00059 | 546 | ***no hit found*** | | | | | |
| EP00202 | Archaeplastida;Chloroplastida;Chlorophyta;"core ch | EP00202 Volvox carteri 2681 | 0,00059 | 2760 | XP 002948287.1 | hypothetical protein VOLCADRAFT 88480 [Volvox carteri f. nagariensis] | 2760 | 0 | 5492,93 | 100 |
| EP00023 | Amoebozoa;Evosea;Eumycetozoa;Dictyostelia;Dicty | EP00023 Dictyostelium discoideum 6306 | 0,0006 | 1751 | XP 640439.1 | hypothetical protein DDB G0282057 [Dictyostelium discoideum AX4] | 1751 | 0 | 3459,46 | 100 |
| EP00120 | Obazoa;Opisthokonta;Holozoa;Filasterea;Capsaspor | EP00120 Capsaspora owczarzaki 7383 | 0,0006 | 2048 | XP 004365651.2 | hypothetical protein CAOG 00780 [Capsaspora owczarzaki ATCC 30864] | 2048 | 0 | 4182,48 | 100 |
| EP00331 | Sar;Alveolata;Ciliophora;Intramacronucleata;"CON: | EP00331 Platyophrya macrostoma 13575 | 0,0006 | 712 | XP 001027006.2 | ubiquitin family protein [Tetrahymena thermophila SB210] | 1021 | 7,76E-19 | 103,219 | 43,2 |
| EP00578 | Sar;Stramenopiles;Gyrista;Ochrophyta;"BD clade";[ | EP00578 Thalassiosira sp NH16 8863 | 0,00061 | 1120 | XP 002286990.1 | predicted protein [Thalassiosira pseudonana CCMP1335] | 913 | 0 | 565,074 | 68,53625171 |
| EP00099 | Obazoa;Opisthokonta;Holozoa;Choanozoa;Metazoa | EP00099 Drosophila melanogaster 10199 | 0,00061 | 618 | NP 001262959.1 | musashi, isoform E [Drosophila melanogaster] | 618 | 0 | 1237,63 | 100 |
| EP00290 | Cryptista;Cryptophyceae;Cryptomonadales;Baffinel | EP00290 Baffinella sp CCMP2293 6043 | 0,00061 | 560 | XP 005825898.1 | hypothetical protein GUITHDRAFT 115021 [Guillardia theta CCMP2712] | 663 | 6,37E-32 | 141,354 | 58,33333333 |
| EP00580 | Sar;Stramenopiles;Gyrista;Ochrophyta;"BD clade";[ | EP00580 Thalassiosira gravida 14543 | 0,00062 | 2089 | XP 002290207.1 | THAPSDRAFT 262487, partial [Thalassiosira pseudonana CCMP1335] | 466 | 0 | 624,009 | 77,65765766 |
| EP00615 | Sar;Stramenopiles;Gyrista;Ochrophyta;Dictyochoph | EP00615 Pteridomonas danica 13748 | 0,00062 | 617 | XP 002999298.1 | replication protein A 70 kDa DNA-binding subunit, putative [Phytophthora infestans T30-4 | 629 | 6,31E-120 | 379,022 | 40,89456869 |
| EP00426 | Sar;Alveolata;Myzozoa;"DP clade";Dinoflagellata;D | EP00426 Karenia brevis 70147 | 0,00062 | 431 | ***no hit found*** | | | | | |
| EP00194 | Archaeplastida;Chloroplastida;Chlorophyta;"core ch | EP00194 Tetraselmis striata 6163 | 0,00062 | 346 | ***no hit found*** | | | | | |
| EP00256 | Archaeplastida;Chloroplastida;Streptophyta;"core s | EP00256 Glycine max 35113 | 0,00063 | 887 | XP 006581383.1 | transcriptional corepressor LEUNIG isoform X2 [Glycine max] >gi\|1590532614\|ref\|XP 02{ | 887 | 0 | 1824,68 | 100 |
| EP00334 | Sar;Alveolata;Ciliophora;Intramacronucleata;"CON: | EP00334 Paramecium tetraurelia 18063 | 0,00064 | 1200 | XP 001441053.1 | hypothetical protein [Paramecium tetraurelia strain d4-2] | 1200 | 0 | 2396,31 | 100 |
| EP00376 | Sar;Alveolata;Myzozoa;"AC clade";Apicomplexa;"cc | EP00376 Eimeria tenella 2486 | 0,00065 | 472 | XP 013230213.1 | hypothetical protein, conserved, partial [Eimeria tenella] | 472 | 0 | 876,7 | 100 |
| EP00158 | Obazoa;Opisthokonta;Nucletmycea;Fungi;Aphelide; | EP00158 Paraphelidium tribonemae 9301 | 0,00065 | 1197 | XP 021881067.1 | hypothetical protein BCR41DRAFT 396657 [Lobosporangium transversale] | 1406 | 0 | 591,652 | 50 |
| EP00442 | Sar;Alveolata;Myzozoa;"DP clade";Dinoflagellata;D | EP00442 Polarella glacialis 17195 | 0,00066 | 884 | XP 002781363.1 | eukaryotic initiation factor 4f, putative [Perkinsus marinus ATCC 50983] | 1474 | 1,16E-56 | 223,787 | 43,63636364 |
| EP00013 | Amoebozoa;Discosea;Flabellinia;Stygamoeba;Styga | EP00013 Stygamoeba regulata 29492 | 0,00066 | 1230 | XP 025175506.1 | hypothetical protein GLOIN 2v1633885 [Rhizophagus irregularis DAOM 181602=DAOM 1 | 1409 | 5,57E-106 | 375,941 | 45,81124072 |
| EP00388 | Sar;Alveolata;Myzozoa;"AC clade";Colpodellida;Col | EP00388 Colpodella angusta 42248 | 0,00068 | 552 | XP 005648604.1 | eukaryotic translation initiation factor 4E [Coccomyxa subellipsoidea C-169] | 218 | 9,73E-23 | 107,071 | 42,95774648 |
| EP00705 | Metamonada;Parabasalia;Trichomonadida;Trichom | EP00705 Trichomonas vaginalis 28625 | 0,00068 | 478 | XP 001305425.1 | hypothetical protein [Trichomonas vaginalis G3] | 478 | 0 | 984,941 | 100 |
| EP00021 | Amoebozoa;Evosea;Eumycetozoa;Dictyostelia;Acyto | EP00021 Heterostelium album 5409 | 0,00068 | 1221 | XP 020432045.1 | hypothetical protein PPL 06745 [Heterostelium album PN500] | 1221 | 0 | 2470,66 | 100 |
| EP00444 | Sar;Alveolata;Myzozoa;"DP clade";Dinoflagellata;D | EP00444 Apocalathium aciculiferum 53398 | 0,00068 | 647 | WP 041084212.1 | RluA family pseudouridine synthase [Thermotoga profunda] | 303 | 1,97E-20 | 102,834 | 55,55555556 |
| EP00044 | Obazoa;Opisthokonta;Holozoa;Choanozoa;Choanof | EP00044 Salpingoeca rosetta 1267 | 0,00068 | 1097 | XP 004988700.1 | hypothetical protein PTSG 10735 [Salpingoeca rosetta] | 1097 | 0 | 2207,56 | 100 |
| EP00738 | Archaeplastida;Chloroplastida;Chlorophyta;Chlorop | EP00738 Chloropicon primus 831 | 0,00068 | 671 | XP 002955457.1 | hypothetical protein VOLCADRAFT 121426 [Volvox carteri f. nagariensis] | 662 | 8,81E-159 | 481,871 | 64,46808511 |
| EP00250 | Archaeplastida;Chloroplastida;Streptophyta;"core s | EP00250 Pteridium aquilinum 20126 | 0,00069 | 581 | XP 024363811.1 | pre-mRNA-processing factor 39-like isoform X4 [Physcomitrella patens] | 773 | 6,75E-174 | 521,161 | 57,28643216 |
| EP00250 | Archaeplastida;Chloroplastida;Streptophyta;"core s | EP00250 Pteridium aquilinum 20127 | 0,00069 | 581 | XP 024363811.1 | pre-mRNA-processing factor 39-like isoform X4 [Physcomitrella patens] | 773 | 6,75E-174 | 521,161 | 57,28643216 |
| EP00250 | Archaeplastida;Chloroplastida;Streptophyta;"core s | EP00250 Pteridium aquilinum 20128 | 0,00069 | 581 | XP 024363811.1 | pre-mRNA-processing factor 39-like isoform X4 [Physcomitrella patens] | 773 | 6,75E-174 | 521,161 | 57,28643216 |
| EP00334 | Sar;Alveolata;Ciliophora;Intramacronucleata;"CON: | EP00334 Paramecium tetraurelia 21805 | 0,00069 | 547 | XP 001444801.1 | hypothetical protein [Paramecium tetraurelia strain d4-2] | 547 | 0 | 1101,27 | 100 |
| EP00445 | Sar;Alveolata;Myzozoa;"DP clade";Dinoflagellata;D | EP00445 Apocalathium hangoei 10190 | 0,0007 | 604 | XP 003880818.1 | hypothetical protein NCLIV 038600 [Neospora caninum Liverpool] | 1060 | 4,15E-56 | 215,312 | 63,29113924 |
| EP00439 | Sar;Alveolata;Myzozoa;"DP clade";Dinoflagellata;D | EP00439 Symbiodinium sp Y106 63730 | 0,0007 | 520 | ***no hit found*** | | | | | |
| EP00747 | Sar;Alveolata;Myzozoa;"DP clade";Dinoflagellata;D | EP00747 Dinoflagellata sp TGD 167021 | 0,00071 | 248 | XP 002772264.1 | hypothetical protein Pmar PMAR023065 [Perkinsus marinus ATCC 50983] | 173 | 2,35E-10 | 67,781 | 39,74358974 |
| EP00573 | Sar;Stramenopiles;Gyrista;Ochrophyta;"BD clade";[ | EP00573 Skeletonema grethae 10816 | 0,00071 | 598 | XP 002290579.1 | predicted protein [Thalassiosira pseudonana CCMP1335] | 649 | 1,20E-156 | 473,396 | 69,74789916 |
| EP00290 | Cryptista;Cryptophyceae;Cryptomonadales;Baffinel | EP00290 Baffinella sp CCMP2293 55320 | 0,00071 | 743 | XP 005825898.1 | hypothetical protein GUITHDRAFT 115021 [Guillardia theta CCMP2712] | 663 | 1,45E-31 | 142,124 | 58,33333333 |
| EP00008 | Amoebozoa;Discosea;Flabellinia;"core flabellinids"; | EP00008 Paramoeba atlantica 303 | 0,00072 | 1451 | XP 002257643.1 | hypothetical protein, conserved in Plasmodium species [Plasmodium knowlesi strain H] | 911 | 2,89E-10 | 77,0258 | 67,03296703 |
| EP00211 | Archaeplastida;Chloroplastida;Chlorophyta;Chlorop | EP00211 Chloroparvula japonica 4884 | 0,00072 | 953 | XP 005850236.1 | hypothetical protein CHLNCDRAFT 141882 [Chlorella variabilis] | 959 | 1,41E-144 | 464,151 | 43,70860927 |
| EP00445 | Sar;Alveolata;Myzozoa;"DP clade";Dinoflagellata;D | EP00445 Apocalathium hangoei 91951 | 0,00072 | 741 | YP 009052475.1 | pp1ab [Ball python nidovirus 1] | 8108 | 1,78E-10 | 76,6406 | 40,34090909 |
| EP00004 | CRuMs;Rigifilida;Rigifila;Rigifila ramosa;strain CCAF | EP00004 Rigifila ramosa 12236 | 0,00073 | 4004 | XP 012187456.1 | hypothetical protein PHSY 001436 [Pseudozyma hubeiensis SY62] | 578 | 4,41E-168 | 542,732 | 56,54648956 |
| EP00364 | Sar;Alveolata;Myzozoa;"AC clade";Apicomplexa;"cc | EP00364 Plasmodium chabaudi 4849 | 0,00074 | 1766 | XP 745298.2 | merozoite surface protein 1 [Plasmodium chabaudi chabaudi] | 1766 | 0 | 3588,51 | 100 |
| EP00023 | Amoebozoa;Evosea;Eumycetozoa;Dictyostelia;Dicty | EP00023 Dictyostelium discoideum 978 | 0,00074 | 540 | XP 629256.1 | hypothetical protein DDB G0293274 [Dictyostelium discoideum AX4] | 540 | 0 | 1067,37 | 100 |
| EP00751 | Sar;Stramenopiles;Gyrista;Ochrophyta;"BD clade";[ | EP00751 Fragilariopsis kerguelensis 46516 | 0,00076 | 1142 | XP 002176587.1 | beta-glucan elicitor receptor [Phaeodactylum tricornutum CCAP 1055/1] | 1208 | 3,48E-148 | 485,723 | 47,26315789 |
| EP00134 | Obazoa;Opisthokonta;Nucletmycea;Fungi;"core Fur | EP00134 Neurospora crassa 4601 | 0,00076 | 329 | XP 957935.1 | hypothetical protein NCU09837 [Neurospora crassa OR74A] | 329 | 0 | 647,121 | 100 |
| EP00572 | Sar;Stramenopiles;Gyrista;Ochrophyta;"BD clade";[ | EP00572 Skeletonema dohrnii 10549 | 0,00076 | 800 | XP 002289786.1 | predicted protein [Thalassiosira pseudonana CCMP1335] | 1191 | 0 | 574,318 | 60,66115702 |
| EP00304 | Haptista;Haptophyta;Pavlovales;Pavlova;Pavlova gy | EP00304 Pavlova gyrans 12336 | 0,00077 | 151 | XP 005788197.1 | secretory protein 31 [Emiliania huxleyi CCMP1516] | 1166 | 1,08E-30 | 126,331 | 50,80645161 |
| EP00730 | Obazoa;Opisthokonta;Holozoa;Choanozoa;Choanof | EP00730 Choanoeca flexa 3932 | 0,00078 | 829 | XP 001746649.1 | hypothetical protein [Monosiga brevicollis MX1] | 1562 | 8,92E-121 | 409,068 | 56,40394089 |
| EP00403 | Sar;Alveolata;Myzozoa;"DP clade";Dinoflagellata;D | EP00403 Amphidinium massartii 9926 | 0,00078 | 877 | XP 004356759.1 | hypothetical protein ACA1 175330 [Acanthamoeba castellanii str. Neff] | 2283 | 4,70E-90 | 324,709 | 38,37689133 |
| EP00315 | Haptista;Haptophyta;Prymnesiophyceae;Calcihapto | EP00315 Gephyrocapsa oceanica 27680 | 0,00078 | 1227 | XP 005766133.1 | hypothetical protein EMIHUDRAFT 470692 [Emiliania huxleyi CCMP1516] | 943 | 3,76E-161 | 514,998 | 94,61966605 |
| EP00405 | Sar;Alveolata;Myzozoa;"DP clade";Dinoflagellata;D | EP00405 Crypthecodinium cohnii 16599 | 0,00078 | 1173 | ***no hit found*** | | | | | |
| EP00365 | Sar;Alveolata;Myzozoa;"AC clade";Apicomplexa;"cc | EP00365 Plasmodium falciparum 237 | 0,00078 | 1811 | XP 001347549.1 | conserved Plasmodium protein, unknown function [Plasmodium falciparum 3D7] | 1811 | 0 | 3476,41 | 100 |
| EP00023 | Amoebozoa;Evosea;Eumycetozoa;Dictyostelia;Dicty | EP00023 Dictyostelium discoideum 12167 | 0,00079 | 1714 | XP 646638.1 | actin binding protein [Dictyostelium discoideum AX4] | 1714 | 0 | 3360,47 | 100 |
| EP00267 | Archaeplastida;Chloroplastida;Streptophyta;"core s | EP00267 Zea mays 43659 | 0,0008 | 240 | XP 020395775.1 | protein transport protein SEC31 homolog B [Zea mays] | 240 | 3,25E-179 | 500,36 | 100 |
| EP00671 | Discoba;Euglenozoa;Kinetoplastea;Metakinetoplast | EP00671 Bodo saltans 17388 | 0,0008 | 769 | XP 819369.1 | hypothetical protein [Trypanosoma cruzi strain CL Brener] | 639 | 3,90E-59 | 222,246 | 56,06060606 |
| EP00393 | Sar;Alveolata;Myzozoa;"DP clade";Dinoflagellata;"( | EP00393 Amoebophrya sp ex-Akashiwo-sanguinea 93367 | 0,0008 | 476 | ***no hit found*** | | | | | |
| EP00366 | Sar;Alveolata;Myzozoa;"AC clade";Apicomplexa;"cc | EP00366 Plasmodium knowlesi 1085 | 0,0008 | 1821 | XP 002258582.1 | merozoite surface protein 1, MSP-1 [Plasmodium knowlesi strain H] | 1821 | 0 | 3690,97 | 100 |
| EP00359 | Sar;Alveolata;Ciliophora;Postciliodesmatophora;He | EP00359 Climacostomum virens 1465 | 0,00081 | 370 | ***no hit found*** | | | | | |
| EP00279 | Cryptista;Cryptophyceae;Cryptomonadales;"lineage | EP00279 Guillardia theta 1972 | 0,00082 | 345 | XP 005819691.1 | hypothetical protein GUITHDRAFT 121109 [Guillardia theta CCMP2712] | 345 | 0 | 678,707 | 100 |
| EP00045 | Obazoa;Opisthokonta;Holozoa;Choanozoa;Choanof | EP00045 Choanoeca perplexa 11245 | 0,00083 | 1458 | XP 004994900.1 | single-stranded RNA 5'-3' exonuclease [Salpingoeca rosetta] | 1780 | 6,15E-143 | 490,73 | 42,49084249 |
| EP00411 | Sar;Alveolata;Myzozoa;"DP clade";Dinoflagellata;D | EP00411 Alexandrium monilatum 41568 | 0,00083 | 428 | ***no hit found*** | | | | | |
| EP00106 | Obazoa;Opisthokonta;Holozoa;Choanozoa;Metazoa | EP00106 Octopus bimaculoides 7064 | 0,00083 | 734 | XP 014774506.1 | PREDICTED: zinc finger protein 708-like [Octopus bimaculoides] | 734 | 0 | 1509,2 | 100 |
| EP00055 | Obazoa;Opisthokonta;Holozoa;Choanozoa;Choanof | EP00055 Hartaetosiga balthica 2816 | 0,00083 | 847 | XP 029108541.1 | protein transport protein Sec31A isoform X4 [Scleropages formosus] | 1228 | 4,32E-75 | 277,33 | 40,27027027 |
| EP00050 | Obazoa;Opisthokonta;Holozoa;Choanozoa;Choanof | EP00050 Salpingoeca kvevrii 85 | 0,00084 | 689 | WP 118691157.1 | hypothetical protein [Clostridium sp. OM05-6BH] | 981 | 1,11E-21 | 112,079 | 57,71812081 |
| EP00401 | Sar;Alveolata;Myzozoa;"DP clade";Dinoflagellata;D | EP00401 Gymnodinium catenatum 72236 | 0,00084 | 1171 | XP 004988728.1 | hypothetical protein PTSG 10764 [Salpingoeca rosetta] | 523 | 9,50E-20 | 105,531 | 35,53299492 |
| EP00276 | Archaeplastida;Glaucophyta;Gloeochaete;Gloeocha | EP00276 Gloeochaete wittrockiana 12291 | 0,00084 | 995 | ***no hit found*** | | | | | |
| EP00515 | Sar;Stramenopiles;Bigyra;Sagenista;Labyrinthulomy | EP00515 Schizochytrium aggregatum 21191 | 0,00084 | 908 | XP 008868552.1 | nicotinate (nicotinamide) nucleotide adenylyltransferase [Aphanomyces invadans] | 213 | 2,94E-16 | 88,9669 | 48,05194805 |
| EP00183 | Archaeplastida;"RR clade";Rhodophyta;Proteorhod | EP00183 Erythrolobus madagascarensis 4794 | 0,00084 | 922 | ***no hit found*** | | | | | |
| EP00636 | Sar;Stramenopiles;Gyrista;Ochrophyta;Pinguiophyc | EP00636 Phaeomonas parva 4450 | 0,00085 | 84 | ***no hit found*** | | | | | |
| EP00426 | Sar;Alveolata;Myzozoa;"DP clade";Dinoflagellata;D | EP00426 Karenia brevis 82102 | 0,00085 | 273 | XP 003873801.1 | conserved hypothetical protein [Leishmania mexicana MHOM/GT/2001/U1103] | 649 | 1,81E-16 | 89,7373 | 50,74626866 |
| EP00485 | Sar;Rhizaria;Retaria;Foraminifera;Globothalamea;R | EP00485 Elphidium margaritaceum 16686 | 0,00085 | 1399 | XP 027024442.1 | X-linked retinitis pigmentosa GTPase regulator-like isoform X3 [Tachysurus fulvidraco] | 701 | 1,93E-22 | 115,931 | 46,17414248 |
| EP00446 | Sar;Alveolata;Myzozoa;"DP clade";Dinoflagellata;D | EP00446 Apocalathium sp SHHI-4 4144 | 0,00085 | 513 | ***no hit found*** | | | | | |
| EP00457 | Sar;Rhizaria;Cercozoa;"crown cercozoa";Imbricatea | EP00457 Paulinella chromatophora 5313 | 0,00086 | 592 | ***no hit found*** | | | | | |
| EP00044 | Obazoa;Opisthokonta;Holozoa;Choanozoa;Choanof | EP00044 Salpingoeca rosetta 1926 | 0,00086 | 1063 | XP 004989359.1 | hypothetical protein PTSG 09106 [Salpingoeca rosetta] | 1063 | 0 | 2194,47 | 100 |
| EP00393 | Sar;Alveolata;Myzozoa;"DP clade";Dinoflagellata;"( | EP00393 Amoebophrya sp ex-Akashiwo-sanguinea 41850 | 0,00087 | 508 | XP 025127094.1 | microtubule-associated protein 1A isoform X1 [Bubalus bubalis] | 3122 | 1,71E-06 | 62,7734 | 38,36477987 |
| EP00457 | Sar;Rhizaria;Cercozoa;"crown cercozoa";Imbricatea | EP00457 Paulinella chromatophora 291 | 0,00087 | 1672 | YP 009052475.1 | pp1ab [Ball python nidovirus 1] | 8108 | 1,17E-11 | 82,4185 | 35,19736842 |
| EP00592 | Sar;Stramenopiles;Gyrista;Ochrophyta;"BD clade";[ | EP00592 Proboscia alata 15396 | 0,00088 | 195 | ***no hit found*** | | | | | |
| EP00376 | Sar;Alveolata;Myzozoa;"AC clade";Apicomplexa;"cc | EP00376 Eimeria tenella 1134 | 0,00088 | 1410 | XP 013228860.1 | ABC transporter, putative [Eimeria tenella] | 1410 | 0 | 2706,4 | 100 |
| EP00485 | Sar;Rhizaria;Retaria;Foraminifera;Globothalamea;R | EP00485 Elphidium margaritaceum 18737 | 0,00089 | 1605 | XP 008870474.1 | hypothetical protein H310 06987 [Aphanomyces invadans] | 846 | 2,04E-86 | 313,153 | 45,85921325 |
| EP00256 | Archaeplastida;Chloroplastida;Streptophyta;"core s | EP00256 Glycine max 13155 | 0,0009 | 892 | XP 003526430.1 | transcriptional corepressor LEUNIG isoform X1 [Glycine max] >gi\|1590532612\|ref\|XP 02{ | 892 | 0 | 1832,77 | 100 |
| EP00331 | Sar;Alveolata;Ciliophora;Intramacronucleata;"CON: | EP00331 Platyophrya macrostoma 2772 | 0,0009 | 423 | XP 009607598.1 | PREDICTED: ubiquitin receptor RAD23c-like isoform X2 [Nicotiana tomentosiformis] >gi\|1( | 365 | 2,14E-62 | 214,927 | 52,98701299 |
| EP00149 | Obazoa;Opisthokonta;Nucletmycea;Fungi;"core Fur | EP00149 Cryptococcus neoformans 5813 | 0,0009 | 1442 | XP 012051953.1 | protein transporter SEC31 [Cryptococcus neoformans var. grubii H99] | 1442 | 0 | 2934,05 | 100 |
| EP00007 | Amoebozoa;Discosea;Flabellinia;"core flabellinids"; | EP00007 Cunea sp BSH-02190019 9851 | 0,0009 | 1196 | XP 025175506.1 | hypothetical protein GLOIN 2v1633885 [Rhizophagus irregularis DAOM 181602=DAOM 1 | 1409 | 1,16E-105 | 374,4 | 45,81124072 |
| EP00445 | Sar;Alveolata;Myzozoa;"DP clade";Dinoflagellata;D | EP00445 Apocalathium hangoei 92858 | 0,00091 | 504 | ***no hit found*** | | | | | |
| EP00536 | Sar;Stramenopiles;Gyrista;Ochrophyta;"BD clade";[ | EP00536 Pseudo-nitzschia multiseries 4481 | 0,00091 | 620 | XP 008912499.1 | hypothetical protein PPTG 16764 [Phytophthora parasitica INRA-310] | 525 | 9,94E-21 | 107,071 | 46,83760684 |
| EP00534 | Sar;Stramenopiles;Gyrista;Ochrophyta;"BD clade";[ | EP00534 Pseudo-nitzschia fraudulenta 4098 | 0,00092 | 394 | ***no hit found*** | | | | | |
| EP00334 | Sar;Alveolata;Ciliophora;Intramacronucleata;"CON: | EP00334 Paramecium tetraurelia 33410 | 0,00092 | 595 | XP 001456424.1 | hypothetical protein [Paramecium tetraurelia strain d4-2] | 595 | 0 | 1212,59 | 100 |
| EP00063 | Obazoa;Opisthokonta;Holozoa;Choanozoa;Metazoa | EP00063 Salmo salar 14561 | 0,00092 | 853 | XP 013989396.1 | PREDICTED: zinc finger protein 37-like isoform X1 [Salmo salar] | 853 | 0 | 1774,22 | 100 |
| EP00011 | Amoebozoa;Discosea;Flabellinia;"core flabellinids"; | EP00011 Vannella sp DIVA3-517-6-12 11026 | 0,00093 | 641 | XP 011265577.2 | arf-GAP domain and FG repeat-containing protein 1 isoform X1 [Camponotus floridanus] | 411 | 5,13E-33 | 141,739 | 41,21863799 |
| EP00548 | Sar;Stramenopiles;Gyrista;Ochrophyta;"BD clade";[ | EP00548 Thalassiothrix antarctica 9223 | 0,00093 | 472 | XP 002186485.1 | predicted protein [Phaeodactylum tricornutum CCAP 1055/1] | 899 | 6,80E-52 | 198,749 | 49,67032967 |
| EP00403 | Sar;Alveolata;Myzozoa;"DP clade";Dinoflagellata;D | EP00403 Amphidinium massartii 6909 | 0,00093 | 607 | XP 025556731.1 | putative NADH-cytochrome b5 reductase [Aspergillus homomorphus CBS 101889] | 320 | 4,95E-16 | 90,1225 | 43,85026738 |
| EP00035 | Obazoa;Opisthokonta;Holozoa;Choanozoa;Choanof | EP00035 Acanthoeca spectabilis 31354 | 0,00094 | 1138 | XP 010159299.1 | PREDICTED: protein transport protein Sec31A isoform X4 [Eurypyga helias] | 1176 | 5,64E-63 | 245,743 | 40,91653028 |
| EP00075 | Obazoa;Opisthokonta;Holozoa;Choanozoa;Metazoa | EP00075 Anas platyrhynchos 22253 | 0,00094 | 1505 | XP 027311506.1 | membrane-associated guanylate kinase, WW and PDZ domain-containing protein 2 [Anas | 1505 | 0 | 3103,15 | 100 |
| EP00490 | Sar;Rhizaria;Retaria;Foraminifera;Tubothalamea;M | EP00490 Sorites sp Unknown 13511 | 0,00094 | 227 | XP 002772264.1 | hypothetical protein Pmar PMAR023065 [Perkinsus marinus ATCC 50983] | 173 | 5,72E-08 | 60,8474 | 48,22695035 |
| EP00432 | Sar;Alveolata;Myzozoa;"DP clade";Dinoflagellata;D | EP00432 Prorocentrum cordatum 20703 | 0,00094 | 446 | XP 002773527.1 | conserved hypothetical protein [Perkinsus marinus ATCC 50983] | 1211 | 2,71E-93 | 315,849 | 58,17805383 |
| EP00361 | Sar;Alveolata;Ciliophora;Postciliodesmatophora;He | EP00361 Fabrea salina 2748 | 0,00095 | 1036 | XP 001440954.1 | hypothetical protein [Paramecium tetraurelia strain d4-2] | 988 | 5,78E-72 | 269,24 | 36,1849711 |
| EP00196 | Archaeplastida;Chloroplastida;Chlorophyta;"core ch | EP00196 Chlamydomonas euryale 9818 | 0,00095 | 486 | ***no hit found*** | | | | | |
| EP00266 | Archaeplastida;Chloroplastida;Streptophyta;"core s | EP00266 Sorghum bicolor 32225 | 0,00095 | 755 | XP 021315051.1 | protein STRUBBELIG-RECEPTOR FAMILY 3 isoform X2 [Sorghum bicolor] | 755 | 0 | 1524,61 | 100 |
| EP00104 | Obazoa;Opisthokonta;Holozoa;Choanozoa;Metazoa | EP00104 Lingula anatina 26233 | 0,00096 | 3292 | XP 013409251.1 | uncharacterized protein LOC106172880 isoform X4 [Lingula anatina] | 3292 | 0 | 6689,36 | 100 |
| EP00532 | Sar;Stramenopiles;Gyrista;Ochrophyta;"BD clade";[ | EP00532 Pseudo-nitzschia australis 2807 | 0,00096 | 592 | XP 002183090.1 | predicted protein [Phaeodactylum tricornutum CCAP 1055/1] | 491 | 2,30E-84 | 281,567 | 53,57142857 |
| EP00578 | Sar;Stramenopiles;Gyrista;Ochrophyta;"BD clade";[ | EP00578 Thalassiosira sp NH16 29976 | 0,00096 | 1114 | XP 002291969.1 | predicted protein [Thalassiosira pseudonana CCMP1335] | 1116 | 1,56E-119 | 405,601 | 56,31868132 |
| EP00403 | Sar;Alveolata;Myzozoa;"DP clade";Dinoflagellata;D | EP00403 Amphidinium massartii 47727 | 0,00097 | 629 | XP 647551.1 | hypothetical protein DDB G0268102 [Dictyostelium discoideum AX4] | 1861 | 4,42E-18 | 100,523 | 96,82539683 |
| EP00258 | Archaeplastida;Chloroplastida;Streptophyta;"core s | EP00258 Populus trichocarpa 6135 | 0,00097 | 900 | XP 002309593.3 | transcriptional corepressor LEUNIG isoform X1 [Populus trichocarpa] | 900 | 0 | 1852,41 | 100 |
| EP00613 | Sar;Stramenopiles;Gyrista;Ochrophyta;Dictyochoph | EP00613 Pedinella sp CCMP2098 55953 | 0,00097 | 970 | XP 009526689.1 | hypothetical protein PHYSODRAFT 331569 [Phytophthora sojae] | 785 | 3,55E-85 | 301,212 | 50 |
| EP00011 | Amoebozoa;Discosea;Flabellinia;"core flabellinids"; | EP00011 Vannella sp DIVA3-517-6-12 6729 | 0,00099 | 1717 | XP 004338529.1 | RhoGEF domain containing protein [Acanthamoeba castellanii str. Neff] | 670 | 2,72E-41 | 174,481 | 36,42384106 |
| EP00445 | Sar;Alveolata;Myzozoa;"DP clade";Dinoflagellata;D | EP00445 Apocalathium hangoei 33382 | 0,00099 | 686 | ***no hit found*** | | | | | |
| EP00025 | Amoebozoa;Evosea;Eumycetozoa;Dictyostelia;Dicty | EP00025 Tieghemostelium lacteum 2789 | 0,00099 | 413 | ***no hit found*** | | | | | |

**Supplementary Table 7.** List of Sea2, Sea3, and Sea4 proteins in studied euglenozoans and their reverse best hits.

| | | | | BLAST against *Naegleria gruberi* | | | | | | BLAST against *Paradiplonema papillatum* | | | | | |
|---|---|---|---|---|---|---|---|---|---|---|---|---|---|---|---|
| | | | | 1st hit | | | 2nd hit | | | 1st hit | | | 2nd hit | | |
| | Organism | Accession number | Sequence | Accession | Description | Evalue | Accession | Description | Evalue | Accession | Description | Evalue | Accession | Description | Evalue |
| **Sea2 (Wdr24)** | *Paradiplonema papillatum* | DIPPA 06078.mRNA.1 | MADSSVAGSCK | XP 002680795.1 | WD repeat domain 24 [Naegleria gruberi] = Sea2 | 1.00E-38 | XP 002678232.1 | uncharacterized protein NAEGRDRAFT 48518 [Naegleria gruberi] | 2.00E-09 | - | | | | | |
| | *Diplonema japonicum* | TRINITY DN10112 c4 g2 i1 | MQDVAKSTVKL | XP 002680795.1 | WD repeat domain 24 [Naegleria gruberi] = Sea2 | 9,00E-47 | XP 002678232.1 | uncharacterized protein NAEGRDRAFT 48518 [Naegleria gruberi] | 3,00E-08 | - | | | | | |
| | *Rhynchopus humris* | TRINITY DN63379 c0 g1 i1 | MSCSSPTGGSG | XP 002680795.1 | WD repeat domain 24 [Naegleria gruberi] = Sea2 | 3,00E-42 | XP 002677128.1 | Peroxin 7 (Pex7), putative [Naegleria gruberi] | 9,00E-09 | - | | | | | |
| | *Lacrimia lanifica* | TRINITY DN12466 c8 g1 i2 | MEAREDSGAQ | XP 002680795.1 | WD repeat domain 24 [Naegleria gruberi] = Sea2 | 4.00E-41 | XP 002676671.1 | uncharacterized protein NAEGRDRAFT 68293 [Naegleria gruberi] | 2.00E-10 | - | | | | | |
| | *Sulcionema specki* | TRINITY DN1214 c0 g2 i1 | MRGMPGGART | XP 002680795.1 | WD repeat domain 24 [Naegleria gruberi] = Sea2 | 6,00E-44 | XP 002677128.1 | Peroxin 7 (Pex7), putative [Naegleria gruberi] | 2,00E-12 | - | | | | | |
| | *Artemidia motanka* | TRINITY DN16143 c4 g1 i3 | MLPVHGRQRQ | XP 002680795.1 | WD repeat domain 24 [Naegleria gruberi] = Sea2 | 9,00E-37 | XP 002677128.1 | Peroxin 7 (Pex7), putative [Naegleria gruberi] | 4,00E-08 | - | | | | | |
| | *Namystynia karyoxenos* | TRINITY DN34409 c0 g1 i1 | MSALDALRPAT | XP 002680795.1 | WD repeat domain 24 [Naegleria gruberi] = Sea2 | 6.00E-38 | XP 002676671.1 | uncharacterized protein NAEGRDRAFT 68293 [Naegleria gruberi] | 1.00E-13 | - | | | | | |
| | *Bodo saltans* | BSAL 18765 | MRNSAAANNS | XP 002680795.1 | WD repeat domain 24 [Naegleria gruberi] = Sea2 | 3.00E-10 | XP 002671379.1 | uncharacterized protein NAEGRDRAFT 73659 [Naegleria gruberi] = Sea3 | 7.00E-05 | DIPPA 06078.mRNA.1 | Sea2 | 2.71E-06 | DIPPA 19733.mRNA.1 | E3 ubiquitin-protein ligase RING1 | 2.93E-03 |
| | *Trypanosoma brucei* | Tb927.1.1880 | MNESHMCHQF | XP 002680795.1 | WD repeat domain 24 [Naegleria gruberi] = Sea2 | 3,00E-08 | XP 002671379.1 | uncharacterized protein NAEGRDRAFT 73659 [Naegleria gruberi] = Sea3 | 0,002 | DIPPA 06078.mRNA.1 | Sea2 | 3,00E-06 | DIPPA 21857.mRNA.1 | Sea4 | 3,29E-04 |
| | *Leishmania major* | LmjF.20.1080 | MPVSLRKNPPR | XP 002680795.1 | WD repeat domain 24 [Naegleria gruberi] = Sea2 | 4,00E-11 | XP 002671379.1 | uncharacterized protein NAEGRDRAFT 73659 [Naegleria gruberi] = Sea3 | 0,002 | DIPPA 06078.mRNA.1 | Sea2 | 1,60E-05 | DIPPA 21857.mRNA.1 | Sea4 | 5,37E-05 |
| | *Euglena gracilis* | GDJR01058097.1 | MNTSGSASSLSI | XP 002680795.1 | WD repeat domain 24 [Naegleria gruberi] = Sea2 | 6.00E-52 | XP 002678232.1 | uncharacterized protein NAEGRDRAFT 48518 [Naegleria gruberi] | 5.00E-17 | - | | | | | |
| | *Euglena longa* | Contig992 | MNLSGDLIGPT | XP 002680795.1 | WD repeat domain 24 [Naegleria gruberi] = Sea2 | 3,00E-52 | XP 002678232.1 | uncharacterized protein NAEGRDRAFT 48518 [Naegleria gruberi] | 8,00E-17 | - | | | | | |
| | *Eutreptiella gymnastica* | Transcript 32132+62298 | MGDEFKDRESY | XP 002680795.1 | WD repeat domain 24 [Naegleria gruberi] = Sea2 | 3,00E-35 | XP 002676671.1 | uncharacterized protein NAEGRDRAFT 68293 [Naegleria gruberi] | 7,00E-10 | - | | | | | |
| **Sea3 (Wdr59)** | *Paradiplonema papillatum* | DIPPA 06252.mRNA.1 | MNDGPGKKLW | - | | | - | | | - | | | | | |
| | *Diplonema japonicum* | TRINITY DN11501 c6 g1 i1 | MPEYNGPGKLL | XP 002671379.1 | uncharacterized protein NAEGRDRAFT 73659 [Naegleria gruberi] = Sea3 | 4.00E-07 | - | | | - | | | | | |
| | *Rhynchopus humris* | TRINITY DN22035 c1 g1 i1 | LCAGGTVASEPA | - | | | - | | | - | | | | | |
| | *Lacrimia lanifica* | TRINITY DN10159 c0 g3 i2 | MTEYDGPGRLL | XP 002671379.1 | uncharacterized protein NAEGRDRAFT 73659 [Naegleria gruberi] = Sea3 | 3,00E-07 | XP 002680795.1 | WD repeat domain 24 [Naegleria gruberi] = Sea2 | 0,014 | - | | | | | |
| | *Sulcionema specki* | TRINITY DN47071 c0 g1 i1 | MGSHEKSSREY | XP 002671379.1 | uncharacterized protein NAEGRDRAFT 73659 [Naegleria gruberi] = Sea3 | 1.90E-02 | - | | | - | | | | | |
| | *Artemidia motanka* | TRINITY DN17178 c8 g2 i1 | MAGRDKGPEKI | XP 002671379.1 | uncharacterized protein NAEGRDRAFT 73659 [Naegleria gruberi] = Sea3 | 1,50E-02 | - | | | - | | | | | |
| | *Namystynia karyoxenos* | TRINITY DN46998 c0 g1 i1 | MGQRPRSDGP | - | | | - | | | - | | | | | |
| | *Bodo saltans* | BSAL 71480 | MTQSIHDDVRK | XP 002671379.1 | uncharacterized protein NAEGRDRAFT 73659 [Naegleria gruberi] = Sea3 | 6.00E-05 | XP 002674131.1 | uncharacterized protein NAEGRDRAFT 80725 [Naegleria gruberi] = Sea4 | 4.00E-04 | DIPPA 21857.mRNA.1 | Sea4 | 9.22E-06 | DIPPA 06252.mRNA.1 | Sea3 | 8.87E-03 |
| | *Trypanosoma brucei* | Tb927.3.1880 | MGPLVVVSPEV | XP 002671379.1 | uncharacterized protein NAEGRDRAFT 73659 [Naegleria gruberi] = Sea3 | 5.00E-04 | - | | | DIPPA 06252.mRNA.1 | Sea3 | 6.89E-03 | DIPPA 21857.mRNA.1 | Sea4 | 2.56E-01 |
| | *Leishmania major* | LmjF.25.1800 | MLSQALLSAVEI | XP 002671379.1 | uncharacterized protein NAEGRDRAFT 73659 [Naegleria gruberi] = Sea3 | 1,00E-06 | XP 002674131.1 | uncharacterized protein NAEGRDRAFT 80725 [Naegleria gruberi] = Sea4 | 1,10E-02 | DIPPA 06252.mRNA.1 | Sea3 | 4,13E-05 | DIPPA 30849.mRNA.1 | hypothetical protein | 3,08E-01 |
| | *Euglena gracilis* | GDJR01069079.1 | MAVAWQLSLR | XP 002671379.1 | uncharacterized protein NAEGRDRAFT 73659 [Naegleria gruberi] = Sea3 | 3,00E-11 | XP 002680795.1 | WD repeat domain 24 [Naegleria gruberi] = Sea2 | 7,00E-06 | - | | | | | |
| | *Euglena longa* | Contig63495+Contig40859+Contig15550 | MAVDWQLSLR | XP 002671379.1 | uncharacterized protein NAEGRDRAFT 73659 [Naegleria gruberi] = Sea3 | 7.00E-11 | XP 002683071.1 | uncharacterized protein NAEGRDRAFT 61206 [Naegleria gruberi] | 3.00E-10 | - | | | | | |
| | *Eutreptiella gymnastica* | Transcript 137785+137784+134183+109285 | NTDRMLPGLGF | XP 002671379.1 | uncharacterized protein NAEGRDRAFT 73659 [Naegleria gruberi] = Sea3 | 8.00E-10 | XP 002680795.1 | WD repeat domain 24 [Naegleria gruberi] = Sea2 | 7.00E-06 | - | | | | | |
| **Sea4** | *Paradiplonema papillatum* | DIPPA 21857.mRNA.1 | MSSAACVGPTL | XP 002674131.1 | uncharacterized protein NAEGRDRAFT 80725 [Naegleria gruberi] = Sea4 | 9,00E-33 | XP 002671379.1 | uncharacterized protein NAEGRDRAFT 73659 [Naegleria gruberi] = Sea3 | 1,00E-06 | - | | | | | |
| | *Diplonema japonicum* | TRINITY DN11332 c4 g1 i1 | MSNAACIGGDI | XP 002674131.1 | uncharacterized protein NAEGRDRAFT 80725 [Naegleria gruberi] = Sea4 | 9.00E-20 | XP 002671379.1 | uncharacterized protein NAEGRDRAFT 73659 [Naegleria gruberi] = Sea3 | 1.00E-07 | - | | | | | |
| | *Rhynchopus humris* | TRINITY DN63991 c0 g1 i1 | VDKEGPLGDRIC | XP 002674131.1 | uncharacterized protein NAEGRDRAFT 80725 [Naegleria gruberi] = Sea4 | 1.00E-29 | XP 002671379.1 | uncharacterized protein NAEGRDRAFT 73659 [Naegleria gruberi] = Sea3 | 1.00E-09 | - | | | | | |
| | *Lacrimia lanifica* | TRINITY DN10718 c0 g1 i1 | MANTAANPGD | XP 002674131.1 | uncharacterized protein NAEGRDRAFT 80725 [Naegleria gruberi] = Sea4 | 3,00E-17 | XP 002671379.1 | uncharacterized protein NAEGRDRAFT 73659 [Naegleria gruberi] = Sea3 | 2,00E-07 | - | | | | | |
| | *Sulcionema specki* | TRINITY DN38070 c0 g1 i1 | MVNRTVCCWV | XP 002674131.1 | uncharacterized protein NAEGRDRAFT 80725 [Naegleria gruberi] = Sea4 | 8,00E-28 | XP 002671379.1 | uncharacterized protein NAEGRDRAFT 73659 [Naegleria gruberi] = Sea3 | 9,00E-08 | - | | | | | |
| | *Artemidia motanka* | TRINITY DN17041 c6 g10 i1 | MQPILCPHPWF | XP 002674131.1 | uncharacterized protein NAEGRDRAFT 80725 [Naegleria gruberi] = Sea4 | 7.00E-15 | XP 002671379.1 | uncharacterized protein NAEGRDRAFT 73659 [Naegleria gruberi] = Sea3 | 2.00E-05 | - | | | | | |
| | *Namystynia karyoxenos* | TRINITY DN21390 c0 g1 i1 | MSSASCVLATPI | XP 002674131.1 | uncharacterized protein NAEGRDRAFT 80725 [Naegleria gruberi] = Sea4 | 2.00E-11 | XP 002671379.1 | uncharacterized protein NAEGRDRAFT 73659 [Naegleria gruberi] = Sea3 | 8.00E-05 | - | | | | | |
| | *Bodo saltans* | BSAL 10500 | MSCKFFLSVPET | XP 002674131.1 | uncharacterized protein NAEGRDRAFT 80725 [Naegleria gruberi] = Sea4 | 5,00E-05 | - | | | DIPPA 21857.mRNA.1 | Sea4 | 2,07E-08 | DIPPA 06078.mRNA.1 | Sea2 | 9,26E-02 |
| | *Trypanosoma brucei* | Tb927.3.1310 | MSFSALSVPSEI | XP 002674131.1 | uncharacterized protein NAEGRDRAFT 80725 [Naegleria gruberi] = Sea4 | 4.00E-11 | XP 002671379.1 | uncharacterized protein NAEGRDRAFT 73659 [Naegleria gruberi] = Sea3 | 3.00E-07 | DIPPA 21857.mRNA.1 | Sea4 | 1.28E-09 | DIPPA 06078.mRNA.1 | Sea2 | 3.65E-04 |
| | *Trypanosoma brucei* | Tb927.3.1320 | MKYFACRLDDH | XP 002674131.1 | uncharacterized protein NAEGRDRAFT 80725 [Naegleria gruberi] = Sea4 | 1.00E-06 | XP 002671379.1 | uncharacterized protein NAEGRDRAFT 73659 [Naegleria gruberi] = Sea3 | 3.00E-03 | DIPPA 21857.mRNA.1 | Sea4 | 3.36E-05 | DIPPA 06078.mRNA.1 | Sea2 | 4.80E-02 |
| | *Leishmania major* | LmjF.25.1250 | MKHFSCQPDSH | XP 002674131.1 | uncharacterized protein NAEGRDRAFT 80725 [Naegleria gruberi] = Sea4 | 9,00E-11 | XP 002671379.1 | uncharacterized protein NAEGRDRAFT 73659 [Naegleria gruberi] = Sea3 | 9,00E-09 | DIPPA 21857.mRNA.1 | Sea4 | 1,58E-08 | DIPPA 06078.mRNA.1 | Sea2 | 5,84E-04 |
| | *Leishmania major* | LmjF.25.1260 | MCVRLLSPSLHF | XP 002674131.1 | uncharacterized protein NAEGRDRAFT 80725 [Naegleria gruberi] = Sea4 | 3,00E-09 | XP 002671379.1 | uncharacterized protein NAEGRDRAFT 73659 [Naegleria gruberi] = Sea3 | 2,00E-05 | DIPPA 21857.mRNA.1 | Sea4 | 6,89E-07 | DIPPA 09941.mRNA.1 | Sec31 | 1,11E-02 |
| | *Euglena gracilis* | GEFR01000988.1 | MQGRVALCLDI | XP 002674131.1 | uncharacterized protein NAEGRDRAFT 80725 [Naegleria gruberi] = Sea4 | 3.00E-23 | XP 002680795.1 | WD repeat domain 24 [Naegleria gruberi] = Sea2 | 3.00E-07 | - | | | | | |
| | *Euglena longa* | Contig40778+40779+41677+18493 | MPDRAALCVDV | XP 002674131.1 | uncharacterized protein NAEGRDRAFT 80725 [Naegleria gruberi] = Sea4 | 1.00E-22 | XP 002680795.1 | WD repeat domain 24 [Naegleria gruberi] = Sea2 | 1.00E-06 | - | | | | | |
| | *Eutreptiella gymnastica* | Transcript 65317+25574+48903+77209 | MQYVLKSPISFN | XP 002674131.1 | uncharacterized protein NAEGRDRAFT 80725 [Naegleria gruberi] = Sea4 | 6,00E-15 | XP 002680795.1 | WD repeat domain 24 [Naegleria gruberi] = Sea2 | 4,70E-02 | - | | | | | |

**Functional differentiation of Sec13 paralogs in the euglenozoan protists**

Drahomíra Faktorová[1,2,#,*], Kristína Záhonová[1,3,4,*], Corinna Benz[1], Joel B. Dacks[1,5,6], Mark C. Field[1,7] and Julius Lukeš[1,2,#]

[1] *Institute of Parasitology, Biology Centre, Czech Academy of Sciences, České Budějovice, Czech Republic*
[2] *Faculty of Sciences, University of South Bohemia, České Budějovice, Czech Republic*
[3] *Faculty of Science, Charles University, BIOCEV, Vestec, Czech Republic*
[4] *Life Science Research Centre, Department of Biology and Ecology, Faculty of Science, University of Ostrava, Ostrava, Czech Republic*
[5] *Division of Infectious Disease, Department of Medicine, Faculty of Medicine and Dentistry, University of Alberta, Edmonton, Canada*
[6] *Centre for Life's Origins and Evolution, Department of Genetics, Evolution, and Environment, University College London, London, UK*
[7] *School of Life Sciences, University of Dundee, Dundee, UK*

*These authors contributed equally to this work.

**Supplementary Data 1.** Whole MS data.

| iBAQ Sec | iBAQ Sec | iBAQ Sec | iBAQ wt- | iBAQ wt- | iBAQ wt- | LFQ inten | LFQ inten | LFQ inten | LFQ inten | LFQ inten | LFQ inten | Student's | Student's | Peptides | Razor + u | Unique pe | Sequence |
|---|---|---|---|---|---|---|---|---|---|---|---|---|---|---|---|---|---|
| 29,0847 | 24,5133 | 24,3809 | 16,8115 | 10,9929 | 13,6753 | 30,692 | 28,1686 | 28,4786 | 17,2726 | 19,3689 | 16,6639 | | | 64 | 64 | 64 | 58,2 |
| 28,7794 | 24,0447 | 23,0781 | 18,6316 | 11,132 | 14,5643 | 30,5215 | 27,602 | 25,8234 | 20,7824 | 19,8794 | 18,9745 | | | 9 | 9 | 9 | 39,8 |
| 25,2039 | 18,8022 | 19,2667 | 15,0266 | 10,5554 | 12,9881 | 27,4821 | 21,9097 | 22,8969 | 18,8131 | 19,0328 | 19,2564 | | | 12 | 12 | 12 | 65,2 |
| 23,9583 | 17,3563 | 18,3644 | 15,7909 | 10,5058 | 13,5451 | 25,6187 | 21,545 | 22,741 | 20,1325 | 19,0353 | 18,765 | | | 10 | 10 | 10 | 39 |
| 20,012 | 16,8956 | 16,0353 | 16,9567 | 11,0027 | 11,1888 | 22,0315 | 21,749 | 21,5482 | 20,7939 | 19,5189 | 18,6276 | | | 13 | 12 | 12 | 38,1 |
| 21,6038 | 12,0574 | 14,9329 | 11,8494 | 13,3785 | 10,866 | 18,6502 | 20,547 | 19,632 | 20,0912 | 20,0935 | 19,6019 | | | 8 | 8 | 8 | 29,1 |
| 22,8901 | 15,497 | 15,4502 | 19,2286 | 11,6466 | 10,5982 | 24,4796 | 19,9192 | 19,9836 | 22,5345 | 19,5867 | 19,0353 | | | 12 | 12 | 12 | 52 |
| 20,7128 | 18,5551 | 18,4331 | 18,7095 | 11,8058 | 15,7721 | 23,3141 | 23,8634 | 23,6516 | 22,5195 | 19,7913 | 22,6129 | | | 19 | 19 | 19 | 37,9 |
| 20,857 | 14,1126 | 13,0426 | 14,8918 | 11,1928 | 11,3367 | 24,0794 | 19,7627 | 20,1581 | 20,3694 | 20,4995 | 18,9294 | | | 8 | 8 | 8 | 35,6 |
| 19,2605 | 14,0226 | 13,168 | 15,3447 | 10,6043 | 10,3865 | 19,7796 | 19,8603 | 19,3002 | 20,3039 | 18,4565 | 18,9433 | | | 6 | 6 | 6 | 16,1 |
| 19,2386 | 15,5044 | 16,8534 | 16,9877 | 11,2624 | 13,7339 | 22,2721 | 19,9172 | 22,576 | 21,8029 | 19,4946 | 18,0571 | | | 5 | 5 | 5 | 26,8 |
| 23,9297 | 14,509 | 14,119 | 20,1672 | 9,98987 | 13,0633 | 26,2319 | 18,6887 | 19,4108 | 23,7793 | 20,4038 | 18,7961 | | | 18 | 18 | 16 | 56 |
| 17,1082 | 12,7156 | 12,238 | 10,9841 | 11,1162 | 10,6548 | 20,5176 | 19,8748 | 19,4416 | 20,2078 | 19,2719 | 20,8895 | | | 13 | 13 | 13 | 4,8 |
| 19,8975 | 16,4003 | 15,7441 | 16,7567 | 9,89016 | 16,1164 | 19,1604 | 18,9831 | 19,635 | 20,3421 | 20,2214 | 18,0672 | | | 7 | 7 | 7 | 38,3 |
| 19,8893 | 15,1263 | 14,7445 | 17,8161 | 11,2849 | 11,381 | 22,8617 | 20,3656 | 21,0036 | 21,3654 | 19,7629 | 19,4056 | | | 11 | 11 | 11 | 30 |
| 22,1103 | 14,2299 | 15,8349 | 17,9254 | 10,5549 | 14,5366 | 24,1474 | 19,827 | 24,8075 | 24,231 | 20,1095 | 23,9989 | | | 5 | 5 | 5 | 21,2 |
| 20,0638 | 16,5258 | 16,3251 | 15,109 | 12,5067 | 16,1538 | 19,0305 | 18,6227 | 19,491 | 20,148 | 20,5536 | 19,6771 | | | 6 | 6 | 6 | 30,6 |
| 22,2553 | 14,8019 | 14,2461 | 18,7574 | 9,47793 | 13,9644 | 23,4083 | 18,1536 | 19,3353 | 21,3711 | 20,5344 | 18,9526 | | | 8 | 8 | 8 | 54,5 |
| 19,4506 | 17,0847 | 15,7304 | 15,3209 | 13,1563 | 14,712 | 22,4907 | 22,3733 | 21,4562 | 20,5384 | 18,8088 | 20,9007 | | | 7 | 7 | 7 | 16,9 |
| 17,2296 | 14,9054 | 16,208 | 16,0097 | 11,1095 | 12,2644 | 21,1915 | 22,1276 | 21,649 | 21,5476 | 19,8492 | 19,191 | | | 10 | 10 | 10 | 25 |
| 18,8665 | 10,0779 | 13,3464 | 12,6837 | 9,3271 | 11,489 | 18,8701 | 19,8284 | 20,3448 | 19,1062 | 20,5168 | 19,1545 | | | 54 | 54 | 54 | 18,9 |
| 20,7946 | 16,8559 | 14,0878 | 18,2689 | 11,3225 | 13,397 | 23,9255 | 18,8248 | 19,7288 | 21,9434 | 19,952 | 19,6948 | | | 23 | 8 | 8 | 48,5 |
| 19,1375 | 16,1245 | 15,5201 | 17,4237 | 9,73775 | 15,1386 | 21,1837 | 21,8152 | 21,6511 | 20,3805 | 20,445 | 18,893 | | | 6 | 6 | 6 | 27 |
| 21,9068 | 16,3222 | 14,2774 | 19,1793 | 12,3235 | 12,6742 | 24,5231 | 21,2987 | 19,0095 | 22,5858 | 19,5435 | 19,7649 | | | 7 | 7 | 7 | 33,3 |
| 21,0236 | 10,5957 | 17,4835 | 18,6095 | 11,14 | 11,0489 | 22,7038 | 19,4496 | 18,813 | 20,5797 | 20,1166 | 20,0067 | | | 3 | 3 | 3 | 17,8 |
| 19,9908 | 12,826 | 14,8128 | 17,8645 | 9,34147 | 12,2704 | 22,0481 | 18,7326 | 19,3418 | 20,47 | 19,5539 | 18,5479 | | | 9 | 9 | 9 | 40,3 |
| 20,1515 | 18,2038 | 16,804 | 20,2538 | 11,0876 | 15,6647 | 22,2933 | 23,0101 | 22,294 | 23,6572 | 19,5952 | 21,8524 | | | 12 | 12 | 12 | 37,3 |
| 23,4501 | 18,2777 | 17,952 | 20,5049 | 14,9538 | 16,1346 | 25,2766 | 22,2627 | 22,7626 | 23,1161 | 20,3365 | 20,7091 | | | 13 | 13 | 13 | 44,3 |
| 22,1726 | 12,533 | 11,1531 | 17,2228 | 11,1484 | 9,64143 | 23,6095 | 18,5791 | 19,1874 | 20,0158 | 18,9246 | 19,211 | | | 12 | 12 | 12 | 68 |
| 21,503 | 16,4942 | 15,3339 | 18,2406 | 12,4175 | 14,9539 | 19,0133 | 19,8689 | 20,2155 | 20,0765 | 20,246 | 19,4228 | | | 3 | 3 | 3 | 30,5 |
| 20,2635 | 13,3673 | 14,5957 | 17,8415 | 10,0443 | 12,6749 | 22,7328 | 19,7486 | 19,0659 | 21,4906 | 19,3157 | 18,0337 | | | 7 | 7 | 7 | 40,9 |

| | | | | | | | | | | | |
|---|---|---|---|---|---|---|---|---|---|---|---|
| 17,0615 | 19,7507 | 19,1873 | 16,7749 | 17,8016 | 18,759 | 19,5238 | 23,6872 | 23,8545 | 21,3533 | 23,0067 | 24,1331 |
| 18,5871 | 12,1658 | 15,0802 | 18,7653 | 10,1317 | 14,2797 | 21,2597 | 20,5171 | 21,1868 | 22,3061 | 19,4832 | 19,2563 |
| 21,4655 | 14,436 | 15,4695 | 20,0926 | 13,5193 | 15,1122 | 23,5142 | 19,9556 | 19,9672 | 23,4284 | 19,1556 | 19,6078 |
| 22,3252 | 20,594 | 20,3214 | 21,2199 | 19,3691 | 20,0155 | 23,8722 | 24,3223 | 24,4207 | 23,4324 | 24,3064 | 24,4347 |
| 18,2368 | 17,148 | 16,8157 | 19,5242 | 14,729 | 15,3139 | 20,9369 | 21,1588 | 21,0399 | 22,1301 | 20,1226 | 19,6664 |
| 18,9422 | 19,9034 | 18,8827 | 19,6866 | 16,8163 | 18,6139 | 21,184 | 23,6401 | 23,4373 | 22,7468 | 22,4972 | 23,1914 |
| 22,7922 | 19,0297 | 18,1681 | 21,5355 | 17,6084 | 18,246 | 25,0446 | 23,6999 | 23,5392 | 24,3004 | 22,9434 | 23,7875 |
| 19,5826 | 20,9187 | 20,1724 | 19,2816 | 19,2769 | 19,5303 | 21,4919 | 24,6184 | 24,5582 | 22,1982 | 23,9546 | 24,2478 |
| 15,3807 | 13,7321 | 13,0355 | 15,037 | 12,6426 | 11,8852 | 19,7169 | 19,8344 | 20,91 | 20,6519 | 18,8724 | 19,9479 |
| 18,1869 | 12,7321 | 13,5122 | 17,0547 | 10,5262 | 14,2696 | 22,3027 | 20,4412 | 19,5162 | 21,336 | 18,9324 | 21,5916 |
| 18,9641 | 9,50269 | 11,409 | 13,8594 | 10,9786 | 12,4661 | 22,6659 | 18,7679 | 18,7344 | 18,369 | 18,9476 | 19,5589 |
| 16,6691 | 17,7066 | 17,8845 | 17,1424 | 15,5473 | 17,0046 | 20,3884 | 23,1419 | 23,3144 | 21,2952 | 22,2981 | 22,5079 |
| 16,6409 | 14,5662 | 14,7177 | 17,8654 | 11,1302 | 14,3652 | 20,5683 | 21,0376 | 20,9543 | 21,9437 | 20,2907 | 20,6103 |
| 18,4035 | 14,7801 | 14,6008 | 18,7933 | 12,7361 | 13,6915 | 20,7639 | 20,6288 | 21,0896 | 22,6584 | 19,3244 | 20,3379 |
| 21,1914 | 19,6318 | 18,8253 | 20,2653 | 18,6211 | 18,2092 | 23,2165 | 24,4239 | 24,1413 | 23,4175 | 24,3522 | 23,7306 |
| 19,0056 | 16,556 | 16,6747 | 19,7828 | 13,5818 | 16,3239 | 21,7522 | 21,9668 | 22,1318 | 23,0697 | 19,3456 | 22,1639 |
| 21,0608 | 17,6136 | 16,8823 | 19,6852 | 16,1747 | 17,1491 | 22,4238 | 21,7932 | 21,3935 | 22,0108 | 21,722 | 21,4657 |
| 20,5028 | 17,7183 | 15,596 | 19,0151 | 15,6711 | 16,6033 | 23,6648 | 22,7563 | 22,3171 | 23,2956 | 21,7409 | 23,0461 |
| 18,688 | 17,2818 | 16,4495 | 17,8687 | 15,771 | 16,2699 | 21,5968 | 22,9616 | 22,5412 | 22,0952 | 22,4076 | 22,7001 |
| 21,2243 | 20,4126 | 19,0844 | 21,3384 | 17,9874 | 18,8879 | 23,649 | 24,1489 | 24,0777 | 24,524 | 23,4288 | 23,7358 |
| 20,2542 | 17,334 | 17,371 | 19,1011 | 15,9556 | 17,4343 | 22,1216 | 22,7814 | 23,0901 | 22,2129 | 22,4731 | 23,019 |
| 20,4542 | 17,6981 | 17,0129 | 19,1976 | 16,3676 | 17,14 | 21,864 | 22,0159 | 22,1458 | 22,2144 | 21,4392 | 22,1828 |
| 21,8063 | 18,7397 | 18,3353 | 20,4426 | 17,7041 | 18,2753 | 24,0444 | 23,8648 | 23,8013 | 23,7183 | 23,9635 | 23,9288 |
| 19,0615 | 17,076 | 16,5775 | 19,2053 | 15,25 | 15,8007 | 21,2255 | 22,3366 | 21,8838 | 22,1156 | 21,8783 | 21,7202 |
| 19,4943 | 17,464 | 17,2238 | 19,1319 | 16,0469 | 16,5555 | 21,9935 | 22,6028 | 22,3661 | 22,6208 | 22,0589 | 22,5204 |
| 24,7954 | 18,9607 | 18,7238 | 22,9664 | 18,0746 | 18,9955 | 26,9271 | 24,1811 | 24,0808 | 25,812 | 24,0772 | 24,2587 |
| 19,0032 | 19,8683 | 18,6282 | 19,4487 | 17,8771 | 17,7364 | 21,4445 | 24,3325 | 23,4031 | 22,2111 | 23,4113 | 22,7334 |
| 16,5321 | 16,6752 | 15,1213 | 19,6783 | 11,372 | 14,8424 | 19,5972 | 21,6158 | 20,3289 | 22,8968 | 20,6643 | 20,4787 |
| 20,3321 | 19,3567 | 19,1655 | 19,6423 | 18,0534 | 18,7358 | 22,8249 | 24,5763 | 24,3995 | 23,2241 | 24,2034 | 24,3679 |
| 15,0784 | 10,6682 | 14,8486 | 13,9443 | 11,6921 | 12,5372 | 20,4124 | 20,235 | 19,5975 | 20,5981 | 19,4862 | 19,2233 |
| 20,083 | 16,8826 | 15,7386 | 19,1707 | 15,4887 | 15,6297 | 23,0216 | 21,7309 | 21,3271 | 22,4048 | 21,0928 | 21,3983 |
| 19,0607 | 16,8137 | 16,4444 | 18,6157 | 15,0132 | 16,2869 | 21,881 | 21,7752 | 21,9198 | 22,0608 | 18,8631 | 22,0063 |
| 22,1124 | 20,9114 | 20,4335 | 21,7184 | 19,1594 | 20,1799 | 24,153 | 25,2312 | 24,778 | 24,1575 | 24,5804 | 24,898 |
| 24,2709 | 21,9977 | 21,9093 | 23,1922 | 21,2025 | 21,3877 | 26,6317 | 26,6178 | 26,996 | 26,2688 | 26,7993 | 26,8135 |
| 23,2028 | 21,5689 | 21,1648 | 22,9015 | 19,8286 | 20,8125 | 26,5557 | 26,3872 | 26,2991 | 27,0848 | 26,2589 | 26,3169 |
| 17,2453 | 12,0658 | 11,6084 | 14,1789 | 12,1934 | 12,1586 | 19,3077 | 18,7288 | 19,8509 | 19,0525 | 19,65 | 19,512 |
| 21,6115 | 11,5669 | 14,3399 | 20,1523 | 11,2253 | 13,7555 | 22,883 | 19,418 | 19,911 | 22,0415 | 19,5962 | 20,397 |
| 25,8485 | 20,8316 | 20,3849 | 24,2883 | 20,1356 | 20,2675 | 27,8875 | 25,0693 | 25,1521 | 27,0943 | 25,1939 | 25,0935 |
| 17,489 | 15,2926 | 14,6771 | 16,6538 | 14,635 | 13,8051 | 21,5704 | 21,7685 | 21,6801 | 21,4793 | 21,993 | 21,5784 |
| 22,1791 | 18,3543 | 18,1516 | 20,869 | 17,3967 | 18,055 | 23,223 | 23,1665 | 23,4906 | 23,6043 | 23,0302 | 23,5047 |
| 20,3333 | 18,5022 | 18,6978 | 20,4769 | 16,9454 | 17,7528 | 22,5309 | 23,2877 | 23,4709 | 22,8103 | 22,6248 | 23,1025 |
| 22,4214 | 15,628 | 18,4611 | 20,3543 | 16,7416 | 17,0599 | 25,2421 | 21,9144 | 22,2207 | 23,6613 | 22,5895 | 22,6812 |
| 24,5577 | 17,5165 | 18,9982 | 22,2769 | 17,6865 | 18,7588 | 26,1881 | 22,1607 | 23,4798 | 24,6768 | 22,8676 | 23,6331 |
| 16,5014 | 17,2535 | 16,5132 | 15,8297 | 15,8541 | 16,2472 | 20,0518 | 22,3633 | 21,9621 | 19,4688 | 21,8345 | 21,6741 |
| 20,5176 | 20,7534 | 18,7601 | 18,8306 | 19,9492 | 18,9146 | 25,9068 | 25,6256 | 26,2955 | 25,354 | 26,3961 | 26,4493 |
| 19,1781 | 11,7967 | 15,5271 | 17,7522 | 12,9113 | 13,5021 | 21,7399 | 19,5353 | 19,6203 | 20,9433 | 20,0838 | 20,8375 |
| 23,134 | 19,5187 | 19,0367 | 22,1509 | 18,1226 | 19,0808 | 24,8418 | 23,9274 | 23,6389 | 24,5979 | 23,3484 | 23,8346 |
| 18,2213 | 13,9124 | 11,3272 | 17,3605 | 10,4061 | 13,3629 | 20,9387 | 20,1987 | 19,5582 | 20,7131 | 20,3275 | 19,0684 |
| 22,9976 | 20,6132 | 20,0193 | 22,7418 | 19,0213 | 19,5443 | 25,2282 | 24,651 | 24,9311 | 25,8358 | 24,4111 | 24,6717 |
| 21,0885 | 11,6081 | 14,3631 | 19,3342 | 10,5679 | 14,8362 | 22,8986 | 20,5703 | 19,2816 | 22,3514 | 20,107 | 18,9688 |
| 17,3712 | 15,8804 | 15,513 | 17,5166 | 13,5784 | 15,3693 | 20,8908 | 21,7046 | 21,9687 | 21,5844 | 21,0782 | 21,6718 |
| 19,1491 | 15,0677 | 15,6937 | 18,5604 | 13,3963 | 15,6644 | 22,6231 | 21,9071 | 21,82 | 22,9661 | 20,8843 | 21,8257 |
| 23,0765 | 20,9057 | 20,4642 | 22,9751 | 19,1872 | 20,0031 | 25,158 | 25,5316 | 25,1992 | 25,7086 | 24,8473 | 24,9681 |

[illegible]

| | | | | | | | | | | | | | | | | | | | | | | | | | | | | | | | | | | | |
|---|---|---|---|---|---|---|---|---|---|---|---|---|---|---|---|---|---|---|---|---|---|---|---|---|---|---|---|---|---|---|---|---|---|---|---|
| 18,2477 | 17,9634 | 17,2671 | 19,5717 | 16,5555 | 17,2367 | 21,2551 | 23,0491 | 22,4922 | 22,5484 | 22,8962 | 22,369 | 20 | 20 | 20 | 32,6 | 32,6 | 32,6 | 85,405 | 0 | 117,23 | 6,7E+07 | 54 | 12 | 11 | 13 | 17 | 8 | 12 | 0,01314 | 1 | 0,0381 | 0,03975 | DIPPA_2 | DIPPA_2 | DIPPA_23975.mRNA.1 Protein disulfide-isomerase like 2-1;DIPPA_23975.mRNA.2 Chaperone protein DnaJ |
| 19,5223 | 16,7433 | 17,1498 | 20,5726 | 16,4325 | 16,3113 | 21,9138 | 22,02 | 22,4863 | 23,6735 | 22,1346 | 22,4135 | 9 | 9 | 9 | 37,6 | 37,6 | 37,6 | 39,354 | 0 | 51,083 | 4,9E+07 | 24 | 5 | 2 | 4 | 9 | 4 | 3 | 0,00658 | 1 | 0,03302 | 0,02005 | DIPPA_3 | DIPPA_3 | DIPPA_35723.mRNA.1 Uncharacterized RNA-binding protein C328.05 |
| 20,6927 | 17,2855 | 17,4865 | 20,5572 | 17,3606 | 17,4597 | 23,1123 | 22,1387 | 21,999 | 23,4873 | 22,9313 | 22,6311 | 9 | 9 | 9 | 31 | 31 | 31 | 42,059 | 0 | 35,481 | 7,1E+07 | 29 | 7 | 4 | 7 | 9 | 5 | 7 | 0,00626 | 1 | 0,02905 | 0,01907 | DIPPA_1 | DIPPA_1 | DIPPA_13986.mRNA.1 26S proteasome non-ATPase regulatory subunit 7 homolog B |
| 24,0874 | 22,5628 | 22,0095 | 23,6158 | 22,367 | 22,5941 | 26,303 | 26,9572 | 26,5986 | 27,0778 | 26,6751 | 27,5913 | 4 | 4 | 4 | 39,5 | 39,5 | 39,5 | 25,78 | 0 | 75,66 | 4,2E+08 | 14 | 2 | 3 | 2 | 2 | 3 | 3 | 0,01248 | 1 | 0,02759 | 0,03777 | DIPPA_0 | DIPPA_0 | DIPPA_01396.mRNA.1 Endo-1 3-1 4-beta-glycanase ExsH |
| 25 | 20,6861 | 20,9852 | 23,7014 | 21,6327 | 21,2581 | 25,9317 | 24,1359 | 24,6547 | 25,6315 | 25,7031 | 25,3376 | 15 | 15 | 15 | 59,9 | 59,9 | 59,9 | 19,739 | 0 | 127,99 | 6,2E+08 | 78 | 14 | 8 | 11 | 14 | 11 | 11 | 0,00545 | 0,99912 | 0,02636 | 0,01663 | DIPPA_1 | DIPPA_1 | DIPPA_12097.mRNA.2 hypothetical protein;DIPPA_12097.mRNA.1 hypothetical protein |
| 17,705 | 16,5939 | 15,8419 | 18,8061 | 14,5494 | 16,7315 | 20,76 | 19,2686 | 21,211 | 21,9495 | 19,2572 | 21,3592 | 5 | 5 | 5 | 35,9 | 35,9 | 35,9 | 25,456 | 0 | 14,322 | 9624400 | 9 | 2 | 3 | 2 | 3 | 1 | 3 | 0,00438 | 0,99853 | 0,01798 | 0,01339 | DIPPA_1 | DIPPA_1 | DIPPA_11490.mRNA.1 hypothetical protein;DIPPA_08902.mRNA.1 hypothetical protein |
| 16,8598 | 14,026 | 13,0039 | 17,4717 | 13,9836 | 12,4026 | 19,6521 | 19,0278 | 19,584 | 20,5808 | 20,7927 | 19,2602 | 4 | 4 | 4 | 17,6 | 17,6 | 17,6 | 39,875 | 0 | 40,524 | 6249700 | 15 | 3 | 2 | 2 | 4 | 1 | 1 | 0,00183 | 1 | 0,0106 | 0,00561 | DIPPA_0 | DIPPA_0 | DIPPA_03829.mRNA.1 Ribose-phosphate pyrophosphokinase A |
| 10,494 | 13,1413 | 12,7391 | 13,4345 | 11,1158 | 11,811 | 18,7087 | 20,8048 | 21,0056 | 19,2488 | 20,916 | 20,5527 | 7 | 7 | 7 | 4,6 | 4,6 | 4,6 | 287,64 | 0 | 20,888 | 4650800 | 12 | 1 | 5 | 3 | 3 | 2 | 3 | 0,00132 | 1 | 0,00435 | 0,00405 | DIPPA_1 | DIPPA_1 | DIPPA_14118.mRNA.1 Inositol-1 4 5-trisphosphate 5-phosphatase 1;DIPPA_14118.mRNA.2 Inositol-1 4 5-trisphosphate 5-phosphatase 1 |
| 22,2846 | 15,7031 | 16,52 | 19,5889 | 17,0474 | 17,8924 | 24,8226 | 21,3388 | 21,9207 | 23,0214 | 22,9192 | 22,5802 | 16 | 16 | 16 | 35,2 | 35,2 | 35,2 | 60,009 | 0 | 92,447 | 1,7E+08 | 48 | 16 | 4 | 6 | 13 | 5 | 9 | 0,00104 | 0,99997 | -0,007 | -0,0032 | DIPPA_1 | DIPPA_1 | DIPPA_10005.mRNA.1 hypothetical protein |
| 19,4463 | 17,1083 | 16,9145 | 19,765 | 16,5428 | 17,1903 | 21,5753 | 22,144 | 22,7062 | 23,1384 | 22,126 | 22,5422 | 7 | 7 | 7 | 59,2 | 59,2 | 59,2 | 23,373 | 0 | 22,301 | 3E+07 | 16 | 6 | 3 | 3 | 5 | 2 | 3 | 0,00247 | 1 | -0,0097 | -0,0076 | DIPPA_3 | DIPPA_3 | DIPPA_31513.mRNA.1 putative p22 protein precursor;DIPPA_31510.mRNA.1 putative p22 protein precursor |
| 23,027 | 16,084 | 13,9503 | 23,301 | 12,498 | 17,2927 | 23,8137 | 20,6174 | 19,9266 | 26,0592 | 18,9279 | 22,2093 | 15 | 15 | 15 | 77,8 | 77,8 | 77,8 | 26,308 | 0 | 180,01 | 2,7E+08 | 42 | 13 | 2 | 1 | 14 | 2 | 6 | 0,0008 | 0,9993 | -0,0102 | -0,0024 | DIPPA_2 | DIPPA_2 | DIPPA_27067.mRNA.1 hypothetical protein unknown function |
| 18,525 | 19,3275 | 18,4049 | 19,6852 | 17,8605 | 18,765 | 21,7454 | 24,0922 | 23,8186 | 23,2554 | 23,9788 | 24,2745 | 22 | 22 | 5 | 30,2 | 30,2 | 8,5 | 110,41 | 0 | 87,545 | 1,3E+08 | 69 | 2 | 1 | 2 | 4 | 0 | 4 | 0,00974 | 1 | -0,0178 | -0,0296 | DIPPA_3 | DIPPA_3 | DIPPA_30608.mRNA.1 Alpha-xylosidase |
| 13,0399 | 12,1016 | 13,7563 | 14,0178 | 13,0507 | 11,8985 | 18,981 | 19,4657 | 19,7165 | 19,4929 | 19,2682 | 18,9074 | 2 | 2 | 2 | 8,1 | 8,1 | 8,1 | 35,923 | 0 | 15,545 | 920690 | 5 | 1 | 0 | 2 | 1 | 1 | 1 | 0,00977 | 1 | -0,0231 | -0,0297 | DIPPA_2 | DIPPA_2 | DIPPA_28945.mRNA.1 Actin-related protein 2/3 complex subunit 2 |
| 17,9834 | 14,0126 | 13,9524 | 16,4342 | 14,4099 | 15,1761 | 19,251 | 20,3954 | 19,8731 | 19,5135 | 20,3277 | 19,7827 | 6 | 6 | 6 | 20,6 | 20,6 | 20,6 | 54,472 | 0 | 18,087 | 8338500 | 6 | 5 | 1 | 1 | 2 | 1 | 1 | 0,00538 | 0,99818 | -0,0239 | -0,0164 | DIPPA_0 | DIPPA_0 | DIPPA_02442.mRNA.1 Quinolinate synthase chloroplastic |
| 20,2613 | 17,1182 | 17,378 | 20,0531 | 16,7559 | 18,0301 | 22,1638 | 21,9238 | 22,2733 | 22,9193 | 22,5501 | 23,1001 | 9 | 9 | 3 | 44,8 | 44,8 | 19,3 | 36,214 | 0 | 69,501 | 4,2E+07 | 29 | 2 | 0 | 1 | 3 | 2 | 2 | 0,00642 | 1 | -0,0272 | -0,0196 | DIPPA_2 | DIPPA_2 | DIPPA_22292.mRNA.1 Protein disulfide-isomerase like 2-1 |
| 19,5969 | 10,8276 | 10,9479 | 19,5136 | 10,7323 | 11,2087 | 21,3515 | 20,2241 | 19,7403 | 21,4933 | 20,4051 | 18,5047 | 5 | 5 | 5 | 27,6 | 27,6 | 27,6 | 25,689 | 0 | 15,67 | 1,9E+07 | 13 | 5 | 0 | 1 | 5 | 0 | 0 | 0,0022 | 1 | -0,0274 | -0,0067 | DIPPA_1 | DIPPA_1 | DIPPA_14792.mRNA.1 hypothetical protein |
| 18,3469 | 14,0253 | 14,2703 | 15,6791 | 16,4688 | 14,5944 | 19,2966 | 20,1104 | 20,881 | 19,7647 | 23,0732 | 21,4337 | 8 | 8 | 8 | 30,3 | 30,3 | 30,3 | 51,031 | 0 | 31,537 | 1,1E+07 | 14 | 7 | 1 | 2 | 3 | 2 | 2 | 0,00727 | 1 | -0,0333 | -0,0221 | DIPPA_1 | DIPPA_1 | DIPPA_14510.mRNA.1 L-rhamnonate dehydratase |
| 18,5083 | 13,5026 | 15,4821 | 17,9284 | 14,1809 | 15,4881 | 21,324 | 19,7647 | 20,8077 | 21,716 | 20,6784 | 21,111 | 5 | 5 | 5 | 22 | 22 | 22 | 37,718 | 0 | 14,262 | 1,3E+07 | 12 | 5 | 1 | 3 | 3 | 2 | 3 | 0,00627 | 1 | -0,0348 | -0,0191 | DIPPA_2 | DIPPA_2 | DIPPA_22316.mRNA.1 Putative monooxygenase Rv1533 |
| 19,2309 | 20,2446 | 19,9415 | 20,2864 | 19,3281 | 19,9362 | 21,6444 | 24,5539 | 24,6317 | 23,4721 | 24,4171 | 24,5966 | 16 | 16 | 16 | 30 | 30 | 30 | 61,661 | 0 | 80,365 | 1,3E+08 | 77 | 10 | 14 | 15 | 13 | 13 | 16 | 0,03678 | 1 | -0,0446 | -0,1085 | DIPPA_0 | DIPPA_0 | DIPPA_05719.mRNA.1 hypothetical protein |
| 18,0643 | 13,6668 | 13,6966 | 18,6487 | 12,8166 | 14,1408 | 20,314 | 19,4499 | 19,3816 | 21,5165 | 19,083 | 20,0494 | 7 | 7 | 7 | 25,3 | 25,3 | 25,3 | 44,244 | 0 | 19,239 | 1,2E+07 | 11 | 5 | 1 | 1 | 6 | 1 | 2 | 0,00854 | 1 | -0,0595 | -0,026 | DIPPA_0 | DIPPA_0 | DIPPA_09865.mRNA.1 RAC family serine/threonine-protein kinase homolog |
| 21,4101 | 15,6617 | 15,8696 | 19,4023 | 16,0298 | 17,7238 | 23,8228 | 20,4537 | 21,2202 | 22,3743 | 20,9551 | 22,0817 | 11 | 11 | 11 | 51,7 | 51,7 | 51,7 | 33,91 | 0 | 38,137 | 6,6E+07 | 30 | 11 | 2 | 4 | 9 | 4 | 6 | 0,01113 | 1 | -0,0715 | -0,0337 | DIPPA_2 | DIPPA_2 | DIPPA_25913.mRNA.1 Uncharacterized protein L780 |
| 20,1814 | 14,1693 | 10,4409 | 19,6231 | 10,7077 | 14,6878 | 22,0347 | 19,3559 | 18,9467 | 21,8672 | 19,5805 | 21,1827 | 8 | 8 | 8 | 42,7 | 42,7 | 42,7 | 34,549 | 0 | 18,211 | 2,7E+07 | 12 | 6 | 1 | 0 | 6 | 0 | 2 | 0,00647 | 1 | -0,0756 | -0,0197 | DIPPA_1 | DIPPA_1 | DIPPA_17900.mRNA.1 Cytochrome c peroxidase mitochondrial |
| 24,0366 | 23,3298 | 23,0329 | 24,2016 | 23,2271 | 23,2026 | 25,8456 | 27,2151 | 27,3655 | 26,7948 | 27,833 | 27,6594 | 59 | 59 | 59 | 77,8 | 77,8 | 77,8 | 88,101 | 0 | 323,31 | 3,4E+09 | 413 | 53 | 47 | 52 | 58 | 52 | 53 | 0,06044 | 1 | -0,0773 | -0,1743 | DIPPA_0 | DIPPA_0 | DIPPA_03893.mRNA.1 Cell division cycle protein 48 homolog |
| 19,8044 | 12,8816 | 11,8486 | 18,5512 | 13,057 | 13,1685 | 23,0236 | 19,1628 | 18,0235 | 22,4665 | 19,8265 | 18,8456 | 10 | 10 | 10 | 24,4 | 24,4 | 24,4 | 68,429 | 0 | 67,035 | 4E+07 | 17 | 9 | 1 | 1 | 7 | 0 | 2 | 0,0086 | 1 | -0,0807 | -0,0261 | DIPPA_2 | DIPPA_2 | DIPPA_29834.mRNA.1 Isocitrate lyase 1 |
| 17,8 | 17,1636 | 17,3741 | 18,7216 | 16,502 | 17,3621 | 21,2945 | 22,637 | 23,0267 | 23,1797 | 22,8551 | 23,1333 | 21 | 21 | 21 | 30,7 | 30,7 | 30,7 | 97,979 | 0 | 62,16 | 6,4E+07 | 58 | 10 | 13 | 16 | 13 | 10 | 17 | 0,04187 | 1 | -0,0827 | -0,1229 | DIPPA_3 | DIPPA_3 | DIPPA_33869.mRNA.1 Kinesin heavy chain |
| 15,8308 | 12,1519 | 13,9539 | 16,9167 | 12,3567 | 12,9245 | 20,2618 | 18,7553 | 20,2878 | 21,2316 | 19,8867 | 19,4392 | 4 | 4 | 4 | 10,4 | 10,4 | 10,4 | 67,455 | 0 | 15,359 | 5346800 | 6 | 2 | 0 | 2 | 4 | 0 | 1 | 0,01619 | 1 | -0,0871 | -0,0488 | DIPPA_1 | DIPPA_1 | DIPPA_19569.mRNA.1 Putative galactocerebrosidase;TR92915__c3_g2_i1__m.19683 #MITO 1 HMMTOP #PFAM PF02057.10 Glyco_hydro_59 Glycosyl hydrolase family 59;2 x PF00652.17 Ricin_B_lectin Ricin-type beta-trefoil lectin domain;PF03498.9 CDtoxinA Cytolethal di |
| 17,7812 | 10,8279 | 12,6905 | 17,3063 | 12,1108 | 12,2147 | 21,5088 | 19,9972 | 19,1889 | 22,1442 | 19,4379 | 19,6872 | 11 | 11 | 11 | 13,8 | 13,8 | 13,8 | 130,28 | 0 | 24,55 | 2,5E+07 | 13 | 9 | 0 | 2 | 9 | 1 | 1 | 0,01359 | 1 | -0,1107 | -0,0411 | DIPPA_1 | DIPPA_1 | DIPPA_17668.mRNA.1 Uncharacterized WD repeat-containing protein alr3466 |
| 19,151 | 19,3862 | 19,3214 | 19,819 | 19,0453 | 19,3512 | 22,6421 | 24,0245 | 24,4605 | 23,4382 | 24,1888 | 24,5621 | 18 | 18 | 18 | 37,4 | 37,4 | 37,4 | 61,775 | 0 | 81,415 | 1,3E+08 | 72 | 11 | 14 | 16 | 14 | 15 | 16 | 0,19373 | 1 | -0,119 | -0,505 | DIPPA_0 | DIPPA_0 | DIPPA_09508.mRNA.1 Polyadenylate-binding protein cytoplasmic and nuclear |
| 16,136 | 14,0122 | 13,807 | 17,403 | 13,2087 | 13,7181 | 20,9255 | 20,1821 | 20,9516 | 22,0526 | 20,252 | 20,9848 | 15 | 15 | 15 | 13,7 | 13,7 | 13,7 | 179,28 | 0 | 113,22 | 1,8E+07 | 22 | 7 | 2 | 4 | 12 | 2 | 4 | 0,02765 | 1 | -0,1249 | -0,0823 | DIPPA_3 | DIPPA_3 | DIPPA_32832.mRNA.1 ABC transporter F family member 4 |
| 15,9035 | 11,4496 | 13,2819 | 15,3533 | 11,6486 | 14,0375 | 19,7353 | 20,0371 | 19,1489 | 19,7768 | 20,4385 | 19,4512 | 4 | 4 | 4 | 11,5 | 11,5 | 11,5 | 62,703 | 0 | 10,565 | 3897900 | 9 | 3 | 0 | 1 | 3 | 0 | 1 | 0,0268 | 1 | -0,1348 | -0,0799 | DIPPA_1 | DIPPA_1 | DIPPA_16631.mRNA.1 hypothetical protein |
| 17,0155 | 19,7178 | 17,1787 | 18,9007 | 17,4474 | 18,0266 | 21,3949 | 23,6796 | 22,5622 | 21,6567 | 22,5713 | 23,0596 | 6 | 6 | 6 | 48,9 | 48,9 | 48,9 | 21,43 | 0 | 17,788 | 2,3E+07 | 11 | 2 | 4 | 3 | 4 | 3 | 6 | 0,05475 | 1 | -0,1542 | -0,1588 | DIPPA_0 | DIPPA_0 | DIPPA_04466.mRNA.1 hypothetical protein |
| 19,3988 | 13,3738 | 14,5515 | 18,1242 | 14,7119 | 14,9552 | 23,025 | 21,2354 | 21,2436 | 21,8419 | 21,5011 | 21,7763 | 13 | 13 | 13 | 15,1 | 15,1 | 15,1 | 111,1 | 0 | 36,373 | 4,8E+07 | 28 | 10 | 2 | 4 | 8 | 4 | 4 | 0,02427 | 1 | -0,1557 | -0,0726 | DIPPA_0 | DIPPA_0 | DIPPA_02798.mRNA.1 hypothetical protein |
| 16,5265 | 12,829 | 11,1883 | 17,6144 | 10,9478 | 12,4496 | 19,5649 | 20,1717 | 19,9474 | 21,5922 | 19,9838 | 19,0687 | 10 | 10 | 10 | 23,5 | 23,5 | 23,5 | 79,588 | 0 | 37,099 | 1,2E+07 | 19 | 6 | 2 | 1 | 9 | 1 | 3 | 0,02027 | 1 | -0,156 | -0,0609 | DIPPA_0 | DIPPA_0 | DIPPA_04189.mRNA.1 Protein SEY1 |
| 17,8718 | 12,0717 | 13,987 | 18,8756 | 11,3806 | 14,1448 | 19,7915 | 19,8635 | 19,3967 | 20,0472 | 19,7001 | 19,2287 | 4 | 4 | 4 | 31,5 | 31,5 | 31,5 | 18,829 | 0 | 33,708 | 8307100 | 9 | 1 | 1 | 1 | 4 | 0 | 1 | 0,0188 | 1 | -0,1568 | -0,0565 | DIPPA_3 | DIPPA_3 | DIPPA_31393.mRNA.1 Ran-binding protein 1 homolog b |
| 18,3597 | 17,4636 | 16,6569 | 19,4342 | 16,7703 | 16,7547 | 20,6378 | 22,2213 | 22,4185 | 21,9095 | 22,5444 | 22,2026 | 8 | 8 | 8 | 23,5 | 23,5 | 23,5 | 40,147 | 0 | 20,965 | 3,1E+07 | 23 | 5 | 4 | 3 | 8 | 3 | 5 | 0,05409 | 1 | -0,1597 | -0,1569 | DIPPA_1 | DIPPA_1 | DIPPA_10221.mRNA.1 hypothetical protein |
| 16,3291 | 13,709 | 11,2759 | 17,1739 | 12,305 | 12,3291 | 22,365 | 19,4658 | 19,5883 | 23,3697 | 19,0231 | 21,0533 | 19 | 19 | 19 | 8,3 | 8,3 | 8,3 | 387,76 | 0 | 128,96 | 5,2E+07 | 29 | 14 | 3 | 1 | 18 | 2 | 5 | 0,0253 | 1 | -0,1646 | -0,0755 | DIPPA_1 | DIPPA_1 | DIPPA_10233.mRNA.1 FACT complex subunit SPT16 |
| 21,3694 | 11,9497 | 15,038 | 18,5794 | 14,4807 | 15,7935 | 23,4814 | 20,2229 | 19,7513 | 20,941 | 20,449 | 20,6176 | 9 | 9 | 9 | 38,3 | 38,3 | 38,3 | 39,086 | 0 | 23,585 | 4,8E+07 | 13 | 9 | 0 | 2 | 3 | 1 | 3 | 0,01819 | 1 | -0,1655 | -0,0547 | DIPPA_0 | DIPPA_0 | DIPPA_01769.mRNA.2 Metacaspase-1;DIPPA_01769.mRNA.1 Metacaspase-1 |
| 20,8667 | 14,1298 | 14,1144 | 19,686 | 14,1698 | 15,7751 | 22,9429 | 20,1965 | 19,3656 | 22,6349 | 19,7573 | 20,4398 | 12 | 12 | 12 | 48,4 | 48,4 | 48,4 | 40,953 | 0 | 102,2 | 5,7E+07 | 37 | 12 | 1 | 3 | 11 | 2 | 4 | 0,02077 | 1 | -0,1734 | -0,0623 | DIPPA_2 | DIPPA_2 | DIPPA_25791.mRNA.1 Putative dTDP-D-glucose 4 6-dehydratase |
| 19,8194 | 15,3939 | 16,3339 | 20,7153 | 14,0812 | 17,2763 | 22,3627 | 19,2142 | 21,996 | 23,8238 | 19,8631 | 22,8314 | 11 | 11 | 11 | 36,3 | 36,3 | 36,3 | 51,625 | 0 | 57,442 | 5,6E+07 | 21 | 9 | 2 | 6 | 10 | 1 | 7 | 0,02506 | 1 | -0,1752 | -0,0748 | DIPPA_2 | DIPPA_2 | DIPPA_20630.mRNA.1 RAC family serine/threonine-protein kinase homolog;DIPPA_11290.mRNA.1 RAC family serine/threonine-protein kinase homolog |
| 19,4304 | 13,0371 | 12,0527 | 20,2368 | 11,0359 | 13,7294 | 22,4386 | 19,9379 | 19,3761 | 24,1625 | 19,903 | 20,7766 | 10 | 10 | 10 | 41,3 | 41,3 | 41,3 | 40,126 | 0 | 41,898 | 4,7E+07 | 18 | 7 | 1 | 0 | 10 | 0 | 2 | 0,01713 | 1 | -0,184 | -0,0516 | DIPPA_0 | DIPPA_0 | DIPPA_00335.mRNA.1 Cycloartenol-C-24-methyltransferase 1 |
| 15,6375 | 10,9515 | 13,4746 | 16,2116 | 11,8471 | 12,558 | 21,1142 | 20,3458 | 22,206 | 22,5205 | 19,801 | 21,6233 | 10 | 10 | 10 | 4,4 | 4,4 | 4,4 | 413,28 | 0 | 24,239 | 2,5E+07 | 19 | 8 | 1 | 5 | 10 | 0 | 3 | 0,03251 | 1 | -0,1844 | -0,0963 | DIPPA_0 | DIPPA_0 | DIPPA_07091.mRNA.1 Protein transport protein Sec24-like CEF |
| 14,3374 | 14,5052 | 13,5023 | 14,9937 | 14,9549 | 12,9495 | 18,9431 | 18,5415 | 19,747 | 19,3024 | 19,5026 | 19,2539 | 3 | 3 | 3 | 8,2 | 8,2 | 8,2 | 52,9 | 0 | 12,412 | 1937700 | 5 | 1 | 2 | 1 | 0 | 1 | 1 | 0,08821 | 1 | -0,1844 | -0,2482 | DIPPA_1 | DIPPA_1 | DIPPA_11434.mRNA.1 hypothetical protein |
| 16,7659 | 11,203 | 13,3652 | 16,6962 | 11,8359 | 13,3841 | 21,1706 | 18,6676 | 21,0064 | 21,8265 | 20,0748 | 21,2098 | 8 | 8 | 8 | 11,4 | 11,4 | 11,4 | 120,06 | 0 | 31,803 | 1,3E+07 | 15 | 4 | 1 | 3 | 7 | 0 | 3 | 0,0302 | 1 | -0,194 | -0,0897 | DIPPA_3 | DIPPA_3 | DIPPA_32844.mRNA.1 Protein transport protein Sec24-like At3g07100 |
| 16,4586 | 12,2303 | 11,3018 | 16,5135 | 11,012 | 13,0498 | 19,8838 | 18,9538 | 18,9618 | 20,6033 | 19,8518 | 20,3708 | 5 | 5 | 5 | 14,7 | 14,7 | 14,7 | 56,183 | 0 | 13,374 | 4582600 | 7 | 3 | 1 | 1 | 5 | 0 | 0 | 0,02902 | 1 | -0,1949 | -0,0863 | DIPPA_1 | DIPPA_1 | DIPPA_15967.mRNA.1 Probable succinyl-CoA:3-ketoacid coenzyme A transferase mitochondrial |
| 14,9235 | 11,7701 | 10,786 | 15,6406 | 10,7768 | 11,6661 | 19,5648 | 18,5839 | 20,7434 | 20,2927 | 19,6086 | 19,748 | 7 | 7 | 7 | 12,8 | 12,8 | 12,8 | 85,312 | 0 | 15,323 | 4251500 | 11 | 5 | 2 | 1 | 6 | 1 | 2 | 0,03496 | 1 | -0,2013 | -0,1034 | DIPPA_2 | DIPPA_2 | DIPPA_20149.mRNA.1 Armadillo repeat-containing kinesin-like protein 3 |
| 18,0852 | 15,8617 | 15,5021 | 18,1126 | 16,1001 | 15,8606 | 19,6992 | 21,88 | 21,9224 | 20,6662 | 22,7123 | 22,0179 | 7 | 7 | 7 | 21,9 | 21,9 | 21,9 | 50,377 | 0 | 10,09 | 1,8E+07 | 18 | 6 | 4 | 3 | 7 | 3 | 4 | 0,06735 | 1 | -0,2081 | -0,193 | DIPPA_2 | DIPPA_2 | DIPPA_23053.mRNA.1 Putative serine/threonine-protein kinase A |
| 22,0838 | 15,2439 | 16,8408 | 20,4125 | 16,6259 | 17,8035 | 23,6382 | 20,2665 | 21,7874 | 22,3764 | 22,3919 | 22,2426 | 8 | 8 | 8 | 63,9 | 63,9 | 63,9 | 20,647 | 0 | 53,136 | 6,3E+07 | 25 | 6 | 2 | 3 | 6 | 3 | 6 | 0,03223 | 1 | -0,2244 | -0,0955 | DIPPA_2 | DIPPA_2 | DIPPA_20982.mRNA.1 hypothetical protein;DIPPA_10870.mRNA.1 hypothetical protein;DIPPA_10854.mRNA.1 hypothetical protein |
| 20,6736 | 19,7205 | 18,823 | 20,5503 | 19,3129 | 20,1104 | 22,2839 | 23,3949 | 23,278 | 23,5284 | 23,9099 | 24,2276 | 7 | 7 | 7 | 51,2 | 51,2 | 51,2 | 19,482 | 0 | 19,872 | 7,6E+07 | 22 | 6 | 5 | 6 | 6 | 6 | 7 | 0,14512 | 1 | -0,2522 | -0,3907 | DIPPA_2 | DIPPA_2 | DIPPA_26143.mRNA.1 Penta-EF hand domain-containing protein 1;DIPPA_26141.mRNA.1 Penta-EF hand domain-containing protein 1 |
| 20,8244 | 11,5917 | 16,4289 | 18,9931 | 14,4647 | 16,1443 | 24,1621 | 18,4164 | 23,5681 | 23,3902 | 21,3511 | 23,1062 | 13 | 13 | 13 | 29,5 | 29,5 | 29,5 | 68,897 | 0 | 47,149 | 8,5E+07 | 26 | 13 | 0 | 2 | 7 | 1 | 3 | 0,0285 | 1 | -0,2524 | -0,0848 | DIPPA_3 | DIPPA_3 | DIPPA_30384.mRNA.1 hypothetical protein |
| 17,5206 | 18,2865 | 16,606 | 19,1835 | 17,0219 | 16,9692 | 21,9725 | 19,4137 | 22,2285 | 23,4395 | 19,312 | 23,2228 | 7 | 7 | 7 | 18,3 | 18,3 | 18,3 | 65,388 | 0 | 32,234 | 3,8E+07 | 16 | 4 | 1 | 3 | 7 | 1 | 4 | 0,10431 | 1 | -0,2538 | -0,2896 | DIPPA_0 | DIPPA_0 | DIPPA_06385.mRNA.1 Protein disulfide-isomerase 2-3 |
| 19,9019 | 18,7946 | 17,7633 | 20,5497 | 17,9042 | 18,7707 | 22,2812 | 23,4307 | 23,0347 | 23,5939 | 23,2372 | 23,1167 | 11 | 11 | 11 | 51,9 | 51,9 | 51,9 | 32 | 0 | 231,92 | 6,2E+07 | 44 | 10 | 7 | 6 | 10 | 7 | 8 | 0,09143 | 1 | -0,255 | -0,2566 | DIPPA_3 | DIPPA_3 | DIPPA_34761.mRNA.1 hypothetical protein;DIPPA_34761.mRNA.2 hypothetical protein |
| 19,7582 | 19,5974 | 17,478 | 22,1669 | 18,1403 | 17,3188 | 21,6305 | 20,0252 | 19,2036 | 23,246 | 19,4826 | 18,2135 | 3 | 3 | 3 | 17,5 | 17,5 | 17,5 | 37,944 | 0 | 50,325 | 2,8E+07 | 7 | 2 | 1 | 1 | 3 | 1 | 1 | 0,05458 | 1 | -0,2641 | -0,1583 | DIPPA_3 | DIPPA_3 | DIPPA_33209.mRNA.1 Cold shock domain-containing protein 4 |
| 17,4741 | 13,503 | 12,1873 | 18,2621 | 12,3735 | 13,3224 | 20,2252 | 19,6316 | 19,6372 | 21,9889 | 19,5136 | 19,8025 | 5 | 5 | 5 | 13 | 13 | 13 | 50,86 | 0 | 17,099 | 1,2E+07 | 7 | 4 | 0 | 1 | 5 | 1 | 2 | 0,03705 | 1 | -0,2645 | -0,1093 | DIPPA_3 | DIPPA_3 | DIPPA_33575.mRNA.1 hypothetical protein AMSG_11957;DIPPA_33575.mRNA.2 hypothetical protein AMSG_11957 |
| 24,5374 | 21,7686 | 21,3671 | 24,4067 | 22,3768 | 21,6903 | 25,6548 | 24,6184 | 25,2635 | 26,358 | 26,2753 | 25,9498 | 13 | 13 | 13 | 54,1 | 54,1 | 54,1 | 22,758 | 0 | 157,13 | 6,2E+08 | 71 | 12 | 11 | 12 | 13 | 12 | 11 | 0,07267 | 1 | -0,2669 | -0,2073 | DIPPA_1 | DIPPA_1 | DIPPA_12072.mRNA.1 hypothetical protein |
| 9,61126 | 13,6081 | 13,4108 | 13,2275 | 12,7054 | 11,5191 | 19,4135 | 20,8518 | 21,0133 | 19,9421 | 18,9841 | 20,3344 | 7 | 7 | 7 | 5,2 | 5,2 | 5,2 | 199,46 | 0 | 18,023 | 3815300 | 14 | 1 | 4 | 5 | 4 | 0 | 3 | 0,06857 | 1 | -0,2739 | -0,1963 | DIPPA_1 | DIPPA_1 | DIPPA_16119.mRNA.1 hypothetical protein |
| 21,2216 | 19,4083 | 20,0859 | 22,1937 | 19,8738 | 19,5002 | 23,0644 | 23,7608 | 24,1659 | 24,4138 | 24,4764 | 23,9372 | 8 | 8 | 8 | 49,3 | 49,3 | 49,3 | 23,232 | 0 | 51,208 | 8,6E+07 | 40 | 6 | 4 | 6 | 7 | 5 | 6 | 0,10267 | 1 | -0,284 | -0,2855 | DIPPA_1 | DIPPA_1 | DIPPA_14508.mRNA.1 Selenoprotein M |
| 12,9421 | 13,5877 | 12,9644 | 15,5776 | 12,1214 | 12,6637 | 18,9071 | 19,6903 | 19,6328 | 20,028 | 19,6712 | 19,4782 | 8 | 8 | 8 | 10,1 | 10,1 | 10,1 | 118 | 0 | 10,64 | 4401000 | 10 | 1 | 2 | 2 | 6 | 1 | 2 | 0,09457 | 1 | -0,2895 | -0,2647 | DIPPA_0 | DIPPA_0 | DIPPA_08161.mRNA.1 Kinesin-like protein KIN-1 |
| 14,4776 | 12,4013 | 11,7486 | 16,8031 | 10,1174 | 12,6198 | 19,7388 | 19,2381 | 19,2681 | 20,8532 | 19,33 | 19,5367 | 4 | 4 | 4 | 13,5 | 13,5 | 13,5 | 56,419 | 0 | 14,761 | 4260200 | 5 | 1 | 0 | 1 | 4 | 0 | 2 | 0,04932 | 1 | -0,3043 | -0,1438 | DIPPA_0 | DIPPA_0 | DIPPA_03265.mRNA.1 6-phosphofructo-2-kinase/fructose-2 6-bisphosphatase |
| 14,7037 | 17,5728 | 16,4397 | 16,2056 | 16,5507 | 16,8952 | 18,7188 | 22,8672 | 22,3967 | 21,9993 | 22,8943 | 23,4435 | 4 | 4 | 4 | 9,8 | 9,8 | 9,8 | 80,249 | 0 | 30,623 | 1,4E+07 | 23 | 1 | 3 | 3 | 2 | 4 | 4 | 0,13393 | 1 | -0,3118 | -0,3635 | TR10862 | TR10862 | TR108626__c4_g1_i1__m.24593 #MITO 1 Predotar #PFAM PF00933.16 Glyco_hydro_3 Glycosyl hydrolase family 3 N terminal domain;PF01915.17 Glyco_hydro_3_C Glycosyl hydrolase family 3 C-terminal domain;PF14310.1 Fn3-like Fibronectin type III-like domain #BLAS |
| 18,2022 | 14,9805 | 15,4391 | 18,4805 | 14,8922 | 16,2126 | 21,8821 | 21,7602 | 22,2143 | 22,4176 | 22,2128 | 22,4273 | 15 | 15 | 15 | 22,5 | 22,5 | 22,5 | 94,393 | 0 | 71,269 | 4,2E+07 | 39 | 9 | 6 | 8 | 14 | 6 | 10 | 0,07786 | 1 | -0,3211 | -0,2211 | DIPPA_0 | DIPPA_0 | DIPPA_01624.mRNA.1 Threonine--tRNA ligase cytoplasmic |
| 10,3536 | 11,4462 | 10,5667 | 13,1423 | 9,86761 | 10,3256 | 18,658 | 20,1072 | 20,1272 | 19,9627 | 19,5414 | 19,1788 | 6 | 6 | 3 | 3,4 | 3,4 | 1,8 | 252,12 | 0 | 11,76 | 2088100 | 8 | 2 | 0 | 0 | 3 | 0 | 0 | 0,10836 | 1 | -0,323 | -0,2999 | DIPPA_1 | DIPPA_1 | DIPPA_19138.mRNA.1 General negative regulator of transcription subunit 1;TR113577__c0_g1_i2__m.26013 #MITO 2 Phobius,HMMTOP #PFAM 3 x PF12842.2 DUF3819 Domain of unknown function (DUF3819);4 x PF09686.5 Plasmid_RAQPRD Plasmid protein of unknown function |
| 13,1984 | 12,2737 | 12,4158 | 15,1882 | 11,6455 | 12,0703 | 20,5157 | 19,4627 | 19,5654 | 20,7099 | 20,451 | 19,7124 | 3 | 3 | 3 | 4,8 | 4,8 | 4,8 | 106,23 | 0 | 24,684 | 3029000 | 3 | 1 | 1 | 1 | 3 | 0 | 0 | 0,10591 | 1 | -0,3387 | -0,2937 | DIPPA_0 | DIPPA_0 | DIPPA_06897.mRNA.1 Aminopeptidase M1 |
| 14,4197 | 12,1973 | 14 | 15,4145 | 11,09 | 15,1515 | 20,3711 | 20,6657 | 21,7904 | 20,9372 | 19,6491 | 22,9031 | 10 | 10 | 10 | 7,2 | 7,2 | 7,2 | 196,75 | 0 | 14,585 | 1,2E+07 | 18 | 5 | 3 | 5 | 8 | 1 | 9 | 0,07839 | 1 | -0,3463 | -0,2224 | DIPPA_1 | DIPPA_1 | DIPPA_13555.mRNA.1 hypothetical protein |
| 21,1134 | 18,4715 | 18,4754 | 22,4436 | 18,012 | 18,6512 | 23,346 | 23,4243 | 23,6358 | 25,4662 | 23,8963 | 23,9409 | 29 | 29 | 28 | 61,1 | 61,1 | 59,9 | 76,295 | 0 | 159,56 | 3,7E+08 | 97 | 17 | 11 | 15 | 28 | 10 | 15 | 0,07475 | 1 | -0,3488 | -0,2128 | DIPPA_1 | DIPPA_1 | DIPPA_17318.mRNA.1 187-kDa microtubule-associated protein AIR9 |
| 15,6033 | 17,8668 | 17,9799 | 18,3579 | 16,5125 | 17,6695 | 18,6209 | 23,6857 | 23,6215 | 22,2933 | 23,2541 | 23,3155 | 50 | 50 | 50 | 33,7 | 33,7 | 33,7 | 239,64 | 0 | 212,3 | 1,3E+08 | 146 | 7 | 30 | 42 | 36 | 24 | 41 | 0,1429 | 1 | -0,3633 | -0,3853 | DIPPA_0 | DIPPA_0 | DIPPA_08914.mRNA.1 ATP-dependent RNA helicase DDX60 |
| 16,3735 | 10,1747 | 12,8413 | 17,4665 | 10,1691 | 12,867 | 20,8284 | 20,6293 | 20,6282 | 22,0306 | 18,0667 | 20,7836 | 8 | 8 | 8 | 12,1 | 12,1 | 12,1 | 101,29 | 0 | 14,096 | 1,3E+07 | 16 | 6 | 1 | 2 | 8 | 1 | 2 | 0,04552 | 1 | -0,371 | -0,1332 | DIPPA_3 | DIPPA_3 | DIPPA_34219.mRNA.1 hypothetical protein;TR53228__c1_g1_i1__m.6802 #MITO 2 Cello,Mitoprot #PFAM 3 x PF01323.15 DSBA DSBA-like thioredoxin domain;2 x PF00440.18 TetR_N Bacterial regulatory proteins, tetR family #BLASTP_UNIREF90 5e-09 Uncharacterized protei |
| 19,0145 | 13,1076 | 12,9993 | 19,1642 | 12,5832 | 14,4963 | 21,845 | 19,176 | 19,2997 | 22,9564 | 18,6968 | 20,9565 | 12 | 12 | 12 | 39,4 | 39,4 | 39,4 | 57,304 | 0 | 55,844 | 3E+07 | 19 | 8 | 0 | 1 | 11 | 1 | 2 | 0,0459 | 1 | -0,3741 | -0,1342 | DIPPA_1 | DIPPA_1 | DIPPA_12058.mRNA.1 hypothetical protein |
| 18,4085 | 14,8292 | 11,398 | 18,2644 | 13,6627 | 13,8834 | 21,4245 | 20,7479 | 18,9644 | 22,1682 | 20,7228 | 20,8743 | 9 | 9 | 9 | 21,1 | 21,1 | 21,1 | 54,347 | 0 | 11,812 | 2,3E+07 | 16 | 8 | 2 | 1 | 9 | 2 | 2 | 0,05357 | 1 | -0,3916 | -0,1555 | DIPPA_3 | DIPPA_3 | DIPPA_35637.mRNA.1 Heat shock protein 70 2 |
| 17,7215 | 17,7093 | 17,2378 | 19,4172 | 17,0841 | 17,3723 | 21,4966 | 22,7772 | 23,1064 | 23,138 | 23,3169 | 23,1112 | 16 | 16 | 16 | 35,7 | 35,7 | 35,7 | 69,81 | 0 | 45,319 | 5,3E+07 | 35 | 7 | 7 | 9 | 13 | 9 | 11 | 0,20668 | 1 | -0,4016 | -0,5345 | DIPPA_0 | DIPPA_0 | DIPPA_01497.mRNA.1 hypothetical protein |
| 21,1499 | 17,3884 | 17,1877 | 20,8593 | 17,8368 | 18,2493 | 22,4864 | 21,3095 | 21,434 | 22,9738 | 22,7253 | 22,3979 | 6 | 6 | 3 | 45,7 | 45,7 | 33,3 | 14,933 | 0 | 18,882 | 3,6E+07 | 23 | 3 | 2 | 1 | 3 | 2 | 3 | 0,09054 | 1 | -0,4065 | -0,2543 | DIPPA_2 | DIPPA_2 | DIPPA_26292.mRNA.1 Histone H2B.1 |
| 18,1999 | 11,6996 | 13,646 | 17,9669 | 11,9554 | 14,8536 | 21,9786 | 19,5079 | 19,7721 | 22,3823 | 18,9518 | 21,7178 | 7 | 7 | 7 | 21,8 | 21,8 | 21,8 | 53,417 | 0 | 16,272 | 1,6E+07 | 11 | 7 | 0 | 1 | 5 | 1 | 2 | 0,05454 | 1 | -0,4101 | -0,1582 | DIPPA_3 | DIPPA_3 | DIPPA_31643.mRNA.1 Cytoplasmic dynein 1 light intermediate chain 1 |
| 21,2223 | 19,8701 | 20,3197 | 22,4156 | 19,8684 | 20,4159 | 24,2686 | 25,0278 | 25,4658 | 26,1642 | 25,2921 | 25,9791 | 18 | 18 | 18 | 39,8 | 39,8 | 39,8 | 58,606 | 0 | 80,292 | 3,3E+08 | 103 | 14 | 15 | 16 | 16 | 14 | 15 | 0,18861 | 1 | -0,4292 | -0,4932 | DIPPA_2 | DIPPA_2 | DIPPA_20303.mRNA.1 Putative serine/threonine-protein kinase A;TR97218__c39_g1_i3__m.20937 #MITO 1 HMMTOP #PFAM PF00069.20 Pkinase Protein kinase domain;PF07714.12 Pkinase_Tyr Protein tyrosine kinase;2 x PF01636.18 APH Phosphotransferase enzyme family # |
| 21,1872 | 10,85 | 11,3351 | 19,7065 | 11,1529 | 13,8337 | 23,5699 | 20,3087 | 19,7971 | 24,0184 | 20,147 | 21,5502 | 18 | 18 | 18 | 42 | 42 | 42 | 67,842 | 0 | 128,21 | 1E+08 | 40 | 18 | 1 | 1 | 12 | 1 | 3 | 0,03539 | 1 | -0,4403 | -0,1046 | DIPPA_1 | DIPPA_1 | DIPPA_18784.mRNA.1 hypothetical protein |
| 21,2062 | 16,1611 | 16,4013 | 19,5092 | 17,5797 | 18,0077 | 23,6741 | 22,62 | 22,6855 | 23,3802 | 23,3065 | 23,7061 | 12 | 12 | 12 | 41,2 | 41,2 | 41,2 | 51,162 | 0 | 34,339 | 8,3E+07 | 36 | 11 | 2 | 5 | 10 | 5 | 8 | 0,09036 | 1 | -0,4427 | -0,2538 | DIPPA_1 | DIPPA_1 | DIPPA_12843.mRNA.1 RNA-binding protein BRN1 |
| 15,7861 | 17,477 | 17,6784 | 17,8011 | 16,7959 | 17,6972 | 20,3979 | 23,5709 | 23,6835 | 22,7892 | 23,4506 | 24,7903 | 50 | 50 | 50 | 16,3 | 16,3 | 16,3 | 474,79 | 0 | 178,28 | 1,6E+08 | 125 | 5 | 23 | 33 | 38 | 25 | 37 | 0,26493 | 1 | -0,4509 | -0,6634 | DIPPA_0 | DIPPA_0 | DIPPA_09931.mRNA.1 hypothetical protein |
| 10,5488 | 14,8579 | 14,1685 | 16,4989 | 10,9516 | 13,5233 | 19,3384 | 22,2929 | 22,1152 | 22,5226 | 19,8414 | 19,5706 | 7 | 7 | 6 | 8,8 | 8,8 | 7,9 | 106,48 | 0 | 11,856 | 8578300 | 9 | 1 | 1 | 1 | 5 | 1 | 1 | 0,07874 | 1 | -0,4662 | -0,2234 | DIPPA_1 | DIPPA_1 | DIPPA_18876.mRNA.1 Kinesin-like protein K39 |
| 22,2179 | 15,2928 | 16,9731 | 20,9469 | 16,7279 | 18,2617 | 24,0346 | 20,1114 | 22,3294 | 23,2103 | 22,0053 | 22,5199 | 10 | 10 | 10 | 57,3 | 57,3 | 57,3 | 28,341 | 0 | 60,709 | 9E+07 | 27 | 10 | 1 | 2 | 10 | 3 | 5 | 0,06991 | 1 | -0,4843 | -0,1999 | DIPPA_2 | DIPPA_2 | DIPPA_29233.mRNA.1 hypothetical protein |
| 14,5505 | 13,8107 | 12,8564 | 16,5816 | 13,3937 | 12,7786 | 19,3044 | 19,3336 | 19,0779 | 19,7407 | 19,8111 | 18,1364 | 3 | 3 | 3 | 8 | 8 | 8 | 56,067 | 0 | 60,375 | 3232800 | 4 | 1 | 2 | 1 | 2 | 1 | 1 | 0,14947 | 1 | -0,5121 | -0,4011 | DIPPA_3 | DIPPA_3 | DIPPA_30884.mRNA.2 hypothetical protein;DIPPA_30884.mRNA.3 hypothetical protein;DIPPA_30884.mRNA.1 hypothetical protein |
| 21,0294 | 12,9927 | 16,6447 | 19,847 | 14,5113 | 17,9117 | 24,7765 | 19,5292 | 22,3508 | 24,1874 | 21,2052 | 23,3453 | 9 | 9 | 9 | 36,9 | 36,9 | 36,9 | 37,778 | 0 | 38,208 | 8E+07 | 26 | 9 | 1 | 2 | 8 | 2 | 4 | 0,06659 | 1 | -0,5344 | -0,191 | DIPPA_0 | DIPPA_0 | DIPPA_02040.mRNA.1 hypothetical protein TraAM80_06864 |
| 19,6513 | 21,6938 | 16,3453 | 20,2651 | 19,302 | 19,7348 | 18,4608 | 24,8901 | 20,3461 | 22,7321 | 23,7124 | 24,2639 | 3 | 3 | 3 | 45,8 | 45,8 | 45,8 | 11,526 | 0 | 13,361 | 3,5E+07 | 16 | 1 | 3 | 2 | 2 | 2 | 2 | 0,12412 | 1 | -0,5372 | -0,3393 | DIPPA_2 | DIPPA_2 | DIPPA_21729.mRNA.1 Cytochrome c |
| 21,6323 | 12,4572 | 14,5985 | 19,5733 | 14,8794 | 15,883 | 23,5979 | 19,5517 | 20,0584 | 21,5275 | 20,5705 | 20,7077 | 12 | 12 | 12 | 56,5 | 56,5 | 56,5 | 37,361 | 0 | 44,775 | 7,9E+07 | 26 | 12 | 0 | 2 | 10 | 2 | 5 | 0,06113 | 1 | -0,5493 | -0,1762 | DIPPA_2 | DIPPA_2 | DIPPA_28118.mRNA.1 Malate dehydrogenase |
| 21,9877 | 13,1919 | 15,5222 | 21,0321 | 16,2009 | 15,1288 | 23,2879 | 19,7807 | 20,4592 | 24,0357 | 21,3683 | 20,5598 | 10 | 10 | 10 | 40 | 40 | 40 | 27,244 | 0 | 49,761 | 9,1E+07 | 19 | 9 | 1 | 3 | 10 | 4 | 2 | 0,05999 | 1 | -0,5533 | -0,1731 | DIPPA_3 | DIPPA_3 | DIPPA_32269.mRNA.1 hypothetical protein |
| 13,6145 | 11,6568 | 12,4769 | 16,2005 | 9,89814 | 13,3333 | 19,4779 | 19,0836 | 20,2887 | 19,6266 | 18,9657 | 19,1172 | 2 | 2 | 2 | 16,6 | 16,6 | 16,6 | 21,771 | 0 | 10,972 | 1454200 | 8 | 1 | 2 | 1 | 2 | 0 | 1 | 0,10608 | 1 | -0,5612 | -0,2941 | DIPPA_1 | DIPPA_1 | DIPPA_14433.mRNA.1 Ras-related protein Rab-5A |
| 20,3293 | 13,3082 | 16,5061 | 20,1497 | 15,2175 | 16,4853 | 22,5942 | 19,8294 | 22,4406 | 22,724 | 20,053 | 22,4478 | 5 | 5 | 4 | 32,9 | 32,9 | 29,2 | 24,111 | 0 | 18,842 | 3E+07 | 10 | 4 | 0 | 1 | 4 | 0 | 1 | 0,08005 | 1 | -0,5696 | -0,2269 | DIPPA_1 | DIPPA_1 | DIPPA_11797.mRNA.1 Glutathione S-transferase 4 |
| 16,9701 | 12,3293 | 10,7468 | 16,1466 | 13,3312 | 12,2857 | 21,21 | 18,6114 | 20,7062 | 21,7888 | 19,2829 | 18,8469 | 12 | 12 | 12 | 11,9 | 11,9 | 11,9 | 154,88 | 0 | 19,042 | 1,5E+07 | 19 | 8 | 2 | 3 | 7 | 0 | 2 | 0,09308 | 1 | -0,5724 | -0,2608 | DIPPA_0 | DIPPA_0 | DIPPA_05461.mRNA.2 Erythroid differentiation-related factor 1 OS_Mus musculus OX_10090 GN_Edrf1 PE_1 SV_1;DIPPA_05461.mRNA.1 Erythroid differentiation-related factor 1 OS_Mus musculus OX_10090 GN_Edrf1 PE_1 SV_1 |
| 20,1835 | 17,123 | 14,1777 | 20,6239 | 16,0009 | 16,5962 | 23,8082 | 21,8777 | 19,6666 | 24,0897 | 21,6685 | 22,4629 | 15 | 15 | 15 | 50 | 50 | 50 | 44,106 | 0 | 125,45 | 7,2E+07 | 32 | 10 | 6 | 4 | 13 | 4 | 7 | 0,0912 | 1 | -0,5789 | -0,256 | DIPPA_2 | DIPPA_2 | DIPPA_21916.mRNA.1 Obg-like ATPase 1 |
| 19,4521 | 10,9234 | 14,3259 | 19,138 | 13,9827 | 13,3473 | 22,4231 | 19,7901 | 21,2661 | 21,6717 | 19,3766 | 19,9338 | 6 | 6 | 6 | 33,6 | 33,6 | 33,6 | 28,985 | 0 | 13,252 | 2,3E+07 | 11 | 4 | 0 | 2 | 6 | 1 | 2 | 0,0666 | 1 | -0,5889 | -0,191 | DIPPA_1 | DIPPA_1 | DIPPA_18304.mRNA.1 Proteasome subunit alpha type-1 |
| 14,5745 | 15,6 | 14,5226 | 16,8856 | 14,0564 | 15,577 | 20,3676 | 23,3214 | 22,4345 | 22,8026 | 23,289 | 23,1533 | 25 | 25 | 25 | 18,6 | 18,6 | 18,6 | 463,15 | 0 | 136,36 | 4,9E+07 | 63 | 4 | 12 | 13 | 22 | 7 | 16 | 0,27385 | 1 | -0,6073 | -0,6827 | DIPPA_0 | DIPPA_0 | DIPPA_00608.mRNA.1 Endoglucanase;DIPPA_00661.mRNA.1 Endoglucanase |
| 13,7789 | 10,9616 | 10,049 | 14,3724 | 10,4603 | 11,7956 | 19,0382 | 20,0848 | 18,2792 | 19,4652 | 19,2204 | 18,9879 | 3 | 3 | 3 | 5,4 | 5,4 | 5,4 | 109,94 | 0 | 25,069 | 1839300 | 4 | 2 | 1 | 1 | 3 | 0 | 0 | 0,14142 | 1 | -0,6129 | -0,3817 | DIPPA_1 | DIPPA_1 | DIPPA_14032.mRNA.1 RNA cytidine acetyltransferase |
| 14,5119 | 16,557 | 17,2802 | 16,8522 | 16,5673 | 16,7783 | 20,5052 | 22,0674 | 23,1818 | 21,5401 | 22,7942 | 22,4954 | 12 | 12 | 12 | 20,7 | 20,7 | 20,7 | 83,256 | 0 | 24,764 | 2,6E+07 | 36 | 3 | 8 | 10 | 7 | 6 | 10 | 0,30046 | 1 | -0,6162 | -0,7395 | DIPPA_3 | DIPPA_3 | DIPPA_35033.mRNA.1 ATP-dependent RNA helicase DBP2 |
| 12,215 | 13,8512 | 13,3518 | 16,4243 | 11,6762 | 13,2231 | 19,4981 | 20,5217 | 20,1481 | 20,776 | 20,4188 | 20,5922 | 7 | 7 | 7 | 14,2 | 14,2 | 14,2 | 66,534 | 0 | 10,2 | 4576900 | 10 | 1 | 2 | 3 | 7 | 1 | 2 | 0,16128 | 1 | -0,6352 | -0,4293 | DIPPA_2 | DIPPA_2 | DIPPA_23063.mRNA.1 ATP-dependent RNA helicase DED1 |
| 16,8817 | 11,0471 | 12,6766 | 15,929 | 13,9238 | 12,6726 | 20,4012 | 20,5703 | 18,7713 | 20,4567 | 20,48 | 19,4072 | 4 | 4 | 4 | 11,4 | 11,4 | 11,4 | 51,261 | 0 | 30,743 | 4235000 | 7 | 3 | 0 | 1 | 3 | 2 | 1 | 0,11764 | 1 | -0,64 | -0,3232 | DIPPA_1 | DIPPA_1 | DIPPA_12153.mRNA.1 COP9 signalosome complex subunit 2;DIPPA_06992.mRNA.1 COP9 signalosome complex subunit 2 |
| 18,0903 | 15,6239 | 16,1283 | 19,9448 | 14,8068 | 17,0335 | 21,645 | 21,0829 | 22,0528 | 24,1015 | 21,085 | 23,5083 | 14 | 14 | 14 | 21,9 | 21,9 | 21,9 | 80,5 | 0 | 28,448 | 5,2E+07 | 30 | 8 | 2 | 6 | 13 | 4 | 9 | 0,14418 | 1 | -0,6475 | -0,3884 | DIPPA_2 | DIPPA_2 | DIPPA_24346.mRNA.1 Long-chain-fatty-acid--CoA ligase 2 |
| 18,3807 | 18,0619 | 17,6163 | 18,989 | 18,2111 | 18,8656 | 21,5561 | 23,6353 | 23,5234 | 22,6878 | 23,5929 | 24,2481 | 7 | 7 | 7 | 21,6 | 21,6 | 21,6 | 47,134 | 0 | 16,214 | 4E+07 | 18 | 5 | 3 | 5 | 5 | 5 | 7 | 0,95556 | 1 | -0,669 | -2,0413 | DIPPA_1 | DIPPA_1 | DIPPA_15671.mRNA.1 Probable leucine aminopeptidase 1 |
| 18,339 | 14,5262 | 15,3758 | 19,3624 | 15,2697 | 15,7725 | 20,8412 | 21,3534 | 22,3419 | 24,1186 | 22,7124 | 22,5145 | 25 | 25 | 24 | 23,1 | 23,1 | 22,3 | 125,93 | 0 | 113,5 | 7,2E+07 | 47 | 10 | 3 | 6 | 24 | 5 | 9 | 0,15594 | 1 | -0,7212 | -0,4166 | TR25560 | TR25560 | TR25560__c0_g1_i1__m.3032 #MITO 0 #PFAM 2 x PF00225.18 Kinesin Kinesin motor domain;2 x PF11351.3 DUF3154 Protein of unknown function (DUF3154) #BLASTP_UNIREF90 6e-91 Kinesin-like protein n=1 Tax=Bodo saltans RepID=A0A0S4KM02_BODSA (length=1017);DIPPA_18 |
| 19,8258 | 10,8969 | 14,4203 | 18,8249 | 14,624 | 13,8922 | 22,5341 | 20,1367 | 19,5964 | 21,8567 | 18,699 | 19,3409 | 5 | 5 | 5 | 34,1 | 34,1 | 34,1 | 29,004 | 0 | 26,389 | 1,9E+07 | 11 | 5 | 0 | 1 | 4 | 1 | 1 | 0,08614 | 1 | -0,7327 | -0,2428 | DIPPA_3 | DIPPA_3 | DIPPA_34022.mRNA.1 hypothetical protein Tco025E_05491 |
| 18,6296 | 13,8882 | 16,1734 | 19,6991 | 14,6763 | 16,5192 | 22,2455 | 20,0621 | 22,7597 | 22,8478 | 22,9363 | 23,3589 | 8 | 8 | 8 | 24 | 24 | 24 | 51,811 | 0 | 14,04 | 3,5E+07 | 20 | 6 | 2 | 4 | 7 | 2 | 5 | 0,13498 | 1 | -0,7345 | -0,3661 | DIPPA_1 | DIPPA_1 | DIPPA_13153.mRNA.1 Probable acyl-CoA dehydrogenase IBR3 |
| 13,3185 | 10,7326 | 12,3083 | 16,4843 | 11,2902 | 10,8816 | 18,0172 | 20,0207 | 20,0204 | 21,3627 | 19,3372 | 20,3602 | 6 | 6 | 6 | 10,7 | 10,7 | 10,7 | 114,54 | 0 | 16,137 | 6204500 | 6 | 1 | 0 | 2 | 6 | 0 | 1 | 0,14558 | 1 | -0,7656 | -0,3918 | DIPPA_2 | DIPPA_2 | DIPPA_21434.mRNA.1 Ubiquitin-like modifier-activating enzyme 1 |
| 18,3381 | 13,3351 | 11,1117 | 17,9472 | 12,7307 | 14,4043 | 18,5634 | 20,4524 | 18,7384 | 20,3725 | 20,6184 | 20,1401 | 10 | 10 | 10 | 33,3 | 33,3 | 33,3 | 59,287 | 0 | 21,545 | 1,8E+07 | 20 | 6 | 2 | 1 | 7 | 2 | 4 | 0,1048 | 1 | -0,7658 | -0,2909 | DIPPA_1 | DIPPA_1 | DIPPA_13676.mRNA.1 hypothetical protein |
| 19,7788 | 16,64 | 16,7821 | 20,5987 | 16,8239 | 18,3089 | 22,6359 | 21,802 | 22,4961 | 24,3388 | 23,0014 | 24,2399 | 31 | 31 | 31 | 47,2 | 47,2 | 47,2 | 100,43 | 0 | 171,05 | 1,6E+08 | 78 | 19 | 4 | 8 | 29 | 6 | 16 | 0,21887 | 1 | -0,8435 | -0,562 | DIPPA_3 | DIPPA_3 | DIPPA_33974.mRNA.1 Major vault protein;DIPPA_33977.mRNA.1 Major vault protein |
| 20,0204 | 13,1672 | 15,1797 | 20,0798 | 14,1525 | 16,6675 | 22,6927 | 18,7409 | 20,9176 | 23,6826 | 20,4873 | 22,2591 | 20 | 20 | 20 | 34,7 | 34,7 | 34,7 | 81,839 | 0 | 92,06 | 9,1E+07 | 45 | 17 | 2 | 4 | 19 | 3 | 8 | 0,1152 | 1 | -0,8442 | -0,3171 | DIPPA_1 | DIPPA_1 | DIPPA_15225.mRNA.1 Elongation factor G mitochondrial |
| 21,5711 | 11,6621 | 16,5399 | 20,4542 | 15,1091 | 16,773 | 22,4364 | 20,142 | 21,3598 | 23,1751 | 21,4375 | 21,913 | 9 | 9 | 9 | 62,3 | 62,3 | 62,3 | 23,08 | 0 | 87,222 | 5,3E+07 | 24 | 7 | 1 | 4 | 7 | 2 | 5 | 0,09333 | 1 | -0,8544 | -0,2615 | DIPPA_2 | DIPPA_2 | DIPPA_22485.mRNA.1 hypothetical protein |
| 16,9231 | 16,1015 | 13,7559 | 17,5708 | 15,5908 | 16,1949 | 20,5731 | 19,3151 | 19,5424 | 21,6038 | 19,8628 | 22,1474 | 8 | 8 | 8 | 25,4 | 25,4 | 25,4 | 52,332 | 0 | 43,314 | 1,2E+07 | 15 | 5 | 1 | 1 | 7 | 1 | 4 | 0,31491 | 1 | -0,8587 | -0,77 | DIPPA_1 | DIPPA_1 | DIPPA_18510.mRNA.1 Sorting nexin 2A |
| 15,8579 | 11,5688 | 13,5764 | 18,0468 | 11,3926 | 14,1569 | 20,3651 | 19,5244 | 20,0782 | 22,6039 | 19,7092 | 20,8215 | 10 | 10 | 10 | 15,5 | 15,5 | 15,5 | 108,5 | 0 | 26,111 | 1,5E+07 | 12 | 3 | 0 | 2 | 10 | 1 | 4 | 0,13942 | 1 | -0,8644 | -0,3769 | DIPPA_1 | DIPPA_1 | DIPPA_19759.mRNA.1 Calcium-transporting ATPase sarcoplasmic/endoplasmic reticulum type |
| 17,7199 | 12,7987 | 15,5732 | 18,0183 | 15,0498 | 15,6719 | 21,1973 | 19,5566 | 21,8073 | 22,116 | 22,4199 | 22,3029 | 14 | 14 | 14 | 21,4 | 21,4 | 21,4 | 92,683 | 0 | 96,536 | 3,2E+07 | 32 | 12 | 1 | 6 | 10 | 5 | 8 | 0,20174 | 1 | -0,8828 | -0,5233 | DIPPA_1 | DIPPA_1 | DIPPA_13373.mRNA.2 hypothetical protein;DIPPA_13373.mRNA.1 hypothetical protein |
| 14,6495 | 11,7861 | 13,0261 | 17,0873 | 10,6991 | 14,4284 | 19,2727 | 18,8858 | 19,1332 | 21,778 | 18,8558 | 21,8014 | 14 | 14 | 14 | 13,1 | 13,1 | 13,1 | 145,67 | 0 | 29,009 | 1,2E+07 | 20 | 5 | 0 | 3 | 12 | 2 | 6 | 0,17095 | 1 | -0,9177 | -0,4521 | DIPPA_6 | DIPPA_6 | DIPPA_65228.mRNA.1 Ubiquitin carrier protein n=1 Tax=Acanthamoeba castellanii str. Neff RepID=L8HHE5_ACACA (length=396) |
| 16,1546 | 16,5192 | 16,5865 | 19,7129 | 15,9904 | 16,3502 | 20,9987 | 22,6229 | 22,7732 | 24,3044 | 22,6934 | 22,5467 | 10 | 10 | 10 | 19,8 | 19,8 | 19,8 | 67,245 | 0 | 19,551 | 3,7E+07 | 18 | 3 | 2 | 3 | 9 | 3 | 5 | 0,31993 | 1 | -0,9311 | -0,7805 | DIPPA_1 | DIPPA_1 | DIPPA_19967.mRNA.1 hypothetical protein |
| 14,0829 | 17,9639 | 17,1819 | 18,5097 | 17,0252 | 16,5161 | 19,8314 | 23,4928 | 22,8895 | 19,7356 | 23,1199 | 22,5401 | 6 | 2 | 2 | 25,8 | 13 | 13 | 40,811 | 0 | 13,801 | 1,3E+07 | 15 | 0 | 2 | 2 | 1 | 2 | 2 | 0,286 | 1 | -0,9408 | -0,7087 | DIPPA_2 | DIPPA_2 | DIPPA_29150.mRNA.1 Proteinase R |
| 15,6799 | 13,7295 | 12,6153 | 18,9439 | 12,2607 | 13,6457 | 18,1663 | 18,0459 | 18,1887 | 24,1939 | 18,2182 | 18,3883 | 19 | 19 | 12 | 17,6 | 17,6 | 12 | 153,53 | 0 | 130,11 | 4E+07 | 26 | 4 | 3 | 2 | 12 | 1 | 3 | 0,15877 | 1 | -0,9419 | -0,4234 | DIPPA_3 | DIPPA_3 | DIPPA_32783.mRNA.1 Multidrug resistance protein 1;DIPPA_20659.mRNA.1 Multidrug resistance protein 1;DIPPA_11051.mRNA.1 Multidrug resistance protein 1 |
| 15,2219 | 13,4418 | 11,0219 | 16,4931 | 12,9568 | 13,1009 | 19,6032 | 19,2997 | 20,1513 | 21,1997 | 19,3818 | 20,0661 | 4 | 4 | 4 | 11,2 | 11,2 | 11,2 | 62,658 | 0 | 22,432 | 4274600 | 8 | 3 | 1 | 0 | 4 | 1 | 2 | 0,22205 | 1 | -0,9551 | -0,5691 | DIPPA_0 | DIPPA_0 | DIPPA_02908.mRNA.1 Minor serine/threonine-protein phosphatase PP2A-1 catalytic subunit |
| 22,5298 | 22,9279 | 21,7485 | 23,1067 | 22,9678 | 23,9972 | 23,4634 | 26,4252 | 25,6824 | 24,7231 | 26,7712 | 27,5114 | 15 | 15 | 15 | 47,9 | 47,9 | 47,9 | 21,914 | 0 | 107,28 | 4,6E+08 | 72 | 12 | 12 | 11 | 12 | 13 | 15 | 0,94414 | 1 | -0,9552 | -2,0182 | DIPPA_3 | DIPPA_3 | DIPPA_30776.mRNA.1 Calcium-binding protein NCSA |
| 18,3397 | 17,9216 | 17,3188 | 20,7731 | 17,5535 | 18,256 | 21,7139 | 22,9228 | 23,0486 | 23,8672 | 23,2915 | 23,8114 | 22 | 22 | 21 | 46,9 | 46,9 | 45,4 | 62,141 | 0 | 61,599 | 8,8E+07 | 60 | 8 | 12 | 12 | 19 | 10 | 17 | 0,41726 | 1 | -1,0008 | -0,9799 | DIPPA_2 | DIPPA_2 | DIPPA_28298.mRNA.1 Kinesin-3 |
| 10,8416 | 12,7977 | 13,3849 | 14,8218 | 12,2789 | 12,9668 | 19,0652 | 21,0054 | 20,8966 | 21,7098 | 20,7397 | 20,9933 | 12 | 12 | 12 | 7,1 | 7,1 | 7,1 | 245,95 | 0 | 30,926 | 7205700 | 19 | 2 | 2 | 4 | 10 | 3 | 5 | 0,39684 | 1 | -1,0144 | -0,9387 | TR35483 | TR35483 | TR35483__c0_g1_i1__m.4373 #MITO 1 HMMTOP #PFAM 2 x PF02889.11 Sec63 Sec63 Brl domain;5 x PF00270.24 DEAD DEAD/DEAH box helicase;2 x PF04851.10 ResIII Type III restriction enzyme, res subunit;4 x PF13401.1 AAA_22 AAA domain;2 x PF00271.26 Helicase_C |
| 12,8371 | 12,0086 | 12,3869 | 17,1411 | 11,6309 | 11,5651 | 18,6478 | 20,2702 | 19,4811 | 20,5234 | 20,0039 | 18,9717 | 7 | 7 | 7 | 15,6 | 15,6 | 15,6 | 104,18 | 0 | 14,155 | 6162700 | 7 | 1 | 1 | 0 | 6 | 1 | 2 | 0,21594 | 1 | -1,0348 | -0,5554 | DIPPA_1 | DIPPA_1 | DIPPA_18552.mRNA.1 hypothetical protein |
| 25,9406 | 12,6043 | 16,2767 | 24,1831 | 16,8587 | 17,0406 | 27,211 | 19,2372 | 21,3871 | 25,0062 | 19,7327 | 21,4661 | 5 | 5 | 5 | 53,8 | 53,8 | 53,8 | 13,26 | 0 | 31,662 | 4,2E+08 | 19 | 4 | 0 | 3 | 5 | 1 | 4 | 0,08265 | 1 | -1,0869 | -0,2337 | DIPPA_3 | DIPPA_3 | DIPPA_33223.mRNA.1 hypothetical protein conserved |
| 17,0169 | 14,3521 | 13,4467 | 17,0828 | 15,2293 | 15,8907 | 21,9989 | 19,8551 | 20,3351 | 21,9512 | 22,6123 | 22,657 | 6 | 6 | 2 | 11,6 | 11,6 | 4,7 | 90,53 | 0 | 19,926 | 1,4E+07 | 7 | 0 | 0 | 0 | 2 | 0 | 0 | 0,39751 | 1 | -1,129 | -0,9401 | TR64081 | TR64081 | TR64081__c1_g1_i1__m.10542 #MITO 1 HMMTOP #PFAM PF13229.1 Beta_helix Right handed beta helix region #BLASTP_UNIREF90 1e-168 Putative uncharacterized protein n=1 Tax=Salpingoeca rosetta (strain ATCC 50818 / BSB-021) RepID=F2UA60_SALR5 (length=948);5e-10 P |
| 6,83643 | 7,46475 | 12,7221 | 12,5734 | 10,7487 | 7,17073 | 19,883 | 18,7218 | 19,1032 | 19,8521 | 19,8602 | 19,8171 | 10 | 10 | 5 | 1,7 | 1,7 | 0,9 | 1061,5 | 0 | 64,814 | 2540200 | 10 | 0 | 0 | 0 | 5 | 0 | 0 | 0,17952 | 1 | -1,1565 | -0,4722 | DIPPA_3 | DIPPA_3 | DIPPA_34706.mRNA.1 hypothetical protein;DIPPA_26125.mRNA.1 hypothetical protein |
| 17,4289 | 16,3067 | 16,4427 | 19,5573 | 17,6448 | 16,457 | 21,8954 | 22,4011 | 23,2738 | 24,2066 | 23,2978 | 23,2161 | 11 | 11 | 11 | 24,9 | 24,9 | 24,9 | 65,85 | 0 | 17,768 | 4E+07 | 22 | 6 | 4 | 7 | 9 | 3 | 7 | 0,52631 | 1 | -1,1602 | -1,1963 | DIPPA_1 | DIPPA_1 | DIPPA_13683.mRNA.1 Serine/threonine-protein phosphatase BSL3 |
| 14,1666 | 11,6683 | 14,6821 | 16,6331 | 13,6338 | 13,7347 | 19,0648 | 18,936 | 18,9183 | 19,1658 | 19,557 | 19,7597 | 3 | 3 | 3 | 21 | 21 | 21 | 26,402 | 0 | 10,289 | 2417200 | 3 | 1 | 0 | 1 | 1 | 1 | 1 | 0,35723 | 1 | -1,1616 | -0,8579 | DIPPA_1 | DIPPA_1 | DIPPA_16065.mRNA.1 Thioredoxin domain-containing protein PLP3B |
| 17,3496 | 13,503 | 13,1787 | 18,2442 | 14,4114 | 15,1046 | 20,0439 | 19,0829 | 18,6115 | 21,841 | 21,2047 | 20,7096 | 6 | 6 | 6 | 18,8 | 18,8 | 18,8 | 49,4 | 0 | 79,262 | 1,2E+07 | 14 | 4 | 1 | 2 | 6 | 3 | 4 | 0,28029 | 1 | -1,243 | -0,6965 | DIPPA_1 | DIPPA_1 | DIPPA_18028.mRNA.1 Protein ABHD14A OS_Mus musculus GN_Abhd14a PE_1 SV_2 |
| 20,8973 | 12,4408 | 15,4 | 19,117 | 16,035 | 17,3526 | 23,1772 | 20,6934 | 21,8634 | 22,9734 | 22,3908 | 22,92 | 11 | 11 | 11 | 32,7 | 32,7 | 32,7 | 45,554 | 0 | 32,043 | 5E+07 | 24 | 10 | 2 | 4 | 7 | 2 | 4 | 0,1815 | 1 | -1,2555 | -0,4767 | DIPPA_2 | DIPPA_2 | DIPPA_24329.mRNA.1 Tyrosine aminotransferase |
| 22,095 | 11,0827 | 16,4057 | 20,9595 | 15,2211 | 17,2261 | 24,1168 | 19,8601 | 21,5058 | 24,3177 | 20,9763 | 21,7473 | 9 | 9 | 7 | 62,3 | 62,3 | 49,3 | 24,138 | 0 | 21,604 | 8,2E+07 | 19 | 7 | 1 | 2 | 6 | 2 | 3 | 0,13019 | 1 | -1,2744 | -0,3543 | DIPPA_1 | DIPPA_1 | DIPPA_19873.mRNA.1 Glutathione s-transferase kappa 2 |
| 18,9306 | 18,7804 | 19,0718 | 21,4347 | 19,1473 | 20,0336 | 22,1242 | 23,9977 | 23,838 | 23,1389 | 23,9131 | 24,5782 | 7 | 7 | 2 | 52,2 | 52,2 | 20,7 | 20,631 | 0 | 20,092 | 6E+07 | 23 | 0 | 0 | 1 | 2 | 1 | 1 | 0,88704 | 1 | -1,2776 | -1,9036 | DIPPA_3 | DIPPA_3 | DIPPA_34939.mRNA.1 ADP-ribosylation factor 1 |
| 14,7652 | 13,277 | 13,0976 | 17,3114 | 13,6317 | 14,0529 | 19,3488 | 19,9385 | 19,7654 | 22,1512 | 19,8514 | 20,1174 | 8 | 8 | 2 | 16,3 | 16,3 | 5 | 75,212 | 0 | 207,08 | 1E+07 | 12 | 0 | 0 | 2 | 2 | 0 | 0 | 0,43044 | 1 | -1,2854 | -1,0064 | TR85819 | TR85819 | TR85819__c2_g2_i2__m.17553 #MITO 3 Predotar,TargetP,Mitoprot #PFAM - #BLASTP_UNIREF90 4e-12 Uncharacterized protein n=2 Tax=Rhizophagus irregularis RepID=A0A015KF33_9GLOM (length=652) |
| 15,427 | 12,2039 | 12,6639 | 15,4948 | 13,4409 | 15,2635 | 20,4439 | 18,838 | 21,0318 | 21,1666 | 22,0807 | 22,6058 | 10 | 8 | 8 | 7,7 | 5,7 | 5,7 | 237,05 | 0 | 10,45 | 9058100 | 12 | 0 | 1 | 2 | 5 | 4 | 8 | 0,47061 | 1 | -1,3015 | -1,0865 | DIPPA_1 | DIPPA_1 | DIPPA_19686.mRNA.1 hypothetical protein |
| 17,5175 | 10,6102 | 13,8591 | 17,6111 | 14,0328 | 14,2482 | 20,0513 | 19,669 | 19,5377 | 20,7277 | 20,6001 | 20,7079 | 7 | 7 | 7 | 24,1 | 24,1 | 24,1 | 50,573 | 0 | 58,189 | 7900200 | 13 | 5 | 0 | 2 | 6 | 2 | 2 | 0,21988 | 1 | -1,3018 | -0,5642 | DIPPA_0 | DIPPA_0 | DIPPA_00219.mRNA.1 CCR4-NOT transcription complex subunit 11 OS_Homo sapiens GN_CNOT11 PE_1 SV_1 |
| 16,8726 | 10,4747 | 13,8477 | 18,2268 | 14,2869 | 13,001 | 20,3891 | 19,1894 | 18,8864 | 22,1577 | 19,2567 | 19,6931 | 8 | 8 | 8 | 24,1 | 24,1 | 24,1 | 55,301 | 0 | 21,262 | 1,4E+07 | 9 | 4 | 0 | 2 | 7 | 1 | 4 | 0,23302 | 1 | -1,4399 | -0,5935 | DIPPA_1 | DIPPA_1 | DIPPA_14281.mRNA.1 hypothetical protein conserved |
| 17,8962 | 11,9478 | 15,107 | 17,3719 | 16,9676 | 14,9437 | 21,5856 | 20,3533 | 23,0704 | 21,5856 | 24,0741 | 23,0987 | 8 | 8 | 8 | 16,1 | 16,1 | 16,1 | 69,946 | 0 | 15,096 | 2,2E+07 | 15 | 8 | 1 | 3 | 6 | 3 | 4 | 0,31496 | 1 | -1,4441 | -0,7701 | DIPPA_0 | DIPPA_0 | DIPPA_04398.mRNA.2 Alpha-actinin-like protein 1;DIPPA_04398.mRNA.1 Alpha-actinin-like protein 1 |
| 19,9184 | 12,7999 | 15,0203 | 19,1139 | 15,8323 | 17,2059 | 21,7704 | 19,6618 | 18,3885 | 21,6929 | 20,023 | 23,2036 | 8 | 8 | 8 | 47,7 | 47,7 | 47,7 | 33,129 | 0 | 18,901 | 2,3E+07 | 15 | 6 | 0 | 2 | 7 | 1 | 4 | 0,25296 | 1 | -1,4712 | -0,6374 | DIPPA_3 | DIPPA_3 | DIPPA_34172.mRNA.1 Thiamine thiazole synthase 2 |
| 11,4269 | 15,4373 | 14,616 | 15,9536 | 14,3061 | 15,639 | 19,0146 | 22,3518 | 21,5764 | 21,0354 | 22,5011 | 22,1051 | 11 | 11 | 11 | 13,2 | 13,2 | 13,2 | 120,85 | 0 | 14,636 | 1,3E+07 | 19 | 1 | 3 | 5 | 8 | 3 | 8 | 0,48402 | 1 | -1,4728 | -1,113 | DIPPA_1 | DIPPA_1 | DIPPA_14412.mRNA.1 Kinesin-like protein unc-104 |
| 17,421 | 13,9554 | 15,3004 | 21,3548 | 14,6295 | 15,38 | 19,7235 | 18,2949 | 19,2366 | 24,9874 | 19,4991 | 19,4112 | 14 | 14 | 14 | 30,7 | 30,7 | 30,7 | 62,949 | 0 | 47,872 | 9,2E+07 | 23 | 6 | 2 | 3 | 14 | 1 | 3 | 0,26499 | 1 | -1,5625 | -0,6636 | DIPPA_1 | DIPPA_1 | DIPPA_19232.mRNA.1 hypothetical protein DQ04_00191280 |
| 18,7284 | 13,835 | 13,7322 | 18,3305 | 15,5159 | 17,2377 | 23,3056 | 20,2227 | 20,3599 | 23,2745 | 23,3152 | 24,2684 | 68 | 68 | 68 | 17 | 17 | 17 | 664,83 | 0 | 323,31 | 2,5E+08 | 136 | 50 | 11 | 14 | 59 | 24 | 49 | 0,3617 | 1 | -1,5962 | -0,8671 | TR67648 | TR67648 | TR67648__c0_g1_i2__m.11458 #MITO 0 #PFAM 22 x PF08309.6 LVIVD LVIVD repeat;8 x PF09826.4 Beta_propel Beta propeller domain;7 x PF01197.13 Ribosomal_L31 Ribosomal protein L31;2 x PF09992.4 DUF2233 Predicted periplasmic protein (DUF2233);PF05345.7 He |
| 19,65 | 11,9803 | 15,8833 | 20,0463 | 15,8811 | 16,718 | 21,826 | 18,9613 | 21,5132 | 22,6661 | 19,9854 | 21,5964 | 6 | 6 | 6 | 42 | 42 | 42 | 21,565 | 0 | 11,481 | 2,8E+07 | 11 | 5 | 0 | 2 | 6 | 1 | 4 | 0,26792 | 1 | -1,7106 | -0,6699 | DIPPA_1 | DIPPA_1 | DIPPA_14995.mRNA.2 hypothetical protein;DIPPA_14995.mRNA.1 hypothetical protein |
| 18,3716 | 11,4382 | 17,0378 | 19,3504 | 15,9845 | 16,793 | 18,9523 | 18,684 | 18,7538 | 19,741 | 19,7733 | 19,4685 | 2 | 2 | 2 | 19,4 | 19,4 | 19,4 | 18,701 | 0 | 74,885 | 5281900 | 7 | 1 | 1 | 2 | 2 | 2 | 2 | 0,30437 | 1 | -1,7601 | -0,7478 | DIPPA_1 | DIPPA_1 | DIPPA_11759.mRNA.2 hypothetical protein;DIPPA_11759.mRNA.1 conserved Plasmodium protein unknown function;DIPPA_26747.mRNA.1 conserved Plasmodium protein unknown function;DIPPA_15071.mRNA.1 conserved Plasmodium protein unknown function |
| 20,2566 | 16,2911 | 11,5411 | 20,7956 | 15,5357 | 17,0496 | 21,9588 | 19,2298 | 20,325 | 23,0901 | 19,0978 | 19,495 | 5 | 5 | 2 | 60 | 60 | 29,5 | 10,753 | 0 | 64,139 | 2E+07 | 11 | 1 | 1 | 0 | 2 | 1 | 1 | 0,23366 | 1 | -1,764 | -0,595 | DIPPA_1 | DIPPA_1 | DIPPA_18299.mRNA.1 hypothetical protein |
| 11,1016 | 11,4357 | 10,3241 | 15,0894 | 10,8756 | 12,2445 | 19,3111 | 20,3275 | 19,1575 | 19,595 | 18,9639 | 18,7694 | 4 | 4 | 4 | 4,6 | 4,6 | 4,6 | 165,08 | 0 | 14,967 | 3105700 | 5 | 2 | 0 | 1 | 4 | 0 | 0 | 0,62466 | 1 | -1,7827 | -1,3884 | DIPPA_3 | DIPPA_3 | DIPPA_33928.mRNA.1 Clustered mitochondria protein homolog;TR122417__c5_g1_i1__m.29049 #MITO 0 #PFAM 4 x PF13424.1 TPR_12 Tetratricopeptide repeat;PF13236.1 CLU Clustered mitochondria;4 x PF13414.1 TPR_11 TPR repeat;6 x PF07719.12 TPR_2 Tetratricopept |
| 14,1906 | 18,3654 | 17,5497 | 18,3233 | 18,4996 | 18,7648 | 19,5379 | 23,4942 | 23,2849 | 22,265 | 24,0384 | 24,2699 | 7 | 7 | 7 | 18,5 | 18,5 | 18,5 | 59,514 | 0 | 26,615 | 3,2E+07 | 18 | 0 | 4 | 3 | 5 | 5 | 6 | 0,6425 | 1 | -1,8273 | -1,4232 | DIPPA_0 | DIPPA_0 | DIPPA_04587.mRNA.1 Tripeptidyl-peptidase sed2;TR125201__c3_g3_i1__m.29678 #MITO 0 #PFAM PF09286.6 Pro-kuma_activ Pro-kumamolisin, activation domain;PF00082.17 Peptidase_S8 Subtilase family #BLASTP_UNIREF90 3e-85 Predicted protein n=1 Tax=Nematostella vec |
| 15,3116 | 19,3869 | 18,4571 | 20,3026 | 19,0885 | 19,3403 | 19,6387 | 23,183 | 22,641 | 22,4063 | 23,7534 | 23,5042 | 7 | 7 | 7 | 29,5 | 29,5 | 29,5 | 33,471 | 0 | 64,136 | 4,3E+07 | 33 | 1 | 5 | 7 | 6 | 6 | 7 | 0,65307 | 1 | -1,8586 | -1,4438 | DIPPA_3 | DIPPA_3 | DIPPA_35556.mRNA.2 hypothetical protein;DIPPA_08165.mRNA.1 hypothetical protein;DIPPA_08164.mRNA.2 hypothetical protein;DIPPA_08156.mRNA.1 hypothetical protein;DIPPA_35556.mRNA.1 hypothetical protein;DIPPA_08164.mRNA.1 hypothetical protein |
| 18,1539 | 12,6077 | 9,33404 | 17,9072 | 14,2586 | 14,0719 | 20,2389 | 19,2317 | 19,2286 | 21,0487 | 19,636 | 21,0626 | 2 | 2 | 2 | 7,2 | 7,2 | 7,2 | 34,692 | 0 | 27,855 | 6966300 | 7 | 2 | 1 | 0 | 2 | 1 | 1 | 0,28923 | 1 | -2,0473 | -0,7156 | DIPPA_1 | DIPPA_1 | DIPPA_18643.mRNA.1 merozoite adhesive erythrocytic binding protein;DIPPA_18643.mRNA.2 merozoite adhesive erythrocytic binding protein;DIPPA_18643.mRNA.3 merozoite adhesive erythrocytic binding protein |
| 20,5056 | 12,5669 | 14,0533 | 21,1885 | 15,2505 | 17,0214 | 21,6353 | 20,1888 | 19,5777 | 22,7655 | 19,6845 | 21,5593 | 4 | 4 | 4 | 49,1 | 49,1 | 49,1 | 13,109 | 0 | 11,021 | 2,4E+07 | 10 | 3 | 0 | 1 | 4 | 1 | 3 | 0,28308 | 1 | -2,1115 | -0,7025 | DIPPA_2 | DIPPA_2 | DIPPA_20803.mRNA.1 hypothetical protein |
| 19,1292 | 16,8925 | 16,6691 | 20,1426 | 19,3613 | 19,5993 | 20,5442 | 20,3166 | 21,5499 | 22,4195 | 24,5788 | 24,2345 | 3 | 3 | 3 | 25,8 | 25,8 | 25,8 | 18,178 | 0 | 16,338 | 2,7E+07 | 15 | 3 | 1 | 2 | 3 | 2 | 3 | 1,22631 | 1 | -2,1374 | -2,6105 | DIPPA_0 | DIPPA_0 | DIPPA_06543.mRNA.1 N-acetyltransferase ats1 |
| 16,1337 | 13,3607 | 14,4596 | 17,0193 | 16,4406 | 17,0628 | 21,2382 | 20,0282 | 22,2979 | 22,6385 | 24,3483 | 24,2802 | 18 | 18 | 18 | 27,7 | 27,7 | 27,7 | 396,54 | 0 | 67,414 | 4,5E+07 | 33 | 6 | 2 | 5 | 13 | 6 | 14 | 1,23771 | 1 | -2,1896 | -2,6356 | DIPPA_1 | DIPPA_1 | DIPPA_17995.mRNA.1 Pectate trisaccharide-lyase |
| 15,3268 | 17,6902 | 17,7932 | 20,8729 | 18,8402 | 18,3421 | 18,6902 | 22,834 | 23,1419 | 23,6988 | 24,4975 | 23,9463 | 20 | 16 | 16 | 40,4 | 34,3 | 34,3 | 79,016 | 0 | 104,24 | 8,5E+07 | 44 | 0 | 7 | 11 | 15 | 11 | 12 | 1,01457 | 1 | -2,415 | -2,1617 | DIPPA_0 | DIPPA_0 | DIPPA_00024.mRNA.1 Xyloglucanase Xgh74A |
| 20,4113 | 13,2073 | 15,201 | 20,9491 | 17,6451 | 17,7514 | 18,4284 | 20,0375 | 19,59 | 18,6998 | 19,3443 | 19,7749 | 5 | 5 | 5 | 42,7 | 42,7 | 42,7 | 21,296 | 0 | 11,022 | 3,1E+07 | 11 | 2 | 1 | 1 | 3 | 1 | 3 | 0,44865 | 1 | -2,5086 | -1,0428 | DIPPA_0 | DIPPA_0 | DIPPA_09620.mRNA.1 Vacuolar protein sorting-associated protein 29 |
| 17,1222 | 11,6181 | 11,2312 | 19,3613 | 13,3137 | 15,3834 | 20,9177 | 19,2644 | 18,5785 | 23,5311 | 19,8459 | 21,42 | 20 | 20 | 19 | 29,8 | 29,8 | 28,5 | 94,663 | 0 | 130,12 | 4,4E+07 | 29 | 7 | 0 | 1 | 19 | 3 | 9 | 0,44532 | 1 | -2,6956 | -1,0361 | DIPPA_1 | DIPPA_1 | DIPPA_16419.mRNA.1 Major vault protein |
| 11,3635 | 12,6423 | 11,0262 | 16,6811 | 12,9446 | 14,3849 | 18,9637 | 17,7536 | 19,193 | 22,5575 | 21,155 | 22,0906 | 20 | 20 | 20 | 8,1 | 8,1 | 8,1 | 415,87 | 0 | 87,609 | 2,2E+07 | 29 | 2 | 2 | 2 | 17 | 3 | 10 | 1,17829 | 1 | -2,9928 | -2,5061 | DIPPA_2 | DIPPA_2 | DIPPA_26123.mRNA.1 hypothetical protein |
| 14,2867 | 12,0111 | 11,4531 | 18,0574 | 13,871 | 15,3319 | 19,7257 | 18,28 | 19,4814 | 21,5359 | 21,4764 | 21,1411 | 5 | 5 | 5 | 10 | 10 | 10 | 79,163 | 0 | 40,008 | 6332500 | 6 | 1 | 0 | 1 | 4 | 2 | 3 | 0,98952 | 1 | -3,1698 | -2,1104 | TR99499 | TR99499 | TR99499__c0_g1_i2__m.21727 #MITO 0 #PFAM PF12849.2 PBP_like_2 PBP superfamily domain;5 x PF13415.1 Kelch_3 Galactose oxidase, central domain;3 x PF13418.1 Kelch_4 Galactose oxidase, central domain;4 x PF01344.20 Kelch_1 Kelch motif;4 x PF13964.1 Ke |
| 12,7315 | 19,6593 | 18,8351 | 19,896 | 20,7473 | 20,7773 | 19,2395 | 24,9577 | 24,63 | 24,2397 | 26,9203 | 26,8287 | 27 | 27 | 19 | 62,2 | 62,2 | 43,9 | 172,53 | 0 | 223,44 | 4,3E+08 | 116 | 0 | 11 | 12 | 15 | 17 | 18 | 0,70344 | 1 | -3,3983 | -1,5419 | DIPPA_2 | DIPPA_2 | DIPPA_27297.mRNA.1 Xyloglucanase Xgh74A;DIPPA_10258.mRNA.1 Xyloglucanase Xgh74A |
| 13,9589 | 11,3935 | 10,5118 | 19,9066 | 14,4259 | 15,6595 | 18,4646 | 19,7184 | 19,0882 | 24,8487 | 21,3887 | 22,2549 | 16 | 16 | 16 | 18,3 | 18,3 | 18,3 | 126,32 | 0 | 49,495 | 5,7E+07 | 20 | 1 | 0 | 1 | 16 | 2 | 5 | 1,13249 | 1 | -4,7093 | -2,4081 | DIPPA_2 | DIPPA_2 | DIPPA_26227.mRNA.1 Ubiquitin-conjugating enzyme E2 10 |

| iBAQ Sec | iBAQ Sec | iBAQ Sec | iBAQ wt- | iBAQ wt- | iBAQ wt- | LFQ inten | LFQ inten | LFQ inten | LFQ inten | LFQ inten | LFQ inten | Student's | Student's | Peptides | Razor + u | Unique p | Sequence | Unique + | Unique se | Mol. weig | Q-value | Score | Intensity | MS/MS c | Unique p | Unique p | Unique p | Unique p | Unique p | Unique p | -Log Stud | Student's | Student's | Student's | Protein ID | Majority | Fasta headers |
|---|---|---|---|---|---|---|---|---|---|---|---|---|---|---|---|---|---|---|---|---|---|---|---|---|---|---|---|---|---|---|---|---|---|---|---|---|---|
| 26,6517 | 23,1134 | 24,4482 | 15,9998 | 11,1551 | 10,5225 | 28,6715 | 25,7404 | 28,4554 | 18,8777 | 18,9596 | 19,0326 | | | 7 | 7 | 7 | 35,5 | 35,5 | 35,5 | 35,549 | 0 | 84,579 | 1,8E+09 | 30 | 7 | 7 | 7 | 1 | 0 | 0 | 2,42312 | 0,22 | 12,1786 | 6,04627 | DIPPA_24 | DIPPA_24 | DIPPA_24951.mRNA.1 Protein transport protein sec13 |
| 25,375 | 20,0358 | 20,5471 | 15,6674 | 12,7183 | 13,9332 | 27,5617 | 19,4688 | 22,2436 | 16,9909 | 18,1147 | 17,1904 | | | 27 | 27 | 27 | 35,2 | 35,2 | 35,2 | 130,25 | 0 | 323,31 | 1,8E+09 | 74 | 27 | 19 | 16 | 2 | 1 | 2 | 1,84175 | 0,572 | 7,87964 | 4,1383 | DIPPA_0 | DIPPA_0 | DIPPA_09941.mRNA.1 Protein transport protein SEC31 |
| 22,3515 | 17,7038 | 18,9901 | 13,3925 | 10,9619 | 12,5367 | 24,6039 | 21,0245 | 22,4891 | 19,4115 | 18,8969 | 17,6569 | | | 12 | 12 | 12 | 50,8 | 50,8 | 50,8 | 40,36 | 0 | 101,16 | 1E+08 | 17 | 12 | 4 | 4 | 0 | 0 | 0 | 2,04424 | 0,46 | 7,3848 | 4,74084 | DIPPA_3 | DIPPA_3 | DIPPA_30206.mRNA.1 Nucleoporin SEH1 |
| 22,5578 | 21,798 | 23,12 | 24,7361 | 11,1551 | 10,9013 | 19,9738 | 17,4588 | 18,6026 | 19,0989 | 18,5325 | 18,1266 | | | 2 | 2 | 2 | 9,2 | 9,2 | 9,2 | 34,524 | 0 | 11,318 | 8,4E+08 | 2 | 2 | 1 | 1 | 1 | 0 | 0 | 0,68368 | 1 | 6,89442 | 1,5034 | DIPPA_2 | DIPPA_2 | DIPPA_29840.mRNA.1 26S proteasome regulatory subunit RPN13 |
| 21,1048 | 18,9067 | 19,1669 | 20,1999 | 14,4164 | 10,0295 | 23,6166 | 21,0288 | 17,4304 | 21,537 | 18,6813 | 17,9771 | | | 3 | 3 | 3 | 21,7 | 21,7 | 21,7 | 19,803 | 0 | 18,562 | 4,1E+07 | 5 | 2 | 3 | 2 | 2 | 1 | 0 | 0,73371 | 1 | 4,84423 | 1,60104 | DIPPA_0 | DIPPA_0 | DIPPA_08083.mRNA.1 Nascent polypeptide-associated complex subunit alpha-like protein;DIPPA_08116.mRNA.1 Nascent polypeptide-associated complex subunit alpha-like protein;DIPPA_08077.mRNA.1 Nascent polypeptide-associated complex subunit alpha-like protein |
| 19,7739 | 14,9358 | 16,1492 | 15,9905 | 10,3959 | 10,4843 | 21,8546 | 18,1793 | 18,991 | 20,3441 | 19,2122 | 18,6257 | | | 5 | 5 | 5 | 21,2 | 21,2 | 21,2 | 38,488 | 0 | 38,604 | 1,5E+07 | 9 | 5 | 2 | 1 | 0 | 0 | 0 | 0,92605 | 1 | 4,66273 | 1,98176 | DIPPA_1 | DIPPA_1 | DIPPA_10103.mRNA.1 hypothetical protein |
| 21,6349 | 14,8954 | 13,7247 | 14,9574 | 10,2464 | 12,7583 | 24,6937 | 17,1958 | 17,4452 | 19,6769 | 18,1307 | 18,0695 | | | 21 | 21 | 21 | 36 | 36 | 36 | 120,29 | 0 | 209,04 | 1,7E+08 | 28 | 21 | 3 | 2 | 0 | 0 | 1 | 0,659 | 1 | 4,09763 | 1,45531 | DIPPA_0 | DIPPA_0 | DIPPA_00542.mRNA.1 hypothetical protein |
| 21,3268 | 13,3838 | 15,7013 | 15,1697 | 11,3293 | 13,4643 | 24,6872 | 17,3026 | 19,8279 | 19,7826 | 19,3672 | 18,5928 | | | 17 | 17 | 17 | 36,8 | 36,8 | 36,8 | 73,667 | 0 | 114,1 | 8,9E+07 | 18 | 17 | 2 | 4 | 0 | 0 | 1 | 0,59777 | 1 | 3,48287 | 1,33603 | DIPPA_2 | DIPPA_2 | DIPPA_27525.mRNA.1 Nuclear pore complex protein Nup98-Nup96 OS_Mus musculus GN_Nup98 PE_1 SV_2 |
| 16,6155 | 12,94 | 15,273 | 15,1318 | 8,86738 | 10,5485 | 20,777 | 17,8752 | 17,5423 | 18,4085 | 19,2465 | 17,7603 | | | 2 | 2 | 2 | 7,5 | 7,5 | 7,5 | 47,134 | 0 | 11,792 | 2660100 | 1 | 1 | 0 | 1 | 0 | 0 | 0 | 0,72703 | 1 | 3,42695 | 1,58797 | DIPPA_1 | DIPPA_1 | DIPPA_15671.mRNA.1 Probable leucine aminopeptidase 1 |
| 21,2815 | 18,7629 | 17,3759 | 18,4642 | 16,9814 | 12,1673 | 23,7614 | 21,9128 | 18,437 | 18,7496 | 21,7327 | 17,2713 | | | 3 | 3 | 3 | 22,6 | 22,6 | 22,6 | 21,393 | 0 | 71,453 | 3,7E+07 | 6 | 3 | 2 | 1 | 1 | 3 | 0 | 0,66856 | 1 | 3,26912 | 1,47394 | DIPPA_1 | DIPPA_1 | DIPPA_11165.mRNA.1 Protein disulfide-isomerase-like protein EhSep2;DIPPA_11165.mRNA.2 4 5-DOPA dioxygenase extradiol |
| 21,1474 | 14,3884 | 10,295 | 13,6875 | 10,662 | 11,8228 | 23,7916 | 18,2796 | 18,9976 | 19,814 | 18,6682 | 17,7395 | | | 11 | 11 | 11 | 24,4 | 24,4 | 24,4 | 82,647 | 0 | 230,13 | 6,3E+07 | 9 | 11 | 2 | 0 | 0 | 0 | 0 | 0,41738 | 1 | 3,21953 | 0,9802 | DIPPA_1 | DIPPA_1 | DIPPA_16310.mRNA.1 hypothetical protein |
| 22,0035 | 18,4113 | 19,5878 | 21,3178 | 10,3037 | 19,1393 | 24,0041 | 21,9522 | 23,0677 | 23,0265 | 18,9988 | 23,0662 | | | 3 | 3 | 3 | 29,9 | 29,9 | 29,9 | 23,232 | 0 | 26,746 | 6,8E+07 | 12 | 3 | 1 | 2 | 2 | 0 | 2 | 0,36447 | 1 | 3,0806 | 0,87279 | DIPPA_1 | DIPPA_1 | DIPPA_14508.mRNA.1 Selenoprotein M |
| 21,2564 | 18,9834 | 18,6151 | 14,7185 | 16,6896 | 18,4568 | 22,9875 | 20,6147 | 17,8983 | 20,2767 | 19,1011 | 19,582 |  |  | 2 | 2 | 2 | 35,9 | 35,9 | 35,9 | 13,26 | 0 | 18,26 | 1,9E+07 | 7 | 2 | 2 | 1 | 0 | 2 | 1 | 1,03529 | 1 | 2,99667 | 2,20444 | DIPPA_3 | DIPPA_3 | DIPPA_33223.mRNA.1 hypothetical protein conserved |
| 20,8919 | 15,7978 | 18,1663 | 18,9828 | 13,2917 | 13,6354 | 23,8432 | 19,8954 | 21,6659 | 19,2206 | 18,8956 | 17,8 | | | 3 | 3 | 2 | 22,8 | 22,8 | 16 | 17,886 | 0 | 33,262 | 3,4E+07 | 5 | 2 | 2 | 2 | 1 | 1 | 0 | 0,56118 | 1 | 2,98203 | 1,26461 | DIPPA_1 | DIPPA_1 | DIPPA_12857.mRNA.1 60S ribosomal protein L6;DIPPA_00907.mRNA.1 60S ribosomal protein L6 |
| 16,7908 | 12,0406 | 14,5992 | 13,764 | 11,6319 | 9,12186 | 20,232 | 16,6806 | 18,0611 | 19,435 | 18,6254 | 19,5982 | | | 6 | 6 | 6 | 16,2 | 16,2 | 16,2 | 61,811 | 0 | 37,012 | 4411300 | 6 | 6 | 0 | 1 | 1 | 0 | 0 | 0,7065 | 1 | 2,97094 | 1,54789 | DIPPA_0 | DIPPA_0 | DIPPA_02852.mRNA.1 T-complex protein 1 subunit eta |
| 19,1309 | 14,3822 | 10,5744 | 15,2433 | 10,0088 | 10,4985 | 20,6901 | 17,2797 | 19,592 | 19,8022 | 19,2017 | 18,6297 | | | 7 | 7 | 7 | 15,4 | 15,4 | 15,4 | 67,423 | 0 | 43,669 | 1,8E+07 | 7 | 7 | 1 | 0 | 1 | 0 | 0 | 0,39296 | 1 | 2,77896 | 0,93088 | DIPPA_3 | DIPPA_3 | DIPPA_33414.mRNA.1 ATP-dependent RNA helicase DBP2 |
| 20,0223 | 12,2068 | 16,2854 | 16,7473 | 11,7218 | 11,8907 | 22,1478 | 16,8096 | 20,788 | 20,6965 | 19,3071 | 17,6425 | | | 3 | 3 | 3 | 17,2 | 17,2 | 17,2 | 33,471 | 0 | 22,363 | 1,5E+07 | 5 | 3 | 0 | 2 | 2 | 0 | 0 | 0,41369 | 1 | 2,71824 | 0,97276 | DIPPA_3 | DIPPA_3 | DIPPA_35556.mRNA.2 hypothetical protein;DIPPA_08165.mRNA.1 hypothetical protein;DIPPA_08164.mRNA.2 hypothetical protein;DIPPA_08156.mRNA.1 hypothetical protein;DIPPA_35556.mRNA.1 hypothetical protein;DIPPA_08164.mRNA.1 hypothetical protein |
| 21,4193 | 15,6164 | 17,3393 | 21,448 | 14,117 | 11,1135 | 22,7718 | 16,1823 | 21,2601 | 21,5135 | 18,4449 | 18,6191 | | | 4 | 4 | 4 | 28,3 | 28,3 | 28,3 | 15,803 | 0 | 29,202 | 2,9E+07 | 8 | 4 | 1 | 1 | 3 | 1 | 1 | 0,29557 | 1 | 2,56549 | 0,72911 | DIPPA_1 | DIPPA_1 | DIPPA_15219.mRNA.2 40S ribosomal protein S24-2;DIPPA_15219.mRNA.1 40S ribosomal protein S24-2 |
| 23,3233 | 16,6756 | 19,0831 | 22,0028 | 10,7228 | 18,752 | 25,5914 | 16,2315 | 18,3957 | 25,0992 | 17,6619 | 18,1219 | | | 3 | 3 | 3 | 27,9 | 27,9 | 27,9 | 22,142 | 0 | 62,737 | 1,9E+08 | 12 | 3 | 1 | 2 | 1 | 0 | 1 | 0,26067 | 1 | 2,53479 | 0,65419 | DIPPA_3 | DIPPA_3 | DIPPA_30006.mRNA.1 Thioredoxin peroxidase |
| 21,7094 | 18,5704 | 19,2914 | 19,3253 | 16,2419 | 16,4141 | 24,6803 | 21,7561 | 23,3008 | 22,2501 | 22,1314 | 21,9338 | | | 7 | 7 | 7 | 30,4 | 30,4 | 30,4 | 27,244 | 0 | 48,978 | 7,4E+07 | 13 | 7 | 2 | 3 | 2 | 2 | 2 | 0,85239 | 1 | 2,52997 | 1,83458 | DIPPA_3 | DIPPA_3 | DIPPA_32269.mRNA.1 hypothetical protein |
| 20,5548 | 11,3309 | 16,414 | 19,9593 | 10,9493 | 10,1097 | 24,1092 | 16,4916 | 22,1688 | 23,4234 | 18,4691 | 18,0358 | | | 8 | 8 | 8 | 26,6 | 26,6 | 26,6 | 50,069 | 0 | 57,137 | 6,4E+07 | 15 | 8 | 0 | 2 | 5 | 0 | 0 | 0,23042 | 1 | 2,42717 | 0,58775 | DIPPA_2 | DIPPA_2 | DIPPA_22663.mRNA.1 ATP-dependent RNA helicase uap56 |
| 20,7319 | 15,9411 | 19,2225 | 19,8742 | 12,9423 | 15,9903 | 24,2394 | 21,7971 | 23,8285 | 22,1959 | 19,4394 | 18,18 | | | 5 | 2 | 2 | 24 | 13 | 13 | 40,811 | 0 | 51,433 | 4,5E+07 | 10 | 2 | 1 | 2 | 2 | 1 | 1 | 0,40873 | 1 | 2,36289 | 0,96276 | DIPPA_2 | DIPPA_2 | DIPPA_29150.mRNA.1 Proteinase R |
| 19,7939 | 15,5242 | 15,0049 | 20,0475 | 13,3216 | 9,87137 | 21,701 | 17,4779 | 18,1833 | 21,4769 | 19,8297 | 17,8147 | | | 2 | 2 | 2 | 11,6 | 11,6 | 11,6 | 14,933 | 0 | 13,808 | 1,5E+07 | 4 | 2 | 1 | 1 | 2 | 1 | 0 | 0,28401 | 1 | 2,3608 | 0,7045 | DIPPA_2 | DIPPA_2 | DIPPA_26292.mRNA.1 Histone H2B.1;TR133520__c8_g2_i1__m.32469 #MITO 1 Mitoprot #PFAM PF00125.19 Histone Core histone H2A/H2B/H3/H4 #BLASTP_UNIREF90 7e-18 Histone H2B n=3 Tax=Bacillariophyta RepID=B7FWR8_PHATC (length=116);DIPPA_26287.mRNA.1 Histone H2B.1 |
| 20,7412 | 12,3885 | 13,8171 | 15,0002 | 11,8315 | 13,1963 | 23,8735 | 15,3599 | 16,9089 | 18,9269 | 18,0424 | 17,6546 | | | 17 | 17 | 17 | 27,8 | 27,8 | 27,8 | 108,64 | 0 | 182,46 | 8,7E+07 | 21 | 17 | 2 | 3 | 0 | 0 | 1 | 0,34971 | 1 | 2,30626 | 0,84241 | DIPPA_2 | DIPPA_2 | DIPPA_22976.mRNA.1 Nuclear pore complex protein Nup107 OS_Homo sapiens GN_NUP107 PE_1 SV_1 |
| 23,974 | 17,5806 | 14,7496 | 23,5411 | 16,2395 | 9,71612 | 25,8923 | 18,6706 | 18,2282 | 23,7821 | 22,2082 | 17,9381 | | | 5 | 5 | 5 | 37,5 | 37,5 | 37,5 | 18,025 | 0 | 32,37 | 2,6E+08 | 7 | 5 | 1 | 1 | 5 | 2 | 0 | 0,17826 | 1 | 2,26919 | 0,46922 | DIPPA_2 | DIPPA_2 | DIPPA_21650.mRNA.1 60S ribosomal protein L21;DIPPA_21649.mRNA.1 60S ribosomal protein L21 |
| 21,8943 | 12,7504 | 15,7605 | 20,6641 | 11,5826 | 11,5047 | 24,3492 | 17,0883 | 17,9917 | 23,1034 | 19,3641 | 18,7245 | | | 6 | 6 | 3 | 31,8 | 31,8 | 13,6 | 24,787 | 0 | 38,635 | 8,4E+07 | 7 | 3 | 0 | 0 | 0 | 0 | 0 | 0,21191 | 1 | 2,21797 | 0,54633 | DIPPA_2 | DIPPA_2 | DIPPA_28055.mRNA.1 GTP-binding nuclear protein spi1 |
| 18,2081 | 11,6456 | 14,9484 | 17,0985 | 10,9269 | 10,1643 | 19,8999 | 15,292 | 18,0657 | 20,1063 | 19,3797 | 19,1409 | | | 2 | 2 | 2 | 16,3 | 16,3 | 16,3 | 23,75 | 0 | 12,771 | 5697100 | 2 | 2 | 0 | 1 | 1 | 0 | 0 | 0,31022 | 1 | 2,20411 | 0,76011 | DIPPA_0 | DIPPA_0 | DIPPA_05541.mRNA.1 Peroxiredoxin 1 |
| 20,8074 | 17,338 | 17,5464 | 20,1953 | 14,6305 | 14,2855 | 23,6195 | 16,7643 | 19,0562 | 22,7349 | 19,0546 | 18,2407 | | | 2 | 2 | 2 | 9,9 | 9,9 | 9,9 | 36,795 | 0 | 84,215 | 4,1E+07 | 6 | 2 | 1 | 1 | 2 | 1 | 1 | 0,42126 | 1 | 2,19352 | 0,98798 | DIPPA_1 | DIPPA_1 | DIPPA_12990.mRNA.1 Chaperone protein dnaJ 2 |
| 19,1303 | 11,6974 | 14,4462 | 14,6488 | 12,5007 | 11,6606 | 22,8467 | 16,8004 | 17,3961 | 20,8936 | 19,6125 | 18,7988 | | | 5 | 5 | 5 | 8,7 | 8,7 | 8,7 | 93,213 | 0 | 77,396 | 1,7E+07 | 8 | 5 | 0 | 1 | 1 | 0 | 0 | 0,38699 | 1 | 2,15461 | 0,91875 | DIPPA_2 | DIPPA_2 | DIPPA_20050.mRNA.1 hypothetical protein |
| 19,3419 | 12,4781 | 11,7953 | 13,5586 | 11,2281 | 12,5882 | 23,951 | 17,2574 | 18,9322 | 19,7523 | 18,509 | 18,0095 | | | 10 | 10 | 10 | 9,4 | 9,4 | 9,4 | 200,37 | 0 | 78,663 | 6,5E+07 | 11 | 10 | 2 | 1 | 0 | 0 | 1 | 0,34424 | 1 | 2,0801 | 0,83111 | DIPPA_1 | DIPPA_1 | DIPPA_17330.mRNA.1 hypothetical protein |
| 16,9612 | 11,7732 | 13,4251 | 13,2959 | 11,1009 | 11,5471 | 21,1021 | 16,4526 | 19,9035 | 18,6083 | 18,6762 | 17,326 | | | 11 | 11 | 11 | 4,4 | 4,4 | 4,4 | 474,79 | 0 | 68,026 | 2,4E+07 | 14 | 11 | 0 | 1 | 1 | 0 | 0 | 0,54878 | 1 | 2,07184 | 1,24035 | DIPPA_0 | DIPPA_0 | DIPPA_09931.mRNA.1 hypothetical protein |
| [illegible] | [illegible] | [illegible] | [illegible] | [illegible] | [illegible] | [illegible] | [illegible] | [illegible] | [illegible] | [illegible] | [illegible] | [illegible] | [illegible] | [illegible] | [illegible] | [illegible] | [illegible] | [illegible] | [illegible] | [illegible] | [illegible] | [illegible] | [illegible] | [illegible] | [illegible] | [illegible] | [illegible] | [illegible] | [illegible] | [illegible] | [illegible] | [illegible] | [illegible] | [illegible] | [illegible] | [illegible] | [illegible] |

| iBAQ PnH | iBAQ PnH | iBAQ PnH | iBAQ wt- | iBAQ wt- | iBAQ wt- | LFQ inten | LFQ inten | LFQ inten | LFQ inten | LFQ inten | LFQ inten | Student's | Student's | Peptides | Razor + u | Unique pe | Sequence | Unique + | Unique se | Mol. weig | Q-value | Score | Intensity | MS/MS co | Unique pe | Unique pe | Unique pe | Unique pe | Unique pe | Unique pe | -Log Stud | Student's | Student's | Student's | Protein IE | Majority | Fasta headers |
|---|---|---|---|---|---|---|---|---|---|---|---|---|---|---|---|---|---|---|---|---|---|---|---|---|---|---|---|---|---|---|---|---|---|---|---|---|---|
| 27,8047 | 28,1599 | 28,238 | 14,5367 | 16,5655 | 15,0484 | 29,986 | 29,9178 | 29,9712 | 19,4894 | 20,5597 | 20,8631 | + | iBAQ PnH | 19 | 19 | 19 | 79,7 | 79,7 | 79,7 | 24,345 | 0 | 323,31 | 1,1E+10 | 106 | 17 | 18 | 17 | 0 | 1 | 0 | 4,46231 | 0 | 12,684 | 20,3411 | DIPPA_0 | DIPPA_0 | DIPPA_09665.mRNA.2 hypothetical protein;DIPPA_09665.mRNA.1 hypothetical protein |
| 25,2802 | 25,7097 | 25,995 | 18,2602 | 16,5086 | 18,8378 | 28,3702 | 28,3426 | 28,4142 | 19,5644 | 20,4315 | 20,6432 | + | iBAQ PnH | 13 | 13 | 13 | 51,9 | 51,9 | 51,9 | 26,941 | 0 | 99,172 | 1,8E+09 | 30 | 10 | 11 | 12 | 1 | 0 | 1 | 3,35934 | 0 | 7,7928 | 10,6689 | DIPPA_3 | DIPPA_3 | DIPPA_31723.mRNA.1 hypothetical protein AMSG_10510 |
| 17,6545 | 18,4557 | 18,0393 | 19,6818 | 20,806 | 20,1905 | 21,6328 | 23,5355 | 23,8521 | 24,796 | 24,5781 | 23,9968 | | | 6 | 6 | 6 | 14,7 | 14,7 | 14,7 | 58,323 | 0 | 39,031 | 1,6E+08 | 11 | 1 | 3 | 4 | 4 | 6 | 4 | 2,26059 | 0,424 | -2,1763 | -5,4551 | DIPPA_2 | DIPPA_2 | DIPPA_20627.mRNA.1 Apoptosis inhibitor 5-like protein API5 |
| 19,5705 | 20,1464 | 20,2131 | 14,085 | 17,1999 | 15,7031 | 23,3261 | 23,3092 | 23,4722 | 20,4488 | 20,2488 | 21,2183 | | | 10 | 10 | 10 | 14,2 | 14,2 | 14,2 | 130,25 | 0 | 87,891 | 1,3E+08 | 25 | 7 | 7 | 10 | 1 | 3 | 0 | 2,02391 | 0,64 | 4,31404 | 4,67763 | DIPPA_0 | DIPPA_0 | DIPPA_09941.mRNA.1 Protein transport protein SEC31 |
| 17,4073 | 17,873 | 18,4978 | 14,4305 | 16,0542 | 14,2112 | 23,5926 | 22,5806 | 23,2929 | 20,0592 | 21,1871 | 20,8279 | | | 4 | 4 | 4 | 5,1 | 5,1 | 5,1 | 195,31 | 0 | 32,474 | 3,9E+07 | 5 | 2 | 3 | 4 | 1 | 1 | 1 | 1,99087 | 0,512 | 3,02741 | 4,57625 | DIPPA_0 | DIPPA_0 | DIPPA_06136.mRNA.1 hypothetical protein DQ04_07291000;TR92002__c0_g1_i1__m.19319 #MITO 2 Predotar,HMMTOP #PFAM 2 x PF04403.8 PqiA Paraquat-inducible protein A;2 x PF08545.5 ACP_syn_III 3-Oxoacyl-[acyl-carrier-protein (ACP)] synthase III #BLASTP_UNIREF90 |
| 19,8167 | 19,3157 | 19,5091 | 20,975 | 22,5662 | 21,6387 | 23,6624 | 23,6094 | 23,3362 | 24,2057 | 24,4969 | 24,0071 | | | 6 | 6 | 6 | 34,9 | 34,9 | 34,9 | 33,626 | 0 | 54,515 | 1,9E+08 | 19 | 2 | 2 | 3 | 4 | 5 | 4 | 1,96681 | 0,44733 | -2,1795 | -4,5034 | DIPPA_0 | DIPPA_0 | DIPPA_08181.mRNA.1 Cytochrome b-c1 complex subunit Rieske-3 mitochondrial |
| 14,5141 | 15,4597 | 14,6155 | 17,0193 | 18,2614 | 16,6406 | 20,8308 | 21,0312 | 20,2951 | 20,8917 | 20,7839 | 20,7688 | | | 3 | 3 | 3 | 7,4 | 7,4 | 7,4 | 55,991 | 0 | 18,444 | 1,3E+07 | 5 | 1 | 1 | 1 | 3 | 3 | 1 | 1,88359 | 0,49314 | -2,444 | -4,258 | DIPPA_0 | DIPPA_0 | DIPPA_07891.mRNA.1 hypothetical protein |
| 20,7117 | 21,1319 | 21,3122 | 22,4869 | 23,9465 | 22,5563 | 23,1558 | 23,3248 | 23,1713 | 24,1248 | 24,1779 | 23,9708 | | | 11 | 11 | 11 | 51,1 | 51,1 | 51,1 | 24,135 | 0 | 157,78 | 4,5E+08 | 39 | 5 | 7 | 7 | 10 | 9 | 9 | 1,73047 | 0,7 | -1,9447 | -3,8311 | DIPPA_1 | DIPPA_1 | DIPPA_15108.mRNA.1 Glutathione S-transferase 1;DIPPA_04328.mRNA.1 Glutathione S-transferase 1 |
| 14,1414 | 14,4544 | 15,3576 | 16,7895 | 17,7236 | 16,1139 | 21,378 | 19,7681 | 20,2269 | 20,8033 | 20,4297 | 20,58 | | | 2 | 2 | 2 | 10,4 | 10,4 | 10,4 | 44,655 | 0 | 14,735 | 1,1E+07 | 2 | 0 | 1 | 1 | 2 | 2 | 1 | 1,70249 | 0,64978 | -2,2245 | -3,7563 | DIPPA_3 | DIPPA_3 | DIPPA_33968.mRNA.1 Actin-related protein 2 |
| 21,5479 | 22,0344 | 21,6254 | 22,4706 | 23,5552 | 22,9669 | 24,2285 | 24,4054 | 23,9895 | 24,3713 | 24,1704 | 24,8856 | | | 8 | 8 | 8 | 67,6 | 67,6 | 67,6 | 16,982 | 0 | 66,483 | 2,6E+08 | 12 | 3 | 4 | 4 | 5 | 7 | 5 | 1,65305 | 0,6928 | -1,2617 | -3,6264 | DIPPA_7 | DIPPA_7 | DIPPA_70185.mRNA.1 hypothetical protein;DIPPA_26697.mRNA.1 40S ribosomal protein S15;DIPPA_26691.mRNA.1 40S ribosomal protein S15 |
| 17,7412 | 19,8904 | 20,1658 | 21,5111 | 23,6444 | 22,7248 | 22,8715 | 22,7061 | 22,873 | 23,734 | 24,5818 | 25,0192 | | | 7 | 7 | 7 | 35,2 | 35,2 | 35,2 | 24,062 | 0 | 46,558 | 3E+08 | 16 | 2 | 4 | 4 | 6 | 7 | 6 | 1,56984 | 0,80509 | -3,3609 | -3,4141 | DIPPA_1 | DIPPA_1 | DIPPA_16372.mRNA.1 Glutathione S-transferase 4 |
| 18,6448 | 18,9575 | 20,4249 | 17,2313 | 17,6119 | 17,486 | 22,6314 | 22,4602 | 23,5852 | 22,9815 | 22,0097 | 23,2512 | | | 4 | 4 | 4 | 11,9 | 11,9 | 11,9 | 63,933 | 0 | 28,024 | 5,4E+07 | 8 | 2 | 2 | 4 | 3 | 3 | 2 | 1,56079 | 0,746 | 1,89933 | 3,3915 | DIPPA_3 | DIPPA_3 | DIPPA_33238.mRNA.1 Chaperone protein DnaJ |
| 22,1912 | 21,9403 | 21,847 | 22,7475 | 24,0123 | 23,1135 | 24,848 | 25,3066 | 24,7942 | 25,4496 | 25,6521 | 25,4083 | | | 7 | 7 | 7 | 32,6 | 32,6 | 32,6 | 27,978 | 0 | 90,186 | 5,5E+08 | 24 | 3 | 5 | 6 | 6 | 7 | 6 | 1,53702 | 0,75538 | -1,2983 | -3,3325 | DIPPA_0 | DIPPA_0 | DIPPA_05558.mRNA.1 hypothetical protein TraAM80_06864 |
| 16,2195 | 17,4988 | 17,3612 | 18,2204 | 19,7844 | 19,0368 | 20,4094 | 20,4441 | 21,123 | 20,3661 | 22,2319 | 22,7852 | | | 2 | 2 | 2 | 12,3 | 12,3 | 12,3 | 44,584 | 0 | 13,615 | 3,9E+07 | 2 | 1 | 1 | 1 | 2 | 2 | 2 | 1,51342 | 0,75257 | -1,9874 | -3,2744 | TR11651 | TR11651 | TR116518__c3_g1_i1__m.27004 #MITO 0 #PFAM PF01053.15 Cys_Met_Meta_PP Cys/Met metabolism PLP-dependent enzyme;PF00155.16 Aminotran_1_2 Aminotransferase class I and II;PF01041.12 DegT_DnrJ_EryC1 DegT/DnrJ/EryC1/StrS aminotransferase family;PF00266.14 A |
| 14,226 | 15,9677 | 14,9427 | 16,6926 | 18,3195 | 17,0017 | 20,848 | 20,1566 | 21,0154 | 21,0272 | 21,1454 | 20,3165 | | | 3 | 3 | 3 | 14,1 | 14,1 | 14,1 | 63,385 | 0 | 20,812 | 1,8E+07 | 3 | 0 | 1 | 1 | 2 | 3 | 1 | 1,49451 | 0,73893 | -2,2925 | -3,2284 | DIPPA_0 | DIPPA_0 | DIPPA_05208.mRNA.1 Dihydroxy-acid dehydratase |
| 18,0653 | 17,8815 | 18,3161 | 19,6183 | 20,4971 | 18,8325 | 21,4303 | 20,1474 | 20,1712 | 20,4657 | 20,0859 | 20,777 | | | 2 | 2 | 2 | 34,8 | 34,8 | 34,8 | 15,417 | 0 | 13,381 | 2,2E+07 | 2 | 1 | 1 | 1 | 2 | 2 | 1 | 1,45877 | 0,7875 | -1,5617 | -3,1423 | DIPPA_0 | DIPPA_0 | DIPPA_05869.mRNA.1 60S ribosomal protein L22;DIPPA_31488.mRNA.1 60S ribosomal protein L22 |
| 16,5264 | 17,1942 | 17,691 | 18,2388 | 19,8071 | 18,5864 | 21,0415 | 20,4579 | 22,008 | 21,5522 | 21,5403 | 21,6498 | | | 3 | 3 | 3 | 10,8 | 10,8 | 10,8 | 52,027 | 0 | 49,653 | 3,3E+07 | 9 | 1 | 1 | 2 | 3 | 3 | 2 | 1,39194 | 0,89553 | -1,7402 | -2,9845 | DIPPA_2 | DIPPA_2 | DIPPA_26538.mRNA.1 hypothetical protein |
| 19,6552 | 20,0478 | 19,6491 | 20,4999 | 21,5696 | 20,5758 | 24,3652 | 24,0498 | 24,2769 | 24,1743 | 24,3917 | 24,0839 | | | 14 | 14 | 14 | 38,9 | 38,9 | 38,9 | 63,418 | 0 | 323,31 | 3,1E+08 | 37 | 5 | 6 | 7 | 10 | 11 | 9 | 1,38778 | 0,86244 | -1,0977 | -2,9749 | DIPPA_2 | DIPPA_2 | DIPPA_20247.mRNA.1 conserved hypothetical protein |
| 14,7861 | 16,3622 | 16,6206 | 17,6149 | 19,1566 | 17,5838 | 21,7109 | 21,6874 | 21,6675 | 21,6532 | 22,2197 | 21,6785 | | | 4 | 4 | 4 | 8,9 | 8,9 | 8,9 | 87,068 | 0 | 26,914 | 6,2E+07 | 7 | 1 | 2 | 2 | 4 | 4 | 4 | 1,32851 | 1 | -2,1955 | -2,8386 | DIPPA_2 | DIPPA_2 | DIPPA_24500.mRNA.1 Probable glutamate--tRNA ligase cytoplasmic |
| 19,5532 | 19,5774 | 19,399 | 20,1258 | 21,7854 | 20,7272 | 23,9259 | 23,2894 | 22,7975 | 23,3012 | 23,6471 | 23,9326 | | | 14 | 14 | 14 | 27,5 | 27,5 | 27,5 | 73,574 | 0 | 181,5 | 3,4E+08 | 35 | 6 | 7 | 10 | 10 | 14 | 9 | 1,31359 | 1 | -1,3696 | -2,8048 | DIPPA_2 | DIPPA_2 | DIPPA_29361.mRNA.1 Alcohol dehydrogenase class-3 |
| 20,1747 | 20,1582 | 20,3321 | 20,9562 | 21,9754 | 20,823 | 23,6461 | 23,4619 | 24,0134 | 24,2598 | 24,3289 | 23,7583 | | | 12 | 12 | 12 | 43,7 | 43,7 | 43,7 | 43,231 | 0 | 88,75 | 2,8E+08 | 34 | 5 | 7 | 9 | 9 | 12 | 7 | 1,31022 | 0,98286 | -1,0298 | -2,7972 | DIPPA_2 | DIPPA_2 | DIPPA_23822.mRNA.1 Sorbitol dehydrogenase |
| 21,2411 | 22,0342 | 21,6149 | 22,0761 | 22,7382 | 22,7274 | 25,3087 | 24,5191 | 24,5724 | 24,382 | 24,1787 | 25,2426 | | | 8 | 6 | 6 | 52,9 | 34 | 34 | 21,382 | 0 | 116,61 | 3,1E+08 | 16 | 4 | 4 | 3 | 6 | 6 | 4 | 1,30672 | 0,95255 | -0,8838 | -2,7893 | DIPPA_2 | DIPPA_2 | DIPPA_27907.mRNA.1 Superoxide dismutase |
| 15,4243 | 14,4621 | 16,1913 | 17,269 | 18,1805 | 16,4793 | 21,608 | 20,782 | 20,0552 | 20,6646 | 20,717 | 20,1437 | | | 2 | 2 | 2 | 4,8 | 4,8 | 4,8 | 52,62 | 0 | 11,669 | 1,4E+07 | 3 | 1 | 1 | 1 | 2 | 2 | 2 | 1,30305 | 0,92609 | -1,9504 | -2,781 | DIPPA_0 | DIPPA_0 | DIPPA_07861.mRNA.1 Serine/threonine-protein phosphatase 2B catalytic subunit |
| 20,2508 | 20,9555 | 20,9064 | 21,9414 | 23,0875 | 21,4853 | 24,383 | 24,3888 | 25,1195 | 25,136 | 25,115 | 24,6408 | | | 9 | 9 | 4 | 29,1 | 29,1 | 18 | 65,686 | 0 | 222,32 | 6,8E+08 | 41 | 2 | 2 | 4 | 4 | 4 | 2 | 1,30216 | 0,8875 | -1,4672 | -2,779 | DIPPA_2 | DIPPA_2 | DIPPA_27311.mRNA.1 Probable protein phosphatase methylesterase 1 |
| 16,8172 | 18,5189 | 17,9298 | 19,1987 | 20,3903 | 19,0822 | 21,3322 | 21,6507 | 21,5117 | 22,0979 | 22,0861 | 21,674 | | | 6 | 6 | 6 | 17,1 | 17,1 | 17,1 | 62,977 | 0 | 55,158 | 9,8E+07 | 12 | 2 | 5 | 4 | 6 | 6 | 6 | 1,29736 | 0,86464 | -1,8017 | -2,7682 | DIPPA_3 | DIPPA_3 | DIPPA_34053.mRNA.1 Acetyl-CoA carboxylase |
| 19,8159 | 19,7233 | 19,83 | 20,3451 | 21,9332 | 20,9089 | 21,1167 | 20,7485 | 20,5241 | 21,176 | 21,4528 | 21,5302 | | | 2 | 2 | 2 | 47,7 | 47,7 | 47,7 | 10,847 | 0 | 104,6 | 3E+07 | 11 | 1 | 2 | 1 | 2 | 2 | 2 | 1,28071 | 0,89369 | -1,2727 | -2,7309 | DIPPA_7 | DIPPA_7 | DIPPA_70188.mRNA.1 hypothetical protein |
| 13,9874 | 16,7765 | 17,0649 | 19,1647 | 20,2236 | 18,0108 | 20,9051 | 22,6376 | 22,7456 | 22,4914 | 22,8611 | 20,8004 | | | 4 | 4 | 4 | 15 | 15 | 15 | 55,724 | 0 | 33,715 | 6E+07 | 6 | 0 | 2 | 2 | 3 | 4 | 1 | 1,27775 | 0,86519 | -3,1901 | -2,7243 | DIPPA_0 | DIPPA_0 | DIPPA_01787.mRNA.1 Importin subunit alpha-4 |
| 19,5612 | 20,4394 | 20,9221 | 21,81 | 22,9342 | 21,2851 | 23,3016 | 23,727 | 23,76 | 24,3729 | 24,912 | 23,7673 | | | 3 | 3 | 2 | 30,8 | 30,8 | 25,4 | 14,693 | 0 | 99,764 | 1,5E+08 | 11 | 1 | 2 | 2 | 2 | 2 | 2 | 1,27017 | 0,86857 | -1,7022 | -2,7074 | DIPPA_0 | DIPPA_0 | DIPPA_09967.mRNA.1 40S ribosomal protein S15a-1;DIPPA_09940.mRNA.1 40S ribosomal protein S15a-1 |
| 15,3178 | 14,8932 | 14,1714 | 15,6623 | 15,5958 | 16,0062 | 21,1053 | 20,2567 | 20,9699 | 20,705 | 21,0892 | 20,6532 | | | 6 | 2 | 2 | 8,4 | 3 | 3 | 102,51 | 0 | 12,922 | 7230300 | 3 | 1 | 0 | 2 | 2 | 2 | 1 | 1,25938 | 0,86841 | -0,9606 | -2,6834 | DIPPA_7 | DIPPA_7 | DIPPA_70006.mRNA.1 hypothetical conserved protein;DIPPA_70005.mRNA.1 hypothetical conserved protein |
| 17,4886 | 18,13 | 18,9263 | 19,7303 | 21,9826 | 19,7315 | 22,5694 | 23,374 | 23,18 | 23,0622 | 23,6557 | 23,1117 | | | 12 | 12 | 12 | 34 | 34 | 34 | 71,867 | 0 | 85,291 | 2,3E+08 | 17 | 3 | 5 | 5 | 6 | 12 | 6 | 1,25794 | 0,84267 | -2,2998 | -2,6802 | DIPPA_0 | DIPPA_0 | DIPPA_03297.mRNA.1 Transketolase |
| 18,9578 | 18,0901 | 19,2727 | 19,5452 | 21,434 | 20,4311 | 22,433 | 22,5294 | 22,6378 | 23,1831 | 23,7703 | 22,9347 | | | 5 | 5 | 5 | 27,5 | 27,5 | 27,5 | 32,534 | 0 | 30,704 | 1,3E+08 | 10 | 3 | 3 | 4 | 4 | 5 | 5 | 1,22583 | 0,9169 | -1,6966 | -2,6095 | DIPPA_5 | DIPPA_5 | DIPPA_51020.mRNA.2 Branched-chain amino acid aminotransferase/4-amino-4-deoxychorismate lyase n=1 Tax=Clostridium aceticum RepID=A0A0D8I9F1_9CLOT (length=283);DIPPA_51020.mRNA.1 Branched-chain amino acid aminotransferase/4-amino-4-deoxychorismate lyase n=1 |
| 12,6852 | 13,2787 | 14,0197 | 15,154 | 17,0813 | 14,4717 | 21,4525 | 20,4107 | 19,5499 | 19,836 | 20,4309 | 21,2775 | | | 2 | 2 | 2 | 2,5 | 2,5 | 2,5 | 114,98 | 0 | 12,491 | 1E+07 | 2 | 0 | 1 | 1 | 0 | 2 | 1 | 1,20848 | 0,91575 | -2,2412 | -2,5716 | TR63214 | TR63214 | TR63214__c2_g1_i1__m.10097 #MITO 1 HMMTOP #PFAM 6 x PF00400.27 WD40 WD domain, G-beta repeat;3 x PF13570.1 YWTD YWTD domain #BLASTP_UNIREF90 2e-36 WD40 repeat-containing protein, putative n=1 Tax=Bodo saltans RepID=A0A0S4J9X3_BODSA (length=885) |
| 16,396 | 14,206 | 16,7378 | 18,5921 | 20,1402 | 17,2869 | 20,663 | 21,0538 | 20,9205 | 21,7388 | 21,996 | 20,8427 | | | 6 | 6 | 6 | 13,3 | 13,3 | 13,3 | 47,306 | 0 | 34,472 | 4,4E+07 | 9 | 2 | 1 | 2 | 4 | 6 | 2 | 1,18868 | 0,99321 | -2,8931 | -2,5286 | DIPPA_1 | DIPPA_1 | DIPPA_18478.mRNA.2 hypothetical protein;DIPPA_18478.mRNA.1 hypothetical protein |
| 13,98 | 15,6906 | 16,4195 | 17,0534 | 18,8933 | 17,1525 | 20,3662 | 20,3238 | 20,209 | 20,7605 | 21,3098 | 21,0743 | | | 2 | 2 | 2 | 8,2 | 8,2 | 8,2 | 42,059 | 0 | 14,347 | 1,6E+07 | 3 | 0 | 1 | 1 | 2 | 2 | 1 | 1,1713 | 1 | -2,3364 | -2,4911 | DIPPA_1 | DIPPA_1 | DIPPA_13986.mRNA.1 26S proteasome non-ATPase regulatory subunit 7 homolog B |
| 17,9918 | 18,1417 | 17,5347 | 18,5786 | 20,2469 | 18,8855 | 22,4388 | 22,0673 | 21,2584 | 21,9054 | 21,8301 | 23,0614 | | | 8 | 8 | 8 | 29,7 | 29,7 | 29,7 | 60,672 | 0 | 51,861 | 8,6E+07 | 11 | 3 | 4 | 4 | 7 | 8 | 3 | 1,1644 | 1 | -1,3476 | -2,4763 | DIPPA_1 | DIPPA_1 | DIPPA_13285.mRNA.1 hypothetical protein |
| 16,3913 | 18,0746 | 18,2996 | 19,7569 | 21,3146 | 18,7734 | 21,6522 | 22,6095 | 22,8851 | 23,0574 | 23,2839 | 22,7188 | | | 11 | 11 | 11 | 29 | 29 | 29 | 58,618 | 0 | 100,32 | 1,3E+08 | 14 | 2 | 5 | 4 | 8 | 10 | 6 | 1,16334 | 1 | -2,3598 | -2,474 | DIPPA_2 | DIPPA_2 | DIPPA_26600.mRNA.1 T-complex protein 1 subunit zeta 1 |
| 19,4444 | 19,0218 | 19,2111 | 19,7461 | 20,7793 | 19,7918 | 20,7261 | 20,5075 | 20,4962 | 20,6417 | 21,8364 | 21,6261 | | | 2 | 2 | 2 | 17,2 | 17,2 | 17,2 | 13,94 | 0 | 12,56 | 3,3E+07 | 4 | 1 | 1 | 1 | 1 | 2 | 2 | 1,15419 | 1 | -0,88 | -2,4544 | DIPPA_1 | DIPPA_1 | DIPPA_12075.mRNA.1 hypothetical protein |
| 15,423 | 17,3443 | 17,2528 | 17,7526 | 19,0808 | 18,6104 | 21,0629 | 21,8758 | 21,4882 | 21,5959 | 21,7855 | 21,7567 | | | 5 | 5 | 5 | 17 | 17 | 17 | 50,993 | 0 | 30,957 | 4E+07 | 9 | 1 | 2 | 4 | 4 | 4 | 4 | 1,154 | 0,98295 | -1,8079 | -2,454 | DIPPA_1 | DIPPA_1 | DIPPA_12962.mRNA.1 Adenylosuccinate lyase |

[illegible]

| | | | | | | | | | | | | | | | | | | | | | | |
|---|---|---|---|---|---|---|---|---|---|---|---|---|---|---|---|---|---|---|---|---|---|---|
| 20,6115 | 21,4652 | 21,7223 | 21,8059 | 23,3647 | 21,3463 | 23,5532 | 24,1079 | 23,8715 | 24,1083 | 24,0893 | 23,7284 | 21 | 21 | 21 | 40,6 | 40,6 | 40,6 | 67,423 | 0 | 246,08 | 7,3E+08 | 73 |
| 18,3898 | 18,1949 | 16,7141 | 17,7369 | 19,6047 | 18,9202 | 22,4567 | 22,2092 | 19,8001 | 21,5059 | 21,5787 | 22,5383 | 3 | 3 | 3 | 12,8 | 12,8 | 12,8 | 38,126 | 0 | 63,764 | 3,8E+07 | 7 |
| 15,7707 | 18,1634 | 17,6874 | 17,6507 | 19,3974 | 18,0699 | 21,3523 | 22,5466 | 22,0473 | 22,0923 | 22,8347 | 22,4305 | 3 | 3 | 3 | 7 | 7 | 7 | 76,377 | 0 | 37,972 | 6,4E+07 | 10 |
| 14,2575 | 17,744 | 18,7021 | 18,1427 | 20,1279 | 18,2384 | 21,5999 | 20,6213 | 20,8727 | 20,8373 | 21,4721 | 20,7794 | 2 | 2 | 2 | 12,1 | 12,1 | 12,1 | 23,227 | 0 | 12,184 | 2,4E+07 | 4 |
| 20,8087 | 21,4869 | 21,8634 | 21,7662 | 23,312 | 21,5454 | 24,0197 | 24,1084 | 24,2803 | 24,2836 | 24,5365 | 24,1013 | 14 | 14 | 14 | 47,9 | 47,9 | 47,9 | 46,187 | 0 | 253,53 | 5,1E+08 | 56 |
| 19,4487 | 14,7196 | 20,6402 | 20,6383 | 22,1122 | 19,6007 | 20,8105 | 20,7872 | 22,7954 | 22,8643 | 23,1021 | 21,4808 | 2 | 2 | 2 | 34,4 | 34,4 | 34,4 | 10,564 | 0 | 11,752 | 5,6E+07 | 3 |
| 16,5019 | 17,9163 | 17,7521 | 17,5483 | 18,144 | 18,5265 | 20,3755 | 21,5476 | 21,341 | 21,334 | 20,6213 | 22,236 | 3 | 3 | 3 | 11,3 | 11,3 | 11,3 | 46,138 | 0 | 19,03 | 2,8E+07 | 7 |
| 16,5522 | 18,8214 | 20,2731 | 19,5744 | 21,1982 | 19,551 | 21,0638 | 20,369 | 19,711 | 21,1287 | 20,0533 | 21,4647 | 2 | 2 | 2 | 13,1 | 13,1 | 13,1 | 25,399 | 0 | 26,521 | 6,9E+07 | 5 |
| 17,8387 | 18,732 | 18,769 | 18,3618 | 21,1464 | 19,2166 | 22,47 | 22,8066 | 22,736 | 22,3781 | 22,6127 | 22,4138 | 3 | 3 | 3 | 19 | 19 | 19 | 31,229 | 0 | 114,26 | 7,9E+07 | 7 |
| 23,8292 | 24,7874 | 24,8899 | 24,7743 | 26,1007 | 24,7723 | 28,2004 | 28,3841 | 28,7275 | 28,543 | 28,7627 | 28,4488 | 16 | 16 | 16 | 47,9 | 47,9 | 47,9 | 41,557 | 0 | 323,31 | 4,3E+09 | 90 |
| 14,5539 | 18,0571 | 18,9598 | 19,4366 | 20,8354 | 17,5959 | 21,644 | 22,0089 | 22,6621 | 22,45 | 22,8145 | 21,9267 | 6 | 6 | 6 | 17,5 | 17,5 | 17,5 | 64,756 | 0 | 51,844 | 9,3E+07 | 12 |
| 14,2967 | 14,2427 | 15,3993 | 14,9782 | 17,2184 | 14,9639 | 20,9561 | 21,2 | 19,8389 | 21,1983 | 21,3909 | 20,6718 | 5 | 5 | 5 | 2,7 | 2,7 | 2,7 | 382,95 | 0 | 34,141 | 3,6E+07 | 6 |
| 19,0694 | 19,5215 | 20,0269 | 19,9231 | 21,7782 | 19,6759 | 22,8011 | 22,958 | 22,8804 | 23,0127 | 23,9326 | 22,9373 | 4 | 4 | 4 | 19,5 | 19,5 | 19,5 | 22,344 | 0 | 22,763 | 9,4E+07 | 7 |
| 18,517 | 20,7516 | 21,014 | 20,748 | 22,7422 | 20,7567 | 22,8925 | 23,2483 | 23,6735 | 23,9593 | 23,99 | 23,218 | 7 | 7 | 7 | 44,8 | 44,8 | 44,8 | 28,757 | 0 | 150,55 | 2,5E+08 | 22 |
| 21,173 | 21,6867 | 22,8118 | 22,2308 | 24,0656 | 22,3191 | 24,3065 | 24,4012 | 24,6972 | 24,6349 | 24,746 | 24,7017 | 4 | 4 | 4 | 39,4 | 39,4 | 39,4 | 17,949 | 0 | 74,617 | 3,7E+08 | 21 |
| 17,416 | 17,4811 | 15,1706 | 14,8364 | 15,8299 | 16,1317 | 22,8242 | 22,558 | 20,4011 | 20,7196 | 20,459 | 20,9252 | 2 | 2 | 2 | 3,4 | 3,4 | 3,4 | 93,05 | 0 | 12,07 | 1,9E+07 | 3 |
| 16,1075 | 18,2101 | 18,4025 | 17,931 | 20,1009 | 18,4515 | 19,9894 | 21,7755 | 21,9268 | 22,1764 | 22,0727 | 21,7428 | 5 | 5 | 5 | 23,5 | 23,5 | 23,5 | 37,336 | 0 | 31,317 | 5,2E+07 | 7 |
| 14,5321 | 15,1571 | 16,5632 | 15,3495 | 18,1488 | 16,6061 | 20,4914 | 20,7553 | 21,3523 | 20,3873 | 21,412 | 21,4392 | 6 | 6 | 6 | 9,1 | 9,1 | 9,1 | 118,35 | 0 | 57,279 | 3,5E+07 | 11 |
| 13,8002 | 15,0128 | 14,8242 | 14,8425 | 18,0725 | 14,991 | 21,255 | 20,9893 | 20,5782 | 20,3415 | 21,5355 | 21,0632 | 4 | 4 | 4 | 7 | 7 | 7 | 115,69 | 0 | 25,791 | 1,9E+07 | 5 |
| 14,7833 | 15,2184 | 16,4705 | 15,1853 | 18,7922 | 16,9112 | 20,7731 | 20,0209 | 20,9579 | 20,7096 | 22,0177 | 21,157 | 2 | 2 | 2 | 4,6 | 4,6 | 4,6 | 80,798 | 0 | 15,8 | 2,3E+07 | 3 |
| 16,6709 | 18,0139 | 17,6943 | 17,6012 | 19,631 | 17,9739 | 20,869 | 21,7697 | 21,6344 | 21,7425 | 22,1602 | 21,9005 | 3 | 3 | 3 | 11,9 | 11,9 | 11,9 | 52,322 | 0 | 41,392 | 4,4E+07 | 11 |
| 14,1343 | 18,3039 | 19,0924 | 19,7556 | 20,5714 | 17,8184 | 20,8202 | 20,8527 | 21,2984 | 21,9029 | 21,4219 | 21,4191 | 2 | 2 | 2 | 18,4 | 18,4 | 18,4 | 18,078 | 0 | 158,97 | 2,5E+07 | 11 |
| 22,2238 | 23,619 | 23,9362 | 23,6894 | 25,5298 | 23,6483 | 24,7198 | 25,1053 | 25,3477 | 24,9869 | 25,5319 | 24,9638 | 9 | 9 | 9 | 67,2 | 67,2 | 67,2 | 15,245 | 0 | 213,76 | 7,6E+08 | 42 |
| 19,3085 | 20,7974 | 20,3786 | 20,28 | 22,2223 | 20,7598 | 23,2216 | 23,6192 | 23,2477 | 23,5337 | 23,8556 | 23,3619 | 6 | 6 | 6 | 19,9 | 19,9 | 19,9 | 55,843 | 0 | 52,316 | 2,2E+08 | 23 |
| 19,0306 | 20,2063 | 20,5344 | 20,313 | 22,2529 | 20,2412 | 23,1475 | 23,6268 | 23,6757 | 23,6455 | 23,7909 | 23,3641 | 6 | 6 | 6 | 32,6 | 32,6 | 32,6 | 34,446 | 0 | 122,91 | 2,2E+08 | 24 |
| 19,9412 | 20,5801 | 20,5098 | 21,2409 | 22,4651 | 20,0605 | 22,152 | 22,2129 | 21,7775 | 23,0506 | 22,7749 | 22,0996 | 5 | 5 | 5 | 15,9 | 15,9 | 15,9 | 24,075 | 0 | 33,965 | 1,2E+08 | 23 |
| 17,2355 | 18,6226 | 19,2829 | 20,1132 | 21,5104 | 18,0004 | 19,2545 | 22,719 | 23,0989 | 22,2438 | 22,7916 | 20,5957 | 2 | 2 | 2 | 14,6 | 14,6 | 14,6 | 28,824 | 0 | 47,136 | 6,7E+07 | 10 |
| 14,3056 | 18,0515 | 18,8114 | 18,6012 | 21,0395 | 17,9001 | 21,8662 | 22,2593 | 21,9077 | 22,4104 | 22,3999 | 22,1122 | 6 | 6 | 6 | 21,6 | 21,6 | 21,6 | 52,092 | 0 | 38,543 | 8,8E+07 | 8 |
| 17,5903 | 19,3239 | 18,058 | 19,0805 | 20,4647 | 18,4213 | 20,3488 | 22,1286 | 20,956 | 21,8005 | 21,8832 | 20,1232 | 3 | 3 | 3 | 15,8 | 15,8 | 15,8 | 34,017 | 0 | 20,802 | 3,5E+07 | 5 |
| 16,4212 | 18,0841 | 17,3764 | 17,0826 | 19,2254 | 18,5756 | 21,075 | 21,6966 | 21,7163 | 21,7323 | 22,6565 | 22,1438 | 7 | 7 | 7 | 10,6 | 10,6 | 10,6 | 72,992 | 0 | 45,327 | 5,5E+07 | 10 |
| 22,3759 | 22,842 | 23,2595 | 22,8155 | 24,608 | 23,2895 | 25,1339 | 25,1632 | 25,3572 | 25,1978 | 25,4937 | 25,4911 | 14 | 14 | 14 | 57,5 | 57,5 | 57,5 | 40,716 | 0 | 199,82 | 1,2E+09 | 67 |
| 14,3367 | 16,0106 | 17,9528 | 18,2255 | 20,2273 | 15,9331 | 20,8926 | 20,8037 | 21,8147 | 22,0868 | 22,7359 | 20,0999 | 5 | 4 | 4 | 15,8 | 13,2 | 13,2 | 58,583 | 0 | 45,607 | 3,8E+07 | 7 |
| 14,9922 | 16,6628 | 16,9479 | 16,9908 | 18,4744 | 16,3855 | 21,3897 | 20,957 | 21,8234 | 21,9543 | 22,0862 | 21,4235 | 2 | 2 | 2 | 4,2 | 4,2 | 4,2 | 101,84 | 0 | 21,606 | 2,6E+07 | 5 |
| 13,6015 | 17,3127 | 16,6344 | 16,8765 | 18,2249 | 16,9965 | 20,953 | 20,9554 | 20,604 | 21,7037 | 21,8331 | 20,9425 | 2 | 2 | 2 | 3,1 | 3,1 | 3,1 | 88,836 | 0 | 12,168 | 2,5E+07 | 2 |
| 18,7772 | 20,219 | 19,8009 | 19,7725 | 21,6375 | 20,0712 | 22,9793 | 23,8021 | 23,0976 | 23,3985 | 23,6545 | 23,3209 | 8 | 8 | 8 | 45,7 | 45,7 | 45,7 | 26,979 | 0 | 58,515 | 1,4E+08 | 28 |
| 20,4724 | 21,6348 | 21,7745 | 21,6424 | 23,2243 | 21,551 | 24,0014 | 24,5101 | 24,5561 | 24,4899 | 24,7365 | 24,5353 | 18 | 18 | 2 | 49,8 | 49,8 | 6,9 | 50,977 | 0 | 148,38 | 8,1E+08 | 47 |
| 15,1774 | 14,8515 | 15,1613 | 17,6097 | 18,2169 | 14,117 | 21,2107 | 20,3837 | 20,0151 | 21,6917 | 22,1097 | 20,9268 | 2 | 2 | 2 | 5,7 | 5,7 | 5,7 | 55,436 | 0 | 10,994 | 1,8E+07 | 2 |
| 19,1616 | 19,6525 | 20,4573 | 20,2742 | 21,8219 | 19,8176 | 23,5683 | 23,1575 | 23,5249 | 23,3786 | 23,3974 | 23,5284 | 8 | 8 | 8 | 25,5 | 25,5 | 25,5 | 66,615 | 0 | 64,516 | 2,7E+08 | 22 |
| 17,4714 | 20,2481 | 20,5993 | 20,4593 | 22,0427 | 20,0912 | 20,7049 | 22,2841 | 22,5399 | 22,6325 | 23,2755 | 22,0356 | 13 | 13 | 13 | 34 | 34 | 34 | 52,822 | 0 | 97,053 | 2,3E+08 | 32 |
| 18,6766 | 19,8565 | 20,4674 | 20,1694 | 21,935 | 19,9308 | 22,4431 | 22,7066 | 22,9457 | 22,6553 | 23,0599 | 22,5368 | 8 | 8 | 8 | 20,9 | 20,9 | 20,9 | 51,638 | 0 | 65,821 | 2E+08 | 25 |
| 24,3736 | 25,1451 | 25,4194 | 25,5257 | 26,8226 | 24,9402 | 26,7397 | 26,9731 | 27,101 | 27,2899 | 27,3259 | 26,8932 | 28 | 28 | 2 | 74,7 | 74,7 | 7,3 | 49,912 | 0 | 323,31 | 6,4E+09 | 172 |
| 16,7785 | 18,1477 | 19,5511 | 18,8 | 21,0098 | 18,7112 | 21,189 | 22,8223 | 22,5858 | 23,2992 | 22,9446 | 22,5593 | 7 | 7 | 7 | 27,2 | 27,2 | 27,2 | 51,021 | 0 | 44,392 | 1E+08 | 9 |
| 14,4082 | 16,7954 | 19,4924 | 18,7481 | 20,9219 | 17,5412 | 20,8924 | 20,8687 | 22,4636 | 22,3784 | 22,9999 | 20,886 | 4 | 4 | 4 | 17,5 | 17,5 | 17,5 | 54,591 | 0 | 31,858 | 6,2E+07 | 10 |
| 17,419 | 19,3825 | 19,6337 | 19,0809 | 21,42 | 19,6005 | 21,2253 | 23,0665 | 23,179 | 22,856 | 23,7108 | 23,1621 | 10 | 10 | 10 | 30,2 | 30,2 | 30,2 | 57,192 | 0 | 67,004 | 1,9E+08 | 22 |
| 17,9978 | 19,976 | 19,8221 | 19,3698 | 21,4772 | 20,1897 | 22,5359 | 23,0807 | 22,8187 | 22,4609 | 23,0833 | 22,8696 | 7 | 7 | 7 | 37,7 | 37,7 | 37,7 | 30,281 | 0 | 44,949 | 1,2E+08 | 12 |
| 13,851 | 14,9857 | 16,6484 | 15,6666 | 18,1835 | 15,844 | 21,3177 | 20,6485 | 20,8319 | 20,864 | 21,0887 | 20,852 | 5 | 5 | 5 | 9,8 | 9,8 | 9,8 | 93,957 | 0 | 92,85 | 2,2E+07 | 10 |
| 17,87 | 17,963 | 17,565 | 17,8852 | 19,5208 | 17,9827 | 20,7275 | 20,4126 | 19,7777 | 21,5159 | 21,728 | 20,9447 | 3 | 3 | 3 | 20,3 | 20,3 | 20,3 | 25,865 | 0 | 19,724 | 2,9E+07 | 9 |
| 16,307 | 13,7721 | 16,9622 | 16,0138 | 18,5611 | 16,9532 | 20,2362 | 20,3941 | 21,2992 | 20,1018 | 20,4006 | 20,5129 | 2 | 2 | 2 | 10,2 | 10,2 | 10,2 | 28,968 | 0 | 57,042 | 1,4E+07 | 5 |
| 16,8135 | 15,2712 | 18,781 | 17,7642 | 21,3449 | 17,5778 | 21,8798 | 20,5524 | 22,3374 | 20,6724 | 23,1644 | 20,921 | 2 | 2 | 2 | 6,6 | 6,6 | 6,6 | 55,267 | 0 | 14,568 | 5,8E+07 | 5 |
| 14,464 | 20,2852 | 21,013 | 20,6715 | 22,7133 | 20,4081 | 21,1542 | 22,1136 | 22,3478 | 22,7957 | 23,1481 | 22,533 | 3 | 3 | 3 | 20 | 20 | 20 | 12,362 | 0 | 18,37 | 6,7E+07 | 6 |
| 13,8526 | 15,0147 | 14,5875 | 15,2216 | 18,0307 | 14,4025 | 21,4129 | 20,901 | 20,6153 | 20,8199 | 21,5833 | 20,9703 | 4 | 4 | 4 | 4,1 | 4,1 | 4,1 | 153,61 | 0 | 24,982 | 2,6E+07 | 4 |
| 15,6151 | 19,0978 | 19,7875 | 19,795 | 21,297 | 18,7982 | 21,2932 | 22,5665 | 22,886 | 22,7635 | 23,1495 | 20,5852 | 3 | 3 | 3 | 21 | 21 | 21 | 21,565 | 0 | 21,479 | 7E+07 | 8 |
| 13,7108 | 18,4633 | 18,5661 | 18,4574 | 19,5527 | 18,6738 | 20,3604 | 20,8368 | 20,8284 | 20,9846 | 20,6028 | 20,0758 | 2 | 2 | 2 | 19,3 | 19,3 | 19,3 | 21,954 | 0 | 12,093 | 1,8E+07 | 4 |
| 13,6802 | 18,1523 | 20,878 | 21,3055 | 22,3365 | 18,0227 | 20,7855 | 20,4733 | 23,6483 | 23,8169 | 23,266 | 19,6996 | 2 | 2 | 2 | 15,2 | 15,2 | 15,2 | 24,498 | 0 | 28,488 | 7,3E+07 | 10 |

| | | | | | | | | | | | | | | | | | | | | | | | | | | | | | | | | | | | |
|---|---|---|---|---|---|---|---|---|---|---|---|---|---|---|---|---|---|---|---|---|---|---|---|---|---|---|---|---|---|---|---|---|---|---|---|
| 17,2805 | 18,5242 | 18,8728 | 18,9456 | 20,8957 | 17,3573 | 22,3087 | 22,4479 | 22,2297 | 22,2185 | 22,7342 | 22,3453 | 7 | 7 | 7 | 13,8 | 13,8 | 13,8 | 75,843 | 0 | 48,754 | 1,2E+08 | 13 | 3 | 5 | 5 | 7 | 7 | 3 | 0,30196 | 0,71906 | -0,8404 | -0,7427 | DIPPA_06 | DIPPA_06 | DIPPA_06919.mRNA.1 hypothetical protein |
| 13,1129 | 19,4696 | 17,9918 | 17,3725 | 17,6698 | 20,2674 | 20,905 | 20,5965 | 20,5277 | 20,7311 | 20,6383 | 21,3537 | 2 | 2 | 2 | 11,2 | 11,2 | 11,2 | 25,812 | 0 | 12,792 | 2,4E+07 | 3 | 0 | 1 | 1 | 1 | 1 | 1 | 0,30132 | 0,71941 | -1,5785 | -0,7413 | DIPPA_34 | DIPPA_34 | DIPPA_34567.mRNA.1 hypothetical protein |
| 18,6641 | 19,935 | 20,3591 | 19,2765 | 21,8078 | 19,9055 | 21,2901 | 22,3326 | 22,4552 | 21,8189 | 23,0233 | 21,2118 | 2 | 2 | 2 | 23,8 | 23,8 | 23,8 | 11,478 | 0 | 86,559 | 4,8E+07 | 6 | 1 | 2 | 2 | 2 | 2 | 1 | 0,30052 | 0,71933 | -0,6772 | -0,7396 | DIPPA_63 | DIPPA_63 | DIPPA_63332.mRNA.1 Glycine-rich protein 2, putative n=1 Tax=Eimeria praecox RepID=U6H6B5_9EIME (length=251) |
| 19,9567 | 20,8597 | 21,5302 | 21,028 | 22,8239 | 20,3963 | 23,2364 | 24,0102 | 23,873 | 23,6432 | 24,0492 | 23,2826 | 14 | 14 | 14 | 63,9 | 63,9 | 63,9 | 38,054 | 0 | 164,48 | 3,5E+08 | 33 | 6 | 7 | 12 | 11 | 12 | 7 | 0,30005 | 0,71861 | -0,6339 | -0,7386 | DIPPA_29 | DIPPA_29 | DIPPA_29380.mRNA.1 Flagellar Member 1 |
| 20,3034 | 21,0574 | 21,0279 | 20,4765 | 22,3482 | 20,9269 | 22,8534 | 23,3801 | 23,0266 | 22,7414 | 23,4 | 22,7834 | 4 | 4 | 4 | 47,3 | 47,3 | 47,3 | 16,743 | 0 | 74,949 | 1E+08 | 19 | 2 | 2 | 3 | 2 | 3 | 3 | 0,29978 | 0,7177 | -0,4543 | -0,738 | DIPPA_31 | DIPPA_31 | DIPPA_31883.mRNA.1 Ubiquitin-conjugating enzyme E2-16 kDa |
| 17,895 | 18,6342 | 18,5909 | 17,9618 | 20,1229 | 18,5573 | 21,0585 | 23,1782 | 20,7886 | 22,6314 | 23,6515 | 22,9831 | 19 | 2 | 2 | 49,7 | 6,9 | 6,9 | 57,288 | 0 | 122,7 | 7,4E+07 | 7 | 1 | 2 | 1 | 2 | 2 | 2 | 0,29968 | 0,71665 | -0,5073 | -0,7378 | DIPPA_10 | DIPPA_10 | DIPPA_10954.mRNA.2 Aldehyde dehydrogenase family 2 member B4 mitochondrial;DIPPA_10954.mRNA.5 Aldehyde dehydrogenase family 2 member B4 mitochondrial;DIPPA_10954.mRNA.1 Putative aldehyde dehydroge[illegible] |
| 17,1248 | 16,9023 | 18,2195 | 17,3299 | 18,2877 | 17,7183 | 20,3126 | 20,4336 | 20,2229 | 20,4317 | 20,5225 | 20,7868 | 2 | 2 | 2 | 8,2 | 8,2 | 8,2 | 37,002 | 0 | 12,083 | 2,3E+07 | 3 | 1 | 1 | 2 | 1 | 1 | 1 | 0,29902 | 0,7166 | -0,3631 | -0,7364 | DIPPA_17 | DIPPA_17 | DIPPA_17507.mRNA.1 hypothetical protein |
| 16,8931 | 19,9706 | 20,2589 | 19,4206 | 22,1887 | 18,8 | 21,1253 | 22,9034 | 22,8008 | 22,6764 | 23,317 | 22,3782 | 8 | 8 | 8 | 36,3 | 36,3 | 36,3 | 36,526 | 0 | 270,16 | 1,3E+08 | 15 | 1 | 4 | 7 | 6 | 8 | 3 | 0,29654 | 0,71809 | -1,0956 | -0,7312 | DIPPA_07 | DIPPA_07 | DIPPA_07446.mRNA.1 hypothetical protein Tco025E_06130 |
| 14,2303 | 18,0594 | 17,4129 | 16,0776 | 18,357 | 18,3477 | 21,3211 | 20,7488 | 21,3606 | 20,6977 | 20,8585 | 20,4212 | 2 | 2 | 2 | 6,4 | 6,4 | 6,4 | 61,22 | 0 | 18,77 | 3,3E+07 | 5 | 0 | 1 | 2 | 2 | 1 | 2 | 0,29618 | 0,71766 | -1,0266 | -0,7304 | DIPPA_11 | DIPPA_11 | DIPPA_11651.mRNA.1 hypothetical protein |
| 14,3609 | 14,1379 | 15,1646 | 14,1959 | 17,3521 | 14,437 | 20,2329 | 20,9332 | 19,978 | 20,6404 | 20,2862 | 20,3859 | 7 | 7 | 7 | 9,4 | 9,4 | 9,4 | 117,63 | 0 | 52,87 | 1,4E+07 | 8 | 1 | 1 | 2 | 1 | 7 | 1 | 0,29566 | 0,71746 | -0,7739 | -0,7293 | DIPPA_22 | DIPPA_22 | DIPPA_22579.mRNA.1 Importin subunit beta-3 |
| 20,0668 | 20,1025 | 17,892 | 19,385 | 20,6515 | 19,818 | 23,268 | 22,6857 | 20,7038 | 21,6781 | 22,8654 | 22,1882 | 6 | 6 | 6 | 47,6 | 47,6 | 47,6 | 18,829 | 0 | 52,478 | 6,3E+07 | 13 | 4 | 4 | 2 | 4 | 4 | 4 | 0,29548 | 0,71674 | -0,5977 | -0,7289 | DIPPA_31 | DIPPA_31 | DIPPA_31393.mRNA.1 Ran-binding protein 1 homolog b |
| 18,0174 | 17,701 | 16,244 | 17,5966 | 19,3306 | 16,9817 | 20,3496 | 21,8289 | 20,7561 | 21,7529 | 20,1975 | 20,6881 | 4 | 4 | 4 | 17 | 17 | 17 | 38,791 | 0 | 27,604 | 3,2E+07 | 6 | 2 | 2 | 1 | 2 | 3 | 1 | 0,29537 | 0,71606 | -0,6488 | -0,7287 | DIPPA_23 | DIPPA_23 | DIPPA_23019.mRNA.1 Glutathione S-transferase 1 isoform C |
| 16,7662 | 18,9386 | 20,134 | 19,8804 | 21,4827 | 17,6924 | 22,6988 | 22,5804 | 23,1122 | 23,0094 | 23,4886 | 22,8634 | 3 | 3 | 3 | 15 | 15 | 15 | 25,016 | 0 | 16,77 | 8,8E+07 | 4 | 2 | 3 | 3 | 3 | 3 | 2 | 0,29434 | 0,71583 | -1,0723 | -0,7265 | DIPPA_26 | DIPPA_26 | DIPPA_26684.mRNA.1 60S ribosomal protein L13 |
| 17,9044 | 18,6106 | 20,6262 | 19,3651 | 21,6604 | 18,7233 | 22,7639 | 23,4068 | 23,3415 | 23,5686 | 23,8606 | 22,9818 | 6 | 6 | 6 | 20,5 | 20,5 | 20,5 | 48,496 | 0 | 71,122 | 1,7E+08 | 16 | 3 | 2 | 4 | 4 | 6 | 3 | 0,291 | 0,71915 | -0,8692 | -0,7194 | DIPPA_16 | DIPPA_16 | DIPPA_16986.mRNA.1 CBL-interacting protein kinase 23 |
| 22,112 | 22,1418 | 20,175 | 20,3605 | 18,5905 | 22,6079 | 24,699 | 24,0676 | 21,3317 | 22,3786 | 19,7444 | 24,1283 | 3 | 3 | 3 | 12,4 | 12,4 | 12,4 | 18,238 | 0 | 30,651 | 1,3E+08 | 13 | 3 | 3 | 3 | 3 | 3 | 3 | 0,29039 | 0,71971 | 0,95663 | 0,7181 | DIPPA_34 | DIPPA_34 | DIPPA_34733.mRNA.1 hypothetical protein |
| 21,2305 | 22,1449 | 20,6817 | 21,4581 | 21,081 | 23,1858 | 26,4044 | 26,2398 | 24,4628 | 24,7855 | 23,65 | 27,0351 | 2 | 2 | 2 | 5 | 5 | 5 | 51,031 | 0 | 15,301 | 4,7E+08 | 10 | 2 | 2 | 2 | 2 | 2 | 2 | 0,28963 | 0,71946 | -0,5559 | -0,7165 | DIPPA_14 | DIPPA_14 | DIPPA_14510.mRNA.1 L-rhamnonate dehydratase |
| 21,0527 | 22,1043 | 22,3736 | 21,8514 | 23,6671 | 21,6382 | 24,9557 | 25,1106 | 25,2411 | 25,1331 | 25,2522 | 24,7642 | 38 | 38 | 38 | 48,7 | 48,7 | 48,7 | 116,42 | 0 | 323,31 | 1,7E+09 | 137 | 11 | 28 | 30 | 29 | 35 | 21 | 0,28832 | 0,7208 | -0,542 | -0,7137 | DIPPA_32 | DIPPA_32 | DIPPA_32821.mRNA.1 Myosin-I heavy chain |
| 14,2246 | 17,5299 | 18,6361 | 16,3685 | 19,7042 | 17,8264 | 21,0897 | 21,6196 | 22,0194 | 21,4027 | 22,3803 | 21,7727 | 4 | 4 | 4 | 10,3 | 10,3 | 10,3 | 58,52 | 0 | 25,561 | 4,6E+07 | 5 | 0 | 3 | 4 | 2 | 4 | 2 | 0,28813 | 0,71989 | -1,1695 | -0,7133 | DIPPA_33 | DIPPA_33 | DIPPA_33179.mRNA.1 hypothetical protein |
| 19,3101 | 20,2126 | 21,3392 | 20,1999 | 22,1773 | 20,3261 | 22,8734 | 23,1187 | 23,5077 | 22,8669 | 23,1785 | 23,2054 | 11 | 11 | 11 | 35,9 | 35,9 | 35,9 | 46,031 | 0 | 150,55 | 2,2E+08 | 18 | 5 | 7 | 10 | 8 | 11 | 7 | 0,28534 | 0,72248 | -0,6138 | -0,7073 | DIPPA_05 | DIPPA_05 | DIPPA_05630.mRNA.1 Fumarylacetoacetase |
| 19,8881 | 18,9086 | 19,1045 | 18,6052 | 19,8079 | 18,3629 | 22,8778 | 20,174 | 20,4051 | 20,9451 | 19,8648 | 20,274 | 4 | 4 | 4 | 21,2 | 21,2 | 21,2 | 29,317 | 0 | 26,276 | 2,9E+07 | 4 | 3 | 2 | 3 | 2 | 3 | 1 | 0,28073 | 0,72926 | 0,37505 | 0,69746 | DIPPA_29 | DIPPA_29 | DIPPA_29193.mRNA.1 translocon-associated protein subunit gamma |
| 13,6218 | 17,1473 | 18,5214 | 17,1985 | 19,4937 | 16,2548 | 20,3648 | 20,6912 | 22,9468 | 21,3403 | 22,4692 | 21,1161 | 4 | 4 | 4 | 12,6 | 12,6 | 12,6 | 62,26 | 0 | 36,124 | 4,6E+07 | 3 | 0 | 2 | 3 | 1 | 4 | 1 | 0,28072 | 0,72822 | -1,2189 | -0,6975 | DIPPA_31 | DIPPA_31 | DIPPA_31238.mRNA.1 Eukaryotic translation initiation factor 3 subunit L |
| 14,9666 | 16,7326 | 16,5516 | 16,8646 | 19,4787 | 14,8566 | 20,8023 | 20,9779 | 20,1998 | 20,9217 | 21,8241 | 20,7385 | 2 | 2 | 2 | 14,4 | 14,4 | 14,4 | 26,953 | 0 | 11,84 | 1,6E+07 | 2 | 0 | 1 | 1 | 2 | 2 | 0 | 0,27144 | 0,74124 | -0,983 | -0,6775 | DIPPA_17 | DIPPA_17 | DIPPA_17612.mRNA.1 Adenylate kinase 1 chloroplastic |
| 13,6872 | 14,9168 | 15,7609 | 14,9451 | 16,5115 | 14,6081 | 20,6478 | 20,139 | 20,7138 | 20,5336 | 20,9306 | 20,979 | 2 | 2 | 2 | 4,5 | 4,5 | 4,5 | 63,233 | 0 | 11,812 | 6475700 | 3 | 0 | 1 | 1 | 0 | 2 | 0 | 0,26988 | 0,74259 | -0,5666 | -0,6741 | DIPPA_31 | DIPPA_31 | DIPPA_31694.mRNA.1 BTB/POZ domain-containing protein At2g30600 |
| 16,9629 | 19,4087 | 18,8684 | 17,9854 | 20,112 | 19,0757 | 21,1391 | 22,9818 | 20,5078 | 22,033 | 22,3744 | 22,653 | 4 | 4 | 4 | 15,4 | 15,4 | 15,4 | 49,366 | 0 | 54,17 | 6,5E+07 | 11 | 1 | 2 | 1 | 2 | 4 | 2 | 0,26761 | 0,74436 | -0,6444 | -0,6692 | DIPPA_28 | DIPPA_28 | DIPPA_28109.mRNA.1 hypothetical protein |
| 15,2185 | 15,6041 | 15,9667 | 15,6728 | 17,914 | 15,0156 | 20,1407 | 20,2429 | 20,4342 | 20,493 | 20,6242 | 20,028 | 2 | 2 | 2 | 5,5 | 5,5 | 5,5 | 82,182 | 0 | 12,089 | 1,7E+07 | 3 | 0 | 1 | 1 | 1 | 2 | 0 | 0,26744 | 0,74345 | -0,6043 | -0,6689 | DIPPA_01 | DIPPA_01 | DIPPA_01194.mRNA.1 Prolyl endopeptidase |
| 18,2615 | 18,691 | 20,3087 | 18,8773 | 21,2725 | 19,0838 | 22,6298 | 22,5583 | 23,0251 | 22,7174 | 23,1042 | 22,7194 | 4 | 4 | 4 | 21,1 | 21,1 | 21,1 | 42,155 | 0 | 66,798 | 1,1E+08 | 12 | 2 | 2 | 3 | 3 | 4 | 3 | 0,26593 | 0,74477 | -0,6575 | -0,6656 | DIPPA_14 | DIPPA_14 | DIPPA_14619.mRNA.1 hypothetical protein |
| 16,8877 | 15,9358 | 15,6457 | 16,0451 | 17,9492 | 15,9687 | 19,9291 | 20,3305 | 20,5647 | 20,0818 | 19,8622 | 20,1434 | 2 | 2 | 2 | 5,4 | 5,4 | 5,4 | 51,52 | 0 | 13,394 | 1,4E+07 | 6 | 1 | 1 | 1 | 1 | 2 | 1 | 0,26573 | 0,74448 | -0,4979 | -0,6652 | DIPPA_25 | DIPPA_25 | DIPPA_25262.mRNA.1 hypothetical protein |
| 21,1048 | 20,8626 | 18,1784 | 19,4994 | 21,1852 | 21,7404 | 25,5259 | 25,7039 | 24,7612 | 24,7911 | 24,0759 | 25,1299 | 7 | 7 | 7 | 25,9 | 25,9 | 25,9 | 45,047 | 0 | 48,054 | 2,4E+08 | 10 | 2 | 2 | 3 | 4 | 7 | 3 | 0,26242 | 0,74693 | -0,7598 | -0,658 | DIPPA_03 | DIPPA_03 | DIPPA_03248.mRNA.1 hypothetical protein |
| 17,5391 | 19,568 | 19,35 | 18,6259 | 20,0558 | 19,2777 | 23,8243 | 23,612 | 22,7298 | 22,2402 | 21,9545 | 23,1961 | 4 | 4 | 4 | 10,1 | 10,1 | 10,1 | 63,631 | 0 | 26,129 | 1,2E+08 | 16 | 1 | 4 | 4 | 3 | 4 | 3 | 0,26107 | 0,74841 | -0,5007 | -0,6551 | DIPPA_04 | DIPPA_04 | DIPPA_04140.mRNA.1 Polypyrimidine tract-binding protein homolog 3 |
| 15,3951 | 15,3237 | 16,0258 | 15,5448 | 18,6045 | 14,8865 | 20,9124 | 19,2546 | 19,9794 | 19,8123 | 20,6307 | 21,7574 | 3 | 3 | 3 | 6,7 | 6,7 | 6,7 | 47,053 | 0 | 17,532 | 1,3E+07 | 3 | 1 | 2 | 2 | 2 | 3 | 1 | 0,26078 | 0,74748 | -0,7637 | -0,6544 | DIPPA_25 | DIPPA_25 | DIPPA_25074.mRNA.1 Saccharopine dehydrogenase |
| 15,5452 | 16,1709 | 17,0155 | 16,4364 | 17,8621 | 15,8662 | 20,2854 | 20,7178 | 22,0339 | 21,5963 | 21,5839 | 20,8068 | 3 | 3 | 3 | 6,1 | 6,1 | 6,1 | 94,019 | 0 | 19,209 | 2,7E+07 | 6 | 1 | 1 | 2 | 2 | 3 | 1 | 0,26048 | 0,74721 | -0,4777 | -0,6538 | DIPPA_27 | DIPPA_27 | DIPPA_27371.mRNA.1 Poly polymerase |
| 16,7606 | 15,4692 | 16,4276 | 16,0458 | 18,3048 | 15,9783 | 23,0362 | 22,4657 | 23,159 | 22,7169 | 22,9341 | 22,925 | 16 | 16 | 5 | 7,3 | 7,3 | 1,9 | 445,94 | 0 | 118,53 | 1,4E+08 | 22 | 2 | 1 | 3 | 0 | 5 | 3 | 0,25882 | 0,75011 | -0,5572 | -0,6502 | DIPPA_23 | DIPPA_23 | DIPPA_23635.mRNA.1 Fatty acid synthase subunit beta (FAS I) |
| [illegible] | [illegible] | [illegible] | [illegible] | [illegible] | [illegible] | [illegible] | [illegible] | [illegible] | [illegible] | [illegible] | [illegible] | [illegible] | [illegible] | [illegible] | [illegible] | [illegible] | [illegible] | [illegible] | [illegible] | [illegible] | [illegible] | [illegible] | [illegible] | [illegible] | [illegible] | [illegible] | [illegible] | [illegible] | [illegible] | [illegible] | [illegible] | [illegible] | [illegible] | [illegible] | [illegible] |

| iBAQ Sec | iBAQ Sec | iBAQ Sec | iBAQ wt | iBAQ wt | iBAQ wt | LFQ inten | LFQ inten | LFQ inten | LFQ inten | LFQ inten | LFQ inten | Student's | Student's | Peptides | Razor + u | Unique p | Sequence | Unique + | Unique se | Mol. weig | Q-value | Score | Intensity | MS/MS c | Unique p | Unique p | Unique p | Unique p | Unique p | Unique p | -Log Stud | Student's | Student's | Student's | Protein ID | Majority | Fasta headers |
|---|---|---|---|---|---|---|---|---|---|---|---|---|---|---|---|---|---|---|---|---|---|---|---|---|---|---|---|---|---|---|---|---|---|---|---|---|---|
| 26,5346 | 26,7905 | 26,3272 | 16,3788 | 13,3829 | 14,1822 | 27,6625 | 27,9845 | 27,3588 | 19,238 | 18,2532 | 18,6265 | + | iBAQ Sec | 17 | 17 | 17 | 74,2 | 74,2 | 74,2 | 34,042 | 0 | 219,79 | 5,3E+09 | 105 | 16 | 17 | 16 | 1 | 0 | 0 | 3,71328 | 0 | 11,9028 | 13,1436 | DIPPA_3 | DIPPA_3 | DIPPA_35934.mRNA.1 Protein transport protein SEC13 |
| 23,3137 | 23,4427 | 22,7879 | 13,4563 | 11,8759 | 13,2234 | 26,1161 | 26,3179 | 25,5481 | 18,7799 | 17,8857 | 19,2622 | + | iBAQ Sec | 26 | 26 | 26 | 20,7 | 20,7 | 20,7 | 193,19 | 0 | 260,45 | 2,1E+09 | 117 | 25 | 25 | 24 | 0 | 0 | 0 | 4,38295 | 0 | 10,3296 | 19,4253 | DIPPA_1 | DIPPA_1 | DIPPA_10282.mRNA.1 hypothetical protein |
| 21,8424 | 20,5642 | 21,3602 | 12,6939 | 12,5136 | 18,2734 | 23,2833 | 17,7597 | 22,5995 | 18,8241 | 18,2429 | 19,1552 | + | iBAQ Sec | 2 | 2 | 2 | 20,3 | 20,3 | 20,3 | 6,8619 | 0 | 14,171 | 4E+07 | 2 | 2 | 1 | 2 | 0 | 0 | 1 | 1,60745 | 0,04153 | 6,76197 | 3,50912 | DIPPA_2 | DIPPA_2 | DIPPA_24015.mRNA.1 hypothetical protein |
| 19,4339 | 19,2277 | 19,3456 | 13,2345 | 11,219 | 13,6223 | 23,8909 | 18,3746 | 23,7781 | 19,7509 | 18,7297 | 18,9502 | + | iBAQ Sec | 4 | 4 | 4 | 11,1 | 11,1 | 11,1 | 72,471 | 0 | 24,767 | 5,9E+07 | 4 | 4 | 1 | 3 | 1 | 0 | 1 | 3,05329 | 0,00886 | 6,64379 | 8,89031 | DIPPA_3 | DIPPA_3 | [illegible] |
| 19,0806 | 19,2681 | 18,5651 | 13,6565 | 11,0527 | 13,2043 | 22,3746 | 22,6938 | 22,2619 | 19,8375 | 17,894 | 18,4204 | + | iBAQ Sec | 12 | 12 | 12 | 22,8 | 22,8 | 22,8 | 95,466 | 0 | 99,857 | 1,2E+08 | 24 | 10 | 12 | 7 | 0 | 0 | 0 | 2,79968 | 0,00586 | 6,33344 | 7,62801 | DIPPA_0 | DIPPA_0 | DIPPA_06252.mRNA.1 hypothetical protein;TR85149__c0_g1_i1__m.17174 #MITO [illegible] |
| 17,9365 | 18,1826 | 17,7803 | 11,8626 | 12,7313 | 13,3135 | 20,876 | 18,6148 | 20,4053 | 19,8332 | 18,4548 | 18,5398 | + | iBAQ Sec | 2 | 2 | 2 | 20,8 | 20,8 | 20,8 | 19,893 | 0 | 13,092 | 1,4E+07 | 3 | 2 | 2 | 2 | 0 | 0 | 0 | 3,5843 | 0 | 5,33073 | 12,1842 | DIPPA_1 | DIPPA_1 | DIPPA_12427.mRNA.1 hypothetical protein;DIPPA_12427.mRNA.2 hypothetical prote[illegible] |
| 18,8155 | 19,0654 | 19,0648 | 13,0611 | 13,5275 | 14,8799 | 21,714 | 22,0981 | 21,991 | 18,9309 | 19,3562 | 17,8326 | + | iBAQ Sec | 2 | 2 | 2 | 19,7 | 19,7 | 19,7 | 25,812 | 0 | 32,472 | 1,8E+07 | 9 | 2 | 2 | 2 | 0 | 0 | 1 | 3,13767 | 0,01058 | 5,15907 | 9,35088 | DIPPA_3 | DIPPA_3 | DIPPA_34567.mRNA.1 hypothetical protein |
| 18,5807 | 14,3937 | 16,3933 | 10,5417 | 11,6102 | 12,3783 | 18,8984 | 19,325 | 19,1979 | 18,742 | 18,5504 | 19,0005 | + | iBAQ Sec | 2 | 2 | 2 | 24,7 | 24,7 | 24,7 | 16,413 | 0 | 18,088 | 3346800 | 2 | 2 | 0 | 1 | 0 | 0 | 0 | 1,69768 | 0,03511 | 4,94583 | 3,74353 | DIPPA_0 | DIPPA_0 | DIPPA_05112.mRNA.1 hypothetical protein;DIPPA_12878.mRNA.1 hypothetical prote[illegible] |
| 18,6917 | 17,9545 | 18,7745 | 11,7366 | 14,2266 | 15,7745 | 20,656 | 20,6808 | 20,8576 | 19,1961 | 18,364 | 18,3897 | + | iBAQ Sec | 4 | 4 | 4 | 28,4 | 28,4 | 28,4 | 28,731 | 0 | 26,363 | 2,1E+07 | 10 | 3 | 3 | 4 | 0 | 1 | 1 | 1,71365 | 0,03429 | 4,56101 | 3,78602 | DIPPA_1 | DIPPA_1 | DIPPA_19789.mRNA.1 NADP-dependent 3-hydroxy acid dehydrogenase YdfG;DIPPA_[illegible] |
| 17,4472 | 17,1016 | 17,8802 | 12,6185 | 11,3356 | 15,4431 | 20,4961 | 19,8285 | 20,8812 | 19,1698 | 18,3358 | 17,9523 | + | iBAQ Sec | 2 | 2 | 2 | 13,6 | 13,6 | 13,6 | 33,986 | 0 | 11,88 | 1,4E+07 | 5 | 2 | 1 | 2 | 0 | 0 | 1 | 1,61182 | 0,04171 | 4,34396 | 3,52025 | DIPPA_0 | DIPPA_0 | DIPPA_00094.mRNA.1 hypothetical protein GUITHDRAFT_154440 |
| 17,291 | 15,3497 | 16,9773 | 12,6889 | 11,9876 | 12,0343 | 20,6867 | 18,9328 | 20,1304 | 18,4061 | 19,0855 | 19,073 | + | iBAQ Sec | 2 | 2 | 2 | 13 | 13 | 13 | 35,846 | 0 | 12,688 | 4305100 | 2 | 2 | 1 | 2 | 0 | 0 | 0 | 2,58629 | 0,00633 | 4,30246 | 6,69275 | DIPPA_1 | DIPPA_1 | DIPPA_10368.mRNA.1 ATP-dependent protease subunit HslV |
| 16,511 | 15,8785 | 16,7125 | 12,9509 | 11,8639 | 11,7748 | 20,2261 | 18,7217 | 19,8309 | 18,597 | 19,3712 | 19,2653 | + | iBAQ Sec | 2 | 2 | 2 | 11,8 | 11,8 | 11,8 | 40,36 | 0,00144 | 11,161 | 6174400 | 2 | 2 | 1 | 1 | 0 | 0 | 0 | 3,10824 | 0,00965 | 4,17078 | 9,18779 | DIPPA_3 | DIPPA_3 | DIPPA_30206.mRNA.1 Nucleoporin SEH1 |
| 17,9696 | 17,5034 | 17,2615 | 12,8545 | 12,6256 | 15,0225 | 21,4791 | 21,4302 | 20,8242 | 19,5544 | 18,5738 | 18,6063 | + | iBAQ Sec | 10 | 10 | 10 | 17,7 | 17,7 | 17,7 | 119,14 | 0 | 80,81 | 4,6E+07 | 21 | 7 | 6 | 7 | 0 | 0 | 0 | 2,1716 | 0,02336 | 4,07729 | 5,15181 | TR10072 | TR10072 | TR100720__c0_g1_i1__m.22092 #MITO 2 TargetP,Mitoprot #PFAM PF13639.1 zf-RIN[illegible] |
| 17,7798 | 17,4313 | 17,8753 | 13,2492 | 13,8309 | 14,0624 | 21,4069 | 20,7685 | 21,2183 | 18,5071 | 19,131 | 19,592 | + | iBAQ Sec | 7 | 7 | 7 | 17,5 | 17,5 | 17,5 | 87,485 | 0 | 51,956 | 2,8E+07 | 16 | 5 | 6 | 5 | 1 | 1 | 1 | 3,8661 | 0 | 3,98125 | 14,3744 | DIPPA_0 | DIPPA_0 | DIPPA_00126.mRNA.1 Alpha,alpha-trehalose-phosphate synthase [UDP-forming] A |
| 16,652 | 16,9318 | 16,3458 | 12,7601 | 13,0398 | 12,9518 | 19,0267 | 21,5232 | 20,7583 | 18,3191 | 18,6798 | 18,848 | + | iBAQ Sec | 2 | 2 | 2 | 6,2 | 6,2 | 6,2 | 87,316 | 0 | 14,697 | 1,3E+07 | 2 | 2 | 2 | 2 | 0 | 0 | 0 | 4,41479 | 0 | 3,72599 | 19,7878 | DIPPA_0 | DIPPA_0 | DIPPA_03259.mRNA.1 hypothetical protein |
| 15,6626 | 15,8863 | 16,2649 | 13,4497 | 11,4245 | 11,9475 | 20,1909 | 20,2418 | 21,0529 | 19,211 | 18,4629 | 18,5222 | + | iBAQ Sec | 5 | 5 | 5 | 8,4 | 8,4 | 8,4 | 123,35 | 0 | 34,265 | 1,3E+07 | 7 | 3 | 3 | 4 | 2 | 1 | 1 | 2,35666 | 0,01263 | 3,66401 | 5,79847 | DIPPA_6 | DIPPA_6 | DIPPA_63080.mRNA.1 Predicted protein n=1 Tax=Phaeodactylum tricornutum (strain[illegible] |
| 18,9315 | 18,4257 | 18,4053 | 16,2704 | 12,3058 | 16,4241 | 21,2489 | 20,4854 | 21,4922 | 20,0881 | 17,7357 | 20,2031 | | | 2 | 2 | 2 | 11,2 | 11,2 | 11,2 | 20,532 | 0 | 12,94 | 1,3E+07 | 5 | 2 | 2 | 2 | 2 | 0 | 2 | 1,23977 | 0,07919 | 3,58739 | 2,64009 | DIPPA_2 | DIPPA_2 | DIPPA_23289.mRNA.1 Flagellar calcium-binding protein;DIPPA_23254.mRNA.1 Flage[illegible] |
| 19,1056 | 18,795 | 18,9289 | 15,178 | 15,0723 | 16,0547 | 23,3727 | 23,1479 | 22,0991 | 21,2518 | 21,1368 | 21,6932 | + | iBAQ Sec | 4 | 4 | 4 | 7,1 | 7,1 | 7,1 | 67,638 | 0 | 26,818 | 6,6E+07 | 11 | 4 | 4 | 4 | 1 | 2 | 3 | 3,38387 | 0,0062 | 3,50819 | 10,825 | DIPPA_0 | DIPPA_0 | DIPPA_01837.mRNA.1 Chaperone protein DnaK |
| 15,9426 | 15,9289 | 17,0557 | 13,1891 | 11,4359 | 13,8534 | 20,4868 | 20,6333 | 20,9305 | 19,3007 | 18,5182 | 19,9586 | + | iBAQ Sec | 5 | 5 | 5 | 11,2 | 11,2 | 11,2 | 95,321 | 0 | 38,616 | 1,5E+07 | 9 | 3 | 2 | 5 | 0 | 1 | 2 | 1,89438 | 0,02844 | 3,48289 | 4,28931 | DIPPA_1 | DIPPA_1 | DIPPA_11662.mRNA.1 Coatomer subunit gamma-2 |
| 16,2101 | 16,6014 | 14,4812 | 11,6556 | 13,9733 | 11,2744 | 20,0894 | 21,4495 | 19,553 | 19,8815 | 19,5064 | 19,4274 | + | iBAQ Sec | 6 | 6 | 6 | 6,1 | 6,1 | 6,1 | 205,74 | 0 | 50,828 | 2,9E+07 | 16 | 6 | 5 | 3 | 1 | 3 | 1 | 1,50336 | 0,0496 | 3,46314 | 3,2499 | DIPPA_5 | DIPPA_5 | DIPPA_53800.mRNA.1 PREDICTED: uncharacterized protein LOC103934149 isoform X[illegible] |
| 16,3596 | 15,7616 | 16,185 | 14,0921 | 11,7521 | 12,2575 | 20,99 | 20,696 | 20,7248 | 17,5987 | 18,2952 | 19,2641 | + | iBAQ Sec | 2 | 2 | 2 | 3,6 | 3,6 | 3,6 | 94,424 | 0 | 30,863 | 9733200 | 7 | 1 | 1 | 1 | 0 | 1 | 2 | 2,0124 | 0,02499 | 3,40149 | 4,64212 | DIPPA_0 | DIPPA_0 | DIPPA_05264.mRNA.1 Importin subunit beta-1;TR129715__c0_g3_i1__m.31138 #MI[illegible] |
| 17,0205 | 16,3701 | 16,9729 | 14,3304 | 12,6752 | 13,1852 | 20,6041 | 20,0344 | 20,5977 | 19,1175 | 18,4245 | 19,6386 | + | iBAQ Sec | 2 | 2 | 2 | 15,5 | 15,5 | 15,5 | 36,105 | 0 | 11,573 | 6889400 | 3 | 2 | 2 | 2 | 1 | 1 | 1 | 2,50669 | 0,00923 | 3,39088 | 6,3704 | DIPPA_3 | DIPPA_3 | DIPPA_32790.mRNA.1 hypothetical protein conserved |
| 16,8018 | 15,3255 | 16,9874 | 12,2039 | 13,3463 | 13,4235 | 20,7199 | 19,5381 | 20,2716 | 19,0638 | 18,9279 | 17,7313 | + | iBAQ Sec | 2 | 2 | 2 | 8,7 | 8,7 | 8,7 | 52,693 | 0 | 15,702 | 8930600 | 4 | 2 | 2 | 2 | 0 | 0 | 1 | 2,16912 | 0,02321 | 3,38034 | 5,14356 | DIPPA_1 | DIPPA_1 | DIPPA_18049.mRNA.1 Alanine aminotransferase 2 |
| 16,6754 | 16,4467 | 16,5648 | 13,7126 | 13,065 | 12,8012 | 18,16 | 19,8417 | 19,9202 | 18,6371 | 17,5106 | 18,8344 | + | iBAQ Sec | 3 | 3 | 3 | 9,5 | 9,5 | 9,5 | 50,86 | 0 | 21,708 | 9368900 | 4 | 2 | 3 | 3 | 1 | 1 | 0 | 3,57099 | 0 | 3,36931 | 12,0892 | DIPPA_3 | DIPPA_3 | DIPPA_33575.mRNA.1 hypothetical protein AMSG_11957 |
| 18,3297 | 15,1461 | 18,8892 | 14,2294 | 13,638 | 14,3943 | 22,6754 | 18,719 | 22,1604 | 19,0214 | 18,4969 | 19,3777 | | | 4 | 4 | 4 | 25,3 | 25,3 | 25,3 | 37,953 | 0 | 28,497 | 1,8E+07 | 5 | 3 | 1 | 4 | 1 | 0 | 1 | 1,32675 | 0,07025 | 3,36778 | 2,83461 | TR24310 | TR24310 | TR24310__c0_g1_i1__m.2847 #MITO 1 Cello #PFAM PF01210.18 NAD_Gly3P_dh_N N[illegible] |
| 15,2633 | 19,1617 | 14,9895 | 13,024 | 14,4611 | 11,8792 | 19,5449 | 18,5262 | 19,5304 | 18,7838 | 19,2252 | 17,9024 | | | 2 | 2 | 2 | 3,8 | 3,8 | 3,8 | 67,933 | 0,00144 | 11,138 | 2,5E+07 | 1 | 1 | 1 | 1 | 0 | 1 | 0 | 1,02077 | 0,11366 | 3,35005 | 2,1745 | DIPPA_0 | DIPPA_0 | DIPPA_09076.mRNA.1 Serine/threonine-protein kinase CTR1 |
| 16,1936 | 13,2708 | 16,3328 | 12,2327 | 10,7691 | 12,7469 | 19,8607 | 19,5618 | 19,923 | 18,1533 | 19,5991 | 19,4787 | | | 2 | 2 | 2 | 10,8 | 10,8 | 10,8 | 48,773 | 0 | 11,91 | 4353000 | 2 | 2 | 0 | 2 | 0 | 0 | 0 | 1,34902 | 0,06755 | 3,34953 | 2,88541 | DIPPA_2 | DIPPA_2 | DIPPA_23120.mRNA.1 Purine nucleoside phosphorylase |
| 18,2294 | 18,0758 | 19,0292 | 16,4541 | 17,3855 | 11,4523 | 18,8329 | 18,6564 | 17,9314 | 18,2642 | 18,5097 | 19,4808 | | | 2 | 2 | 2 | 26,8 | 26,8 | 26,8 | 14,394 | 0,00141 | 10,765 | 1,3E+07 | 2 | 1 | 2 | 2 | 1 | 1 | 0 | 0,83199 | 0,16854 | 3,3475 | 1,79417 | DIPPA_0 | DIPPA_0 | DIPPA_08877.mRNA.1 hypothetical protein |
| 19,034 | 18,582 | 18,5763 | 14,5493 | 15,3151 | 16,481 | 21,9893 | 21,3926 | 21,8827 | 19,3184 | 19,157 | 19,8257 | + | iBAQ Sec | 13 | 13 | 13 | 23,4 | 23,4 | 23,4 | 104,62 | 0 | 91,316 | 1,1E+08 | 30 | 11 | 13 | 11 | 3 | 6 | 9 | 2,31357 | 0,01659 | 3,28228 | 5,64236 | DIPPA_0 | DIPPA_0 | DIPPA_02118.mRNA.1 hypothetical protein |
| 17,0964 | 15,8506 | 16,2443 | 12,6747 | 11,8313 | 14,9261 | 20,2692 | 18,6432 | 20,2282 | 19,1816 | 18,9408 | 19,7289 | + | iBAQ Sec | 3 | 3 | 3 | 10,3 | 10,3 | 10,3 | 40,612 | 0 | 16,935 | 7391600 | 4 | 3 | 2 | 2 | 0 | 0 | 2 | 1,51246 | 0,04859 | 3,25305 | 3,2721 | DIPPA_0 | DIPPA_0 | DIPPA_08118.mRNA.1 Eukaryotic translation initiation factor 5 |
| 18,8875 | 16,7149 | 18,4604 | 16,9385 | 10,4159 | 16,992 | 21,4536 | 18,588 | 20,4418 | 19,592 | 18,4178 | 19,8735 | | | 2 | 2 | 2 | 31,9 | 31,9 | 31,9 | 10,27 | 0 | 12,196 | 1E+07 | 3 | 2 | 1 | 2 | 2 | 0 | 2 | 0,64047 | 0,23054 | 3,23879 | 1,41923 | DIPPA_1 | DIPPA_1 | DIPPA_11283.mRNA.1 hypothetical protein |
| [illegible] | | | | | | | | | | | | | | | | | | | | | | | | | | | | | | | | | | | | | |

| | | | | | | | | | | | | | | | | | | | | | | | | | | | | | | | | | | | | | |
|---|---|---|---|---|---|---|---|---|---|---|---|---|---|---|---|---|---|---|---|---|---|---|---|---|---|---|---|---|---|---|---|---|---|---|---|---|---|
| [illegible] | | | | | | | | | | | | | | | | | | | | | | | | | | | | | | | | | | | | | |
| 13,7537 | 14,8565 | 15,2611 | 13,0284 | 11,9133 | 14,5178 | 19,1202 | 17,3815 | 19,5866 | 18,2669 | 19,7251 | 18,4495 | | | 2 | 2 | 2 | 30,7 | 30,7 | 30,7 | 20,192 | 0,00073 | 11,273 | 1118400 | 4 | 0 | 1 | 1 | 0 | 0 | 0 | 0,7708 | 0,18294 | 1,47059 | 1,67367 | DIPPA_2 | DIPPA_2 | DIPPA_25282.mRNA.1 Caltractin |
| 19,5226 | 18,3771 | 19,6845 | 18,1726 | 16,7207 | 18,2892 | 22,2221 | 21,4061 | 22,9047 | 21,6163 | 20,8861 | 21,8957 |  |  | 6 | 6 | 6 | 18,7 | 18,7 | 18,7 | 57,518 | 0 | 77,27 | 7,7E+07 | 22 | 6 | 5 | 6 | 4 | 3 | 5 | 1,05904 | 0,10417 | 1,46726 | 2,25367 | DIPPA_0 | DIPPA_0 | DIPPA_04617.mRNA.1 6-phosphogluconate dehydrogenase decarboxylating |
| 15,2093 | 14,9479 | 14,6626 | 13,354 | 14,4846 | 12,5881 | 19,5495 | 18,087 | 19,09 | 19,1485 | 18,618 | 18,8262 | | | 2 | 2 | 2 | 6 | 6 | 6 | 71,729 | 0 | 36,222 | 4941400 | 4 | 2 | 2 | 2 | 1 | 1 | 0 | 1,20107 | 0,0858 | 1,46435 | 2,55545 | DIPPA_1 | DIPPA_1 | DIPPA_15230.mRNA.1 DNA-directed RNA polymerase II subunit 2;TR113034__c0_g1_i1__m.25836 #MITO 0 #PFAM PF00562.23 RNA_pol_Rpb2_6 RNA polymerase Rpb2, domain 6;PF04563.10 RNA_pol_Rpb2_1 RNA polymerase beta subunit;PF04561.9 RNA_pol_Rpb2_2 RNA polymerase |
| 14,1016 | 13,8549 | 14,3795 | 12,3445 | 13,4259 | 12,1761 | 18,8547 | 17,8421 | 19,0231 | 18,3182 | 18,6758 | 19,3022 | + | iBAQ Sec | 2 | 2 | 2 | 3,6 | 3,6 | 3,6 | 121,35 | 0 | 14,462 | 3325800 | 6 | 2 | 2 | 2 | 2 | 0 | 1 | 1,59785 | 0,04231 | 1,46311 | 3,48472 | DIPPA_0 | DIPPA_0 | DIPPA_04359.mRNA.1 Eukaryotic translation initiation factor 4G |
| 22,4841 | 20,5061 | 22,2772 | 20,5492 | 19,4042 | 20,937 | 24,8049 | 23,0803 | 24,6267 | 23,5291 | 22,5886 | 23,8637 |  |  | 18 | 18 | 18 | 49,7 | 49,7 | 49,7 | 50,563 | 0 | 187,47 | 3,6E+08 | 68 | 18 | 15 | 17 | 14 | 10 | 15 | 0,87252 | 0,15476 | 1,45896 | 1,87459 | DIPPA_0 | DIPPA_0 | DIPPA_07423.mRNA.1 Probable citrate synthase mitochondrial |
| 17,8659 | 16,4494 | 17,8522 | 16,4289 | 14,6403 | 16,7285 | 21,161 | 19,7693 | 21,0901 | 19,037 | 17,8295 | 18,8317 | | | 2 | 2 | 2 | 20,7 | 20,7 | 20,7 | 24,384 | 0 | 13,76 | 9676000 | 7 | 2 | 1 | 2 | 1 | 0 | 1 | 0,84127 | 0,16407 | 1,45655 | 1,81253 | DIPPA_2 | DIPPA_2 | DIPPA_21605.mRNA.1 Glutathione S-transferase |
| 21,6516 | 19,9016 | 20,777 | 19,3591 | 18,6206 | 19,9934 | 23,5664 | 22,5142 | 23,3659 | 22,4307 | 21,9637 | 23,0694 | | | 12 | 6 | 6 | 59,9 | 32,2 | 32,2 | 33,987 | 0 | 44,444 | 1,5E+08 | 24 | 6 | 6 | 6 | 5 | 4 | 6 | 1,06261 | 0,10349 | 1,45236 | 2,26111 | DIPPA_1 | DIPPA_1 | DIPPA_15475.mRNA.1 Phosphoenolpyruvate carboxykinase |
| 20,2098 | 18,1092 | 19,9112 | 18,2775 | 17,2004 | 18,4255 | 22,422 | 20,69 | 22,3737 | 20,976 | 20,3071 | 20,943 | | | 5 | 5 | 5 | 31,4 | 31,4 | 31,4 | 34,683 | 0 | 36,409 | 5,9E+07 | 15 | 5 | 3 | 4 | 5 | 3 | 4 | 0,88252 | 0,15198 | 1,44229 | 1,89453 | DIPPA_3 | DIPPA_3 | DIPPA_34617.mRNA.1 Putative NADP-dependent oxidoreductase YfmJ;TR115258__c0_g1_i1__m.26742 #MITO 1 HMMTOP #PFAM PF00107.21 ADH_zinc_N Zinc-binding dehydrogenase;2 x PF13602.1 ADH_zinc_N_2 Zinc-binding dehydrogenase;PF01370.16 Epimerase NAD dependent ep |
| 15,724 | 15,8028 | 14,1753 | 12,0877 | 15,1117 | 14,2027 | 20,011 | 18,4319 | 19,317 | 20,0103 | 18,163 | 19,3151 | | | 2 | 2 | 2 | 11,1 | 11,1 | 11,1 | 49,598 | 0 | 12,151 | 6391700 | 1 | 2 | 2 | 1 | 0 | 1 | 1 | 0,61887 | 0,24171 | 1,43328 | 1,37715 | DIPPA_3 | DIPPA_3 | DIPPA_35125.mRNA.1 hypothetical protein |
| 18,7275 | 18,8489 | 18,5136 | 16,622 | 17,2108 | 17,9787 | 22,5303 | 22,603 | 22,2157 | 21,4714 | 21,3532 | 21,989 | + | iBAQ Sec | 5 | 5 | 5 | 11,5 | 11,5 | 11,5 | 95,05 | 0 | 34,991 | 8,9E+07 | 17 | 5 | 5 | 5 | 4 | 3 | 5 | 1,61287 | 0,04146 | 1,42618 | 3,52292 | DIPPA_7 | DIPPA_7 | DIPPA_70132.mRNA.1 hypothetical protein |
| 19,4726 | 18,8139 | 18,9544 | 18,2117 | 18,1497 | 17,5927 | 21,6988 | 21,8426 | 22,6904 | 21,6975 | 21,4641 | 21,3282 | + | iBAQ Sec | 9 | 9 | 9 | 22,8 | 22,8 | 22,8 | 63,628 | 0 | 70,596 | 1E+08 | 20 | 9 | 7 | 8 | 6 | 7 | 6 | 1,80288 | 0,03083 | 1,42557 | 4,02919 | DIPPA_0 | DIPPA_0 | DIPPA_03244.mRNA.1 Guanine nucleotide-binding protein-like 3 homolog |
| 17,4226 | 15,695 | 17,3988 | 16,503 | 14,1208 | 15,6228 | 20,9082 | 18,1462 | 20,8108 | 18,8951 | 19,0369 | 19,0415 | | | 2 | 2 | 2 | 10 | 10 | 10 | 33,91 | 0 | 14,026 | 9420400 | 2 | 2 | 2 | 2 | 2 | 0 | 2 | 0,72331 | 0,19694 | 1,42328 | 1,58071 | DIPPA_2 | DIPPA_2 | DIPPA_25913.mRNA.1 Uncharacterized protein L780 |
| 18,4499 | 18,436 | 19,3676 | 16,8686 | 18,2842 | 16,8433 | 21,1898 | 21,1939 | 21,9366 | 20,6071 | 21,3378 | 20,4452 | | | 5 | 5 | 5 | 16 | 16 | 16 | 56,514 | 0 | 34,924 | 4E+07 | 18 | 4 | 5 | 5 | 3 | 5 | 4 | 1,1763 | 0,08896 | 1,41911 | 2,50185 | DIPPA_1 | DIPPA_1 | DIPPA_10818.mRNA.1 hypothetical protein |
| 16,1255 | 16,4636 | 17,1524 | 14,9161 | 15,5593 | 15,0127 | 19,6822 | 19,9485 | 20,3482 | 19,9097 | 19,2527 | 19,6765 | + | iBAQ Sec | 6 | 6 | 6 | 10,8 | 10,8 | 10,8 | 105,87 | 0 | 70,204 | 2,3E+07 | 18 | 4 | 5 | 6 | 2 | 5 | 4 | 1,76003 | 0,03241 | 1,4178 | 3,91115 | DIPPA_0 | DIPPA_0 | DIPPA_02471.mRNA.1 Coatomer subunit beta-1 |
| 17,4301 | 18,112 | 17,8554 | 15,4521 | 16,963 | 16,73 | 18,8865 | 22,0227 | 21,7297 | 19,1744 | 21,603 | 19,3953 | | | 4 | 4 | 4 | 5,5 | 5,5 | 5,5 | 108,6 | 0 | 23,563 | 5,1E+07 | 6 | 2 | 4 | 3 | 1 | 3 | 2 | 1,30239 | 0,07366 | 1,41748 | 2,7795 | DIPPA_2 | DIPPA_2 | DIPPA_20069.mRNA.1 hypothetical protein |
| 18,8034 | 17,6551 | 18,2458 | 16,742 | 16,7371 | 16,9741 | 21,6354 | 20,6403 | 21,2973 | 19,6088 | 19,4711 | 19,7839 | + | iBAQ Sec | 2 | 2 | 2 | 17,9 | 17,9 | 17,9 | 28,337 | 0 | 19,113 | 2,1E+07 | 12 | 2 | 2 | 2 | 2 | 2 | 2 | 1,84938 | 0,0293 | 1,41699 | 4,15996 | DIPPA_1 | DIPPA_1 | DIPPA_14880.mRNA.1 Predicted protein OS_Naegleria gruberi OX_5762 GN_NAEGRDRAFT_29414 PE_4 SV_1 |
| 18,4745 | 16,1633 | 18,5552 | 17,0873 | 14,7365 | 17,126 | 22,7263 | 21,1409 | 22,7794 | 21,4027 | 20,2827 | 21,5917 | | | 12 | 12 | 12 | 18,1 | 18,1 | 18,1 | 105,45 | 0 | 141,64 | 6,4E+07 | 22 | 10 | 6 | 12 | 5 | 5 | 8 | 0,56422 | 0,27279 | 1,41439 | 1,27055 | DIPPA_0 | DIPPA_0 | DIPPA_04546.mRNA.1 Staphylococcal nuclease domain-containing protein 1 |
| 20,3443 | 18,0296 | 19,9647 | 18,2434 | 17,224 | 18,6322 | 22,9577 | 21,3102 | 22,694 | 21,755 | 20,6619 | 21,9845 | | | 20 | 20 | 20 | 38,9 | 38,9 | 38,9 | 96,141 | 0 | 164,82 | 2E+08 | 44 | 20 | 11 | 18 | 13 | 9 | 14 | 0,78477 | 0,17968 | 1,413 | 1,70109 | DIPPA_1 | DIPPA_1 | DIPPA_19939.mRNA.1 Aconitate hydratase B |
| 16,9358 | 20,2464 | 19,8235 | 17,349 | 18,8699 | 16,5499 | 19,17 | 22,1542 | 19,0286 | 21,6281 | 18,5778 | 18,1665 | | | 3 | 3 | 3 | 25,4 | 25,4 | 25,4 | 26,238 | 0 | 20,263 | 4,6E+07 | 4 | 1 | 3 | 3 | 2 | 2 | 1 | 0,49577 | 0,31473 | 1,41234 | 1,13622 | DIPPA_0 | DIPPA_0 | DIPPA_02340.mRNA.1 Ribosome biogenesis regulatory protein homolog |
| 25,2327 | 23,4889 | 24,4654 | 23,4636 | 22,407 | 23,0796 | 25,5592 | 23,9781 | 24,8732 | 24,2797 | 23,2842 | 23,8105 | | | 6 | 6 | 2 | 71,9 | 71,9 | 24,7 | 9,5803 | 0 | 145,98 | 4,6E+08 | 43 | 2 | 2 | 2 | 2 | 2 | 2 | 1,12264 | 0,09505 | 1,41226 | 2,38718 | DIPPA_1 | DIPPA_1 | DIPPA_15691.mRNA.1 hypothetical protein;DIPPA_15704.mRNA.1 hypothetical protein;DIPPA_15702.mRNA.1 hypothetical protein;DIPPA_15697.mRNA.1 hypothetical protein;DIPPA_15689.mRNA.1 hypothetical protein;DIPPA_15686.mRNA.1 hypothetical protein;DIPPA_08178.mRNA |
| 21,068 | 19,0719 | 20,4526 | 18,913 | 17,8909 | 19,5544 | 23,8431 | 21,5342 | 22,8803 | 22,1881 | 21,3148 | 22,3909 | | | 7 | 7 | 7 | 25,1 | 25,1 | 25,1 | 37,383 | 0 | 67,541 | 1,1E+08 | 18 | 7 | 6 | 6 | 5 | 3 | 6 | 0,85946 | 0,15992 | 1,41141 | 1,84863 | DIPPA_3 | DIPPA_3 | DIPPA_34173.mRNA.1 Transaldolase 1 |
| 19,3537 | 17,9803 | 19,3624 | 18,1385 | 17,0003 | 17,3395 | 23,146 | 22,0548 | 23,1477 | 22,1981 | 21,2829 | 22,456 | | | 9 | 9 | 2 | 14,7 | 14,7 | 3,7 | 122,03 | 0 | 64,161 | 1E+08 | 26 | 2 | 0 | 2 | 2 | 1 | 1 | 1,16018 | 0,0902 | 1,40599 | 2,46719 | TR97238 | TR97238 | TR97238__c7_g3_i5__m.20955 #MITO 1 Predotar #PFAM PF07748.8 Glyco_hydro_38C Glycosyl hydrolases family 38 C-terminal domain;PF01074.17 Glyco_hydro_38 Glycosyl hydrolases family 38 N-terminal domain;2 x PF09261.6 Alpha-mann_mid Alpha mannosidase, middle |
| 17,2355 | 17,2324 | 17,689 | 15,4161 | 16,3834 | 16,1432 | 21,1699 | 21,3877 | 21,7366 | 20,7325 | 21,0389 | 20,6335 | + | iBAQ Sec | 5 | 5 | 5 | 12,2 | 12,2 | 12,2 | 99,769 | 0 | 39,502 | 4,2E+07 | 22 | 5 | 5 | 5 | 2 | 3 | 4 | 1,89225 | 0,02831 | 1,40473 | 4,28313 | DIPPA_0 | DIPPA_0 | DIPPA_01300.mRNA.1 WD repeat-containing protein 3 homolog |
| 21,3401 | 17,1412 | 20,9431 | 18,2782 | 18,1839 | 18,7521 | 21,9864 | 19,1563 | 21,9028 | 21,1175 | 20,5034 | 21,1635 | | | 5 | 5 | 5 | 64,4 | 64,4 | 64,4 | 14,342 | 0 | 34,667 | 3,9E+07 | 15 | 5 | 1 | 4 | 2 | 2 | 2 | 0,44709 | 0,35457 | 1,40343 | 1,03968 | DIPPA_2 | DIPPA_2 | DIPPA_23705.mRNA.1 hypothetical protein |
| 20,105 | 19,8749 | 20,2137 | 18,341 | 18,5339 | 19,1098 | 23,1983 | 23,0716 | 23,1969 | 22,6436 | 22,305 | 22,7893 | + | iBAQ Sec | 7 | 7 | 7 | 27,8 | 27,8 | 27,8 | 45,108 | 0 | 120,39 | 1,4E+08 | 25 | 6 | 6 | 6 | 3 | 5 | 5 | 2,29503 | 0,01879 | 1,40295 | 5,57625 | DIPPA_2 | DIPPA_2 | DIPPA_28257.mRNA.1 hypothetical protein;TR85811__c2_g1_i1__m.17543 #MITO 0 #PFAM - #BLASTP_UNIREF90 7e-06 Putative uncharacterized protein n=1 Tax=Prorocentrum minimum RepID=E8Z6S1_PROMN (length=241) |
| 19,6024 | 19,3448 | 19,5798 | 17,789 | 18,4125 | 18,1246 | 22,6896 | 22,4919 | 22,6368 | 21,5384 | 21,6992 | 21,923 | + | iBAQ Sec | 9 | 9 | 9 | 27,8 | 27,8 | 27,8 | 56,196 | 0 | 65,861 | 9,9E+07 | 25 | 6 | 8 | 6 | 4 | 7 | 5 | 2,67506 | 0,00606 | 1,40025 | 7,06881 | DIPPA_2 | DIPPA_2 | DIPPA_20017.mRNA.1 Polypyrimidine tract-binding protein homolog 3;DIPPA_20017.mRNA.2 Protein elav |
| 18,078 | 18,4186 | 18,9166 | 16,5514 | 17,3437 | 17,3231 | 20,9764 | 21,2496 | 21,9974 | 20,9889 | 21,149 | 20,8164 | + | iBAQ Sec | 10 | 10 | 10 | 26,7 | 26,7 | 26,7 | 85,462 | 0 | 77,713 | 8E+07 | 24 | 8 | 10 | 9 | 4 | 6 | 6 | 1,76314 | 0,03288 | 1,39834 | 3,91964 | DIPPA_1 | DIPPA_1 | DIPPA_16800.mRNA.3 Ribosome biogenesis protein BOP1 homolog;DIPPA_16800.mRNA.2 Ribosome biogenesis protein BOP1 homolog;DIPPA_16800.mRNA.1 Ribosome biogenesis protein BOP1 homolog |
| 21,0668 | 19,8816 | 20,7199 | 19,2928 | 17,517 | 19,7695 | 22,6769 | 21,1647 | 22,4496 | 21,5684 | 20,4968 | 22,0456 | | | 9 | 9 | 6 | 45,8 | 45,8 | 34,2 | 25,812 | 0 | 73,323 | 1,1E+08 | 24 | 6 | 4 | 6 | 4 | 3 | 5 | 0,67352 | 0,21713 | 1,39633 | 1,4836 | DIPPA_3 | DIPPA_3 | DIPPA_31391.mRNA.1 Glutathione S-transferase 1 |
| 17,1425 | 16,5377 | 17,2048 | 15,6914 | 15,2774 | 15,737 | 20,3364 | 19,5223 | 20,3768 | 19,2881 | 18,9304 | 19,364 | + | iBAQ Sec | 2 | 2 | 2 | 10,1 | 10,1 | 10,1 | 39,043 | 0 | 29,521 | 1,2E+07 | 14 | 2 | 2 | 2 | 1 | 1 | 1 | 2,2438 | 0,01861 | 1,39305 | 5,39681 | DIPPA_0 | DIPPA_0 | DIPPA_08492.mRNA.1 Probable nitronate monooxygenase |
| 18,0831 | 17,3095 | 17,9273 | 15,7525 | 16,7064 | 16,6917 | 21,6257 | 21,0699 | 21,2601 | 20,4044 | 20,3992 | 20,7776 | + | iBAQ Sec | 4 | 4 | 4 | 16,2 | 16,2 | 16,2 | 60,009 | 0 | 29,304 | 3,2E+07 | 12 | 3 | 3 | 4 | 2 | 3 | 3 | 1,61404 | 0,0411 | 1,38981 | 3,52591 | DIPPA_1 | DIPPA_1 | DIPPA_10005.mRNA.1 hypothetical protein |
| 19,918 | 18,3197 | 19,8709 | 18,6043 | 17,8901 | 17,4515 | 22,2573 | 21,362 | 22,1645 | 21,6241 | 18,1765 | 18,3025 | | | 2 | 2 | 2 | 27,3 | 27,3 | 27,3 | 19,409 | 0 | 24,074 | 3,7E+07 | 15 | 2 | 2 | 2 | 2 | 1 | 1 | 1,04571 | 0,10783 | 1,38756 | 2,22601 | DIPPA_0 | DIPPA_0 | DIPPA_08077.mRNA.1 Nascent polypeptide-associated complex subunit alpha-like protein;DIPPA_08116.mRNA.1 Nascent polypeptide-associated complex subunit alpha-like protein;DIPPA_08083.mRNA.1 Nascent polypeptide-associated complex subunit alpha-like protein |
| 16,5811 | 17,6534 | 17,3271 | 15,3095 | 16,313 | 15,7869 | 19,8467 | 20,7824 | 20,6626 | 18,4859 | 19,529 | 19,5507 | | | 3 | 3 | 3 | 12,6 | 12,6 | 12,6 | 36,74 | 0 | 24,732 | 1,2E+07 | 12 | 2 | 3 | 2 | 1 | 3 | 2 | 1,49133 | 0,05357 | 1,38406 | 3,22065 | DIPPA_0 | DIPPA_0 | DIPPA_04995.mRNA.1 Cell differentiation protein rcd1 |
| 19,5141 | 17,7048 | 18,9678 | 16,9495 | 17,3257 | 17,7678 | 18,8941 | 19,08 | 19,3976 | 17,9595 | 18,8221 | 20,0374 | | | 3 | 3 | 3 | 48,3 | 48,3 | 48,3 | 16,643 | 0 | 22,948 | 1,8E+07 | 4 | 1 | 1 | 1 | 1 | 1 | 1 | 1,10905 | 0,09714 | 1,38125 | 2,35845 | DIPPA_2 | DIPPA_2 | DIPPA_23696.mRNA.1 Caltractin |
| 22,0306 | 20,1854 | 21,8134 | 19,8794 | 19,1383 | 20,8791 | 23,3145 | 21,9272 | 23,0568 | 22,206 | 21,4268 | 22,6762 |  |  | 13 | 13 | 13 | 46,5 | 46,5 | 46,5 | 40,716 | 0 | 141,96 | 2,6E+08 | 60 | 12 | 11 | 12 | 8 | 10 | 11 | 0,82901 | 0,1687 | 1,37758 | 1,78828 | DIPPA_3 | DIPPA_3 | DIPPA_30805.mRNA.1 Fructose-bisphosphate aldolase glycosomal;DIPPA_23086.mRNA.1 Fructose-bisphosphate aldolase glycosomal |
| 19,2393 | 20,1289 | 19,4371 | 17,9255 | 18,3071 | 18,464 | 21,862 | 22,2464 | 21,982 | 21,6176 | 21,982 | 21,8443 | + | iBAQ Sec | 9 | 9 | 9 | 36,7 | 36,7 | 36,7 | 37,455 | 0 | 114,29 | 1,1E+08 | 32 | 7 | 8 | 8 | 6 | 6 | 7 | 1,92142 | 0,02793 | 1,36958 | 4,36837 | DIPPA_1 | DIPPA_1 | DIPPA_14424.mRNA.1 Serine-threonine kinase receptor-associated protein |
| 16,6045 | 16,4698 | 16,9378 | 14,6214 | 15,4442 | 15,8442 | 19,6864 | 19,7383 | 20,5367 | 18,7884 | 19,25 | 19,5033 | + | iBAQ Sec | 2 | 2 | 2 | 8,6 | 8,6 | 8,6 | 37,718 | 0 | 14,066 | 1,1E+07 | 4 | 2 | 2 | 2 | 1 | 2 | 2 | 1,62098 | 0,04055 | 1,36743 | 3,54367 | DIPPA_2 | DIPPA_2 | DIPPA_22316.mRNA.1 Putative monooxygenase Rv1533 |
| 16,8962 | 16,1723 | 17,865 | 11,8057 | 17,552 | 17,4744 | 24,4941 | 19,891 | 24,6136 | 18,1218 | 25,0767 | 25,7131 | | | 3 | 3 | 3 | 1 | 1 | 1 | 448,97 | 0 | 21,01 | 1,6E+08 | 4 | 2 | 1 | 2 | 1 | 3 | 3 | 0,27996 | 0,54526 | 1,36712 | 0,69581 | DIPPA_1 | DIPPA_1 | DIPPA_10720.mRNA.1 hypothetical protein;TR67648__c0_g1_i2__m.11458 #MITO 0 #PFAM 22 x PF08309.6 LVIVD LVIVD repeat;8 x PF09826.4 Beta_propel Beta propeller domain;7 x PF01197.13 Ribosomal_L31 Ribosomal protein L31;2 x PF09992.4 DUF2233 Predicted peri |
| 19,217 | 18,6134 | 19,2583 | 17,4918 | 17,6798 | 17,8164 | 22,2797 | 21,9436 | 22,3474 | 21,296 | 21,4323 | 21,547 | + | iBAQ Sec | 6 | 6 | 6 | 32,6 | 32,6 | 32,6 | 37,361 | 0 | 39,607 | 4,9E+07 | 12 | 6 | 6 | 5 | 4 | 5 | 5 | 2,40479 | 0,01279 | 1,36689 | 5,97708 | DIPPA_2 | DIPPA_2 | DIPPA_28118.mRNA.1 Malate dehydrogenase |
| 14,7459 | 15,3321 | 15,7919 | 13,339 | 14,6817 | 13,7507 | 18,9919 | 19,0126 | 18,3194 | 19,3795 | 17,9516 | 18,0157 | | | 2 | 2 | 2 | 5,1 | 5,1 | 5,1 | 72,095 | 0,00144 | 11,11 | 7242100 | 2 | 1 | 2 | 1 | 1 | 1 | 1 | 1,28293 | 0,07615 | 1,36616 | 2,73584 | DIPPA_3 | DIPPA_3 | DIPPA_31987.mRNA.1 Probable U3 small nucleolar RNA-associated protein 7 |
| 20,9027 | 20,6095 | 20,9552 | 19,5591 | 19,4487 | 19,3793 | 23,0962 | 22,9267 | 23,1308 | 22,2936 | 22,2879 | 22,229 | + | iBAQ Sec | 12 | 12 | 12 | 41,3 | 41,3 | 41,3 | 50,069 | 0 | 140,13 | 2,4E+08 | 53 | 11 | 12 | 11 | 9 | 10 | 10 | 3,46669 | 0,00653 | 1,36012 | 11,3682 | DIPPA_2 | DIPPA_2 | DIPPA_22663.mRNA.1 ATP-dependent RNA helicase uap56 |
| 18,3085 | 16,1358 | 18,5755 | 16,7812 | 15,5611 | 16,6055 | 21,3167 | 20,6075 | 21,914 | 20,5638 | 18,2423 | 20,8604 | | | 4 | 4 | 4 | 10,2 | 10,2 | 10,2 | 55,652 | 0 | 25,739 | 2,6E+07 | 9 | 4 | 3 | 4 | 3 | 1 | 2 | 0,72079 | 0,19778 | 1,35729 | 1,57579 | DIPPA_1 | DIPPA_1 | DIPPA_15560.mRNA.1 Propionyl-CoA carboxylase beta chain |
| 19,3119 | 19,4683 | 20,1067 | 18,3035 | 18,5597 | 17,9541 | 21,3264 | 22,0645 | 22,1004 | 21,3618 | 21,3965 | 21,1347 | + | iBAQ Sec | 5 | 5 | 5 | 26,9 | 26,9 | 26,9 | 30,052 | 0 | 52,675 | 5,3E+07 | 11 | 5 | 5 | 5 | 3 | 3 | 3 | 1,97378 | 0,02533 | 1,35655 | 4,52444 | DIPPA_1 | DIPPA_1 | DIPPA_15566.mRNA.1 Ribosome biogenesis protein NSA2 homolog |
| 18,9089 | 19,101 | 19,9227 | 17,5721 | 18,7483 | 17,557 | 21,3576 | 21,748 | 22,1526 | 21,0505 | 21,302 | 20,998 | | | 10 | 10 | 10 | 41,8 | 41,8 | 41,8 | 46,519 | 0 | 66,908 | 8E+07 | 24 | 5 | 7 | 8 | 4 | 9 | 4 | 1,26264 | 0,07833 | 1,35175 | 2,69066 | DIPPA_0 | DIPPA_0 | DIPPA_00815.mRNA.1 Ribosome biogenesis protein WDR12 homolog |
| 17,6069 | 13,8583 | 17,4717 | 16,5479 | 12,2942 | 16,0505 | 20,532 | 18,4316 | 20,558 | 20,2748 | 18,2093 | 18,1689 | | | 2 | 2 | 2 | 6,7 | 6,7 | 6,7 | 54,077 | 0 | 14,222 | 8782600 | 5 | 2 | 1 | 2 | 2 | 1 | 1 | 0,30116 | 0,51738 | 1,34807 | 0,74098 | DIPPA_1 | DIPPA_1 | DIPPA_11715.mRNA.1 Tripeptidyl-peptidase 1;DIPPA_11706.mRNA.1 Tripeptidyl-peptidase 1;DIPPA_11703.mRNA.1 Tripeptidyl-peptidase 1;DIPPA_11708.mRNA.1 Tripeptidyl-peptidase 1;DIPPA_11702.mRNA.1 Tripeptidyl-peptidase 1 |
| 17,9094 | 17,6641 | 18,8942 | 15,9976 | 17,0512 | 17,3797 | 21,9936 | 21,907 | 22,468 | 20,8106 | 20,665 | 21,285 | | | 4 | 4 | 4 | 14,2 | 14,2 | 14,2 | 64,698 | 0 | 36,883 | 5,4E+07 | 14 | 3 | 3 | 4 | 2 | 4 | 4 | 1,12802 | 0,09473 | 1,34638 | 2,39859 | DIPPA_0 | DIPPA_0 | DIPPA_02925.mRNA.1 EH domain-containing protein 1 |
| 18,5262 | 19,2298 | 19,7724 | 17,5925 | 17,7051 | 18,2104 | 24,7105 | 24,7026 | 24,9234 | 23,3663 | 23,4968 | 23,8953 | + | iBAQ Sec | 25 | 25 | 25 | 22,4 | 22,4 | 22,4 | 195,34 | 0 | 245,58 | 3,4E+08 | 81 | 19 | 21 | 22 | 14 | 22 | 16 | 1,51848 | 0,04757 | 1,34011 | 3,28683 | DIPPA_1 | DIPPA_1 | DIPPA_18917.mRNA.1 Clathrin heavy chain |
| 17,1768 | 16,8901 | 17,7805 | 15,7748 | 16,2783 | 15,7784 | 20,6899 | 20,3644 | 20,8311 | 20,1563 | 19,9312 | 20,2671 | + | iBAQ Sec | 10 | 10 | 10 | 12,7 | 12,7 | 12,7 | 117,37 | 0 | 91,126 | 4,6E+07 | 33 | 7 | 8 | 9 | 6 | 5 | 9 | 1,89868 | 0,02881 | 1,33862 | 4,3018 | DIPPA_2 | DIPPA_2 | DIPPA_21452.mRNA.1 Acetyl-CoA carboxylase 1 |
| 23,1423 | 22,1551 | 23,0297 | 21,5431 | 20,8038 | 21,9652 | 25,3302 | 24,5315 | 25,2442 | 24,2779 | 23,6078 | 24,686 | | | 32 | 32 | 2 | 69,9 | 69,9 | 4,5 | 60,361 | 0 | 323,31 | 1,2E+09 | 171 | 2 | 2 | 2 | 1 | 1 | 2 | 1,3567 | 0,06694 | 1,33836 | 2,90302 | DIPPA_0 | DIPPA_0 | DIPPA_01753.mRNA.1 Chaperonin HSP60, mitochondrial |
| 18,4041 | 18,3952 | 19,4278 | 17,8187 | 17,1823 | 17,2115 | 22,0937 | 22,591 | 23,0167 | 22,4074 | 21,753 | 21,736 | + | iBAQ Sec | 7 | 7 | 7 | 20,3 | 20,3 | 20,3 | 67,512 | 0 | 51,608 | 7,9E+07 | 22 | 5 | 4 | 6 | 5 | 3 | 3 | 1,54025 | 0,04606 | 1,3382 | 3,34046 | DIPPA_3 | DIPPA_3 | DIPPA_31501.mRNA.1 ATP-dependent RNA helicase HAS1 |
| 20,2175 | 19,3198 | 19,9969 | 18,2856 | 18,4319 | 18,8062 | 24,0273 | 23,48 | 24,1013 | 22,8751 | 22,6744 | 23,1027 | + | iBAQ Sec | 22 | 2 | 2 | 50,2 | 5,5 | 5,5 | 52,758 | 0 | 71,05 | 1,3E+08 | 26 | 2 | 2 | 2 | 1 | 2 | 2 | 1,89567 | 0,02869 | 1,33683 | 4,29304 | DIPPA_3 | DIPPA_3 | DIPPA_34891.mRNA.4 Elongation factor 1-alpha;DIPPA_34891.mRNA.2 Elongation factor 1-alpha;DIPPA_15736.mRNA.1 Elongation factor 1-alpha;DIPPA_15734.mRNA.1 Elongation factor 1-alpha;DIPPA_15729.mRNA.1 Elongation factor 1-alpha;DIPPA_15746.mRNA.1 Elongation f |
| 17,6796 | 17,2156 | 17,5394 | 16,0637 | 15,8421 | 16,5229 | 20,1433 | 18,8626 | 18,9081 | 19,41 | 18,3142 | 17,955 | + | iBAQ Sec | 2 | 2 | 2 | 7,3 | 7,3 | 7,3 | 38,696 | 0 | 12,022 | 2,1E+07 | 2 | 2 | 1 | 2 | 1 | 2 | 2 | 2,27175 | 0,01891 | 1,33531 | 5,49413 | DIPPA_6 | DIPPA_6 | DIPPA_62031.mRNA.1 Alcohol dehydrogenase (NADP+)-like protein n=6 Tax=Boreoeutheria RepID=F7CBN0_HORSE (length=324) |
| 17,2356 | 18,1675 | 17,871 | 16,5105 | 16,4711 | 16,2915 | 20,7172 | 20,9047 | 20,9336 | 20,3661 | 19,9606 | 20,4399 | + | iBAQ Sec | 7 | 7 | 7 | 25,4 | 25,4 | 25,4 | 50,35 | 0 | 48,485 | 3,7E+07 | 19 | 3 | 7 | 5 | 4 | 5 | 3 | 2,03497 | 0,02508 | 1,33365 | 4,71194 | DIPPA_1 | DIPPA_1 | DIPPA_10453.mRNA.1 Kynurenine--oxoglutarate transaminase |
| 16,0998 | 14,1423 | 15,5318 | 14,1187 | 11,8058 | 15,8507 | 19,504 | 19,3116 | 19,5578 | 18,382 | 19,0885 | 19,3404 | | | 2 | 2 | 2 | 5,3 | 5,3 | 5,3 | 104,18 | 0 | 14,812 | 6714300 | 3 | 2 | 0 | 1 | 0 | 0 | 2 | 0,43675 | 0,36359 | 1,33288 | 1,01904 | DIPPA_1 | DIPPA_1 | DIPPA_18552.mRNA.1 hypothetical protein |
| 17,4198 | 14,7635 | 16,9711 | 12,9655 | 16,1049 | 16,0906 | 19,4983 | 19,0973 | 19,7938 | 18,2676 | 18,2034 | 19,4514 | | | 2 | 2 | 2 | 20,4 | 20,4 | 20,4 | 28,511 | 0 | 14,775 | 4884700 | 4 | 2 | 0 | 1 | 0 | 1 | 2 | 0,42836 | 0,37229 | 1,33115 | 1,00223 | DIPPA_1 | DIPPA_1 | DIPPA_12744.mRNA.1 hypothetical protein |
| 20,9625 | 19,0174 | 20,8801 | 18,791 | 18,4884 | 19,5872 | 22,5164 | 21,0213 | 22,7717 | 21,8476 | 20,7218 | 21,7788 | | | 9 | 9 | 9 | 41,3 | 41,3 | 41,3 | 35,276 | 0 | 93,809 | 1,1E+08 | 25 | 9 | 5 | 8 | 5 | 6 | 8 | 0,86654 | 0,15782 | 1,33114 | 1,8627 | DIPPA_1 | DIPPA_1 | DIPPA_16518.mRNA.1 Guanine nucleotide-binding protein subunit beta-like protein;DIPPA_16498.mRNA.1 Guanine nucleotide-binding protein subunit beta-like protein |
| 20,2848 | 18,9498 | 20,1304 | 18,1439 | 18,5283 | 18,7162 | 21,669 | 21,0708 | 21,6444 | 21,027 | 20,8454 | 20,5327 | | | 3 | 3 | 3 | 56,1 | 56,1 | 56,1 | 16,743 | 0 | 26,062 | 3,4E+07 | 18 | 3 | 3 | 3 | 2 | 2 | 3 | 1,36385 | 0,06597 | 1,32554 | 2,91947 | DIPPA_3 | DIPPA_3 | DIPPA_31883.mRNA.1 Ubiquitin-conjugating enzyme E2-16 kDa |
| [illegible] | | | | | | | | | | | | | | | | | | | | | | | | | | | | | | | | | | | | | |

[illegible]

| | | | | | | | | | | | | | | | | | | | | | | | | | | |
|---|---|---|---|---|---|---|---|---|---|---|---|---|---|---|---|---|---|---|---|---|---|---|---|---|---|---|
| 22,2927 | 21,7199 | 22,4313 | 21,6026 | 21,4416 | 21,5309 | 24,0127 | 23,721 | 23,9809 | 23,5793 | 23,5299 | 23,7416 | | | 5 | 5 | 5 | 44 | 44 | 44 | 15,689 | 0 | 65,82 | 1,7E+08 | 21 | 5 | 5 |
| 16,4655 | 14,3873 | 16,5028 | 15,7035 | 13,8652 | 15,9264 | 21,5987 | 20,2092 | 22,0146 | 22,2104 | 21,2087 | 21,6728 | | | 5 | 5 | 5 | 9,4 | 9,4 | 9,4 | 396,54 | 0 | 46,339 | 3,6E+07 | 10 | 4 | 3 |
| 18,2428 | 15,2726 | 16,7672 | 15,8833 | 15,6928 | 16,8465 | 20,9897 | 18,7227 | 18,9371 | 19,28 | 19,3916 | 19,7761 | | | 3 | 3 | 3 | 15,1 | 15,1 | 15,1 | 31,94 | 0 | 17,661 | 1,3E+07 | 4 | 3 | 1 |
| 19,7567 | 18,1345 | 19,5785 | 18,731 | 18,168 | 18,7414 | 22,413 | 20,6641 | 21,8036 | 21,6272 | 20,4751 | 21,3116 | | | 6 | 6 | 6 | 27,7 | 27,7 | 27,7 | 30,006 | 0 | 37,125 | 6,9E+07 | 16 | 6 | 5 |
| 19,8638 | 19,5297 | 20,2626 | 19,3738 | 18,9953 | 19,4641 | 22,2724 | 22,072 | 22,7047 | 22,3613 | 22,3251 | 22,4841 | | | 8 | 6 | 6 | 33,6 | 24,7 | 24,7 | 30,756 | 0 | 43,465 | 1E+08 | 22 | 6 | 6 |
| 17,0391 | 18,1993 | 17,6859 | 16,8991 | 17,1825 | 17,0379 | 19,1651 | 21,0439 | 19,6218 | 20,7717 | 18,093 | 17,9 | | | 3 | 3 | 3 | 20,1 | 20,1 | 20,1 | 23,98 | 0 | 21,319 | 2,3E+07 | 3 | 2 | 3 |
| 22,682 | 22,27 | 22,8217 | 21,7383 | 22,1001 | 22,134 | 24,1334 | 24,279 | 24,611 | 24,2807 | 24,4041 | 24,3748 | | | 10 | 10 | 10 | 52,8 | 52,8 | 52,8 | 21,845 | 0 | 96,512 | 4,8E+08 | 31 | 8 | 7 |
| 15,2544 | 14,8453 | 16,0969 | 14,3321 | 15,1135 | 14,9589 | 19,3789 | 18,6282 | 19,5607 | 19,7222 | 19,0507 | 18,4333 | | | 2 | 2 | 2 | 10,9 | 10,9 | 10,9 | 36,304 | 0 | 12,617 | 5444600 | 4 | 1 | 1 |
| 22,6225 | 21,8382 | 22,6773 | 21,8509 | 21,5441 | 21,9521 | 24,3933 | 23,8577 | 24,3707 | 24,0917 | 23,6441 | 24,174 | | | 4 | 4 | 4 | 39,4 | 39,4 | 39,4 | 17,949 | 0 | 58,613 | 3,1E+08 | 28 | 4 | 3 |
| 22,3688 | 22,4932 | 22,3733 | 21,6957 | 22,0562 | 21,7162 | 24,2513 | 24,5765 | 24,4835 | 24,3122 | 24,5323 | 24,2394 | + | iBAQ Sec | 15 | 15 | 3 | 56,5 | 56,5 | 15,6 | 33,735 | 0 | 157,84 | 7,7E+08 | 83 | 3 | 3 |
| 19,2498 | 17,3312 | 19,4965 | 18,3334 | 17,4805 | 18,5015 | 22,2907 | 20,7067 | 22,0743 | 21,4674 | 20,2354 | 21,944 | | | 7 | 7 | 7 | 24,2 | 24,2 | 24,2 | 51,02 | 0 | 151,22 | 7E+07 | 23 | 6 | 4 |
| 20,1132 | 20,265 | 19,9749 | 19,7177 | 19,1525 | 19,721 | 22,8341 | 22,8052 | 22,7538 | 23,1662 | 22,4361 | 22,7544 | | | 9 | 9 | 9 | 30,7 | 30,7 | 30,7 | 52,177 | 0 | 113,99 | 1,3E+08 | 33 | 5 | 7 |
| 18,3858 | 18,3436 | 18,7896 | 17,5089 | 18,3927 | 17,8628 | 21,7076 | 21,9668 | 22,2899 | 21,5597 | 21,9419 | 21,4961 | | | 8 | 8 | 8 | 21,8 | 21,8 | 21,8 | 76,826 | 0 | 55,442 | 8E+07 | 29 | 5 | 5 |
| 22,417 | 22,2538 | 22,8275 | 21,9982 | 21,946 | 21,8 | 23,2126 | 23,3811 | 23,8729 | 23,4658 | 23,4486 | 23,5472 | | | 7 | 7 | 7 | 42,9 | 42,9 | 42,9 | 23,334 | 0 | 62,826 | 2,8E+08 | 32 | 7 | 7 |
| 16,7556 | 12,4402 | 15,5366 | 15,2742 | 12,3283 | 15,379 | 19,9795 | 18,1145 | 19,531 | 18,7372 | 17,4903 | 18,7383 | | | 2 | 2 | 2 | 9,5 | 9,5 | 9,5 | 50,023 | 0 | 16,755 | 3125500 | 2 | 2 | 0 |
| 20,3593 | 20,8454 | 21,0128 | 20,1747 | 20,1523 | 20,1402 | 23,5407 | 23,9938 | 23,7207 | 23,7139 | 23,3084 | 23,4169 | | | 22 | 22 | 22 | 31,9 | 31,9 | 31,9 | 115,79 | 0 | 201,92 | 6,6E+08 | 107 | 18 | 18 |
| 20,3383 | 20,1932 | 20,0608 | 19,416 | 19,8006 | 19,6255 | 22,9769 | 23,5012 | 22,9717 | 23,1064 | 23,1189 | 22,9872 | + | iBAQ Sec | 12 | 12 | 12 | 29,7 | 29,7 | 29,7 | 74,288 | 0 | 112,76 | 2,2E+08 | 56 | 11 | 11 |
| 18,6961 | 15,8352 | 18,0029 | 16,567 | 16,4201 | 17,7991 | 22,251 | 20,2412 | 21,3066 | 21,1083 | 20,689 | 21,1086 | | | 10 | 10 | 10 | 23,3 | 23,3 | 23,3 | 61,404 | 0 | 62,981 | 4,3E+07 | 19 | 9 | 2 |
| 19,0857 | 18,9226 | 18,3097 | 17,761 | 18,6198 | 18,1946 | 22,0949 | 21,4559 | 21,3861 | 21,3084 | 21,4252 | 21,4498 | | | 9 | 9 | 9 | 36,6 | 36,6 | 36,6 | 57,988 | 0 | 112,89 | 7,6E+07 | 31 | 5 | 8 |
| 17,7186 | 18,5209 | 19,4642 | 18,1835 | 18,2707 | 17,5093 | 21,298 | 21,8299 | 22,3124 | 21,4233 | 21,7307 | 21,0992 | | | 22 | 22 | 22 | 24,1 | 24,1 | 24,1 | 127,89 | 0 | 155,27 | 1,8E+08 | 50 | 13 | 18 |
| 19,832 | 18,3195 | 20,8249 | 18,834 | 18,6396 | 19,7763 | 21,9844 | 21,864 | 22,9473 | 22,6302 | 22,154 | 23,0016 | | | 7 | 4 | 4 | 29,6 | 21,7 | 21,7 | 27,277 | 0 | 32,95 | 6,4E+07 | 20 | 3 | 4 |
| 23,5496 | 23,8505 | 24,6527 | 23,3817 | 23,6025 | 23,3465 | 25,322 | 25,7122 | 26,4593 | 25,9606 | 25,9858 | 25,7415 | | | 11 | 11 | 11 | 61,7 | 61,7 | 61,7 | 16,687 | 0 | 236,19 | 1,2E+09 | 66 | 7 | 9 |
| 15,5763 | 15,4337 | 16,3272 | 13,3466 | 16,7316 | 15,5425 | 19,5604 | 19,0263 | 20,7182 | 17,6158 | 21,19 | 20,3913 | | | 2 | 2 | 2 | 10,5 | 10,5 | 10,5 | 58,75 | 0 | 16,104 | 1,2E+07 | 8 | 1 | 1 |
| 21,7083 | 21,0929 | 22,5583 | 21,8134 | 20,4028 | 21,437 | 22,2704 | 21,5171 | 22,6714 | 22,2515 | 21,6295 | 21,7233 | | | 6 | 6 | 5 | 94,4 | 94,4 | 94,4 | 10,847 | 0 | 178,94 | 7,7E+07 | 42 | 3 | 3 |
| 13,0715 | 17,1615 | 16,6691 | 13,8449 | 15,5062 | 15,8469 | 20,3572 | 20,8198 | 18,8624 | 20,4278 | 18,6709 | 18,7737 | | | 3 | 3 | 3 | 12 | 12 | 12 | 41,731 | 0 | 20,409 | 8771300 | 4 | 0 | 2 |
| 21,6563 | 20,9225 | 21,8508 | 21,2555 | 20,7875 | 20,686 | 23,4846 | 23,3005 | 23,5584 | 23,5419 | 23,3785 | 23,3042 | | | 2 | 2 | 2 | 19,8 | 19,8 | 19,8 | 14,65 | 0 | 13,948 | 9,3E+07 | 6 | 2 | 2 |
| 22,3639 | 22,1334 | 22,856 | 21,8869 | 21,7521 | 22,0228 | 24,1877 | 24,0165 | 24,543 | 24,1 | 23,9169 | 24,2025 | | | 5 | 5 | 5 | 31,8 | 31,8 | 31,8 | 17,411 | 0 | 36,802 | 3,5E+08 | 30 | 5 | 5 |
| 20,9882 | 21,3385 | 21,3817 | 20,5174 | 20,7044 | 20,7969 | 23,8258 | 24,2302 | 24,2082 | 23,8344 | 23,903 | 24,0949 | + | iBAQ Sec | 43 | 43 | 17 | 39,9 | 39,9 | 20,1 | 156,24 | 0 | 323,31 | 1,2E+09 | 197 | 17 | 16 |
| 15,2733 | 14,6968 | 15,8323 | 14,3914 | 15,7375 | 14,0008 | 20,2363 | 18,3321 | 20,7946 | 18,2436 | 18,7053 | 18,7284 | | | 2 | 2 | 2 | 4,5 | 4,5 | 4,5 | 93,501 | 0 | 14,558 | 1,1E+07 | 8 | 2 | 1 |
| 21,6591 | 22,2434 | 20,9584 | 21,021 | 21,0927 | 21,0766 | 23,5666 | 23,951 | 23,2538 | 23,7419 | 23,3617 | 24,014 | | | 10 | 10 | 10 | 58,6 | 58,6 | 58,6 | 32,834 | 0 | 138,8 | 4,2E+08 | 58 | 8 | 8 |
| 17,4072 | 17,8079 | 17,133 | 17,0588 | 16,5451 | 17,075 | 20,0515 | 20,9018 | 20,9724 | 20,383 | 20,9736 | 20,8877 | | | 9 | 9 | 9 | 21,4 | 21,4 | 21,4 | 75,435 | 0 | 119,9 | 4,4E+07 | 22 | 5 | 7 |
| 14,31 | 13,551 | 14,1435 | 14,1858 | 14,2707 | 11,8816 | 18,1656 | 18,4314 | 18,8597 | 19,1353 | 17,8605 | 17,997 | | | 3 | 2 | 2 | 5,4 | 3,2 | 3,2 | 100,38 | 0,00142 | 10,875 | 1E+07 | 2 | 1 | 1 |
| 15,4347 | 14,6212 | 15,7268 | 14,7381 | 14,4468 | 14,9335 | 19,8182 | 18,5277 | 18,8309 | 18,7382 | 18,4375 | 18,6569 | | | 3 | 2 | 2 | 11,9 | 8,8 | 8,8 | 43,81 | 0 | 12,628 | 4912100 | 5 | 1 | 1 |
| 22,0479 | 22,1024 | 22,5543 | 21,6119 | 21,7525 | 21,6826 | 22,9766 | 23,102 | 23,3761 | 22,9269 | 23,0815 | 22,9215 | + | iBAQ Sec | 8 | 8 | 8 | 51,7 | 51,7 | 51,7 | 16,485 | 0 | 57,093 | 3,4E+08 | 40 | 8 | 8 |
| 17,8531 | 17,7719 | 18,5386 | 17,4159 | 17,4623 | 17,63 | 20,8378 | 21,3094 | 21,6214 | 20,827 | 21,1344 | 20,9289 | | | 3 | 3 | 3 | 13,8 | 13,8 | 13,8 | 51,282 | 0 | 21,795 | 3,8E+07 | 15 | 3 | 3 |
| 17,3157 | 15,6755 | 18,7433 | 16,7402 | 16,1207 | 17,2278 | 18,4928 | 18,7508 | 20,9727 | 19,8516 | 17,5571 | 19,9739 | | | 2 | 2 | 2 | 16,3 | 16,3 | 16,3 | 31,066 | 0 | 16,46 | 8989000 | 2 | 1 | 1 |
| 21,4675 | 20,5763 | 21,3856 | 20,4273 | 20,643 | 20,7164 | 22,8602 | 22,3075 | 22,8485 | 22,6269 | 22,6142 | 22,6414 | | | 13 | 13 | 13 | 67,8 | 67,8 | 67,8 | 26,308 | 0 | 93,722 | 2,1E+08 | 37 | 11 | 11 |
| 19,4389 | 17,2757 | 19,0964 | 18,2152 | 17,2078 | 18,7474 | 22,2125 | 20,4786 | 21,8027 | 21,578 | 20,4297 | 22,0292 | | | 5 | 5 | 5 | 14,2 | 14,2 | 14,2 | 80,249 | 0 | 164,14 | 6,2E+07 | 30 | 5 | 4 |
| 19,3308 | 18,0472 | 20,0174 | 18,3453 | 17,8094 | 19,6006 | 21,837 | 20,913 | 22,5323 | 21,8046 | 20,8414 | 22,5708 | | | 5 | 5 | 5 | 29,1 | 29,1 | 29,1 | 35,651 | 0 | 85,671 | 6,2E+07 | 21 | 4 | 4 |
| 19,5129 | 19,6365 | 19,6143 | 18,8106 | 19,6145 | 18,6988 | 21,0758 | 21,3472 | 21,1062 | 18,8428 | 21,8813 | 18,6665 | | | 2 | 2 | 2 | 21,6 | 21,6 | 21,6 | 12,569 | 0 | 14,135 | 2,8E+07 | 7 | 2 | 2 |
| 23,7318 | 23,9869 | 24,3971 | 23,4904 | 23,5838 | 23,4063 | 25,6931 | 26,121 | 26,4239 | 26,1697 | 26,2578 | 26,0341 | | | 27 | 27 | 27 | 51 | 51 | 51 | 64,309 | 0 | 323,31 | 4,2E+09 | 174 | 23 | 24 |
| 21,655 | 20,6663 | 22,2948 | 21,0336 | 21,0453 | 20,9082 | 23,0526 | 22,4722 | 23,7692 | 23,2528 | 22,7645 | 22,7759 | | | 6 | 6 | 6 | 72 | 72 | 72 | 11,002 | 0 | 82,469 | 1,5E+08 | 22 | 5 | 5 |
| 18,6437 | 12,7028 | 18,234 | 16,9236 | 13,1363 | 17,9 | 19,0119 | 18,2204 | 18,9591 | 18,4756 | 18,3536 | 18,3427 | | | 2 | 2 | 2 | 35,6 | 35,6 | 35,6 | 11,654 | 0 | 11,695 | 7069000 | 2 | 2 | 0 |
| 19,6643 | 20,0735 | 20,0402 | 19,2306 | 19,578 | 19,3518 | 21,5994 | 22,1947 | 21,8594 | 21,6037 | 21,8435 | 21,5121 | | | 2 | 2 | 2 | 11,7 | 11,7 | 11,7 | 23,265 | 0 | 14,01 | 5,8E+07 | 9 | 2 | 2 |
| 21,1319 | 21,7583 | 22,1646 | 20,7419 | 21,4893 | 21,2121 | 22,4522 | 22,9121 | 23,1713 | 22,69 | 22,8702 | 22,8029 | | | 4 | 4 | 4 | 34,1 | 34,1 | 34,1 | 15,803 | 0 | 44,994 | 1,3E+08 | 28 | 4 | 4 |
| 19,8937 | 18,4598 | 20,108 | 19,1214 | 18,3897 | 19,3417 | 22,5752 | 21,5056 | 22,8591 | 22,4282 | 21,54 | 22,548 | | | 9 | 9 | 9 | 23,2 | 23,2 | 23,2 | 63,888 | 0 | 64,405 | 1,6E+08 | 27 | 9 | 8 |
| 22,0062 | 22,3613 | 22,63 | 21,5202 | 21,9996 | 21,8689 | 24,0032 | 24,5574 | 24,5611 | 24,366 | 24,0894 | 24,0103 | | | 9 | 9 | 9 | 43 | 43 | 43 | 37,357 | 0 | 187,44 | 5,3E+08 | 55 | 7 | 8 |
| 17,8783 | 17,246 | 17,6598 | 17,0192 | 17,5019 | 16,6544 | 20,7485 | 19,2174 | 18,9165 | 19,7088 | 20,9125 | 20,2867 | | | 2 | 2 | 2 | 15,3 | 15,3 | 15,3 | 20,746 | 0 | 13,566 | 1,8E+07 | 5 | 2 | 1 |
| 21,6365 | 21,9641 | 22,6105 | 21,5096 | 21,779 | 21,339 | 22,9301 | 23,0042 | 23,3092 | 22,8511 | 23,3141 | 22,476 | | | 5 | 5 | 5 | 34,6 | 34,6 | 34,6 | 18,03 | 0 | 32,622 | 1,7E+08 | 21 | 4 | 5 |
| 21,0735 | 21,0811 | 21,0253 | 19,7834 | 20,983 | 20,834 | 22,7651 | 22,0653 | 22,8333 | 22,0977 | 22,4609 | 22,5517 | | | 4 | 4 | 3 | 27,6 | 27,6 | 27,6 | 14,032 | 0 | 24,056 | 7,5E+07 | 13 | 1 | 3 |
| 19,2206 | 18,4846 | 20,2736 | 18,5775 | 18,8496 | 18,9911 | 23,0424 | 22,5048 | 23,6387 | 22,4847 | 22,6167 | 22,498 | | | 6 | 6 | 6 | 40,9 | 40,9 | 40,9 | 70,838 | 0 | 57,566 | 1,3E+08 | 21 | 5 | 4 |
| 17,0925 | 12,0292 | 15,7435 | 14,4283 | 13,4857 | 15,4026 | 19,4172 | 18,8745 | 20,1356 | 17,7599 | 17,9322 | 18,0737 | | | 2 | 2 | 2 | 11,4 | 11,4 | 11,4 | 53,584 | 0 | 12,67 | 7123200 | 4 | 2 | 0 |

[illegible]

[illegible]
